# An Endogenous Aryl Hydrocarbon Receptor Ligand Induces Preeclampsia-like Phenotypes: Transcriptome, Phosphoproteome, and Cell Functions

**DOI:** 10.1101/2023.12.20.572271

**Authors:** Ying-jie Zhao, Chi Zhou, Ying-ying Wei, Si-yan Zhang, Jay S. Mishra, Hui-hui Li, Wei Lei, Kai Wang, Sathish Kumar, Jing Zheng

## Abstract

**Background:** Preeclampsia (PE) is one hypertensive disorder and a leading cause of maternal and fetal mortality and morbidity during human pregnancy. Aryl hydrocarbon receptor (AhR) is a transcription factor, which regulates vascular functions. Exogenous and endogenous AhR ligands can induce hypertension in animals. However, if dysregulation of endogenous AhR ligands contributes to the pathophysiology of PE remains elusive.

**Methods:** We measured AhR activities in human maternal and umbilical vein sera. We also applied physiological, cellular, and molecular approaches to dissect the role of endogenous AhR ligands in vascular functions during pregnancy using pregnant rats and primary human umbilical vein endothelial cells (HUVECs) as models.

**Results:** PE elevated AhR activities in human umbilical vein sera. Exposure of pregnant rats to an endogenous AhR ligand, 2-(1’H-indole-3’-carbonyl)-thiazole-4-carboxylic acid methyl ester (ITE) increased blood pressure and proteinuria, while decreased uteroplacental blood flow and reduced fetal and placental weights, all of which are hallmarks of PE. ITE dampened vascular growth and fetal sex-specifically altered immune cell infiltration in rat placentas. ITE also decreased cell proliferation and cell monolayer integrity in HUVECs *in vitro*. RNA sequencing analysis revealed that ITE dysregulated transcriptome in rat placentas and HUVECs in a fetal sex-specific manner. Bottom-up phosphoproteomics showed that ITE disrupted phosphoproteome in HUVECs. These ITE-dysregulated genes and phosphoproteins were enriched in biological functions and pathways which are highly relevant to diseases of heart, liver, and kidney, vascular functions, inflammation responses, cell death, and kinase inhibition.

**Conclusions:** Dysregulation of endogenous AhR ligands during pregnancy may lead to the development of PE with underlying impaired vascular functions, fetal sex-specific immune cell infiltration and transcriptome, and phosphoproteome. Thus, this study has provided a novel mechanism for the development of PE and potentially other forms of hypertensive pregnancies. These AhR ligand-activated genes and phosphoproteins might represent promising therapeutic and fetal sex-specific targets for PE-impaired vascular functions.

## INTRODUCTION

Preeclampsia (PE) is a hypertensive disorder and a leading cause of maternal and fetal mortality and morbidity during human pregnancy^1,2^. Children born to mothers with PE also face a higher risk of adult-onset cardiovascular diseases^3^. While causing widespread maternal vascular dysfunction, PE also impairs fetal vascular functions^1^. Specifically, PE reduces fetal blood flow along with increased vascular resistance and vascular permeability in fetoplacentas^4^. Additionally, human umbilical vein endothelial cells (HUVECs) from PE exhibit increased cell permeability^5^ and reduced nitric oxide (NO) production^6^. We have also reported that PE dysregulates transcriptome and angiogenic responses in HUVECs in a fetal sex-specific manner^7,8^, supporting the importance of sexually dimorphic regulation of endothelial functions in PE. Currently, the etiology of PE is known although many factors may contribute to the development of PE^2^. Only by improving our understanding of the pathophysiology of PE, can we develop appropriate diagnostic tools and therapeutic strategies for PE.

Aryl hydrocarbon receptor (AhR) is a ligand-activated transcription factor, which is involved in metabolizing xenobiotics e., dioxin (TCDD)^9^. AhR also participates in mediating vascular development and functions^10,11^ and immune responses^12,13^. For instance, the AhR-mutant mice are defects in fetal vasculature (patent ductus venosus and hyaloid artery)^14^. Adult AhR null mice develop hypertension with elevated angiotensin II and endothelin 1^15,16^. In contrast, endothelial cell-specific AhR-null mice are hypotensive accompanying with increases in endothelial NO synthase (eNOS) activity and NO production^17^. Moreover, treating male mice with an exogenous AhR ligand (3-methylcholanthrene; 3MC) induces hypertension^18^. Activation of AhR by an endogenous AhR ligand (6-formylindolo[3,2-b] carbazole [FICZ]), also induces pulmonary arterial hypertension in male rats^19^. Collectively, these data suggest that the AhR pathway is critical in controlling blood pressure.

Many endogenous AhR ligands have been identified, including a tryptophan (Trp)-metabolite, 2-(1’H-indole-3’-carbonyl)-thiazole-4-carboxylic acid methyl ester (ITE)^20,21^. ITE suppresses the expansion and functions of T helper cells and promotes differentiation of T-regulatory cells^12,22^. ITE also inhibits endothelial growth without altering activation of ERK1/2, AKT1, JNK, and p38^23^. Together with the observation that PE elevates indole-3-lactic acid (another Trp-derived AhR ligand) levels in maternal and fetal serum^24^, these data imply that Trp-derived AhR-ligands may contribute to the pathophysiology of PE.

In this study, we tested the hypothesis that dysregulation of endogenous AhR ligands impairs vascular functions during pregnancy, leading to the development of PE using rats and HUVECs as models. We showed that PE elevated AhR activities in fetal serum. ITE induced typical PE-like phenotypes in association with defective placental vasculature and immune cell infiltration in rats. We found that ITE dysregulated transcriptome in rat placentas and HUVECs in a fetal sex dependent manner with dysregulation of biological functions, disease-associated functions, canonical pathways, and gene networks. We also observed that ITE dysregulated phosphoproteome in HUVECs. This data provides a novel mechanism underlying PE. The identified AhR ligand target genes, phosphoproteins, and pathways may serve as therapeutic targets for PE.

## METHODS

All detailed methods are available in the online Data Supplements.

### Data Availability

RNA sequencing (RNA-seq) data have been deposited in NCBI Gene Expression Omnibus (GEO) database (GEO accession: GSE250296 for rat placentas and GSE250196 for HUVECs). Phosphoproteomics data are deposited in ProteomeXchange (PXD047834).

## RESULTS

### Elevated AhR activities in PE-fetal serum

We determined AhR activities in maternal and umbilical vein blood samples from PE and normotensive (NT) pregnancy by measuring CYP1A1/B1 mRNA levels in HUVECs using reverse transcription-quantitative PCR (RT-qPCR). The patients’ demographics and clinical characteristics for AhR activities in human sera are shown in Table S1. Maternal age and BMI were similar between PE and NT. Gestational ages and fetal birth weights were lower in PE vs. NT. Fetuses from PE were not growth restricted as their weights were above the 10th percentile for gestational age^25^. The levels of systolic, diastolic, and mean artery pressure were higher in PE vs. NT. All PE-patients had proteinuria.

Compared with control, umbilical, but not maternal vein serum increased CYP1A1/B1 mRNA levels in HUVECs (Fig. 1A and B). Compared with NT, PE further elevated CYP1A1 mRNA levels in umbilical vein serum at 12 hr (Fig. 1A). This PE-increased CYP1A1 mRNA was blocked by CH-223191, an AhR antagonist (Fig. 1C), verifying the AhR activities.

**Fig. 1.**
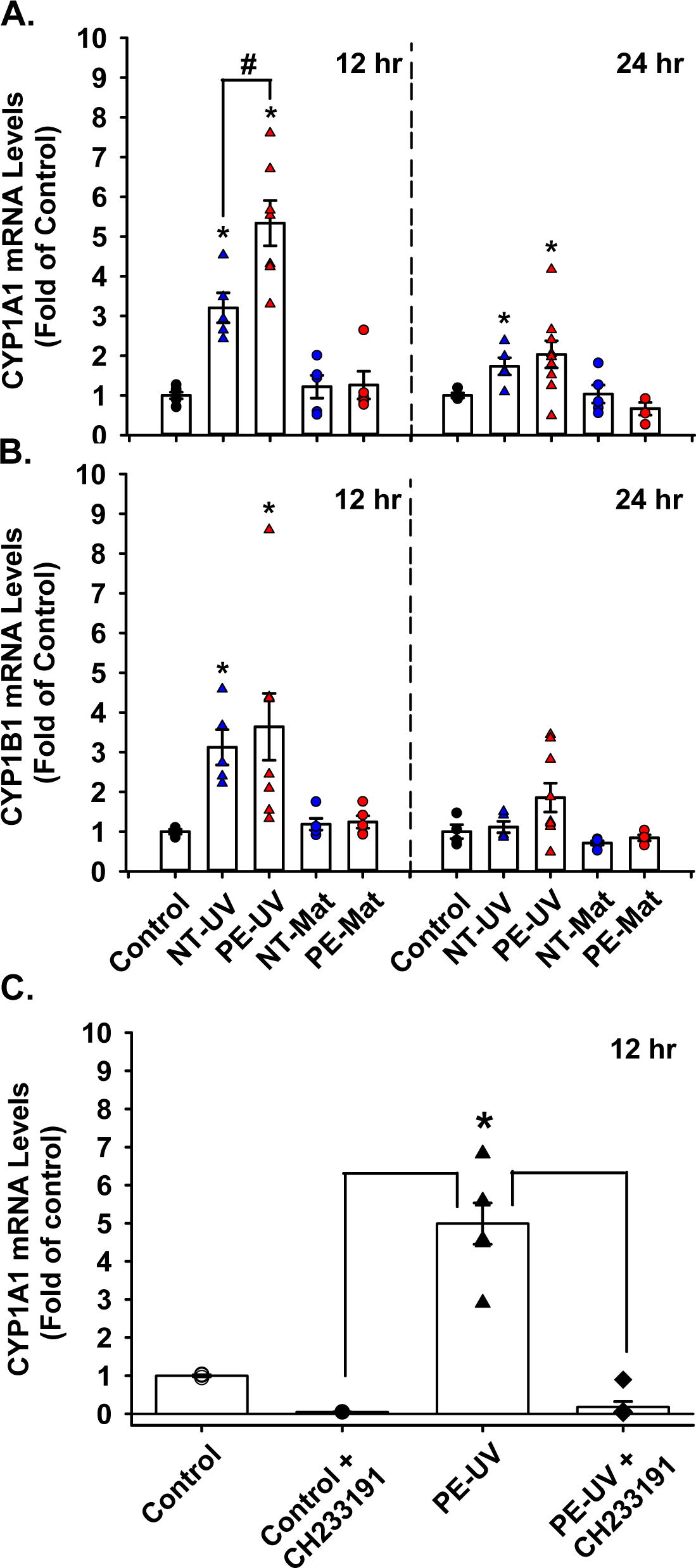
PE elevates AhR activities in fetal serum. HUVECs were treated with 20% maternal (Mat) or umbilical vein (UV) sera in ECM for 12 or 24 hr. RT-qPCR was conducted to determine changes in CYP1A1 (**A**) and CYP1B1 (**B**) mRNA, indicatives of AhR activities. Additional cells were treated with 20% PE-UV serum in the absence or presence of 10 μM CH-233191 for 12 hr (**C**). Control: ECM with 0.01% vehicle control (DMSO). The Holm-Sidak test was performed for all pairwise multiple comparisons. In **A** and **B**, \**P* < 0.03 vs. control; #*P* < 0.004 vs. NT-UV. n = 3 for control; n = 4-7 for serum samples. In **C**, \**P* < 0.0004 vs. control; ##*P* < 0.0001 vs. PE-UV. n = 3 in control; n = 6 in serum samples.

### ITE induces PE-like phenotypes in pregnant rats

To dissect the role of endogenous AhR ligands during pregnancy, pregnant rats were treated with ITE from gestational days 10-19. No animal death occurred throughout the study. ITE did not alter body weights of dams from GD10 to GD20 (Fig. 2A). On GD20, ITE increased weights of maternal lung, kidney, and liver, but not heart (Fig. 2B).

**Fig. 2.**
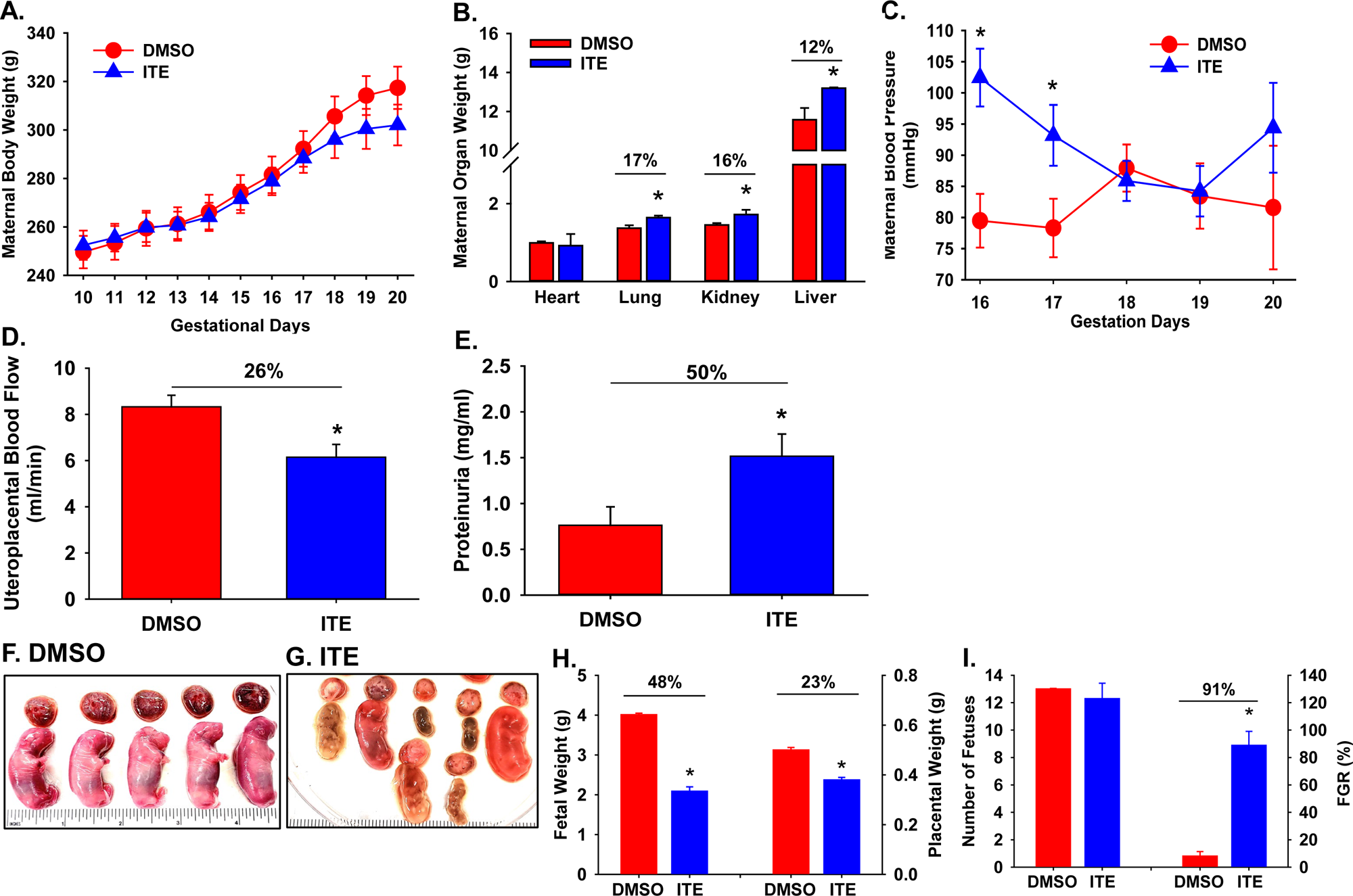
ITE dysregulates tissue growth and cardiovascular functions in pregnant rats. Pregnant rats were treated with ITE (5 mg/kg body weight/day) or vehicle control (DMSO) from GD10 to GD19. Maternal body (**A**) and organ weights (**B**) were recorded. n = 9 and 10 for ITE and DMSO groups, respectively. Maternal blood pressure (**C**; n = 6 and 5 for DMSO and ITE groups, respectively) and uteroplacental blood (**D**; n =4/group) were determined using the tail-cuff method and Vevo 2100 ultrasound system, respectively. Maternal proteinuria was measured (**E;** n =3/group). Placentas and fetuses were collected (n = 9 and 10 dams for ITE and DMSO groups, respectively) and weighed (**F-I**). The Mann-Whitney rank-sum test or Student *t*-test was performed to compare differences between ITE and DMSO. \**P* < 0.05 vs. DMSO (**B, D, E, H, I**) or DMSO at each corresponding time point (**C**).

The mean maternal blood pressure values in DEMO group were consistent with those previous reported^26,27^. ITE elevated maternal blood pressure by 22% and 16% on GD16 and GD17, respectively (Fig. 2C). ITE decreased uteroplacental blood flow, while elevated proteinuria (Fig. 2D) on GD20, indicating that ITE induced PE-like phenotypes in pregnant rats.

Compared to dimethyl sulfoxide (DMSO, the vehicle control), ITE caused morphological changes in placentas and fetuses (Figs. 2F-I). Many placentas and fetuses were smaller in size in ITE group (Figs. 2F and G). Some fetuses were degenerated or started to degenerate as indicated by their smaller size and dark appearance (Figs. 2F and G). The number of fetuses and placentas was similar between ITE and DMSO (Fig. 2I). However, ITE reduced fetal and placental weights on GD20 (Fig. 2H), which likely contributed to slight decreases in dams’ body weights on GD18-20 (Fig. 2A). When fetal body weights below 10th percentile of mean fetal weight (3.55 g) in DMSO group were designated as fetal growth restriction (FGR), ITE increased the FGR rate (Fig. 2I).

### ITE decrease vasculature in rat placentas

Compared to DMSO, ITE did not alter relative thicknesses of labyrinth and basal zones (Figs. 3a-c). However, ITE decreased relative areas of labyrinth zone, while increased (*P* < 0.05) relative areas of basal zone (Fig. 3d). ITE caused placental lesions with apparent decreases in the number of blood vessels and enlarged maternal sinusoids (Figs. 3e and f). ITE also induced formation of edematous stroma and patches of densely packed death cells and immune cell infiltration in labyrinth zone (Figs. 3e and f). No significant morphological change was seen in basal zone. Semi-quantitative analysis for CD31 immunostaining revealed that ITE reduced vascular area density and CD31 staining intensity in labyrinth zone (Fig. 3g). The TUNEL assay showed that abundant positive staining was present in rat placentas treated with DNase I (a positive control, Fig. 3h), but not in those treated without DNase I (Fig. 3i). A sporadic distribution of apoptotic cells was observed in a few placental tissue sections from ITE group, primarily present in basal zone (Fig. 3j). These data suggest that ITE mainly acts cells in labyrinth zone, primarily on endothelial cells and vasculature, leading to decreased cellular growth in labyrinth zone.

**Fig. 3.**
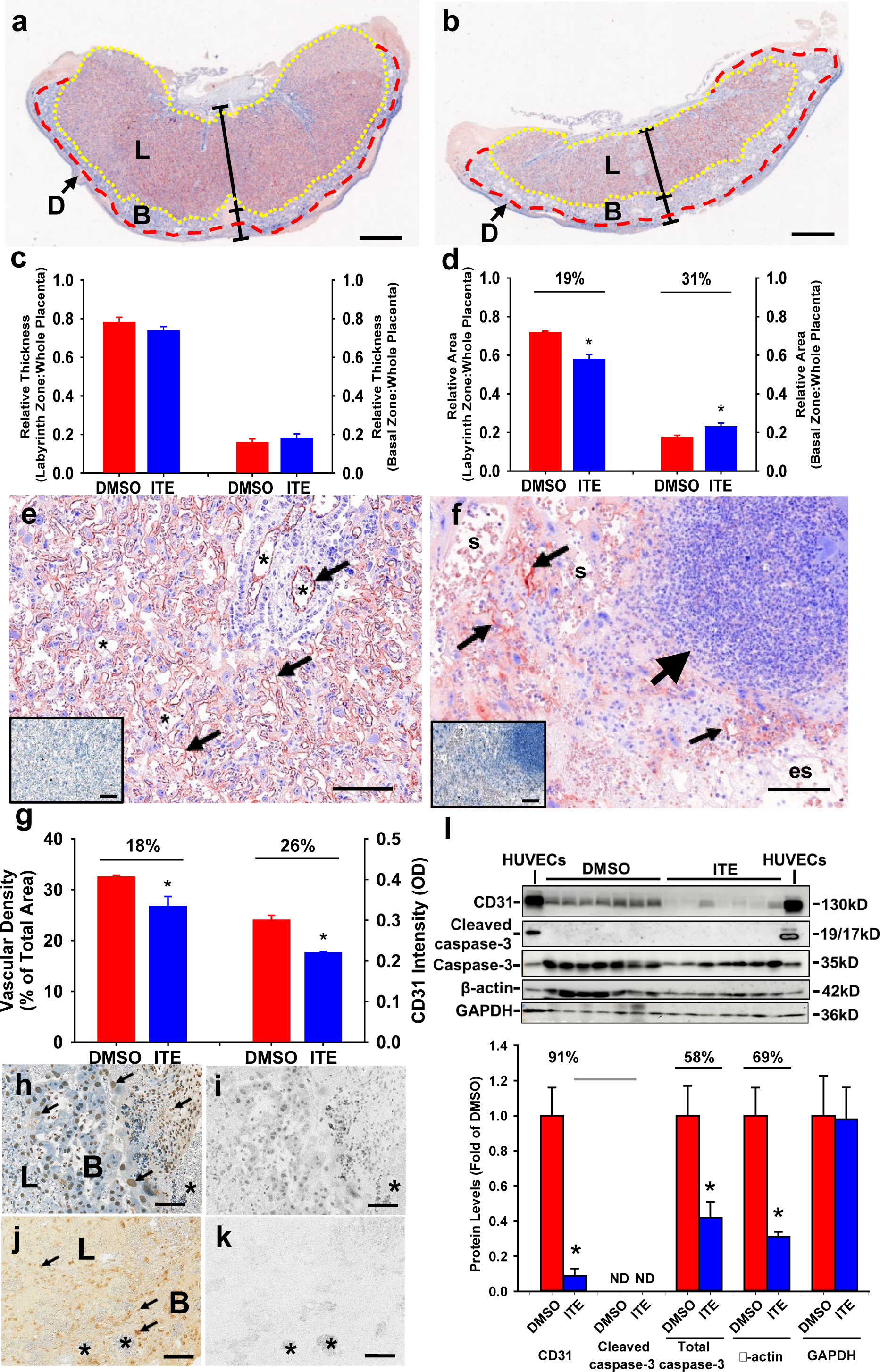
ITE decreases vascular densities and induces apoptosis in rat placentas. Pregnant rats were treated with ITE or DMSO from GD10 to GD19. Placentas were collected on GD20. (**a)**-(**g**) Placental morphological changes. The placental tissue sections were subjected to CD31 immunostaining followed by hematoxylin counterstaining. The representative images of L zones from ITE (**a**) and DMSO (**b**) groups are shown. Relative thicknesses and areas of L and B zones were quantified (**c, d**). Reddish indicates CD31 positive staining. Inserts in (**e**) and (**f**) are the adjacent tissue sections treated with normal goat IgG as a control. Blue: hematoxylin counterstaining; D: decidua basalis. Bar = 1 mm. (**e)**-(**g**) Vascular density and CD31 staining intensity in L zone. The representative images of CD31 immunostaining in L from ITE (**e**) and DMSO (**f**) groups were shown (**e**)-(**f**). Vascular area density and CD31 staining intensity were quantified (**g**). Small arrows: vascular endothelial cells. Large arrow: an area with a dense of death cells and immune cell infiltration. s: maternal sinusoids. es: edematous stoma. (**h**)-(**k**) Cell apoptosis. The tissue sections were subjected to TUNEL assay. The representative images of two adjacent placental sections from DMSO group (**h**)-(**i**). The sections were pre-treated with DNase I (a positive control) (**h**) or without (a negative control) (**i**) followed by slightly counterstaining with hematoxylin. (**j**) and (**k**) The representative images of two adjacent placental sections from ITE with (**j**) or without recombinant terminal deoxynucleotidyl transferase (**k**; a negative control). Arrows: apoptotic cells. *: blood vessels. Brownish nuclear DAB staining: positive apoptotic cells. Blue: hematoxylin counter staining. L: labyrinth zone. B: basal zone * *P* < 0.05 vs. DMSO. Bar = 100 µm. n = 6 placentas from different dams/group. Western blotting analysis for CD31 and cleaved caspase-3 (**l**). Data normalized to GAPDH are expressed as means ± SEM fold of the control. HUVECs at caspase-3 bands were treated with staurosporine (200 nM; a positive control for cleaved caspase-3). The Mann-Whitney rank-sum test or Student *t*-test was performed to compare differences between DMSO and ITE. \**P* < 0.05 vs. DMSO. n = 7 placentas from different dam/group.

Data from Western blotting is shown in Fig. 3l. Compared to DMSO, ITE reduced CD31 protein levels in placentas, consistent with decreases in vascular density and CD31 immunostaining intensity (Fig. 3g). ITE did not induce formation of cleaved caspase-3, while decreased total caspase-3. ITE reduced β-actin, but not GAPDH levels.

### ITE dysregulates transcriptome in rat placentas depending on placental sex

RNA-seq was performed to determine ITE-altered transcriptomic profiles. Compared to DMSO, ITE up- and down-regulated 409 and 907 genes in female (F) placentas, respectively, among which 69 are located on X chromosome (Fig. 4A and B; Table S2). ITE up- and down-regulated 571 and 1449 genes in male (M) placentas, respectively, among which 114 are located on X chromosome, and none is on Y chromosome. ITE commonly induced 926 differentially expressed genes (DEGs) in F and M placentas (Table S2). Of these DEGs, 188 and 731 were up- and down-regulated, respectively, in both ITE-F and ITE-M; 6 (LOC690483, Ugt2b15, LOC100910565, AABR07057442.1, U1, and Rfxapl1) were down-regulated in M, but up-regulated in F; one (ZFP280B) was down-regulated in F, but up-regulated in M; 136 are located on X chromosome (Table S2).

**Fig. 4.**
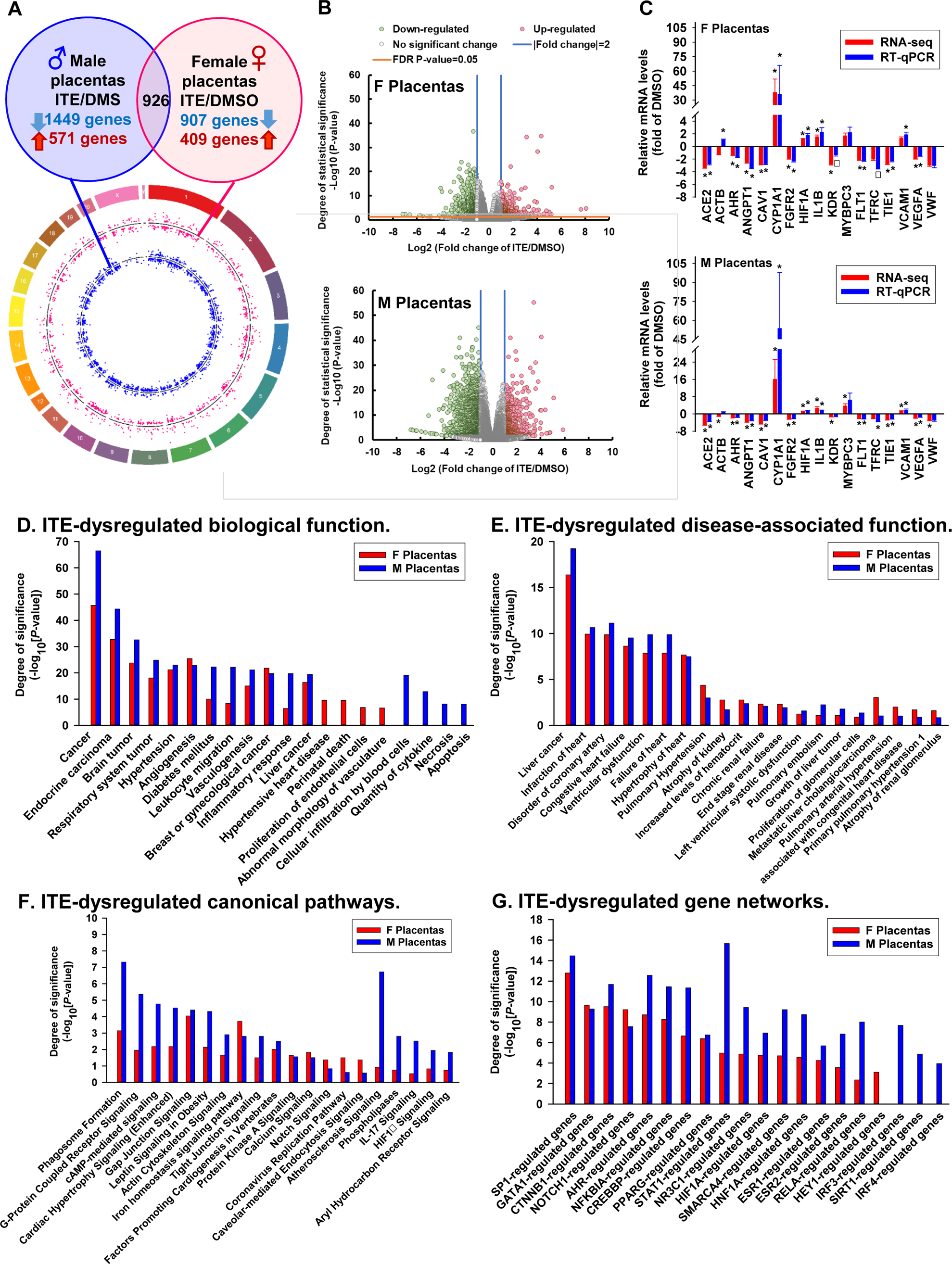
ITE dysregulates transcriptome and pathways in rat placentas. (**A**) Circos plot illustrating the chromosomal position of DEGs in F (red dots) and M (blue dots) placentas. Each dot represents one gene. The numbers and letters in the outer ring indicate the chromosomal location. For each scatter plot track, dots outside and inside of the centerline are up- and down-regulated genes, respectively. (**B**) Volcano plots showing DEGs in F and M placentas. Grey dots: no significant difference; Red and green dots: >2-fold up- and down-regulation, respectively (FDR-adjusted *P* < 0.05) in ITE vs. DMSO; *Differ (*P* < 0.05) from DMSO, n = 13, 16, 8, and 13 for DMSO-F, DMSO-M, ITE-F, and ITE-M, respectively. (**C**) RT-qPCR validation of ITE-dysregulated genes in F and M placentas. *Means differ (FDR-adjusted *P* < 0.05) from DMSO; ^γ^Means differ (0.1 > FDR-adjusted *P* > 0.05) from DMSO. n = 4/fetal sex/group. (**D**) Biological functions. (**E**) Disease-associated biological functions. (**G**) Canonical pathways-associated genes. **(F)** Gene networks. Significant enrichments were determined using IPA software (*P* < 0.05, Fisher exact test). Dotted line: *P* = 0.05. F: female; M: Male.

Data from RT-qPCR and RNA-seq analyses were highly correlated (r = 0.999 and 0.982 for F and M placentas, respectively. *P* < 0.001) (Fig. 4C). In F placentas, ITE upregulated CYP1A1, HIF1A, and IL1B, whereas down-regulated ACE2, AHR, ANGPT1, CAV1, FGFR2, SFLT1, TIE1, and VEGFA (Fig. 4C). ITE did not alter the expression of MYBPC3, TFRC, and VWF. RNA-seq, but not RT-qPCR analysis showed that ITE down-regulated ACTB and KDR. RT-qPCR, but not RNA-seq analysis showed that ITE upregulated VCAM1. In M placentas (Fig. 4C), ITE up-regulated CYP1A1, HIF1A, IL1B, MYBPC3 and VCAM1, whereas down-regulated ACE2, AHR, ANGPT1, CAV1, FGFR2, SFLT1, TFRC, TIE1, VEGFA, and VWF. RNA-seq, but not RT-qPCR analysis revealed that ITE attenuated expression of ACTB and KDR in M placentas. It is noted that both RNA-seq and RT-qPCR analyses revealed that ITE robustly increased levels of CYP1A1 and decreased AhR in F and M placentas, indicating activation of the AhR pathway.

Bioinformatics analysis on ITE-induced placental DEGs revealed that 637 biological functions were enriched (Fig. 4D; Table S3). Of these 637 functions, 363 (e.g., cancer, hypertension, angiogenesis, diabetes mellitus, vasculogenesis, leukocyte migration, inflammatory response, and vaso-occlusion) were commonly enriched in F and M placentas; 137 each were uniquely enriched in F (e.g., hypertensive heart disease, abnormal morphology of vasculature, perinatal death, and proliferation of endothelial cells) and M (e.g., cellular infiltration by blood cells, quantity of cytokine, necrosis, and apoptosis) placentas.

The ITE-induced placental DEGs were enriched in 249 diseases-associated functions (Fig. 4E; Table S4). Of these 249 functions, 132 were commonly enriched in F and M placentas, while 53 and 62 were only in F and M placentas, respectively. For example, diseases-associated functions including liver cancer, disorder of coronary artery, congenital heart failure, failure of heart, pulmonary hypertension, and atrophy of kidney were commonly enriched in F and M placentas (Fig. 4E). However, pulmonary embolism and growth of liver tumor were only enriched in F placentas, whereas pulmonary artery hypertension and atrophy of renal glomerulus were only enriched in M placentas (Fig. 4E).

The ITE-induced placental DEGs were enriched in 95 canonical pathways including phagosome formation, g-protein coupled receptor signaling, camp-mediated signaling, and gap junction signaling were enriched in placentas (Fig. 4F, Table S5). Of these 95 pathways, 35 were commonly enriched in both F and M placentas. These pathways included phagosome formation, g-protein coupled receptor signaling, cardiac hypertrophy signaling (enhanced), LPS/IL-1 mediated inhibition of RXR functions, gap junction signaling, tight junction signaling, protein kinase a signaling, and calcium signaling). Moreover, 3 pathways (notch signaling, coronavirus replication pathway, and caveolar-mediated endocytosis signaling) were solely enriched in F placentas, while 55 pathways including atherosclerosis signaling, phospholipases, IL-17 signaling, HIFα signaling, and aryl hydrocarbon receptor signaling were only enriched in M placentas.

Upstream regulator analysis identified 96 enriched gene networks, of which 81 were commonly enriched in F and M placentas, 1 and 14 were uniquely enriched in F and M placentas, respectively (Fig. 4G; Table S6). These common gene networks included NOTCH1, AHR-, HIF1A-, ESR1-, ESR2-, CREBBP-regulated genes, in which ESR1- (90 vs. 116 in F and M placentas) and CREBBP (86 vs. 108)-regulated genes contained the most abundant numbers of target genes (Table S6). In addition, HEY1-regulated genes were uniquely enriched in F placentas; IRF3-, SIRT1-, and IRF4-regulated genes were only enriched in M placentas (Fig. 4G).

### ITE dysregulates immune cell infiltration in rat placentas

Since PE is a pro-inflammatory state with aberrant immune cell functions and cytokine production^1^, we conducted CYBERSORTx analysis on the ITE-induced DEGs. CYBERSORTx analysis predicted that the top 8 immune cells with a fraction ≥ 2.8% in DMSO-treated placentas were dendritic cells resting, NK cells resting, macrophages M0, B cells naïve, T cells follicular helper, macrophages M2, T cells CD4 memory resting, and eosinophils (Tables S7 and S8). Six subtypes of cells (monocytes, plasma cells, T cells CD4 naïve, T cells regulatory, T cells CD4 memory activated, and mast cells resting) were ranging from 1.3% to 2.1%. Eight subtypes of cells (macrophages M1, mast cells activated, neutrophils, NK cells activated, dendritic cells activated, T cells gamma delta, T cells CD8, and B cells memory) were less than 1% or absent. Additionally, the fraction of macrophages M2 in NT-M and ITE-M was 44% and 38% higher than those in NT-F and ITE-F, respectively (Table S8).

ITE-dysregulated immune cell infiltration in placentas is shown in Fig. S1 and Table S8. Compared with DMSO, ITE increased fractions of plasma cells (2.1-fold of DMSO), monocytes (2.7-fold), macrophages M0 (1.2-fold), macrophages M2 (2.3-fold), and eosinophils (1.7-fold), whereas decreased T cells follicular helper (0.6-fold), NK cells resting (0.8-fold), and dendritic cells resting (0.6-fold) in rat placentas (Table S8). Within each placenta sex, ITE-altered immune cell fractions were like those in combined F and M placentas (Fig. S1), though changes in 3 cell subtypes (i.e., plasma cells and eosinophils in either F or M cells, and macrophages M2 in F cells) did not reach significant. This loss of significance is likely due to the reduced sample size in each placenta sex.

### ITE dysregulates cellular responses and transcriptome in HUVECs

We also investigated ITE’s effects on cellular function and profiled transcriptomic changes in HUVECs. The patients’ demographics are shown in Table S9. All clinical characteristics were similar between F and M cells. All patients except one in PE group underwent cesarean section delivery.

ITE similarly decreased cell proliferation and electrical resistance (or termed cell monolayer Integrity) in F and M HUVECs. As such, data from F and M HUVECs were pooled (Fig. 5). ITE dose-dependently inhibited cell proliferation, starting at 10 nM (87% of vehicle control) and reaching the maximum effect at 10 µM (76%) (Fig. 5A). The ITE-decreased electrical resistance was dose- and time-dependent (Fig. 5B), beginning at 15 and 20 hr for 1 and 10 µM of ITE, respectively and reaching the maximum effect up to 40 hr (Fig. 5B), indicating that ITE reduced cell monolayer integrity in HUVECs.

**Fig. 5.**
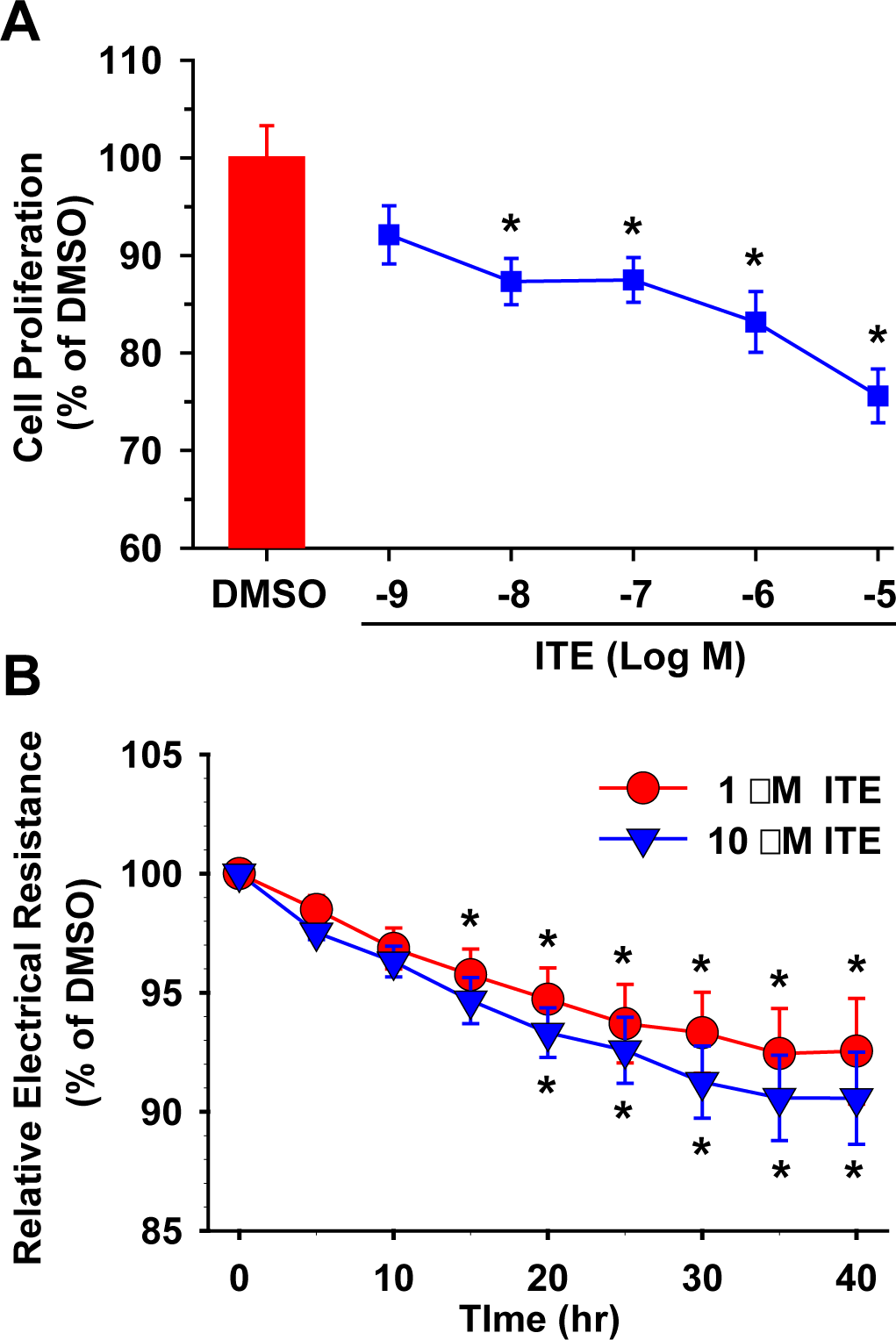
ITE decreases cell proliferation and monolayer integrity in HUVECs. (**A**) Cell proliferation. Sub-confluent cells seeded in 96 well plates were treated daily with ITE or DMSO in ECM for 48 h. Cell proliferation was determined using the crystal violet method. n = 6 individual cell preparations/cell sex. (**B**) Cell monolayer integrity. Confluent cells were treated with ITE or DMSO in ECM for up to 40 hr. Electrical resistance at 4000Hz was constantly recorded using the ECIS system. n = 4 individual cell preparations/cell sex. The Holm-Sidak test was performed for all pairwise multiple comparisons. *Means differ from DMSO (cell proliferation) or DMSO at corresponding time point (monolayer integrity). *P* < 0.05.

Compared with DMSO, ITE up- and down-regulated 93 and 47 genes in F HUVECs, respectively, of which 5 are located on X chromosome (Fig. 6A and B; Table S10). ITE up- and down-regulated 57 and 23 genes in M HUVECs, respectively. None of these DEGs is located on Y chromosome. ITE commonly induced 44 DEGs in F and M HUVECs. Of these 44 DEGs, 34 and 8 were up- and down-regulated, respectively, in F and M HUVECs; 1 (PALM2-AKAP2) was up-regulated in M, but down-regulated in F; 1 (AC048382.6, a IncRNA) was down-regulated in F, but up-regulated in M; one (DRP2) is located on X chromosome.

**Fig. 6.**
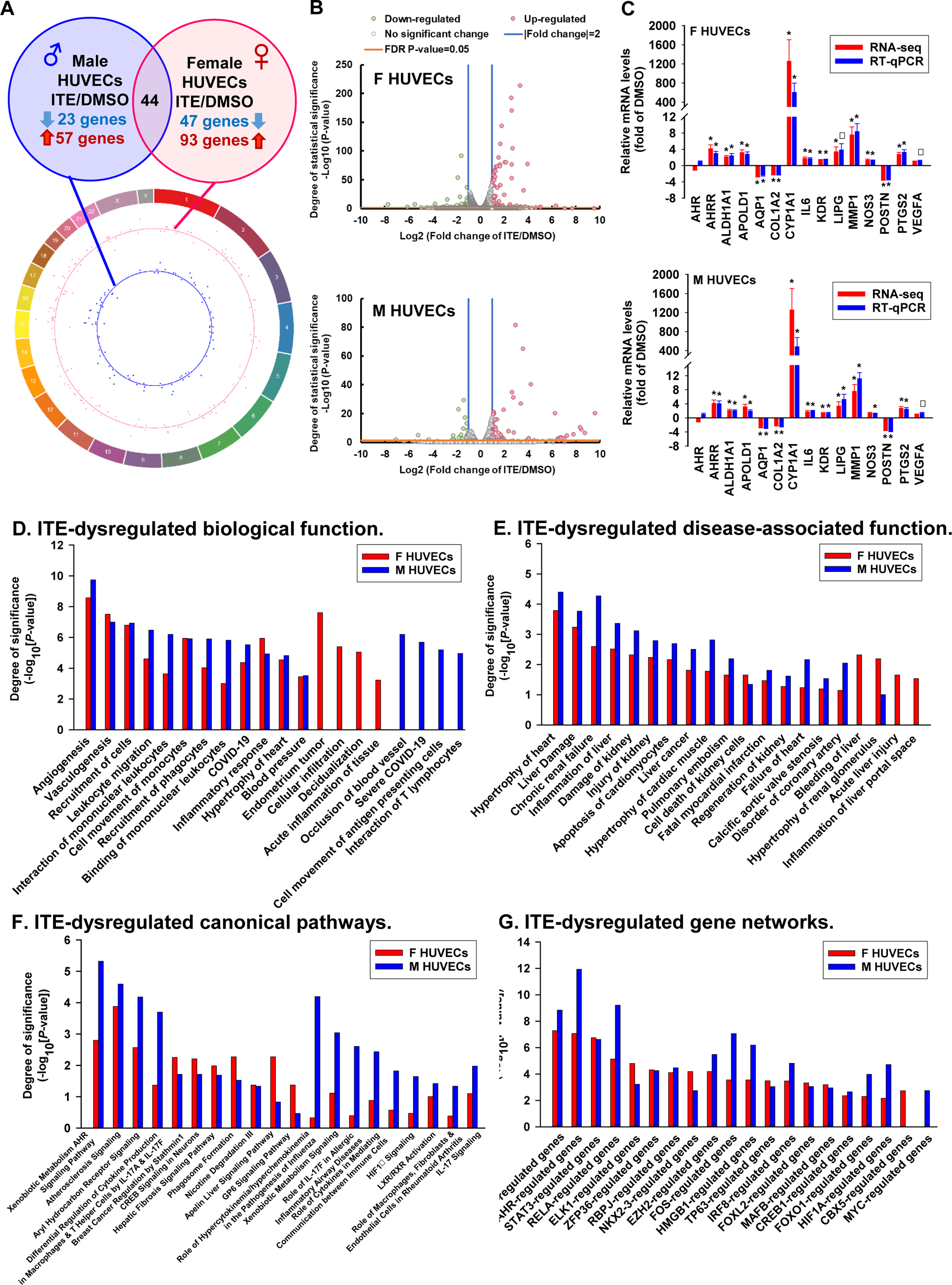
ITE dysregulates transcriptome and pathways in HUVECs. (**A**) Circos plot illustrating the chromosomal position of DEGs in F (red dots) and M (blue dots) cells. Each dot represents one gene. The numbers and letters in the outer ring indicate the chromosomal location. For each scatter plot track, dots outside and inside of the centerline are up- and down-regulated genes, respectively. (**B**). Volcano plots showing DEGs in F and M HUVECs. Grey dots: no significant difference; Red and green dots: > 2-fold up- and down-regulation, respectively (FDR-adjusted *P* < 0.05) in ITE vs. DMSO; n = 6 and 5 for F and M, respectively. (**C**) RT-qPCR validation of ITE-dysregulated genes in F and M HUVECs. *Means differ (FDR-adjusted *P* < 0.05) from DMSO, ^γ^Means differ (0.08 > *P* > 0.05) from DMSO, n = 4-6/fetal sex/group. (**D**) Biological functions, (**E**) Disease-associated biological functions, (**F**) Canonical pathways-associated genes, and (**G**). Gene networks. Significant enrichments were determined using IPA software (*P* < 0.05, Fisher exact test). Dotted line: *P* = 0.05. F: female; M: Male.

Data from RT-qPCR and RNA-seq analyses were completely correlated (r = 1.000 for F and M HUVECs. *P* < 0.001) (Fig. 6C). In F and M HUVECs, ITE upregulated AHRR, ALDH1A1,

APOLD1, CYP1A1, IL6, KDR, MMP1, NOS3, and PTGS, whereas down-regulated AQP1, COL1A2, and POSTN. RT-qPCR and RNA-seq analysis showed that ITE upregulated LIPG. RT-qPCR, but no RNA-seq analysis showed that ITE upregulated VEGFA. ITE did not alter expression of AhR. Either RT-qPCR or RNA-seq analysis showed that ITE elevated levels of CYP1A1 over 400-fold over vehicle control, indicating activation of the AhR pathway.

Biological function enrichment analysis on ITE-induced DEGs revealed that 719 biological functions were enriched in HUVECs (Fig. 6D; Table S11). Of these functions, 281 (e.g., angiogenesis, leukocyte migration, interaction of mononuclear leukocytes, binding of mononuclear leukocytes, inflammatory response, and hypertrophy of heart) were commonly enriched in F and M cells; 219 each were uniquely enriched in F (e.g., cellular infiltration, and acute inflammation of tissue) and M (e.g., occlusion of blood vessel, severe covid-19, cell movement of antigen presenting cells, and interaction of T lymphocytes) cells.

Diseases-associated biological function enrichment analysis revealed that the ITE-induced DEGs in HUVECs were enriched in 175 gene networks, many of which were associated with heart, liver, lung, and kidney diseases (Fig. 6E; Table S12). These gene networks included 88 commonly enriched in F and M HUVECs, 24 and 63 each solely in F and M HUVECs, respectively.

Canonical pathways enrichment analysis indicated that 35 pathways were enriched in HUVECs (Fig. 6F, Table S13). Of these 35 pathways, 9 (e.g., atherosclerosis signaling, aryl hydrocarbon receptor signaling) were enriched in both F and M HUVECs, while 2 (apelin liver signaling pathway and role of hypercytokinemia/hyperchemokinemia in the pathogenesis of influenza) and 24 (e.g., role of cytokines in mediating communication between immune cells, HIF1α signaling, and IL-17 signaling) exclusively in F and M HUVECs, respectively.

Upstream regulator analysis on ITE-induced DEGs in HUVECs predicted 20 enriched gene networks (Fig. 6G; Table S14). Of these 20, 18 were common between F and M HUVECs, including AHR-, HIF1A-, and STAT3-regulated genes; CBX5- and MYC-regulated genes were only enriched in F and M placentas, respectively. Three sets of key ITE-dysregulated signaling pathway genes (ITE-dysregulated AhR signaling pathway genes, cytokine production in macrophages and T helper cells by IL-17A and IL-17F pathway genes, and IL17 signaling pathway genes) are listed in Tables S15-17.

### ITE and PE alter immune cell-like phenotypes in HUVECs

To predict ITE-altered immune cell-like phenotypes in HUVECs, CYBERSORTx analysis was performed on the ITE-dysregulated genes in HUVECs. In DMSO-treated NT-HUVECs, 8 immune cell fractions were greater than 1% including T cells CD4 memory resting, dendritic cells activated, T cells CD4 naïve, mast cells resting, NK cells activated, macrophages M2, B cells naïve, and monocytes (Tables S18 and S19). However, compared with DMSO, ITE did not alter any immune cell fraction in HUVECs (Table S19).

We also reanalyzed the previously a published RNA-seq dataset from PE and NT HUVECs^7^. CYBERSORTx analysis inferred that in passage 0 (P0) NT-HUVECs, 11 immune cell fractions were greater than 1% (Tables S20 and S21). Of these immune cell-like phenotypes, 7 (T cells CD4 memory resting, dendritic cells activated, T cells CD4 naïve, mast cells activated, macrophages M2, B cells naïve, and monocytes) were identical to those in vehicle control-treated NT-HUVECs, implying that HUVECs have phenotypes of both innate and adaptive immune cells. Compared with NT, PE decreased the fractions of T cells CD4 naïve (0.6-fold of NT cells), whereas increased dendritic cells activated (1.7-fold) and master cells activated (2.9-fold) in F, but not M HUVECs (Fig. S2). Compared with their F counterparts, M HUVECs had a decrease in T cells CD4 naïve (0.6-fold of F cells) and an increase in dendritic cells activated (1.4-fold) (Fig. S2).

### ITE dysregulates common genes and pathways in rat placentas and HUVECs

We compared the ITE-induced common DEGs and pathways in rat placentas and HUVECs (Tables S22-26). Overall, 21 DEGs were commonly dysregulated in placentas and HUVECs (Table S22A). Of these DEGs, 9 and 5 were up- and down-regulated, respectively in placentas and HUVECs; 7 had different regulation directions in placentas and HUVECs. Additionally, 12 (Table S22B) and 20 (Table S22C) common genes were found in F and M placentas and HUVECs, respectively. However, the direction of regulation of these genes varied depending on tissues and cells. For example, CYP1A1/B1 and AHRR were both upregulated in placentas and HUVECs; PTGS2 was down- and up-regulated in placentas and HUVECs, respectively.

Of 316 biological functions identified in rat placentas and HUVECs, 118 were identical in both F and M placentas and HUVECs, including angiogenesis, blood pressure, carcinoma, cell movement, diabetes mellitus, disorder of pregnancy, vasculogenesis, and vaso-occlusion (Table S23). Eighty disease-associated functions were predicted in rat placentas and HUVECs (Table S24), including 31 overlapped in both F and M placentas and HUVECs (e.g., damage of kidney, enlargement of heart, fibrosis of heart, liver cancer, and left ventricular dysfunction). Nineteen canonical pathways were commonly enriched in placentas and HUVECs (Table S25). These pathways comprised 4 commonly enriched in both F and M placentas and HUVECs (e.g., breast cancer regulation by stathmin1, CREB signaling in neurons, hepatic fibrosis signaling pathway, and phagosome formation), and 14 (e.g., HIF1α signaling, HMGB1 signaling, IL-17 signaling, and LXR/RXR activation) solely in male placentas and HUVECs. Five gene networks (AHR-, CBX5-, HIF1A-, NKX2-3-, and REAL-regulated genes) were commonly enriched in placentas and HUVECs (Table S26).

### ITE dysregulates phosphoproteins and phospho-signaling pathways in HUVECs

To profile the ITE-regulated protein phosphorylation in HUVECs, bottom-up phosphoproteomic analysis was performed. The bottom-up phosphoproteomics detected 15,624 proteins (not shown), including a total of 2906 phosphosites with overall 78% at serine sites and 1285 phosphoproteins (Table 1). Top 10 of ITE dysregulated phosphosites in HUVECs at 4 and 24 hr are shown in Table 2. Of 3 phosphoproteins selected for verification (Table S27), Western blotting analysis only detected phospho(p) LMNAa/c (Ser390) and pHP1γ (Ser90) (Fig. S3A), but not pEEF2 (Thr57) in HUVECs. We observed that ITE significantly increased pLMNAα (S390) by 1.77-fold at 4 hr and pHP1γ (Ser90) by 1.61- and 2.14-fold at 4 and 24 hr, respectively. ITE did not alter pLMNAα (S390) at 24 hr, total (t) LMNAa/c and tHP1γ at 4 and 24 hr (Fig. S3B). Thus, p/t LMNA a/c and HP1γ data at 4 and 24 hr were used to determine the correlation between phophosproteomics and Western blotting analyses. The regression analysis showed that Western blotting and phjosphoproteomic data were highly correlated (r = 0.879; *P* < 0.0004) (Fig. S3B).

**Table 1.** ITE-regulated protein phosphorylation in HUVECs.

|  | 4 hr | 24 hr |
| --- | --- | --- |
| Total number of phosphosites/proteins detected | 2906/1286 |  |
| Number of up-regulated phosphosites | 86 (S=67, T=5, S/T=14) | 112 (S = 92; T=12, S/T= 8) |
| Number of upregulated phosphoproteins | 72 | 84 |
| Number of downregulated phosphosites | 21 (S=15, T=3, S/T=3) | 27 (S=20; T=6, S/T=1) |
| Number downregulated phosphoproteins | 20 | 21 |
| Commonly upregulated phosphoproteins/phosphosites at 4 and 24 hr |  |  |
| Commonly upregulated | 1. Death-associated protein 1 (DAP1), S51<br>2. Microtubule-associated protein 1B (MAP1B), S1396; S1400<br>3. Ras GTPase-activating protein-binding protein (G3BP1), S<br>4. Tight junction protein ZO-2 (TJP2), S163<br>5. 40S ribosomal protein S6 (RPS6), S240<br>6. Chromobox protein homolog 3 (CBX3), S93<br>7. Nuclear fragile X mental retardation-interacting protein 2 (NUFIP2), S629<br>8. Nestin (NES), S680<br>9. MAP7 domain-containing protein 1 (MAP7D1), S753<br>10. Eukaryotic translation initiation factor 4 gamma 1 (EIF4G1), S1068<br>11. Microtubule-associated protein RP/EB family member 1 (MAPRE1), S155<br>12. Serine/threonine-protein kinase Nek9 (NEK9), S<br>13. A-kinase anchor protein 12 (AKAP12), S/T<br>14. A-kinase anchor protein 12 (AKAP12), S696<br>15. Microtubule-associated protein 1B (MAP1B), S<br>16. A-kinase anchor protein 12 (AKAP12), S1331 |  |
| Commonly downregulated | 1. Programmed cell death protein 4 (PDCD4), S94<br>2. Vimentin (VIM), S430 |  |

\*HUVECs were treated with 1 μM ITE or equivalent DMSO for 4 and 24 hr. Phosphoproteins in lysates were enriched and subjected to the bottom-up phosphoproteomics. S: serine; T: threonine; S/T: serine or threonine. Fold change (ITE/DMSO) ≥ |1.2|, n = 4 cell preparations (2 female and 2 male cell preparations). Adjusted $P < 0.05$ .

**Table 2.**
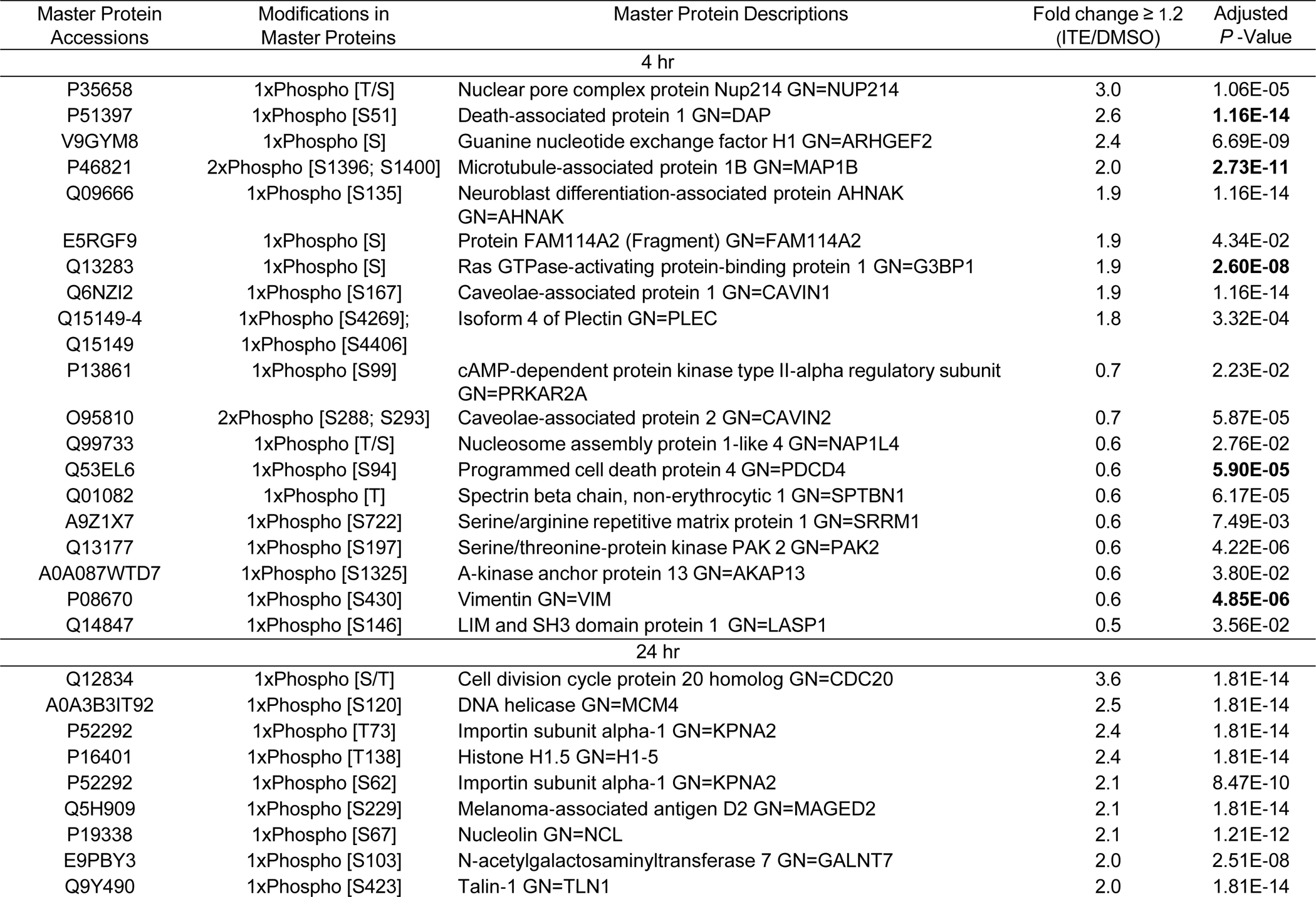
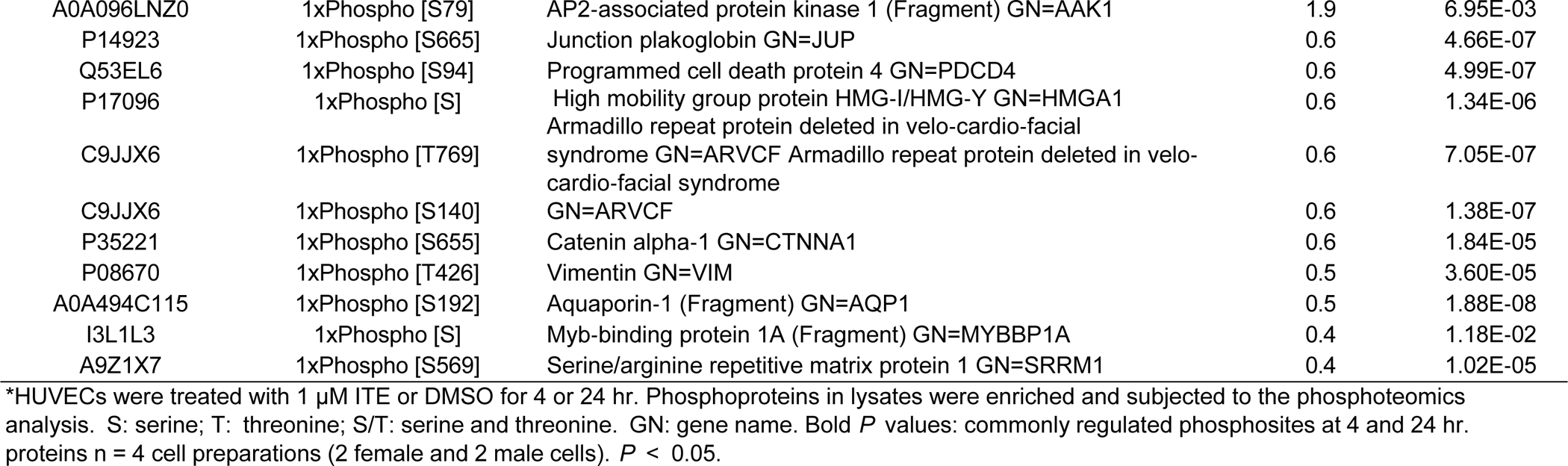
Top 10 of ITE-upregulated and -downregulated phosphosites in HUVECs at 4 and 24 hr.*

| Master Protein<br>Accessions | Modifications in<br>Master Proteins | Master Protein Descriptions | Fold change $\geq 1.2$<br>(ITE/DMSO) | Adjusted<br>P-Value |
| --- | --- | --- | --- | --- |
| 4 hr |  |  |  |  |
| P35658 | 1xPhospho [T/S] | Nuclear pore complex protein Nup214 GN=NUP214 | 3.0 | 1.06E-05 |
| P51397 | 1xPhospho [S51] | Death-associated protein 1 GN=DAP | 2.6 | **1.16E-14** |
| V9GYM8 | 1xPhospho [S] | Guanine nucleotide exchange factor H1 GN=ARHGEF2 | 2.4 | 6.69E-09 |
| P46821 | 2xPhospho [S1396; S1400] | Microtubule-associated protein 1B GN=MAP1B | 2.0 | **2.73E-11** |
| Q09666 | 1xPhospho [S135] | Neuroblast differentiation-associated protein AHNAK<br>GN=AHNAK | 1.9 | 1.16E-14 |
| E5RGF9 | 1xPhospho [S] | Protein FAM114A2 (Fragment) GN=FAM114A2 | 1.9 | 4.34E-02 |
| Q13283 | 1xPhospho [S] | Ras GTPase-activating protein-binding protein 1 GN=G3BP1 | 1.9 | **2.60E-08** |
| Q6NZI2 | 1xPhospho [S167] | Caveolae-associated protein 1 GN=CAVIN1 | 1.9 | 1.16E-14 |
| Q15149-4 | 1xPhospho [S4269]; | Isoform 4 of Plectin GN=PLEC | 1.8 | 3.32E-04 |
| Q15149 | 1xPhospho [S4406] |  |  |  |
| P13861 | 1xPhospho [S99] | cAMP-dependent protein kinase type II-alpha regulatory subunit<br>GN=PRKAR2A | 0.7 | 2.23E-02 |
| O95810 | 2xPhospho [S288; S293] | Caveolae-associated protein 2 GN=CAVIN2 | 0.7 | 5.87E-05 |
| Q99733 | 1xPhospho [T/S] | Nucleosome assembly protein 1-like 4 GN=NAP1L4 | 0.6 | 2.76E-02 |
| Q53EL6 | 1xPhospho [S94] | Programmed cell death protein 4 GN=PDCD4 | 0.6 | **5.90E-05** |
| Q01082 | 1xPhospho [T] | Spectrin beta chain, non-erythrocytic 1 GN=SPTBN1 | 0.6 | 6.17E-05 |
| A9Z1X7 | 1xPhospho [S722] | Serine/arginine repetitive matrix protein 1 GN=SRRM1 | 0.6 | 7.49E-03 |
| Q13177 | 1xPhospho [S197] | Serine/threonine-protein kinase PAK 2 GN=PAK2 | 0.6 | 4.22E-06 |
| A0A087WTD7 | 1xPhospho [S1325] | A-kinase anchor protein 13 GN=AKAP13 | 0.6 | 3.80E-02 |
| P08670 | 1xPhospho [S430] | Vimentin GN=VIM | 0.6 | **4.85E-06** |
| Q14847 | 1xPhospho [S146] | LIM and SH3 domain protein 1 GN=LASP1 | 0.5 | 3.56E-02 |
| 24 hr |  |  |  |  |
| Q12834 | 1xPhospho [S/T] | Cell division cycle protein 20 homolog GN=CDC20 | 3.6 | 1.81E-14 |
| A0A3B3IT92 | 1xPhospho [S120] | DNA helicase GN=MCM4 | 2.5 | 1.81E-14 |
| P52292 | 1xPhospho [T73] | Importin subunit alpha-1 GN=KPNA2 | 2.4 | 1.81E-14 |
| P16401 | 1xPhospho [T138] | Histone H1.5 GN=H1-5 | 2.4 | 1.81E-14 |
| P52292 | 1xPhospho [S62] | Importin subunit alpha-1 GN=KPNA2 | 2.1 | 8.47E-10 |
| Q5H909 | 1xPhospho [S229] | Melanoma-associated antigen D2 GN=MAGED2 | 2.1 | 1.81E-14 |
| P19338 | 1xPhospho [S67] | Nucleolin GN=NCL | 2.1 | 1.21E-12 |
| E9PBY3 | 1xPhospho [S103] | N-acetylgalactosaminyltransferase 7 GN=GALNT7 | 2.0 | 2.51E-08 |
| Q9Y490 | 1xPhospho [S423] | Talin-1 GN=TLN1 | 2.0 | 1.81E-14 |
| A0A096LNZ0 | 1xPhospho [S79] | AP2-associated protein kinase 1 (Fragment) GN=AAK1 | 1.9 | 6.95E-03 |
| P14923 | 1xPhospho [S665] | Junction plakoglobin GN=JUP | 0.6 | 4.66E-07 |
| Q53EL6 | 1xPhospho [S94] | Programmed cell death protein 4 GN=PDCD4 | 0.6 | 4.99E-07 |
| P17096 | 1xPhospho [S] | High mobility group protein HMG-I/HMG-Y GN=HMGA1 | 0.6 | 1.34E-06 |
| C9JJX6 | 1xPhospho [T769] | Armadillo repeat protein deleted in velo-cardio-facial syndrome GN=ARVCF | 0.6 | 7.05E-07 |
| C9JJX6 | 1xPhospho [S140] | Armadillo repeat protein deleted in velo-cardio-facial syndrome GN=ARVCF | 0.6 | 1.38E-07 |
| P35221 | 1xPhospho [S655] | Catenin alpha-1 GN=CTNNA1 | 0.6 | 1.84E-05 |
| P08670 | 1xPhospho [T426] | Vimentin GN=VIM | 0.5 | 3.60E-05 |
| A0A494C115 | 1xPhospho [S192] | Aquaporin-1 (Fragment) GN=AQP1 | 0.5 | 1.88E-08 |
| I3L1L3 | 1xPhospho [S] | Myb-binding protein 1A (Fragment) GN=MYBBP1A | 0.4 | 1.18E-02 |
| A9Z1X7 | 1xPhospho [S569] | Serine/arginine repetitive matrix protein 1 GN=SRRM1 | 0.4 | 1.02E-05 |

\*HUVECs were treated with 1 μM ITE or DMSO for 4 or 24 hr. Phosphoproteins in lysates were enriched and subjected to the phosphoteomics analysis. S: serine; T: threonine; S/T: serine and threonine. GN: gene name. Bold *P* values: commonly regulated phosphosites at 4 and 24 hr. proteins n = 4 cell preparations (2 female and 2 male cells). *P* < 0.05.

ITE time-dependently regulated protein phosphorylation (Tables S28 and S29). At 4 hr, ITE increased phosphorylation of 72 proteins, while decreased phosphorylation of 20 proteins (Table S28). At 24 hr, ITE enhanced phosphorylation of 84 proteins, while suppressed phosphorylation of 27 proteins (Table S29). Of these phosphoproteins, 16 and 2 were commonly up- and down-regulated, respectively at the same phosphosites at 4 and 24 hr, including proteins that are associated with microtubule, tight junction, kinase, and apoptosis (Table 1; Tables S28 and S29). The top 10 of these ITE-dysregulated phosphoproteins at 4 and 24 hr are shown in Tables 2 and 3, respectively. While most proteins exhibited 1 or 2 phosphosites induced by ITE, A-kinase anchor protein 12 (AKAP12) at 24 hr had 9 phosphosites, the largest number of phosphosites detected (Tables S28 and S29). No significant difference was detected in any total protein (Tables S30).

PANTHER analysis revealed that these ITE-dysregulated phosphoproteins were functionally classified into molecular functions, biological processes, cellular components, protein class, and pathway (Figs. S3 and S4). Many of these phosphoproteins were overrepresented (*P* < 0.05) in biological process, molecular functions, cellular component, and reactome pathway (Tables S31 and S32). Many of the notable ones are critical to cytoskeleton organization, cell division/death, protein binding, and signaling pathways (Tables S31 and S32).

## DISCUSSION

We have provided novel evidence that PE increases AhR activities in human fetal serum, and that ITE induces PE-like phenotypes in rats. ITE also dysregulates transcriptome and function- and diseases-associated genes and pathways in rat placentas and HUVECs in a fetal sex-specific manner. Moreover, ITE disrupts phosphoproteome and function-associated pathways in HUVECs. Collectively, these data suggest that dysregulation of endogenous AhR ligands may contribute to the pathophysiology of PE, though no longitudinal change in AhR activities is known during pregnancy.

The current finding on elevated AhR activities in PE-umbilical, but not maternal vein, serum indicates that the placenta is the major source for these increases, possibly via enhancing local synthesis and/or transportation of AhR ligands into the fetus. To date, it is unknown what exactly AhR ligands contribute to these PE-elevated AhR activities. However, given being primarily metabolized in placentas, beside livers and guts, increased Trp-derived AhR ligands^24,28,29^ may partially account for these increases^20,30^. Nonetheless, we should be cautious about these PE-elevated AhR activities in umbilical veins as the different gestational ages (Table S1) may be attributed to these increases.

Contradictorily to previous reports that a single dose of ITE failed to adversely affect fetuses and placentas in rats^31^ and mice ^32^, our data indicate that multiple doses of ITE reduce fetal and placental weights, and decreases placental vascular density and blood flow in rats as TCDD did in laboratory animals^33–36^. Thus, like TCDD, repeated exposure to endogenous AhR ligands impairs vascular growth in rat placentas.

Our finding that ITE induced PE-like phenotypes in rats suggests that dysregulation of endogenous AhR ligands could be a common contributor to the development of hypertension in general because AhR ligands also induce hypertension in M mice^18^ and rats^19^, and are involved in pulmonary artery hypertension in humans^19^. However, as PE does not alter vascular growth in human placentas^37,38^, different mechanisms may govern PE and ITE-induced hypertension in rats.

The ITE induced-transient increase blood pressure in pregnant rats observed in this study is dissimilar to the previous reports that repeated treatment of 3MC and FICZ elevated blood pressure for up to 4-8 weeks in M mice^18^ and rats^19^. Different AhR ligands with distinct potentials for AhR activation might partially explain these discrepancies^9^. However, it is more likely that sex- and pregnancy state-specific regulation of AhR activities in blood pressure exists since sex steroid hormones (e.g., estrogen and androgen) can crosstalk to the AhR pathway^9,39^. For example, increased estrogen during late pregnancy could antagonize AhR ligand-elevated blood pressure by interfering AhR transcriptional activity^9,39^ and enhancing vasodilatation^40^. Moreover, while ITE did not elevate blood pressure in near term rats, ITE did decrease uteroplacental blood flow, which could be due to the reduced placental vasculature observed.

The current study reveals the sexual dimorphisms of ITE-dysregulated transcriptome in placentas and HUVECs. Intriguingly, only 5% and 0% of DEGs are located on X chromosomes in F and M rat placentas, respectively (Fig. 4 and Table S2), and none of DEGs is on Y chromosomes. The same DEGs distribution on sex chromosomes is also seen in HUVECs (Fig. 6 and Table S10). Thus, genes on sex chromosome may not significantly contribute to the ITE-induced transcriptomic differences between F and M placentas and HUVECs as PE-induced transcriptomic differences^7^.

For rat placentas, one dose of TCDD has been shown to induce 182 DEGs using representational difference analysis and microarray technology^41^. The current study identified a set of ITE-induced fetal-sex specific DEGs, expanding the list of AhR ligand-induced DEGs in placentas. Additionally, of 182 TCDD-induced DEGs^41^, 29 are also identified in ITE-treated rat placentas with only 8 having the same regulatory trends (upregulated: *Cxcl10, Cxcl2, Cyp1a1, Cyp1b1, Mx1, Mx2,* and *Tgfbi;* downregulated: *Gata1*), suggesting differential regulation of ITE and TCDD in placental transcriptome.

It is noteworthy that M rat placentas had 54% more ITE-induced DEGs than their F counterparts, implying that M placentas are more susceptible or adaptive to the stimuli of endogenous AhR ligands. Additionally, the majority of these DEGs are downregulated in M (72%) and F (69%) placentas and are critical to vascular functions. For instance, decreases in angiotensin II (ANG II, a potent vasoconstrictor) receptors (*Agtr1a* and *Agtr2*) and *Ace2* (Fig. 4 and Table S2) might reduce ANG II activities, decreasing ITE-elevated blood pressure (Fig. 2C). Moreover, decreased expression of key endothelial regulators (e.g., *Angpt1, Cav1, Fgfr2, Kdr, Tie1,* and *Vegfa*) (Fig. 4 and Table S2) may contribute to the reduced placental vascular density.

Our bioinformatics analysis predicted that ITE differentially dysregulated many biological processes, disease-associated functions, pathways-associated genes, and gene networks between F and M placentas. Top biological processes include those relevant to cancer, tumor, hypertension, vascular formation and growth angiogenesis, and inflammatory response. ITE also dysregulated common genes between F and M placentas, which are primarily enriched in liver cancer, cardiovascular diseases, and kidney disease and are associated with many common signaling pathways (e.g., cardiac hypertrophy signaling [enhanced] and gap junction signaling). Notable is that ESR1 and CREBBP are the 2 most prominent hub genes in ITE-dysregulated gene networks (Table S6). Conversely, many other ITE-dysregulated processes and pathways were enriched uniquely in F and M placentas. For example, hypertensive heart disease was only enriched in F, while cellular infiltration by blood cells was predicted solely in M placentas. Moreover, notch signaling and caveolar-mediated endocytosis signaling are enriched in F, while atherosclerosis signaling, IL-17 signaling, and HIF1α signaling were in M placentas.

This study confirms ITE’s anti-cell proliferation activities^23^ and further shows ITE’s anti-endothelial integrity actions, possibly contributing to increased vascular edema in PE^1^. However, it is unknown why these ITE-induced cell responses are not cell sex-dependent.

Compared with rat placentas, ITE induced a fewer number of DEGs (80 vs 140 in M and F cells, respectively) in HUVECs. Contradicting to rat placentas, F HUVECs had 75% more DEGs than M cells, consistent with those in PE F HUVECs^7^. These data demonstrate that sex-specific susceptibility to endogenous AhR ligands differs between rat placentas and HUVECs. However, like those in rat placentas, the majority of DEGs in HUVECs were downregulated in M (71%) and F (66%) cells, although unlike rat placentas, 77% of 44 common DEGs are upregulated in HUVECs. Many of these DEGs are highly relevant to vascular functions. Specifically, ITE increased expression of AHRR (also elevated in rat placentas; Table S2), KDR, LIPG, MMP1, NOS3, PTGS2, and VEGFA (Fig. 6 and Table S10), most of which are pro-angiogenic. These data suggest their compensating roles in ITE-induced endothelial dysfunction. Besides these angiogenesis related DEGs, ITE also promoted expression of DEGs involved in inflammation, including IL6, IL1B, and PTGS2, which are indicative of immune response in HUVECs.

Like those in rat placentas, the ITE-dysregulated biological processes in HUVECs include angiogenesis, blood pressure, immune responses, and heart growth. This dysregulation is linked to disease-associated functions, canonical pathways, and gene networks. Specifically, ITE did dysregulate HIF1α signaling, STAT3 signaling, cytokine production, and IL-17 signaling, the latter of which is a crucial contributor to the pathogenesis of PE^42^.

A small portion of ITE-induced DEGs, pathways, and gene networks in HUVECs are shared with those in rat placentas (Tables S22-26). This is foreseeable since HUVECs used were a highly pure, single population of cells as opposed to placentas, a multicellular organ. However, these common DEGs, pathways, and gene networks are vital for AhR ligands-regulated biological processes. For example, CYP1A1/B1 are essential enzymes for initiating AhR ligands’ actions, while AHRR suppresses AhR activities^9,39^. Moreover, PTGS2 (also termed cyclooxygenase-2 [COX2]) (Table S22A) is well known for its roles in mediating inflammation, cardiovascular functions, and cancer development^43^.

The immune cell distribution in human placentas has been predicted using single-cell RNA-seq^44^, which projected the presence of T-cells resting and activated, NK cells, monocytes, and macrophages 1/2 with the most abundant amount of T-cells-activated (∼15-30% of total immune cells). Our current CIBERSORTx analysis also predicted a relatively high proportion of NK cells resting (15.5%), monocytes (2.1%), and macrophages M2 (5.6%) in vehicle control-treated rat placentas (Table S7). The current study, however, found that dendritic cells resting accounted for 29.8% of total immune cells with macrophages M0 (15.5%) and NK cells resting (15.5%) for another 31%. Together with the absence of mast cells activated, neutrophils, NK cells activated, and dendritic cells activated in rat placentas (Fig. S1, Tables S7 and S8), these data imply th at different distributions of immune cells exist between human and rat placentas and that rat near term placentas may represent a state of strong tissue repair.

Endogenous AhR ligands including ITE regulate proliferation and differentiation of immune cells (e.g., Th1, Th17, Treg, and dendritic cells) in mice^12,13^. Similarly, our data have shown that ITE alters infiltration of innate (e.g., monocytes, macrophages 0/2, NK cells resting, dendritic cells resting, and eosinophils) and adaptive (e.g., plasma cells and T cells follicular helper) immune cells in rat placentas. Our observation that ITE increased monocytes and plasma cells is consistent with the previous finding in PE-maternal circulation^45,46^. Notably, ITE decreased Tregs by ∼ 3-fold (Table S8), though this decrease did not reach significant, agreeing with the reduction of Tregs in circulations and tissues in PE^45,46^. The novel observation that ITE-increased macrophages 0/2 is important as macrophages 2 may participate in placental tissue repair after ITE induced tissue damage. Moreover, contradicting to decreased eosinophils in PE-maternal circulation^47^, ITE increased eosinophils in rat placentas. Thus, ITE likely partially mimics PE-altered immune cell distribution. Additionally, ITE appears more prone to maintain immune responses by increasing monocytes, plasma cells, and eosinophils, while decreasing Tregs in placentas. However, caution should be taken when interpreting these data since the signature matrix LM22 is based on a human dataset, and humans and rat share only 25% genomes though they share almost all disease-associated genes^48^. Additional limitations include that we cannot distinguish fetal and maternal origins of immune cells and classify these immune cells into subsets within each subtype.

One important finding of this study is that vehicle control-treated- and P0 NT-HUVECs had similar gene signatures of immune cells (Tables S18 and S21). Aside from having part of gene signatures of dendritic cells^49,50^, HUVECs exhibit part of gene signatures of T cells CD4 memory resting and naïve. Along the same line, HUVECs might also possess a small portion of gene signatures of other immune cells (e.g., mast cells, monocytes, macrophages, NK cells, and B cells) as HUVECs have a relative fraction of each of these immune cells greater than 1% (Tables S18 and S21). These data indicate the dynamic and plastic features of HUVECs and suggest that these endothelial cells might act partly as immune cells, particularly in producing cytokines and growth factors and affiliating antigen presentation, as suggested^49,50^.

Another interesting observation is that PE altered immune cell gene signatures of HUVECs in three subtypes of immune cells in F, but not M cells (Fig. S2). However, the increases in dendritic cells activated and mast cells activated in F cells suggest that PE might sex-specifically promotes pro-inflammatory activities of HUVECs. Additionally, NT M HUVECs had a gene signature of dendritic cells activated like that in PE F HUVECs, implying that NT M HUVECs may already possess activities of dendritic cells activated, comparable to that in PE F HUVECs. Unexpectedly, ITE did not alter immune cell gene signatures in HUVECs. One possible explanation is that a short duration (48 hr) of ITE treatment is insufficient to induce the changes in HUVECs seen in PE.

Mechanisms governing sexual dimorphisms of ITE-dysregulated placental and HUVECs responses remain elusive. Sex steroid hormones may partially control these dimorphisms because of differential expressions of aromatase between F and M placentas^51^ and differential distributions of circulating levels of gonadal hormones between F and M fetueses^52^.

Our current finding indicates that ITE rapidly altered phosphoprotein profiles in HUVECs. Notably, most of these ITE-induced changes are upregulation (∼80%; Table 1). In line with our previous report^23^, we also did not detect any significant changes in phosphorylation of ERK1/2, AKT1, c-Jun N-terminal kinase (JNK), and p38 mitogen-activated protein kinase, which could be due to a relative prolonged treatment time (e.g., hours rather than minutes). Our data suggest that phosphorylation of other proteins is also critically involved in the ITE-induced cellular responses. These proteins include those highly relevant to cell morphology and communication (e.g., microtubule-associated protein 1B and tight junction protein ZO-2, and vimentin), kinase activation (e.g., Serine/threonine-protein kinase Nek9 and A-kinase anchor protein 12), and cell growth (e.g., Ras GTPase-activating protein-binding protein and programmed cell death protein 4) (Table 1; Tables S27 and S28), which are supported by the bioinformatics analysis (Figs. S3 and S4) in biological process, molecular functions, cellular component, and reactome pathway (Tables S30 and S31), many of which are critical to cytoskeleton organization, cell division/death, aggrephagy, protein binding, and kinase inhibition (Tables S30 and S31).

In conclusion, we have shown that dysregulation of endogenous AhR ligands induces PE-like phenotypes in rats and impairs human endothelial function. These adverse effects are associated with sexually dimorphic dysregulation of transcriptome in placentas and HUVECs, and disrupted phosphoproteome in HUVECs. These data imply that the AhR ligand pathways might represent promising therapeutic and sex-specific targets for PE-associated vascular dysfunction.

## Acknowledgments

We would like to thank Ms. Lori Uttech-Hanson, from the Office of Grant Writing and Collaborative Project Development, University of Wisconsin–Madison for English editing. We are grateful to Qi-zi He, M.D., Ph.D., a pathologist at Department of Pathology, Shanghai First Maternity and Infant Hospital, Tongji University School of Medicine, Shanghai, China, for evaluating pathology of rat placental sections.

## Sources of Funding

This study was supported by the American Heart Association award 19CDA34660348 (CZ). The project was also supported by Translational Basic and Clinical Pilot Award (CZ, SK, and JZ) from the UW Institute for Clinical and Translational Research and the Clinical and Translational Science Award program, through the NIH National Center for Advancing Translational Sciences, grant UL1TR002373.

## Author contributions

YJZ, CZ, WL, KW, SK, and JZ conceived and designed this study. YJZ, CZ, YYW, SYZ, JSM, HHL acquired and analyzed data. YJZ, CZ, YYW, SYZ, JSM, KW, SK, and JZ interpreted data. YJZ, CZ, and JZ drafted the manuscript. HHL, WL, KW, and SK critically revised the manuscript. All authors approved the final version of the manuscript and agree to be accountable for all aspects of the work.

## Disclosures

None.

## Supplemental Materials

Supplemental Detailed Materials and Methods

Tables S1-S33

Figures S1-S4

Unedited gel blot images and bar chart.

## Graphic abstract

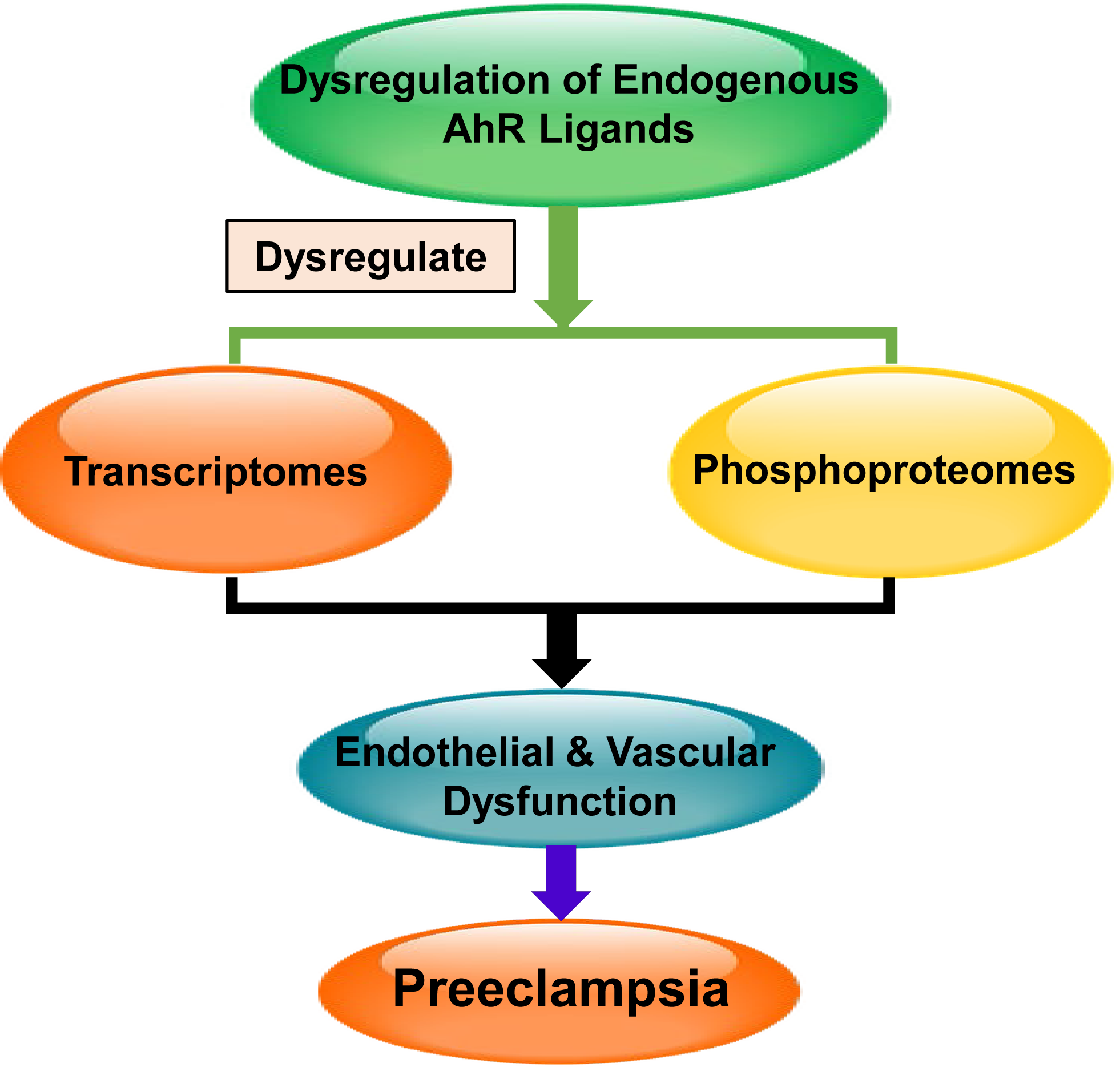

## Notes

### Competing Interest Statement

The authors have declared no competing interest.

### Summary of Updates

1. We combined tables 2 and 3 into a new Table 2. 2. We added a phrase "the period of maximal vascular adaptations" and a reference at Line 6-7, page 3 in supplementary materials. The order of reference is changed accordingly.

