## Supplementary materials for "An Endogenous Aryl Hydrocarbon Receptor Ligand Induces Preeclampsia-like Phenotypes: Transcriptome, Phosphoproteome, and Cell Functions"

**Short title:** AhR ligands induce preeclampsia.

**Key words:** AhR ligand, Preeclampsia, Vasculature, Endothelial cells, Sexual-dimorphism, Transcriptomics, Phosphoproteomics.

#### Supplementary Detailed Material and Methods

##### Ethical Approval

All procedures for human tissue collection were conducted in accordance to the Declaration of Helsinki. All subjects gave written, informed consent. Preeclampsia was defined according to the standard American College of Obstetricians and Gynecologists criteria. The blood sample collection was approved by the Scientific and Ethical Committee of Shanghai First Maternity and Infant Hospital, Tongji University, China (Protocol number KS21358). The umbilical cord collection was approved by the Institutional Review Board of UnityPoint Health-Meriter Hospital (Madison, WI) and the Health Sciences Institutional Review Boards of the University of Wisconsin-Madison (Protocol number 2004-006). The obstetricians at the UnityPoint Health-Meriter Hospital Birth Center where the umbilical cords were collected made the clinical diagnosis of PE and NT.

##### AhR activities in maternal and umbilical vein sera

Maternal and umbilical vein blood samples were collected from singleton pregnant women with NT and PE (Table S1) by the staff from the Tissue Bank of Shanghai First Maternity and Infant Hospital as described[29]. Maternal blood was collected from patients before delivery via venipuncture. Umbilical blood was collected immediately after delivery. Serum samples were prepared and stored at -80 $^{o}$C until assayed.[24]

RT-qPCR was conducted to determine expression of cytochrome P450 A1 and B1 (CYP1A1/B1) mRNA in HUVECs[23] as increased expression of CYP1A1/B1 is indicative of AhR activation[53]. Subconfluent HUVECs[54] were treated with 20% serum samples in endothelial cell media (ECM) for 12 or 24 hr. This ECM consists of ECM-b media (ScienCell, Cat # 1001-b) supplemented with 2% fetal bovine serum, 1% penicillin/streptomycin, and 37.5 mg/L endothelial cell growth supplement. To confirm AhR activities in sera, additional HUVECs were treated with 20% umbilical vein serum in the presence of 10 μM CH223191 (an AhR antagonist, cat # C8124; Sigma-Aldrich, St. Louis, MO) or dimethyl sulfoxide (DMSO, vehicle control, 0.01% v/v; the concentration used in the final CH223191 solutions) in ECM for 12 hr. Total RNA was extracted using TRIzol (Tiangen Biotech, Beijing, China) and quantified by a NanoDrop 2000C instrument (Thermo Scientific). Samples of total RNA (1 μg) from each treatment were reverse transcribed to cDNA. The reverse transcription was carried out using PrimeScript RT reagent Kit (TaKaRa, Dalian, China) for 15 min at 37 °C and 5 s at 85 °C in a 10 μl reaction volume. RT-qPCR was performed using SuperReal PreMix Plus (SYBR Green) (Tiangen Biotech). Primers for CYP1A1 (Sense: 5'-TCGGCCACGGAGTTTCTTC-3'; Antisense: 5'-GGTCAGCATGTGCCCAATCA-3'), CYP1B1 (Sense: 5'-AAGTTCTTGAGGCACTGCGAA-3'; Antisense:5'-GGCCGGTACGTTCTCCAAAT-3'), and GAPDH (Sense: 5'-GGAGCGAGATCCCTCCAAAAT-3'; Antisense: 5'-GGCTGTTGTCATACTTCTCATGG-3') were synthesized by Sangon Biotechnology (Shanghai, China). The RT-qPCR reaction was carried out as follows: 30 s at 95 °C for initial denaturation, and then 15 s at 95 °C and 32 s at 56 °C for 40 cycles. To confirm the amplification specificity, the PCR products were subjected to a melting curve analysis. Relative mRNA levels were analyzed using $2^{-\triangle\triangle CT}$ method and normalized to GAPDH.

##### Animals

All procedures of animal care and use were in accordance with National Institutes of Health guidelines (NIH Publication No. 85-23, revised 1996) with approval by the Institutional Animal Care and Use Committee at the University of Wisconsin at Madison (Protocol ID V005847). Timed-pregnant Sprague-Dawley rats (6-8 weeks old; n = 20) purchased from (Envigo Bioproducts, Madison, WI). All animals were housed in designated animal rooms under controlled conditions (lights on from 07.00 hours to 19.00 hours at 22°C-23°C). Standard rodent chow and water were available ad libitum throughout the experimental protocols. After acclimatization on gestational day (GD) 10, rats were randomly divided into treatment and control groups. Rats in the treatment group were daily injected via tail vein with ITE (5 mg/kg body weight [bw] per day, n = 10) in DMSO from GD 10 to GD 19. The control group received an equivalent of DMSO (n = 10). The physiological concentration of ITE is unknown in any species. The dose of ITE we used likely represents a pharmacological level since ITE levels are very low in tissues[55]. However, this dose is comparable to those that regulate placental expression of angiogenic factors[31,56], immune cell differentiation[12,22], and tumor growth[56]. Dam weights were recorded daily from GD 10 to 20. Blood pressure was monitored in rats (n = 6/group) from GD16 to GD20, the period of maximal vascular adaptations, using the tail-cuff method using a computerized CODA system (Kent Scientific Torrington, CT)[26,27]. For this set of animals, one rat was not pregnant in the ITE group and was removed from the study. Uteroplacental blood flow was measured on GD20 rats (n = 4/group) under isoflurane anesthesia and on a heated platform using a 30-MHz transducer and Vevo 2100 ultrasound system (Visual Sonics, Toronto, ON, Canada)[26]. Urine samples were collected on GD20 before blood flow measurement and stored at -80 $^{o}$C[57]. Uterine samples were centrifuged at 2500rpm for 3 min at 4 $^{o}$C. Urinary albumin levels were determined by ELISA using rat albumin ELISA kit (cat#: NR002, Nephrat, Exocell, Philadelphia, PA) according to the manufacturer's protocol.

**Rat tissue preparation**

Dams were sacrificed on GD 20 by $CO_2$ inhalation. Following euthanasia, maternal heart, liver, lung, kidney as well as each placenta and fetus were immediately collected and weighed. Fetal viability was not determined to avoid RNA degradation. Placental tissue samples were fixed in 4% buffered formaldehyde or snap frozen in liquid nitrogen and then stored at -80 $^{o}$C for the subsequent experiments.

**HUVECs for RNA sequencing (RNA-seq), functional assays, and phosphoproteomics**

HUVECs were isolated from umbilical veins of women with NT (Table S9) as described[7,23]. After isolation, cells were purified using CD31 Dynabeads and cultured in ECM under 37◦C, 5% CO2, and 95% air. HUVECs at passage 2 were daily treated with ITE (1 μM) or DMSO in ECM for 48 or up to 40 hr for RNA-seq[7] and functional assays[23], respectively, and 4 or 24 hr for phosphoproteomics.

**RNA isolation and quality control for RNA sequencing**

Total RNA samples were isolated from rat placentas and HUVECs using a RNeasy Mini Kit (Qiagen, Valencia, CA)[7,58]. The concentration and quality of each RNA sample were assessed using NanoDrop ND-2000 spectrophotometer (NanoDrop Technologies; Wilmington, DE) and Agilent 2100-bioanalyzer (Agilent Technologies; Santa Clara, CA). Only RNA samples with a high RNA integrity number (> 7.0) were used for RNA-seq and RT-qPCR analyses.

**Sex-typing of rat placentas**

RT-qPCR method was used to detect Y chromosome-specific sex-determining region Y (SRY) gene in rat placentas. The positive expression of SRY gene indicated the M placenta. RNA samples from both F and M placentas were prepared for RNA-seq.

**RNA-seq analysis and RT-qPCR verification**

RNA-seq was conducted at the UW-Madison Biotechnology Center. RNA libraries were generated using the Illumina TruSeq Stranded Total RNA Sample Preparation Kit (Illumina, San Diego, CA). In brief, 300ng of total RNA of each sample was fragmented using divalent cations under elevated temperature and purified. Double-stranded complementary DNAs (cDNAs) were synthesized and ligated to unique dual indexes adapters and cleaned up with two rounds of AMPure XP beads (0.8X). Adapter ligated DNA was amplified by PCR and cleaned up with AMPure XP beads (0.8X).

The quality and quantity of the libraries were assessed using an Agilent DNA1000 series chip assay (Agilent Technologies, Santa Clara, CA) and Invitrogen Qubit HS cDNA Kit (Invitrogen), respectively. Libraries were standardized to 2nM. Paired end 150bp sequencing was performed, using standard SBS chemistry on an Illumina NovaSeq6000 sequencer. Images were analyzed using bcl2fastq v2.17 software.

To validate RNA-seq data, RT-qPCR analysis was conducted using TaqMan Gene Expression Assays (Tables S33 and S34), TaqMan Fast Advanced Master Mix, and a StepOnePlus qPCR system (Life Technologies; Carlsbad, CA). For rat placentas and HUVECs, 15 genes with different expression patterns were selected. Data were normalized to internal housekeeping genes (GAPDH, YWHAZ, and RPLP0 for rat placentas; GAPDH, YWHAZ, and ACTB for HUVECs). The normalized data were analyzed using the $2^{-\Delta\Delta CT}$ method[7].

**Functional genomic analysis**

We obtained 67 to 121 million reads per sample from rat placentas and HUVECs, respectively, from the University of Wisconsin-Madison Biotechnology Sequencing Center. Reads were trimmed off TruSeq adapters using Skewer software with default parameters. The quality of individual trimmed sequence reads was evaluated by the University of Wisconsin-Madison Bioinformatics Resource Center. Phred quality score from per base sequence quality analysis averaged approximately 37, which is a conventional threshold denoting high-quality base calls. Trimmed reads were aligned to the GRCH38 genome using the STAR (Spliced Transcripts Alignment to a Reference) aligner, which is a splice-junction aware RNA-seq alignment algorithm. Consistent with sample-level RNA QC steps (Bioanalyzer 2100), gene body coverage indicated no evidence of RNA degradation. Mapped reads in each sample were counted using RNASeq by Expectation Maximization (RSEM) software. Empirical Analysis of Digital Gene Expression Data in R (EdgeR) software was utilized to determine the differentially expressed genes (DEGs; fold change (FC) > |2|, and Benjamini-Hochberg false discovery rate (FDR)-adjusted $P$-value < 0.05). Circos plots showing the location DEGs on chromosome sites were generated using Circa software (Circa Information Technologies, Downey, CA). Gene ontology analysis was performed to determine the enriched biological functions, diseases/canonical pathways, and gene networks using Ingenuity Pathway Analysis (IPA; www.qiagenbioinformatics.com).

**Analyses of immune cell infiltration and immune cell-like phenotypes by CIBERSORTx**

To infer ITE-altered immune cell infiltration in placentas, RNA-seq data on rat placentas were subjected to CIBERSORTx algorithm. As endothelial cells are considered conditional innate immune cells (i.e., antigen-presenting cells)[49,50], two RNA-seq data sets on HUVECs from the current study and our previous report for unpassaged (P0) NT and PE HUVECs[7] were analyzed to predict immune cell-like phenotypes of HUVECs. The signature matrix LM22 was used to define 22 subtypes of immune cells. RNA-seq data were uploaded to the CIBERSORTx web portal (https://cibersortx.stanford.edu/). The algorithm was run under B-mode with Batch correction set at 1000 permutations.

**Morphological and immunohistochemical analyses**

Fixed placentas were processed and embedded in paraffin. Immunolocalization of CD31 was detected as described. Two adjacent tissue sections (5-μm thick)/sample were mounted on one glass slide. The sections were deparaffinized, dehydrated, and boiled in a 10 mM citrate buffer solution (pH 6.5) in a microwave for 10 min for antigen retrieval. Endogenous peroxidase activity was quenched by immersing the tissue array in 3% hydrogen peroxide in methanol for 10 min. After blocking nonspecific binding sites, one tissue section per slide was probed with a CD31 goat polyclonal antibody (4 μg/ml, cat # AF3628, Novus Biologicals, Littleton, CO) for 1 hr. The second tissue section on the same slide was probed with the pre-immune goat IgG (4 μg/ml; cat # 55926, MP Biomedicals, Irvine, CA) as a negative control. After washing, the tissue sections were incubated with a biotinylated horse anti-goat antibody for 30 min. The specific immunoreactivity was visualized by 3-amino-9-ethylcarbazole (SK-4200, Vector Laboratories, Burlingame, CA).

The whole tissue sections were scanned at 20X using the Aperio Digital Pathology Slide Scanner system (Aperio-AT2, Leica Biosystems, Buffalo Grove, IL). Placental thickness of labyrinth (L) and basal (B) zones at the middle of cross-section of each tissue as well as areas of L (marked by yellow dotted lines) and B (marked by red dotted lines) were measured using NIH Image J software. Vascular area density (areas occupied by vessel wells) and CD31 staining intensity were analyzed using InForm 2.4 imaging software, a machine learning software (PerknElmer, Waltham, MA). For each tissue section, 10 areas were randomly photographed at 20X objective, which represented 21% of total L area. The whole tissue scanning, and CD31 image acquisition and imaging analysis were performed at Translational Research Initiatives in Pathology laboratory, University of Wisconsin-Madison.

**TUNEL assay**

Apoptotic cells in placental tissues were assayed using the DeadEnd Colorimetric TUNEL System (cat # G7130, Promega, Madison, WI). Briefly, the deparaffinized tissue sections were fixed in 10% buffered formalin followed by antigen retrieval (100 µg/ml proteinase K solution for 30 min). Tissue sections were then incubated with recombinant terminal deoxynucleotidyl transferase reaction mixture for 60 min at 37 C. Reaction was stopped by 2 × saline sodium citrate buffer. Endogenous horseradish peroxidase (HRP) was blocked by 1% hydrogen peroxide. HRP-streptavidin was added and incubated for 30 min. The tissue sections were incubated with diaminobenzidine until development of brownish staining (5-10 min).

**Cell proliferation**

Cell viability was assessed as described[56]. HUVECs seeded in 96-well plates (1000-2000 cells/well) were treated with 0.1 nM to 10 µM of ITE or vehicle control, dimethyl sulfoxide (DMSO, 0.01% v/v; the maximum concentration used in the final ITE solutions, Sigma, St. Louis, MO) in ECM for 2 days with daily change of media containing ITE or DMSO. The number of cells was determined using the crystal violet method.

**Cell monolayer integrity assay**

Cell monolayer integrity was determined using the ECIS Zθ+96-well array station (Applied BioPhysics, Troy, NY) using 96-well plates (96W10idf)[7,58]. Cells seeded (2 × $10^4$/well) were cultured for ~ 48 hr to allow cells to reach confluence. Cells were daily treated with ITE (1µM and 10µM) or DMSO in ECM (6 wells/treatment). The electrical resistance (indicating of the monolayer integrity) was real-time monitored for up to 25 hr.

**Bottom-up phosphoproteomics analysis**

Bottom-up phosphoproteomic analysis on HUVECs was performed at the UW-Madison Biotechnology Center. Subconfluent HUVECs at passage 3 (n = 4; 2 cell preparations/cell sex) were cultured in ECM. After 16 hr serum starvation, cells were treated with 1 µM ITE or DMSO in ECM for 4 and 24 hr to determine acute and prolonged effects of ITE on protein phosphorylation. Cells were lyzed in lysis buffer (cat. # 78501; ThermoFisher Scientific, Waltham, MA) supplemented with protease inhibits (cat. # PIA32953 ThermoFisher Scientific) and protein phosphatase inhibitors (cat. # 490684500; Sigma-Aldrich). Lysate proteins (100 µg/sample) were enzymatically digested using trypsin and LysC proteases and labeled using the TMTpro 16plex Label Reagent Set (cat #; A44521;ThermoFisher Scientific). A fraction of each enzymatically digested sample was pooled and used for total protein profiling. The majority of the samples were used for phosphopeptide enrichment with combinatorial TiO2 and ZrIMAC magnetic resins. Total and enriched phosphopeptides were scanned for abundance and identifications on the Thermo Scientific Orbitrap Fusion Lumos Tribrid Mass Spectrometer. Database searching, quantification and statistical analyses were performed on the Proteome Discoverer (version 2.5.0.400) informatics platform. Benjamini-Hochberg false discovery rate (FDR)-adjusted *P*-value < 0.05) was used to determine differential expression of phosphosites induced by ITE. ITE-regulated biological pathways were identified using the PANTHER Pathways Statistical Overrepresentation (Fisher's exact test, FDR *P* < 0.05) and the PANTHER list analysis Tools (PANTHER version 17.0 at http://www.pantherdb.org).

**Western blotting**

Western blotting was conducted as described[58]. Samples of rat placentas and HUVECs were lysed in lysis buffer (cat #: 39000, Thermo Fisher Scientific, Waltham, MA). Lysates were collected and centrifuged. Protein samples (20 or 25 µg/sample) were separated on 10 or 15% SDS-PAGE gels, and electrically transferred to polyvinylidene difluoride membranes. Membranes were probed with primary antibodies against target proteins (Table S27). Proteins were visualized by enhanced chemiluminescence (ECL) or ECL plus (Thermo Fisher Scientific) using Biomolecular Imager (Amersham ImageQuant 800, Cytiva, Marlborough, MA). Absorbance was measured using Image J (NIH, Bethesda. MD). Data were normalized to GAPDH (for rat placental tissue) or β-actin or GAPDH (for HUVECs). When appropriate, pooled HUVECs[23,59] were used as positive controls. For phosphoproteins, data were normalized to the total protein and GAPDH.

The number of commercially available antibodies against a specific phosphosite identified in HUVECs (Table S27) was extremely limited. To validate phosphoproteomics data, three antibodies against phospho-lamin a/c (LMNAa/c) (Ser390), -heterochromatin protein 1γ (HP1γ)/chromobox homolog 3 (CBX3) (Ser93), and -eukaryotic translation elongation factor 2 (EEF2) (Thr57) were chosen (Table S27). The corresponding total proteins were also probed.

**Statistical analyses**

All data except RNA-seq and phosphoproteomics data were analyzed using SigmaPlot software (Systat Software, San Jose, CA). For comparing two groups, data analyses were performed using the Mann-Whitney rank-sum test or Student *t*-test. For comparing multiple groups, data were analyzed using one-way or two ANOVA. When an F-test was significant, data were analyzed using the Dunnett's Method (Multiple comparisons versus control group) or the Holm-Sidak or Dunn's test (pairwise comparisons) when appropriate. The Benjamini-Hochberg FDR-adjusted *P*-value was used to decide significance for RT-qPCR and phosphoproteomics data. Correlations between RNA-seq and RT-qPCR data as well as phosphoproteomics and verification Western blotting data were analyzed using Pearson Product Moment Correlation Analysis. Data are represented as means ± SEM. Differences and correlations were considered significant when *P* < 0.05 unless stated otherwise.

**Supplementary Results**

**Supplementary figure legend**

**Fig. S1. ITE dysregulates immune cellular infiltration in rat placentas.** (**A**) Estimated fractions of 22 subtypes of immune cells in rat placentas treated with ITE or DMSO. (**B** and **C**) ITE-dysregulated infiltration in 8 subtypes of immune cells. Data are expressed as means ± SEM relative cell fractions. F: female; M: Male. The Holm-Sidak test was performed for all pairwise multiple comparisons. $^{\varepsilon}P < 0.03$ and $^{\gamma}P < 0.001$ vs. corresponding DMSO. $^{\delta}P < 0.001$ vs. corresponding female. n = 13, 16, and 8, and 13 for DMSO-F, DMSO-M, ITE-F, and ITE-M, respectively.

**Fig. S2. PE dysregulates immune cell-like phenotypes in P0-HUVECs.** (**A**) Estimated proportions of 22 subtypes of immune cells in P0-HUVECs from NT and PE using CIBERSORTx.

(**B**) PE-dysregulated immune cell phenotype changes. Data are expressed as means ± SEM relative cell fractions. The Two-way ANOVA with all pairwise multiple comparison was performed using the Holm-Sidak test. $^{\varepsilon}P < 0.03$ and $^{\gamma}P < 0.001$ vs. corresponding control (DMSO). $^{\delta}P < 0.001$ vs. corresponding female. n = 8, 8, 5, and 6 for DMSO-F, DMSO-M, ITE-F, and ITE-M, respectively. F: female; M: Male.

**Fig. S3. Western blotting verification of phosphoproteomics data in HUVECs.** Subconfluent cells were treated with 1 µM ITE or DMSO in ECM for 4 or 24 hr. Lysate proteins (20 µg/sample) were subjected to Western blotting for pLMNA ac (Ser 390) and pHP1γ/CBX3 (Ser 90) as well as their total proteins. Protein intensities were quantified. For total proteins, data were normalized to GAPDH. For phosphoproteins, data were normalized to corresponding total protein and GAPDH. All data are expressed as fold of DMSO ± SEM. Two-way t-tests were conducted. T: total; p: phosphor. HU1-4: HUVECs preparations 1-4. CON: DMSO. n = 4 cell preparations. $P < 0.05$ vs. corresponding DMSO.

**Fig. S4. Functional classification for ITE-dysregulated phosphoproteins in HUVECs.** Subconfluent cells were treated with ITE or DMSO in ECM for 4 (**I**) or 24 (**II**) hr. Lysate proteins (100 µg/sample) were enzymatically digested and labeled. Phosphopeptides were enriched and subjected to bottom-up proteomics analysis. The PANTHER software was used for further bioinformatics analysis. Gene ontology analysis was performed using Statistical Overrepresentation test (Fisher's exact test, FDR $P < 0.05$). Chart tooltips are read as Category name (Accession); # genes; Percent of gene hit against total # genes; Percent of gene hit against total # Function hits. n = 4 cell preparations. FDR $P < 0.05$.

Fig. S1. ITE-treated rat placentas.

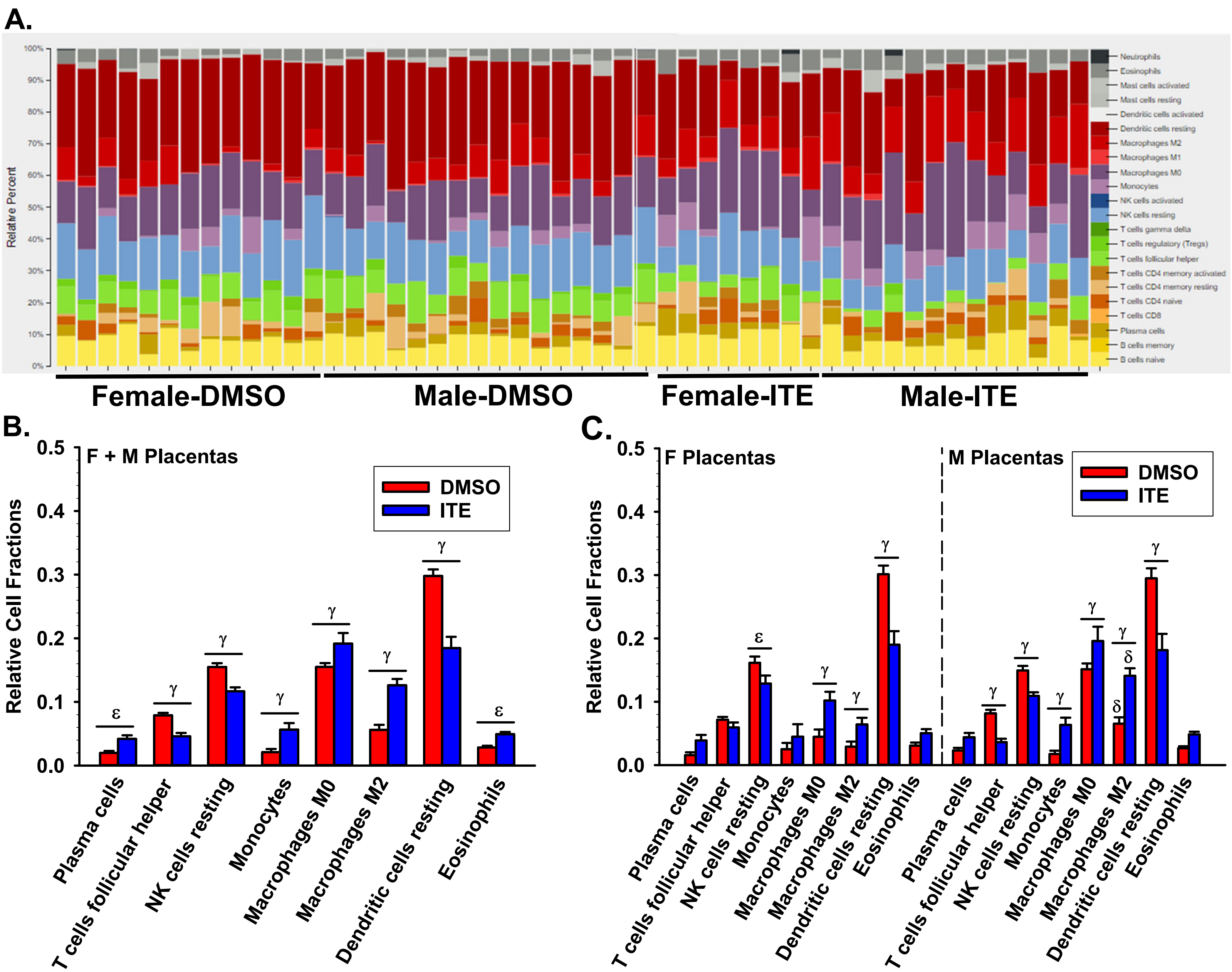

Fig.S2

A.

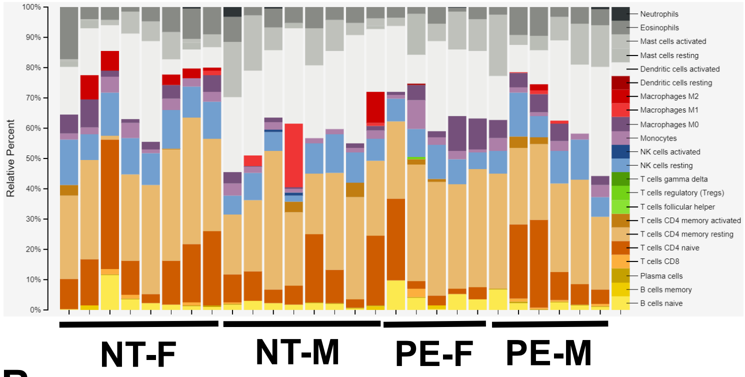
Relative Percent
100%
90%
80%
70%
60%
50%
40%
30%
20%
10%
0%
Neutrophils
Eosinophils
Mast cells activated
Mast cells resting
Dendritic cells activated
Dendritic cells resting
Macrophages M2
Macrophages M1
Macrophages M0
Monocytes
NK cells activated
NK cells resting
T cells gamma delta
T cells regulatory (Tregs)
T cells follicular helper
T cells CD4 memory activated
T cells CD4 memory resting
T cells CD4 naive
T cells CD8
Plasma cells
B cells memory
B cells naive
NT-F
NT-M
PE-F
PE-M

B.

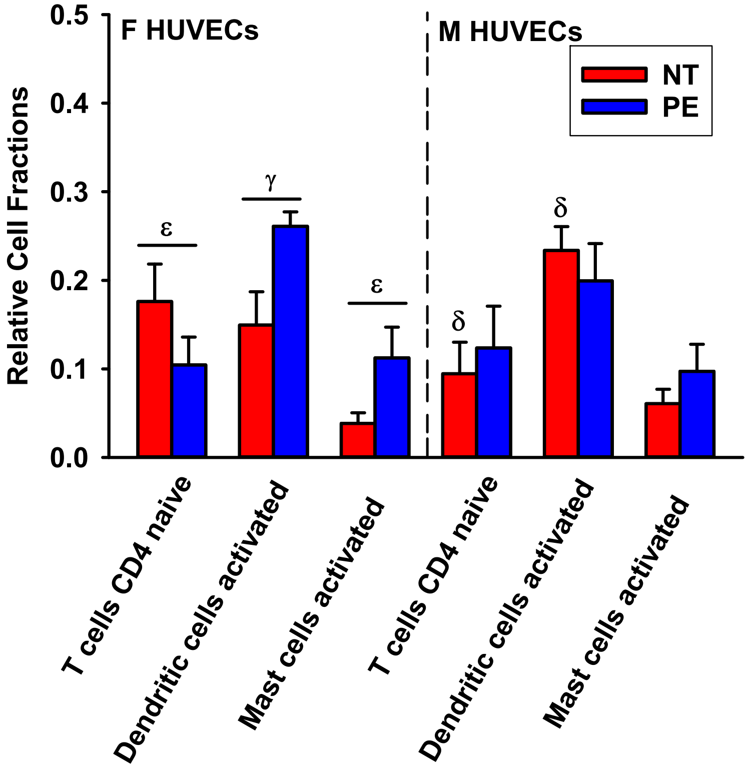
Relative Cell Fractions
0.5
0.4
0.3
0.2
0.1
0.0
F HUVECs
M HUVECs
NT
PE
ε
γ
ε
δ
δ
T cells CD4 naive
Dendritic cells activated
Mast cells activated
T cells CD4 naive
Dendritic cells activated
Mast cells activated

Fig. S3

**A. Western blot verification of phosphoproteomics data**

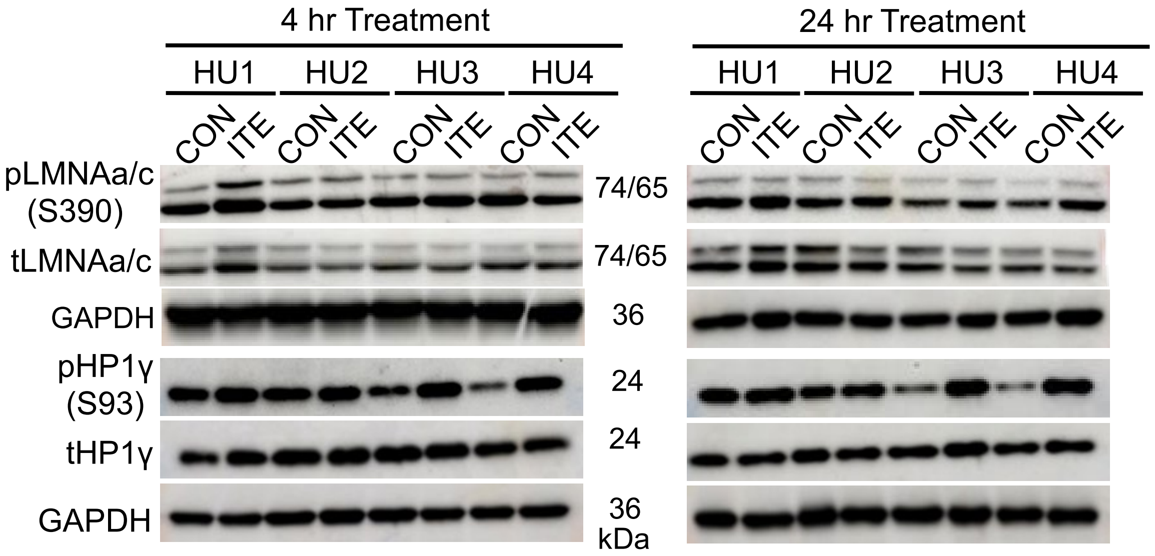


**B. Correlation**

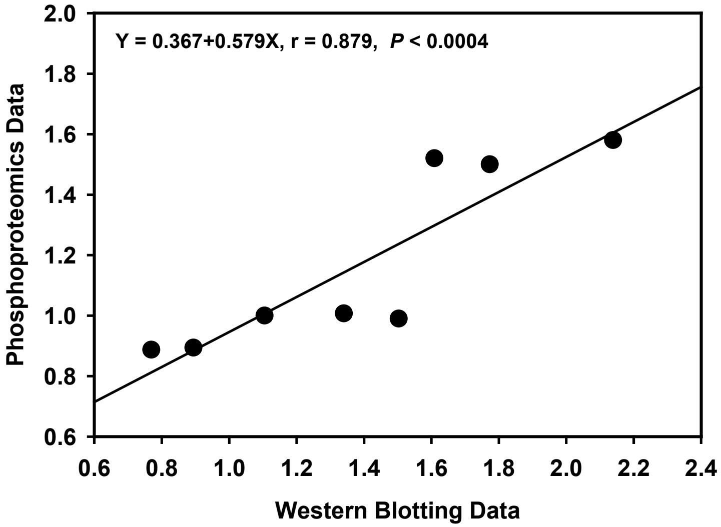

Fig. S4

## I. 4 hr

##### A. Molecular function

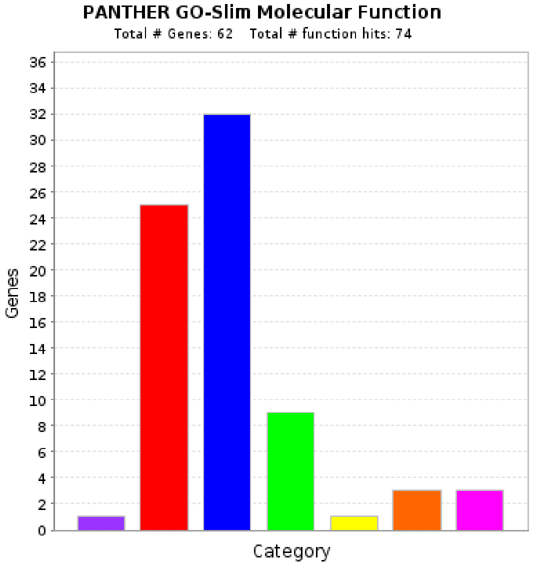


- ATP-dependent activity (GO:0140657): 1; 1.6%; 1.4%.
- No PANTHER category is assigned (UNCLASSIFIED): 25; 40.3%; 33.8%
- binding (GO:0005488) : 32; 51.6%; 43.2%
- catalytic activity (GO:0003824) : 9; 14.5%; 12.2%
- cytoskeletal motor activity (GO:0003774) : 1; 1.6%; 1.4%.
- molecular function regulator (GO:0098772) : 3; 4.8%; 4.1%
- structural molecule activity (GO:0005198) : 3; 4.8%; 4.1%

##### B. Biological process

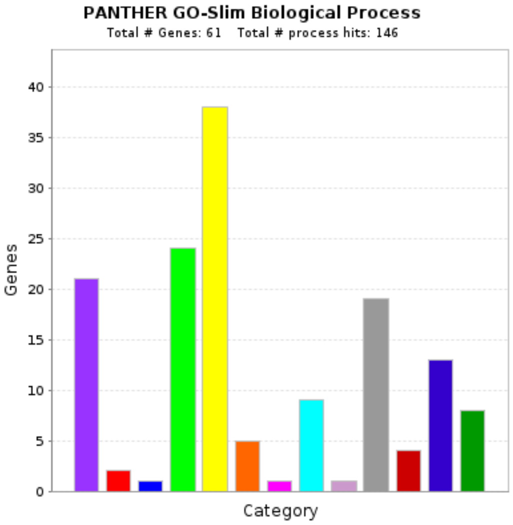


- No PANTHER category is assigned (UNCLASSIFIED) : 21; 34.4%; 14.4%
- biological adhesion (GO:0022610) : 2; 3.3%; 1.6%
- biological process involved in interspecies interaction between organisms (GO:0044419) : 1; 1.6%, 0.8%
- biological regulation (GO:0065007) : 24; 39.3%; 19.2%
- cellular process (GO:0009987) : 38; 62.3%; 30.4%
- developmental process (GO:0032502) : 5; 8.2%; 4.0%
- immune system process (GO:0002376) : 1; 1.6%; 0.8%
- localization (GO:0051179) : 9; 14.8%; 7.2%
- locomotion (GO:0040011) : 1; 1.6%; 0.8%
- metabolic process (GO:0008152) : 19; 31.1%; 15.2%
- multicellular organismal process (GO:0032501) : 4; 6.6%; 2.7%
- response to stimulus (GO:0050896) : 13; 21.3%; 8.9%
- signaling (GO:0023052) : 8; 13.1; 5.5%

#### C. Cellular component

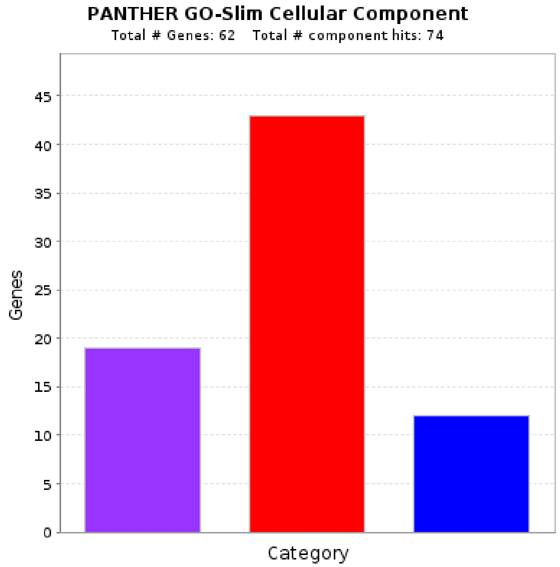


- No PANTHER category is assigned (UNCLASSIFIED): **19; 30.6%; 25.7%**
- cellular anatomical entity (GO:0110165): **43; 69.4%; 58.1%**
- protein-containing complex (GO:0032991): **12; 19.4%; 16.2%**

#### D. Protein class

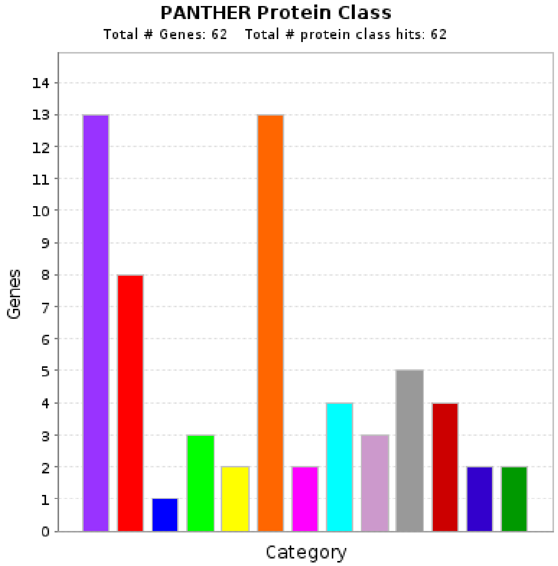


- No PANTHER category is assigned (UNCLASSIFIED) **: 14; 22.2%; 22.2%**
- RNA metabolism protein (PC00031) **: 8; 12.7%; 12.7%**
- cell junction protein (PC00070)**: 1; 1.6%; 1.6%**
- chaperone (PC00072) **: 3; 4.8%; 4.8%**
- chromatin/chromatin-binding, or -regulatory protein (PC00077)**: 2; 3.2%; 3.2%**
- cytoskeletal protein (PC00085)**: 13; 20.6%; 20.6%**
- membrane traffic protein (PC00150)**: 2; 3.2%; 3.2%**
- metabolite interconversion enzyme (PC00262)**: 4; 6.3%;6.3%**
- protein modifying enzyme (PC00260)**: 3; 4.8%; 4.8%**
- protein-binding activity modulator (PC00095)**: 5; 7.9%; 7.9%**
- scaffold/adaptor protein (PC00226)**: 4; 6.3%; 6.3%**
- translational protein (PC00263) **: 2; 3.2%; 3.2%**
- transporter (PC00227) **: 2; 3.2%; 3.2%**

#### E. Pathway

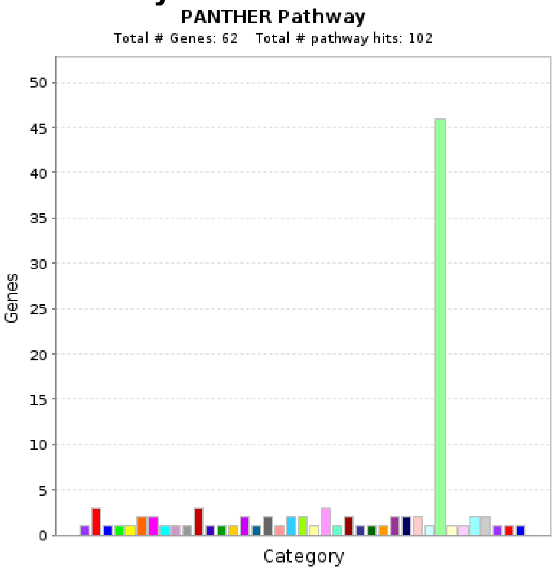


- 5HT1 type receptor mediated signaling pathway (P04373): 1; 1.60%; 1.00%
- Angiogenesis (P00005) : 3; 4.80%; 2.90%
- Axon guidance mediated by netrin (P00009) : 1; 1.60%; 1.00%
- Beta1 adrenergic receptor signaling pathway (P04377) : 1; 1.60%; 1.00%
- Beta2 adrenergic receptor signaling pathway (P04378) : 1; 1.60%; 1.00%
- CCKR signaling map (P06959) : 2; 3.20%; 1.90%
- Cytoskeletal regulation by Rho GTPase (P00016) : 2; 3.20%; 1.90%
- De novo pyrimidine ribonucleotides biosythesis (P02740): 1; 1.60%; 1.00%
- Dopamine receptor mediated signaling pathway (P05912) : 1; 1.60%; 1.00%
- EGF receptor signaling pathway (P00018) : 1; 1.60%; 1.00%
- Endothelin signaling pathway (P00019) : 3; 4.80%; 2.90%
- Enkephalin release (P05913) : 1; 1.60%; 1.00%
- FAS signaling pathway (P00020): 1; 1.60%; 1.00%
- FGF signaling pathway (P00021) : 1; 1.60%; 1.00%
- GABA-B receptor II signaling (P05731): 2; 3.20%; 1.90%
- General transcription by RNA polymerase I (P00022): 1; 1.60%; 1.00%
- Glycolysis (P00024) : 2; 3.20%; 1.90%
- Gonadotropin-releasing hormone receptor pathway (P06664) : 1; 1.60%; 1.00%
- Hedgehog signaling pathway (P00025): 2; 3.20%; 1.90%
- Heterotrimeric G-protein signaling pathway-Gi alpha and Gs alpha mediated pathway (P00026): 2; 3.20%; 1.90%
- Histamine H2 receptor mediated signaling pathway (P04386) : 1; 1.60%; 1.00%
- Huntington disease (P00029) : 3; 4.80%; 2.90%
- Hypoxia response via HIF activation (P00030): 1; 1.60%; 1.00%
- Inflammation mediated by chemokine and cytokine signaling pathway (P00031): 2; 3.20%; 1.9%
- Insulin/IGF pathway-protein kinase B signaling cascade (P00033) : 1; 1.60%; 1.00%
- Integrin signalling pathway (P00034): 1; 1.60%; 1.00%
- Metabotropic glutamate receptor group I pathway (P00041): 1; 1.60%; 1.00%
- Metabotropic glutamate receptor group II pathway (P00040): 2; 3.20%; 1.9%
- Metabotropic glutamate receptor group III pathway (P00039) :2; 3.20%; 1.9%
- Muscarinic acetylcholine receptor 2 and 4 signaling pathway (P00043):2; 3.20%; 1.9%
- Nicotinic acetylcholine receptor signaling pathway (P00044) : 1; 1.60%; 1.00%
- No PANTHER category is assigned (UNCLASSIFIED) : 47; 74.60% 45.60%
- Ras Pathway (P04393) : 1; 1.60%; 1.00%
- T cell activation (P00053) : 1; 1.60%; 1.00%
- Transcription regulation by bZIP transcription factor (P00055) : 2; 3.20%; 1.90%
- VEGF signaling pathway (P00056) : 2 ;3.20%; 1.90%
- p38 MAPK pathway (P05918) : 1; 1.60%; 1.00%
- p53 pathway feedback loops 2 (P04398) : 1; 1.60%; 1.00%
- p53 pathway (P00059) : 1; 1.60%; 1.00%

Fig. S4

## II. 24 hr

##### A. Molecular function

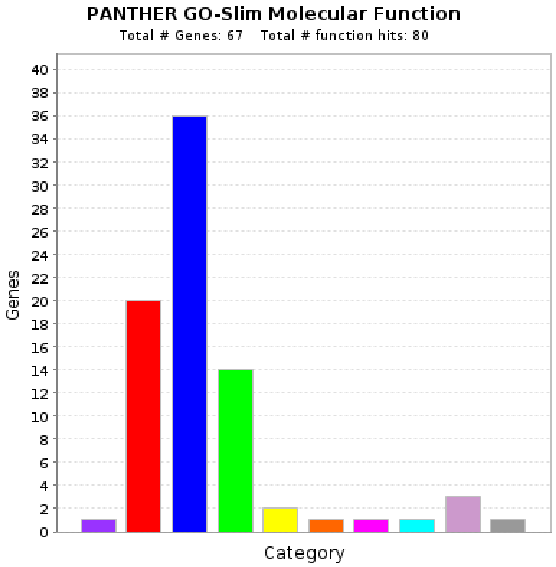


- ATP-dependent activity (GO:0140657): **1; 1.5%; 1.30%**
- No PANTHER category is assigned (UNCLASSIFIED): **20; 29.9%; 25.00%**
- binding (GO:0005488): **36; 53.7%; 45.00%**
- catalytic activity (GO:0003824): **14; 20.9%; 17.50%**
- molecular adaptor activity (GO:0060090): **2; 3%; 2.50%**
- molecular function regulator (GO:0098772): **1; 1.5%; 1.30%**
- molecular transducer activity (GO:0060089): **1; 1.5%; 1.30%**
- structural molecule activity (GO:0005198): **1; 1.5%; 1.30%**
- transcription regulator activity (GO:0140110): **1; 1.5%; 1.30%**
- translation regulator activity (GO:0045182): **3; 4.5%; 3.80%**

##### B. Biological process

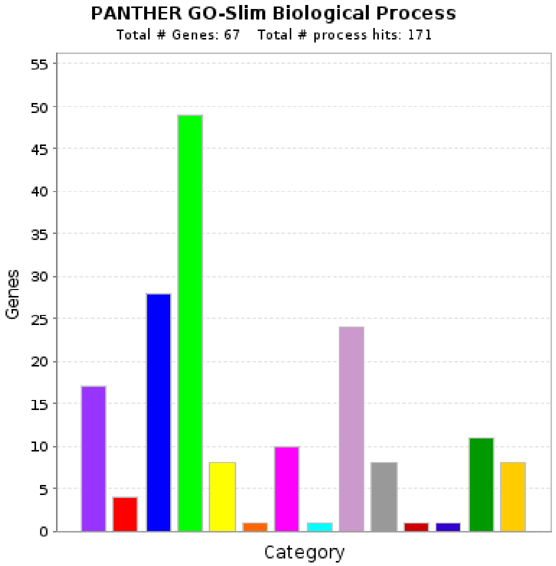


- No PANTHER category is assigned (UNCLASSIFIED): **17; 25.4%; 9.9%**
- biological adhesion (GO:0022610): **4; 6%; 2.3%**
- biological regulation (GO:0065007): **28; 41.8%; 16.4%**
- cellular process (GO:0009987): **49; 73.1%; 28.7%**
- developmental process (GO:0032502): **8; 11.9%; 4.7%**
- growth (GO:0040007): **1; 1.5%; 0.6%**
- localization (GO:0051179): **10; 14.9%; 5.8%**
- locomotion (GO:0040011): **1; 1.5%; 0.6%**
- metabolic process (GO:0008152): **24; 35.8%; 14%**
- multicellular organismal process (GO:0032501):**8; 11.9%; 4.7%**
- reproduction (GO:0000003): **1; 1.5%; 0.6%**
- reproductive process (GO:0022414): **1; 1.5%; 0.6%**
- response to stimulus (GO:0050896): **11; 16.4%; 6.4%**
- signaling (GO:0023052): **8; 11.9%; 4.7%**

**C. Cellular component**

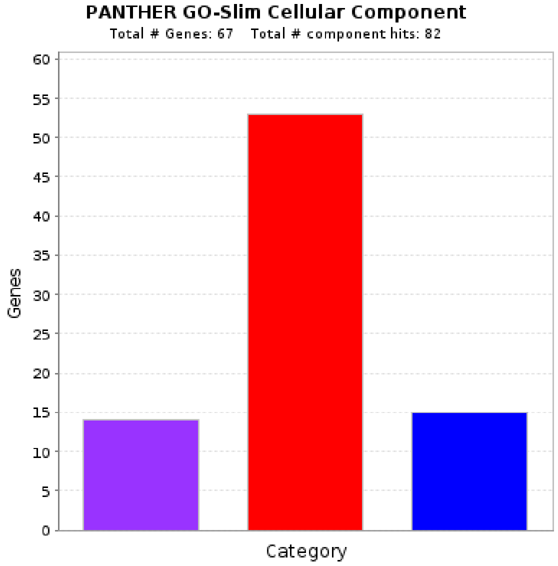


No PANTHER category is assigned (UNCLASSIFIED): 14; 20.9%; 17.7%
cellular anatomical entity (GO:0110165): 53; 79.1%; 64.6%
protein-containing complex (GO:0032991): 15; 22.4%; 18.3%

**D. Protein class**

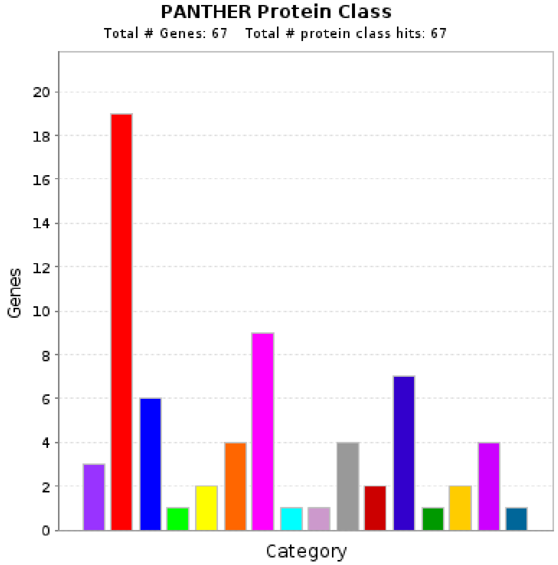


DNA metabolism protein (PC00009): 3; 4.5%; 4.5%
No PANTHER category is assigned (UNCLASSIFIED): 19; 28.4%; 28.4%
RNA metabolism protein (PC00031): 6; 9%; 9%
cell junction protein (PC00070): 1; 1.5%; 1.5%
chaperone (PC00072): 2; 3%; 3%
chromatin/chromatin-binding, or -regulatory protein (PC00077): 4; 6%; 6%
cytoskeletal protein (PC00085): 9; 13.4%; 13.4%
defense/immunity protein (PC00090): 1; 1.5%; 1.5%
gene-specific transcriptional regulator (PC00264): 1; 1.5%; 1.5%
membrane traffic protein (PC00150): 4; 6%; 6%
metabolite interconversion enzyme (PC00262): 2; 3%; 3%
protein modifying enzyme (PC00260): 7; 10.4%; 10.4%
protein-binding activity modulator (PC00095): 1; 1.5%; 1.5%
scaffold/adaptor protein (PC00226): 2; 3%; 3%
translational protein (PC00263): 4; 6%; 6%
transporter (PC00227): 1; 1.5%; 1.5%

#### E. Pathway

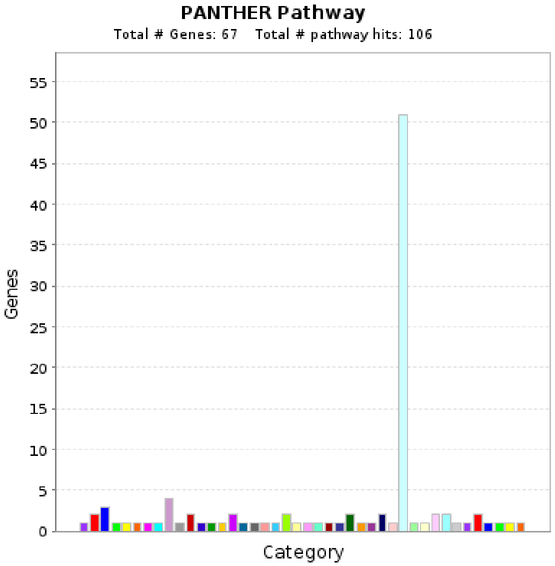


- 5HT1 type receptor mediated signaling pathway (P04373): 1; 1.5%; 0.9%
- Alzheimer disease-presenilin pathway (P00004): 2; 3%; 1.9%
- Angiogenesis (P00005): 3; 4.5%; 2.8%
- Angiotensin II-stimulated signaling through G proteins and beta-arrestin (P05911): 1; 1.5%; 0.9%
- Axon guidance mediated by semaphorins (P00007): 1; 1.5%; 0.9%
- B cell activation (P00010) : 1; 1.5%; 0.9%
- Beta1 adrenergic receptor signaling pathway (P04377): 1; 1.5%; 0.9%
- Beta2 adrenergic receptor signaling pathway (P04378): 1; 1.5%; 0.9%
- CCKR signaling map (P06959): 4; 6%; 3.8%
- Cadherin signaling pathway (P00012): 1; 1.5%; 0.9%
- Cytoskeletal regulation by Rho GTPase (P00016): 2; 3%; 1.9%
- De novo pyrimidine ribonucleotides biosythesis (P02740): 1; 1.5%; 0.9%
- Dopamine receptor mediated signaling pathway (P05912): 1; 1.5%; 0.9%
- EGF receptor signaling pathway (P00018): 1; 1.5%; 0.9%
- Endothelin signaling pathway (P00019): 2; 3%; 1.9%
- Enkephalin release (P05913): 1; 1.5%; 0.9%
- FAS signaling pathway (P00020): 1; 1.5%; 0.9%
- FGF signaling pathway (P00021): 1; 1.5%; 0.9%
- GABA-B receptor II signaling (P05731): 1; 1.5%; 0.9%
- Gonadotropin-releasing hormone receptor pathway (P06664): 2; 3%; 1.9%
- Hedgehog signaling pathway (P00025): 1; 1.5%; 0.9%
- Heterotrimeric G-protein signaling pathway-Gi alpha and Gs alpha mediated pathway (P00026): 1; 1.5%; 0.9%
- Histamine H2 receptor mediated signaling pathway (P04386): 1; 1.5%; 0.9%
- Inflammation mediated by chemokine and cytokine signaling pathway (P00031): 1; 1.5%; 0.9%
- Insulin/IGF pathway-mitogen activated protein kinase kinase/MAP kinase cascade (P00032): 1; 1.5%; 0.9%
- Integrin signalling pathway (P00034): 2; 3%; 1.9%
- Metabotropic glutamate receptor group II pathway (P00040): 1; 1.5%; 0.9%
- Metabotropic glutamate receptor group III pathway (P00039): 1; 1.5%; 0.9%
- Muscarinic acetylcholine receptor 2 and 4 signaling pathway (P00043): 2; 3%; 1.9%
- Nicotinic acetylcholine receptor signaling pathway (P00044): 1; 1.5%; 0.9%
- No PANTHER category is assigned (UNCLASSIFIED): 51; 76.1%; 48.1%
- PDGF signaling pathway (P00047): 1; 1.5%; 0.9%
- Pyrimidine Metabolism (P02771): 1; 1.5%; 0.9%
- Ras Pathway (P04393): 2; 3%; 1.9%
- T cell activation (P00053): 2; 3%; 1.9%
- Toll receptor signaling pathway (P00054): 1; 1.5%; 0.9%
- Transcription regulation by bZIP transcription factor (P00055): 1; 1.5%; 0.9%
- VEGF signaling pathway (P00056): 2; 3%; 1.9%
- Wnt signaling pathway (P00057): 1; 1.5%; 0.9%
- p38 MAPK pathway (P05918): 1; 1.5%; 0.9%
- p53 pathway feedback loops 2 (P04398): 1; 1.5%; 0.9%
- p53 pathway (P00059): 1; 1.5%; 0.9%

**Original Western blot images for Fig. 3I**

1. CD31

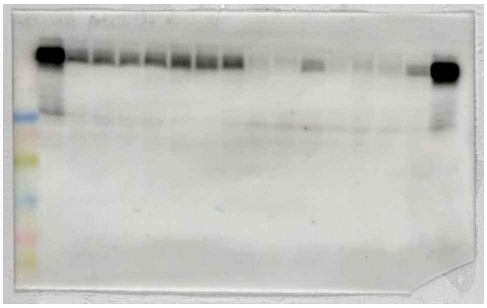

–130kD

1st Ab: CD31, 1:500 (R&D systems, cat #: AF3628); 2nd Ab: anti-goat 1:1000 (R&D systems, cat #: HAF017)

2. Cleaved caspase-3

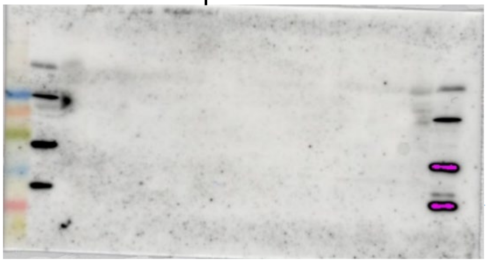

–19/17kD

1st Ab: cleaved cspase-3, 1:1000 (Cell signaling, cat #: 9661); 2nd Ab: anti-rabbit, 1:1000 (Sigma-Aldrich, cat #: NA934V)

3. Caspase-3

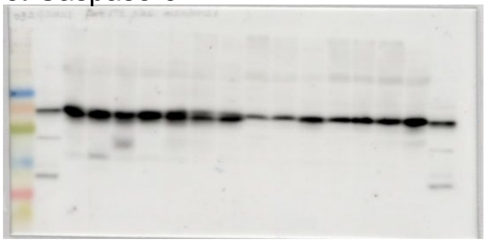

–35kD

1st Ab: total caspase-3 1:1000 (Cell signaling, cat #: 9665); 2nd Ab: anti-rabbit 1:1000 (Sigma-Aldrich, cat #: NA934V)

4. β-actin

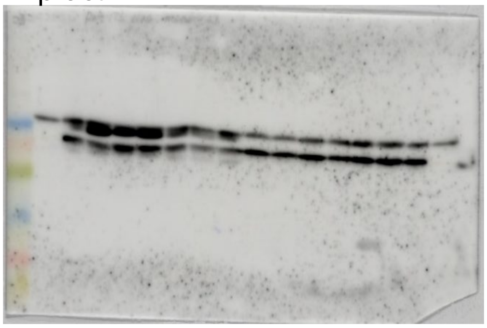

– 42kD

1st Ab: β-actin 1:4000 (Cell signaling, cat #: 4967); 2nd Ab: anti-rabbit 1:2000 (Sigma, cat #: NA934V)

5. GAPDH

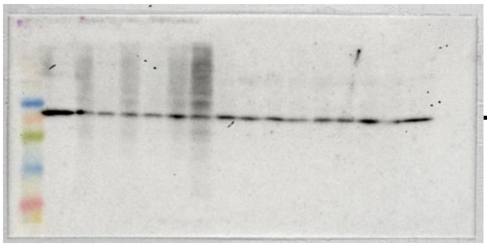

–36kD

1st Ab: rGAPDH , 1:3000 (Cell signaling, cat #: 5714; 2nd Ab: anti-rabbit, 1:10000 (Sigma, cat #: NA934V)

### Original Western blot images for Fig. S3

a. pLMNA a/c (S390). b. tLMNA a/c & GAPDH.

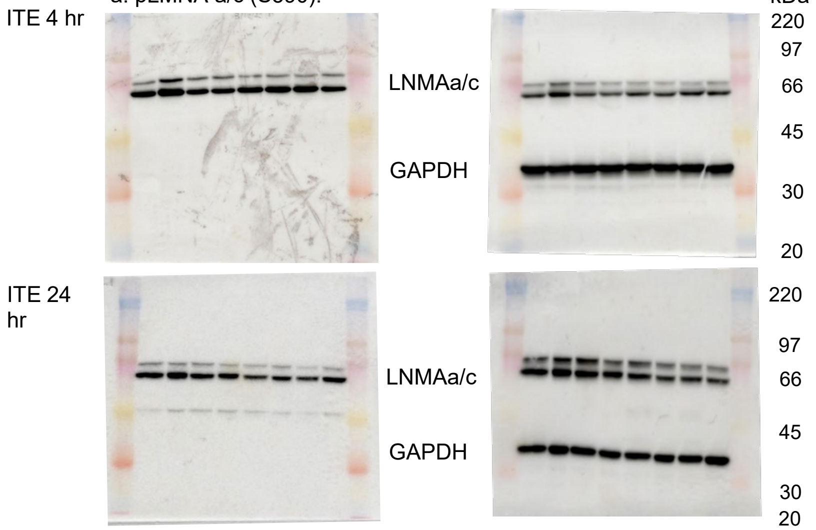


10% gel. Electrophoresis was run on 03/15/23. tLMNA a/c & GAPDH on 03/16/2023. After stripping, reprobed for pLMNA on 04/04/2023

12% gel. Electrophoresis was run on 04/11/23. pLMNA a/c After stripping, reprobed for tLMNA a/c & GAPDH on 04/12/2023.

c. phospho HP1γ (S93), total HP1, and GAPDH.

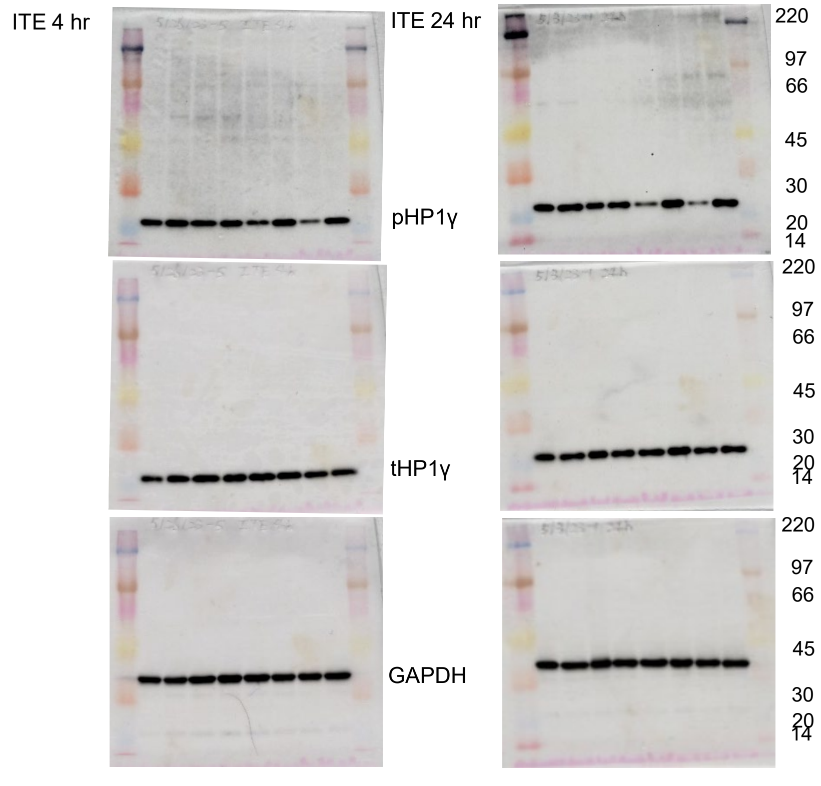


12% gel. Electrophoresis was run on 05/03/23 and 05/28/23 for pHP1 After striping, reprobed for tHP1 and GAPDH on 6/7/2023.

**For pLMNA (S390):**
$1^{O}$ ab: Rabbit pAB (Millipore-Sigma ABT1388 ) 1: 1000.
$2^{O}$ ab: Cytiva/Amrisham NNA934: goat-anti-rabbit IgG HRP-linked whole antibody 1:5000 or 10,000. 1 hr at RT.
**For tLMNA +GAPDH:**
$1^{O}$ ab: mouse monoclonal LMNAa/c ab (Sigma cat # SAB4200236): 1:5000 + mouse monoclonal GAPDH ab (Abnova cat H00002597-MO1) 1:5000 . 4C overnight.
$2^{O}$ ab : Cytiva/Amrisham NXA931V: sheep-anti-mouse IgG HRP-linked whole antibody 1:10,000K.1 hr at RT

**For pHP1γ (S93):**
$1^{O}$ ab: rabbit polyclonal pHP1γ (S93) (Abcam cat # ab45270): 1: 1000 . 4C overnight.
**For tHP1γ:**
$1^{O}$ ab: rabbit polyclonal tHP1γ (Cell Signaling cat # 2619): 1: 1;1000 tHP1γ ab ab . 4C overnight.
$2^{O}$ ab : Cytiva/Amrisham NA934V: donkey-anti-rabbit IgG HRP-linked whole antibody 1:10,000. 1 hr at RT;
**For GAPDH:** 1:10,000 as above
**Rainbow marker**: Cytiva High-Range Rainbow Molecular Weight Markers RPN756: 5 or 10 ul/lane
Images were recorded on AmershaM ImageQuant 800, Cytiva, Marlborough, MA.

**Original CIBERSORTx screenshots for Fig. S1A**

DMSO- and ITE-treated rat placentas.

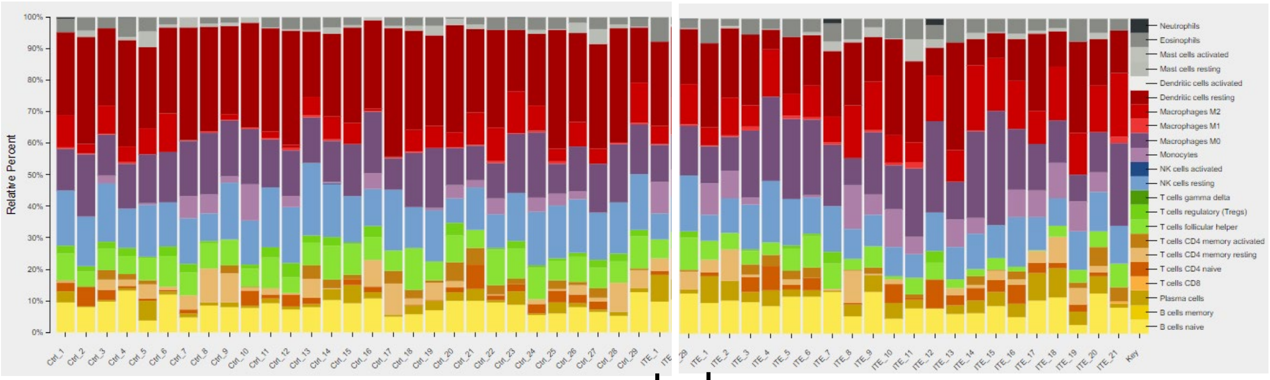


Stitched in the manuscript

Table S1. Patient demographic and clinical characteristics for serum AhR activity analysis*

| | Maternal Serum | | | Umbilical Vein Serum | | |
|---|---|---|---|---|---|---|
| Characteristics | NT (n = 7) | PE (n = 6) | *P* value | NT (n = 8) | PE (n = 12) | *P* value |
| Maternal age (years) | 30.9 ±1.86 | 33.5 ± 0.76 | 0.366 | 31.2 ± 1.35 | 31.9 ± 0.88 | 0.670 |
| Pre-pregnancy BMI (kg/m$^2$) | 21.3 ± 0.91 | 23.1 ± 0.61 | 0.139 | 20.9 ± 0.50 | 21.7 ± 0.51 | 0.297 |
| Gestational age (weeks) | 39.3 ± 0.23 | 36.4 ± 0.69 | 0.001 | 39.5 ± 0.32 | 37.8 ± 0.54 | 0.035 |
| Male fetus, n (%) | 2 (28.6%) | 3 (50.0%) | 0.428 | 3 (37.5%) | 4 (33.3%) | 1.000 |
| Birth weight (grams) | 3362.1 ± 147.64 | 2376.7± 196.42 | 0.002 | 3513.8 ±140.38 | 2941.7 ± 136.18 | 0.011 |
| Cesarean section, n (%) | 7(100%) | 6(100%) | - | 8(100%) | 12(100%) | - |
| Blood pressure (mmHg) | | | | | | |
| Systolic | 118.0 ± 1.94 | 155.2 ± 5.58 | 0.001 | 113.9 ± 3.94 | 141.8 ± 3.47 | 0.001 |
| Diastolic | 71.7 ±3.72 | 99.0 ± 3.63 | 0.001 | 69.6 ± 3.92 | 91.1 ± 1.92 | 0.001 |
| MAP | 87.1 ± 2.93 | 117.7 ± 3.51 | 0.001 | 84.4 ± 3.68 | 108.0 ± 2.28 | 0.001 |
| Proteinuria (mg/24hr) | - | 300.0 ± 63.25 | - | - | 683.3 ± 368.25 | - |

*All patients are Han Chinese without current or history of other major complications. All data are presented as means ± SEM or n (%). NT: normotensive pregnancy; PE: preeclampsia. BMI: Body mass index; MAP: mean artery pressure.

Table S2. ITE-dysregulated genes in rat placentas.*

| Gene ID | Gene location on sex chromosomes | Gene symbol | Female placentas | | Male placentas | | Comments |
|---|---|---|---|---|---|---|---|
| | | | log2 (ITE/Ctrl | *P*-Value | log2 (ITE/Ctrl) | *P*-Value | |
| ENSRNOG00000019500 | | CYP1A1 | 5.25 | 4.75E-14 | 4.01 | 1.37E-12 | DE in both female and male placentas |
| ENSRNOG00000055843 | | ZIK1 | 4.89 | 9.07E-03 | 4.73 | 1.51E-03 | DE in both female and male placentas |
| ENSRNOG00000037852 | | PIRT | 4.80 | 1.93E-04 | 3.41 | 4.17E-04 | DE in both female and male placentas |
| ENSRNOG00000040287 | | CYP1B1 | 4.34 | 2.85E-35 | 4.09 | 6.16E-42 | DE in both female and male placentas |
| ENSRNOG00000052033 | | SNORA73 | 4.14 | 9.41E-05 | 2.63 | 7.68E-03 | DE in both female and male placentas |
| ENSRNOG00000014721 | | AHRR | 4.05 | 5.18E-29 | 3.62 | 8.70E-32 | DE in both female and male placentas |
| ENSRNOG00000024709 | | VWDE | 4.01 | 5.48E-08 | 2.87 | 1.27E-05 | DE in both female and male placentas |
| ENSRNOG00000006367 | | SLC5A8 | 3.79 | 3.10E-08 | 2.05 | 6.71E-04 | DE in both female and male placentas |
| ENSRNOG00000020939 | | MS4A8 | 3.73 | 5.25E-05 | 2.27 | 3.35E-03 | DE in both female and male placentas |
| ENSRNOG00000048659 | | LOC498316 | 3.31 | 5.13E-06 | 4.07 | 1.66E-12 | DE in both female and male placentas |
| ENSRNOG00000054314 | | KCNG1 | 3.31 | 4.26E-06 | 4.42 | 8.15E-13 | DE in both female and male placentas |
| ENSRNOG00000020953 | | MS4A7 | 3.16 | 6.03E-35 | 3.45 | 6.54E-56 | DE in both female and male placentas |
| ENSRNOG00000007778 | | ALOX15B | 3.02 | 7.76E-04 | 2.19 | 4.29E-03 | DE in both female and male placentas |
| ENSRNOG00000018285 | | KCNA2 | 3.02 | 6.50E-14 | 2.49 | 8.49E-14 | DE in both female and male placentas |
| ENSRNOG00000004470 | | IL24 | 2.94 | 3.79E-05 | 5.10 | 1.71E-17 | DE in both female and male placentas |
| ENSRNOG00000030154 | | CYP4A2 | 2.93 | 4.19E-05 | 1.62 | 3.15E-02 | DE in both female and male placentas |

| | | | | | | | |
|---|---|---|---|---|---|---|---|
| ENSRNOG00000038211 | | AABR070328 56.1 | 2.92 | 2.29E-03 | 2.84 | 2.37E-04 | DE in both female and male placentas |
| ENSRNOG00000004229 | | TAC3 | 2.81 | 1.51E-02 | 4.04 | 1.31E-07 | DE in both female and male placentas |
| ENSRNOG00000019432 | | LYPD5 | 2.76 | 2.13E-03 | 3.48 | 1.17E-06 | DE in both female and male placentas |
| ENSRNOG00000016253 | | SLC6A18 | 2.73 | 1.55E-03 | 1.49 | 3.85E-02 | DE in both female and male placentas |
| ENSRNOG00000030719 | | CSMD1 | 2.70 | 3.01E-13 | 2.55 | 7.80E-17 | DE in both female and male placentas |
| ENSRNOG00000019317 | | CPLX3 | 2.65 | 7.60E-04 | 1.38 | 4.73E-02 | DE in both female and male placentas |
| ENSRNOG00000026196 | | LRRD1 | 2.53 | 2.19E-02 | 1.77 | 3.29E-02 | DE in both female and male placentas |
| ENSRNOG00000012477 | | EEF1A2 | 2.53 | 2.19E-09 | 2.85 | 4.02E-16 | DE in both female and male placentas |
| ENSRNOG00000015049 | | SCN5A | 2.45 | 7.13E-04 | 1.84 | 1.19E-03 | DE in both female and male placentas |
| ENSRNOG00000054203 | | SIM2 | 2.35 | 9.28E-06 | 1.86 | 2.22E-05 | DE in both female and male placentas |
| ENSRNOG00000005215 | X | KLHL4 | 2.33 | 2.05E-03 | 1.62 | 1.05E-02 | DE in both female and male placentas |
| ENSRNOG00000000201 | | GSTA5 | 2.29 | 2.19E-05 | 2.42 | 4.76E-08 | DE in both female and male placentas |
| ENSRNOG00000021260 | | PRND | 2.27 | 1.20E-03 | 3.36 | 2.52E-10 | DE in both female and male placentas |
| ENSRNOG00000024382 | | FCGR3A | 2.20 | 1.43E-09 | 2.86 | 8.63E-22 | DE in both female and male placentas |
| ENSRNOG00000016446 | | GPR45 | 2.19 | 3.00E-04 | 2.56 | 5.51E-08 | DE in both female and male placentas |
| ENSRNOG00000019479 | | KLF17 | 2.17 | 3.62E-03 | 2.67 | 8.72E-06 | DE in both female and male placentas |
| ENSRNOG00000028992 | | ACAN | 2.10 | 8.31E-12 | 2.04 | 4.14E-16 | DE in both female and male placentas |
| ENSRNOG00000003715 | X | SRPX2 | 2.10 | 1.52E-05 | 3.13 | 2.20E-15 | DE in both female and male placentas |

| | | | | | | | |
|---|---|---|---|---|---|---|---|
| ENSRNOG00000024693 | | ZFP831 | 2.09 | 1.35E-07 | 1.38 | 4.28E-05 | DE in both female and male placentas |
| ENSRNOG00000033618 | | COL28A1 | 2.03 | 1.81E-03 | 2.65 | 4.96E-07 | DE in both female and male placentas |
| ENSRNOG00000004062 | | C4BPA | 1.98 | 3.56E-05 | 2.32 | 1.54E-09 | DE in both female and male placentas |
| ENSRNOG00000059605 | | DDN | 1.98 | 6.14E-03 | 2.11 | 2.42E-04 | DE in both female and male placentas |
| ENSRNOG00000029768 | | CCL12 | 1.97 | 2.65E-05 | 3.60 | 3.04E-21 | DE in both female and male placentas |
| ENSRNOG00000050177 | X | LOC100911002 | 1.97 | 2.44E-02 | 2.38 | 3.76E-04 | DE in both female and male placentas |
| ENSRNOG00000022256 | | CXCL10 | 1.95 | 1.94E-05 | 3.81 | 4.38E-24 | DE in both female and male placentas |
| ENSRNOG00000007159 | | CCL2 | 1.94 | 1.08E-08 | 3.15 | 1.42E-29 | DE in both female and male placentas |
| ENSRNOG00000005286 | | COCH | 1.94 | 1.22E-07 | 2.22 | 9.95E-14 | DE in both female and male placentas |
| ENSRNOG00000056359 | | AABR07035722.1 | 1.93 | 3.83E-05 | 1.65 | 1.21E-05 | DE in both female and male placentas |
| ENSRNOG00000038047 | | MT1 | 1.91 | 1.24E-10 | 2.45 | 1.10E-23 | DE in both female and male placentas |
| ENSRNOG00000031443 | | HAVCR2 | 1.90 | 1.91E-05 | 2.22 | 1.83E-10 | DE in both female and male placentas |
| ENSRNOG00000033195 | | A1CF | 1.90 | 8.92E-06 | 1.35 | 1.38E-04 | DE in both female and male placentas |
| ENSRNOG00000016262 | | DNASE2B | 1.88 | 6.84E-07 | 3.38 | 6.99E-28 | DE in both female and male placentas |
| ENSRNOG00000023943 | | GBP1 | 1.86 | 7.61E-03 | 3.26 | 7.39E-11 | DE in both female and male placentas |
| ENSRNOG00000017485 | | CD72 | 1.86 | 2.32E-03 | 1.61 | 5.40E-04 | DE in both female and male placentas |
| ENSRNOG00000024832 | | GPR158 | 1.86 | 1.33E-02 | 2.77 | 7.81E-09 | DE in both female and male placentas |
| ENSRNOG00000029637 | | ACOT6 | 1.81 | 1.50E-03 | 1.57 | 9.88E-04 | DE in both female and male placentas |

| | | | | | | |
|---|---|---|---|---|---|---|
| ENSRNOG00000011517 | TNFRSF21 | 1.81 | 1.24E-22 | 1.92 | 3.94E-36 | DE in both female and male placentas |
| ENSRNOG00000001959 | MX1 | 1.79 | 2.02E-07 | 2.66 | 2.18E-21 | DE in both female and male placentas |
| ENSRNOG00000059698 | GPR141 | 1.79 | 2.21E-02 | 2.68 | 1.90E-07 | DE in both female and male placentas |
| ENSRNOG00000003866 | CXCR4 | 1.77 | 2.22E-12 | 2.05 | 2.49E-23 | DE in both female and male placentas |
| ENSRNOG00000025297 | GPR171 | 1.75 | 1.36E-07 | 1.13 | 7.36E-05 | DE in both female and male placentas |
| ENSRNOG00000059625 | RGD1560281 | 1.75 | 1.09E-02 | 1.93 | 2.73E-04 | DE in both female and male placentas |
| ENSRNOG00000023089 | AABR07067520.1 | 1.74 | 5.06E-03 | 1.23 | 3.37E-02 | DE in both female and male placentas |
| ENSRNOG00000027240 | KNDC1 | 1.74 | 6.49E-10 | 2.08 | 1.69E-19 | DE in both female and male placentas |
| ENSRNOG00000002782 | EPGN | 1.72 | 2.94E-02 | 2.73 | 1.70E-06 | DE in both female and male placentas |
| ENSRNOG00000043323 | IL36B | 1.72 | 2.61E-04 | 2.43 | 9.34E-11 | DE in both female and male placentas |
| ENSRNOG00000003365 | CADM3 | 1.70 | 4.28E-02 | 2.94 | 2.62E-06 | DE in both female and male placentas |
| ENSRNOG00000020864 | KIRREL2 | 1.69 | 7.30E-03 | 1.37 | 1.09E-02 | DE in both female and male placentas |
| ENSRNOG00000039954 | RGD1308751 | 1.67 | 3.47E-08 | 1.27 | 3.92E-07 | DE in both female and male placentas |
| ENSRNOG00000030983 | B3GALT5 | 1.66 | 8.72E-03 | 1.17 | 3.06E-02 | DE in both female and male placentas |
| ENSRNOG00000003709 | KMO | 1.63 | 5.94E-03 | 1.50 | 1.01E-03 | DE in both female and male placentas |
| ENSRNOG00000000768 | UBD | 1.63 | 4.25E-02 | 2.42 | 3.73E-06 | DE in both female and male placentas |
| ENSRNOG00000043098 | MT2A | 1.61 | 2.94E-15 | 2.00 | 7.26E-33 | DE in both female and male placentas |
| ENSRNOG00000058780 | IGFBP1 | 1.61 | 3.49E-02 | 1.41 | 2.17E-02 | DE in both female and male placentas |

| | | | | | | |
|---|---|---|---|---|---|---|
| ENSRNOG00000006732 | SPO11 | 1.60 | 9.99E-03 | 1.60 | 2.63E-03 | DE in both female and male placentas |
| ENSRNOG00000020560 | AABR070058 21.1 | 1.55 | 2.55E-03 | 1.25 | 2.71E-03 | DE in both female and male placentas |
| ENSRNOG00000010932 | MATN1 | 1.54 | 1.94E-03 | 1.14 | 3.73E-03 | DE in both female and male placentas |
| ENSRNOG00000018092 | CD83 | 1.53 | 1.86E-14 | 1.52 | 1.50E-20 | DE in both female and male placentas |
| ENSRNOG00000000239 | CCL7 | 1.52 | 3.99E-04 | 2.80 | 3.63E-17 | DE in both female and male placentas |
| ENSRNOG00000012168 | FBXL21 | 1.49 | 1.19E-06 | 1.40 | 2.21E-08 | DE in both female and male placentas |
| ENSRNOG00000006963 | CTSE | 1.49 | 1.08E-07 | 1.42 | 3.92E-10 | DE in both female and male placentas |
| ENSRNOG00000034190 | IGHM | 1.46 | 4.53E-05 | 1.96 | 3.54E-12 | DE in both female and male placentas |
| ENSRNOG00000005871 | IL1RN | 1.45 | 2.73E-07 | 2.41 | 1.96E-26 | DE in both female and male placentas |
| ENSRNOG00000029427 | GRHL3 | 1.44 | 3.22E-07 | 1.43 | 2.10E-10 | DE in both female and male placentas |
| ENSRNOG00000010797 | ESM1 | 1.44 | 3.40E-05 | 1.35 | 1.39E-06 | DE in both female and male placentas |
| ENSRNOG00000032009 | LOC499643 | 1.44 | 7.09E-03 | 1.02 | 3.69E-02 | DE in both female and male placentas |
| ENSRNOG00000033017 | LOC1009106 69 | 1.44 | 8.71E-04 | 2.36 | 7.43E-13 | DE in both female and male placentas |
| ENSRNOG00000026613 | AADACL3 | 1.42 | 9.90E-03 | 1.21 | 2.63E-03 | DE in both female and male placentas |
| ENSRNOG00000061739 | KLRK1 | 1.42 | 1.44E-02 | 1.80 | 8.44E-06 | DE in both female and male placentas |
| ENSRNOG00000058446 | IL7R | 1.41 | 1.38E-04 | 2.26 | 9.10E-16 | DE in both female and male placentas |
| ENSRNOG00000019587 | PTPRN | 1.40 | 9.09E-04 | 3.00 | 3.75E-20 | DE in both female and male placentas |
| ENSRNOG00000003105 | KIF19 | 1.40 | 3.76E-05 | 1.51 | 2.59E-08 | DE in both female and male placentas |

| | | | | | | |
|---|---|---|---|---|---|---|
| ENSRNOG00000054964 | AOAH | 1.40 | 6.48E-04 | 1.54 | 8.28E-07 | DE in both female and male placentas |
| ENSRNOG00000001011 | RFLNA | 1.37 | 5.48E-07 | 1.25 | 6.98E-09 | DE in both female and male placentas |
| ENSRNOG00000001691 | CLDN14 | 1.37 | 9.55E-03 | 1.67 | 7.05E-05 | DE in both female and male placentas |
| ENSRNOG00000019862 | PINLYP | 1.37 | 9.97E-05 | 2.04 | 1.37E-13 | DE in both female and male placentas |
| ENSRNOG00000046134 | OLR1551 | 1.37 | 2.19E-05 | 1.42 | 1.11E-07 | DE in both female and male placentas |
| ENSRNOG00000010824 | PARP8 | 1.36 | 2.75E-07 | 1.57 | 1.04E-13 | DE in both female and male placentas |
| ENSRNOG00000034228 | PIK3C2G | 1.34 | 6.49E-09 | 1.35 | 6.63E-14 | DE in both female and male placentas |
| ENSRNOG00000028841 | MT1M | 1.34 | 2.51E-02 | 2.20 | 4.64E-09 | DE in both female and male placentas |
| ENSRNOG00000001926 | CLDN1 | 1.34 | 6.18E-05 | 1.41 | 9.00E-08 | DE in both female and male placentas |
| ENSRNOG00000027433 | LOC100910708 | 1.34 | 4.35E-02 | 1.03 | 4.51E-02 | DE in both female and male placentas |
| ENSRNOG00000038319 | AKR1C19 | 1.33 | 3.47E-06 | 1.18 | 3.51E-07 | DE in both female and male placentas |
| ENSRNOG00000008478 | MMP13 | 1.33 | 4.54E-04 | 2.23 | 3.86E-15 | DE in both female and male placentas |
| ENSRNOG00000002978 | FASLG | 1.32 | 1.23E-07 | 1.50 | 1.17E-13 | DE in both female and male placentas |
| ENSRNOG00000022950 | DPPA3 | 1.31 | 3.11E-05 | 1.87 | 1.69E-14 | DE in both female and male placentas |
| ENSRNOG00000053848 | LOC102552344 | 1.31 | 4.04E-02 | 1.98 | 1.23E-05 | DE in both female and male placentas |
| ENSRNOG00000000536 | MDGA1 | 1.31 | 9.76E-04 | 1.50 | 2.48E-06 | DE in both female and male placentas |
| ENSRNOG00000026306 | CLEC5A | 1.30 | 4.72E-04 | 1.87 | 3.07E-11 | DE in both female and male placentas |
| ENSRNOG00000014578 | FXYD4 | 1.30 | 1.35E-03 | 1.77 | 7.40E-09 | DE in both female and male placentas |

| | | | | | | |
|---|---|---|---|---|---|---|
| ENSRNOG00000006767 | CST7 | 1.29 | 1.92E-02 | 2.81 | 2.14E-13 | DE in both female and male placentas |
| ENSRNOG00000003800 | RGS9 | 1.28 | 8.37E-03 | 1.34 | 5.19E-04 | DE in both female and male placentas |
| ENSRNOG00000000640 | EGR2 | 1.28 | 3.08E-03 | 1.72 | 6.37E-08 | DE in both female and male placentas |
| ENSRNOG00000000500 | SCUBE3 | 1.28 | 1.69E-02 | 1.92 | 7.88E-07 | DE in both female and male placentas |
| ENSRNOG00000028043 | CXCL3 | 1.28 | 2.10E-02 | 1.43 | 6.67E-04 | DE in both female and male placentas |
| ENSRNOG00000005007 | SCN3A | 1.26 | 4.28E-02 | 1.23 | 1.45E-02 | DE in both female and male placentas |
| ENSRNOG00000011205 | CCL3 | 1.26 | 2.32E-02 | 1.92 | 1.54E-06 | DE in both female and male placentas |
| ENSRNOG00000020907 | ZP1 | 1.26 | 2.70E-04 | 1.43 | 1.44E-07 | DE in both female and male placentas |
| ENSRNOG00000048036 | NCAN | 1.24 | 1.72E-04 | 1.41 | 5.95E-08 | DE in both female and male placentas |
| ENSRNOG00000048967 | LOC688459 | 1.24 | 1.18E-02 | 2.28 | 4.33E-11 | DE in both female and male placentas |
| ENSRNOG00000046482 | ZNF750 | 1.24 | 2.06E-04 | 2.08 | 1.88E-16 | DE in both female and male placentas |
| ENSRNOG00000000383 | MYPN | 1.23 | 2.44E-08 | 1.72 | 2.55E-23 | DE in both female and male placentas |
| ENSRNOG00000006314 | ZBP1 | 1.23 | 1.44E-02 | 1.24 | 4.46E-04 | DE in both female and male placentas |
| ENSRNOG00000017803 | APBB1IP | 1.23 | 4.41E-06 | 1.11 | 2.94E-07 | DE in both female and male placentas |
| ENSRNOG00000042975 | TMEM45A | 1.22 | 6.59E-04 | 1.20 | 2.05E-05 | DE in both female and male placentas |
| ENSRNOG00000028622 | PNPLA1 | 1.22 | 1.72E-04 | 1.97 | 2.75E-15 | DE in both female and male placentas |
| ENSRNOG00000008716 | NEFH | 1.22 | 5.41E-03 | 1.42 | 2.45E-05 | DE in both female and male placentas |
| ENSRNOG00000058645 | TNC | 1.22 | 9.85E-05 | 1.82 | 7.49E-14 | DE in both female and male placentas |

| | | | | | | |
|---|---|---|---|---|---|---|
| ENSRNOG00000009350 | SEZ6 | 1.21 | 2.50E-06 | 1.36 | 2.00E-11 | DE in both female and male placentas |
| ENSRNOG00000017089 | PRL6A1 | 1.20 | 3.37E-09 | 1.12 | 1.10E-11 | DE in both female and male placentas |
| ENSRNOG00000031599 | SH2D1B | 1.20 | 2.95E-02 | 1.10 | 8.20E-03 | DE in both female and male placentas |
| ENSRNOG00000011054 | LAPTM5 | 1.20 | 1.30E-11 | 1.23 | 1.16E-17 | DE in both female and male placentas |
| ENSRNOG00000005299 | KIF5A | 1.20 | 1.16E-04 | 1.21 | 9.67E-07 | DE in both female and male placentas |
| ENSRNOG00000037409 | SCIMP | 1.19 | 4.39E-03 | 1.85 | 2.04E-09 | DE in both female and male placentas |
| ENSRNOG00000037632 | LOC688553 | 1.19 | 1.05E-09 | 1.48 | 2.38E-21 | DE in both female and male placentas |
| ENSRNOG00000019440 | KCNN4 | 1.18 | 3.95E-02 | 1.96 | 1.14E-06 | DE in both female and male placentas |
| ENSRNOG00000002792 | AHRR | 1.17 | 4.67E-02 | 1.59 | 1.69E-04 | DE in both female and male placentas |
| ENSRNOG00000028750 | TUBAL3 | 1.17 | 9.63E-03 | 1.00 | 4.36E-03 | DE in both female and male placentas |
| ENSRNOG00000021468 | GRM8 | 1.17 | 1.86E-02 | 1.37 | 3.01E-04 | DE in both female and male placentas |
| ENSRNOG00000009734 | AKR1B8 | 1.16 | 7.58E-06 | 1.52 | 9.35E-14 | DE in both female and male placentas |
| ENSRNOG00000012450 | DYNLRB2 | 1.16 | 1.02E-07 | 1.29 | 8.87E-14 | DE in both female and male placentas |
| ENSRNOG00000023337 | SEMA3A | 1.16 | 1.46E-02 | 1.28 | 3.60E-04 | DE in both female and male placentas |
| ENSRNOG00000004019 | PHLDA1 | 1.15 | 4.96E-06 | 1.31 | 4.55E-11 | DE in both female and male placentas |
| ENSRNOG00000029756 | P2RY13 | 1.14 | 2.12E-04 | 1.06 | 2.45E-05 | DE in both female and male placentas |
| ENSRNOG00000011424 | CLDN23 | 1.14 | 2.97E-04 | 1.75 | 4.83E-13 | DE in both female and male placentas |
| ENSRNOG00000001963 | MX2 | 1.14 | 1.07E-03 | 1.75 | 4.58E-11 | DE in both female and male placentas |

| | | | | | | | |
|---|---|---|---|---|---|---|---|
| ENSRNOG00000021084 | | AABR070063 10.1 | 1.14 | 3.10E-07 | 1.51 | 8.50E-18 | DE in both female and male placentas |
| ENSRNOG00000017519 | | LOC1025572 44 | 1.14 | 1.21E-10 | 1.13 | 3.34E-15 | DE in both female and male placentas |
| ENSRNOG00000021009 | | GAPDHS | 1.14 | 1.04E-03 | 1.25 | 5.71E-06 | DE in both female and male placentas |
| ENSRNOG00000014465 | | OCA2 | 1.14 | 3.59E-03 | 1.26 | 9.58E-06 | DE in both female and male placentas |
| ENSRNOG00000023257 | | ADAMTS9 | 1.13 | 4.02E-10 | 1.11 | 2.13E-14 | DE in both female and male placentas |
| ENSRNOG00000057072 | | SLC12A3 | 1.13 | 1.43E-02 | 1.68 | 5.45E-07 | DE in both female and male placentas |
| ENSRNOG00000046566 | | TUB | 1.11 | 1.79E-02 | 1.25 | 3.01E-04 | DE in both female and male placentas |
| ENSRNOG00000028768 | | GBP4 | 1.10 | 2.87E-03 | 2.52 | 5.09E-20 | DE in both female and male placentas |
| ENSRNOG00000024390 | | OSM | 1.10 | 6.41E-05 | 1.50 | 4.37E-12 | DE in both female and male placentas |
| ENSRNOG00000016483 | | MYO16 | 1.10 | 3.83E-04 | 1.95 | 6.65E-17 | DE in both female and male placentas |
| ENSRNOG00000008736 | | SLAMF8 | 1.10 | 1.19E-02 | 1.95 | 2.17E-10 | DE in both female and male placentas |
| ENSRNOG00000059140 | | MYO1G | 1.09 | 5.53E-05 | 1.52 | 5.60E-13 | DE in both female and male placentas |
| ENSRNOG00000029106 | X | USP26 | 1.09 | 2.32E-06 | 1.04 | 7.38E-08 | DE in both female and male placentas |
| ENSRNOG00000012772 | | NQO1 | 1.09 | 2.68E-15 | 1.51 | 1.10E-41 | DE in both female and male placentas |
| ENSRNOG00000054142 | | SLC30A2 | 1.09 | 5.85E-12 | 1.14 | 3.31E-19 | DE in both female and male placentas |
| ENSRNOG00000006940 | | NCF4 | 1.09 | 2.16E-04 | 1.67 | 4.48E-14 | DE in both female and male placentas |
| ENSRNOG00000012393 | | S100A13 | 1.08 | 6.70E-06 | 1.34 | 1.12E-12 | DE in both female and male placentas |
| ENSRNOG00000019422 | | EGR1 | 1.07 | 1.20E-04 | 1.70 | 2.11E-15 | DE in both female and male placentas |

| | | | | | | | |
|---|---|---|---|---|---|---|---|
| ENSRNOG00000009211 | | C3AR1 | 1.07 | 2.92E-03 | 1.49 | 5.11E-08 | DE in both female and male placentas |
| ENSRNOG00000004964 | | ERBB3 | 1.06 | 3.25E-15 | 1.33 | 4.69E-34 | DE in both female and male placentas |
| ENSRNOG00000003741 | | NPTX1 | 1.06 | 4.74E-05 | 1.17 | 1.76E-08 | DE in both female and male placentas |
| ENSRNOG00000005700 | | NSG1 | 1.06 | 5.92E-05 | 1.01 | 1.35E-06 | DE in both female and male placentas |
| ENSRNOG00000003435 | X | MCF2 | 1.06 | 3.60E-11 | 1.23 | 5.58E-22 | DE in both female and male placentas |
| ENSRNOG00000021424 | | CD300LF | 1.06 | 5.91E-03 | 1.51 | 7.93E-08 | DE in both female and male placentas |
| ENSRNOG00000012216 | | TGFBI | 1.06 | 4.21E-04 | 1.38 | 2.25E-09 | DE in both female and male placentas |
| ENSRNOG00000029514 | | SLC26A9 | 1.05 | 1.27E-03 | 1.21 | 3.47E-06 | DE in both female and male placentas |
| ENSRNOG00000039891 | | ADGRG5 | 1.04 | 2.00E-02 | 1.89 | 1.03E-09 | DE in both female and male placentas |
| ENSRNOG00000047606 | | BCL2A1 | 1.04 | 3.71E-06 | 1.34 | 1.34E-14 | DE in both female and male placentas |
| ENSRNOG00000011947 | | TIFAB | 1.04 | 2.20E-03 | 1.74 | 6.97E-12 | DE in both female and male placentas |
| ENSRNOG00000000552 | | COL13A1 | 1.04 | 3.79E-04 | 1.27 | 1.76E-08 | DE in both female and male placentas |
| ENSRNOG00000052070 | | ALDH1A3 | 1.03 | 1.32E-02 | 1.17 | 2.26E-04 | DE in both female and male placentas |
| ENSRNOG00000026091 | | SLC10A4 | 1.03 | 3.61E-05 | 1.39 | 1.42E-12 | DE in both female and male placentas |
| ENSRNOG00000048186 | | F2RL3 | 1.03 | 9.89E-06 | 1.36 | 3.67E-13 | DE in both female and male placentas |
| ENSRNOG00000060518 | | AABR07015057.1 | 1.03 | 1.02E-06 | 1.30 | 6.66E-15 | DE in both female and male placentas |
| ENSRNOG00000047630 | X | LOC100911047 | 1.03 | 2.95E-02 | 1.72 | 1.08E-06 | DE in both female and male placentas |
| ENSRNOG00000047110 | X | XLR3A | 1.03 | 2.95E-02 | 1.72 | 1.08E-06 | DE in both female and male placentas |

| | | | | | | | |
|---|---|---|---|---|---|---|---|
| ENSRNOG00000003585 | X | SYTL5 | 1.03 | 3.11E-04 | 1.03 | 4.32E-06 | DE in both female and male placentas |
| ENSRNOG00000049743 | | GSTM5 | 1.03 | 3.81E-05 | 1.31 | 1.62E-11 | DE in both female and male placentas |
| ENSRNOG00000006583 | | HPGDS | 1.02 | 7.22E-04 | 1.30 | 2.58E-08 | DE in both female and male placentas |
| ENSRNOG00000013791 | | ENPP3 | 1.01 | 1.21E-06 | 1.46 | 7.33E-19 | DE in both female and male placentas |
| ENSRNOG00000027288 | | PSGB1 | 1.01 | 2.24E-06 | 1.06 | 4.19E-10 | DE in both female and male placentas |
| ENSRNOG00000012817 | | LRRC17 | 1.01 | 1.13E-03 | 1.09 | 4.03E-06 | DE in both female and male placentas |
| ENSRNOG00000008346 | | ITGB6 | 1.01 | 1.21E-10 | 1.00 | 3.17E-15 | DE in both female and male placentas |
| ENSRNOG00000027935 | | LRRC34 | 1.00 | 3.13E-03 | 1.04 | 1.32E-04 | DE in both female and male placentas |
| ENSRNOG00000009004 | | LRRIQ4 | 1.00 | 6.48E-03 | 1.48 | 2.51E-07 | DE in both female and male placentas |
| ENSRNOG00000013674 | | MEGF10 | 1.00 | 2.78E-04 | 1.33 | 2.21E-10 | DE in both female and male placentas |
| ENSRNOG00000028382 | | RFXAPL1 | 8.12 | 2.19E-05 | -5.58 | 9.87E-04 | DE in both female and male placentas |
| ENSRNOG00000061082 | | U1 | 7.67 | 4.96E-05 | -6.70 | 8.78E-05 | DE in both female and male placentas |
| ENSRNOG00000049283 | | AABR07057442.1 | 5.07 | 6.11E-03 | -4.71 | 4.63E-03 | DE in both female and male placentas |
| ENSRNOG00000033358 | | LOC100910565 | 3.90 | 4.46E-02 | -3.90 | 2.29E-02 | DE in both female and male placentas |
| ENSRNOG00000046540 | | UGT2B15 | 2.60 | 2.48E-02 | -2.90 | 1.68E-03 | DE in both female and male placentas |
| ENSRNOG00000032990 | | LOC690483 | 2.42 | 1.58E-02 | -2.26 | 2.26E-02 | DE in both female and male placentas |
| ENSRNOG00000053807 | | ZFP280B | -5.52 | 3.33E-02 | 4.83 | 1.19E-03 | DE in both female and male placentas |
| ENSRNOG00000019147 | | STOM | -1.00 | 2.82E-05 | -1.28 | 1.36E-12 | DE in both female and male placentas |

| | | | | | | |
|---|---|---|---|---|---|---|
| ENSRNOG00000022448 | HTRA2 | -1.00 | 2.01E-11 | -1.02 | 1.93E-18 | DE in both female and male placentas |
| ENSRNOG00000015370 | SLC35G2 | -1.00 | 1.62E-04 | -1.19 | 5.09E-09 | Differentially expressed in both female and male |
| ENSRNOG00000062220 | CENPP | -1.00 | 4.03E-02 | -1.86 | 1.95E-08 | DE in both female and male placentas |
| ENSRNOG00000019503 | REC8 | -1.00 | 4.75E-02 | -2.53 | 5.17E-12 | DE in both female and male placentas |
| ENSRNOG00000058039 | ACTA2 | -1.00 | 1.98E-06 | -1.69 | 2.54E-25 | DE in both female and male placentas |
| ENSRNOG00000056550 | EPB41L4B | -1.00 | 7.50E-05 | -1.26 | 6.31E-11 | DE in both female and male placentas |
| ENSRNOG00000024365 | ECT2 | -1.01 | 9.84E-05 | -1.40 | 3.74E-13 | DE in both female and male placentas |
| ENSRNOG00000023148 | COL11A1 | -1.01 | 9.92E-03 | -1.59 | 3.70E-08 | DE in both female and male placentas |
| ENSRNOG00000023576 | ECRG4 | -1.01 | 4.28E-03 | -1.16 | 9.01E-06 | DE in both female and male placentas |
| ENSRNOG00000039582 | RGD1561161 | -1.01 | 2.07E-05 | -1.72 | 1.57E-21 | DE in both female and male placentas |
| ENSRNOG00000001215 | CFAP410 | -1.01 | 2.18E-08 | -1.06 | 1.81E-14 | DE in both female and male placentas |
| ENSRNOG00000020918 | CCND1 | -1.01 | 6.20E-09 | -1.16 | 1.00E-17 | DE in both female and male placentas |
| ENSRNOG00000045553 | PROSER2 | -1.01 | 3.91E-11 | -1.25 | 1.16E-25 | DE in both female and male placentas |
| ENSRNOG00000017164 | AFAP1L2 | -1.01 | 1.21E-06 | -1.15 | 6.90E-13 | DE in both female and male placentas |
| ENSRNOG00000000818 | NRM | -1.02 | 3.50E-06 | -1.55 | 2.05E-20 | DE in both female and male placentas |
| ENSRNOG00000014504 | IL1R1 | -1.02 | 6.52E-05 | -1.51 | 4.82E-15 | DE in both female and male placentas |
| ENSRNOG00000012036 | PCSK5 | -1.02 | 1.76E-06 | -1.57 | 9.16E-21 | DE in both female and male placentas |
| ENSRNOG00000016538 | ITGA8 | -1.02 | 9.92E-10 | -1.15 | 1.06E-18 | DE in both female and male placentas |

| | | | | | | | |
|---|---|---|---|---|---|---|---|
| ENSRNOG00000000186 | | TST | -1.02 | 3.38E-02 | -1.05 | 2.66E-03 | DE in both female and male placentas |
| ENSRNOG00000017260 | | CDR2 | -1.02 | 6.03E-18 | -1.17 | 6.96E-36 | DE in both female and male placentas |
| ENSRNOG00000016099 | | ID4 | -1.02 | 3.83E-04 | -1.27 | 5.11E-09 | DE in both female and male placentas |
| ENSRNOG00000025383 | X | PDHA1 | -1.03 | 5.65E-10 | -1.32 | 1.61E-24 | DE in both female and male placentas |
| ENSRNOG00000059237 | | CLUL1 | -1.03 | 9.99E-03 | -1.26 | 1.71E-05 | DE in both female and male placentas |
| ENSRNOG00000011559 | | CNN3 | -1.03 | 7.47E-10 | -1.27 | 2.29E-22 | DE in both female and male placentas |
| ENSRNOG00000024178 | | CENPT | -1.03 | 4.09E-03 | -1.81 | 1.20E-12 | DE in both female and male placentas |
| ENSRNOG00000059750 | | AC141102.1 | -1.03 | 2.81E-02 | -1.65 | 4.93E-07 | DE in both female and male placentas |
| ENSRNOG00000022391 | X | ZFP711 | -1.03 | 1.55E-04 | -1.41 | 5.94E-11 | DE in both female and male placentas |
| ENSRNOG00000043422 | | LOC100359777 | -1.03 | 1.32E-03 | -1.47 | 4.89E-10 | DE in both female and male placentas |
| ENSRNOG00000017976 | | SLCO2B1 | -1.03 | 5.69E-07 | -1.35 | 1.98E-17 | DE in both female and male placentas |
| ENSRNOG00000014585 | | CFAP53 | -1.03 | 2.40E-04 | -1.07 | 3.93E-07 | DE in both female and male placentas |
| ENSRNOG00000012557 | | LGALS5 | -1.03 | 3.32E-03 | -1.91 | 1.75E-13 | DE in both female and male placentas |
| ENSRNOG00000047369 | | LOC100910057 | -1.03 | 4.84E-02 | -2.33 | 2.14E-09 | DE in both female and male placentas |
| ENSRNOG00000000782 | | TRIM10 | -1.04 | 2.10E-02 | -1.94 | 1.22E-08 | DE in both female and male placentas |
| ENSRNOG00000024208 | | PRSS48 | -1.04 | 4.70E-04 | -1.04 | 5.81E-06 | DE in both female and male placentas |
| ENSRNOG00000000614 | | BICC1 | -1.04 | 2.37E-08 | -1.26 | 2.07E-18 | DE in both female and male placentas |
| ENSRNOG00000008401 | | CARD10 | -1.04 | 1.57E-05 | -1.20 | 8.95E-11 | DE in both female and male placentas |

| | | | | | | | |
|---|---|---|---|---|---|---|---|
| ENSRNOG00000050766 | | RD3 | -1.04 | 5.40E-03 | -1.15 | 3.71E-05 | DE in both female and male placentas |
| ENSRNOG00000059507 | | LAMC3 | -1.04 | 3.87E-04 | -1.40 | 1.02E-10 | DE in both female and male placentas |
| ENSRNOG00000018254 | | SNCAIP | -1.04 | 7.87E-04 | -1.46 | 2.71E-10 | DE in both female and male placentas |
| ENSRNOG00000008738 | | TP53I11 | -1.04 | 7.22E-08 | -1.03 | 5.56E-12 | DE in both female and male placentas |
| ENSRNOG00000033123 | | PCDHB20 | -1.05 | 9.41E-03 | -1.22 | 3.20E-05 | DE in both female and male placentas |
| ENSRNOG00000021800 | | ZFP541 | -1.05 | 3.51E-02 | -1.23 | 5.32E-04 | DE in both female and male placentas |
| ENSRNOG00000014367 | | EPHB6 | -1.05 | 1.93E-03 | -1.45 | 5.73E-09 | DE in both female and male placentas |
| ENSRNOG00000028090 | | ARHGEF18 | -1.05 | 5.58E-05 | -1.41 | 9.74E-13 | DE in both female and male placentas |
| ENSRNOG00000016260 | | POLR3G | -1.05 | 9.07E-07 | -1.41 | 1.28E-17 | DE in both female and male placentas |
| ENSRNOG00000049849 | X | FAM199X | -1.05 | 1.59E-04 | -1.08 | 3.46E-07 | DE in both female and male placentas |
| ENSRNOG00000000033 | | TMCC2 | -1.06 | 1.02E-08 | -1.07 | 1.43E-13 | DE in both female and male placentas |
| ENSRNOG00000015473 | | PHACTR2 | -1.06 | 3.47E-06 | -1.43 | 2.08E-16 | DE in both female and male placentas |
| ENSRNOG00000047143 | | FKBP1B | -1.06 | 6.94E-05 | -1.34 | 5.33E-11 | DE in both female and male placentas |
| ENSRNOG00000036571 | | CES2C | -1.06 | 3.77E-06 | -1.11 | 2.70E-10 | DE in both female and male placentas |
| ENSRNOG00000008235 | | MYLK2 | -1.06 | 3.09E-02 | -1.23 | 2.56E-03 | DE in both female and male placentas |
| ENSRNOG00000015421 | | SLC27A3 | -1.06 | 1.31E-08 | -1.04 | 8.13E-13 | DE in both female and male placentas |
| ENSRNOG00000048751 | | AABR07065531.3 | -1.06 | 6.56E-05 | -1.70 | 3.32E-17 | DE in both female and male placentas |
| ENSRNOG00000002751 | X | ZDHHC15 | -1.07 | 8.95E-07 | -1.54 | 1.56E-19 | DE in both female and male placentas |

| | | | | | | | |
|---|---|---|---|---|---|---|---|
| ENSRNOG00000011750 | | FAM180A | -1.07 | 2.40E-05 | -1.23 | 1.89E-10 | DE in both female and male placentas |
| ENSRNOG00000028845 | | EBF1 | -1.07 | 1.49E-06 | -1.35 | 3.35E-15 | DE in both female and male placentas |
| ENSRNOG00000008012 | | ABCB4 | -1.07 | 3.49E-05 | -1.42 | 4.21E-13 | DE in both female and male placentas |
| ENSRNOG00000004281 | | COBL | -1.07 | 3.61E-04 | -1.85 | 2.30E-16 | DE in both female and male placentas |
| ENSRNOG00000007426 | | TMEM64 | -1.07 | 4.36E-05 | -1.21 | 2.29E-09 | DE in both female and male placentas |
| ENSRNOG00000015654 | | GHR | -1.07 | 6.67E-07 | -1.62 | 3.04E-22 | DE in both female and male placentas |
| ENSRNOG00000002097 | | RASL11B | -1.07 | 3.27E-16 | -1.08 | 3.13E-25 | DE in both female and male placentas |
| ENSRNOG00000039593 | | ECSCR | -1.08 | 2.69E-03 | -1.59 | 1.02E-09 | DE in both female and male placentas |
| ENSRNOG00000011716 | | DEGS2 | -1.08 | 1.61E-02 | -1.12 | 5.78E-04 | DE in both female and male placentas |
| ENSRNOG00000054764 | | FLT3 | -1.08 | 1.47E-03 | -1.72 | 6.79E-12 | DE in both female and male placentas |
| ENSRNOG00000015756 | | AABR070536 87.1 | -1.08 | 4.06E-09 | -1.29 | 1.10E-19 | DE in both female and male placentas |
| ENSRNOG00000047663 | X | GATA1 | -1.08 | 8.57E-04 | -1.35 | 1.99E-08 | DE in both female and male placentas |
| ENSRNOG00000001621 | | C2CD2 | -1.08 | 6.58E-13 | -1.28 | 1.94E-27 | DE in both female and male placentas |
| ENSRNOG00000059166 | | LDB3 | -1.09 | 2.96E-06 | -1.42 | 1.64E-15 | DE in both female and male placentas |
| ENSRNOG00000016550 | | DCLK2 | -1.09 | 7.65E-05 | -1.16 | 6.99E-08 | DE in both female and male placentas |
| ENSRNOG00000020562 | | SLC22A18 | -1.09 | 7.97E-05 | -1.28 | 8.03E-10 | DE in both female and male placentas |
| ENSRNOG00000003229 | X | TSPAN7 | -1.09 | 4.35E-06 | -1.54 | 1.97E-17 | DE in both female and male placentas |
| ENSRNOG00000039063 | X | LOC681355 | -1.09 | 8.09E-06 | -1.31 | 4.71E-12 | DE in both female and male placentas |

| | | | | | | |
|---|---|---|---|---|---|---|
| ENSRNOG00000001793 | HEG1 | -1.09 | 4.33E-10 | -1.40 | 1.13E-24 | DE in both female and male placentas |
| ENSRNOG00000019365 | ABLIM3 | -1.09 | 1.95E-03 | -1.60 | 9.17E-10 | DE in both female and male placentas |
| ENSRNOG00000014549 | ARHGEF26 | -1.09 | 1.40E-02 | -2.19 | 4.57E-12 | DE in both female and male placentas |
| ENSRNOG00000001214 | PFKL | -1.09 | 5.00E-10 | -1.07 | 5.55E-15 | DE in both female and male placentas |
| ENSRNOG00000038980 | LYPD6 | -1.09 | 5.12E-03 | -1.49 | 1.68E-07 | DE in both female and male placentas |
| ENSRNOG00000001769 | TNK2 | -1.09 | 1.04E-08 | -1.08 | 2.79E-13 | DE in both female and male placentas |
| ENSRNOG00000054049 | PRELID2 | -1.10 | 3.74E-08 | -1.03 | 1.44E-11 | DE in both female and male placentas |
| ENSRNOG00000055328 | AABR070172 68.1 | -1.10 | 4.76E-02 | -1.45 | 7.05E-04 | DE in both female and male placentas |
| ENSRNOG00000000940 | SFLT1 (HSOULD BE [illegible] | -1.10 | 1.09E-08 | -1.32 | 1.02E-18 | DE in both female and male placentas |
| ENSRNOG00000009980 | PLPP1 | -1.10 | 2.58E-09 | -1.14 | 2.25E-15 | DE in both female and male placentas |
| ENSRNOG00000008674 | PRDM11 | -1.10 | 9.02E-07 | -1.54 | 4.48E-19 | DE in both female and male placentas |
| ENSRNOG00000016648 | GULO | -1.10 | 5.53E-04 | -1.62 | 9.53E-12 | DE in both female and male placentas |
| ENSRNOG00000027891 | DHRS11 | -1.10 | 3.01E-07 | -1.12 | 2.27E-11 | DE in both female and male placentas |
| ENSRNOG00000050190 | ENG | -1.10 | 1.60E-09 | -1.05 | 1.32E-13 | DE in both female and male placentas |
| ENSRNOG00000008539 | LGALS2 | -1.10 | 2.12E-02 | -1.08 | 2.40E-03 | DE in both female and male placentas |
| ENSRNOG00000024008 | CDC25C | -1.10 | 5.84E-13 | -1.20 | 8.57E-24 | DE in both female and male placentas |
| ENSRNOG00000004320 | TGM4 | -1.11 | 2.22E-14 | -1.06 | 2.16E-21 | DE in both female and male placentas |
| ENSRNOG00000002061 | PTPN13 | -1.11 | 5.82E-08 | -1.26 | 1.05E-15 | DE in both female and male placentas |

| | | | | | | | |
|---|---|---|---|---|---|---|---|
| ENSRNOG00000008989 | | CRYL1 | -1.11 | 5.36E-06 | -1.28 | 6.65E-12 | DE in both female and male placentas |
| ENSRNOG00000029141 | | TRABD2B | -1.11 | 5.35E-04 | -1.34 | 3.55E-08 | DE in both female and male placentas |
| ENSRNOG00000000274 | | PHYHIPL | -1.11 | 7.58E-06 | -1.18 | 4.23E-10 | DE in both female and male placentas |
| ENSRNOG00000056562 | | OLFML1 | -1.11 | 9.02E-07 | -1.46 | 9.32E-17 | DE in both female and male placentas |
| ENSRNOG00000000986 | | CAMSAP3 | -1.11 | 3.26E-07 | -1.38 | 1.60E-16 | DE in both female and male placentas |
| ENSRNOG00000025371 | | SPRY1 | -1.11 | 4.43E-11 | -1.14 | 5.79E-18 | DE in both female and male placentas |
| ENSRNOG00000013280 | | OLFM4 | -1.11 | 1.37E-03 | -1.79 | 7.79E-12 | Differentially expressed in both female and male |
| ENSRNOG00000047531 | | CCDC57 | -1.12 | 7.58E-05 | -1.74 | 4.41E-16 | DE in both female and male placentas |
| ENSRNOG00000025484 | | TMEM177 | -1.12 | 5.47E-08 | -1.65 | 5.31E-25 | DE in both female and male placentas |
| ENSRNOG00000016879 | | LDLRAD4 | -1.12 | 1.99E-08 | -1.47 | 3.53E-21 | DE in both female and male placentas |
| ENSRNOG00000002511 | | FLT4 | -1.12 | 6.09E-04 | -1.52 | 4.03E-10 | DE in both female and male placentas |
| ENSRNOG00000061147 | | FAM189A2 | -1.12 | 1.56E-07 | -1.14 | 2.87E-12 | DE in both female and male placentas |
| ENSRNOG00000054581 | | TMEM101 | -1.12 | 1.22E-06 | -1.48 | 7.34E-17 | DE in both female and male placentas |
| ENSRNOG00000010238 | | CHMP4C | -1.12 | 3.10E-08 | -1.23 | 4.50E-15 | DE in both female and male placentas |
| ENSRNOG00000002215 | | MYLK | -1.12 | 4.65E-11 | -1.53 | 2.77E-30 | DE in both female and male placentas |
| ENSRNOG00000028805 | | CFAP77 | -1.12 | 4.20E-02 | -1.37 | 3.72E-04 | DE in both female and male placentas |
| ENSRNOG00000004330 | X | CHRDL1 | -1.12 | 1.28E-05 | -1.50 | 1.98E-14 | DE in both female and male placentas |
| ENSRNOG00000001652 | | ERG | -1.13 | 2.91E-04 | -1.48 | 1.99E-10 | DE in both female and male placentas |

| | | | | | | | |
|---|---|---|---|---|---|---|---|
| ENSRNOG00000014398 | | SCARA5 | -1.13 | 1.35E-06 | -1.13 | 3.28E-10 | DE in both female and male placentas |
| ENSRNOG00000028743 | | GAL3ST3 | -1.13 | 2.67E-02 | -2.58 | 5.36E-12 | DE in both female and male placentas |
| ENSRNOG00000001976 | | TMPRSS2 | -1.13 | 3.41E-04 | -1.96 | 1.39E-16 | DE in both female and male placentas |
| ENSRNOG00000007025 | | EVC2 | -1.13 | 3.55E-10 | -1.23 | 1.88E-18 | DE in both female and male placentas |
| ENSRNOG00000021086 | | DTX4 | -1.13 | 4.50E-06 | -1.51 | 5.15E-15 | DE in both female and male placentas |
| ENSRNOG00000011702 | X | FOXP3 | -1.13 | 2.43E-03 | -1.31 | 3.01E-06 | DE in both female and male placentas |
| ENSRNOG00000011193 | | HMGCLL1 | -1.13 | 2.12E-03 | -1.27 | 1.68E-05 | DE in both female and male placentas |
| ENSRNOG00000018355 | | MSX2 | -1.13 | 1.02E-05 | -1.57 | 1.01E-15 | DE in both female and male placentas |
| ENSRNOG00000050539 | | FBLN5 | -1.13 | 2.18E-06 | -1.81 | 8.41E-22 | DE in both female and male placentas |
| ENSRNOG00000017198 | | HIF3A | -1.13 | 6.17E-03 | -1.15 | 1.88E-04 | DE in both female and male placentas |
| ENSRNOG00000014142 | | OGFRL1 | -1.14 | 1.73E-05 | -1.17 | 7.36E-09 | DE in both female and male placentas |
| ENSRNOG00000019778 | | CAVIN1 | -1.14 | 5.37E-21 | -1.13 | 3.16E-32 | DE in both female and male placentas |
| ENSRNOG00000017767 | | MRVI1 | -1.14 | 3.80E-12 | -1.22 | 1.19E-21 | DE in both female and male placentas |
| ENSRNOG00000026978 | | OSCP1 | -1.14 | 1.10E-07 | -1.69 | 3.48E-24 | DE in both female and male placentas |
| ENSRNOG00000013934 | | DENND2B | -1.14 | 2.38E-07 | -1.50 | 1.11E-18 | DE in both female and male placentas |
| ENSRNOG00000060775 | | LMO7 | -1.14 | 2.07E-05 | -1.88 | 5.33E-20 | DE in both female and male placentas |
| ENSRNOG00000017511 | | DCDC2 | -1.14 | 2.63E-02 | -2.22 | 8.49E-10 | DE in both female and male placentas |
| ENSRNOG00000030981 | | SCNN1B | -1.14 | 1.98E-03 | -1.75 | 1.11E-10 | DE in both female and male placentas |

| | | | | | | | |
|---|---|---|---|---|---|---|---|
| ENSRNOG00000007682 | X | GRIA3 | -1.15 | 1.45E-03 | -1.67 | 2.25E-09 | DE in both female and male placentas |
| ENSRNOG00000027557 | | KLF14 | -1.15 | 1.90E-05 | -1.24 | 1.54E-09 | DE in both female and male placentas |
| ENSRNOG00000019098 | | CA5A | -1.15 | 3.54E-02 | -1.35 | 4.09E-04 | DE in both female and male placentas |
| ENSRNOG00000010350 | | RCAN2 | -1.15 | 3.09E-07 | -1.19 | 1.53E-11 | DE in both female and male placentas |
| ENSRNOG00000012060 | | GUCY1B1 | -1.15 | 2.31E-07 | -1.34 | 6.61E-15 | DE in both female and male placentas |
| ENSRNOG00000018709 | | PATZ1 | -1.15 | 2.11E-15 | -1.11 | 5.40E-22 | DE in both female and male placentas |
| ENSRNOG00000009157 | | FUT4 | -1.15 | 1.60E-07 | -1.49 | 1.76E-18 | DE in both female and male placentas |
| ENSRNOG00000055293 | | PTPRB | -1.15 | 1.80E-03 | -1.91 | 2.48E-12 | DE in both female and male placentas |
| ENSRNOG00000016848 | | FZD4 | -1.15 | 1.39E-05 | -1.76 | 3.87E-18 | DE in both female and male placentas |
| ENSRNOG00000043085 | | CLSTN2 | -1.16 | 1.52E-04 | -1.76 | 1.41E-13 | DE in both female and male placentas |
| ENSRNOG00000010593 | | CTNNAL1 | -1.16 | 4.64E-08 | -1.50 | 6.90E-20 | DE in both female and male placentas |
| ENSRNOG00000010205 | | MTURN | -1.16 | 2.37E-08 | -1.68 | 4.81E-25 | DE in both female and male placentas |
| ENSRNOG00000057865 | | PRSS16 | -1.16 | 2.02E-02 | -1.83 | 1.53E-07 | DE in both female and male placentas |
| ENSRNOG00000009785 | | CDKN3 | -1.16 | 1.75E-05 | -1.61 | 5.63E-15 | DE in both female and male placentas |
| ENSRNOG00000008931 | | SLC25A21 | -1.16 | 1.39E-03 | -1.71 | 5.46E-10 | DE in both female and male placentas |
| ENSRNOG00000020309 | | GFRA3 | -1.16 | 1.74E-06 | -1.02 | 6.36E-08 | DE in both female and male placentas |
| ENSRNOG00000008380 | | ASIC3 | -1.17 | 2.52E-02 | -1.69 | 3.85E-06 | DE in both female and male placentas |
| ENSRNOG00000006784 | | SEMA4F | -1.17 | 1.52E-05 | -1.39 | 1.22E-11 | DE in both female and male placentas |

| | | | | | | | |
|---|---|---|---|---|---|---|---|
| ENSRNOG00000027276 | | XKR4 | -1.17 | 6.22E-04 | -1.13 | 1.65E-05 | DE in both female and male placentas |
| ENSRNOG00000010725 | | CPA1 | -1.17 | 1.40E-05 | -1.37 | 2.21E-11 | DE in both female and male placentas |
| ENSRNOG00000018122 | | TSPAN17 | -1.17 | 6.21E-08 | -1.09 | 7.18E-11 | DE in both female and male placentas |
| ENSRNOG00000049537 | | AABR07021544.1 | -1.17 | 5.48E-08 | -1.68 | 9.17E-24 | DE in both female and male placentas |
| ENSRNOG00000000981 | | SCARB1 | -1.17 | 3.05E-11 | -1.23 | 2.56E-19 | DE in both female and male placentas |
| ENSRNOG00000012294 | X | HEPH | -1.17 | 8.55E-03 | -1.79 | 2.12E-08 | DE in both female and male placentas |
| ENSRNOG00000020401 | | ADCY4 | -1.17 | 1.12E-06 | -1.03 | 2.73E-08 | DE in both female and male placentas |
| ENSRNOG00000023812 | | RAVER2 | -1.17 | 1.11E-08 | -1.33 | 8.37E-16 | DE in both female and male placentas |
| ENSRNOG00000011417 | | PDE3B | -1.17 | 1.12E-03 | -1.75 | 4.73E-11 | DE in both female and male placentas |
| ENSRNOG00000025695 | | TNS3 | -1.17 | 3.37E-13 | -1.41 | 3.89E-29 | DE in both female and male placentas |
| ENSRNOG00000023951 | | TBC1D30 | -1.17 | 3.17E-02 | -1.77 | 4.87E-06 | DE in both female and male placentas |
| ENSRNOG00000040108 | | RGD1565355 | -1.17 | 6.21E-04 | -1.96 | 2.03E-14 | DE in both female and male placentas |
| ENSRNOG00000019014 | | NDST1 | -1.17 | 2.77E-11 | -1.25 | 9.45E-20 | DE in both female and male placentas |
| ENSRNOG00000021134 | | ANXA9 | -1.17 | 1.37E-04 | -1.29 | 4.34E-08 | DE in both female and male placentas |
| ENSRNOG00000010716 | | ATOH8 | -1.17 | 1.88E-03 | -1.40 | 6.74E-07 | DE in both female and male placentas |
| ENSRNOG00000003687 | | RGS2 | -1.17 | 4.55E-05 | -1.38 | 3.42E-10 | DE in both female and male placentas |
| ENSRNOG00000003732 | | FLRT2 | -1.18 | 1.18E-06 | -1.32 | 1.16E-12 | DE in both female and male placentas |
| ENSRNOG00000001216 | | TRPM2 | -1.18 | 2.10E-04 | -1.72 | 5.50E-13 | DE in both female and male placentas |

| | | | | | | |
|---|---|---|---|---|---|---|
| ENSRNOG00000007256 | NECAB1 | -1.18 | 1.20E-02 | -1.99 | 2.57E-08 | DE in both female and male placentas |
| ENSRNOG00000061910 | IGFBP3 | -1.18 | 4.58E-07 | -1.59 | 1.54E-18 | DE in both female and male placentas |
| ENSRNOG00000015075 | STC1 | -1.18 | 5.15E-05 | -1.15 | 2.58E-07 | DE in both female and male placentas |
| ENSRNOG00000056944 | ARHGAP24 | -1.18 | 2.34E-07 | -1.63 | 3.16E-20 | DE in both female and male placentas |
| ENSRNOG00000054391 | SNURF | -1.18 | 3.22E-07 | -1.24 | 1.86E-12 | DE in both female and male placentas |
| ENSRNOG00000060329 | EMB | -1.18 | 7.12E-04 | -1.77 | 9.96E-12 | DE in both female and male placentas |
| ENSRNOG00000008602 | STEAP4 | -1.19 | 5.47E-07 | -1.09 | 2.68E-09 | DE in both female and male placentas |
| ENSRNOG00000016618 | TMEM52 | -1.19 | 6.13E-04 | -1.27 | 2.04E-06 | DE in both female and male placentas |
| ENSRNOG00000018059 | IHH | -1.19 | 1.96E-04 | -1.12 | 4.05E-06 | DE in both female and male placentas |
| ENSRNOG00000013654 | CBLN2 | -1.19 | 3.81E-02 | -1.20 | 5.56E-03 | DE in both female and male placentas |
| ENSRNOG00000010960 | ANKH | -1.19 | 1.06E-09 | -1.54 | 4.46E-24 | DE in both female and male placentas |
| ENSRNOG00000011927 | SDC3 | -1.19 | 2.97E-17 | -1.20 | 9.57E-27 | DE in both female and male placentas |
| ENSRNOG00000014243 | PEAR1 | -1.19 | 7.74E-09 | -1.15 | 9.00E-13 | DE in both female and male placentas |
| ENSRNOG00000033915 | GPT | -1.20 | 4.94E-06 | -1.24 | 5.10E-10 | DE in both female and male placentas |
| ENSRNOG00000000588 | SLC16A10 | -1.20 | 3.05E-04 | -1.77 | 1.25E-12 | DE in both female and male placentas |
| ENSRNOG00000007637 | ACER2 | -1.20 | 4.80E-08 | -1.51 | 7.04E-19 | DE in both female and male placentas |
| ENSRNOG00000023452 | AABR070504 49.1 | -1.20 | 1.15E-02 | -1.49 | 8.51E-06 | DE in both female and male placentas |
| ENSRNOG00000013920 | ARHGAP26 | -1.20 | 5.24E-07 | -1.62 | 1.52E-18 | DE in both female and male placentas |

| | | | | | | | |
|---|---|---|---|---|---|---|---|
| ENSRNOG00000016945 | | PLA2G2A | -1.20 | 1.67E-05 | -1.36 | 1.82E-10 | DE in both female and male placentas |
| ENSRNOG00000002449 | X | MAGED2 | -1.20 | 2.35E-21 | -1.16 | 6.65E-31 | DE in both female and male placentas |
| ENSRNOG00000056184 | | LRP2 | -1.20 | 1.59E-03 | -1.65 | 4.62E-09 | DE in both female and male placentas |
| ENSRNOG00000009332 | | REC114 | -1.21 | 1.12E-04 | -1.63 | 9.82E-12 | DE in both female and male placentas |
| ENSRNOG00000009620 | | CYBRD1 | -1.21 | 4.22E-14 | -1.63 | 1.08E-37 | DE in both female and male placentas |
| ENSRNOG00000017283 | | KCNT1 | -1.21 | 3.47E-07 | -1.01 | 4.27E-08 | DE in both female and male placentas |
| ENSRNOG00000011134 | | LAMA2 | -1.21 | 1.16E-07 | -1.32 | 1.05E-13 | DE in both female and male placentas |
| ENSRNOG00000024799 | | CD93 | -1.21 | 3.49E-05 | -1.45 | 6.77E-11 | DE in both female and male placentas |
| ENSRNOG00000004110 | | TRIB2 | -1.21 | 2.58E-09 | -1.54 | 4.01E-22 | DE in both female and male placentas |
| ENSRNOG00000000566 | | PCBD1 | -1.22 | 1.53E-06 | -1.39 | 8.27E-13 | DE in both female and male placentas |
| ENSRNOG00000007243 | | HAVCR1 | -1.22 | 1.24E-02 | -1.34 | 8.89E-05 | DE in both female and male placentas |
| ENSRNOG00000021713 | | KIF18B | -1.22 | 5.17E-05 | -1.30 | 5.32E-09 | DE in both female and male placentas |
| ENSRNOG00000000563 | | ADAMTS14 | -1.22 | 7.78E-09 | -1.27 | 1.24E-14 | DE in both female and male placentas |
| ENSRNOG00000013364 | | HEY2 | -1.22 | 1.04E-04 | -1.31 | 5.80E-08 | DE in both female and male placentas |
| ENSRNOG00000016474 | | TSPAN3 | -1.22 | 8.40E-22 | -1.35 | 7.30E-41 | DE in both female and male placentas |
| ENSRNOG00000010658 | | LOC103691744 | -1.22 | 1.05E-02 | -1.57 | 4.56E-06 | DE in both female and male placentas |
| ENSRNOG00000001217 | | LRRC3 | -1.22 | 1.51E-06 | -1.07 | 1.67E-07 | DE in both female and male placentas |
| ENSRNOG00000000010 | | CBLN1 | -1.22 | 2.25E-04 | -1.78 | 3.69E-12 | DE in both female and male placentas |

| | | | | | | |
|---|---|---|---|---|---|---|
| ENSRNOG00000020927 | TULP2 | -1.22 | 2.58E-08 | -1.40 | 2.11E-16 | DE in both female and male placentas |
| ENSRNOG00000021276 | BMP2 | -1.22 | 4.94E-08 | -1.15 | 2.71E-11 | DE in both female and male placentas |
| ENSRNOG00000052025 | TLE5 | -1.22 | 2.14E-13 | -1.03 | 2.00E-15 | DE in both female and male placentas |
| ENSRNOG00000046151 | TUBB1 | -1.22 | 8.60E-06 | -1.39 | 5.40E-11 | DE in both female and male placentas |
| ENSRNOG00000014745 | AABR070266 54.1 | -1.22 | 6.97E-11 | -1.18 | 4.25E-16 | DE in both female and male placentas |
| ENSRNOG00000013428 | ATP6V0A4 | -1.22 | 7.24E-03 | -2.05 | 4.07E-10 | DE in both female and male placentas |
| ENSRNOG00000018577 | NEIL1 | -1.23 | 2.07E-05 | -1.95 | 9.25E-19 | DE in both female and male placentas |
| ENSRNOG00000011334 | TMEM63C | -1.23 | 2.67E-07 | -1.23 | 2.10E-11 | DE in both female and male placentas |
| ENSRNOG00000010695 | PDGFC | -1.23 | 8.83E-07 | -1.57 | 3.80E-16 | DE in both female and male placentas |
| ENSRNOG00000021381 | CCDC68 | -1.23 | 2.78E-04 | -1.32 | 3.37E-07 | DE in both female and male placentas |
| ENSRNOG00000058752 | POPDC2 | -1.23 | 1.45E-06 | -1.62 | 2.47E-16 | DE in both female and male placentas |
| ENSRNOG00000017914 | CAVIN3 | -1.23 | 1.24E-11 | -1.05 | 1.67E-13 | DE in both female and male placentas |
| ENSRNOG00000018338 | VWA1 | -1.23 | 7.62E-06 | -1.40 | 2.59E-11 | DE in both female and male placentas |
| ENSRNOG00000000547 | TSPYL4 | -1.23 | 5.33E-07 | -1.75 | 4.52E-18 | DE in both female and male placentas |
| ENSRNOG00000036631 | PTGDR2 | -1.23 | 1.08E-04 | -1.16 | 1.18E-06 | DE in both female and male placentas |
| ENSRNOG00000051690 | CLEC9A | -1.23 | 5.31E-03 | -1.88 | 8.85E-09 | DE in both female and male placentas |
| ENSRNOG00000004532 | DIPK1B | -1.23 | 7.30E-12 | -1.04 | 8.41E-14 | DE in both female and male placentas |
| ENSRNOG00000004874 | FLRT3 | -1.24 | 4.12E-08 | -1.40 | 7.81E-16 | DE in both female and male placentas |

| | | | | | | |
|---|---|---|---|---|---|---|
| ENSRNOG00000038202 | CALML4 | -1.24 | 3.91E-05 | -1.27 | 2.84E-08 | DE in both female and male placentas |
| ENSRNOG00000050949 | TTC39C | -1.24 | 1.06E-08 | -1.54 | 6.55E-20 | DE in both female and male placentas |
| ENSRNOG00000003374 | CCDC85A | -1.24 | 1.68E-04 | -1.65 | 2.98E-10 | DE in both female and male placentas |
| ENSRNOG00000047158 | SELENBP1 | -1.24 | 1.04E-02 | -2.07 | 2.55E-09 | DE in both female and male placentas |
| ENSRNOG00000029830 | ADM2 | -1.24 | 2.86E-02 | -1.04 | 6.01E-03 | DE in both female and male placentas |
| ENSRNOG00000028072 | CHIT1 | -1.24 | 1.20E-05 | -1.35 | 7.26E-10 | DE in both female and male placentas |
| ENSRNOG00000027230 | FHOD3 | -1.24 | 4.62E-04 | -1.88 | 1.49E-12 | DE in both female and male placentas |
| ENSRNOG00000006470 | CAMK1G | -1.24 | 1.55E-05 | -1.11 | 3.91E-07 | DE in both female and male placentas |
| ENSRNOG00000010731 | GPM6A | -1.24 | 1.26E-02 | -1.11 | 4.80E-03 | DE in both female and male placentas |
| ENSRNOG00000021004 | RASIP1 | -1.25 | 4.66E-10 | -1.31 | 5.11E-17 | DE in both female and male placentas |
| ENSRNOG00000047914 | GNG11 | -1.25 | 3.48E-07 | -1.24 | 5.28E-11 | DE in both female and male placentas |
| ENSRNOG00000005987 | SUOX | -1.25 | 1.56E-07 | -1.76 | 1.44E-21 | DE in both female and male placentas |
| ENSRNOG00000027724 | PLEKHF1 | -1.25 | 9.05E-11 | -1.01 | 1.23E-11 | DE in both female and male placentas |
| ENSRNOG00000012302 | GUCY1A1 | -1.25 | 7.03E-05 | -2.01 | 6.82E-17 | DE in both female and male placentas |
| ENSRNOG00000006970 | NTN3 | -1.26 | 6.80E-03 | -1.14 | 1.09E-03 | DE in both female and male placentas |
| ENSRNOG00000007726 | MCAM | -1.26 | 4.04E-06 | -1.59 | 2.51E-14 | DE in both female and male placentas |
| ENSRNOG00000049952 | GUCD1 | -1.26 | 2.16E-14 | -1.37 | 2.74E-26 | DE in both female and male placentas |
| ENSRNOG00000032238 | PLCD1 | -1.26 | 1.28E-10 | -1.51 | 4.58E-23 | DE in both female and male placentas |

| | | | | | | | |
|---|---|---|---|---|---|---|---|
| ENSRNOG00000050964 | | LOC100911068 | -1.26 | 1.95E-04 | -1.17 | 6.29E-06 | DE in both female and male placentas |
| ENSRNOG00000055251 | | PRTG | -1.26 | 4.05E-05 | -2.20 | 1.07E-20 | DE in both female and male placentas |
| ENSRNOG00000003613 | X | MID1 | -1.26 | 3.41E-14 | -1.24 | 3.59E-21 | DE in both female and male placentas |
| ENSRNOG00000012659 | | VAMP5 | -1.27 | 4.17E-07 | -1.01 | 1.58E-07 | DE in both female and male placentas |
| ENSRNOG00000026238 | | INSYN1 | -1.27 | 6.06E-12 | -1.07 | 1.40E-13 | DE in both female and male placentas |
| ENSRNOG00000009802 | | FAM124A | -1.27 | 1.62E-04 | -1.98 | 1.40E-13 | DE in both female and male placentas |
| ENSRNOG00000028353 | | SOX13 | -1.27 | 1.08E-12 | -1.50 | 1.54E-26 | DE in both female and male placentas |
| ENSRNOG00000059764 | | SNRPN | -1.27 | 2.48E-10 | -1.13 | 4.79E-13 | DE in both female and male placentas |
| ENSRNOG00000047175 | | CRYAA | -1.27 | 2.53E-02 | -1.08 | 9.71E-03 | DE in both female and male placentas |
| ENSRNOG00000021855 | | KCP | -1.27 | 4.82E-04 | -1.01 | 3.16E-04 | DE in both female and male placentas |
| ENSRNOG00000014597 | | IRS1 | -1.28 | 9.58E-10 | -1.47 | 1.50E-19 | DE in both female and male placentas |
| ENSRNOG00000000561 | | PALD1 | -1.28 | 2.44E-15 | -1.13 | 4.64E-19 | DE in both female and male placentas |
| ENSRNOG00000060356 | | KIF15 | -1.28 | 2.03E-08 | -1.84 | 2.31E-25 | DE in both female and male placentas |
| ENSRNOG00000017249 | | ZFP366 | -1.28 | 2.77E-03 | -2.04 | 1.21E-10 | DE in both female and male placentas |
| ENSRNOG00000005903 | | ST14 | -1.28 | 2.49E-14 | -1.27 | 7.39E-22 | DE in both female and male placentas |
| ENSRNOG00000000636 | | RTKN2 | -1.29 | 2.04E-05 | -1.98 | 3.13E-17 | DE in both female and male placentas |
| ENSRNOG00000010947 | | MMP14 | -1.29 | 2.18E-37 | -1.15 | 1.14E-45 | DE in both female and male placentas |
| ENSRNOG00000015093 | | SPARCL1 | -1.29 | 1.50E-04 | -1.99 | 9.58E-15 | DE in both female and male placentas |

| | | | | | | | |
|---|---|---|---|---|---|---|---|
| ENSRNOG00000025051 | | TAL1 | -1.29 | 2.79E-05 | -1.67 | 1.85E-12 | DE in both female and male placentas |
| ENSRNOG00000017307 | | PRSS23 | -1.29 | 3.48E-07 | -1.54 | 2.44E-15 | DE in both female and male placentas |
| ENSRNOG00000030763 | | DPP4 | -1.29 | 3.24E-04 | -2.06 | 3.35E-14 | DE in both female and male placentas |
| ENSRNOG00000032208 | X | EDA | -1.29 | 2.44E-03 | -1.25 | 1.80E-04 | DE in both female and male placentas |
| ENSRNOG00000007740 | | TACSTD2 | -1.30 | 2.32E-04 | -1.46 | 3.98E-08 | DE in both female and male placentas |
| ENSRNOG00000002800 | X | GDPD2 | -1.30 | 1.95E-02 | -1.83 | 7.38E-06 | DE in both female and male placentas |
| ENSRNOG00000060348 | | ZDHHC23 | -1.30 | 3.58E-06 | -1.83 | 1.77E-17 | DE in both female and male placentas |
| ENSRNOG00000006410 | | AKAP5 | -1.30 | 1.82E-04 | -2.28 | 6.33E-18 | DE in both female and male placentas |
| ENSRNOG00000060528 | | CACNA1G | -1.30 | 2.40E-05 | -1.79 | 6.43E-14 | DE in both female and male placentas |
| ENSRNOG00000008336 | | TNFRSF11B | -1.30 | 9.51E-05 | -1.27 | 6.54E-07 | DE in both female and male placentas |
| ENSRNOG00000058609 | | PALMD | -1.31 | 4.31E-04 | -1.88 | 2.25E-11 | DE in both female and male placentas |
| ENSRNOG00000061650 | | AC128059.5 | -1.31 | 1.92E-02 | -1.42 | 3.46E-04 | DE in both female and male placentas |
| ENSRNOG00000019388 | | EGFL7 | -1.31 | 2.57E-05 | -1.17 | 1.08E-06 | DE in both female and male placentas |
| ENSRNOG00000032703 | | RASGRP3 | -1.31 | 2.89E-08 | -1.47 | 1.05E-15 | DE in both female and male placentas |
| ENSRNOG00000001924 | | TP63 | -1.31 | 6.06E-03 | -2.24 | 1.13E-10 | DE in both female and male placentas |
| ENSRNOG00000011631 | | FST | -1.31 | 1.07E-03 | -1.60 | 8.70E-08 | DE in both female and male placentas |
| ENSRNOG00000026217 | | ARMC2 | -1.31 | 2.39E-05 | -1.02 | 2.09E-05 | DE in both female and male placentas |
| ENSRNOG00000004589 | | GALNT16 | -1.31 | 3.92E-10 | -1.36 | 1.03E-16 | DE in both female and male placentas |

| | | | | | | | |
|---|---|---|---|---|---|---|---|
| ENSRNOG00000008466 | | LBX2 | -1.31 | 1.38E-02 | -2.11 | 1.97E-08 | DE in both female and male placentas |
| ENSRNOG00000018020 | | APBB1 | -1.31 | 1.03E-16 | -1.18 | 2.57E-21 | DE in both female and male placentas |
| ENSRNOG00000039560 | | LOC1036901 16 | -1.32 | 1.32E-02 | -2.11 | 7.93E-08 | DE in both female and male placentas |
| ENSRNOG00000060262 | X | PRPS1 | -1.32 | 3.58E-07 | -1.83 | 6.15E-20 | DE in both female and male placentas |
| ENSRNOG00000019342 | | SULT1A1 | -1.32 | 7.57E-06 | -1.90 | 5.55E-17 | DE in both female and male placentas |
| ENSRNOG00000004280 | | TCN2 | -1.32 | 1.96E-08 | -1.82 | 2.95E-23 | DE in both female and male placentas |
| ENSRNOG00000012119 | | AABR070692 82.1 | -1.33 | 6.21E-06 | -1.35 | 2.11E-09 | DE in both female and male placentas |
| ENSRNOG00000058666 | | AABR070433 65.1 | -1.33 | 1.66E-04 | -1.75 | 1.51E-11 | DE in both female and male placentas |
| ENSRNOG00000017602 | | IL34 | -1.34 | 1.56E-08 | -1.58 | 1.67E-17 | DE in both female and male placentas |
| ENSRNOG00000008394 | | PRG2 | -1.34 | 1.67E-03 | -1.55 | 3.93E-06 | DE in both female and male placentas |
| ENSRNOG00000007955 | | TIMP4 | -1.34 | 1.14E-05 | -2.19 | 3.99E-19 | DE in both female and male placentas |
| ENSRNOG00000010389 | | NDRG2 | -1.34 | 2.03E-05 | -2.32 | 1.60E-21 | DE in both female and male placentas |
| ENSRNOG00000001219 | | TSPEAR | -1.34 | 3.00E-02 | -2.45 | 2.65E-08 | DE in both female and male placentas |
| ENSRNOG00000053766 | | RAMP3 | -1.35 | 1.21E-03 | -1.70 | 3.72E-08 | DE in both female and male placentas |
| ENSRNOG00000009835 | | TRAM1L1 | -1.35 | 6.23E-03 | -1.73 | 5.18E-06 | DE in both female and male placentas |
| ENSRNOG00000012791 | | AJUBA | -1.35 | 5.58E-09 | -1.72 | 1.25E-21 | DE in both female and male placentas |
| ENSRNOG00000046204 | | TSPAN15 | -1.35 | 1.02E-04 | -1.97 | 7.16E-14 | DE in both female and male placentas |
| ENSRNOG00000060460 | | CLEC1A | -1.35 | 2.67E-05 | -1.95 | 2.42E-15 | DE in both female and male placentas |

| | | | | | | |
|---|---|---|---|---|---|---|
| ENSRNOG00000029286 | HBE1 | -1.35 | 3.01E-02 | -2.28 | 1.72E-07 | DE in both female and male placentas |
| ENSRNOG00000002305 | SLC15A2 | -1.36 | 7.60E-04 | -2.13 | 1.84E-12 | DE in both female and male placentas |
| ENSRNOG00000030210 | FNDC1 | -1.36 | 1.36E-04 | -2.22 | 4.80E-16 | DE in both female and male placentas |
| ENSRNOG00000029876 | GUCY1A2 | -1.36 | 1.21E-07 | -2.03 | 1.36E-23 | DE in both female and male placentas |
| ENSRNOG00000036677 | SLC16A3 | -1.36 | 2.70E-07 | -1.52 | 6.84E-14 | DE in both female and male placentas |
| ENSRNOG00000000275 | FAM13C | -1.36 | 3.87E-09 | -1.28 | 4.67E-13 | DE in both female and male placentas |
| ENSRNOG00000006033 | LOC100910790 | -1.36 | 8.37E-08 | -1.38 | 2.07E-12 | DE in both female and male placentas |
| ENSRNOG00000021823 | AABR07018318.1 | -1.36 | 2.86E-02 | -2.75 | 4.07E-10 | DE in both female and male placentas |
| ENSRNOG00000019609 | B4GALT2 | -1.37 | 1.24E-10 | -1.69 | 7.07E-23 | DE in both female and male placentas |
| ENSRNOG00000023485 | USHBP1 | -1.37 | 2.48E-06 | -1.10 | 1.23E-06 | DE in both female and male placentas |
| ENSRNOG00000012058 | EGFLAM | -1.37 | 1.28E-06 | -1.65 | 3.61E-14 | DE in both female and male placentas |
| ENSRNOG00000005854 | ANGPT1 | -1.37 | 9.20E-08 | -1.95 | 3.52E-22 | Differentially expressed in both female and male |
| ENSRNOG00000008706 | TBX3 | -1.37 | 1.08E-05 | -1.94 | 4.11E-16 | DE in both female and male placentas |
| ENSRNOG00000055765 | RASL10B | -1.38 | 8.66E-03 | -1.82 | 6.15E-07 | DE in both female and male placentas |
| ENSRNOG00000013053 | TRPM6 | -1.38 | 1.51E-05 | -1.90 | 4.76E-15 | DE in both female and male placentas |
| ENSRNOG00000042771 | APOL3 | -1.38 | 1.14E-07 | -1.08 | 7.08E-08 | DE in both female and male placentas |
| ENSRNOG00000024569 | GIMAP9 | -1.38 | 1.32E-04 | -1.10 | 1.26E-04 | DE in both female and male placentas |
| ENSRNOG00000011794 | TCEA3 | -1.38 | 3.22E-05 | -1.90 | 4.74E-14 | DE in both female and male placentas |

| | | | | | | |
|---|---|---|---|---|---|---|
| ENSRNOG00000001752 | NRROS | -1.38 | 5.29E-19 | -1.24 | 5.95E-24 | DE in both female and male placentas |
| ENSRNOG00000018346 | AGTR1A | -1.38 | 5.15E-10 | -1.25 | 3.69E-13 | DE in both female and male placentas |
| ENSRNOG00000026919 | RGD1565367 | -1.38 | 2.75E-04 | -1.65 | 9.24E-09 | DE in both female and male placentas |
| ENSRNOG00000003273 | SPATA20 | -1.38 | 7.29E-07 | -1.77 | 1.15E-16 | DE in both female and male placentas |
| ENSRNOG00000001979 | RCAN1 | -1.38 | 2.32E-07 | -1.67 | 4.21E-16 | DE in both female and male placentas |
| ENSRNOG00000004810 | PLCB1 | -1.38 | 7.79E-07 | -1.63 | 5.23E-14 | DE in both female and male placentas |
| ENSRNOG00000052296 | SHANK3 | -1.38 | 4.36E-06 | -1.67 | 6.04E-13 | DE in both female and male placentas |
| ENSRNOG00000047734 | CHST2 | -1.39 | 1.27E-09 | -1.24 | 2.15E-12 | DE in both female and male placentas |
| ENSRNOG00000026902 | LYVE1 | -1.39 | 3.33E-03 | -2.21 | 1.86E-10 | DE in both female and male placentas |
| ENSRNOG00000027954 | CDH15 | -1.39 | 5.38E-03 | -1.87 | 9.36E-07 | DE in both female and male placentas |
| ENSRNOG00000038496 | TEX21 | -1.39 | 3.90E-04 | -1.71 | 1.06E-08 | DE in both female and male placentas |
| ENSRNOG00000024210 | NLRC3 | -1.39 | 1.90E-03 | -1.30 | 1.52E-04 | DE in both female and male placentas |
| ENSRNOG00000020196 | SCGB1A1 | -1.39 | 9.14E-04 | -1.42 | 9.76E-06 | DE in both female and male placentas |
| ENSRNOG00000042309 | KIAA0408L | -1.39 | 1.38E-06 | -1.75 | 1.32E-14 | DE in both female and male placentas |
| ENSRNOG00000005370 | S1PR4 | -1.39 | 9.16E-07 | -1.34 | 1.21E-09 | DE in both female and male placentas |
| ENSRNOG00000024751 | MACROH2A2 | -1.39 | 5.93E-06 | -1.90 | 9.79E-16 | DE in both female and male placentas |
| ENSRNOG00000050675 | MYL4 | -1.39 | 1.01E-09 | -1.25 | 8.74E-13 | DE in both female and male placentas |
| ENSRNOG00000020733 | CAMP | -1.39 | 1.69E-02 | -1.23 | 1.93E-03 | DE in both female and male placentas |

| | | | | | | |
|---|---|---|---|---|---|---|
| ENSRNOG00000010050 | STARD9 | -1.39 | 2.03E-15 | -1.70 | 3.26E-34 | DE in both female and male placentas |
| ENSRNOG00000018839 | NTRK2 | -1.40 | 4.33E-06 | -2.03 | 1.71E-17 | DE in both female and male placentas |
| ENSRNOG00000002599 | GRAP | -1.40 | 2.36E-08 | -1.28 | 3.27E-11 | DE in both female and male placentas |
| ENSRNOG00000008356 | MYO5C | -1.40 | 2.32E-07 | -2.17 | 3.89E-24 | DE in both female and male placentas |
| ENSRNOG00000005359 | CSRNP3 | -1.40 | 1.74E-03 | -2.32 | 4.77E-12 | DE in both female and male placentas |
| ENSRNOG00000016491 | FAM110D | -1.41 | 4.41E-03 | -1.67 | 5.68E-06 | DE in both female and male placentas |
| ENSRNOG00000059618 | SCARF1 | -1.41 | 7.25E-06 | -1.17 | 8.31E-07 | DE in both female and male placentas |
| ENSRNOG00000008257 | MFAP2 | -1.41 | 6.22E-06 | -1.49 | 4.05E-10 | DE in both female and male placentas |
| ENSRNOG00000011044 | CLMN | -1.41 | 1.62E-04 | -2.38 | 7.47E-17 | DE in both female and male placentas |
| ENSRNOG00000006144 | CNTN3 | -1.41 | 1.33E-03 | -2.31 | 5.46E-12 | DE in both female and male placentas |
| ENSRNOG00000011263 | PLAC9 | -1.41 | 3.21E-08 | -1.91 | 1.24E-20 | DE in both female and male placentas |
| ENSRNOG00000007636 | CADPS2 | -1.42 | 4.81E-04 | -1.57 | 4.63E-07 | DE in both female and male placentas |
| ENSRNOG00000027193 | AABR070513 76.1 | -1.42 | 5.29E-07 | -1.81 | 1.24E-16 | DE in both female and male placentas |
| ENSRNOG00000027736 | CNN1 | -1.42 | 2.58E-06 | -1.02 | 1.25E-05 | DE in both female and male placentas |
| ENSRNOG00000057256 | SLC19A3 | -1.42 | 2.26E-05 | -1.54 | 1.42E-09 | DE in both female and male placentas |
| ENSRNOG00000005708 | MMP16 | -1.42 | 9.87E-05 | -1.66 | 1.87E-09 | DE in both female and male placentas |
| ENSRNOG00000025143 | ICAM2 | -1.42 | 1.98E-04 | -1.92 | 2.36E-11 | DE in both female and male placentas |
| ENSRNOG00000010262 | HDC | -1.42 | 1.38E-02 | -1.99 | 1.65E-06 | DE in both female and male placentas |

| | | | | | | |
|---|---|---|---|---|---|---|
| ENSRNOG00000016543 | TRIM63 | -1.42 | 1.25E-02 | -1.77 | 1.68E-05 | DE in both female and male placentas |
| ENSRNOG00000039388 | PRRT4 | -1.42 | 1.07E-03 | -1.40 | 5.39E-05 | DE in both female and male placentas |
| ENSRNOG00000005619 | MISP3 | -1.42 | 6.62E-03 | -1.79 | 4.97E-06 | DE in both female and male placentas |
| ENSRNOG00000012108 | THSD1 | -1.42 | 2.28E-11 | -1.61 | 2.45E-22 | DE in both female and male placentas |
| ENSRNOG00000019700 | PROZ | -1.42 | 3.71E-02 | -2.24 | 3.40E-06 | DE in both female and male placentas |
| ENSRNOG00000011200 | BHMT | -1.42 | 7.45E-03 | -2.44 | 2.31E-10 | DE in both female and male placentas |
| ENSRNOG00000006604 | THY1 | -1.42 | 4.45E-05 | -1.79 | 1.50E-11 | DE in both female and male placentas |
| ENSRNOG00000014714 | TRPV6 | -1.42 | 3.77E-02 | -2.02 | 2.29E-05 | DE in both female and male placentas |
| ENSRNOG00000002930 | PPL | -1.42 | 2.02E-07 | -1.84 | 3.38E-18 | DE in both female and male placentas |
| ENSRNOG00000037137 | SVIP | -1.43 | 2.67E-07 | -1.93 | 3.16E-18 | DE in both female and male placentas |
| ENSRNOG00000029792 | NEWGENE_1308171 | -1.43 | 2.69E-04 | -2.30 | 5.32E-14 | DE in both female and male placentas |
| ENSRNOG00000020585 | TBXA2R | -1.43 | 5.03E-04 | -1.56 | 8.24E-07 | DE in both female and male placentas |
| ENSRNOG00000015794 | FAM83D | -1.43 | 2.04E-17 | -1.56 | 7.44E-31 | DE in both female and male placentas |
| ENSRNOG00000001232 | SLC19A1 | -1.43 | 4.95E-11 | -1.28 | 3.11E-14 | DE in both female and male placentas |
| ENSRNOG00000014851 | COL4A4 | -1.43 | 1.98E-02 | -2.40 | 4.08E-08 | DE in both female and male placentas |
| ENSRNOG00000061777 | AABR07024203.3 | -1.44 | 3.23E-03 | -2.34 | 1.74E-09 | DE in both female and male placentas |
| ENSRNOG00000031249 | SLCO1A2 | -1.44 | 9.11E-04 | -2.25 | 1.81E-11 | DE in both female and male placentas |
| ENSRNOG00000013426 | MRGPRF | -1.44 | 1.93E-04 | -1.84 | 1.90E-10 | DE in both female and male placentas |

| | | | | | | |
|---|---|---|---|---|---|---|
| ENSRNOG00000009709 | AC118772.1 | -1.44 | 2.33E-03 | -2.32 | 2.59E-10 | DE in both female and male placentas |
| ENSRNOG00000017901 | ACY3 | -1.44 | 3.55E-03 | -1.16 | 1.65E-03 | DE in both female and male placentas |
| ENSRNOG00000001419 | CLDN15 | -1.44 | 2.54E-08 | -1.35 | 2.75E-11 | DE in both female and male placentas |
| ENSRNOG00000012879 | FABP3 | -1.44 | 2.33E-12 | -1.52 | 1.47E-21 | DE in both female and male placentas |
| ENSRNOG00000013683 | S1PR1 | -1.44 | 3.10E-05 | -1.79 | 9.39E-12 | DE in both female and male placentas |
| ENSRNOG00000033338 | GIMAP6 | -1.44 | 1.69E-06 | -2.19 | 1.60E-20 | DE in both female and male placentas |
| ENSRNOG00000038607 | TMEM86B | -1.44 | 8.16E-05 | -1.70 | 9.94E-10 | DE in both female and male placentas |
| ENSRNOG00000050469 | LOC100912034 | -1.44 | 6.61E-05 | -2.65 | 4.56E-18 | DE in both female and male placentas |
| ENSRNOG00000027595 | TFAP2E | -1.44 | 1.39E-02 | -2.73 | 1.92E-10 | DE in both female and male placentas |
| ENSRNOG00000018895 | MCHR1 | -1.45 | 1.63E-02 | -1.76 | 5.93E-05 | DE in both female and male placentas |
| ENSRNOG00000037275 | TLCD2 | -1.45 | 1.13E-04 | -1.59 | 1.91E-08 | DE in both female and male placentas |
| ENSRNOG00000028531 | CCL25 | -1.45 | 2.54E-02 | -2.70 | 1.22E-08 | DE in both female and male placentas |
| ENSRNOG00000012061 | PRKCB | -1.45 | 5.03E-08 | -2.00 | 2.68E-22 | DE in both female and male placentas |
| ENSRNOG00000026490 | DNM3 | -1.45 | 3.92E-06 | -1.54 | 4.57E-10 | DE in both female and male placentas |
| ENSRNOG00000001874 | SLC7A4 | -1.46 | 5.75E-08 | -1.24 | 4.15E-09 | Differentially expressed in both female and male |
| ENSRNOG00000059215 | SLC66A2 | -1.46 | 2.79E-16 | -1.15 | 1.24E-16 | DE in both female and male placentas |
| ENSRNOG00000016890 | TPPP3 | -1.46 | 2.97E-06 | -1.96 | 5.84E-16 | DE in both female and male placentas |
| ENSRNOG00000057834 | LOC102553018 | -1.46 | 5.85E-04 | -1.08 | 1.52E-03 | DE in both female and male placentas |

| | | | | | | |
|---|---|---|---|---|---|---|
| ENSRNOG00000005271 | RAPGEF5 | -1.46 | 3.04E-07 | -1.99 | 2.27E-19 | DE in both female and male placentas |
| ENSRNOG00000021750 | ID1 | -1.46 | 1.10E-07 | -1.69 | 1.86E-15 | DE in both female and male placentas |
| ENSRNOG00000021793 | TLCD5 | -1.47 | 1.05E-09 | -2.06 | 1.09E-27 | DE in both female and male placentas |
| ENSRNOG00000053769 | SOAT2 | -1.47 | 9.32E-03 | -1.21 | 4.19E-03 | DE in both female and male placentas |
| ENSRNOG00000027357 | SOX17 | -1.47 | 8.85E-08 | -1.26 | 3.36E-09 | DE in both female and male placentas |
| ENSRNOG00000003952 | AABR07020879.1 | -1.47 | 3.23E-07 | -1.53 | 4.81E-12 | DE in both female and male placentas |
| ENSRNOG00000003669 | MYOCD | -1.48 | 7.50E-07 | -1.85 | 8.66E-16 | DE in both female and male placentas |
| ENSRNOG00000015406 | PGM5 | -1.48 | 2.06E-10 | -1.54 | 3.82E-17 | DE in both female and male placentas |
| ENSRNOG00000001469 | ELN | -1.48 | 4.88E-03 | -1.84 | 1.69E-06 | DE in both female and male placentas |
| ENSRNOG00000020173 | TIE1 | -1.49 | 5.21E-11 | -1.56 | 7.81E-19 | DE in both female and male placentas |
| ENSRNOG00000038948 | STAB2 | -1.49 | 8.83E-05 | -1.49 | 3.17E-07 | DE in both female and male placentas |
| ENSRNOG00000008161 | PDZK1IP1 | -1.50 | 6.04E-19 | -1.23 | 6.60E-21 | DE in both female and male placentas |
| ENSRNOG00000032728 | LOC689230 | -1.50 | 1.58E-03 | -1.27 | 3.82E-04 | DE in both female and male placentas |
| ENSRNOG00000012991 | ADGRA2 | -1.50 | 3.24E-18 | -1.18 | 1.30E-18 | DE in both female and male placentas |
| ENSRNOG00000050425 | RGD1561777 | -1.50 | 2.16E-03 | -1.57 | 1.66E-05 | DE in both female and male placentas |
| ENSRNOG00000037712 | MFSD6L | -1.50 | 4.18E-04 | -1.43 | 1.88E-05 | DE in both female and male placentas |
| ENSRNOG00000057713 | CAV2 | -1.50 | 1.79E-06 | -2.31 | 2.16E-21 | DE in both female and male placentas |
| ENSRNOG00000042634 | SERPINA12 | -1.50 | 5.97E-03 | -2.46 | 7.10E-10 | DE in both female and male placentas |

| | | | | | | | |
|---|---|---|---|---|---|---|---|
| ENSRNOG00000002917 | | SEC14L5 | -1.50 | 4.65E-04 | -2.82 | 1.66E-14 | DE in both female and male placentas |
| ENSRNOG00000024501 | | RGS3 | -1.50 | 1.34E-19 | -1.31 | 3.69E-24 | DE in both female and male placentas |
| ENSRNOG00000019943 | | SLC7A6 | -1.50 | 7.08E-10 | -2.18 | 6.19E-30 | DE in both female and male placentas |
| ENSRNOG00000001378 | | IQCD | -1.50 | 1.11E-03 | -2.12 | 7.48E-10 | DE in both female and male placentas |
| ENSRNOG00000059539 | | LOC1036935 64 | -1.50 | 5.18E-04 | -1.48 | 6.24E-06 | DE in both female and male placentas |
| ENSRNOG00000018053 | | FECH | -1.50 | 1.04E-13 | -1.68 | 3.61E-26 | DE in both female and male placentas |
| ENSRNOG00000025581 | | VWA2 | -1.51 | 1.36E-09 | -2.26 | 4.44E-29 | DE in both female and male placentas |
| ENSRNOG00000012575 | | FAT2 | -1.51 | 9.68E-03 | -3.03 | 2.37E-12 | DE in both female and male placentas |
| ENSRNOG00000025476 | | TMEM252 | -1.51 | 2.69E-02 | -2.93 | 1.38E-09 | DE in both female and male placentas |
| ENSRNOG00000014973 | | TEKT1 | -1.51 | 1.08E-04 | -1.47 | 5.48E-07 | DE in both female and male placentas |
| ENSRNOG00000018316 | | GRAP2 | -1.51 | 7.32E-04 | -1.09 | 1.27E-03 | DE in both female and male placentas |
| ENSRNOG00000002814 | | HIGD1B | -1.51 | 4.37E-04 | -2.31 | 1.60E-12 | DE in both female and male placentas |
| ENSRNOG00000003597 | | TUBA4A | -1.52 | 2.66E-07 | -1.90 | 4.90E-17 | DE in both female and male placentas |
| ENSRNOG00000011016 | | SLC7A2 | -1.52 | 2.04E-05 | -2.34 | 1.34E-17 | DE in both female and male placentas |
| ENSRNOG00000030868 | X | AABR070373 07.1 | -1.52 | 7.74E-03 | -3.00 | 2.44E-12 | DE in both female and male placentas |
| ENSRNOG00000056836 | | CAV1 | -1.52 | 1.15E-09 | -2.23 | 8.76E-30 | DE in both female and male placentas |
| ENSRNOG00000015423 | | CCNA2 | -1.52 | 3.17E-07 | -1.85 | 2.65E-16 | DE in both female and male placentas |
| ENSRNOG00000013847 | | NOVA2 | -1.53 | 6.37E-04 | -2.34 | 3.15E-11 | DE in both female and male placentas |

| | | | | | | | |
|---|---|---|---|---|---|---|---|
| ENSRNOG00000054889 | | WDR72 | -1.53 | 9.57E-12 | -1.95 | 6.43E-29 | DE in both female and male placentas |
| ENSRNOG00000013794 | | RBP1 | -1.53 | 5.23E-05 | -2.14 | 9.27E-14 | DE in both female and male placentas |
| ENSRNOG00000025705 | X | ARMCX2 | -1.53 | 8.16E-19 | -1.53 | 2.22E-29 | DE in both female and male placentas |
| ENSRNOG00000000638 | | ZFP365 | -1.53 | 3.48E-13 | -1.47 | 4.48E-19 | DE in both female and male placentas |
| ENSRNOG00000002050 | | IGFBP7 | -1.53 | 5.37E-14 | -1.86 | 3.67E-31 | DE in both female and male placentas |
| ENSRNOG00000029399 | | BCAM | -1.53 | 2.37E-19 | -1.70 | 1.05E-36 | DE in both female and male placentas |
| ENSRNOG00000055527 | | ARAP3 | -1.53 | 6.40E-11 | -1.40 | 1.27E-14 | DE in both female and male placentas |
| ENSRNOG00000036692 | | GCGR | -1.53 | 9.93E-09 | -1.95 | 7.21E-21 | DE in both female and male placentas |
| ENSRNOG00000013092 | X | LONRF3 | -1.54 | 5.27E-05 | -2.21 | 1.93E-14 | DE in both female and male placentas |
| ENSRNOG00000015948 | | SLC1A5 | -1.54 | 1.88E-07 | -1.87 | 1.55E-16 | DE in both female and male placentas |
| ENSRNOG00000015353 | | PRSS12 | -1.54 | 6.97E-04 | -1.78 | 2.10E-07 | DE in both female and male placentas |
| ENSRNOG00000028426 | | MCF2L | -1.55 | 1.32E-12 | -1.94 | 1.20E-29 | DE in both female and male placentas |
| ENSRNOG00000022760 | | RUBCNL | -1.55 | 2.46E-07 | -1.14 | 7.49E-07 | DE in both female and male placentas |
| ENSRNOG00000023208 | | MYORG | -1.55 | 1.35E-08 | -1.50 | 8.74E-13 | DE in both female and male placentas |
| ENSRNOG00000039323 | X | DGKK | -1.55 | 1.45E-02 | -2.90 | 2.46E-10 | DE in both female and male placentas |
| ENSRNOG00000051307 | | AOC3 | -1.56 | 8.62E-06 | -2.50 | 7.54E-20 | DE in both female and male placentas |
| ENSRNOG00000014573 | | CKMT1 | -1.56 | 3.98E-05 | -2.10 | 3.15E-13 | DE in both female and male placentas |
| ENSRNOG00000002229 | | ADCY5 | -1.56 | 2.35E-12 | -1.73 | 4.01E-23 | DE in both female and male placentas |

| | | | | | | |
|---|---|---|---|---|---|---|
| ENSRNOG00000029644 | CLEC2L | -1.56 | 1.30E-02 | -1.70 | 2.33E-04 | DE in both female and male placentas |
| ENSRNOG00000053687 | ZNF660 | -1.56 | 8.82E-04 | -1.37 | 1.45E-04 | DE in both female and male placentas |
| ENSRNOG00000051619 | ASB2 | -1.56 | 9.27E-10 | -1.11 | 7.86E-09 | DE in both female and male placentas |
| ENSRNOG00000022136 | LOC685680 | -1.56 | 1.33E-05 | -1.84 | 1.30E-11 | DE in both female and male placentas |
| ENSRNOG00000013257 | HECW2 | -1.56 | 1.62E-07 | -1.86 | 4.45E-16 | DE in both female and male placentas |
| ENSRNOG00000000572 | CHST3 | -1.56 | 4.78E-06 | -1.62 | 6.12E-10 | DE in both female and male placentas |
| ENSRNOG00000013306 | PCDH20 | -1.56 | 2.29E-02 | -1.76 | 8.94E-04 | DE in both female and male placentas |
| ENSRNOG00000021796 | RGD1565166 | -1.56 | 3.14E-03 | -1.93 | 1.23E-06 | DE in both female and male placentas |
| ENSRNOG00000006119 | SLC7A15 | -1.57 | 2.63E-03 | -1.32 | 2.37E-04 | DE in both female and male placentas |
| ENSRNOG00000017448 | EXOC3L2 | -1.57 | 3.90E-02 | -2.02 | 1.40E-04 | DE in both female and male placentas |
| ENSRNOG00000004899 | KCNS3 | -1.57 | 8.96E-06 | -1.60 | 4.95E-09 | DE in both female and male placentas |
| ENSRNOG00000019310 | EID2 | -1.57 | 1.66E-02 | -1.01 | 4.36E-02 | DE in both female and male placentas |
| ENSRNOG00000025757 | MYH6 | -1.57 | 1.13E-03 | -1.97 | 9.27E-08 | DE in both female and male placentas |
| ENSRNOG00000029012 | SHISA6 | -1.57 | 1.64E-02 | -2.56 | 1.12E-07 | DE in both female and male placentas |
| ENSRNOG00000020904 | CDC42EP2 | -1.57 | 2.72E-21 | -1.47 | 2.99E-29 | DE in both female and male placentas |
| ENSRNOG00000003338 | PMP22 | -1.57 | 1.65E-21 | -1.68 | 6.62E-38 | DE in both female and male placentas |
| ENSRNOG00000022711 | SLCO4C1 | -1.58 | 7.26E-03 | -2.93 | 1.17E-11 | DE in both female and male placentas |
| ENSRNOG00000032150 | ADCY2 | -1.58 | 6.17E-15 | -1.33 | 3.08E-17 | Differentially expressed in both female and male [illegible] |

| | | | | | | | |
|---|---|---|---|---|---|---|---|
| ENSRNOG00000021400 | X | MAGEE2 | -1.58 | 3.73E-02 | -2.54 | 2.38E-05 | DE in both female and male placentas |
| ENSRNOG00000001544 | | CYYR1 | -1.58 | 1.23E-02 | -2.66 | 5.24E-09 | DE in both female and male placentas |
| ENSRNOG00000017680 | | DENND2D | -1.58 | 4.05E-05 | -2.39 | 5.54E-16 | DE in both female and male placentas |
| ENSRNOG00000024882 | | PAX1 | -1.59 | 2.20E-02 | -2.72 | 3.49E-08 | DE in both female and male placentas |
| ENSRNOG00000005955 | | NAALAD2 | -1.59 | 1.09E-04 | -1.90 | 3.95E-09 | DE in both female and male placentas |
| ENSRNOG00000004966 | | SLC38A11 | -1.59 | 5.93E-03 | -2.14 | 9.98E-07 | DE in both female and male placentas |
| ENSRNOG00000049906 | | FOXF1 | -1.60 | 8.81E-09 | -1.80 | 9.46E-17 | DE in both female and male placentas |
| ENSRNOG00000011060 | | UNC119 | -1.60 | 1.70E-06 | -2.09 | 4.13E-16 | DE in both female and male placentas |
| ENSRNOG00000002950 | | LYL1 | -1.60 | 1.21E-08 | -1.31 | 2.35E-09 | DE in both female and male placentas |
| ENSRNOG00000015614 | | PPP1R16B | -1.60 | 3.24E-06 | -1.89 | 1.09E-12 | DE in both female and male placentas |
| ENSRNOG00000038473 | | GOT1L1 | -1.60 | 1.20E-02 | -2.19 | 2.00E-06 | DE in both female and male placentas |
| ENSRNOG00000042533 | | ACCSL | -1.60 | 2.39E-03 | -2.48 | 1.62E-09 | DE in both female and male placentas |
| ENSRNOG00000032339 | | RASL12 | -1.60 | 5.57E-15 | -1.45 | 2.30E-20 | DE in both female and male placentas |
| ENSRNOG00000007591 | | SLC45A3 | -1.61 | 1.74E-09 | -1.47 | 1.18E-12 | DE in both female and male placentas |
| ENSRNOG00000040109 | | TREML1 | -1.61 | 3.91E-07 | -1.97 | 3.55E-15 | DE in both female and male placentas |
| ENSRNOG00000047741 | | E2F2 | -1.61 | 5.33E-12 | -1.55 | 2.16E-17 | DE in both female and male placentas |
| ENSRNOG00000002525 | | PTGS2 | -1.61 | 1.87E-04 | -2.73 | 1.38E-16 | DE in both female and male placentas |
| ENSRNOG00000020246 | | MYL9 | -1.61 | 7.06E-15 | -1.89 | 4.64E-31 | DE in both female and male placentas |

| | | | | | | |
|---|---|---|---|---|---|---|
| ENSRNOG00000010280 | PDE8B | -1.62 | 8.54E-04 | -2.78 | 3.28E-13 | DE in both female and male placentas |
| ENSRNOG00000000442 | NOTCH4 | -1.62 | 7.18E-10 | -1.60 | 4.21E-15 | DE in both female and male placentas |
| ENSRNOG00000046171 | AABR07035839.1 | -1.62 | 6.67E-06 | -2.47 | 3.58E-18 | DE in both female and male placentas |
| ENSRNOG00000010263 | CLDN11 | -1.62 | 1.86E-02 | -2.20 | 4.01E-06 | DE in both female and male placentas |
| ENSRNOG00000037762 | SERPINA1F | -1.63 | 1.21E-02 | -1.18 | 1.77E-02 | DE in both female and male placentas |
| ENSRNOG00000001261 | FTCD | -1.63 | 1.27E-02 | -2.69 | 6.31E-07 | DE in both female and male placentas |
| ENSRNOG00000033466 | APON | -1.64 | 3.28E-02 | -2.30 | 1.84E-05 | DE in both female and male placentas |
| ENSRNOG00000022239 | TRARG1 | -1.64 | 3.51E-05 | -2.14 | 1.33E-12 | DE in both female and male placentas |
| ENSRNOG00000013380 | RHOV | -1.64 | 2.91E-11 | -1.74 | 6.24E-21 | DE in both female and male placentas |
| ENSRNOG00000017484 | GJA5 | -1.64 | 1.80E-07 | -1.95 | 8.46E-16 | DE in both female and male placentas |
| ENSRNOG00000002372 | SGCD | -1.65 | 4.07E-04 | -2.21 | 2.53E-09 | DE in both female and male placentas |
| ENSRNOG00000018131 | SLC16A4 | -1.65 | 9.13E-05 | -2.54 | 2.44E-15 | DE in both female and male placentas |
| ENSRNOG00000029401 | ACTG2 | -1.65 | 8.99E-10 | -1.68 | 6.02E-16 | DE in both female and male placentas |
| ENSRNOG00000018606 | OLR59 | -1.65 | 5.77E-05 | -2.95 | 9.81E-16 | DE in both female and male placentas |
| ENSRNOG00000022946 | SLC22A3 | -1.65 | 1.13E-03 | -2.68 | 1.63E-12 | DE in both female and male placentas |
| ENSRNOG00000033026 | DCLK3 | -1.65 | 5.45E-06 | -1.82 | 1.07E-10 | DE in both female and male placentas |
| ENSRNOG00000005480 | YBX3 | -1.66 | 4.07E-13 | -1.73 | 2.19E-22 | DE in both female and male placentas |
| ENSRNOG00000007827 | COX4I2 | -1.66 | 1.86E-11 | -1.89 | 1.71E-22 | DE in both female and male placentas |

| | | | | | | | |
|---|---|---|---|---|---|---|---|
| ENSRNOG00000054986 | | AABR07018050.2 | -1.66 | 2.03E-02 | -3.01 | 4.14E-09 | DE in both female and male placentas |
| ENSRNOG00000043219 | | FBN2 | -1.66 | 7.79E-12 | -1.80 | 1.60E-21 | DE in both female and male placentas |
| ENSRNOG00000027767 | X | SLC38A5 | -1.66 | 5.47E-03 | -2.11 | 2.53E-06 | DE in both female and male placentas |
| ENSRNOG00000045797 | | LEP | -1.66 | 4.92E-02 | -3.26 | 7.10E-08 | DE in both female and male placentas |
| ENSRNOG00000017282 | | TESK2 | -1.66 | 1.24E-05 | -2.28 | 5.15E-15 | DE in both female and male placentas |
| ENSRNOG00000037707 | X | ARMCX6 | -1.66 | 2.71E-11 | -1.89 | 6.95E-22 | DE in both female and male placentas |
| ENSRNOG00000012049 | | SOX7 | -1.66 | 2.05E-13 | -1.50 | 1.15E-17 | DE in both female and male placentas |
| ENSRNOG00000057153 | | PLA1A | -1.67 | 8.99E-10 | -1.62 | 1.18E-14 | DE in both female and male placentas |
| ENSRNOG00000003929 | X | PCDH19 | -1.67 | 4.92E-06 | -2.18 | 8.42E-15 | DE in both female and male placentas |
| ENSRNOG00000029071 | | UNC5C | -1.67 | 4.39E-04 | -2.09 | 5.22E-09 | DE in both female and male placentas |
| ENSRNOG00000014847 | | RASSF10 | -1.67 | 2.60E-02 | -1.31 | 2.43E-02 | DE in both female and male placentas |
| ENSRNOG00000019921 | | RHBDL1 | -1.67 | 2.34E-04 | -1.06 | 3.57E-03 | DE in both female and male placentas |
| ENSRNOG00000021474 | | SIGLEC5 | -1.68 | 8.44E-05 | -2.30 | 2.14E-12 | DE in both female and male placentas |
| ENSRNOG00000045952 | | EPHB4 | -1.68 | 9.87E-13 | -1.82 | 2.52E-23 | DE in both female and male placentas |
| ENSRNOG00000008170 | | JPH2 | -1.68 | 6.66E-05 | -1.11 | 5.54E-04 | DE in both female and male placentas |
| ENSRNOG00000059445 | | AIFM2 | -1.68 | 5.68E-09 | -2.15 | 9.11E-22 | DE in both female and male placentas |
| ENSRNOG00000047977 | | TCIM | -1.68 | 3.89E-07 | -1.86 | 2.16E-13 | DE in both female and male placentas |
| ENSRNOG00000019217 | | ADRB2 | -1.68 | 3.02E-05 | -2.60 | 4.79E-16 | DE in both female and male placentas |

| | | | | | | | |
|---|---|---|---|---|---|---|---|
| ENSRNOG00000011330 | | CES2A | -1.69 | 7.09E-06 | -2.39 | 5.57E-14 | DE in both female and male placentas |
| ENSRNOG00000009790 | | KCNK3 | -1.69 | 6.43E-06 | -2.11 | 7.50E-14 | DE in both female and male placentas |
| ENSRNOG00000006320 | | PTGES | -1.69 | 3.37E-06 | -2.05 | 1.36E-13 | DE in both female and male placentas |
| ENSRNOG00000001296 | | P2RX7 | -1.69 | 3.95E-04 | -2.16 | 1.84E-09 | DE in both female and male placentas |
| ENSRNOG00000005575 | | ABI3 | -1.70 | 9.81E-18 | -1.52 | 1.05E-22 | DE in both female and male placentas |
| ENSRNOG00000008416 | | GIMAP5 | -1.70 | 5.30E-07 | -1.82 | 3.74E-12 | DE in both female and male placentas |
| ENSRNOG00000054136 | X | XKRX | -1.70 | 1.12E-03 | -2.77 | 1.68E-12 | DE in both female and male placentas |
| ENSRNOG00000004089 | | ENPP2 | -1.70 | 2.40E-02 | -2.91 | 5.12E-08 | DE in both female and male placentas |
| ENSRNOG00000004928 | | SNTG2 | -1.70 | 1.73E-03 | -2.39 | 1.25E-07 | DE in both female and male placentas |
| ENSRNOG00000003537 | | SPTA1 | -1.70 | 1.83E-06 | -2.14 | 5.72E-15 | DE in both female and male placentas |
| ENSRNOG00000017628 | | TAGLN | -1.70 | 2.25E-23 | -1.62 | 1.43E-33 | DE in both female and male placentas |
| ENSRNOG00000009260 | | CNR2 | -1.71 | 4.59E-10 | -1.58 | 1.38E-13 | DE in both female and male placentas |
| ENSRNOG00000030034 | | SOX11 | -1.71 | 1.17E-05 | -1.83 | 1.08E-09 | DE in both female and male placentas |
| ENSRNOG00000006104 | | TG | -1.71 | 1.93E-11 | -1.84 | 3.98E-20 | DE in both female and male placentas |
| ENSRNOG00000024838 | | NA | -1.71 | 2.96E-02 | -2.82 | 6.21E-06 | DE in both female and male placentas |
| ENSRNOG00000053136 | | AABR07026557.1 | -1.72 | 2.11E-03 | -1.35 | 2.39E-03 | DE in both female and male placentas |
| ENSRNOG00000024221 | | METTL24 | -1.72 | 5.77E-03 | -2.46 | 1.88E-07 | DE in both female and male placentas |
| ENSRNOG00000028273 | | TMEM150B | -1.72 | 8.95E-05 | -1.10 | 1.24E-03 | DE in both female and male placentas |

| | | | | | | |
|---|---|---|---|---|---|---|
| ENSRNOG00000053675 | DHH | -1.72 | 2.32E-02 | -1.54 | 6.53E-03 | DE in both female and male placentas |
| ENSRNOG00000015615 | TNFRSF11A | -1.72 | 2.12E-10 | -1.53 | 2.10E-13 | DE in both female and male placentas |
| ENSRNOG00000020685 | ATP12A | -1.72 | 2.02E-02 | -2.09 | 6.08E-05 | Differentially expressed in both female and male |
| ENSRNOG00000002434 | TMEM100 | -1.72 | 8.23E-03 | -2.53 | 8.58E-08 | DE in both female and male placentas |
| ENSRNOG00000017197 | PDGFB | -1.72 | 2.81E-13 | -1.83 | 4.02E-23 | DE in both female and male placentas |
| ENSRNOG00000009892 | ADAMTS15 | -1.73 | 9.15E-15 | -1.85 | 3.29E-26 | DE in both female and male placentas |
| ENSRNOG00000001739 | MELTF | -1.73 | 5.01E-03 | -2.53 | 2.03E-08 | DE in both female and male placentas |
| ENSRNOG00000056941 | AABR07066529.3 | -1.73 | 1.61E-03 | -2.73 | 1.32E-10 | DE in both female and male placentas |
| ENSRNOG00000033444 | F10 | -1.74 | 3.69E-02 | -1.58 | 9.06E-03 | DE in both female and male placentas |
| ENSRNOG00000004566 | ARHGEF15 | -1.74 | 2.76E-07 | -2.23 | 1.69E-17 | DE in both female and male placentas |
| ENSRNOG00000056153 | TENT5B | -1.74 | 3.41E-09 | -1.72 | 1.40E-14 | DE in both female and male placentas |
| ENSRNOG00000006588 | MEOX2 | -1.74 | 1.14E-02 | -2.71 | 5.56E-08 | DE in both female and male placentas |
| ENSRNOG00000013439 | CPN1 | -1.74 | 4.55E-03 | -1.97 | 1.29E-05 | DE in both female and male placentas |
| ENSRNOG00000004346 | NOTCH3 | -1.74 | 5.85E-08 | -1.81 | 1.99E-13 | DE in both female and male placentas |
| ENSRNOG00000011346 | EHD2 | -1.74 | 4.91E-16 | -1.62 | 3.18E-22 | DE in both female and male placentas |
| ENSRNOG00000013932 | RBP2 | -1.74 | 4.66E-02 | -1.95 | 1.20E-03 | DE in both female and male placentas |
| ENSRNOG00000033261 | FAM107A | -1.74 | 6.44E-05 | -2.48 | 9.60E-14 | DE in both female and male placentas |
| ENSRNOG00000012876 | SLC6A13 | -1.74 | 4.81E-04 | -2.31 | 6.86E-10 | DE in both female and male placentas |

| | | | | | | | |
|---|---|---|---|---|---|---|---|
| ENSRNOG00000036837 | | NFE2 | -1.75 | 2.69E-09 | -1.78 | 7.02E-15 | DE in both female and male placentas |
| ENSRNOG00000057040 | | SPNS2 | -1.75 | 3.06E-05 | -2.39 | 7.55E-14 | DE in both female and male placentas |
| ENSRNOG00000059006 | X | AABR07042557.1 | -1.75 | 2.83E-04 | -2.05 | 1.75E-08 | DE in both female and male placentas |
| ENSRNOG00000001174 | | PDE9A | -1.76 | 2.40E-05 | -2.54 | 1.81E-15 | DE in both female and male placentas |
| ENSRNOG00000008656 | | SNCA | -1.76 | 2.18E-05 | -2.31 | 3.96E-13 | DE in both female and male placentas |
| ENSRNOG00000031665 | X | ACE2 | -1.76 | 1.45E-07 | -2.38 | 4.99E-20 | DE in both female and male placentas |
| ENSRNOG00000008369 | | GIMAP4 | -1.76 | 7.34E-06 | -2.62 | 5.74E-17 | DE in both female and male placentas |
| ENSRNOG00000011471 | | FGF9 | -1.76 | 1.00E-03 | -2.63 | 6.29E-11 | DE in both female and male placentas |
| ENSRNOG00000002878 | | AFM | -1.78 | 3.58E-05 | -1.11 | 9.67E-04 | DE in both female and male placentas |
| ENSRNOG00000048011 | X | AABR07040695.1 | -1.78 | 4.16E-04 | -2.50 | 1.07E-10 | DE in both female and male placentas |
| ENSRNOG00000057710 | | CCND2 | -1.78 | 1.48E-09 | -2.04 | 5.19E-19 | DE in both female and male placentas |
| ENSRNOG00000000439 | | AGER | -1.79 | 2.59E-07 | -2.25 | 5.18E-17 | DE in both female and male placentas |
| ENSRNOG00000007830 | | APOLD1 | -1.79 | 1.02E-06 | -1.86 | 2.82E-11 | DE in both female and male placentas |
| ENSRNOG00000026124 | | ID3 | -1.79 | 2.91E-16 | -1.33 | 5.92E-15 | DE in both female and male placentas |
| ENSRNOG00000052541 | | LOC103690023 | -1.79 | 1.08E-04 | -2.66 | 6.79E-14 | DE in both female and male placentas |
| ENSRNOG00000057061 | | HOXA13 | -1.80 | 7.72E-07 | -1.86 | 1.04E-10 | DE in both female and male placentas |
| ENSRNOG00000016659 | | HABP2 | -1.80 | 1.20E-02 | -1.66 | 3.30E-03 | DE in both female and male placentas |
| ENSRNOG00000008133 | | MFNG | -1.80 | 2.53E-10 | -1.90 | 8.85E-18 | DE in both female and male placentas |

| | | | | | | |
|---|---|---|---|---|---|---|
| ENSRNOG00000058077 | AABR07001068.1 | -1.81 | 4.52E-03 | -1.80 | 4.45E-04 | DE in both female and male placentas |
| ENSRNOG00000024428 | KIF20A | -1.81 | 2.11E-11 | -2.40 | 1.20E-29 | DE in both female and male placentas |
| ENSRNOG00000000017 | STEAP1 | -1.82 | 3.33E-07 | -2.19 | 1.26E-15 | DE in both female and male placentas |
| ENSRNOG00000018874 | PHF19 | -1.82 | 3.72E-12 | -2.22 | 4.99E-27 | DE in both female and male placentas |
| ENSRNOG00000005054 | PROX2 | -1.83 | 4.28E-07 | -1.01 | 4.15E-04 | DE in both female and male placentas |
| ENSRNOG00000027319 | EPHX3 | -1.83 | 6.92E-04 | -1.85 | 3.70E-06 | DE in both female and male placentas |
| ENSRNOG00000019745 | ACTN3 | -1.83 | 2.91E-06 | -1.81 | 1.84E-09 | DE in both female and male placentas |
| ENSRNOG00000018570 | C1QTNF3 | -1.83 | 7.85E-05 | -1.63 | 4.62E-06 | DE in both female and male placentas |
| ENSRNOG00000003544 | TBX4 | -1.83 | 1.29E-14 | -1.76 | 1.44E-21 | DE in both female and male placentas |
| ENSRNOG00000052190 | AABR07072602.1 | -1.83 | 4.40E-08 | -1.56 | 1.12E-09 | DE in both female and male placentas |
| ENSRNOG00000010213 | FGD5 | -1.84 | 2.71E-11 | -2.19 | 3.40E-24 | DE in both female and male placentas |
| ENSRNOG00000010253 | CD163 | -1.84 | 8.57E-05 | -2.92 | 6.62E-16 | DE in both female and male placentas |
| ENSRNOG00000037203 | TGM7 | -1.84 | 4.73E-02 | -1.76 | 8.68E-03 | DE in both female and male placentas |
| ENSRNOG00000022697 | CLEC14A | -1.84 | 2.76E-03 | -2.78 | 9.95E-10 | DE in both female and male placentas |
| ENSRNOG00000020415 | RAMP2 | -1.84 | 3.16E-08 | -2.18 | 2.68E-17 | DE in both female and male placentas |
| ENSRNOG00000047786 | OGN | -1.84 | 1.02E-02 | -2.15 | 5.00E-05 | DE in both female and male placentas |
| ENSRNOG00000051977 | MMRN2 | -1.84 | 1.85E-09 | -2.03 | 1.05E-17 | DE in both female and male placentas |
| ENSRNOG00000020169 | GIMAP8 | -1.85 | 1.96E-05 | -2.19 | 3.24E-11 | DE in both female and male placentas |

| | | | | | | | |
|---|---|---|---|---|---|---|---|
| ENSRNOG00000002997 | | SLC9A3R2 | -1.85 | 1.49E-20 | -1.59 | 1.57E-24 | DE in both female and male placentas |
| ENSRNOG00000033940 | | ADGRL4 | -1.85 | 2.96E-06 | -2.63 | 8.71E-18 | DE in both female and male placentas |
| ENSRNOG00000053656 | | LOC690020 | -1.86 | 8.13E-03 | -4.10 | 2.04E-13 | DE in both female and male placentas |
| ENSRNOG00000012344 | | SLC7A9 | -1.86 | 1.39E-02 | -2.41 | 9.82E-06 | DE in both female and male placentas |
| ENSRNOG00000001066 | | ITIH2 | -1.86 | 1.91E-02 | -2.11 | 2.95E-04 | DE in both female and male placentas |
| ENSRNOG00000037718 | | CFAP52 | -1.87 | 1.25E-03 | -2.95 | 1.18E-11 | DE in both female and male placentas |
| ENSRNOG00000007932 | | GCM1 | -1.87 | 1.96E-04 | -2.65 | 2.78E-12 | DE in both female and male placentas |
| ENSRNOG00000022296 | | PNPLA5 | -1.87 | 4.28E-02 | -1.68 | 1.13E-02 | DE in both female and male placentas |
| ENSRNOG00000038258 | | UPB1 | -1.89 | 1.02E-02 | -1.74 | 1.30E-03 | DE in both female and male placentas |
| ENSRNOG00000032332 | | KRT4 | -1.89 | 4.98E-03 | -1.97 | 6.27E-05 | DE in both female and male placentas |
| ENSRNOG00000047940 | | MMP28 | -1.90 | 1.43E-04 | -2.40 | 2.47E-10 | DE in both female and male placentas |
| ENSRNOG00000061101 | X | RIPPLY1 | -1.91 | 9.90E-04 | -2.83 | 8.01E-11 | DE in both female and male placentas |
| ENSRNOG00000028114 | | HBQ1B | -1.91 | 4.92E-04 | -1.97 | 8.85E-06 | DE in both female and male placentas |
| ENSRNOG00000053942 | | PRAMEF20 | -1.92 | 3.97E-03 | -2.22 | 5.35E-06 | DE in both female and male placentas |
| ENSRNOG00000025895 | | CAVIN2 | -1.92 | 2.41E-03 | -2.89 | 7.58E-10 | DE in both female and male placentas |
| ENSRNOG00000001399 | | TBX5 | -1.92 | 1.12E-08 | -1.69 | 2.56E-11 | DE in both female and male placentas |
| ENSRNOG00000027831 | X | MAP7D3 | -1.93 | 2.34E-04 | -2.39 | 2.22E-07 | DE in both female and male placentas |
| ENSRNOG00000039668 | | COL8A1 | -1.93 | 1.88E-08 | -2.67 | 1.08E-22 | DE in both female and male placentas |

| | | | | | | | |
|---|---|---|---|---|---|---|---|
| ENSRNOG00000011313 | | SORCS1 | -1.94 | 3.95E-04 | -2.37 | 4.91E-09 | DE in both female and male placentas |
| ENSRNOG00000006674 | | RFLNB | -1.95 | 2.05E-13 | -1.99 | 3.92E-22 | DE in both female and male placentas |
| ENSRNOG00000009173 | | SMAD6 | -1.95 | 1.14E-10 | -1.97 | 4.44E-17 | DE in both female and male placentas |
| ENSRNOG00000008587 | | TEK | -1.96 | 4.54E-04 | -2.69 | 1.48E-10 | DE in both female and male placentas |
| ENSRNOG00000018952 | | SEMA3G | -1.96 | 1.69E-05 | -2.38 | 6.77E-12 | DE in both female and male placentas |
| ENSRNOG00000060338 | | LIPC | -1.97 | 8.53E-05 | -2.40 | 2.73E-10 | Differentially expressed in both female and male |
| ENSRNOG00000004317 | | VIPR2 | -1.97 | 3.36E-14 | -2.53 | 3.67E-31 | DE in both female and male placentas |
| ENSRNOG00000048688 | | NEWGENE_ 1307313 | -1.98 | 1.70E-02 | -2.18 | 2.75E-04 | DE in both female and male placentas |
| ENSRNOG00000006802 | | LRRN1 | -1.99 | 2.24E-03 | -3.15 | 2.01E-10 | DE in both female and male placentas |
| ENSRNOG00000054695 | | CALCRL | -1.99 | 3.65E-09 | -2.53 | 7.69E-22 | DE in both female and male placentas |
| ENSRNOG00000019305 | | SH3TC2 | -1.99 | 3.50E-05 | -2.89 | 4.50E-15 | DE in both female and male placentas |
| ENSRNOG00000031912 | | LOC367195 | -1.99 | 1.24E-02 | -2.60 | 1.24E-05 | DE in both female and male placentas |
| ENSRNOG00000048699 | | VPS37D | -1.99 | 7.67E-09 | -1.74 | 9.14E-11 | DE in both female and male placentas |
| ENSRNOG00000057880 | | MYH11 | -2.00 | 6.37E-07 | -2.63 | 2.11E-17 | DE in both female and male placentas |
| ENSRNOG00000012324 | | SOGA3 | -2.02 | 1.38E-05 | -1.63 | 1.26E-05 | DE in both female and male placentas |
| ENSRNOG00000024505 | | PPP1R35 | -2.02 | 5.70E-04 | -1.63 | 2.12E-04 | DE in both female and male placentas |
| ENSRNOG00000017775 | | SLC5A1 | -2.02 | 1.71E-02 | -3.38 | 5.54E-08 | DE in both female and male placentas |
| ENSRNOG00000000167 | X | ALAS2 | -2.02 | 4.12E-07 | -2.60 | 3.04E-17 | DE in both female and male placentas |

| | | | | | | |
|---|---|---|---|---|---|---|
| ENSRNOG00000005874 | TLE2 | -2.02 | 4.74E-08 | -2.04 | 1.13E-13 | DE in both female and male placentas |
| ENSRNOG00000009436 | HEMGN | -2.02 | 3.18E-06 | -2.90 | 1.41E-17 | DE in both female and male placentas |
| ENSRNOG00000027942 | EPS8L1 | -2.03 | 1.14E-06 | -2.29 | 6.06E-13 | DE in both female and male placentas |
| ENSRNOG00000018086 | SLC22A8 | -2.03 | 2.37E-07 | -2.23 | 1.48E-13 | DE in both female and male placentas |
| ENSRNOG00000017676 | PLVAP | -2.04 | 1.21E-03 | -2.51 | 7.00E-08 | DE in both female and male placentas |
| ENSRNOG00000017606 | P2RX1 | -2.04 | 2.40E-10 | -2.29 | 7.14E-20 | DE in both female and male placentas |
| ENSRNOG00000045989 | HBA-A3 | -2.04 | 1.66E-08 | -2.19 | 2.82E-15 | DE in both female and male placentas |
| ENSRNOG00000061100 | TMEM150A | -2.04 | 6.95E-09 | -2.44 | 4.34E-19 | DE in both female and male placentas |
| ENSRNOG00000060629 | LOC103694855 | -2.05 | 4.40E-08 | -2.90 | 4.84E-23 | DE in both female and male placentas |
| ENSRNOG00000061544 | SPOCK2 | -2.05 | 1.52E-05 | -2.79 | 3.28E-14 | DE in both female and male placentas |
| ENSRNOG00000015403 | CD52 | -2.05 | 1.47E-07 | -2.72 | 1.91E-18 | DE in both female and male placentas |
| ENSRNOG00000051548 | LMOD1 | -2.06 | 6.80E-12 | -2.43 | 5.72E-25 | DE in both female and male placentas |
| ENSRNOG00000019112 | CHP2 | -2.06 | 3.58E-09 | -1.55 | 1.59E-08 | DE in both female and male placentas |
| ENSRNOG00000031851 | NDUFA4L2 | -2.07 | 5.65E-10 | -2.41 | 3.06E-20 | DE in both female and male placentas |
| ENSRNOG00000009354 | NRARP | -2.07 | 3.17E-06 | -2.48 | 3.67E-13 | DE in both female and male placentas |
| ENSRNOG00000052061 | PPEF2 | -2.07 | 3.91E-02 | -1.63 | 4.35E-02 | DE in both female and male placentas |
| ENSRNOG00000024426 | AABR07032520.1 | -2.07 | 1.75E-04 | -2.03 | 1.18E-06 | DE in both female and male placentas |
| ENSRNOG00000016275 | TTR | -2.08 | 1.69E-02 | -2.25 | 3.29E-04 | DE in both female and male placentas |

| | | | | | | | |
|---|---|---|---|---|---|---|---|
| ENSRNOG00000043419 | | HIST3H2BA | -2.08 | 9.22E-08 | -1.80 | 3.90E-09 | DE in both female and male placentas |
| ENSRNOG00000011184 | | SLC13A4 | -2.09 | 6.04E-04 | -2.98 | 7.29E-11 | DE in both female and male placentas |
| ENSRNOG00000023643 | | MMP17 | -2.10 | 9.99E-09 | -2.27 | 7.44E-16 | DE in both female and male placentas |
| ENSRNOG00000008529 | | FOXS1 | -2.10 | 1.06E-11 | -1.28 | 1.95E-08 | DE in both female and male placentas |
| ENSRNOG00000020879 | | NAGS | -2.10 | 1.18E-08 | -1.22 | 1.95E-05 | DE in both female and male placentas |
| ENSRNOG00000060320 | | LIN28A | -2.11 | 3.86E-05 | -2.98 | 5.59E-14 | DE in both female and male placentas |
| ENSRNOG00000046484 | X | CCDC160 | -2.12 | 3.11E-09 | -3.22 | 1.27E-26 | DE in both female and male placentas |
| ENSRNOG00000021248 | | CDC25B | -2.12 | 3.75E-15 | -2.59 | 6.94E-34 | DE in both female and male placentas |
| ENSRNOG00000018241 | | ANK1 | -2.13 | 9.13E-09 | -2.96 | 1.44E-23 | DE in both female and male placentas |
| ENSRNOG00000058775 | | LY49S7 | -2.13 | 1.27E-05 | -1.82 | 7.72E-07 | DE in both female and male placentas |
| ENSRNOG00000046949 | | KCNB1 | -2.14 | 1.36E-06 | -2.70 | 3.32E-15 | DE in both female and male placentas |
| ENSRNOG00000005269 | | SRL | -2.14 | 2.05E-03 | -2.81 | 7.29E-08 | DE in both female and male placentas |
| ENSRNOG00000009662 | | RSPO4 | -2.16 | 3.54E-03 | -2.03 | 4.09E-04 | DE in both female and male placentas |
| ENSRNOG00000011649 | | EPB42 | -2.16 | 3.96E-07 | -3.13 | 1.94E-20 | DE in both female and male placentas |
| ENSRNOG00000048976 | | LOC501467 | -2.16 | 3.16E-02 | -1.84 | 1.98E-02 | DE in both female and male placentas |
| ENSRNOG00000003884 | | ACMSD | -2.16 | 2.46E-06 | -2.89 | 7.51E-14 | DE in both female and male placentas |
| ENSRNOG00000042229 | | NA | -2.17 | 2.44E-03 | -1.63 | 4.23E-03 | DE in both female and male placentas |
| ENSRNOG00000030814 | | KB23 | -2.18 | 2.36E-02 | -2.00 | 1.01E-02 | DE in both female and male placentas |

| | | | | | | | |
|---|---|---|---|---|---|---|---|
| ENSRNOG00000010451 | | SLC34A3 | -2.20 | 4.23E-02 | -2.33 | 2.24E-03 | DE in both female and male placentas |
| ENSRNOG00000045614 | | AABR070494 55.1 | -2.20 | 3.83E-02 | -2.57 | 1.16E-03 | DE in both female and male placentas |
| ENSRNOG00000004794 | | RTN1 | -2.21 | 2.62E-10 | -2.67 | 2.08E-22 | DE in both female and male placentas |
| ENSRNOG00000020165 | | AHSP | -2.22 | 1.07E-12 | -3.01 | 6.94E-34 | DE in both female and male placentas |
| ENSRNOG00000052810 | | CYP2C11 | -2.22 | 2.40E-02 | -3.43 | 6.10E-04 | DE in both female and male placentas |
| ENSRNOG00000008988 | | CYP2D2 | -2.22 | 2.28E-02 | -1.62 | 3.17E-02 | DE in both female and male placentas |
| ENSRNOG00000026787 | | DISP2 | -2.22 | 5.46E-05 | -3.27 | 1.46E-14 | DE in both female and male placentas |
| ENSRNOG00000052368 | | CLCNKA | -2.22 | 2.35E-03 | -1.57 | 8.76E-03 | DE in both female and male placentas |
| ENSRNOG00000003746 | X | GJB1 | -2.22 | 2.33E-02 | -2.61 | 2.22E-04 | DE in both female and male placentas |
| ENSRNOG00000061904 | | LOC1036948 64 | -2.23 | 1.72E-07 | -2.40 | 2.81E-13 | DE in both female and male placentas |
| ENSRNOG00000009694 | | BMP4 | -2.24 | 1.70E-07 | -2.95 | 8.60E-19 | DE in both female and male placentas |
| ENSRNOG00000062297 | | KLRA5 | -2.25 | 8.66E-07 | -1.99 | 1.01E-08 | DE in both female and male placentas |
| ENSRNOG00000018853 | | SLC28A3 | -2.25 | 5.95E-06 | -3.01 | 5.91E-15 | DE in both female and male placentas |
| ENSRNOG00000016429 | | GRM5 | -2.25 | 5.70E-03 | -1.61 | 5.27E-03 | DE in both female and male placentas |
| ENSRNOG00000050121 | | RHAG | -2.26 | 4.09E-05 | -2.65 | 9.80E-10 | DE in both female and male placentas |
| ENSRNOG00000028207 | | COLGALT2 | -2.26 | 1.04E-16 | -1.92 | 1.45E-19 | DE in both female and male placentas |
| ENSRNOG00000046751 | | LOC688649 | -2.27 | 1.91E-02 | -2.99 | 1.61E-05 | DE in both female and male placentas |
| ENSRNOG00000003870 | | C1QTNF2 | -2.28 | 1.49E-24 | -2.12 | 5.42E-35 | DE in both female and male placentas |

| | | | | | | |
|---|---|---|---|---|---|---|
| ENSRNOG00000042820 | PRAMEF17 | -2.30 | 5.65E-05 | -3.07 | 2.34E-12 | DE in both female and male placentas |
| ENSRNOG00000020546 | LIPE | -2.30 | 3.06E-17 | -2.24 | 1.89E-26 | DE in both female and male placentas |
| ENSRNOG00000001711 | PLAAT1 | -2.31 | 4.69E-14 | -1.88 | 5.40E-16 | DE in both female and male placentas |
| ENSRNOG00000020420 | PKLR | -2.32 | 3.68E-07 | -2.72 | 1.24E-14 | DE in both female and male placentas |
| ENSRNOG00000006911 | SPTB | -2.32 | 5.24E-17 | -2.64 | 8.61E-34 | DE in both female and male placentas |
| ENSRNOG00000002728 | BTC | -2.33 | 7.72E-07 | -2.64 | 2.38E-13 | DE in both female and male placentas |
| ENSRNOG00000000335 | ERMAP | -2.37 | 5.70E-07 | -2.10 | 2.47E-08 | DE in both female and male placentas |
| ENSRNOG00000058105 | HBB | -2.38 | 5.78E-09 | -2.88 | 1.31E-19 | DE in both female and male placentas |
| ENSRNOG00000020951 | SLC4A1 | -2.38 | 1.37E-08 | -3.24 | 1.10E-22 | Differentially expressed in both female and male |
| ENSRNOG00000010158 | MAGEL2 | -2.38 | 3.15E-18 | -2.78 | 1.08E-37 | DE in both female and male placentas |
| ENSRNOG00000051160 | AABR070440 17.1 | -2.38 | 1.28E-05 | -2.14 | 3.16E-07 | DE in both female and male placentas |
| ENSRNOG00000020877 | PYY | -2.39 | 6.82E-03 | -1.62 | 1.76E-02 | DE in both female and male placentas |
| ENSRNOG00000009191 | CCM2L | -2.39 | 8.71E-05 | -2.23 | 7.10E-07 | DE in both female and male placentas |
| ENSRNOG00000002548 | TNN | -2.40 | 6.19E-04 | -2.50 | 1.56E-06 | DE in both female and male placentas |
| ENSRNOG00000061527 | GCK | -2.40 | 1.27E-02 | -3.37 | 4.25E-06 | DE in both female and male placentas |
| ENSRNOG00000061299 | LOC1001348 71 | -2.42 | 6.69E-10 | -2.93 | 8.63E-22 | DE in both female and male placentas |
| ENSRNOG00000026679 | SCN4B | -2.43 | 4.32E-05 | -4.12 | 2.53E-13 | DE in both female and male placentas |
| ENSRNOG00000017002 | ADRB1 | -2.43 | 1.00E-06 | -2.15 | 9.40E-09 | DE in both female and male placentas |

| | | | | | | | |
|---|---|---|---|---|---|---|---|
| ENSRNOG00000003500 | | APOA2 | -2.44 | 7.47E-03 | -2.75 | 3.13E-05 | DE in both female and male placentas |
| ENSRNOG00000015691 | | INKA2 | -2.44 | 5.53E-07 | -2.83 | 4.26E-14 | DE in both female and male placentas |
| ENSRNOG00000018369 | | PRX | -2.44 | 7.62E-22 | -2.71 | 1.10E-41 | DE in both female and male placentas |
| ENSRNOG00000012238 | | CLEC3A | -2.47 | 2.56E-05 | -3.26 | 2.38E-13 | Differentially expressed in both female and male |
| ENSRNOG00000036680 | | NOTUM | -2.48 | 5.73E-05 | -2.15 | 2.40E-06 | DE in both female and male placentas |
| ENSRNOG00000017130 | | RHD | -2.49 | 3.06E-07 | -3.40 | 8.39E-19 | DE in both female and male placentas |
| ENSRNOG00000020406 | | TMED6 | -2.50 | 1.97E-18 | -1.51 | 1.42E-12 | DE in both female and male placentas |
| ENSRNOG00000005857 | | LRRC3B | -2.50 | 2.43E-02 | -4.05 | 2.66E-06 | DE in both female and male placentas |
| ENSRNOG00000047491 | X | AABR07037520.1 | -2.52 | 1.08E-07 | -3.09 | 3.70E-17 | DE in both female and male placentas |
| ENSRNOG00000004713 | | KCNJ16 | -2.56 | 9.86E-03 | -5.31 | 1.71E-10 | DE in both female and male placentas |
| ENSRNOG00000015682 | | KEL | -2.57 | 8.69E-09 | -3.48 | 3.77E-22 | DE in both female and male placentas |
| ENSRNOG00000047098 | | HBB-BS | -2.58 | 3.70E-09 | -3.23 | 4.38E-21 | DE in both female and male placentas |
| ENSRNOG00000010698 | | CAR1 | -2.59 | 6.08E-05 | -3.98 | 1.81E-14 | DE in both female and male placentas |
| ENSRNOG00000029886 | | HBA-A1 | -2.59 | 1.18E-10 | -2.89 | 1.88E-20 | DE in both female and male placentas |
| ENSRNOG00000048275 | | RGD1309350 | -2.68 | 2.08E-06 | -3.53 | 1.64E-15 | DE in both female and male placentas |
| ENSRNOG00000050006 | X | AGTR2 | -2.68 | 8.72E-03 | -1.96 | 5.18E-03 | DE in both female and male placentas |
| ENSRNOG00000016070 | | TRPC3 | -2.69 | 2.73E-08 | -3.02 | 4.26E-16 | Differentially expressed in both female and male |
| ENSRNOG00000020405 | | INS2 | -2.69 | 4.34E-03 | -1.79 | 1.19E-02 | DE in both female and male placentas |

| | | | | | | | |
|---|---|---|---|---|---|---|---|
| ENSRNOG00000052549 | | CHGA | -2.70 | 2.27E-07 | -3.49 | 1.10E-17 | DE in both female and male placentas |
| ENSRNOG00000009656 | | RSPO1 | -2.71 | 6.75E-07 | -2.80 | 3.85E-11 | DE in both female and male placentas |
| ENSRNOG00000026501 | | SLC6A19 | -2.74 | 4.49E-04 | -2.75 | 2.55E-06 | DE in both female and male placentas |
| ENSRNOG00000013162 | X | ZCCHC12 | -2.74 | 4.69E-07 | -3.19 | 3.34E-14 | DE in both female and male placentas |
| ENSRNOG00000025987 | | LOC498236 | -2.74 | 3.90E-03 | -3.28 | 1.86E-06 | DE in both female and male placentas |
| ENSRNOG00000010761 | | SFTPB | -2.75 | 2.93E-07 | -3.15 | 1.03E-15 | DE in both female and male placentas |
| ENSRNOG00000024330 | | NGP | -2.75 | 7.37E-06 | -3.15 | 1.41E-11 | DE in both female and male placentas |
| ENSRNOG00000047321 | | HBA-A2 | -2.77 | 2.94E-10 | -3.15 | 2.73E-20 | DE in both female and male placentas |
| ENSRNOG00000009563 | | KRT2 | -2.81 | 1.86E-03 | -2.54 | 1.94E-05 | DE in both female and male placentas |
| ENSRNOG00000057995 | | LOC1025560 51 | -2.84 | 4.24E-02 | -3.15 | 8.42E-04 | DE in both female and male placentas |
| ENSRNOG00000062101 | | ACE | -2.85 | 1.30E-09 | -3.54 | 5.58E-22 | DE in both female and male placentas |
| ENSRNOG00000003443 | | KLF1 | -2.90 | 2.20E-16 | -2.40 | 1.12E-18 | DE in both female and male placentas |
| ENSRNOG00000054582 | | LOC1036948 57 | -2.91 | 5.27E-05 | -3.74 | 1.07E-11 | DE in both female and male placentas |
| ENSRNOG00000012189 | | LECT2 | -2.92 | 2.18E-02 | -2.44 | 1.52E-03 | Differentially expressed in both female and male |
| ENSRNOG00000030879 | | HBG1 | -2.99 | 1.08E-04 | -4.42 | 6.70E-13 | DE in both female and male placentas |
| ENSRNOG00000005770 | | SOSTDC1 | -3.02 | 1.33E-02 | -2.58 | 1.74E-03 | DE in both female and male placentas |
| ENSRNOG00000055454 | X | ARL13A | -3.05 | 9.95E-07 | -3.61 | 4.10E-14 | DE in both female and male placentas |
| ENSRNOG00000058883 | | NEWGENE_ 1589866 | -3.05 | 1.20E-02 | -2.80 | 1.43E-03 | DE in both female and male placentas |

| | | | | | | | |
|---|---|---|---|---|---|---|---|
| ENSRNOG00000021130 | | ABCC8 | -3.11 | 5.77E-04 | -3.55 | 1.54E-07 | DE in both female and male placentas |
| ENSRNOG00000005781 | | WNT16 | -3.13 | 8.31E-05 | -1.70 | 1.07E-02 | DE in both female and male placentas |
| ENSRNOG00000028085 | X | TMEM255A | -3.19 | 3.34E-12 | -4.12 | 1.44E-28 | DE in both female and male placentas |
| ENSRNOG00000052271 | X | AABR07041794.1 | -3.33 | 3.56E-03 | -3.88 | 5.68E-05 | DE in both female and male placentas |
| ENSRNOG00000055855 | | AABR07027254.1 | -3.56 | 1.57E-05 | -1.97 | 3.58E-04 | DE in both female and male placentas |
| ENSRNOG00000016295 | | LYZL1 | -3.64 | 6.28E-03 | -2.74 | 5.42E-04 | DE in both female and male placentas |
| ENSRNOG00000023917 | | AABR07034362.1 | -3.69 | 2.00E-02 | -2.48 | 1.34E-02 | DE in both female and male placentas |
| ENSRNOG00000000012 | | TCF15 | -3.92 | 2.54E-12 | -3.83 | 5.84E-17 | DE in both female and male placentas |
| ENSRNOG00000046734 | | EVX1 | -4.02 | 7.71E-14 | -4.98 | 8.34E-25 | DE in both female and male placentas |
| ENSRNOG00000018148 | | LOC108348081 | -6.59 | 9.99E-03 | -5.91 | 5.06E-04 | DE in both female and male placentas |
| ENSRNOG00000062289 | | AABR07055846.2 | 5.19 | 5.27E-03 | | | DE only in female placentas |
| ENSRNOG00000030596 | | LOC100362149 | 5.02 | 6.49E-04 | | | DE only in female placentas |
| ENSRNOG00000030345 | | LOC100362684 | 5.02 | 6.49E-04 | | | DE only in female placentas |
| ENSRNOG00000009768 | | LOC100912228 | 4.84 | 9.88E-03 | | | DE only in female placentas |
| ENSRNOG00000055570 | | AABR07034940.2 | 4.58 | 4.72E-03 | | | DE only in female placentas |
| ENSRNOG00000049740 | | AABR07034273.1 | 4.33 | 2.25E-02 | | | DE only in female placentas |
| ENSRNOG00000060541 | | AABR07017825.5 | 4.18 | 1.03E-02 | | | DE only in female placentas |
| ENSRNOG00000050713 | | LOC108348050 | 3.96 | 3.07E-02 | | | DE only in female placentas |
| ENSRNOG00000025088 | | TMC3 | 3.94 | 2.29E-04 | | | DE only in female placentas |

| | | | | |
|---|---|---|---|---|
| ENSRNOG00000004875 | CCER1 | 3.74 | 2.67E-02 | DE only in female placentas |
| ENSRNOG00000030150 | LOC1003621 09 | 3.68 | 8.49E-03 | DE only in female placentas |
| ENSRNOG00000057504 | AABR070723 74.2 | 3.66 | 4.32E-02 | DE only in female placentas |
| ENSRNOG00000047336 | AABR070672 46.1 | 3.65 | 6.24E-03 | DE only in female placentas |
| ENSRNOG00000045802 | OLR1374 | 3.62 | 7.66E-03 | DE only in female placentas |
| ENSRNOG00000030840 | CADM2 | 3.55 | 1.89E-03 | DE only in female placentas |
| ENSRNOG00000058599 | AC119462.3 | 3.50 | 9.13E-03 | DE only in female placentas |
| ENSRNOG00000042697 | AABR070732 16.1 | 3.38 | 2.58E-02 | DE only in female placentas |
| ENSRNOG00000057392 | AABR070428 59.1 | 3.33 | 1.89E-02 | DE only in female placentas |
| ENSRNOG00000024288 | ETV2 | 3.31 | 2.13E-02 | DE only in female placentas |
| ENSRNOG00000009209 | SLITRK1 | 3.31 | 2.96E-02 | DE only in female placentas |
| ENSRNOG00000047646 | AABR070086 70.1 | 3.27 | 3.35E-03 | DE only in female placentas |
| ENSRNOG00000059904 | AABR070701 61.4 | 3.27 | 3.36E-02 | DE only in female placentas |
| ENSRNOG00000062263 | AABR070315 21.2 | 3.20 | 2.46E-02 | DE only in female placentas |
| ENSRNOG00000032508 | ACOT5 | 3.20 | 1.30E-02 | DE only in female placentas |
| ENSRNOG00000057463 | AABR070082 93.4 | 3.17 | 4.78E-02 | DE only in female placentas |
| ENSRNOG00000057214 | AABR070435 10.1 | 3.10 | 4.92E-02 | DE only in female placentas |
| ENSRNOG00000059074 | LOC691661 | 3.06 | 3.75E-02 | DE only in female placentas |
| ENSRNOG00000057765 | AABR070344 57.1 | 3.05 | 3.85E-02 | DE only in female placentas |
| ENSRNOG00000060606 | AC098180.1 | 3.04 | 4.46E-02 | DE only in female placentas |
| ENSRNOG00000060517 | AABR070690 08.2 | 2.97 | 1.63E-04 | DE only in female placentas |
| ENSRNOG00000051410 | AABR070433 47.1 | 2.90 | 1.28E-02 | DE only in female placentas |
| ENSRNOG00000029173 | AABR070537 16.1 | 2.89 | 2.02E-02 | DE only in female placentas |

| | | | | | |
|---|---|---|---|---|---|
| ENSRNOG00000049239 | | AABR07017006.1 | 2.88 | 4.68E-02 | DE only in female placentas |
| ENSRNOG00000022863 | | LINGO4 | 2.84 | 3.85E-02 | DE only in female placentas |
| ENSRNOG00000054065 | | AABR07058976.1 | 2.83 | 3.14E-02 | DE only in female placentas |
| ENSRNOG00000021266 | | PROKR2 | 2.80 | 3.16E-02 | DE only in female placentas |
| ENSRNOG00000047100 | X | LOC367975 | 2.75 | 1.51E-02 | DE only in female placentas |
| ENSRNOG00000005609 | | NEUROD1 | 2.61 | 3.78E-02 | DE only in female placentas |
| ENSRNOG00000057757 | | AABR07058501.1 | 2.60 | 2.90E-02 | DE only in female placentas |
| ENSRNOG00000026989 | | GAPT | 2.60 | 4.54E-02 | DE only in female placentas |
| ENSRNOG00000061785 | | AABR07065124.2 | 2.56 | 2.53E-02 | DE only in female placentas |
| ENSRNOG00000046790 | X | AABR07040887.1 | 2.52 | 4.81E-02 | DE only in female placentas |
| ENSRNOG00000048545 | | LOC100910483 | 2.52 | 1.16E-02 | DE only in female placentas |
| ENSRNOG00000009771 | | CDH10 | 2.48 | 4.55E-02 | DE only in female placentas |
| ENSRNOG00000056817 | | MUC6 | 2.46 | 4.50E-02 | DE only in female placentas |
| ENSRNOG00000016136 | | PDCD1LG2 | 2.44 | 2.35E-02 | DE only in female placentas |
| ENSRNOG00000055069 | X | 7SK | 2.41 | 7.72E-03 | DE only in female placentas |
| ENSRNOG00000062160 | | AC108616.2 | 2.40 | 1.40E-03 | DE only in female placentas |
| ENSRNOG00000030253 | | TAFA3 | 2.38 | 3.89E-02 | DE only in female placentas |
| ENSRNOG00000011269 | | SULT1C3 | 2.36 | 4.55E-02 | DE only in female placentas |
| ENSRNOG00000039762 | X | GPR82 | 2.36 | 4.65E-02 | DE only in female placentas |
| ENSRNOG00000033540 | | CTS7 | 2.34 | 7.17E-03 | DE only in female placentas |
| ENSRNOG00000052547 | | LOC103694082 | 2.28 | 1.97E-02 | DE only in female placentas |
| ENSRNOG00000060077 | | AABR07065485.1 | 2.25 | 7.76E-03 | DE only in female placentas |
| ENSRNOG00000055401 | | KCNC1 | 2.23 | 2.08E-04 | DE only in female placentas |
| ENSRNOG00000048686 | | AABR07066379.1 | 2.19 | 1.24E-03 | DE only in female placentas |
| ENSRNOG00000037368 | | AABR07049862.1 | 2.16 | 2.11E-02 | DE only in female placentas |

| | | | | | |
|---|---|---|---|---|---|
| ENSRNOG00000055486 | | AABR07025316.1 | 2.12 | 1.88E-02 | DE only in female placentas |
| ENSRNOG00000054763 | | LOC108348322 | 2.08 | 4.49E-02 | DE only in female placentas |
| ENSRNOG00000030481 | | LOC100362110 | 2.08 | 4.48E-02 | DE only in female placentas |
| ENSRNOG00000052774 | | AC123253.2 | 2.08 | 4.84E-03 | DE only in female placentas |
| ENSRNOG00000018832 | | RGD1564854 | 2.06 | 8.94E-03 | DE only in female placentas |
| ENSRNOG00000051951 | | AABR07034592.1 | 2.03 | 1.93E-04 | DE only in female placentas |
| ENSRNOG00000031445 | | LOC100361866 | 2.03 | 1.06E-02 | DE only in female placentas |
| ENSRNOG00000053229 | | AABR07007905.2 | 2.00 | 2.04E-04 | DE only in female placentas |
| ENSRNOG00000047765 | | AABR07010682.1 | 1.94 | 4.25E-04 | DE only in female placentas |
| ENSRNOG00000060196 | | AABR07035926.2 | 1.93 | 2.06E-02 | DE only in female placentas |
| ENSRNOG00000047712 | X | DCX | 1.92 | 3.35E-02 | DE only in female placentas |
| ENSRNOG00000042741 | | ADGB | 1.90 | 2.30E-06 | DE only in female placentas |
| ENSRNOG00000052461 | | AABR07072759.2 | 1.90 | 3.19E-02 | DE only in female placentas |
| ENSRNOG00000057101 | | AABR07050652.1 | 1.88 | 1.32E-02 | DE only in female placentas |
| ENSRNOG00000060803 | | AABR07028984.1 | 1.86 | 3.95E-03 | DE only in female placentas |
| ENSRNOG00000007061 | | ARHGEF33 | 1.86 | 3.02E-02 | DE only in female placentas |
| ENSRNOG00000009299 | | ADRA2C | 1.85 | 2.72E-02 | DE only in female placentas |
| ENSRNOG00000029450 | | AABR07058422.1 | 1.85 | 1.41E-02 | DE only in female placentas |
| ENSRNOG00000057738 | | AC104053.1 | 1.85 | 5.50E-04 | DE only in female placentas |
| ENSRNOG00000061578 | | AABR07057333.1 | 1.82 | 3.16E-03 | DE only in female placentas |
| ENSRNOG00000048904 | | AABR07004232.1 | 1.82 | 3.55E-02 | DE only in female placentas |

| | | | | |
|---|---|---|---|---|
| ENSRNOG00000053367 | AABR07064257.1 | 1.80 | 3.04E-04 | DE only in female placentas |
| ENSRNOG00000007808 | NAP1L5 | 1.80 | 1.00E-03 | DE only in female placentas |
| ENSRNOG00000059013 | AABR07060519.1 | 1.77 | 2.30E-02 | DE only in female placentas |
| ENSRNOG00000054029 | AABR07033540.1 | 1.76 | 1.46E-02 | DE only in female placentas |
| ENSRNOG00000032828 | MAPK8IP2 | 1.75 | 3.70E-02 | DE only in female placentas |
| ENSRNOG00000055867 | AC105577.1 | 1.74 | 4.46E-02 | DE only in female placentas |
| ENSRNOG00000056748 | AC229945.1 | 1.74 | 3.13E-03 | DE only in female placentas |
| ENSRNOG00000033162 | CHIA | 1.74 | 3.25E-03 | DE only in female placentas |
| ENSRNOG00000008299 | DCDC2C | 1.68 | 2.23E-03 | DE only in female placentas |
| ENSRNOG00000029649 | PRSS45 | 1.68 | 1.50E-02 | DE only in female placentas |
| ENSRNOG00000034030 | AC130162.1 | 1.66 | 3.80E-03 | DE only in female placentas |
| ENSRNOG00000002468 | TNR | 1.66 | 6.45E-04 | DE only in female placentas |
| ENSRNOG00000016656 | NKX2-3 | 1.65 | 4.30E-02 | DE only in female placentas |
| ENSRNOG00000056358 | CCDC27 | 1.65 | 4.29E-02 | DE only in female placentas |
| ENSRNOG00000058819 | AABR07056767.1 | 1.63 | 1.69E-02 | DE only in female placentas |
| ENSRNOG00000056490 | AABR07013566.1 | 1.63 | 1.22E-03 | DE only in female placentas |
| ENSRNOG00000050416 | KCNA10 | 1.62 | 2.24E-03 | DE only in female placentas |
| ENSRNOG00000014143 | COL24A1 | 1.61 | 1.58E-04 | DE only in female placentas |
| ENSRNOG00000052486 | KCNA6 | 1.60 | 1.48E-02 | DE only in female placentas |
| ENSRNOG00000054338 | AC094221.4 | 1.59 | 2.39E-02 | DE only in female placentas |
| ENSRNOG00000060618 | AABR07027267.1 | 1.58 | 4.26E-02 | DE only in female placentas |
| ENSRNOG00000047117 | AABR07001037.1 | 1.57 | 1.54E-02 | DE only in female placentas |
| ENSRNOG00000000855 | LST1 | 1.56 | 1.06E-04 | DE only in female placentas |
| ENSRNOG00000057501 | FAM81A | 1.56 | 1.13E-02 | DE only in female placentas |
| ENSRNOG00000060718 | AABR07064253.1 | 1.56 | 4.17E-04 | DE only in female placentas |
| ENSRNOG00000057396 | AABR07005581.1 | 1.55 | 2.39E-02 | DE only in female placentas |

| | | | | | |
|---|---|---|---|---|---|
| ENSRNOG00000049961 | X | AABR07037923.1 | 1.55 | 1.98E-02 | DE only in female placentas |
| ENSRNOG00000061389 | | AABR07058233.2 | 1.55 | 8.14E-03 | DE only in female placentas |
| ENSRNOG00000032803 | X | RPL21 | 1.54 | 3.26E-05 | DE only in female placentas |
| ENSRNOG00000049314 | | AABR07069524.1 | 1.53 | 3.06E-02 | DE only in female placentas |
| ENSRNOG00000007544 | | IL23R | 1.52 | 1.94E-04 | DE only in female placentas |
| ENSRNOG00000051664 | | AABR07010878.1 | 1.52 | 8.53E-03 | DE only in female placentas |
| ENSRNOG00000058408 | | AC108588.1 | 1.51 | 1.33E-03 | DE only in female placentas |
| ENSRNOG00000004795 | X | POF1B | 1.50 | 2.16E-02 | DE only in female placentas |
| ENSRNOG00000008683 | | ALK | 1.48 | 3.06E-02 | DE only in female placentas |
| ENSRNOG00000054519 | | AABR07053179.1 | 1.48 | 5.48E-03 | DE only in female placentas |
| ENSRNOG00000025198 | | GAS2L3 | 1.47 | 5.58E-11 | DE only in female placentas |
| ENSRNOG00000061280 | | AABR07068018.2 | 1.46 | 3.78E-02 | DE only in female placentas |
| ENSRNOG00000049178 | | AABR07028414.1 | 1.45 | 4.26E-02 | DE only in female placentas |
| ENSRNOG00000045839 | | AABR07011857.1 | 1.45 | 4.30E-02 | DE only in female placentas |
| ENSRNOG00000012176 | | RAB27B | 1.44 | 1.01E-06 | DE only in female placentas |
| ENSRNOG00000049504 | | AABR07009221.1 | 1.44 | 5.25E-03 | DE only in female placentas |
| ENSRNOG00000028082 | | TAL2 | 1.43 | 2.80E-02 | DE only in female placentas |
| ENSRNOG00000008027 | | CAVIN4 | 1.43 | 1.69E-03 | DE only in female placentas |
| ENSRNOG00000061488 | | AABR07026557.2 | 1.41 | 9.45E-03 | DE only in female placentas |
| ENSRNOG00000030269 | | ATP2B2 | 1.40 | 3.29E-04 | DE only in female placentas |
| ENSRNOG00000019589 | | TAS1R3 | 1.39 | 1.34E-02 | DE only in female placentas |
| ENSRNOG00000018286 | | CHRNA1 | 1.39 | 1.09E-02 | DE only in female placentas |
| ENSRNOG00000059554 | | AABR07058907.2 | 1.38 | 3.17E-02 | DE only in female placentas |
| ENSRNOG00000002014 | | STAP1 | 1.37 | 1.49E-02 | DE only in female placentas |

| | | | | | |
|---|---|---|---|---|---|
| ENSRNOG00000058425 | | AABR07000385.2 | 1.37 | 4.20E-03 | DE only in female placentas |
| ENSRNOG00000017788 | | KATNAL2 | 1.37 | 2.39E-02 | DE only in female placentas |
| ENSRNOG00000015168 | | LOC365238 | 1.36 | 2.07E-02 | DE only in female placentas |
| ENSRNOG00000059974 | | AABR07053283.1 | 1.36 | 3.30E-03 | DE only in female placentas |
| ENSRNOG00000061317 | | AABR07011733.1 | 1.35 | 4.87E-02 | DE only in female placentas |
| ENSRNOG00000057709 | | SNORD8 | 1.35 | 3.89E-02 | DE only in female placentas |
| ENSRNOG00000016041 | | ZC3H12D | 1.35 | 8.90E-03 | DE only in female placentas |
| ENSRNOG00000056331 | | AC127640.1 | 1.34 | 7.57E-04 | DE only in female placentas |
| ENSRNOG00000046027 | | AABR07034730.1 | 1.33 | 4.28E-02 | DE only in female placentas |
| ENSRNOG00000016510 | | PRL7D1 | 1.32 | 3.22E-08 | DE only in female placentas |
| ENSRNOG00000060733 | | AABR07004101.2 | 1.32 | 4.18E-02 | DE only in female placentas |
| ENSRNOG00000054768 | | AABR07050487.1 | 1.31 | 9.74E-04 | DE only in female placentas |
| ENSRNOG00000042714 | | RGD1559459 | 1.31 | 1.29E-04 | DE only in female placentas |
| ENSRNOG00000057039 | | AC124874.1 | 1.30 | 1.17E-02 | DE only in female placentas |
| ENSRNOG00000051812 | | AC129370.1 | 1.30 | 3.60E-02 | DE only in female placentas |
| ENSRNOG00000050523 | | AABR07049579.2 | 1.30 | 4.87E-02 | DE only in female placentas |
| ENSRNOG00000061404 | | AABR07050484.1 | 1.29 | 5.68E-06 | DE only in female placentas |
| ENSRNOG00000000451 | | RT1-BA | 1.29 | 1.67E-02 | DE only in female placentas |
| ENSRNOG00000034287 | | AABR07035916.1 | 1.29 | 8.72E-03 | DE only in female placentas |
| ENSRNOG00000054196 | | AABR07048228.1 | 1.29 | 6.11E-04 | DE only in female placentas |
| ENSRNOG00000059056 | | AABR07048637.1 | 1.28 | 9.60E-03 | DE only in female placentas |
| ENSRNOG00000055364 | X | AC141028.4 | 1.27 | 2.66E-02 | DE only in female placentas |

| | | | | | |
|---|---|---|---|---|---|
| ENSRNOG00000057318 | | AABR07073295.1 | 1.26 | 7.66E-03 | DE only in female placentas |
| ENSRNOG00000059468 | | AABR07034573.3 | 1.26 | 4.60E-02 | DE only in female placentas |
| ENSRNOG00000038422 | | LOC691352 | 1.25 | 1.49E-04 | DE only in female placentas |
| ENSRNOG00000059087 | | LOC102546889 | 1.25 | 5.43E-03 | DE only in female placentas |
| ENSRNOG00000027847 | | NXPE2 | 1.24 | 4.10E-02 | DE only in female placentas |
| ENSRNOG00000051456 | | AABR07013729.1 | 1.24 | 2.97E-03 | DE only in female placentas |
| ENSRNOG00000038471 | | LOC500148 | 1.24 | 4.97E-02 | DE only in female placentas |
| ENSRNOG00000053770 | | NA | 1.23 | 3.53E-02 | DE only in female placentas |
| ENSRNOG00000052694 | X | AABR07037878.2 | 1.23 | 3.86E-02 | DE only in female placentas |
| ENSRNOG00000006804 | | PPP1R42 | 1.23 | 2.54E-03 | DE only in female placentas |
| ENSRNOG00000061900 | | AABR07047771.1 | 1.23 | 3.19E-02 | DE only in female placentas |
| ENSRNOG00000051954 | | AC242863.1 | 1.22 | 4.35E-02 | DE only in female placentas |
| ENSRNOG00000059786 | | AC107531.3 | 1.22 | 5.87E-03 | DE only in female placentas |
| ENSRNOG00000057296 | | AC112001.1 | 1.21 | 1.93E-02 | DE only in female placentas |
| ENSRNOG00000022601 | | LOC100360821 | 1.21 | 9.11E-03 | DE only in female placentas |
| ENSRNOG00000052405 | | CDH4 | 1.21 | 2.87E-02 | DE only in female placentas |
| ENSRNOG00000061053 | | AABR07007997.1 | 1.21 | 4.77E-02 | DE only in female placentas |
| ENSRNOG00000036672 | | SECTM1A | 1.20 | 4.84E-04 | DE only in female placentas |
| ENSRNOG00000003748 | X | RGD1565785 | 1.20 | 1.37E-03 | DE only in female placentas |
| ENSRNOG00000023720 | | NTM | 1.19 | 9.93E-04 | DE only in female placentas |
| ENSRNOG00000025601 | | TOPAZ1 | 1.19 | 4.72E-02 | DE only in female placentas |
| ENSRNOG00000061869 | | AABR07035012.1 | 1.19 | 2.75E-02 | DE only in female placentas |
| ENSRNOG00000061273 | | AABR07021972.2 | 1.18 | 5.96E-03 | DE only in female placentas |
| ENSRNOG00000011028 | | AABR07054096.1 | 1.17 | 9.56E-03 | DE only in female placentas |

| | | | | | |
|---|---|---|---|---|---|
| ENSRNOG00000026891 | | AC093995.1 | 1.17 | 8.58E-03 | DE only in female placentas |
| ENSRNOG00000008560 | | CCDC30 | 1.16 | 2.78E-09 | DE only in female placentas |
| ENSRNOG00000010345 | | SSMEM1 | 1.15 | 4.21E-03 | DE only in female placentas |
| ENSRNOG00000013670 | | PAX3 | 1.15 | 5.13E-03 | DE only in female placentas |
| ENSRNOG00000055384 | | PRL3D4 | 1.14 | 1.26E-09 | DE only in female placentas |
| ENSRNOG00000032989 | | LRRN4 | 1.13 | 3.20E-02 | DE only in female placentas |
| ENSRNOG00000052935 | | AABR07057382.1 | 1.12 | 9.05E-04 | DE only in female placentas |
| ENSRNOG00000059506 | X | AC124926.2 | 1.12 | 4.73E-02 | DE only in female placentas |
| ENSRNOG00000038101 | | LOC102557319 | 1.12 | 2.61E-10 | DE only in female placentas |
| ENSRNOG00000052315 | | AC244225.1 | 1.11 | 1.39E-03 | DE only in female placentas |
| ENSRNOG00000050137 | | AJAP1 | 1.11 | 4.79E-02 | DE only in female placentas |
| ENSRNOG00000030276 | | CTSM | 1.11 | 1.43E-06 | DE only in female placentas |
| ENSRNOG00000039725 | | CES1A | 1.11 | 4.14E-08 | DE only in female placentas |
| ENSRNOG00000056947 | | AABR07025272.1 | 1.10 | 2.53E-04 | DE only in female placentas |
| ENSRNOG00000040026 | X | RHOX10 | 1.10 | 4.03E-02 | DE only in female placentas |
| ENSRNOG00000051964 | | LOC108348190 | 1.09 | 3.33E-02 | DE only in female placentas |
| ENSRNOG00000003251 | | B3GALT2 | 1.09 | 1.13E-04 | DE only in female placentas |
| ENSRNOG00000023612 | | DOXL1 | 1.09 | 1.33E-04 | DE only in female placentas |
| ENSRNOG00000061539 | | LOC102550577 | 1.09 | 1.79E-02 | DE only in female placentas |
| ENSRNOG00000007765 | | FRZB | 1.08 | 1.08E-12 | DE only in female placentas |
| ENSRNOG00000040300 | | RAET1E | 1.07 | 8.77E-03 | DE only in female placentas |

| | | | | | |
|---|---|---|---|---|---|
| ENSRNOG00000047077 | | LOC681341 | 1.07 | 4.45E-02 | DE only in female placentas |
| ENSRNOG00000019770 | | BNC1 | 1.07 | 3.07E-06 | DE only in female placentas |
| ENSRNOG00000008678 | | ANTXR1 | 1.07 | 3.16E-05 | DE only in female placentas |
| ENSRNOG00000027607 | | CEACAM3 | 1.05 | 6.14E-07 | DE only in female placentas |
| ENSRNOG00000048235 | X | AABR07039778.1 | 1.05 | 5.17E-03 | DE only in female placentas |
| ENSRNOG00000053576 | | AC126530.2 | 1.04 | 9.73E-03 | DE only in female placentas |
| ENSRNOG00000010283 | | CD28 | 1.04 | 1.30E-03 | DE only in female placentas |
| ENSRNOG00000019447 | | ECEL1 | 1.04 | 4.66E-02 | DE only in female placentas |
| ENSRNOG00000002381 | | BMP3 | 1.03 | 1.96E-02 | DE only in female placentas |
| ENSRNOG00000031536 | | SYNDIG1 | 1.03 | 2.28E-02 | DE only in female placentas |
| ENSRNOG00000061152 | | AABR07018038.3 | 1.02 | 1.04E-03 | DE only in female placentas |
| ENSRNOG00000012535 | | ADGRF4 | 1.02 | 3.89E-02 | DE only in female placentas |
| ENSRNOG00000000605 | | HS3ST5 | 1.02 | 6.58E-04 | DE only in female placentas |
| ENSRNOG00000051095 | | AABR07021528.1 | 1.02 | 3.54E-03 | DE only in female placentas |
| ENSRNOG00000052652 | | LOC100910500 | 1.01 | 3.59E-03 | DE only in female placentas |
| ENSRNOG00000050953 | | OLR1280 | 1.01 | 3.59E-03 | DE only in female placentas |
| ENSRNOG00000046648 | | LOC100910640 | 1.01 | 3.59E-03 | DE only in female placentas |
| ENSRNOG00000016463 | | PRL2A1 | 1.01 | 1.74E-16 | DE only in female placentas |

| | | | | | |
|---|---|---|---|---|---|
| ENSRNOG00000031275 | | AABR07054361.1 | 1.00 | 3.40E-02 | DE only in female placentas |
| ENSRNOG00000031202 | | TRIM30C | 1.00 | 2.43E-02 | DE only in female placentas |
| ENSRNOG00000014863 | | RECK | -1.00 | 3.37E-11 | DE only in female placentas |
| ENSRNOG00000048722 | | LOC103690365 | -1.00 | 4.67E-02 | DE only in female placentas |
| ENSRNOG00000046844 | | LOC108348150 | -1.00 | 4.67E-02 | DE only in female placentas |
| ENSRNOG00000007408 | | EBF4 | -1.00 | 3.92E-02 | DE only in female placentas |
| ENSRNOG00000062053 | | AABR07030527.2 | -1.00 | 2.56E-02 | DE only in female placentas |
| ENSRNOG00000046434 | | HIST1H3B | -1.00 | 4.68E-02 | DE only in female placentas |
| ENSRNOG00000006877 | X | EFNB1 | -1.00 | 5.50E-11 | DE only in female placentas |
| ENSRNOG00000020955 | | MYL3 | -1.00 | 2.56E-03 | DE only in female placentas |
| ENSRNOG00000004206 | | GLRX5 | -1.00 | 9.66E-12 | DE only in female placentas |
| ENSRNOG00000010077 | | SMARCD3 | -1.00 | 2.00E-04 | DE only in female placentas |
| ENSRNOG00000000956 | | RASL11A | -1.00 | 4.20E-05 | DE only in female placentas |
| ENSRNOG00000049605 | | ZFP296 | -1.00 | 8.34E-03 | DE only in female placentas |
| ENSRNOG00000022417 | X | PRICKLE3 | -1.01 | 8.94E-08 | DE only in female placentas |
| ENSRNOG00000020981 | | PRSS46 | -1.01 | 2.16E-02 | DE only in female placentas |
| ENSRNOG00000000041 | | SLC26A1 | -1.01 | 3.58E-03 | DE only in female placentas |
| ENSRNOG00000016828 | | CMTM5 | -1.01 | 5.63E-04 | DE only in female placentas |

| | | | | |
|---|---|---|---|---|
| ENSRNOG00000002916 | CA4 | -1.01 | 3.53E-04 | DE only in female placentas |
| ENSRNOG00000004483 | PTPRR | -1.02 | 1.76E-04 | DE only in female placentas |
| ENSRNOG00000000597 | CCN6 | -1.02 | 7.49E-03 | DE only in female placentas |
| ENSRNOG00000021170 | PLEKHO1 | -1.02 | 1.95E-10 | DE only in female placentas |
| ENSRNOG00000010457 | VASH1 | -1.02 | 7.61E-06 | DE only in female placentas |
| ENSRNOG00000025843 | CCDC102A | -1.02 | 8.66E-06 | DE only in female placentas |
| ENSRNOG00000022071 | ITGA2B | -1.03 | 2.47E-06 | DE only in female placentas |
| ENSRNOG00000003722 | DUSP27 | -1.03 | 1.54E-08 | DE only in female placentas |
| ENSRNOG00000020839 | LRFN3 | -1.03 | 7.07E-03 | DE only in female placentas |
| ENSRNOG00000048838 | GJC1 | -1.03 | 1.78E-06 | DE only in female placentas |
| ENSRNOG00000004781 | CRMP1 | -1.03 | 2.15E-03 | DE only in female placentas |
| ENSRNOG00000016967 | HFE | -1.03 | 5.19E-03 | DE only in female placentas |
| ENSRNOG00000004344 | ZDHHC3 | -1.03 | 1.40E-19 | DE only in female placentas |
| ENSRNOG00000027592 | RERG | -1.04 | 5.84E-06 | DE only in female placentas |
| ENSRNOG00000032561 | SLC52A2 | -1.04 | 1.70E-07 | DE only in female placentas |
| ENSRNOG00000027990 | CRIP1 | -1.04 | 7.67E-07 | DE only in female placentas |
| ENSRNOG00000008904 | FLI1 | -1.04 | 3.74E-08 | DE only in female placentas |
| ENSRNOG00000026686 | DDX4 | -1.04 | 2.81E-02 | DE only in female placentas |

| | | | | |
|---|---|---|---|---|
| ENSRNOG00000059950 | AABR070141<br>58.1 | -1.04 | 1.18E-02 | DE only in female placentas |
| ENSRNOG00000006255 | NIPAL4 | -1.05 | 1.79E-05 | DE only in female placentas |
| ENSRNOG00000008178 | NXN | -1.05 | 5.05E-07 | DE only in female placentas |
| ENSRNOG00000045978 | HIST2H4 | -1.05 | 2.68E-02 | DE only in female placentas |
| ENSRNOG00000048642 | HIST1H4B | -1.05 | 2.68E-02 | DE only in female placentas |
| ENSRNOG00000048735 | HIST1H2AO | -1.05 | 2.68E-02 | DE only in female placentas |
| ENSRNOG00000017960 | DEAF1 | -1.05 | 3.05E-18 | DE only in female placentas |
| ENSRNOG00000010977 | IGFBP6 | -1.05 | 6.69E-03 | DE only in female placentas |
| ENSRNOG00000012714 | RGD1564149 | -1.05 | 3.87E-02 | DE only in female placentas |
| ENSRNOG00000059572 | TSEN34L1 | -1.05 | 2.59E-07 | DE only in female placentas |
| ENSRNOG00000050564 | AABR070605<br>85.1 | -1.05 | 8.29E-03 | DE only in female placentas |
| ENSRNOG00000025735 | WDR86 | -1.05 | 1.15E-03 | DE only in female placentas |
| ENSRNOG00000014347 | SLC4A2 | -1.05 | 1.83E-12 | DE only in female placentas |
| ENSRNOG00000010629 | NOD1 | -1.05 | 2.30E-11 | DE only in female placentas |
| ENSRNOG00000015822 | KLF13 | -1.05 | 4.14E-02 | DE only in female placentas |
| ENSRNOG00000008065 | WWC1 | -1.06 | 1.93E-11 | DE only in female placentas |
| ENSRNOG00000028690 | LOC1003623<br>66 | -1.06 | 1.60E-02 | DE only in female placentas |
| ENSRNOG00000010146 | NDN | -1.06 | 1.83E-12 | DE only in female placentas |

| | | | | |
|---|---|---|---|---|
| ENSRNOG00000018445 | AGT | -1.06 | 1.80E-02 | DE only in female placentas |
| ENSRNOG00000019985 | ASIC4 | -1.07 | 5.98E-05 | DE only in female placentas |
| ENSRNOG00000002773 | RGS4 | -1.07 | 2.29E-06 | DE only in female placentas |
| ENSRNOG00000027286 | LRRC75A | -1.07 | 8.13E-03 | DE only in female placentas |
| ENSRNOG00000019771 | TBX6 | -1.08 | 4.51E-02 | DE only in female placentas |
| ENSRNOG00000003494 | PPFIA4 | -1.09 | 1.51E-03 | DE only in female placentas |
| ENSRNOG00000011901 | RRAD | -1.09 | 4.81E-03 | DE only in female placentas |
| ENSRNOG00000000786 | RPP21 | -1.09 | 2.55E-12 | DE only in female placentas |
| ENSRNOG00000046313 | BASP1 | -1.09 | 1.45E-04 | DE only in female placentas |
| ENSRNOG00000016700 | TCF21 | -1.09 | 7.05E-04 | DE only in female placentas |
| ENSRNOG00000032224 | RGD1562378 | -1.10 | 1.65E-03 | DE only in female placentas |
| ENSRNOG00000022686 | ZDHHC2 | -1.10 | 7.29E-08 | DE only in female placentas |
| ENSRNOG00000017558 | TUBB2A | -1.10 | 1.90E-17 | DE only in female placentas |
| ENSRNOG00000018616 | CDC42EP5 | -1.11 | 4.12E-05 | DE only in female placentas |
| ENSRNOG00000019990 | REX1BD | -1.11 | 2.08E-02 | DE only in female placentas |
| ENSRNOG00000004461 | NEWGENE_6497122 | -1.11 | 3.24E-03 | DE only in female placentas |
| ENSRNOG00000023296 | AABR07002945.1 | -1.11 | 2.97E-09 | DE only in female placentas |
| ENSRNOG00000050220 | NRXN1 | -1.12 | 5.94E-04 | DE only in female placentas |

| | | | | |
|---|---|---|---|---|
| ENSRNOG00000019850 | SPEG | -1.12 | 5.57E-04 | DE only in female placentas |
| ENSRNOG00000014205 | KLF2 | -1.13 | 5.62E-03 | DE only in female placentas |
| ENSRNOG00000018822 | SLC5A5 | -1.13 | 2.89E-05 | DE only in female placentas |
| ENSRNOG00000011201 | CMTM8 | -1.13 | 2.00E-09 | DE only in female placentas |
| ENSRNOG00000021526 | SLC25A34 | -1.13 | 7.42E-03 | DE only in female placentas |
| ENSRNOG00000020942 | PLEKHA4 | -1.13 | 2.07E-06 | DE only in female placentas |
| ENSRNOG00000030155 | BMP8A | -1.13 | 2.65E-06 | DE only in female placentas |
| ENSRNOG00000001325 | IL3RA | -1.13 | 1.09E-04 | DE only in female placentas |
| ENSRNOG00000020906 | POLA2 | -1.13 | 1.37E-16 | DE only in female placentas |
| ENSRNOG00000026324 | STARD6 | -1.14 | 5.95E-03 | DE only in female placentas |
| ENSRNOG00000051891 | LOC100912564 | -1.15 | 7.32E-04 | DE only in female placentas |
| ENSRNOG00000052555 | HIST2H2AA2 | -1.16 | 1.51E-02 | DE only in female placentas |
| ENSRNOG00000047898 | HIST2H2AA3 | -1.16 | 1.51E-02 | DE only in female placentas |
| ENSRNOG00000006324 | TRPC6 | -1.16 | 1.83E-06 | DE only in female placentas |
| ENSRNOG00000020416 | ZFR2 | -1.16 | 5.75E-03 | DE only in female placentas |
| ENSRNOG00000050183 | RT1-CE1 | -1.17 | 4.78E-05 | DE only in female placentas |
| ENSRNOG00000010607 | RNF208 | -1.17 | 6.90E-04 | DE only in female placentas |
| ENSRNOG00000006278 | TSPYL5 | -1.17 | 3.51E-02 | DE only in female placentas |

| | | | | |
|---|---|---|---|---|
| ENSRNOG00000057690 | LOC1009125<br>64 | -1.17 | 1.89E-02 | DE only in female placentas |
| ENSRNOG00000010359 | BIK | -1.18 | 1.64E-02 | DE only in female placentas |
| ENSRNOG00000047322 | CRB3 | -1.18 | 1.30E-14 | DE only in female placentas |
| ENSRNOG00000001336 | ORAI1 | -1.18 | 9.23E-08 | DE only in female placentas |
| ENSRNOG00000001888 | ARVCF | -1.19 | 7.54E-16 | DE only in female placentas |
| ENSRNOG00000004310 | CASKIN2 | -1.20 | 1.67E-11 | DE only in female placentas |
| ENSRNOG00000015304 | TMEM160 | -1.21 | 5.01E-06 | DE only in female placentas |
| ENSRNOG00000015131 | UBE2C | -1.22 | 1.94E-05 | DE only in female placentas |
| ENSRNOG00000020455 | CST6 | -1.23 | 5.69E-14 | DE only in female placentas |
| ENSRNOG00000014232 | P2RY1 | -1.24 | 1.75E-02 | DE only in female placentas |
| ENSRNOG00000050933 | LOC1009125<br>64 | -1.24 | 4.95E-04 | DE only in female placentas |
| ENSRNOG00000051078 | ZFP37 | -1.25 | 3.72E-02 | DE only in female placentas |
| ENSRNOG00000056955 | BICDL1 | -1.25 | 6.07E-11 | DE only in female placentas |
| ENSRNOG00000005132 | SLC52A3 | -1.25 | 6.50E-14 | DE only in female placentas |
| ENSRNOG00000003781 | ATP10B | -1.26 | 5.74E-09 | DE only in female placentas |
| ENSRNOG00000001177 | ACADS | -1.26 | 5.50E-11 | DE only in female placentas |
| ENSRNOG00000008857 | ADAMTS10 | -1.28 | 6.46E-12 | DE only in female placentas |
| ENSRNOG00000005569 | PHOSPHO1 | -1.28 | 1.28E-09 | DE only in female placentas |

| | | | | | |
|---|---|---|---|---|---|
| ENSRNOG00000003737 | | BEST2 | -1.29 | 1.72E-04 | DE only in female placentas |
| ENSRNOG00000054709 | | AABR070613 82.2 | -1.29 | 3.77E-02 | DE only in female placentas |
| ENSRNOG00000046636 | | HIST1H4M | -1.30 | 9.68E-03 | DE only in female placentas |
| ENSRNOG00000029478 | | CYP4F39 | -1.30 | 2.55E-02 | DE only in female placentas |
| ENSRNOG00000018378 | | CACNB2 | -1.30 | 1.04E-03 | DE only in female placentas |
| ENSRNOG00000056457 | | GPD1 | -1.30 | 6.36E-03 | DE only in female placentas |
| ENSRNOG00000048169 | | TUBA8 | -1.31 | 7.56E-07 | DE only in female placentas |
| ENSRNOG00000055564 | | RGD1564664 | -1.32 | 2.36E-04 | DE only in female placentas |
| ENSRNOG00000047817 | | ENOX1 | -1.32 | 1.48E-03 | DE only in female placentas |
| ENSRNOG00000008544 | | B3GNT4 | -1.32 | 3.23E-02 | DE only in female placentas |
| ENSRNOG00000046001 | | AABR070308 23.1 | -1.35 | 5.31E-03 | DE only in female placentas |
| ENSRNOG00000055049 | | ALDH1A2 | -1.38 | 4.78E-02 | DE only in female placentas |
| ENSRNOG00000013578 | | TREM2 | -1.38 | 6.86E-06 | DE only in female placentas |
| ENSRNOG00000054539 | | HIST1H2AIL 1 | -1.38 | 7.89E-03 | DE only in female placentas |
| ENSRNOG00000050805 | | LOC1025504 56 | -1.39 | 5.19E-05 | DE only in female placentas |
| ENSRNOG00000058444 | | LOC1009125 64 | -1.40 | 7.63E-03 | DE only in female placentas |
| ENSRNOG00000050869 | | CEBPD | -1.41 | 4.63E-06 | DE only in female placentas |
| ENSRNOG00000037845 | X | CYSLTR1 | -1.41 | 3.86E-02 | DE only in female placentas |

| | | | | |
|---|---|---|---|---|
| ENSRNOG00000020676 | PPP1R14A | -1.42 | 2.53E-04 | DE only in female placentas |
| ENSRNOG00000021098 | RASGRP2 | -1.43 | 6.76E-12 | DE only in female placentas |
| ENSRNOG00000047864 | LOC108348072 | -1.45 | 1.77E-07 | DE only in female placentas |
| ENSRNOG00000006855 | PADI4 | -1.45 | 2.99E-07 | DE only in female placentas |
| ENSRNOG00000020400 | MEF2B | -1.45 | 9.48E-03 | DE only in female placentas |
| ENSRNOG00000050554 | LOC100362342 | -1.46 | 9.74E-08 | DE only in female placentas |
| ENSRNOG00000030927 | GRID2IP | -1.48 | 5.20E-03 | DE only in female placentas |
| ENSRNOG00000018217 | SYT5 | -1.49 | 1.55E-03 | DE only in female placentas |
| ENSRNOG00000023474 | DAZL | -1.50 | 4.53E-05 | DE only in female placentas |
| ENSRNOG00000057700 | HIST1H2AN | -1.51 | 7.45E-03 | DE only in female placentas |
| ENSRNOG00000039876 | LOC681410 | -1.53 | 1.11E-03 | DE only in female placentas |
| ENSRNOG00000020835 | HIPK4 | -1.56 | 1.62E-02 | DE only in female placentas |
| ENSRNOG00000024294 | AABR07019083.1 | -1.59 | 5.95E-03 | DE only in female placentas |
| ENSRNOG00000052594 | HIST1H2AK | -1.60 | 1.96E-02 | DE only in female placentas |
| ENSRNOG00000016734 | EMILIN3 | -1.60 | 2.16E-04 | DE only in female placentas |
| ENSRNOG00000015156 | GAL | -1.60 | 3.60E-03 | DE only in female placentas |
| ENSRNOG00000032300 | RPL39L | -1.63 | 5.32E-04 | DE only in female placentas |
| ENSRNOG00000058329 | PRRX2 | -1.65 | 3.22E-03 | DE only in female placentas |

| | | | | | |
|---|---|---|---|---|---|
| ENSRNOG00000062143 | | LOC100911830 | -1.66 | 2.24E-05 | DE only in female placentas |
| ENSRNOG00000021041 | | FAM171A2 | -1.71 | 5.34E-08 | DE only in female placentas |
| ENSRNOG00000027109 | | ODF3L1 | -1.76 | 2.48E-02 | DE only in female placentas |
| ENSRNOG00000050207 | | UTF1 | -1.81 | 1.91E-03 | DE only in female placentas |
| ENSRNOG00000021920 | | PRAMEL6 | -1.90 | 2.02E-02 | DE only in female placentas |
| ENSRNOG00000010600 | | CYSRT1 | -1.91 | 4.52E-04 | DE only in female placentas |
| ENSRNOG00000060366 | | LOC102548682 | -2.00 | 2.76E-04 | DE only in female placentas |
| ENSRNOG00000032231 | | MROH6 | -2.01 | 4.36E-05 | DE only in female placentas |
| ENSRNOG00000026039 | | TSPAN32 | -2.02 | 1.07E-03 | DE only in female placentas |
| ENSRNOG00000021447 | | PRR7 | -2.06 | 3.39E-04 | DE only in female placentas |
| ENSRNOG00000047281 | X | LOC108348065 | -2.19 | 5.80E-03 | DE only in female placentas |
| ENSRNOG00000011593 | | HEY1 | -2.21 | 1.83E-04 | DE only in female placentas |
| ENSRNOG00000001057 | | CTXN1 | -2.21 | 1.89E-05 | DE only in female placentas |
| ENSRNOG00000000590 | | NAGLT1 | -2.37 | 4.06E-02 | DE only in female placentas |
| ENSRNOG00000021079 | | FXYD1 | -2.37 | 4.36E-05 | DE only in female placentas |
| ENSRNOG00000033609 | | IRX1 | -2.40 | 5.66E-04 | DE only in female placentas |
| ENSRNOG00000021225 | | OXT | -2.52 | 5.95E-03 | DE only in female placentas |
| ENSRNOG00000021434 | X | USP51 | -2.63 | 3.84E-02 | DE only in female placentas |

| | | | | | | | |
|---|---|---|---|---|---|---|---|
| ENSRNOG00000046021 | X | LOC1083481 72 | -2.68 | 3.31E-02 | | | DE only in female placentas |
| ENSRNOG00000060864 | | AABR070593 72.1 | -2.71 | 4.23E-02 | | | DE only in female placentas |
| ENSRNOG00000007668 | | MAFA | -2.75 | 4.50E-02 | | | DE only in female placentas |
| ENSRNOG00000017330 | | AC103221.1 | -2.85 | 2.92E-02 | | | DE only in female placentas |
| ENSRNOG00000008365 | | HOXB2 | -2.92 | 2.80E-02 | | | DE only in female placentas |
| ENSRNOG00000048850 | | AABR070584 41.1 | -3.04 | 2.75E-02 | | | DE only in female placentas |
| ENSRNOG00000028179 | X | AABR070385 52.1 | -3.51 | 3.73E-02 | | | DE only in female placentas |
| ENSRNOG00000034088 | | DNAJB8 | -3.77 | 1.49E-03 | | | DE only in female placentas |
| ENSRNOG00000030925 | | AABR070279 25.2 | -4.07 | 3.23E-02 | | | DE only in female placentas |
| ENSRNOG00000028996 | | KRT1 | -4.13 | 1.71E-02 | | | DE only in female placentas |
| ENSRNOG00000047214 | | AABR070067 75.1 | -4.90 | 3.77E-02 | | | DE only in female placentas |
| ENSRNOG00000045654 | | LOC1083481 08 | -5.67 | 1.72E-02 | | | DE only in female placentas |
| ENSRNOG00000050535 | X | RRAGB | -5.78 | 2.47E-02 | | | DE only in female placentas |
| ENSRNOG00000054513 | | ZMIZ2 | -6.94 | 6.83E-03 | | | DE only in female placentas |
| ENSRNOG00000060344 | | AABR070730 45.1 | -7.16 | 5.29E-03 | | | DE only in female placentas |
| ENSRNOG00000050922 | | NUPR1L1 | | | 3.37 | 2.99E-02 | DE only in male placentas |
| ENSRNOG00000038129 | | DEFB52 | | | 4.12 | 5.74E-03 | DE only in male placentas |
| ENSRNOG00000055138 | | CLDN16 | | | 2.50 | 1.27E-02 | DE only in male placentas |

| | | | | | |
|---|---|---|---|---|---|
| ENSRNOG00000016073 | | TAAR1 | 3.06 | 4.18E-03 | DE only in male placentas |
| ENSRNOG00000002154 | | MEPE | 3.07 | 8.53E-03 | DE only in male placentas |
| ENSRNOG00000031344 | | AABR070487 70.1 | 2.97 | 3.16E-02 | DE only in male placentas |
| ENSRNOG00000029861 | | GSTA2 | 2.61 | 4.72E-03 | DE only in male placentas |
| ENSRNOG00000030212 | | CERKL | 2.87 | 1.79E-03 | DE only in male placentas |
| ENSRNOG00000000487 | | GRM4 | 2.23 | 1.24E-04 | DE only in male placentas |
| ENSRNOG00000010278 | | IL6 | 3.65 | 1.97E-08 | DE only in male placentas |
| ENSRNOG00000050243 | | AKR1C12L1 | 2.52 | 2.12E-04 | DE only in male placentas |
| ENSRNOG00000027331 | X | PPEF1 | 1.37 | 4.41E-02 | DE only in male placentas |
| ENSRNOG00000007231 | | SPAM1 | 1.57 | 3.94E-02 | DE only in male placentas |
| ENSRNOG00000036984 | | SLCO1A1 | 1.63 | 1.48E-02 | DE only in male placentas |
| ENSRNOG00000062121 | | AABR070206 63.2 | 2.02 | 4.86E-02 | DE only in male placentas |
| ENSRNOG00000018394 | | CHRDL2 | 1.42 | 3.54E-02 | DE only in male placentas |
| ENSRNOG00000043088 | | CEACAM19 | 2.89 | 3.93E-07 | DE only in male placentas |
| ENSRNOG00000028708 | | NTSR1 | 2.07 | 1.64E-04 | DE only in male placentas |
| ENSRNOG00000019110 | | OCSTAMP | 2.58 | 3.62E-04 | DE only in male placentas |
| ENSRNOG00000013196 | | DOK5 | 1.66 | 1.78E-02 | DE only in male placentas |
| ENSRNOG00000059170 | | LOC1025513 56 | 1.26 | 3.64E-02 | DE only in male placentas |

| | | | | |
|---|---|---|---|---|
| ENSRNOG00000015992 | CCL20 | 2.62 | 2.94E-04 | DE only in male placentas |
| ENSRNOG00000033099 | DCC | 2.11 | 6.82E-05 | DE only in male placentas |
| ENSRNOG00000000410 | NEPN | 1.51 | 2.78E-03 | DE only in male placentas |
| ENSRNOG00000025805 | MS4A14 | 1.85 | 1.07E-02 | DE only in male placentas |
| ENSRNOG00000061014 | AABR070668 70.1 | 2.42 | 1.25E-02 | DE only in male placentas |
| ENSRNOG00000022298 | CXCL11 | 3.10 | 2.27E-11 | DE only in male placentas |
| ENSRNOG00000014414 | QRFPR | 1.67 | 1.07E-02 | DE only in male placentas |
| ENSRNOG00000053260 | LILRB3A | 1.81 | 2.69E-05 | DE only in male placentas |
| ENSRNOG00000053124 | OLR1L | 1.92 | 1.78E-03 | DE only in male placentas |
| ENSRNOG00000026247 | AABR070183 26.1 | 1.97 | 4.28E-02 | DE only in male placentas |
| ENSRNOG00000047901 | ACP7 | 2.09 | 4.08E-06 | DE only in male placentas |
| ENSRNOG00000033983 | SOHLH1 | 2.99 | 6.90E-03 | DE only in male placentas |
| ENSRNOG00000011406 | CCL4 | 2.78 | 5.76E-11 | DE only in male placentas |
| ENSRNOG00000018958 | MT3 | 1.77 | 9.82E-04 | DE only in male placentas |
| ENSRNOG00000009425 | FGF7 | 3.52 | 4.08E-05 | DE only in male placentas |
| ENSRNOG00000013222 | CD207 | 1.97 | 1.95E-04 | DE only in male placentas |
| ENSRNOG00000004180 | TAFA2 | 1.09 | 2.54E-02 | DE only in male placentas |
| ENSRNOG00000046683 | LILRB3 | 1.88 | 5.62E-04 | DE only in male placentas |

| | | | | |
|---|---|---|---|---|
| ENSRNOG00000055716 | OSCAR | 1.87 | 3.17E-06 | DE only in male placentas |
| ENSRNOG00000029371 | FAM71A | 2.35 | 4.97E-03 | DE only in male placentas |
| ENSRNOG00000019744 | AABR070320 97.1 | 1.36 | 3.06E-04 | DE only in male placentas |
| ENSRNOG00000019318 | SYT3 | 1.40 | 7.99E-03 | DE only in male placentas |
| ENSRNOG00000025254 | QRICH2 | 1.54 | 1.12E-02 | DE only in male placentas |
| ENSRNOG00000016301 | DMRT2 | 1.22 | 1.92E-03 | DE only in male placentas |
| ENSRNOG00000026192 | AABR070363 24.1 | 3.05 | 1.29E-02 | DE only in male placentas |
| ENSRNOG00000014496 | CORO6 | 1.60 | 1.41E-17 | DE only in male placentas |
| ENSRNOG00000036880 | ARL5C | 1.28 | 5.93E-04 | DE only in male placentas |
| ENSRNOG00000009574 | CRCT1 | 3.95 | 2.81E-05 | DE only in male placentas |
| ENSRNOG00000046155 | LOC498675 | 1.37 | 2.64E-04 | DE only in male placentas |
| ENSRNOG00000028812 | TRIM80 | 2.07 | 7.88E-04 | DE only in male placentas |
| ENSRNOG00000017980 | ITGAL | 1.40 | 3.19E-15 | DE only in male placentas |
| ENSRNOG00000022640 | AABR070032 35.1 | 1.56 | 1.31E-04 | DE only in male placentas |
| ENSRNOG00000045829 | THBS1 | 1.28 | 1.06E-14 | DE only in male placentas |
| ENSRNOG00000058422 | PIRB | 1.86 | 3.06E-08 | DE only in male placentas |
| ENSRNOG00000020991 | MS4A6A | 1.79 | 1.18E-13 | DE only in male placentas |
| ENSRNOG00000056151 | AABR070076 42.1 | 1.39 | 4.57E-15 | DE only in male placentas |

| | | | | | |
|---|---|---|---|---|---|
| ENSRNOG00000029212 | | VCAN | 1.72 | 5.02E-14 | DE only in male placentas |
| ENSRNOG00000009385 | | PIK3CG | 1.13 | 1.60E-09 | DE only in male placentas |
| ENSRNOG00000007810 | | GDF6 | 1.27 | 7.71E-03 | DE only in male placentas |
| ENSRNOG00000008668 | | VSIG8 | 2.47 | 8.68E-03 | DE only in male placentas |
| ENSRNOG00000031167 | | AABR070543 19.1 | 2.91 | 5.78E-14 | DE only in male placentas |
| ENSRNOG00000009329 | | NR1D1 | 1.27 | 5.89E-10 | DE only in male placentas |
| ENSRNOG00000008045 | | SLAMF9 | 1.77 | 9.95E-14 | DE only in male placentas |
| ENSRNOG00000007490 | | GABRR2 | 1.30 | 4.52E-18 | DE only in male placentas |
| ENSRNOG00000014124 | | NOD2 | 1.21 | 8.25E-04 | DE only in male placentas |
| ENSRNOG00000037231 | | PLA2G4D | 1.24 | 3.04E-13 | DE only in male placentas |
| ENSRNOG00000048161 | | TLR9 | 1.02 | 2.22E-04 | DE only in male placentas |
| ENSRNOG00000010266 | | CD180 | 1.05 | 2.04E-03 | DE only in male placentas |
| ENSRNOG00000002802 | | CXCL1 | 1.75 | 4.23E-14 | DE only in male placentas |
| ENSRNOG00000004698 | | CD244 | 1.16 | 4.55E-04 | DE only in male placentas |
| ENSRNOG00000023005 | X | LOC681325 | 1.74 | 1.05E-05 | DE only in male placentas |
| ENSRNOG00000032328 | | DIRAS2 | 1.14 | 1.21E-09 | DE only in male placentas |
| ENSRNOG00000060863 | | AABR070171 45.2 | 1.36 | 2.51E-07 | DE only in male placentas |
| ENSRNOG00000042717 | | CIART | 1.40 | 5.64E-08 | DE only in male placentas |

| | | | | | |
|---|---|---|---|---|---|
| ENSRNOG00000001252 | | CHST12 | 1.36 | 1.16E-08 | DE only in male placentas |
| ENSRNOG00000049633 | | FLVCR1 | 1.66 | 2.07E-26 | DE only in male placentas |
| ENSRNOG00000050996 | | KCTD4 | 1.18 | 4.25E-08 | DE only in male placentas |
| ENSRNOG00000001187 | | OASL | 1.48 | 1.24E-15 | DE only in male placentas |
| ENSRNOG00000027811 | | LILRB4 | 1.41 | 4.34E-09 | DE only in male placentas |
| ENSRNOG00000061394 | | CLEC4E | 1.34 | 4.13E-05 | DE only in male placentas |
| ENSRNOG00000057092 | | SLFN4 | 1.76 | 7.16E-14 | DE only in male placentas |
| ENSRNOG00000003887 | | LGI2 | 1.37 | 6.43E-04 | DE only in male placentas |
| ENSRNOG00000000522 | | CPNE5 | 1.31 | 5.46E-21 | DE only in male placentas |
| ENSRNOG00000009513 | | AKR1B1 | 1.26 | 1.13E-13 | DE only in male placentas |
| ENSRNOG00000053075 | | B4GALNT4 | 1.35 | 8.82E-20 | DE only in male placentas |
| ENSRNOG00000032240 | | GBP5 | 2.16 | 2.21E-14 | DE only in male placentas |
| ENSRNOG00000059857 | | RND1 | 1.19 | 8.17E-11 | DE only in male placentas |
| ENSRNOG00000003095 | | FGFBP1 | 1.04 | 6.26E-19 | DE only in male placentas |
| ENSRNOG00000052688 | | DNAH2 | 1.58 | 2.99E-18 | DE only in male placentas |
| ENSRNOG00000006698 | X | RAB33A | 1.51 | 1.09E-02 | DE only in male placentas |
| ENSRNOG00000031671 | | RASGEF1A | 1.27 | 1.69E-03 | DE only in male placentas |
| ENSRNOG00000003480 | | AIM2 | 1.65 | 8.97E-05 | DE only in male placentas |

| | | | | | |
|---|---|---|---|---|---|
| ENSRNOG00000028632 | | BCL2L14 | 1.15 | 1.02E-22 | DE only in male placentas |
| ENSRNOG00000031598 | | ATP8B4 | 1.38 | 2.90E-07 | DE only in male placentas |
| ENSRNOG00000050968 | | PTGER2 | 1.22 | 1.18E-03 | DE only in male placentas |
| ENSRNOG00000053891 | | AABR070180 78.1 | 1.42 | 2.71E-08 | DE only in male placentas |
| ENSRNOG00000024630 | | IGSF23 | 1.40 | 6.54E-15 | DE only in male placentas |
| ENSRNOG00000026647 | | CXCL16 | 1.18 | 8.76E-10 | DE only in male placentas |
| ENSRNOG00000047279 | | PGA5 | 1.21 | 2.30E-10 | DE only in male placentas |
| ENSRNOG00000029582 | X | LRCH2 | 1.24 | 1.56E-03 | DE only in male placentas |
| ENSRNOG00000002414 | | TFCP2L1 | 1.22 | 1.49E-07 | DE only in male placentas |
| ENSRNOG00000015868 | | INSL6 | 1.27 | 1.16E-06 | DE only in male placentas |
| ENSRNOG00000042287 | | MUCL3 | 2.01 | 2.39E-03 | DE only in male placentas |
| ENSRNOG00000061379 | | C7 | 1.66 | 6.82E-18 | DE only in male placentas |
| ENSRNOG00000032554 | | SCD4 | 1.64 | 1.13E-02 | DE only in male placentas |
| ENSRNOG00000020045 | | DRGX | 2.03 | 7.35E-05 | DE only in male placentas |
| ENSRNOG00000020687 | | DPF1 | 1.35 | 9.60E-09 | DE only in male placentas |
| ENSRNOG00000025615 | | RNASE1L1 | 1.43 | 6.31E-03 | DE only in male placentas |
| ENSRNOG00000004939 | | TTLL6 | 1.37 | 8.38E-04 | DE only in male placentas |
| ENSRNOG00000047783 | | TMEM200A | 1.71 | 5.88E-05 | DE only in male placentas |

| | | | | |
|---|---|---|---|---|
| ENSRNOG00000019065 | ZFP385B | 1.16 | 3.26E-04 | DE only in male placentas |
| ENSRNOG00000012972 | ALOX5 | 1.33 | 1.28E-04 | DE only in male placentas |
| ENSRNOG00000047800 | C5AR1 | 1.12 | 1.14E-07 | DE only in male placentas |
| ENSRNOG00000009589 | ANGPTL7 | 1.26 | 3.11E-07 | DE only in male placentas |
| ENSRNOG00000022565 | LRRC25 | 1.19 | 4.11E-08 | DE only in male placentas |
| ENSRNOG00000007604 | IGSF8 | 1.21 | 2.28E-09 | DE only in male placentas |
| ENSRNOG00000036819 | HSPB9 | 1.72 | 7.22E-04 | DE only in male placentas |
| ENSRNOG00000012307 | MYBPC3 | 1.92 | 7.11E-08 | DE only in male placentas |
| ENSRNOG00000060934 | PTX4 | 1.27 | 4.39E-06 | DE only in male placentas |
| ENSRNOG00000061561 | AABR070183 73.1 | 1.14 | 2.56E-03 | DE only in male placentas |
| ENSRNOG00000031523 | AABR070712 87.1 | 1.66 | 1.67E-02 | DE only in male placentas |
| ENSRNOG00000018505 | CIDEA | 1.42 | 1.68E-03 | DE only in male placentas |
| ENSRNOG00000001006 | NPTX2 | 1.02 | 3.32E-05 | DE only in male placentas |
| ENSRNOG00000017874 | CD53 | 1.24 | 7.81E-12 | DE only in male placentas |
| ENSRNOG00000019495 | GBX2 | 1.20 | 2.83E-05 | DE only in male placentas |
| ENSRNOG00000000137 | LY86 | 1.59 | 2.25E-09 | DE only in male placentas |
| ENSRNOG00000042524 | MILR1 | 1.40 | 7.40E-06 | DE only in male placentas |
| ENSRNOG00000003280 | GRIN2C | 1.05 | 3.46E-04 | DE only in male placentas |

| | | | | |
|---|---|---|---|---|
| ENSRNOG00000050509 | EBI3 | 1.22 | 1.39E-04 | DE only in male placentas |
| ENSRNOG00000020444 | HCN3 | 1.05 | 4.29E-06 | DE only in male placentas |
| ENSRNOG00000022691 | ST8SIA5 | 1.76 | 1.36E-02 | DE only in male placentas |
| ENSRNOG00000039754 | RAB7B | 1.19 | 3.64E-04 | DE only in male placentas |
| ENSRNOG00000036856 | AC097183.1 | 3.17 | 7.79E-10 | DE only in male placentas |
| ENSRNOG00000055393 | AABR070172<br>08.1 | 1.45 | 3.80E-02 | DE only in male placentas |
| ENSRNOG00000008078 | KCNH6 | 1.15 | 6.48E-12 | DE only in male placentas |
| ENSRNOG00000006418 | SAMHD1 | 1.21 | 2.52E-09 | DE only in male placentas |
| ENSRNOG00000019430 | CORO1A | 1.32 | 6.04E-19 | DE only in male placentas |
| ENSRNOG00000043071 | LINC00176 | 1.00 | 1.29E-02 | DE only in male placentas |
| ENSRNOG00000026235 | HK3 | 1.81 | 1.01E-10 | DE only in male placentas |
| ENSRNOG00000049642 | SMIM5 | 1.07 | 1.08E-09 | DE only in male placentas |
| ENSRNOG00000057522 | LOC1036899<br>68 | 1.22 | 1.61E-02 | DE only in male placentas |
| ENSRNOG00000018851 | PPP2R2B | 1.45 | 4.52E-06 | DE only in male placentas |
| ENSRNOG00000000412 | SLC35F1 | 1.17 | 9.64E-09 | DE only in male placentas |
| ENSRNOG00000022859 | TREM1 | 1.38 | 5.49E-05 | DE only in male placentas |
| ENSRNOG00000031743 | GBP2 | 1.67 | 2.91E-21 | DE only in male placentas |
| ENSRNOG00000023085 | PMEL | 1.07 | 1.17E-05 | DE only in male placentas |

| | | | | | |
|---|---|---|---|---|---|
| ENSRNOG00000004133 | X | SLC7A3 | 2.10 | 2.95E-04 | DE only in male placentas |
| ENSRNOG00000004410 | | PAX6 | 1.12 | 2.76E-03 | DE only in male placentas |
| ENSRNOG00000045992 | X | TLR8 | 1.03 | 1.59E-06 | DE only in male placentas |
| ENSRNOG00000054418 | | AABR07005779.3 | 1.93 | 3.17E-03 | DE only in male placentas |
| ENSRNOG00000004772 | | CYTIP | 1.02 | 1.85E-06 | DE only in male placentas |
| ENSRNOG00000008182 | | HTRA3 | 1.12 | 5.80E-07 | DE only in male placentas |
| ENSRNOG00000023008 | | FAM131C | 1.45 | 8.29E-08 | DE only in male placentas |
| ENSRNOG00000009331 | | HCK | 1.58 | 2.35E-14 | DE only in male placentas |
| ENSRNOG00000057724 | | AC118957.1 | 1.42 | 2.61E-02 | DE only in male placentas |
| ENSRNOG00000024494 | | AABR07056633.1 | 1.39 | 1.53E-02 | DE only in male placentas |
| ENSRNOG00000013586 | | SNAI3 | 1.17 | 7.27E-05 | DE only in male placentas |
| ENSRNOG00000024159 | | FCER1G | 1.33 | 1.12E-13 | DE only in male placentas |
| ENSRNOG00000017783 | | SFRP1 | 1.26 | 7.78E-08 | DE only in male placentas |
| ENSRNOG00000020784 | | KCNK7 | 1.79 | 4.82E-08 | DE only in male placentas |
| ENSRNOG00000051027 | X | AABR07038886.1 | 1.12 | 3.92E-07 | DE only in male placentas |
| ENSRNOG00000010047 | | DDIT4L2 | 1.54 | 1.74E-02 | DE only in male placentas |
| ENSRNOG00000001439 | | SRRM3 | 1.02 | 9.08E-03 | DE only in male placentas |
| ENSRNOG00000012779 | | MSR1 | 1.28 | 9.66E-07 | DE only in male placentas |

| | | | | |
|---|---|---|---|---|
| ENSRNOG00000049942 | RGD1564899 | 1.07 | 2.45E-09 | DE only in male placentas |
| ENSRNOG00000054250 | FAM186B | 2.26 | 7.90E-03 | DE only in male placentas |
| ENSRNOG00000055156 | LOC1036943 80 | 1.03 | 2.17E-02 | DE only in male placentas |
| ENSRNOG00000027900 | GLT6D1 | 1.18 | 3.98E-02 | DE only in male placentas |
| ENSRNOG00000009528 | SDCBP2 | 1.29 | 8.06E-09 | DE only in male placentas |
| ENSRNOG00000047928 | AABR070486 53.1 | 1.64 | 7.34E-04 | DE only in male placentas |
| ENSRNOG00000018141 | RGD1307443 | 1.61 | 2.79E-06 | DE only in male placentas |
| ENSRNOG00000011809 | AABR070724 49.1 | 1.01 | 1.73E-03 | DE only in male placentas |
| ENSRNOG00000003972 | TSHR | 1.00 | 6.40E-03 | DE only in male placentas |
| ENSRNOG00000021018 | BATF2 | 1.57 | 3.61E-02 | DE only in male placentas |
| ENSRNOG00000022242 | CXCL9 | 1.77 | 1.90E-03 | DE only in male placentas |
| ENSRNOG00000024931 | CCDC88B | 1.31 | 4.59E-13 | DE only in male placentas |
| ENSRNOG00000027833 | FNIP2 | 1.20 | 8.91E-19 | DE only in male placentas |
| ENSRNOG00000013825 | RAP1GAP | 1.60 | 1.41E-16 | DE only in male placentas |
| ENSRNOG00000046864 | ACOT4 | 1.75 | 2.09E-03 | DE only in male placentas |
| ENSRNOG00000016112 | CD274 | 1.51 | 7.48E-12 | DE only in male placentas |
| ENSRNOG00000021220 | CPXM1 | 1.23 | 3.28E-06 | DE only in male placentas |
| ENSRNOG00000007350 | RAC2 | 1.41 | 1.76E-11 | DE only in male placentas |

| | | | | | |
|---|---|---|---|---|---|
| ENSRNOG00000009465 | | SFRP2 | 1.22 | 1.21E-02 | DE only in male placentas |
| ENSRNOG00000015157 | | SMTNL2 | 1.03 | 8.28E-16 | DE only in male placentas |
| ENSRNOG00000004649 | | IL1B | 1.47 | 1.59E-09 | DE only in male placentas |
| ENSRNOG00000014202 | | SNX20 | 1.01 | 3.99E-04 | DE only in male placentas |
| ENSRNOG00000052725 | | SHISAL1 | 1.04 | 2.84E-09 | DE only in male placentas |
| ENSRNOG00000003947 | | NTN1 | 1.12 | 4.29E-09 | DE only in male placentas |
| ENSRNOG00000051136 | X | AABR070409 51.1 | 2.63 | 4.10E-02 | DE only in male placentas |
| ENSRNOG00000007030 | | EPHA7 | 1.02 | 1.53E-07 | DE only in male placentas |
| ENSRNOG00000025624 | | ARHGAP20 | 1.33 | 7.66E-13 | DE only in male placentas |
| ENSRNOG00000047091 | | PSMB11 | 2.47 | 2.15E-02 | DE only in male placentas |
| ENSRNOG00000012749 | | C1QB | 1.25 | 8.64E-12 | DE only in male placentas |
| ENSRNOG00000007989 | | CHST1 | 1.23 | 1.25E-04 | DE only in male placentas |
| ENSRNOG00000021802 | | ISG15 | 1.24 | 8.98E-17 | DE only in male placentas |
| ENSRNOG00000048222 | | NLRC5 | 1.12 | 8.30E-10 | DE only in male placentas |
| ENSRNOG00000016776 | | EXTL1 | 1.12 | 4.34E-12 | DE only in male placentas |
| ENSRNOG00000009912 | | FGR | 1.21 | 1.01E-14 | DE only in male placentas |
| ENSRNOG00000046744 | | ANKRD29 | 1.30 | 6.68E-05 | DE only in male placentas |
| ENSRNOG00000038835 | | CD86 | 1.11 | 1.23E-04 | DE only in male placentas |

| | | | | | |
|---|---|---|---|---|---|
| ENSRNOG00000022839 | | IFIT3 | 1.24 | 1.19E-11 | DE only in male placentas |
| ENSRNOG00000000700 | | TMEM119 | 1.33 | 4.10E-07 | DE only in male placentas |
| ENSRNOG00000030914 | | LEXM | 2.31 | 1.55E-08 | DE only in male placentas |
| ENSRNOG00000009471 | | EPSTI1 | 1.49 | 1.76E-08 | DE only in male placentas |
| ENSRNOG00000003806 | | GPR65 | 1.57 | 5.62E-04 | DE only in male placentas |
| ENSRNOG00000017539 | | MMP9 | 1.16 | 4.80E-11 | DE only in male placentas |
| ENSRNOG00000042581 | | PTCRA | 2.67 | 9.05E-05 | DE only in male placentas |
| ENSRNOG00000003622 | X | CYBB | 1.11 | 1.86E-09 | DE only in male placentas |
| ENSRNOG00000038722 | | TLR1 | 1.38 | 8.00E-10 | DE only in male placentas |
| ENSRNOG00000002776 | | SELL | 1.00 | 8.44E-07 | DE only in male placentas |
| ENSRNOG00000032596 | | RT1-T24-1 | 1.36 | 8.91E-04 | DE only in male placentas |
| ENSRNOG00000057996 | | PHACTR3 | 1.24 | 5.28E-03 | DE only in male placentas |
| ENSRNOG00000021586 | | GPR39 | 1.09 | 6.16E-07 | DE only in male placentas |
| ENSRNOG00000032708 | | RT1-BB | 1.11 | 8.59E-05 | DE only in male placentas |
| ENSRNOG00000019728 | | ITGAM | 1.33 | 1.82E-12 | DE only in male placentas |
| ENSRNOG00000001492 | | SLC8A2 | 3.11 | 6.18E-04 | DE only in male placentas |
| ENSRNOG00000011550 | | KCNAB2 | 1.15 | 2.96E-04 | DE only in male placentas |
| ENSRNOG00000046184 | | RGD1561143 | 1.03 | 1.68E-02 | DE only in male placentas |

| | | | | |
|---|---|---|---|---|
| ENSRNOG00000059266 | AABR070026 74.1 | 1.37 | 3.84E-03 | DE only in male placentas |
| ENSRNOG00000025590 | ARL14EPL | 1.04 | 7.59E-04 | DE only in male placentas |
| ENSRNOG00000029658 | RNF213 | 1.11 | 7.34E-12 | DE only in male placentas |
| ENSRNOG00000002303 | KCNJ12 | 1.15 | 3.97E-07 | DE only in male placentas |
| ENSRNOG00000052064 | PARVG | 1.20 | 4.10E-06 | DE only in male placentas |
| ENSRNOG00000000655 | PTPRC | 1.00 | 3.16E-11 | DE only in male placentas |
| ENSRNOG00000049282 | OAS2 | 1.04 | 1.82E-06 | DE only in male placentas |
| ENSRNOG00000023969 | HERC6 | 1.03 | 1.58E-08 | DE only in male placentas |
| ENSRNOG00000010519 | CYP39A1 | 1.69 | 3.15E-07 | DE only in male placentas |
| ENSRNOG00000025948 | INAVA | 1.21 | 6.98E-08 | DE only in male placentas |
| ENSRNOG00000005900 | SLC2A6 | 1.00 | 1.42E-11 | DE only in male placentas |
| ENSRNOG00000050155 | RGD1560112 | 1.29 | 2.91E-02 | DE only in male placentas |
| ENSRNOG00000012807 | C1QA | 1.10 | 3.15E-10 | DE only in male placentas |
| ENSRNOG00000024277 | AABR070354 70.1 | 1.82 | 1.79E-09 | DE only in male placentas |
| ENSRNOG00000004192 | ARHGAP30 | 1.22 | 5.12E-12 | DE only in male placentas |
| ENSRNOG00000020686 | DKKL1 | 1.02 | 2.24E-10 | DE only in male placentas |
| ENSRNOG00000001514 | CDCA7 | 1.03 | 4.91E-12 | DE only in male placentas |
| ENSRNOG00000019050 | IFIT1 | 1.01 | 1.28E-02 | DE only in male placentas |

| | | | | | |
|---|---|---|---|---|---|
| ENSRNOG00000054973 | | SNORA65 | 1.60 | 4.77E-02 | DE only in male placentas |
| ENSRNOG00000024331 | | UPK1A | 1.21 | 2.19E-02 | DE only in male placentas |
| ENSRNOG00000020355 | | TWIST2 | 1.19 | 2.45E-03 | DE only in male placentas |
| ENSRNOG00000043296 | | ABCA14 | 2.55 | 1.27E-02 | DE only in male placentas |
| ENSRNOG00000057189 | | LOC1036903 26 | 1.80 | 2.91E-03 | DE only in male placentas |
| ENSRNOG00000010018 | | CLEC4A3 | 1.04 | 1.95E-05 | DE only in male placentas |
| ENSRNOG00000000187 | | CSF2RB | 1.08 | 2.89E-11 | DE only in male placentas |
| ENSRNOG00000054524 | | TNIP3 | 1.69 | 1.05E-03 | DE only in male placentas |
| ENSRNOG00000031168 | | ARHGAP15 | 1.16 | 1.51E-02 | DE only in male placentas |
| ENSRNOG00000047223 | | ARRDC5 | 1.00 | 1.42E-02 | DE only in male placentas |
| ENSRNOG00000020845 | | TYROBP | 1.22 | 1.35E-11 | DE only in male placentas |
| ENSRNOG00000010603 | | GZMA | 1.19 | 1.10E-03 | DE only in male placentas |
| ENSRNOG00000039390 | | SLC37A2 | 1.00 | 3.19E-07 | DE only in male placentas |
| ENSRNOG00000029195 | | UBA7 | 1.43 | 6.97E-14 | DE only in male placentas |
| ENSRNOG00000010208 | X | TIMP1 | 1.21 | 8.31E-14 | DE only in male placentas |
| ENSRNOG00000011971 | | C1S | 1.29 | 2.09E-15 | DE only in male placentas |
| ENSRNOG00000021856 | | LAT2 | 1.17 | 5.51E-03 | DE only in male placentas |
| ENSRNOG00000056850 | | LOC1009097 04 | 2.00 | 3.22E-05 | DE only in male placentas |

| | | | | |
|---|---|---|---|---|
| ENSRNOG00000000457 | TAP1 | 1.11 | 1.84E-12 | DE only in male placentas |
| ENSRNOG00000024230 | TNFAIP8L3 | 1.80 | 6.28E-04 | DE only in male placentas |
| ENSRNOG00000002470 | IFI47 | 1.00 | 9.20E-07 | DE only in male placentas |
| ENSRNOG00000036604 | IFIT2 | 1.21 | 1.10E-09 | DE only in male placentas |
| ENSRNOG00000003895 | RGS1 | 1.74 | 3.37E-09 | DE only in male placentas |
| ENSRNOG00000005787 | ERP27 | 1.67 | 8.03E-03 | DE only in male placentas |
| ENSRNOG00000058560 | COL2A1 | 1.24 | 4.95E-03 | DE only in male placentas |
| ENSRNOG00000053272 | CHI3L1 | 1.71 | 1.06E-10 | DE only in male placentas |
| ENSRNOG00000020263 | ATP1A3 | 1.56 | 2.06E-05 | DE only in male placentas |
| ENSRNOG00000013269 | TNFSF10 | 1.16 | 5.28E-04 | DE only in male placentas |
| ENSRNOG00000037814 | CXCL17 | 3.08 | 2.89E-06 | DE only in male placentas |
| ENSRNOG00000014156 | FUT7 | 1.79 | 2.40E-03 | DE only in male placentas |
| ENSRNOG00000053658 | AABR070445 69.1 | 1.88 | 1.13E-02 | DE only in male placentas |
| ENSRNOG00000026387 | MS4A12 | 1.82 | 2.27E-02 | DE only in male placentas |
| ENSRNOG00000014333 | VCAM1 | 1.44 | 5.47E-17 | DE only in male placentas |
| ENSRNOG00000056783 | CD69 | 1.33 | 9.74E-04 | DE only in male placentas |
| ENSRNOG00000037085 | XIRP1 | 1.50 | 3.41E-12 | DE only in male placentas |
| ENSRNOG00000009922 | PRLHR | 1.52 | 6.63E-06 | DE only in male placentas |

| | | | | |
|---|---|---|---|---|
| ENSRNOG00000045560 | GVIN1 | 1.09 | 3.41E-06 | DE only in male placentas |
| ENSRNOG00000003486 | MNDA | 1.34 | 5.03E-06 | DE only in male placentas |
| ENSRNOG00000021644 | SLC15A3 | 1.26 | 5.77E-15 | DE only in male placentas |
| ENSRNOG00000006268 | IFNB1 | 2.60 | 5.41E-03 | DE only in male placentas |
| ENSRNOG00000009730 | CYP7B1 | 1.48 | 1.21E-06 | DE only in male placentas |
| ENSRNOG00000001480 | NCF1 | 1.15 | 3.17E-09 | DE only in male placentas |
| ENSRNOG00000059207 | OAS3 | 1.86 | 2.21E-09 | DE only in male placentas |
| ENSRNOG00000037931 | PLAUR | 1.13 | 3.25E-15 | DE only in male placentas |
| ENSRNOG00000046007 | CLDN3 | 1.27 | 1.04E-06 | DE only in male placentas |
| ENSRNOG00000051986 | PLCG2 | 1.29 | 5.53E-16 | DE only in male placentas |
| ENSRNOG00000004400 | AVPR1A | 1.26 | 1.93E-03 | DE only in male placentas |
| ENSRNOG00000013829 | CHRNA3 | 1.44 | 3.12E-06 | DE only in male placentas |
| ENSRNOG00000008518 | GSG1 | 1.93 | 4.99E-08 | DE only in male placentas |
| ENSRNOG00000017819 | CD14 | 1.14 | 3.19E-16 | DE only in male placentas |
| ENSRNOG00000023428 | PIK3R5 | 1.27 | 1.40E-08 | DE only in male placentas |
| ENSRNOG00000019627 | MYBPC2 | 2.00 | 4.07E-02 | DE only in male placentas |
| ENSRNOG00000007561 | GLB1L2 | 1.05 | 7.14E-04 | DE only in male placentas |
| ENSRNOG00000021116 | BNIPL | 1.10 | 8.81E-03 | DE only in male placentas |

| | | | | |
|---|---|---|---|---|
| ENSRNOG00000023318 | TIGD3 | 1.38 | 3.20E-03 | DE only in male placentas |
| ENSRNOG00000051235 | C2 | 1.11 | 1.17E-09 | DE only in male placentas |
| ENSRNOG00000028879 | SLC4A8 | 1.21 | 7.19E-04 | DE only in male placentas |
| ENSRNOG00000014893 | WDR63 | 1.61 | 2.84E-04 | DE only in male placentas |
| ENSRNOG00000019661 | GDF15 | 1.51 | 2.04E-03 | DE only in male placentas |
| ENSRNOG00000039204 | PRAM1 | 1.16 | 2.53E-03 | DE only in male placentas |
| ENSRNOG00000023227 | AABR07052585.2 | 2.06 | 4.21E-03 | DE only in male placentas |
| ENSRNOG00000000699 | SELPLG | 1.13 | 1.35E-07 | DE only in male placentas |
| ENSRNOG00000012804 | C1QC | 1.16 | 7.45E-09 | DE only in male placentas |
| ENSRNOG00000052569 | AABR07030834.1 | 1.35 | 2.72E-03 | DE only in male placentas |
| ENSRNOG00000000693 | SVOP | 1.47 | 1.92E-02 | DE only in male placentas |
| ENSRNOG00000052121 | AC117065.1 | 1.47 | 4.15E-05 | DE only in male placentas |
| ENSRNOG00000037134 | SHC4 | 1.44 | 3.43E-05 | DE only in male placentas |
| ENSRNOG00000027960 | AC094055.1 | 1.19 | 3.89E-02 | DE only in male placentas |
| ENSRNOG00000049517 | TNFAIP3 | 1.07 | 3.21E-06 | DE only in male placentas |
| ENSRNOG00000014464 | TNFSF13B | 1.18 | 3.07E-03 | DE only in male placentas |
| ENSRNOG00000029191 | LOC685067 | 1.19 | 1.48E-08 | DE only in male placentas |
| ENSRNOG00000019675 | PYCARD | 1.21 | 1.80E-06 | DE only in male placentas |

| | | | | |
|---|---|---|---|---|
| ENSRNOG00000014956 | SLC11A1 | 1.22 | 8.32E-07 | DE only in male placentas |
| ENSRNOG00000000459 | PSMB9 | 1.47 | 1.46E-20 | DE only in male placentas |
| ENSRNOG00000060142 | AABR070726 64.1 | 1.96 | 8.26E-03 | DE only in male placentas |
| ENSRNOG00000006132 | FAM71F1 | 1.29 | 2.09E-03 | DE only in male placentas |
| ENSRNOG00000016997 | AABR070444 42.1 | 1.66 | 1.26E-03 | DE only in male placentas |
| ENSRNOG00000001164 | TFF1 | 2.01 | 9.48E-03 | DE only in male placentas |
| ENSRNOG00000036951 | SLC2A7 | 1.04 | 7.04E-03 | DE only in male placentas |
| ENSRNOG00000036588 | AC121209.1 | 2.05 | 4.49E-02 | DE only in male placentas |
| ENSRNOG00000014166 | SMOC2 | 1.42 | 1.16E-07 | DE only in male placentas |
| ENSRNOG00000019854 | NAPSA | 1.12 | 7.39E-06 | DE only in male placentas |
| ENSRNOG00000048782 | WNT7A | 1.37 | 7.88E-05 | DE only in male placentas |
| ENSRNOG00000043451 | SPP1 | 1.20 | 3.69E-05 | DE only in male placentas |
| ENSRNOG00000003020 | SLC25A47 | 1.84 | 1.16E-03 | DE only in male placentas |
| ENSRNOG00000017487 | OTOA | 1.01 | 3.39E-03 | DE only in male placentas |
| ENSRNOG00000050647 | HSPA1B | 1.05 | 5.80E-07 | DE only in male placentas |
| ENSRNOG00000020778 | CHRNB2 | 1.26 | 7.05E-03 | DE only in male placentas |
| ENSRNOG00000019141 | CH25H | 1.14 | 8.42E-06 | DE only in male placentas |
| ENSRNOG00000001860 | CCDC116 | 1.35 | 3.49E-03 | DE only in male placentas |

| | | | | |
|---|---|---|---|---|
| ENSRNOG00000016177 | SCARA3 | 1.08 | 4.16E-04 | DE only in male placentas |
| ENSRNOG00000033010 | AKR1C12 | 2.88 | 1.75E-02 | DE only in male placentas |
| ENSRNOG00000012172 | SPI1 | 1.35 | 6.78E-09 | DE only in male placentas |
| ENSRNOG00000006019 | G0S2 | 1.12 | 2.28E-04 | DE only in male placentas |
| ENSRNOG00000002134 | GBP6 | 1.04 | 3.55E-05 | DE only in male placentas |
| ENSRNOG00000023870 | SH2D7 | 1.15 | 1.01E-04 | DE only in male placentas |
| ENSRNOG00000028548 | CCL9 | 1.33 | 7.00E-04 | DE only in male placentas |
| ENSRNOG00000000889 | AABR070110 57.1 | 1.67 | 1.26E-04 | DE only in male placentas |
| ENSRNOG00000020980 | AABR070062 78.1 | 1.32 | 2.26E-02 | DE only in male placentas |
| ENSRNOG00000000456 | PSMB8 | 1.04 | 2.71E-09 | DE only in male placentas |
| ENSRNOG00000052587 | LOC1009094 74 | 4.57 | 2.09E-03 | DE only in male placentas |
| ENSRNOG00000018236 | CYM | 2.24 | 6.21E-06 | DE only in male placentas |
| ENSRNOG00000017302 | SLC6A3 | 1.42 | 2.45E-11 | DE only in male placentas |
| ENSRNOG00000000837 | TNF | 5.87 | 9.58E-05 | DE only in male placentas |
| ENSRNOG00000018915 | TAGAP | 1.03 | 3.34E-05 | DE only in male placentas |
| ENSRNOG00000009342 | FCNB | 1.16 | 3.40E-04 | DE only in male placentas |
| ENSRNOG00000052542 | PILRB | 2.14 | 4.52E-02 | DE only in male placentas |
| ENSRNOG00000009919 | ACOD1 | 2.31 | 2.65E-09 | DE only in male placentas |

| | | | | | |
|---|---|---|---|---|---|
| ENSRNOG00000042649 | | A4GNT | 1.90 | 1.75E-03 | DE only in male placentas |
| ENSRNOG00000055032 | | AABR070707 74.1 | 2.65 | 4.42E-02 | DE only in male placentas |
| ENSRNOG00000050210 | | RT1-CE10 | 1.25 | 1.78E-05 | DE only in male placentas |
| ENSRNOG00000015366 | | NEURL3 | 1.18 | 2.64E-05 | DE only in male placentas |
| ENSRNOG00000037149 | | BCL2L15 | 1.09 | 1.14E-02 | DE only in male placentas |
| ENSRNOG00000011174 | | FPR1 | 1.74 | 1.57E-02 | DE only in male placentas |
| ENSRNOG00000002964 | | XCL1 | 1.06 | 9.04E-03 | DE only in male placentas |
| ENSRNOG00000061639 | | AABR070444 04.1 | 1.23 | 3.63E-03 | DE only in male placentas |
| ENSRNOG00000015616 | | RGS14 | 1.23 | 2.52E-03 | DE only in male placentas |
| ENSRNOG00000058834 | | LOC1036924 71 | 1.20 | 1.55E-03 | DE only in male placentas |
| ENSRNOG00000002923 | X | ZCCHC13 | 2.14 | 8.70E-03 | DE only in male placentas |
| ENSRNOG00000050743 | | LOC1009117 96 | 1.23 | 3.26E-02 | DE only in male placentas |
| ENSRNOG00000011033 | | TEKT2 | 2.14 | 1.30E-02 | DE only in male placentas |
| ENSRNOG00000024257 | | IRGC | 1.60 | 3.93E-04 | DE only in male placentas |
| ENSRNOG00000008431 | | GABBR2 | 1.12 | 2.29E-04 | DE only in male placentas |
| ENSRNOG00000057301 | | LOC1009098 69 | 3.56 | 1.82E-02 | DE only in male placentas |
| ENSRNOG00000039255 | | GPR31 | 1.76 | 2.86E-02 | DE only in male placentas |
| ENSRNOG00000012390 | | NPW | 1.61 | 5.02E-03 | DE only in male placentas |

| | | | | | |
|---|---|---|---|---|---|
| ENSRNOG00000055956 | | AABR070150 78.1 | 1.80 | 1.21E-04 | DE only in male placentas |
| ENSRNOG00000060896 | | AABR070634 24.1 | 1.25 | 7.85E-03 | DE only in male placentas |
| ENSRNOG00000056747 | | AABR070150 55.1 | 1.32 | 3.73E-03 | DE only in male placentas |
| ENSRNOG00000059586 | | AABR070150 80.2 | 1.32 | 3.73E-03 | DE only in male placentas |
| ENSRNOG00000046600 | | AABR070150 66.1 | 1.30 | 4.35E-03 | DE only in male placentas |
| ENSRNOG00000050485 | | GAS1 | 1.18 | 4.40E-02 | DE only in male placentas |
| ENSRNOG00000057769 | | OAS1H | 1.63 | 4.20E-02 | DE only in male placentas |
| ENSRNOG00000053699 | | AC096809.1 | 2.79 | 4.70E-02 | DE only in male placentas |
| ENSRNOG00000024030 | | GPR35 | 1.82 | 3.54E-02 | DE only in male placentas |
| ENSRNOG00000010906 | | CCL5 | 1.18 | 1.05E-02 | DE only in male placentas |
| ENSRNOG00000020747 | | NKX2-5 | 3.54 | 3.00E-04 | DE only in male placentas |
| ENSRNOG00000059837 | | CDKN2A | 1.37 | 7.02E-04 | DE only in male placentas |
| ENSRNOG00000013622 | | AABR070328 21.1 | 2.26 | 1.84E-03 | DE only in male placentas |
| ENSRNOG00000018854 | | NPPC | 1.77 | 2.59E-02 | DE only in male placentas |
| ENSRNOG00000053717 | | METAZOA_S RP | 1.17 | 4.08E-02 | DE only in male placentas |
| ENSRNOG00000010785 | | HELT | 2.19 | 1.58E-02 | DE only in male placentas |
| ENSRNOG00000015137 | | SPATA6L | 1.93 | 4.06E-02 | DE only in male placentas |
| ENSRNOG00000002429 | X | RTL3 | 3.00 | 2.15E-02 | DE only in male placentas |

| | | | | | |
|---|---|---|---|---|---|
| ENSRNOG00000033748 | | AABR070606 10.1 | 4.53 | 2.28E-03 | DE only in male placentas |
| ENSRNOG00000060370 | | AABR070281 13.1 | -3.57 | 1.23E-02 | DE only in male placentas |
| ENSRNOG00000053818 | X | AABR070409 06.1 | -2.90 | 4.16E-02 | DE only in male placentas |
| ENSRNOG00000059261 | | AABR070017 00.2 | -2.87 | 3.14E-02 | DE only in male placentas |
| ENSRNOG00000057798 | | AC102976.1 | -1.25 | 3.43E-02 | DE only in male placentas |
| ENSRNOG00000032490 | | CDH7 | -1.98 | 1.85E-02 | DE only in male placentas |
| ENSRNOG00000011823 | | TFAP2B | -2.09 | 2.08E-02 | DE only in male placentas |
| ENSRNOG00000060173 | | AABR070445 20.1 | -2.94 | 1.08E-02 | DE only in male placentas |
| ENSRNOG00000050042 | | MYH8 | -2.81 | 4.40E-02 | DE only in male placentas |
| ENSRNOG00000022595 | | LOC1003629 65 | -2.76 | 2.45E-02 | DE only in male placentas |
| ENSRNOG00000048765 | | AABR070027 74.3 | -2.80 | 4.81E-02 | DE only in male placentas |
| ENSRNOG00000054506 | | NCCRP1 | -2.52 | 1.88E-02 | DE only in male placentas |
| ENSRNOG00000002484 | | CHRNA9 | -3.15 | 2.28E-03 | DE only in male placentas |
| ENSRNOG00000050885 | | LOC1009105 26 | -2.78 | 4.52E-02 | DE only in male placentas |
| ENSRNOG00000049523 | | AC097153.2 | -2.85 | 3.20E-02 | DE only in male placentas |
| ENSRNOG00000004623 | | RAG2 | -2.91 | 2.53E-02 | DE only in male placentas |
| ENSRNOG00000048493 | | AABR070082 93.3 | -2.31 | 6.54E-04 | DE only in male placentas |
| ENSRNOG00000058829 | | AABR070311 50.1 | -3.08 | 4.08E-02 | DE only in male placentas |

| | | | | | |
|---|---|---|---|---|---|
| ENSRNOG00000048672 | | TMEM207 | -1.94 | 1.37E-02 | DE only in male placentas |
| ENSRNOG00000001329 | | GJC3 | -2.73 | 4.67E-02 | DE only in male placentas |
| ENSRNOG00000043258 | | BPIFA6 | -1.70 | 1.51E-02 | DE only in male placentas |
| ENSRNOG00000062251 | | AABR07055844.1 | -4.66 | 5.92E-03 | DE only in male placentas |
| ENSRNOG00000019199 | | CRACR2B | -5.18 | 2.16E-03 | DE only in male placentas |
| ENSRNOG00000042950 | | SLC5A9 | -2.42 | 4.34E-02 | DE only in male placentas |
| ENSRNOG00000052680 | | AABR07014630.1 | -3.47 | 1.68E-02 | DE only in male placentas |
| ENSRNOG00000021896 | | MMP20 | -1.14 | 2.86E-02 | DE only in male placentas |
| ENSRNOG00000003883 | X | LOC317456 | -1.23 | 9.06E-03 | DE only in male placentas |
| ENSRNOG00000056752 | | LOC689519 | -1.64 | 3.51E-03 | DE only in male placentas |
| ENSRNOG00000018978 | | SCART1 | -3.49 | 1.47E-02 | DE only in male placentas |
| ENSRNOG00000042690 | | ZMAT4 | -1.36 | 7.13E-03 | DE only in male placentas |
| ENSRNOG00000022181 | | LOC103693984 | -3.14 | 1.63E-02 | DE only in male placentas |
| ENSRNOG00000003836 | X | FAM47A | -3.42 | 3.49E-03 | DE only in male placentas |
| ENSRNOG00000058699 | | AABR07036452.1 | -2.67 | 7.62E-03 | DE only in male placentas |
| ENSRNOG00000049162 | | PRR15L | -2.05 | 6.92E-05 | DE only in male placentas |
| ENSRNOG00000039476 | | PCDHB2 | -2.44 | 1.57E-02 | DE only in male placentas |
| ENSRNOG00000011132 | | LOC100362814 | -1.34 | 2.66E-02 | DE only in male placentas |

| | | | | |
|---|---|---|---|---|
| ENSRNOG00000061917 | AABR070428<br>40.1 | -2.07 | 4.24E-02 | DE only in male placentas |
| ENSRNOG00000013484 | GSTA1 | -1.07 | 2.61E-03 | DE only in male placentas |
| ENSRNOG00000048232 | AABR070443<br>02.1 | -1.86 | 2.82E-02 | DE only in male placentas |
| ENSRNOG00000003601 | RGD1564463 | -2.32 | 1.79E-02 | DE only in male placentas |
| ENSRNOG00000054544 | AABR070213<br>57.1 | -1.42 | 2.21E-02 | DE only in male placentas |
| ENSRNOG00000010741 | TIGD4 | -1.84 | 9.08E-03 | DE only in male placentas |
| ENSRNOG00000040128 | CD209A | -2.26 | 3.17E-02 | DE only in male placentas |
| ENSRNOG00000050229 | LOC1025503<br>14 | -1.09 | 1.98E-02 | DE only in male placentas |
| ENSRNOG00000005038 | UBE2T | -1.21 | 8.56E-03 | DE only in male placentas |
| ENSRNOG00000054702 | AABR070681<br>98.2 | -1.79 | 9.98E-04 | DE only in male placentas |
| ENSRNOG00000007410 | DAB1 | -1.46 | 4.42E-02 | DE only in male placentas |
| ENSRNOG00000048575 | VOM2R1 | -1.85 | 3.45E-03 | DE only in male placentas |
| ENSRNOG00000042918 | RGD1304770 | -1.18 | 4.99E-02 | DE only in male placentas |
| ENSRNOG00000050795 | AABR070080<br>16.1 | -1.43 | 7.20E-04 | DE only in male placentas |
| ENSRNOG00000049822 | LOC1036900<br>64 | -3.83 | 2.73E-02 | DE only in male placentas |
| ENSRNOG00000053118 | LOC1036948<br>67 | -4.21 | 1.32E-02 | DE only in male placentas |
| ENSRNOG00000058357 | GSTM3 | -5.02 | 2.87E-03 | DE only in male placentas |
| ENSRNOG00000048562 | AC109542.1 | -6.31 | 2.30E-04 | DE only in male placentas |

| | | | | |
|---|---|---|---|---|
| ENSRNOG00000045653 | LOC1009124 98 | -6.62 | 1.25E-04 | DE only in male placentas |
| ENSRNOG00000048353 | APEG3 | -6.79 | 8.78E-05 | DE only in male placentas |
| ENSRNOG00000011635 | CES2E | -2.26 | 7.59E-03 | DE only in male placentas |
| ENSRNOG00000061630 | AABR070098 34.1 | -1.43 | 6.81E-03 | DE only in male placentas |
| ENSRNOG00000017538 | LRRC27 | -1.43 | 2.00E-02 | DE only in male placentas |
| ENSRNOG00000033929 | STFA2L1 | -2.11 | 5.54E-03 | DE only in male placentas |
| ENSRNOG00000059734 | AABR070435 64.1 | -2.85 | 4.34E-03 | DE only in male placentas |
| ENSRNOG00000023588 | DMGDH | -1.29 | 3.92E-04 | DE only in male placentas |
| ENSRNOG00000000920 | PHKG1 | -1.09 | 4.04E-02 | DE only in male placentas |
| ENSRNOG00000017998 | S100Z | -1.89 | 7.25E-05 | DE only in male placentas |
| ENSRNOG00000005519 | GRM3 | -3.65 | 2.59E-05 | DE only in male placentas |
| ENSRNOG00000056092 | HEPHL1 | -1.00 | 3.90E-02 | DE only in male placentas |
| ENSRNOG00000011754 | MYOM2 | -1.58 | 1.98E-03 | DE only in male placentas |
| ENSRNOG00000007202 | SEMA3D | -1.00 | 3.16E-03 | DE only in male placentas |
| ENSRNOG00000036661 | RAB40B | -1.01 | 3.41E-07 | DE only in male placentas |
| ENSRNOG00000020059 | PCDHB14 | -1.13 | 3.37E-02 | DE only in male placentas |
| ENSRNOG00000039025 | AABR070519 47.1 | -1.33 | 3.53E-02 | DE only in male placentas |
| ENSRNOG00000014018 | ST8SIA1 | -1.74 | 1.42E-02 | DE only in male placentas |

| | | | | | |
|---|---|---|---|---|---|
| ENSRNOG00000005853 | | TACR1 | -1.47 | 2.43E-03 | DE only in male placentas |
| ENSRNOG00000032426 | | SLC10A5 | -1.09 | 1.38E-02 | DE only in male placentas |
| ENSRNOG00000022937 | | BPIFC | -1.09 | 3.25E-02 | DE only in male placentas |
| ENSRNOG00000039624 | | AABR070247 32.1 | -3.18 | 1.26E-02 | DE only in male placentas |
| ENSRNOG00000014937 | | LRRC71 | -1.41 | 1.15E-06 | DE only in male placentas |
| ENSRNOG00000023920 | | ZFP791 | -1.01 | 3.02E-06 | DE only in male placentas |
| ENSRNOG00000016306 | | ADGRV1 | -1.01 | 3.48E-04 | DE only in male placentas |
| ENSRNOG00000006086 | | LYNX1 | -1.44 | 3.18E-02 | DE only in male placentas |
| ENSRNOG00000060278 | | CPG1 | -1.50 | 2.12E-02 | DE only in male placentas |
| ENSRNOG00000012183 | | GLRX | -1.16 | 2.46E-03 | DE only in male placentas |
| ENSRNOG00000039336 | | HRCT1 | -1.10 | 4.48E-03 | DE only in male placentas |
| ENSRNOG00000017149 | | FAM131B | -1.05 | 9.67E-03 | DE only in male placentas |
| ENSRNOG00000009172 | | CPVL | -1.60 | 3.76E-02 | DE only in male placentas |
| ENSRNOG00000026585 | | LRRC66 | -1.15 | 3.84E-02 | DE only in male placentas |
| ENSRNOG00000059876 | | FSD1L | -1.15 | 9.90E-06 | DE only in male placentas |
| ENSRNOG00000023299 | | HFM1 | -2.28 | 4.01E-02 | DE only in male placentas |
| ENSRNOG00000038132 | X | VSIG4 | -2.06 | 4.68E-03 | DE only in male placentas |
| ENSRNOG00000007951 | | GALNT14 | -1.05 | 6.12E-03 | DE only in male placentas |

| | | | | | |
|---|---|---|---|---|---|
| ENSRNOG00000018094 | | SV2C | -1.30 | 1.95E-03 | DE only in male placentas |
| ENSRNOG00000023686 | | UPK3B | -1.92 | 7.90E-03 | DE only in male placentas |
| ENSRNOG00000061026 | | NPY4R | -1.05 | 3.98E-03 | DE only in male placentas |
| ENSRNOG00000024245 | | ZKSCAN2 | -1.40 | 3.45E-02 | DE only in male placentas |
| ENSRNOG00000042978 | | NCALD | -1.38 | 6.16E-08 | DE only in male placentas |
| ENSRNOG00000032375 | | CAPN13 | -1.86 | 6.60E-05 | DE only in male placentas |
| ENSRNOG00000003705 | X | BMX | -1.77 | 1.06E-04 | DE only in male placentas |
| ENSRNOG00000011548 | | EBF2 | -1.09 | 3.24E-04 | DE only in male placentas |
| ENSRNOG00000057022 | | AABR070637 40.2 | -1.90 | 4.06E-03 | DE only in male placentas |
| ENSRNOG00000054246 | | LOC679711 | -1.12 | 4.15E-02 | DE only in male placentas |
| ENSRNOG00000056621 | | AC099384.2 | -1.11 | 8.76E-04 | DE only in male placentas |
| ENSRNOG00000001656 | | KCNJ15 | -1.08 | 5.51E-10 | DE only in male placentas |
| ENSRNOG00000008930 | | TNFSF15 | -1.65 | 3.26E-05 | DE only in male placentas |
| ENSRNOG00000007573 | | HOXB9 | -1.89 | 2.13E-02 | DE only in male placentas |
| ENSRNOG00000031790 | | H2BU1 | -1.00 | 5.06E-04 | DE only in male placentas |
| ENSRNOG00000043445 | | THEM5 | -2.34 | 3.91E-04 | DE only in male placentas |
| ENSRNOG00000056793 | | HSF5 | -1.84 | 5.77E-05 | DE only in male placentas |
| ENSRNOG00000049693 | | LOC1083481 30 | -3.11 | 1.58E-02 | DE only in male placentas |

| | | | | | |
|---|---|---|---|---|---|
| ENSRNOG00000056507 | | AABR070646 13.2 | -1.13 | 9.76E-05 | DE only in male placentas |
| ENSRNOG00000003927 | | CD55 | -1.03 | 4.37E-08 | DE only in male placentas |
| ENSRNOG00000007927 | | METTL7B | -1.69 | 3.59E-08 | DE only in male placentas |
| ENSRNOG00000054742 | | AABR070661 88.1 | -3.61 | 2.38E-12 | DE only in male placentas |
| ENSRNOG00000016043 | | AQP4 | -3.23 | 9.92E-05 | DE only in male placentas |
| ENSRNOG00000059203 | | AC133265.2 | -2.91 | 1.37E-02 | DE only in male placentas |
| ENSRNOG00000010545 | | MRAP2 | -1.99 | 1.05E-03 | DE only in male placentas |
| ENSRNOG00000021384 | | ANKRD44 | -1.21 | 3.23E-09 | DE only in male placentas |
| ENSRNOG00000054322 | | AABR070250 51.3 | -1.81 | 3.64E-02 | DE only in male placentas |
| ENSRNOG00000042971 | | LOC363337 | -1.51 | 3.98E-03 | DE only in male placentas |
| ENSRNOG00000058193 | | SLC27A6 | -1.33 | 9.37E-03 | DE only in male placentas |
| ENSRNOG00000038832 | | AABR070644 15.1 | -1.40 | 2.49E-03 | DE only in male placentas |
| ENSRNOG00000017258 | | ICA1L | -1.15 | 2.10E-02 | DE only in male placentas |
| ENSRNOG00000059363 | X | AC130862.3 | -2.70 | 2.47E-02 | DE only in male placentas |
| ENSRNOG00000010819 | | HSPA4L | -1.37 | 1.12E-05 | DE only in male placentas |
| ENSRNOG00000006365 | | ASB15 | -1.80 | 8.60E-04 | DE only in male placentas |
| ENSRNOG00000003949 | X | AABR070379 95.1 | -1.15 | 1.01E-12 | DE only in male placentas |
| ENSRNOG00000017223 | | PLG | -1.02 | 2.54E-02 | DE only in male placentas |

| | | | | | |
|---|---|---|---|---|---|
| ENSRNOG00000001806 | | FETUB | -1.78 | 4.46E-03 | DE only in male placentas |
| ENSRNOG00000010812 | | OSBPL6 | -1.34 | 1.16E-09 | DE only in male placentas |
| ENSRNOG00000017244 | | RNF186 | -1.28 | 3.09E-02 | DE only in male placentas |
| ENSRNOG00000013684 | | F11 | -1.60 | 1.57E-03 | DE only in male placentas |
| ENSRNOG00000005573 | | NTN4 | -1.09 | 1.20E-07 | DE only in male placentas |
| ENSRNOG00000005687 | X | SLC6A14 | -1.29 | 1.09E-05 | DE only in male placentas |
| ENSRNOG00000040166 | | ANKRD34B | -1.78 | 4.95E-07 | DE only in male placentas |
| ENSRNOG00000003872 | | NEWGENE_ 620180 | -1.02 | 5.69E-12 | DE only in male placentas |
| ENSRNOG00000052188 | | AABR070070 26.1 | -2.07 | 1.02E-08 | DE only in male placentas |
| ENSRNOG00000003134 | | SLC4A4 | -1.41 | 2.59E-10 | DE only in male placentas |
| ENSRNOG00000025360 | | EXD1 | -1.00 | 9.78E-05 | DE only in male placentas |
| ENSRNOG00000017917 | | CDHR2 | -1.03 | 1.09E-03 | DE only in male placentas |
| ENSRNOG00000056413 | | SPATA31D3 | -1.35 | 3.95E-02 | DE only in male placentas |
| ENSRNOG00000027770 | | TRPM3 | -2.26 | 4.33E-05 | DE only in male placentas |
| ENSRNOG00000022139 | | SERPINB9D | -1.02 | 2.24E-06 | DE only in male placentas |
| ENSRNOG00000015408 | | VIL1 | -1.07 | 3.21E-05 | DE only in male placentas |
| ENSRNOG00000025587 | | PLAGL1 | -1.08 | 6.60E-11 | DE only in male placentas |
| ENSRNOG00000049874 | | P3R3URF | -1.13 | 1.84E-03 | DE only in male placentas |

| | | | | |
|---|---|---|---|---|
| ENSRNOG00000036960 | ABCC9 | -1.03 | 1.66E-05 | DE only in male placentas |
| ENSRNOG00000042944 | CENPW | -1.05 | 1.09E-03 | DE only in male placentas |
| ENSRNOG00000031136 | NTNG1 | -2.02 | 6.58E-03 | DE only in male placentas |
| ENSRNOG00000004703 | MUC15 | -1.22 | 2.15E-04 | DE only in male placentas |
| ENSRNOG00000057687 | AC141334.3 | -2.66 | 8.35E-03 | DE only in male placentas |
| ENSRNOG00000038572 | NCAPG | -1.04 | 5.60E-06 | DE only in male placentas |
| ENSRNOG00000025522 | ATF7IP2 | -1.20 | 7.37E-08 | DE only in male placentas |
| ENSRNOG00000030547 | DPP6 | -1.48 | 2.93E-03 | DE only in male placentas |
| ENSRNOG00000030149 | ADGRL3 | -1.11 | 1.36E-06 | DE only in male placentas |
| ENSRNOG00000061085 | FANCD2 | -1.14 | 2.73E-07 | DE only in male placentas |
| ENSRNOG00000052150 | LOC1025499 48 | -1.67 | 8.63E-06 | DE only in male placentas |
| ENSRNOG00000029735 | PID1 | -1.21 | 7.87E-11 | DE only in male placentas |
| ENSRNOG00000014686 | KCND3 | -1.18 | 5.04E-15 | DE only in male placentas |
| ENSRNOG00000016783 | FSHR | -1.55 | 2.41E-02 | DE only in male placentas |
| ENSRNOG00000026958 | CYREN | -1.07 | 4.05E-09 | DE only in male placentas |
| ENSRNOG00000015232 | PIP5K1B | -1.18 | 6.31E-11 | DE only in male placentas |
| ENSRNOG00000045764 | CXXC4 | -1.20 | 1.68E-02 | DE only in male placentas |
| ENSRNOG00000021017 | CA11 | -1.15 | 3.76E-07 | DE only in male placentas |

| | | | | | |
|---|---|---|---|---|---|
| ENSRNOG00000060847 | | KRTAP2-1 | -1.37 | 2.13E-02 | DE only in male placentas |
| ENSRNOG00000053113 | | AABR070013 89.1 | -1.14 | 3.47E-06 | DE only in male placentas |
| ENSRNOG00000017702 | | GPLD1 | -1.31 | 7.21E-10 | DE only in male placentas |
| ENSRNOG00000007782 | | GHRH | -1.02 | 5.69E-09 | DE only in male placentas |
| ENSRNOG00000023861 | | SNAP91 | -1.31 | 4.31E-08 | DE only in male placentas |
| ENSRNOG00000004200 | | SYBU | -1.17 | 1.66E-02 | DE only in male placentas |
| ENSRNOG00000028541 | | BTNL2 | -1.49 | 2.45E-02 | DE only in male placentas |
| ENSRNOG00000051958 | X | AABR070379 43.1 | -1.05 | 4.08E-08 | DE only in male placentas |
| ENSRNOG00000006845 | | PSAPL1 | -2.32 | 3.06E-02 | DE only in male placentas |
| ENSRNOG00000001173 | | CABP1 | -1.62 | 6.41E-03 | DE only in male placentas |
| ENSRNOG00000003878 | | THSD7B | -2.26 | 8.78E-03 | DE only in male placentas |
| ENSRNOG00000049452 | | NEBL | -1.33 | 1.80E-04 | DE only in male placentas |
| ENSRNOG00000059055 | | HSBP1L1 | -1.14 | 2.92E-03 | DE only in male placentas |
| ENSRNOG00000006670 | X | RAI2 | -1.04 | 9.05E-04 | DE only in male placentas |
| ENSRNOG00000029244 | | PCDHB10 | -2.60 | 4.84E-05 | DE only in male placentas |
| ENSRNOG00000053026 | | SHCBP1 | -1.02 | 1.33E-04 | DE only in male placentas |
| ENSRNOG00000002396 | | SERPINB8 | -1.23 | 3.17E-04 | DE only in male placentas |
| ENSRNOG00000022116 | | GJB6 | -1.39 | 1.34E-06 | DE only in male placentas |

| | | | | | |
|---|---|---|---|---|---|
| ENSRNOG00000046053 | X | NUDT10 | -1.38 | 3.97E-05 | DE only in male placentas |
| ENSRNOG00000011861 | | AADAT | -1.27 | 1.38E-03 | DE only in male placentas |
| ENSRNOG00000046547 | | RBM24 | -1.21 | 3.82E-03 | DE only in male placentas |
| ENSRNOG00000049696 | | RSPH4A | -1.25 | 8.03E-03 | DE only in male placentas |
| ENSRNOG00000057433 | | AABR070275 81.1 | -1.50 | 1.05E-04 | DE only in male placentas |
| ENSRNOG00000027522 | | SLC25A54 | -2.49 | 2.23E-02 | DE only in male placentas |
| ENSRNOG00000031612 | | GLS2 | -1.22 | 4.89E-11 | DE only in male placentas |
| ENSRNOG00000009153 | | CIDEC | -1.25 | 8.52E-03 | DE only in male placentas |
| ENSRNOG00000027341 | | TENM2 | -1.20 | 2.39E-03 | DE only in male placentas |
| ENSRNOG00000005076 | X | BCORL1 | -1.25 | 1.31E-08 | DE only in male placentas |
| ENSRNOG00000043304 | | APCDD1 | -1.34 | 2.00E-04 | DE only in male placentas |
| ENSRNOG00000021984 | | RGS7 | -1.86 | 8.35E-03 | DE only in male placentas |
| ENSRNOG00000046714 | | LOC301165 | -3.26 | 1.33E-02 | DE only in male placentas |
| ENSRNOG00000039110 | | LSMEM1 | -1.27 | 4.26E-05 | DE only in male placentas |
| ENSRNOG00000003950 | | LVRN | -1.13 | 2.83E-02 | DE only in male placentas |
| ENSRNOG00000009431 | | TBC1D4 | -1.15 | 1.62E-11 | DE only in male placentas |
| ENSRNOG00000006873 | | DNASE1 | -1.72 | 1.04E-02 | DE only in male placentas |
| ENSRNOG00000003472 | X | ATP11C | -1.03 | 3.17E-15 | DE only in male placentas |

| | | | | | |
|---|---|---|---|---|---|
| ENSRNOG00000002537 | X | WNK3 | -1.40 | 4.10E-14 | DE only in male placentas |
| ENSRNOG00000013262 | | MYL1 | -2.18 | 2.17E-02 | DE only in male placentas |
| ENSRNOG00000015303 | | FBLL1 | -1.75 | 2.44E-02 | DE only in male placentas |
| ENSRNOG00000012512 | | NEXN | -1.08 | 2.96E-25 | DE only in male placentas |
| ENSRNOG00000026415 | | COL14A1 | -1.31 | 1.32E-06 | DE only in male placentas |
| ENSRNOG00000005176 | X | MAP7D2 | -1.39 | 3.81E-07 | DE only in male placentas |
| ENSRNOG00000004101 | | 44256 | -1.37 | 5.38E-05 | DE only in male placentas |
| ENSRNOG00000014975 | | ZFP483 | -1.27 | 1.24E-06 | DE only in male placentas |
| ENSRNOG00000000958 | | AABR070559 43.1 | -1.22 | 4.88E-02 | DE only in male placentas |
| ENSRNOG00000004171 | | DNAH9 | -1.27 | 1.75E-06 | DE only in male placentas |
| ENSRNOG00000020203 | | PTPN20 | -1.69 | 2.86E-02 | DE only in male placentas |
| ENSRNOG00000021091 | | TRANK1 | -1.00 | 5.67E-21 | DE only in male placentas |
| ENSRNOG00000004725 | | CELA1 | -1.14 | 1.45E-05 | DE only in male placentas |
| ENSRNOG00000023007 | | COL6A6 | -1.03 | 1.52E-02 | DE only in male placentas |
| ENSRNOG00000005561 | | BRINP1 | -1.88 | 3.37E-06 | DE only in male placentas |
| ENSRNOG00000019704 | | RESP18 | -1.35 | 1.35E-04 | DE only in male placentas |
| ENSRNOG00000024115 | | C6 | -1.06 | 4.65E-04 | DE only in male placentas |
| ENSRNOG00000012375 | | INPP1 | -1.11 | 6.24E-10 | DE only in male placentas |

| | | | | | |
|---|---|---|---|---|---|
| ENSRNOG00000002848 | X | MAOA | -1.23 | 1.99E-10 | DE only in male placentas |
| ENSRNOG00000040195 | | RGD1559960 | -1.77 | 4.20E-04 | DE only in male placentas |
| ENSRNOG00000039469 | | PCDHB19 | -1.42 | 2.40E-06 | DE only in male placentas |
| ENSRNOG00000018627 | | PLEKHB1 | -1.25 | 3.19E-10 | DE only in male placentas |
| ENSRNOG00000033680 | | CYP2B1 | -2.31 | 5.13E-05 | DE only in male placentas |
| ENSRNOG00000020443 | | TSKS | -1.35 | 2.54E-02 | DE only in male placentas |
| ENSRNOG00000010015 | | GP9 | -1.20 | 7.37E-06 | DE only in male placentas |
| ENSRNOG00000016046 | | HECW1 | -1.44 | 2.22E-03 | DE only in male placentas |
| ENSRNOG00000017672 | | AKR1C14 | -1.14 | 2.06E-21 | DE only in male placentas |
| ENSRNOG00000043128 | X | ARMCX4 | -1.00 | 5.30E-11 | DE only in male placentas |
| ENSRNOG00000012878 | | ATP7B | -1.04 | 2.15E-05 | DE only in male placentas |
| ENSRNOG00000025428 | X | RP2 | -1.04 | 3.77E-06 | DE only in male placentas |
| ENSRNOG00000058157 | | LDLRAD3 | -1.18 | 1.18E-15 | DE only in male placentas |
| ENSRNOG00000033883 | X | STARD8 | -1.20 | 2.43E-10 | DE only in male placentas |
| ENSRNOG00000021029 | | HAMP | -1.46 | 4.16E-03 | DE only in male placentas |
| ENSRNOG00000049873 | | AP1S3 | -1.05 | 2.62E-11 | DE only in male placentas |
| ENSRNOG00000026430 | | TRIM58 | -1.38 | 1.42E-03 | DE only in male placentas |
| ENSRNOG00000058891 | | CYS1 | -1.00 | 4.60E-03 | DE only in male placentas |

| | | | | | |
|---|---|---|---|---|---|
| ENSRNOG00000055714 | | AK7 | -1.05 | 1.83E-05 | DE only in male placentas |
| ENSRNOG00000021433 | | ARHGEF39 | -1.12 | 1.19E-13 | DE only in male placentas |
| ENSRNOG00000001765 | | SLC51A | -1.07 | 4.01E-05 | DE only in male placentas |
| ENSRNOG00000005404 | | RASGRP1 | -1.04 | 3.87E-06 | DE only in male placentas |
| ENSRNOG00000047605 | | PTPRK | -1.01 | 3.05E-12 | DE only in male placentas |
| ENSRNOG00000012719 | | TDRD12 | -1.51 | 1.87E-04 | DE only in male placentas |
| ENSRNOG00000007189 | | TTC22 | -1.07 | 1.90E-02 | DE only in male placentas |
| ENSRNOG00000014648 | | EFNB2 | -1.10 | 6.95E-19 | DE only in male placentas |
| ENSRNOG00000060687 | | SLC24A3 | -1.26 | 3.15E-11 | DE only in male placentas |
| ENSRNOG00000046858 | X | ARXES2 | -1.68 | 3.70E-04 | DE only in male placentas |
| ENSRNOG00000017528 | | GPR157 | -1.00 | 7.63E-10 | DE only in male placentas |
| ENSRNOG00000023162 | | CA14 | -1.10 | 5.17E-06 | DE only in male placentas |
| ENSRNOG00000017105 | | DPYD | -1.54 | 1.86E-03 | DE only in male placentas |
| ENSRNOG00000016872 | | PLPPR4 | -1.41 | 7.65E-03 | DE only in male placentas |
| ENSRNOG00000053288 | | ANK3 | -1.08 | 1.73E-13 | DE only in male placentas |
| ENSRNOG00000006729 | | SLC24A4 | -1.46 | 6.09E-07 | DE only in male placentas |
| ENSRNOG00000003785 | | USP43 | -1.15 | 6.42E-06 | DE only in male placentas |
| ENSRNOG00000047466 | | BDNF | -1.50 | 7.08E-09 | DE only in male placentas |

| | | | | | |
|---|---|---|---|---|---|
| ENSRNOG00000023312 | | LONRF2 | -1.40 | 1.63E-10 | DE only in male placentas |
| ENSRNOG00000049695 | | MYH4 | -1.78 | 1.16E-11 | DE only in male placentas |
| ENSRNOG00000047204 | | GP6 | -1.22 | 6.07E-06 | DE only in male placentas |
| ENSRNOG00000002862 | X | CLCN5 | -1.04 | 2.22E-15 | DE only in male placentas |
| ENSRNOG00000010655 | | MTTP | -1.39 | 1.53E-02 | DE only in male placentas |
| ENSRNOG00000027089 | | ELL2 | -1.10 | 5.89E-19 | DE only in male placentas |
| ENSRNOG00000008758 | | TSPAN18 | -1.53 | 2.89E-13 | DE only in male placentas |
| ENSRNOG00000033479 | | PCDHB21 | -1.95 | 7.34E-06 | DE only in male placentas |
| ENSRNOG00000028526 | | MANSC1 | -1.23 | 5.88E-05 | DE only in male placentas |
| ENSRNOG00000009145 | | KLF12 | -1.33 | 6.45E-12 | DE only in male placentas |
| ENSRNOG00000056945 | | LOC102549408 | -1.30 | 1.14E-03 | DE only in male placentas |
| ENSRNOG00000010780 | | DLC1 | -1.09 | 1.35E-19 | DE only in male placentas |
| ENSRNOG00000010748 | | MTUS1 | -1.00 | 1.17E-17 | DE only in male placentas |
| ENSRNOG00000000775 | | MOG | -1.05 | 1.46E-07 | DE only in male placentas |
| ENSRNOG00000039906 | | ABO3 | -1.05 | 1.78E-02 | DE only in male placentas |
| ENSRNOG00000054379 | | LOC108348134 | -3.88 | 1.42E-02 | DE only in male placentas |
| ENSRNOG00000010463 | | CPA4 | -1.00 | 5.20E-06 | DE only in male placentas |
| ENSRNOG00000015577 | | LPAR6 | -1.07 | 3.16E-08 | DE only in male placentas |

| | | | | | |
|---|---|---|---|---|---|
| ENSRNOG00000033064 | | EDIL3 | -1.37 | 8.89E-11 | DE only in male placentas |
| ENSRNOG00000023548 | | SNED1 | -1.00 | 5.85E-16 | DE only in male placentas |
| ENSRNOG00000007172 | | MAP4K3 | -1.01 | 3.08E-31 | DE only in male placentas |
| ENSRNOG00000047503 | | APOC3 | -1.32 | 6.42E-07 | DE only in male placentas |
| ENSRNOG00000015357 | | BMPER | -1.07 | 4.19E-06 | DE only in male placentas |
| ENSRNOG00000037982 | | TMEM236 | -2.40 | 1.03E-02 | DE only in male placentas |
| ENSRNOG00000022066 | | FAM117B | -1.13 | 7.56E-31 | DE only in male placentas |
| ENSRNOG00000039470 | | AC131360.1 | -1.05 | 7.56E-03 | DE only in male placentas |
| ENSRNOG00000016057 | | KCNJ13 | -1.06 | 3.19E-05 | DE only in male placentas |
| ENSRNOG00000027520 | X | PLS3 | -1.00 | 1.12E-18 | DE only in male placentas |
| ENSRNOG00000051534 | | PRR33 | -1.29 | 3.16E-02 | DE only in male placentas |
| ENSRNOG00000015637 | | LOC690507 | -1.89 | 2.80E-03 | DE only in male placentas |
| ENSRNOG00000027225 | | LY6G6F | -1.09 | 7.93E-04 | DE only in male placentas |
| ENSRNOG00000052469 | | BMPR1A | -1.02 | 1.62E-23 | DE only in male placentas |
| ENSRNOG00000049880 | | TMEM200C | -1.67 | 4.18E-05 | DE only in male placentas |
| ENSRNOG00000004222 | X | S100G | -1.05 | 7.58E-07 | DE only in male placentas |
| ENSRNOG00000018525 | | SLC4A9 | -2.04 | 6.70E-05 | DE only in male placentas |
| ENSRNOG00000001153 | | PLA2G1B | -1.81 | 1.83E-03 | DE only in male placentas |

| | | | | | |
|---|---|---|---|---|---|
| ENSRNOG00000013148 | | RAD54L | -1.43 | 4.46E-16 | DE only in male placentas |
| ENSRNOG00000007377 | | SLIT3 | -1.03 | 2.84E-14 | DE only in male placentas |
| ENSRNOG00000002459 | | FBXO40 | -2.21 | 2.23E-07 | DE only in male placentas |
| ENSRNOG00000009079 | | PRKAR2B | -1.67 | 8.28E-13 | DE only in male placentas |
| ENSRNOG00000007081 | | XDH | -1.00 | 8.72E-12 | DE only in male placentas |
| ENSRNOG00000003088 | | ARHGAP31 | -1.12 | 6.82E-18 | DE only in male placentas |
| ENSRNOG00000055518 | | AABR070243 03.1 | -1.88 | 9.23E-05 | DE only in male placentas |
| ENSRNOG00000004311 | | GPR182 | -1.38 | 3.15E-05 | DE only in male placentas |
| ENSRNOG00000051346 | X | AABR070388 49.2 | -1.06 | 1.59E-04 | DE only in male placentas |
| ENSRNOG00000026265 | X | MAGEC2 | -1.11 | 2.27E-15 | DE only in male placentas |
| ENSRNOG00000007496 | | MTMR4 | -1.05 | 1.07E-18 | DE only in male placentas |
| ENSRNOG00000003893 | | CCNJL | -1.02 | 1.74E-11 | DE only in male placentas |
| ENSRNOG00000002425 | | PCTP | -1.08 | 1.24E-10 | DE only in male placentas |
| ENSRNOG00000003587 | X | VEGFD | -1.01 | 4.37E-09 | DE only in male placentas |
| ENSRNOG00000014361 | | EDN1 | -1.25 | 1.69E-06 | DE only in male placentas |
| ENSRNOG00000008203 | | SYNPR | -1.33 | 7.90E-04 | DE only in male placentas |
| ENSRNOG00000015800 | | ELMO3 | -1.15 | 7.34E-09 | DE only in male placentas |
| ENSRNOG00000019403 | | AFAP1L1 | -1.07 | 1.14E-17 | DE only in male placentas |

| | | | | | |
|---|---|---|---|---|---|
| ENSRNOG00000046764 | | TMPRSS11A | -2.63 | 6.58E-04 | DE only in male placentas |
| ENSRNOG00000057644 | X | LOC1036944<br>88 | -1.38 | 1.13E-07 | DE only in male placentas |
| ENSRNOG00000000068 | | PPP2R5A | -1.18 | 8.71E-18 | DE only in male placentas |
| ENSRNOG00000006368 | | LRRN3 | -2.16 | 6.06E-07 | DE only in male placentas |
| ENSRNOG00000053996 | | AABR070444<br>92.1 | -2.38 | 4.13E-02 | DE only in male placentas |
| ENSRNOG00000040301 | | GALNTL5 | -1.25 | 3.31E-02 | DE only in male placentas |
| ENSRNOG00000006709 | | PZP | -1.37 | 7.75E-03 | DE only in male placentas |
| ENSRNOG00000000875 | X | FHL1 | -1.13 | 1.74E-12 | DE only in male placentas |
| ENSRNOG00000038215 | | AC120448.1 | -1.20 | 2.94E-02 | DE only in male placentas |
| ENSRNOG00000004150 | X | SLC9A7 | -1.07 | 5.76E-09 | DE only in male placentas |
| ENSRNOG00000023458 | X | DCAF12L1 | -2.06 | 7.44E-06 | DE only in male placentas |
| ENSRNOG00000013340 | | TWSG1 | -1.06 | 2.44E-23 | DE only in male placentas |
| ENSRNOG00000010625 | | DNMT3B | -1.25 | 7.60E-17 | DE only in male placentas |
| ENSRNOG00000000096 | | PDZK1 | -1.00 | 8.51E-06 | DE only in male placentas |
| ENSRNOG00000019550 | | SLC11A2 | -1.12 | 1.21E-09 | DE only in male placentas |
| ENSRNOG00000007139 | | TTPA | -2.34 | 2.74E-04 | DE only in male placentas |
| ENSRNOG00000029543 | | CISH | -1.06 | 3.52E-07 | DE only in male placentas |
| ENSRNOG00000033740 | | LURAP1L | -1.13 | 1.12E-08 | DE only in male placentas |

| | | | | | |
|---|---|---|---|---|---|
| ENSRNOG00000015503 | | OTUD7A | -1.37 | 9.79E-04 | DE only in male placentas |
| ENSRNOG00000006570 | | PLEKHG3 | -1.08 | 1.71E-17 | DE only in male placentas |
| ENSRNOG00000012303 | | APOBEC2 | -1.95 | 7.25E-06 | DE only in male placentas |
| ENSRNOG00000060568 | | PTPN21 | -1.07 | 7.47E-19 | DE only in male placentas |
| ENSRNOG00000003386 | | RBFOX3 | -1.30 | 2.35E-04 | DE only in male placentas |
| ENSRNOG00000001152 | | GLP1R | -1.67 | 5.60E-04 | DE only in male placentas |
| ENSRNOG00000002583 | | SERPINB11 | -1.41 | 4.70E-05 | DE only in male placentas |
| ENSRNOG00000003566 | | APOH | -1.68 | 2.49E-02 | DE only in male placentas |
| ENSRNOG00000008443 | | FAM234B | -1.20 | 1.08E-13 | DE only in male placentas |
| ENSRNOG00000037156 | | MRGPRX1 | -1.68 | 3.83E-04 | DE only in male placentas |
| ENSRNOG00000036659 | | FN3K | -2.38 | 2.10E-05 | DE only in male placentas |
| ENSRNOG00000005667 | | ASTN1 | -1.56 | 3.62E-06 | DE only in male placentas |
| ENSRNOG00000009117 | | OTUB2 | -1.09 | 9.38E-19 | DE only in male placentas |
| ENSRNOG00000061122 | | LOC1025516<br>83 | -1.64 | 1.08E-02 | DE only in male placentas |
| ENSRNOG00000053086 | | SELENOP | -1.41 | 2.56E-09 | DE only in male placentas |
| ENSRNOG00000003370 | X | OTC | -1.24 | 1.78E-03 | DE only in male placentas |
| ENSRNOG00000012663 | | MFSD6 | -1.71 | 9.47E-09 | DE only in male placentas |
| ENSRNOG00000060063 | X | NAA10 | -1.05 | 8.21E-16 | DE only in male placentas |

| | | | | | |
|---|---|---|---|---|---|
| ENSRNOG00000011654 | | PLK4 | -1.24 | 1.41E-16 | DE only in male placentas |
| ENSRNOG00000000172 | | SQOR | -1.08 | 3.19E-10 | DE only in male placentas |
| ENSRNOG00000008519 | | DIPK2A | -1.27 | 6.67E-16 | DE only in male placentas |
| ENSRNOG00000030889 | | CLDN22 | -1.39 | 4.34E-03 | DE only in male placentas |
| ENSRNOG00000046366 | X | DMD | -1.04 | 1.94E-12 | DE only in male placentas |
| ENSRNOG00000017054 | | KCNE3 | -1.65 | 2.66E-08 | DE only in male placentas |
| ENSRNOG00000017421 | | ALPK2 | -1.34 | 3.62E-05 | DE only in male placentas |
| ENSRNOG00000001827 | | MASP1 | -1.23 | 1.09E-12 | DE only in male placentas |
| ENSRNOG00000002106 | | USP46 | -1.27 | 5.76E-16 | DE only in male placentas |
| ENSRNOG00000000936 | | STX2 | -1.18 | 4.78E-15 | DE only in male placentas |
| ENSRNOG00000011533 | | IRX3 | -1.12 | 1.53E-15 | DE only in male placentas |
| ENSRNOG00000050910 | X | LOC688583 | -1.09 | 1.49E-05 | DE only in male placentas |
| ENSRNOG00000032884 | | SCN11A | -2.24 | 8.21E-12 | DE only in male placentas |
| ENSRNOG00000005758 | | BTBD11 | -1.09 | 1.15E-05 | DE only in male placentas |
| ENSRNOG00000032798 | | SLCO3A1 | -1.03 | 1.41E-06 | DE only in male placentas |
| ENSRNOG00000027787 | | CDC6 | -1.15 | 2.27E-09 | DE only in male placentas |
| ENSRNOG00000007324 | | PLXNA2 | -1.06 | 3.79E-11 | DE only in male placentas |
| ENSRNOG00000056575 | | AABR070300<br>86.2 | -1.48 | 2.49E-04 | DE only in male placentas |

| | | | | |
|---|---|---|---|---|
| ENSRNOG00000009014 | SLC35F2 | -1.19 | 4.19E-13 | DE only in male placentas |
| ENSRNOG00000012095 | PKIA | -1.01 | 5.01E-06 | DE only in male placentas |
| ENSRNOG00000018382 | INPP4B | -1.16 | 3.13E-16 | DE only in male placentas |
| ENSRNOG00000037211 | KIF14 | -1.55 | 1.40E-06 | DE only in male placentas |
| ENSRNOG00000007104 | ITPR1 | -1.03 | 1.91E-13 | DE only in male placentas |
| ENSRNOG00000033226 | AABR070435<br>57.1 | -1.62 | 4.78E-02 | DE only in male placentas |
| ENSRNOG00000019662 | TM6SF1 | -1.03 | 2.31E-25 | DE only in male placentas |
| ENSRNOG00000011521 | FILIP1 | -1.08 | 6.79E-19 | DE only in male placentas |
| ENSRNOG00000015812 | TM4SF1 | -1.32 | 4.21E-08 | DE only in male placentas |
| ENSRNOG00000048043 | F2R | -1.10 | 1.79E-19 | DE only in male placentas |
| ENSRNOG00000010152 | RGD1306233 | -1.44 | 4.58E-03 | DE only in male placentas |
| ENSRNOG00000004328 | DEPTOR | -1.15 | 5.88E-10 | DE only in male placentas |
| ENSRNOG00000004629 | FKBP3 | -1.09 | 2.34E-21 | DE only in male placentas |
| ENSRNOG00000018238 | NUGGC | -1.24 | 7.66E-07 | DE only in male placentas |
| ENSRNOG00000013097 | LOC691485 | -1.79 | 1.69E-03 | DE only in male placentas |
| ENSRNOG00000013048 | PDE7A | -1.38 | 1.98E-19 | DE only in male placentas |
| ENSRNOG00000009581 | LCE1M | -2.20 | 1.04E-02 | DE only in male placentas |
| ENSRNOG00000016321 | ENTPD4 | -1.10 | 1.51E-32 | DE only in male placentas |

| | | | | | |
|---|---|---|---|---|---|
| ENSRNOG00000038877 | | BTN2A2 | -1.19 | 3.95E-10 | DE only in male placentas |
| ENSRNOG00000008638 | | ANGPTL3 | -1.60 | 9.50E-03 | DE only in male placentas |
| ENSRNOG00000005608 | | TEAD4 | -1.25 | 6.35E-23 | DE only in male placentas |
| ENSRNOG00000005615 | | GADD45A | -1.08 | 3.02E-24 | DE only in male placentas |
| ENSRNOG00000010344 | | B4GALNT3 | -1.08 | 2.15E-12 | DE only in male placentas |
| ENSRNOG00000042751 | X | AABR070385 56.1 | -1.18 | 1.94E-02 | DE only in male placentas |
| ENSRNOG00000043154 | | PRR19 | -1.08 | 1.49E-02 | DE only in male placentas |
| ENSRNOG00000004120 | X | RGD1565844 | -1.81 | 1.31E-02 | DE only in male placentas |
| ENSRNOG00000033706 | | KLK14 | -1.47 | 8.98E-04 | DE only in male placentas |
| ENSRNOG00000002829 | | PPBP | -1.33 | 5.50E-03 | DE only in male placentas |
| ENSRNOG00000010031 | | VTN | -1.39 | 1.59E-06 | DE only in male placentas |
| ENSRNOG00000015027 | | IL18R1 | -1.33 | 1.40E-06 | DE only in male placentas |
| ENSRNOG00000057598 | | AABR070325 20.4 | -2.96 | 2.12E-03 | DE only in male placentas |
| ENSRNOG00000048973 | | NEWGENE_ 1306399 | -1.27 | 5.98E-23 | DE only in male placentas |
| ENSRNOG00000014297 | | SDC4 | -1.10 | 1.69E-17 | DE only in male placentas |
| ENSRNOG00000016502 | | DSCAML1 | -1.16 | 3.16E-02 | DE only in male placentas |
| ENSRNOG00000008237 | | UNC13B | -1.08 | 3.86E-16 | DE only in male placentas |
| ENSRNOG00000046245 | | AABR070574 23.1 | -1.03 | 1.40E-08 | DE only in male placentas |

| | | | | |
|---|---|---|---|---|
| ENSRNOG00000003407 | PTPN14 | -1.11 | 3.50E-18 | DE only in male placentas |
| ENSRNOG00000012546 | FRMPD1 | -1.29 | 1.68E-10 | DE only in male placentas |
| ENSRNOG00000000634 | CABCOCO1 | -1.48 | 9.34E-06 | DE only in male placentas |
| ENSRNOG00000016724 | EPB41L3 | -1.49 | 1.55E-16 | DE only in male placentas |
| ENSRNOG00000024433 | FBXL7 | -1.37 | 6.44E-14 | DE only in male placentas |
| ENSRNOG00000013391 | SORBS2 | -1.11 | 5.14E-17 | DE only in male placentas |
| ENSRNOG00000061429 | RPH3AL | -1.31 | 1.77E-08 | DE only in male placentas |
| ENSRNOG00000006076 | STEAP2 | -1.17 | 1.37E-19 | DE only in male placentas |
| ENSRNOG00000011815 | SGK1 | -1.04 | 3.16E-18 | DE only in male placentas |
| ENSRNOG00000038297 | PLEKHD1 | -2.25 | 2.09E-04 | DE only in male placentas |
| ENSRNOG00000002919 | GFAP | -1.05 | 3.20E-04 | DE only in male placentas |
| ENSRNOG00000023538 | ALDH5A1 | -1.61 | 1.36E-13 | DE only in male placentas |
| ENSRNOG00000013541 | SH2D4A | -1.46 | 1.21E-12 | DE only in male placentas |
| ENSRNOG00000008615 | MAL2 | -1.99 | 8.93E-18 | DE only in male placentas |
| ENSRNOG00000012019 | B3GALNT1 | -1.43 | 3.12E-16 | DE only in male placentas |
| ENSRNOG00000014135 | RAB11FIP4 | -1.15 | 8.47E-12 | DE only in male placentas |
| ENSRNOG00000042111 | SULT1C2A | -2.10 | 1.12E-07 | DE only in male placentas |
| ENSRNOG00000001404 | AGFG2 | -1.54 | 2.31E-08 | DE only in male placentas |

| | | | | | |
|---|---|---|---|---|---|
| ENSRNOG00000016371 | | SLC18B1 | -1.01 | 2.22E-15 | DE only in male placentas |
| ENSRNOG00000007817 | | KCTD6 | -1.11 | 1.22E-17 | DE only in male placentas |
| ENSRNOG00000003468 | | CAPN8 | -1.21 | 1.33E-12 | DE only in male placentas |
| ENSRNOG00000011161 | | SLC2A12 | -1.34 | 3.88E-14 | DE only in male placentas |
| ENSRNOG00000026974 | | DBNDD1 | -1.27 | 8.95E-05 | DE only in male placentas |
| ENSRNOG00000053244 | | SPP2 | -1.46 | 2.36E-02 | DE only in male placentas |
| ENSRNOG00000027730 | | NXPE1 | -1.64 | 7.80E-10 | DE only in male placentas |
| ENSRNOG00000014443 | | PDE5A | -1.16 | 1.58E-16 | DE only in male placentas |
| ENSRNOG00000051704 | X | ZFP185 | -1.40 | 4.53E-14 | DE only in male placentas |
| ENSRNOG00000032048 | | ZFP462 | -1.17 | 4.14E-12 | DE only in male placentas |
| ENSRNOG00000004870 | | DYRK3 | -1.23 | 8.40E-12 | DE only in male placentas |
| ENSRNOG00000043176 | | AABR07059308.1 | -1.45 | 2.48E-09 | DE only in male placentas |
| ENSRNOG00000012942 | | SATB1 | -1.23 | 3.37E-11 | DE only in male placentas |
| ENSRNOG00000048874 | | GCKR | -1.39 | 1.07E-03 | DE only in male placentas |
| ENSRNOG00000002708 | X | PHF8 | -1.39 | 1.80E-17 | DE only in male placentas |
| ENSRNOG00000005094 | | C1QTNF7 | -1.78 | 2.77E-07 | DE only in male placentas |
| ENSRNOG00000018824 | | SLC7A5 | -1.08 | 1.45E-16 | DE only in male placentas |
| ENSRNOG00000021174 | | MACROD1 | -1.44 | 4.79E-12 | DE only in male placentas |

| | | | | |
|---|---|---|---|---|
| ENSRNOG00000007655 | ZFP174 | -1.17 | 3.51E-25 | DE only in male placentas |
| ENSRNOG00000053728 | AABR070067 24.1 | -1.58 | 4.08E-05 | DE only in male placentas |
| ENSRNOG00000021919 | RHOJ | -1.01 | 6.82E-11 | DE only in male placentas |
| ENSRNOG00000009204 | IL17RE | -1.16 | 4.46E-13 | DE only in male placentas |
| ENSRNOG00000024986 | MMRN1 | -1.29 | 5.09E-10 | DE only in male placentas |
| ENSRNOG00000007433 | CYB561 | -1.29 | 1.00E-27 | DE only in male placentas |
| ENSRNOG00000004459 | SDR9C7 | -1.77 | 1.56E-05 | DE only in male placentas |
| ENSRNOG00000028891 | CYP2T1 | -1.31 | 5.74E-07 | DE only in male placentas |
| ENSRNOG00000007748 | TEX15 | -1.51 | 2.74E-09 | DE only in male placentas |
| ENSRNOG00000004626 | SLC34A2 | -1.19 | 1.09E-12 | DE only in male placentas |
| ENSRNOG00000003784 | RILP | -1.10 | 8.43E-03 | DE only in male placentas |
| ENSRNOG00000058940 | RAB11FIP1 | -1.23 | 8.14E-25 | DE only in male placentas |
| ENSRNOG00000060587 | AABR070026 27.1 | -1.39 | 9.62E-19 | DE only in male placentas |
| ENSRNOG00000016451 | CD1D1 | -1.19 | 8.75E-10 | DE only in male placentas |
| ENSRNOG00000006853 | ELAVL2 | -2.28 | 6.98E-09 | DE only in male placentas |
| ENSRNOG00000058120 | TINAG | -1.92 | 1.19E-02 | DE only in male placentas |
| ENSRNOG00000026297 | SERPINA16 | -1.42 | 2.12E-02 | DE only in male placentas |
| ENSRNOG00000008376 | LOC1009095 95 | -1.06 | 6.81E-08 | DE only in male placentas |

| | | | | | |
|---|---|---|---|---|---|
| ENSRNOG00000060675 | | AABR070558<br>01.1 | -1.17 | 2.05E-04 | DE only in male placentas |
| ENSRNOG00000045998 | | SEMA6B | -1.08 | 1.72E-10 | DE only in male placentas |
| ENSRNOG00000011228 | | LAYN | -1.14 | 2.51E-10 | DE only in male placentas |
| ENSRNOG00000011713 | | LOC691113 | -1.06 | 2.63E-12 | DE only in male placentas |
| ENSRNOG00000017842 | | SCNN1G | -1.33 | 1.82E-07 | DE only in male placentas |
| ENSRNOG00000018266 | | SLC23A3 | -1.84 | 1.58E-08 | DE only in male placentas |
| ENSRNOG00000048088 | | MEST | -1.20 | 9.22E-14 | DE only in male placentas |
| ENSRNOG00000015858 | | HYAL1 | -1.29 | 3.36E-09 | DE only in male placentas |
| ENSRNOG00000024237 | | SCEL | -1.06 | 2.09E-07 | DE only in male placentas |
| ENSRNOG00000000001 | | AABR070132<br>55.1 | -1.02 | 1.47E-07 | DE only in male placentas |
| ENSRNOG00000015596 | | EFHD1 | -1.80 | 1.87E-12 | DE only in male placentas |
| ENSRNOG00000012424 | | ADAM23 | -1.29 | 1.35E-09 | DE only in male placentas |
| ENSRNOG00000030954 | | FAT1 | -1.30 | 3.50E-18 | DE only in male placentas |
| ENSRNOG00000048039 | | LOC679149 | -1.10 | 4.75E-08 | DE only in male placentas |
| ENSRNOG00000049519 | | AABR070437<br>20.1 | -3.69 | 2.03E-04 | DE only in male placentas |
| ENSRNOG00000032590 | | TTC28 | -1.10 | 4.31E-16 | DE only in male placentas |
| ENSRNOG00000032729 | X | BEX2 | -1.35 | 3.31E-02 | DE only in male placentas |
| ENSRNOG00000021274 | | FERMT1 | -1.53 | 9.92E-09 | DE only in male placentas |

| | | | | | |
|---|---|---|---|---|---|
| ENSRNOG00000005046 | | TSPAN13 | -1.22 | 2.98E-12 | DE only in male placentas |
| ENSRNOG00000059801 | | LIMA1 | -1.23 | 3.99E-17 | DE only in male placentas |
| ENSRNOG00000015480 | | PRKCZ | -1.16 | 4.70E-13 | DE only in male placentas |
| ENSRNOG00000008151 | | PLSCR4 | -1.57 | 1.66E-12 | DE only in male placentas |
| ENSRNOG00000001726 | | TMEM44 | -1.78 | 1.25E-06 | DE only in male placentas |
| ENSRNOG00000051688 | | SYT15 | -1.91 | 5.13E-05 | DE only in male placentas |
| ENSRNOG00000022910 | | EMCN | -1.41 | 2.93E-07 | DE only in male placentas |
| ENSRNOG00000028627 | | HMCN1 | -1.77 | 7.14E-16 | DE only in male placentas |
| ENSRNOG00000005810 | | NLRC4 | -1.41 | 2.09E-10 | DE only in male placentas |
| ENSRNOG00000060054 | | DOC2B | -1.19 | 9.58E-03 | DE only in male placentas |
| ENSRNOG00000004595 | | POU6F1 | -1.04 | 8.89E-08 | DE only in male placentas |
| ENSRNOG00000024602 | | PLEKHA7 | -1.08 | 8.47E-18 | DE only in male placentas |
| ENSRNOG00000036831 | | GTSF1 | -2.14 | 8.78E-05 | DE only in male placentas |
| ENSRNOG00000002145 | | TMPRSS7 | -1.65 | 9.25E-03 | DE only in male placentas |
| ENSRNOG00000029749 | | PABPC4L | -1.88 | 8.64E-11 | DE only in male placentas |
| ENSRNOG00000009788 | | SERPINA4 | -1.66 | 4.53E-02 | DE only in male placentas |
| ENSRNOG00000003526 | X | SYTL4 | -1.03 | 2.56E-08 | DE only in male placentas |
| ENSRNOG00000007037 | | CFAP57 | -1.28 | 2.34E-02 | DE only in male placentas |

| | | | | | |
|---|---|---|---|---|---|
| ENSRNOG00000014476 | | EVL | -1.14 | 1.47E-07 | DE only in male placentas |
| ENSRNOG00000006867 | | ETV1 | -1.27 | 7.98E-12 | DE only in male placentas |
| ENSRNOG00000013057 | | PRC1 | -1.11 | 6.35E-23 | DE only in male placentas |
| ENSRNOG00000023261 | X | UBE2Q2L | -1.11 | 7.33E-19 | DE only in male placentas |
| ENSRNOG00000003205 | | LDB2 | -1.34 | 8.39E-11 | DE only in male placentas |
| ENSRNOG00000011342 | | TDH | -1.08 | 3.70E-02 | DE only in male placentas |
| ENSRNOG00000052887 | | DNAJC6 | -1.33 | 3.27E-04 | DE only in male placentas |
| ENSRNOG00000009815 | | PNMA2 | -1.25 | 8.07E-08 | DE only in male placentas |
| ENSRNOG00000027157 | | LY6G6E | -2.05 | 3.68E-02 | DE only in male placentas |
| ENSRNOG00000020932 | | MS4A10 | -1.20 | 4.43E-02 | DE only in male placentas |
| ENSRNOG00000016381 | | UST | -1.04 | 3.74E-10 | DE only in male placentas |
| ENSRNOG00000003476 | | SLC6A4 | -1.29 | 3.39E-08 | DE only in male placentas |
| ENSRNOG00000021000 | | NAALADL1 | -1.01 | 2.55E-05 | DE only in male placentas |
| ENSRNOG00000013248 | | WWC2 | -1.29 | 9.75E-28 | DE only in male placentas |
| ENSRNOG00000007246 | | ATXN7 | -1.06 | 2.16E-21 | DE only in male placentas |
| ENSRNOG00000003762 | X | SMARCA1 | -1.40 | 7.04E-14 | DE only in male placentas |
| ENSRNOG00000014276 | | PLCE1 | -1.33 | 2.08E-17 | DE only in male placentas |
| ENSRNOG00000003164 | | PLA2G10 | -1.00 | 1.15E-06 | DE only in male placentas |

| | | | | | |
|---|---|---|---|---|---|
| ENSRNOG00000025287 | | RGD1565611 | -2.14 | 1.24E-03 | DE only in male placentas |
| ENSRNOG00000016505 | | FAM169A | -1.47 | 1.28E-19 | DE only in male placentas |
| ENSRNOG00000013616 | | TMEM213 | -2.23 | 5.23E-04 | DE only in male placentas |
| ENSRNOG00000008855 | | GJB2 | -1.53 | 4.45E-09 | DE only in male placentas |
| ENSRNOG00000001957 | | SULT1E1 | -1.25 | 4.72E-04 | DE only in male placentas |
| ENSRNOG00000055476 | | AABR070178 68.3 | -1.44 | 4.97E-02 | DE only in male placentas |
| ENSRNOG00000009605 | X | TFE3 | -1.07 | 2.20E-20 | DE only in male placentas |
| ENSRNOG00000002984 | X | BMP15 | -1.09 | 1.85E-02 | DE only in male placentas |
| ENSRNOG00000011875 | | SLC2A2 | -1.38 | 4.58E-02 | DE only in male placentas |
| ENSRNOG00000019680 | | MST1 | -1.55 | 1.11E-03 | DE only in male placentas |
| ENSRNOG00000016374 | | FGFR2 | -1.30 | 9.24E-20 | DE only in male placentas |
| ENSRNOG00000042668 | | MSRB3 | -1.60 | 1.16E-19 | DE only in male placentas |
| ENSRNOG00000055300 | | NCAPD2 | -1.73 | 8.95E-11 | DE only in male placentas |
| ENSRNOG00000012756 | X | SLC25A43 | -1.09 | 1.77E-11 | DE only in male placentas |
| ENSRNOG00000015911 | | LRP5 | -1.19 | 3.01E-14 | DE only in male placentas |
| ENSRNOG00000018184 | | TPM1 | -1.05 | 2.52E-29 | DE only in male placentas |
| ENSRNOG00000002255 | | FAM162A | -1.20 | 1.17E-23 | DE only in male placentas |
| ENSRNOG00000053303 | | BEND4 | -1.11 | 6.86E-23 | DE only in male placentas |

| | | | | | |
|---|---|---|---|---|---|
| ENSRNOG00000020484 | | TRPM5 | -1.80 | 1.04E-03 | DE only in male placentas |
| ENSRNOG00000019549 | | AKAP12 | -1.19 | 1.30E-24 | DE only in male placentas |
| ENSRNOG00000049063 | | AABR070447 59.1 | -1.35 | 9.62E-11 | DE only in male placentas |
| ENSRNOG00000015695 | | SULT5A1 | -2.22 | 3.13E-09 | DE only in male placentas |
| ENSRNOG00000002936 | | GOLT1A | -1.03 | 1.86E-02 | DE only in male placentas |
| ENSRNOG00000019598 | | VEGFA | -1.14 | 2.33E-18 | DE only in male placentas |
| ENSRNOG00000018780 | | SH3RF2 | -1.07 | 1.41E-14 | DE only in male placentas |
| ENSRNOG00000002428 | X | LPAR4 | -1.38 | 3.01E-11 | DE only in male placentas |
| ENSRNOG00000027014 | | SPATA31D1 | -1.51 | 2.12E-05 | DE only in male placentas |
| ENSRNOG00000003675 | X | PTCHD1 | -1.61 | 2.42E-03 | DE only in male placentas |
| ENSRNOG00000010529 | | THBS2 | -1.57 | 1.35E-15 | DE only in male placentas |
| ENSRNOG00000031222 | X | RGD1561327 | -2.28 | 2.81E-03 | DE only in male placentas |
| ENSRNOG00000015257 | | CORO2B | -1.28 | 4.91E-12 | DE only in male placentas |
| ENSRNOG00000032977 | | AABR070258 25.1 | -1.56 | 1.95E-05 | DE only in male placentas |
| ENSRNOG00000015061 | | IGFALS | -1.72 | 2.40E-04 | DE only in male placentas |
| ENSRNOG00000004487 | | NEK2 | -1.38 | 1.04E-11 | DE only in male placentas |
| ENSRNOG00000010128 | | SLC27A2 | -1.93 | 2.37E-07 | DE only in male placentas |
| ENSRNOG00000022225 | | LRRC19 | -1.25 | 9.67E-03 | DE only in male placentas |

| | | | | | |
|---|---|---|---|---|---|
| ENSRNOG00000036829 | | NCKAP1L | -1.32 | 1.30E-14 | DE only in male placentas |
| ENSRNOG00000019075 | | STAT5B | -1.02 | 2.36E-14 | DE only in male placentas |
| ENSRNOG00000029342 | | SCN7A | -2.05 | 6.31E-07 | DE only in male placentas |
| ENSRNOG00000054438 | | LOC292543 | -1.10 | 2.09E-07 | DE only in male placentas |
| ENSRNOG00000019560 | | PDE2A | -1.29 | 1.23E-12 | DE only in male placentas |
| ENSRNOG00000037865 | | HORMAD2 | -1.29 | 3.91E-13 | DE only in male placentas |
| ENSRNOG00000019368 | | SCNN1A | -1.75 | 1.08E-15 | DE only in male placentas |
| ENSRNOG00000017021 | | GALNT18 | -1.80 | 5.53E-07 | DE only in male placentas |
| ENSRNOG00000018716 | | DENND2C | -1.19 | 6.22E-23 | DE only in male placentas |
| ENSRNOG00000000873 | X | BRS3 | -2.01 | 7.65E-08 | DE only in male placentas |
| ENSRNOG00000033653 | | MPP3 | -1.75 | 7.22E-08 | DE only in male placentas |
| ENSRNOG00000033933 | | STFA3 | -2.27 | 2.65E-03 | DE only in male placentas |
| ENSRNOG00000038540 | | NEWGENE_2724 | -1.26 | 8.97E-04 | DE only in male placentas |
| ENSRNOG00000009845 | | ACADM | -1.17 | 3.12E-20 | DE only in male placentas |
| ENSRNOG00000005345 | | VSNL1 | -1.37 | 3.11E-02 | DE only in male placentas |
| ENSRNOG00000045663 | X | LOC102546678 | -1.79 | 2.67E-05 | DE only in male placentas |
| ENSRNOG00000047005 | | KCNK5 | -1.32 | 1.64E-10 | DE only in male placentas |
| ENSRNOG00000013867 | | FGF1 | -1.59 | 6.91E-11 | DE only in male placentas |

| | | | | | |
|---|---|---|---|---|---|
| ENSRNOG00000007610 | | GDF11 | -1.39 | 8.39E-14 | DE only in male placentas |
| ENSRNOG00000039571 | X | GLOD5 | -1.61 | 2.29E-09 | DE only in male placentas |
| ENSRNOG00000042120 | | RGD1564129 | -2.13 | 9.10E-04 | DE only in male placentas |
| ENSRNOG00000001431 | | RASA4 | -1.02 | 3.41E-11 | DE only in male placentas |
| ENSRNOG00000022268 | | PNPLA3 | -1.52 | 1.04E-08 | DE only in male placentas |
| ENSRNOG00000011705 | | STMN2 | -1.09 | 2.11E-02 | DE only in male placentas |
| ENSRNOG00000012258 | | RRAS2 | -1.36 | 1.13E-23 | DE only in male placentas |
| ENSRNOG00000009629 | | CAR2 | -1.15 | 3.01E-17 | DE only in male placentas |
| ENSRNOG00000000618 | | MDGA2 | -2.03 | 1.15E-02 | DE only in male placentas |
| ENSRNOG00000021924 | | CYP2C22 | -1.60 | 1.56E-02 | DE only in male placentas |
| ENSRNOG00000016837 | | CKM | -2.39 | 1.27E-04 | DE only in male placentas |
| ENSRNOG00000000850 | | APOM | -1.45 | 1.77E-02 | DE only in male placentas |
| ENSRNOG00000028377 | | MPL | -1.59 | 3.08E-04 | DE only in male placentas |
| ENSRNOG00000015518 | | RBP4 | -1.59 | 1.50E-02 | DE only in male placentas |
| ENSRNOG00000010349 | | GHRL | -1.65 | 1.50E-02 | DE only in male placentas |
| ENSRNOG00000014652 | | AQP8 | -1.62 | 4.25E-06 | DE only in male placentas |
| ENSRNOG00000042344 | | SMIM22 | -1.02 | 3.31E-02 | DE only in male placentas |
| ENSRNOG00000014502 | | TRIML1 | -3.22 | 2.19E-03 | DE only in male placentas |

| | | | | |
|---|---|---|---|---|
| ENSRNOG00000021518 | TRIM31 | -2.90 | 1.42E-04 | DE only in male placentas |
| ENSRNOG00000024119 | GHSR | -3.73 | 9.19E-03 | DE only in male placentas |
| ENSRNOG00000046319 | AABR070000 46.1 | -1.76 | 3.16E-04 | DE only in male placentas |
| ENSRNOG00000010094 | KCNMB2 | -2.42 | 6.16E-04 | DE only in male placentas |
| ENSRNOG00000004679 | FIGN | -1.51 | 5.52E-05 | DE only in male placentas |
| ENSRNOG00000005780 | ADAMTS13 | -1.17 | 1.92E-02 | DE only in male placentas |
| ENSRNOG00000029047 | CUBN | -1.69 | 7.74E-04 | DE only in male placentas |
| ENSRNOG00000001766 | TFRC | -2.17 | 2.94E-07 | DE only in male placentas |
| ENSRNOG00000025408 | DUX4 | -2.25 | 4.06E-09 | DE only in male placentas |
| ENSRNOG00000055271 | AABR070431 90.1 | -2.55 | 7.27E-06 | DE only in male placentas |
| ENSRNOG00000025781 | SEC31B | -1.66 | 1.64E-04 | DE only in male placentas |
| ENSRNOG00000047457 | AABR070717 65.1 | -1.85 | 1.16E-03 | DE only in male placentas |
| ENSRNOG00000046635 | CDCA5 | -1.15 | 1.73E-03 | DE only in male placentas |
| ENSRNOG00000030771 | DGKB | -2.72 | 3.65E-11 | DE only in male placentas |
| ENSRNOG00000032374 | PAQR9 | -2.70 | 5.54E-10 | DE only in male placentas |
| ENSRNOG00000010497 | RGD1305807 | -1.41 | 5.76E-03 | DE only in male placentas |
| ENSRNOG00000025648 | DHRS7L1 | -2.08 | 2.48E-07 | DE only in male placentas |
| ENSRNOG00000018603 | CARNS1 | -1.18 | 5.65E-03 | DE only in male placentas |

| | | | | | |
|---|---|---|---|---|---|
| ENSRNOG00000036622 | | GLYATL1 | -3.63 | 2.56E-03 | DE only in male placentas |
| ENSRNOG00000013950 | | AADAC | -2.63 | 8.73E-06 | DE only in male placentas |
| ENSRNOG00000008467 | | AGBL2 | -1.31 | 1.23E-03 | DE only in male placentas |
| ENSRNOG00000030910 | | GRIK4 | -2.66 | 1.26E-07 | DE only in male placentas |
| ENSRNOG00000004009 | X | XPNPEP2 | -2.20 | 8.27E-06 | DE only in male placentas |
| ENSRNOG00000021782 | | ZSCAN10 | -1.49 | 2.33E-03 | DE only in male placentas |
| ENSRNOG00000040201 | | ATP6AP1L | -1.83 | 7.54E-05 | DE only in male placentas |
| ENSRNOG00000061921 | | AABR07018064.1 | -2.36 | 3.21E-02 | DE only in male placentas |
| ENSRNOG00000005542 | | APOB | -1.82 | 5.50E-03 | DE only in male placentas |
| ENSRNOG00000023721 | | LOC502684 | -1.44 | 2.87E-02 | DE only in male placentas |
| ENSRNOG00000020157 | | WNT8A | -1.37 | 5.44E-04 | DE only in male placentas |
| ENSRNOG00000049303 | | MARCO | -1.72 | 4.32E-02 | DE only in male placentas |
| ENSRNOG00000054395 | | AC114452.1 | -3.33 | 2.38E-03 | DE only in male placentas |
| ENSRNOG00000013771 | | CLCA2 | -1.94 | 3.38E-04 | DE only in male placentas |
| ENSRNOG00000004216 | | LRR1 | -1.48 | 1.07E-02 | DE only in male placentas |
| ENSRNOG00000034191 | | FMO1 | -2.15 | 4.17E-04 | DE only in male placentas |
| ENSRNOG00000036827 | | PPP1R1A | -1.81 | 3.88E-04 | DE only in male placentas |
| ENSRNOG00000000036 | | KLHDC8A | -1.85 | 4.56E-05 | DE only in male placentas |

| | | | | | |
|---|---|---|---|---|---|
| ENSRNOG00000014492 | | PGC | -1.89 | 3.58E-04 | DE only in male placentas |
| ENSRNOG00000014070 | | KRT13 | -2.41 | 3.43E-02 | DE only in male placentas |
| ENSRNOG00000009033 | | CNTN2 | -1.69 | 8.83E-04 | DE only in male placentas |
| ENSRNOG00000054204 | | GRIA2 | -2.18 | 1.42E-02 | DE only in male placentas |
| ENSRNOG00000017933 | | CCDC3 | -2.11 | 3.03E-05 | DE only in male placentas |
| ENSRNOG00000014661 | | ZFP385D | -2.03 | 8.11E-05 | DE only in male placentas |
| ENSRNOG00000026759 | | CCR8 | -1.50 | 3.54E-03 | DE only in male placentas |
| ENSRNOG00000023508 | | MFSD2B | -1.84 | 6.08E-03 | DE only in male placentas |
| ENSRNOG00000010261 | | RNASE10 | -1.44 | 1.17E-02 | DE only in male placentas |
| ENSRNOG00000007495 | X | SMPX | -1.88 | 1.84E-05 | DE only in male placentas |
| ENSRNOG00000046264 | | PCDHB7 | -1.79 | 1.45E-02 | DE only in male placentas |
| ENSRNOG00000013137 | | CLEC4F | -2.12 | 1.92E-04 | DE only in male placentas |
| ENSRNOG00000012819 | | GDNF | -2.14 | 2.72E-02 | DE only in male placentas |
| ENSRNOG00000015306 | | SLC9A4 | -3.18 | 6.81E-03 | DE only in male placentas |
| ENSRNOG00000047740 | | AABR070528 94.1 | -2.41 | 6.32E-04 | DE only in male placentas |
| ENSRNOG00000060598 | | AABR070558 64.2 | -3.80 | 3.05E-05 | DE only in male placentas |
| ENSRNOG00000015438 | | LOC501233 | -2.01 | 3.06E-05 | DE only in male placentas |
| ENSRNOG00000015365 | | COL4A3 | -2.61 | 5.17E-07 | DE only in male placentas |

| | | | | | |
|---|---|---|---|---|---|
| ENSRNOG00000050658 | | TACR2 | -2.94 | 1.56E-05 | DE only in male placentas |
| ENSRNOG00000005375 | | RGD1564053 | -2.24 | 5.18E-03 | DE only in male placentas |
| ENSRNOG00000015008 | | GCM2 | -1.53 | 2.97E-03 | DE only in male placentas |
| ENSRNOG00000029778 | X | MAOB | -2.10 | 1.01E-05 | DE only in male placentas |
| ENSRNOG00000025990 | | IGLL1 | -1.95 | 1.82E-04 | DE only in male placentas |
| ENSRNOG00000014907 | | ALDH8A1 | -2.52 | 3.05E-05 | DE only in male placentas |
| ENSRNOG00000017689 | | ITIH3 | -1.54 | 1.60E-02 | DE only in male placentas |
| ENSRNOG00000053843 | | AABR070565 03.1 | -4.03 | 1.53E-04 | DE only in male placentas |
| ENSRNOG00000003262 | | CACNG4 | -1.41 | 2.80E-03 | DE only in male placentas |
| ENSRNOG00000004084 | | LRATD1 | -1.52 | 1.75E-03 | DE only in male placentas |
| ENSRNOG00000014163 | | ZFP536 | -1.40 | 1.99E-02 | DE only in male placentas |
| ENSRNOG00000057304 | | LOC1025523 98 | -2.26 | 4.73E-04 | DE only in male placentas |
| ENSRNOG00000028707 | | NP4 | -1.98 | 4.94E-02 | DE only in male placentas |
| ENSRNOG00000005592 | | BRINP2 | -1.57 | 1.42E-02 | DE only in male placentas |
| ENSRNOG00000009438 | | SERPINA6 | -2.06 | 5.40E-04 | DE only in male placentas |
| ENSRNOG00000009239 | | ENTPD8 | -2.09 | 5.28E-03 | DE only in male placentas |
| ENSRNOG00000029465 | | SLC26A10 | -2.74 | 2.10E-07 | DE only in male placentas |
| ENSRNOG00000059367 | | U1 | -2.97 | 1.17E-03 | DE only in male placentas |

| | | | | | |
|---|---|---|---|---|---|
| ENSRNOG00000029798 | | LRRC4C | -2.08 | 5.13E-04 | DE only in male placentas |
| ENSRNOG00000002350 | | EAF2 | -1.73 | 3.84E-04 | DE only in male placentas |
| ENSRNOG00000016179 | | ACOXL | -2.25 | 2.42E-06 | DE only in male placentas |
| ENSRNOG00000012851 | | DNAJC5B | -3.01 | 3.26E-07 | DE only in male placentas |
| ENSRNOG00000009043 | | PIWIL4 | -3.03 | 7.12E-09 | DE only in male placentas |
| ENSRNOG00000032560 | | CYP3A23/3A1 | -2.57 | 1.05E-02 | DE only in male placentas |
| ENSRNOG00000058275 | | SPEF2 | -1.84 | 7.63E-03 | DE only in male placentas |
| ENSRNOG00000005367 | | SLC12A1 | -2.41 | 2.04E-04 | DE only in male placentas |
| ENSRNOG00000007700 | | INHBC | -2.61 | 2.02E-02 | DE only in male placentas |
| ENSRNOG00000061769 | | AABR07026044.1 | -1.50 | 2.67E-02 | DE only in male placentas |
| ENSRNOG00000047013 | X | PRRG3 | -1.53 | 5.30E-03 | DE only in male placentas |
| ENSRNOG00000037188 | | MUG1 | -1.56 | 2.68E-02 | DE only in male placentas |
| ENSRNOG00000050884 | | AABR07058745.1 | -2.95 | 4.11E-07 | DE only in male placentas |
| ENSRNOG00000057082 | | AABR07031740.1 | -1.56 | 5.57E-03 | DE only in male placentas |
| ENSRNOG00000026762 | | RT1-DOA | -2.35 | 7.64E-04 | DE only in male placentas |
| ENSRNOG00000027212 | | TEKT3 | -1.69 | 1.07E-03 | DE only in male placentas |
| ENSRNOG00000018402 | | APOC2 | -2.01 | 2.97E-03 | DE only in male placentas |
| ENSRNOG00000059519 | | BBOX1 | -2.21 | 1.12E-03 | DE only in male placentas |

| | | | | |
|---|---|---|---|---|
| ENSRNOG00000060041 | AABR07002677.2 | -1.52 | 5.46E-03 | DE only in male placentas |
| ENSRNOG00000046521 | LOC100911572 | -2.97 | 6.80E-04 | DE only in male placentas |
| ENSRNOG00000047551 | LOC100911615 | -6.35 | 2.14E-04 | DE only in male placentas |
| ENSRNOG00000001865 | SERPIND1 | -1.49 | 1.43E-02 | DE only in male placentas |
| ENSRNOG00000012286 | IL20RA | -2.00 | 2.97E-03 | DE only in male placentas |
| ENSRNOG00000011250 | INMT | -1.67 | 4.64E-03 | DE only in male placentas |
| ENSRNOG00000021128 | KCNJ11 | -1.91 | 2.14E-02 | DE only in male placentas |
| ENSRNOG00000019417 | HSD3B5 | -2.72 | 1.03E-04 | DE only in male placentas |
| ENSRNOG00000060697 | AABR07061261.2 | -3.52 | 1.94E-05 | DE only in male placentas |
| ENSRNOG00000002164 | NMU | -2.98 | 5.34E-03 | DE only in male placentas |
| ENSRNOG00000006807 | ALDOB | -2.07 | 3.10E-04 | DE only in male placentas |
| ENSRNOG00000012613 | F13B | -3.58 | 3.10E-05 | DE only in male placentas |
| ENSRNOG00000051612 | AABR07044570.2 | -2.47 | 1.15E-02 | DE only in male placentas |
| ENSRNOG00000046294 | RGD1624210 | -2.15 | 3.10E-03 | DE only in male placentas |
| ENSRNOG00000050761 | LOC100909524 | -3.53 | 8.98E-03 | DE only in male placentas |
| ENSRNOG00000012387 | GLYATL2 | -2.62 | 9.72E-04 | DE only in male placentas |
| ENSRNOG00000012222 | CDH16 | -1.64 | 5.79E-03 | DE only in male placentas |
| ENSRNOG00000019689 | VWF | -2.78 | 1.55E-05 | DE only in male placentas |

| | | | | |
|---|---|---|---|---|
| ENSRNOG00000056622 | 7SK | -2.56 | 9.50E-03 | DE only in male placentas |
| ENSRNOG00000019510 | TM4SF5 | -1.63 | 1.91E-02 | DE only in male placentas |
| ENSRNOG00000009203 | RGD1561795 | -2.97 | 7.60E-07 | DE only in male placentas |
| ENSRNOG00000015763 | NAT8F3 | -3.61 | 2.39E-03 | DE only in male placentas |
| ENSRNOG00000061610 | AABR070441 48.1 | -3.26 | 2.74E-06 | DE only in male placentas |
| ENSRNOG00000003902 | FABP6 | -2.12 | 4.33E-02 | DE only in male placentas |
| ENSRNOG00000058626 | KCNH8 | -2.24 | 2.43E-02 | DE only in male placentas |
| ENSRNOG00000057315 | KCNH3 | -2.20 | 5.95E-03 | DE only in male placentas |
| ENSRNOG00000016838 | PLA2G5 | -1.74 | 4.71E-03 | DE only in male placentas |
| ENSRNOG00000061664 | LOC502908 | -2.90 | 1.51E-02 | DE only in male placentas |
| ENSRNOG00000001379 | CYP3A62 | -2.03 | 5.01E-03 | DE only in male placentas |
| ENSRNOG00000022669 | CD177 | -2.35 | 7.93E-03 | DE only in male placentas |
| ENSRNOG00000049425 | FAM167B | -2.36 | 5.19E-04 | DE only in male placentas |
| ENSRNOG00000002555 | SERPINB7 | -2.21 | 1.73E-02 | DE only in male placentas |
| ENSRNOG00000028108 | CYTL1 | -3.40 | 6.29E-06 | DE only in male placentas |
| ENSRNOG00000048451 | APOL2 | -2.52 | 3.22E-02 | DE only in male placentas |
| ENSRNOG00000048970 | LOC1009118 04 | -2.52 | 3.22E-02 | DE only in male placentas |
| ENSRNOG00000030538 | SLCO1B2 | -3.13 | 1.31E-04 | DE only in male placentas |

| | | | | |
|---|---|---|---|---|
| ENSRNOG00000050034 | LOC501297 | -2.09 | 5.46E-03 | DE only in male placentas |
| ENSRNOG00000028832 | SORCS3 | -2.31 | 3.55E-03 | DE only in male placentas |
| ENSRNOG00000029128 | CYP2D5 | -2.36 | 1.44E-02 | DE only in male placentas |
| ENSRNOG00000001571 | CLDN8 | -4.21 | 1.95E-08 | DE only in male placentas |
| ENSRNOG00000022509 | AABR070662 01.1 | -3.02 | 3.04E-03 | DE only in male placentas |
| ENSRNOG00000050595 | MUP5 | -3.25 | 1.92E-04 | DE only in male placentas |
| ENSRNOG00000052789 | AABR070445 62.1 | -3.47 | 2.09E-05 | DE only in male placentas |
| ENSRNOG00000013407 | SLC5A11 | -2.23 | 2.99E-02 | DE only in male placentas |
| ENSRNOG00000021949 | PRAMEL7 | -3.66 | 6.48E-04 | DE only in male placentas |
| ENSRNOG00000059443 | LOC1036900 28 | -4.83 | 4.34E-03 | DE only in male placentas |
| ENSRNOG00000018710 | PIDD1 | -3.37 | 4.77E-02 | DE only in male placentas |

*Ctrl: vehicle control (DMSO); *P*-value: Benjamini Hochberg adjusted *P*-value. DE: differentially expressed. $P < 0.05$ is considered significantly differentially expressed. n = 13, 16, 8, and 13 for female-DMSO, male-DMSO, female-ITE, and male-ITE rat placentas,

S3. ITE-dysregulated biological function in rat placentas*

| | Biological Function | *P*-value [Female placentas] | *P*-value [Male placentas] |
|---|---|---|---|
| 1. | Non-melanoma solid tumor | **4.30E-54** | **9.50E-75** |
| 2. | Tumorigenesis of tissue | **2.01E-51** | **1.32E-74** |
| 3. | Epithelial neoplasm | **1.01E-51** | **2.19E-73** |
| 4. | Non-hematological solid tumor | **2.02E-52** | **2.66E-73** |
| 5. | Nonhematologic malignant neoplasm | **5.75E-50** | **7.88E-71** |
| 6. | Carcinoma | **5.72E-49** | **1.39E-70** |
| 7. | Extrapancreatic malignant tumor | **9.08E-45** | **1.06E-69** |
| 8. | Cancer | **2.09E-46** | **3.38E-67** |
| 9. | Solid tumor | **1.35E-47** | **4.12E-67** |
| 10. | Abdominal neoplasm | **1.27E-41** | **1.17E-66** |
| 11. | Malignant solid tumor | **2.41E-46** | **2.11E-66** |
| 12. | Extracranial solid tumor | **9.86E-45** | **1.67E-65** |
| 13. | Abdominal carcinoma | **6.75E-39** | **1.24E-62** |
| 14. | Head and neck tumor | **3.03E-43** | **1.57E-62** |
| 15. | Abdominal cancer | **1.22E-38** | **2.13E-62** |
| 16. | Skin lesion | **3.36E-36** | **8.63E-62** |
| 17. | Adenocarcinoma | **1.33E-37** | **1.22E-61** |
| 18. | Melanoma | **2.87E-35** | **1.42E-61** |
| 19. | Intraabdominal organ tumor | **1.73E-38** | **5.97E-61** |
| 20. | Skin tumor | **4.24E-35** | **2.26E-60** |
| 21. | Abdominal adenocarcinoma | **9.26E-37** | **3.96E-60** |
| 22. | Digestive organ tumor | **5.70E-36** | **4.31E-60** |
| 23. | Formation of solid tumor | **3.38E-40** | **6.27E-60** |
| 24. | Skin cancer | **5.65E-35** | **1.32E-59** |
| 25. | Malignant solid organ tumor | **2.80E-34** | **3.66E-58** |
| 26. | Malignant connective or soft tissue neoplasm | **3.65E-34** | **3.97E-58** |
| 27. | Cutaneous melanoma | **1.02E-34** | **1.64E-57** |
| 28. | Digestive system cancer | **1.49E-34** | **2.03E-57** |
| 29. | Connective or soft tissue tumor | **6.35E-35** | **2.84E-57** |
| 30. | Soft tissue neoplasm | **3.19E-34** | **1.88E-56** |
| 31. | Malignant soft tissue neoplasm | **3.65E-34** | **1.88E-56** |
| 32. | Gastrointestinal carcinoma | **4.07E-35** | **2.26E-52** |
| 33. | Head and neck cancer | **2.35E-38** | **4.64E-52** |
| 34. | Cancer of secretory structure | **4.58E-38** | **6.05E-52** |
| 35. | Gastrointestinal tract cancer | **1.44E-34** | **1.91E-51** |
| 36. | Gastrointestinal tumor | **4.07E-35** | **1.99E-51** |
| 37. | Endocrine gland tumor | **3.83E-34** | **1.37E-49** |
| 38. | Gastrointestinal adenocarcinoma | **4.99E-29** | **8.00E-49** |
| 39. | Head and neck carcinoma | **3.55E-34** | **4.69E-48** |
| 40. | Large intestine carcinoma | **2.05E-31** | **6.58E-48** |
| 41. | Large intestine adenocarcinoma | **8.67E-31** | **1.86E-46** |

| | | | |
|---|---|---|---|
| 42. | Malignant neoplasm of large intestine | **1.09E-30** | **1.90E-46** |
| 43. | Large intestine neoplasm | **5.50E-31** | **8.45E-46** |
| 44. | Intestinal tumor | **5.50E-31** | **1.03E-45** |
| 45. | Endocrine carcinoma | **1.68E-33** | **4.94E-45** |
| 46. | Endocrine cancer | **1.68E-33** | **6.24E-45** |
| 47. | Nonpituitary endocrine tumor | **2.25E-30** | **9.67E-44** |
| 48. | Neck neoplasm | **6.26E-33** | **1.92E-42** |
| 49. | Thyroid carcinoma | **1.16E-30** | **1.68E-40** |
| 50. | Thyroid gland tumor | **4.05E-30** | **2.10E-40** |
| 51. | Incidence of tumor | **7.04E-29** | **3.09E-40** |
| 52. | Glioma | **2.22E-28** | **2.80E-37** |
| 53. | Nervous system neoplasm | **1.08E-27** | **9.13E-37** |
| 54. | Central nervous system solid tumor | **2.31E-26** | **4.62E-36** |
| 55. | Frequency of tumor | **1.27E-23** | **1.20E-35** |
| 56. | Glioma cancer | **9.04E-28** | **3.21E-35** |
| 57. | Brain lesion | **1.07E-25** | **4.44E-35** |
| 58. | Pelvic tumor | **4.39E-24** | **5.00E-35** |
| 59. | Tumorigenesis of epithelial neoplasm | **2.42E-22** | **2.05E-34** |
| 60. | Development of malignant tumor | **3.82E-22** | **3.23E-34** |
| 61. | Central nervous system cancer | **4.21E-26** | **3.64E-34** |
| 62. | Brain glioma | **1.72E-25** | **3.98E-34** |
| 63. | Pelvic cancer | **6.37E-23** | **2.28E-33** |
| 64. | Brain tumor | **1.63E-24** | **2.68E-33** |
| 65. | Genital tumor | **3.82E-22** | **4.68E-33** |
| 66. | Malignant genitourinary solid tumor | **7.55E-20** | **1.52E-32** |
| 67. | Development of carcinoma | **2.04E-20** | **2.09E-32** |
| 68. | Genitourinary tumor | **8.41E-20** | **2.33E-32** |
| 69. | Genital tract cancer | **1.26E-20** | **2.44E-31** |
| 70. | Genitourinary carcinoma | **1.20E-17** | **3.81E-31** |
| 71. | Cell movement | **6.75E-20** | **1.62E-30** |
| 72. | Anogenital cancer | **7.51E-19** | **3.12E-30** |
| 73. | Neoplasia of cells | **2.10E-15** | **1.31E-29** |
| 74. | Migration of cells | **1.68E-18** | **1.85E-29** |
| 75. | Pelvic carcinoma | **8.61E-18** | **2.79E-28** |
| 76. | Development of digestive organ tumor | **6.35E-19** | **3.72E-28** |
| 77. | Liver lesion | **3.51E-19** | **4.92E-26** |
| 78. | Quantity of cells | **4.12E-18** | **5.85E-26** |
| 79. | Pelvic adenocarcinoma | **9.19E-16** | **6.08E-26** |
| 80. | Genitourinary adenocarcinoma | **2.02E-15** | **7.09E-26** |
| 81. | Colon tumor | **1.67E-17** | **9.60E-26** |
| 82. | Respiratory system tumor | **8.33E-19** | **1.39E-25** |
| 83. | Astrocytoma | **2.09E-20** | **7.13E-25** |
| 84. | High grade astrocytoma | **3.58E-20** | **1.57E-24** |
| 85. | Quantity of blood cells | **1.78E-15** | **1.62E-24** |
| 86. | Disorder of blood pressure | **2.76E-21** | **1.80E-24** |

| | | | |
|---|---|---|---|
| 87. | Cancer of cells | **1.87E-15** | **2.40E-24** |
| 88. | Malignant neoplasm of respiratory system | **2.64E-17** | **3.07E-24** |
| 89. | Colon cancer | **1.59E-15** | **4.18E-24** |
| 90. | Colorectal lesion | **1.68E-18** | **5.23E-24** |
| 91. | Cell movement of blood cells | **4.95E-10** | **5.41E-24** |
| 92. | Activation of cells | **1.37E-08** | **5.84E-24** |
| 93. | Colorectal tumor | **2.79E-18** | **8.46E-24** |
| 94. | Breast or colorectal cancer | **1.46E-16** | **1.04E-23** |
| 95. | Hypertension | **6.88E-22** | **1.07E-23** |
| 96. | Quantity of leukocytes | **5.63E-13** | **1.38E-23** |
| 97. | Lung tumor | **9.42E-17** | **1.46E-23** |
| 98. | Angiogenesis | **3.57E-26** | **1.50E-23** |
| 99. | Intrathoracic organ tumor | **4.83E-17** | **1.52E-23** |
| 100. | Development of vasculature | **5.65E-28** | **1.98E-23** |
| 101. | Lung cancer | **2.86E-16** | **5.10E-23** |
| 102. | Colorectal cancer | **1.42E-16** | **5.40E-23** |
| 103. | Diabetes mellitus | **9.91E-11** | **5.93E-23** |
| 104. | Leukocyte migration | **4.19E-09** | **7.22E-23** |
| 105. | Quantity of protein in blood | **1.61E-07** | **9.43E-23** |
| 106. | Prostatic tumor | **3.99E-15** | **1.14E-22** |
| 107. | Cell movement of leukocytes | **2.74E-08** | **1.68E-22** |
| 108. | Development of colorectal tumor | **1.26E-14** | **2.02E-22** |
| 109. | Rheumatic Disease | **1.04E-09** | **2.08E-22** |
| 110. | Prostate cancer | **1.00E-14** | **2.34E-22** |
| 111. | Colon carcinoma | **2.36E-14** | **2.50E-22** |
| 112. | Grade 1-4 astrocytoma | **4.02E-17** | **3.00E-22** |
| 113. | Tumorigenesis of gastrointestinal tumor | **1.55E-14** | **3.08E-22** |
| 114. | Male genital neoplasm | **2.06E-14** | **3.38E-22** |
| 115. | Glucose metabolism disorder | **9.57E-14** | **3.38E-22** |
| 116. | Grade 3-4 glioma cancer | **2.61E-18** | **3.38E-22** |
| 117. | Colorectal carcinoma | **2.78E-14** | **4.25E-22** |
| 118. | Malignant neoplasm of male genital organ | **4.51E-14** | **4.26E-22** |
| 119. | Transport of molecule | **3.99E-15** | **4.87E-22** |
| 120. | Lung carcinoma | **6.35E-16** | **5.40E-22** |
| 121. | Breast or gynecological cancer | **9.43E-16** | **7.02E-22** |
| 122. | Brain astrocytoma | **1.21E-16** | **7.43E-22** |
| 123. | Recruitment of cells | **1.77E-07** | **1.06E-21** |
| 124. | Extraadrenal retroperitoneal tumor | **4.94E-13** | **1.39E-21** |
| 125. | Grade 4 astrocytoma | **8.01E-16** | **1.47E-21** |
| 126. | Grade 4 high grade glioma | **8.01E-16** | **1.47E-21** |
| 127. | Grade 4 malignant glioma | **8.01E-16** | **1.47E-21** |
| 128. | Inflammation of joint | **6.53E-11** | **1.87E-21** |
| 129. | Cellular homeostasis | **1.85E-09** | **1.87E-21** |
| 130. | Liver tumor | **1.54E-17** | **2.07E-21** |
| 131. | Morphology of body cavity | **3.42E-23** | **2.79E-21** |

| | | | |
|---|---|---|---|
| 132. | Systemic autoimmune syndrome | **8.92E-08** | **4.60E-21** |
| 133. | Female genital neoplasm | **2.42E-15** | **5.48E-21** |
| 134. | Pancreatic lesion | **1.30E-11** | **5.67E-21** |
| 135. | Hepatobiliary system cancer | **6.24E-17** | **6.82E-21** |
| 136. | Recruitment of blood cells | **1.28E-06** | **9.20E-21** |
| 137. | Tumorigenesis of reproductive tract | **8.89E-15** | **1.08E-20** |
| 138. | Colon adenocarcinoma | **2.40E-13** | **1.20E-20** |
| 139. | Hepatobiliary carcinoma | **5.80E-17** | **1.24E-20** |
| 140. | Pancreatic tumor | **1.64E-11** | **1.27E-20** |
| 141. | Vasculogenesis | **1.60E-22** | **1.70E-20** |
| 142. | Inflammatory response | **3.86E-07** | **1.96E-20** |
| 143. | Non-colon gastrointestinal cancer | **7.60E-16** | **2.14E-20** |
| 144. | Cell movement of myeloid cells | **9.07E-06** | **2.25E-20** |
| 145. | Liver carcinoma | **1.71E-17** | **3.23E-20** |
| 146. | Liver cancer | **4.10E-17** | **3.90E-20** |
| 147. | Colorectal adenocarcinoma | **4.52E-13** | **6.45E-20** |
| 148. | Female genital tract cancer | **2.10E-14** | **7.76E-20** |
| 149. | Pancreatobiliary tumor | **1.48E-11** | **9.53E-20** |
| 150. | Recruitment of leukocytes | **3.80E-06** | **1.04E-19** |
| 151. | Upper gastrointestinal tract tumor | **1.11E-15** | **1.84E-19** |
| 152. | Upper gastrointestinal tract cancer | **9.83E-16** | **2.33E-19** |
| 153. | Development of body trunk | **4.31E-21** | **3.34E-19** |
| 154. | Lymphohematopoietic cancer | **7.86E-11** | **3.38E-19** |
| 155. | Upper gastrointestinal carcinoma | **1.47E-15** | **3.82E-19** |
| 156. | Quantity of myeloid cells | **4.31E-11** | **4.19E-19** |
| 157. | Hematologic cancer | **1.20E-10** | **4.72E-19** |
| 158. | Lymphohematopoietic neoplasia | **8.06E-11** | **7.21E-19** |
| 159. | Development of genital tumor | **2.61E-14** | **1.13E-18** |
| 160. | Hematopoietic neoplasm | **1.36E-10** | **1.16E-18** |
| 161. | Uterine tumor | **8.95E-14** | **1.53E-18** |
| 162. | Neoplasia of blood cells | **1.16E-09** | **1.61E-18** |
| 163. | Breast or pancreatic cancer | **5.70E-12** | **2.14E-18** |
| 164. | Ion homeostasis of cells | **1.32E-08** | **3.03E-18** |
| 165. | Abnormal morphology of body cavity | **9.77E-19** | **3.43E-18** |
| 166. | Myeloid or lymphoid neoplasm | **5.09E-10** | **3.92E-18** |
| 167. | Squamous-cell carcinoma | **6.10E-10** | **5.42E-18** |
| 168. | Recruitment of granulocytes | **2.97E-06** | **5.69E-18** |
| 169. | Quantity of connective tissue | **5.17E-14** | **6.99E-18** |
| 170. | Abnormality of endometrium | **2.07E-12** | **7.43E-18** |
| 171. | Abnormal growth in endometrium | **1.63E-12** | **9.11E-18** |
| 172. | Prostatic carcinoma | **1.95E-10** | **1.57E-17** |
| 173. | Uterine cancer | **5.47E-13** | **1.59E-17** |
| 174. | Gastroesophageal adenocarcinoma | **1.67E-10** | **1.59E-17** |
| 175. | Recruitment of neutrophils | **6.36E-06** | **2.10E-17** |
| 176. | Endometrium tumor | **1.59E-12** | **2.16E-17** |

| | | | |
|---|---|---|---|
| 177. | Endometrial cancer | **2.23E-12** | **2.99E-17** |
| 178. | Extrapancreatic neuroendocrine tumor | **1.40E-11** | **3.20E-17** |
| 179. | Insulin-dependent diabetes mellitus | **8.13E-06** | **3.27E-17** |
| 180. | Development of adenocarcinoma | **7.92E-11** | **3.28E-17** |
| 181. | Recruitment of myeloid cells | **5.34E-06** | **3.31E-17** |
| 182. | Neuroendocrine tumor | **8.55E-12** | **3.81E-17** |
| 183. | Inflammation of absolute anatomical region | **9.91E-06** | **3.94E-17** |
| 184. | Vaso-occlusion | **4.80E-10** | **4.78E-17** |
| 185. | Chronic inflammatory disorder | **3.69E-08** | **4.79E-17** |
| 186. | Occlusion of blood vessel | **2.75E-10** | **5.72E-17** |
| 187. | Quantity of phagocytes | **2.37E-08** | **6.44E-17** |
| 188. | Malignant neoplasm of retroperitoneum | **9.88E-12** | **7.18E-17** |
| 189. | Gastroesophageal cancer | **1.30E-13** | **8.69E-17** |
| 190. | Blood pressure | **7.09E-09** | **9.27E-17** |
| 191. | Gastro-esophageal carcinoma | **8.41E-14** | **9.86E-17** |
| 192. | Transport of ion | **3.03E-14** | **1.14E-16** |
| 193. | Pancreatic cancer | **7.83E-11** | **1.22E-16** |
| 194. | Occlusion of artery | **1.72E-09** | **1.25E-16** |
| 195. | Cancer of head | **5.48E-11** | **1.40E-16** |
| 196. | Inflammation of organ | **3.84E-06** | **1.55E-16** |
| 197. | Hematologic cancer of cells | **4.14E-10** | **1.90E-16** |
| 198. | Inflammation of respiratory system | **1.90E-08** | **2.94E-16** |
| 199. | Prostatic adenocarcinoma | **5.43E-09** | **3.16E-16** |
| 200. | Atherosclerosis | **3.12E-09** | **3.87E-16** |
| 201. | Neoplasia of leukocytes | **2.09E-09** | **5.53E-16** |
| 202. | Inflammation of respiratory system component | **3.65E-08** | **6.26E-16** |
| 203. | Female genital tract adenocarcinoma | **1.90E-11** | **6.31E-16** |
| 204. | Uterine carcinoma | **7.57E-10** | **1.46E-15** |
| 205. | Non-traumatic arthropathy | **4.97E-08** | **1.62E-15** |
| 206. | Tumorigenesis of lymphocytes | **1.49E-09** | **1.82E-15** |
| 207. | Quantity of metal | **3.76E-10** | **2.67E-15** |
| 208. | Lymphoproliferative disorder | **9.32E-10** | **3.20E-15** |
| 209. | Gastric lesion | **3.09E-13** | **3.67E-15** |
| 210. | Lymphoid cancer | **1.20E-09** | **4.72E-15** |
| 211. | Lymphocytic cancer | **1.63E-09** | **5.38E-15** |
| 212. | Homing of cells | **5.63E-06** | **6.19E-15** |
| 213. | Lymphocytic neoplasm | **1.83E-09** | **6.40E-15** |
| 214. | Pancreatic carcinoma | **1.56E-09** | **7.83E-15** |
| 215. | Lymphoreticular neoplasm | **7.89E-08** | **7.91E-15** |
| 216. | Lymphatic system tumor | **2.11E-09** | **1.33E-14** |
| 217. | Skin carcinoma | **9.06E-08** | **1.55E-14** |
| 218. | Stomach tumor | **1.09E-12** | **1.86E-14** |
| 219. | Non-small cell lung carcinoma | **3.38E-10** | **1.98E-14** |

| | | | |
|---|---|---|---|
| 220. | Diabetic complication | **1.37E-08** | **2.24E-14** |
| 221. | Quantity of granulocytes | **1.58E-06** | **2.65E-14** |
| 222. | Extrapulmonary squamous cell carcinoma | **1.66E-08** | **2.89E-14** |
| 223. | Peripheral vascular disease | **3.15E-08** | **3.09E-14** |
| 224. | Pancreatic adenocarcinoma | **3.10E-09** | **3.29E-14** |
| 225. | Gastric cancer | **1.58E-12** | **3.69E-14** |
| 226. | Blue round small cell tumor | **3.70E-09** | **4.95E-14** |
| 227. | Gastric carcinoma | **1.42E-12** | **5.57E-14** |
| 228. | Malignant neuroendocrine neoplasm | **3.22E-09** | **6.51E-14** |
| 229. | Neuroendocrine carcinoma | **3.65E-09** | **6.51E-14** |
| 230. | Endometrial carcinoma | **1.50E-07** | **9.36E-14** |
| 231. | Morbidity or mortality | **3.06E-14** | **9.67E-14** |
| 232. | Quantity of lymphatic system cells | **3.15E-08** | **1.09E-13** |
| 233. | Development of epithelial tissue | **4.71E-12** | **1.39E-13** |
| 234. | Cell movement of lymphatic system cells | **6.09E-06** | **1.48E-13** |
| 235. | Concentration of lipid | **5.48E-06** | **1.53E-13** |
| 236. | Cerebral disorder | **7.68E-10** | **1.80E-13** |
| 237. | Small-cell carcinoma | **7.57E-09** | **1.99E-13** |
| 238. | Brain cancer | **5.69E-08** | **2.05E-13** |
| 239. | Quantity of lymphocytes | **2.44E-09** | **2.09E-13** |
| 240. | Quantity of metal ion | **4.62E-08** | **2.09E-13** |
| 241. | Neuroendocrine lung cancer | **9.78E-09** | **2.37E-13** |
| 242. | Quantity of mononuclear leukocytes | **1.40E-09** | **2.43E-13** |
| 243. | Small cell lung carcinoma | **1.26E-08** | **2.83E-13** |
| 244. | Infarction | **1.07E-09** | **3.12E-13** |
| 245. | Gastric adenocarcinoma | **8.04E-10** | **3.33E-13** |
| 246. | Dysfunction of heart | **8.54E-10** | **3.51E-13** |
| 247. | Pancreatic ductal adenocarcinoma | **1.90E-08** | **3.57E-13** |
| 248. | Lymphoma | **2.21E-08** | **5.47E-13** |
| 249. | Fibrosis | **2.54E-08** | **6.67E-13** |
| 250. | Development of lung tumor | **1.21E-08** | **6.90E-13** |
| 251. | Organismal death | **4.52E-14** | **8.17E-13** |
| 252. | Benign lesion | **6.02E-09** | **8.31E-13** |
| 253. | Preeclampsia | **1.69E-12** | **8.71E-13** |
| 254. | Morphology of digestive system | **2.11E-08** | **9.90E-13** |
| 255. | Non-Hodgkin lymphoma | **1.09E-09** | **1.63E-12** |
| 256. | Transport of cation | **1.66E-09** | **1.69E-12** |
| 257. | Morphology of bone | **2.77E-08** | **2.06E-12** |
| 258. | Benign solid tumor | **1.49E-09** | **2.17E-12** |
| 259. | Transport of metal ion | **5.81E-09** | **2.31E-12** |
| 260. | Ventricular dysfunction | **5.57E-09** | **2.66E-12** |
| 261. | Endothelial cell development | **6.89E-12** | **2.79E-12** |
| 262. | Lung adenocarcinoma | **1.28E-08** | **2.94E-12** |
| 263. | Transport of metal | **1.24E-09** | **3.44E-12** |
| 264. | Cerebrovascular dysfunction | **1.64E-08** | **3.45E-12** |

| | | | |
|---|---|---|---|
| 265. | Assembly of cells | **2.26E-12** | **3.58E-12** |
| 266. | Abnormality of left ventricle | **1.08E-10** | **3.72E-12** |
| 267. | Gliomatosis cerebri | **5.90E-08** | **3.79E-12** |
| 268. | Abnormal morphology of abdomen | **5.98E-09** | **4.40E-12** |
| 269. | Disorder of pregnancy | **1.03E-11** | **4.54E-12** |
| 270. | Disorder of coronary artery | **1.11E-10** | **4.71E-12** |
| 271. | Development of cardiovascular tissue | **1.04E-11** | **5.42E-12** |
| 272. | B cell cancer | **4.58E-07** | **6.33E-12** |
| 273. | Ductal carcinoma | **3.48E-08** | **6.39E-12** |
| 274. | Endometrial adenocarcinoma | **5.40E-07** | **6.90E-12** |
| 275. | Endometrioid carcinoma | **3.18E-06** | **7.95E-12** |
| 276. | Development of abdomen | **5.46E-07** | **9.12E-12** |
| 277. | Diabetic nephropathy | **1.34E-07** | **1.04E-11** |
| 278. | Inflammation of lung | **5.48E-07** | **1.13E-11** |
| 279. | Edema | **3.33E-13** | **1.15E-11** |
| 280. | Obesity | **9.37E-07** | **1.20E-11** |
| 281. | Liquid tumor | **3.04E-06** | **1.24E-11** |
| 282. | Morphology of cardiovascular system | **3.02E-16** | **1.26E-11** |
| 283. | B-cell neoplasm | **4.87E-08** | **1.26E-11** |
| 284. | Vascular lesion | **6.86E-07** | **1.47E-11** |
| 285. | Infarction of heart | **7.92E-11** | **1.58E-11** |
| 286. | Endometrioid endometrial adenocarcinoma | **6.69E-07** | **1.66E-11** |
| 287. | Weight gain | **2.38E-07** | **1.80E-11** |
| 288. | Left heart disease | **3.82E-09** | **2.01E-11** |
| 289. | Left ventricular dysfunction | **1.81E-08** | **2.10E-11** |
| 290. | Myocardial infarction | **1.28E-10** | **2.15E-11** |
| 291. | Skin squamous cell carcinoma | **2.15E-07** | **2.15E-11** |
| 292. | Uterine endometrioid carcinoma | **7.75E-07** | **2.35E-11** |
| 293. | Head and neck squamous cell carcinoma | **2.11E-06** | **2.40E-11** |
| 294. | Transmigration of cells | **4.15E-09** | **2.94E-11** |
| 295. | Multiple cancers | **4.11E-07** | **3.53E-11** |
| 296. | Hypersensitive reaction | **4.27E-07** | **3.98E-11** |
| 297. | Abnormality of heart ventricle | **1.74E-10** | **5.00E-11** |
| 298. | Transport of inorganic cation | **4.68E-08** | **5.30E-11** |
| 299. | Breast or ovarian cancer | **3.74E-07** | **5.61E-11** |
| 300. | Quantity of T lymphocytes | **1.23E-06** | **5.61E-11** |
| 301. | Abnormality of limb | **3.04E-06** | **6.16E-11** |
| 302. | Mature lymphocytic neoplasm | **1.49E-07** | **6.65E-11** |
| 303. | Osteoarthritis | **1.38E-06** | **7.97E-11** |
| 304. | Quantity of Ca2+ | **2.96E-08** | **8.30E-11** |
| 305. | Myelodysplastic syndrome or leukemia | **1.03E-05** | **8.39E-11** |
| 306. | Leukemia | **1.05E-05** | **9.38E-11** |
| 307. | Failure of heart | **1.76E-08** | **1.19E-10** |
| 308. | Stimulation of cells | **2.96E-08** | **1.22E-10** |

| | | | |
|---|---|---|---|
| 309. | Stroke | **3.96E-06** | **1.36E-10** |
| 310. | Abnormal morphology of thoracic cavity | **6.23E-13** | **1.40E-10** |
| 311. | Congestive heart failure | **2.42E-09** | **2.86E-10** |
| 312. | Heart rate | **4.97E-09** | **2.98E-10** |
| 313. | Abnormal morphology of cardiovascular system | **7.68E-14** | **3.20E-10** |
| 314. | Interaction of tumor cell lines | **1.04E-05** | **5.08E-10** |
| 315. | Size of bone | **8.48E-06** | **5.23E-10** |
| 316. | Proliferation of lymphatic system cells | **2.67E-06** | **5.26E-10** |
| 317. | Valvulopathy | **1.77E-06** | **5.64E-10** |
| 318. | Cardiovascular death | **2.68E-08** | **6.17E-10** |
| 319. | Asthma | **1.30E-08** | **7.30E-10** |
| 320. | Morphology of heart ventricle | **1.60E-09** | **7.35E-10** |
| 321. | Morphology of vasculature | **4.36E-13** | **1.17E-09** |
| 322. | Quantity of osteoclasts | **8.26E-07** | **1.28E-09** |
| 323. | Female genital tract serous carcinoma | **1.19E-12** | **1.28E-09** |
| 324. | Pelvic serous carcinoma | **1.19E-12** | **1.28E-09** |
| 325. | Cell movement of endothelial cells | **2.28E-08** | **1.37E-09** |
| 326. | Bleeding | **2.57E-10** | **1.48E-09** |
| 327. | Proliferation of blood cells | **3.31E-06** | **1.63E-09** |
| 328. | Enlargement of heart ventricle | **2.96E-08** | **1.76E-09** |
| 329. | Quantity of adipose tissue | **3.03E-06** | **1.79E-09** |
| 330. | Tauopathy | **1.36E-05** | **1.88E-09** |
| 331. | Mature B cell malignant tumor | **5.58E-07** | **2.95E-09** |
| 332. | Breast or ovarian carcinoma | **3.49E-06** | **2.99E-09** |
| 333. | Development of genitourinary system | **1.74E-10** | **3.09E-09** |
| 334. | Development of sensory organ | **3.03E-07** | **3.10E-09** |
| 335. | Quantity of bone | **1.31E-06** | **3.28E-09** |
| 336. | Mass of organism | **1.46E-06** | **3.36E-09** |
| 337. | Muscle contraction | **4.44E-08** | **3.43E-09** |
| 338. | Sensation | **6.15E-06** | **3.52E-09** |
| 339. | Morphology of heart | **3.41E-11** | **3.78E-09** |
| 340. | Secretion of molecule | **3.02E-06** | **4.01E-09** |
| 341. | Abnormal morphology of digestive system | **3.96E-06** | **4.03E-09** |
| 342. | Ovarian tumor | **8.19E-07** | **4.21E-09** |
| 343. | Mature B-cell neoplasm | **4.70E-07** | **4.29E-09** |
| 344. | Flux of cation | **7.27E-06** | **4.32E-09** |
| 345. | Coronary occlusion | **2.08E-06** | **4.95E-09** |
| 346. | Serous adenocarcinoma | **1.52E-10** | **5.29E-09** |
| 347. | Flux of ion | **1.26E-05** | **5.36E-09** |
| 348. | Proliferation of immune cells | **7.39E-06** | **6.20E-09** |
| 349. | Familial cardiovascular disease | **2.21E-08** | **6.81E-09** |
| 350. | Flux of inorganic cation | **1.22E-05** | **7.02E-09** |
| 351. | Growth of epithelial tissue | **1.85E-08** | **7.20E-09** |
| 352. | Abnormal morphology of heart | **8.68E-11** | **7.55E-09** |

| | | | |
|---|---|---|---|
| 353. | Nociception | **1.29E-05** | **7.60E-09** |
| 354. | Gonadal tumor | **5.77E-07** | **7.81E-09** |
| 355. | Discomfort | **1.16E-05** | **8.94E-09** |
| 356. | Transmigration of leukocytes | **4.06E-07** | **8.99E-09** |
| 357. | Development of connective tissue cells | **2.97E-07** | **9.56E-09** |
| 358. | Contraction of heart | **1.26E-05** | **1.11E-08** |
| 359. | Coronary artery disease | **4.67E-06** | **1.12E-08** |
| 360. | Morphology of vessel | **8.49E-11** | **1.13E-08** |
| 361. | Acute respiratory disorder | **7.60E-07** | **1.20E-08** |
| 362. | Development of head | **1.65E-10** | **1.23E-08** |
| 363. | Uterine serous papillary cancer | **1.03E-09** | **1.37E-08** |
| 364. | Maternal hypertension | **1.20E-12** | 1 |
| 365. | Primary hypertension | **6.07E-11** | 1 |
| 366. | Uterine serous carcinoma | **6.09E-11** | 1 |
| 367. | Cardiogenesis | **1.48E-10** | 1 |
| 368. | Hypertensive heart disease | **3.04E-10** | 1 |
| 369. | Abnormal morphology of vasculature | **3.23E-10** | 1 |
| 370. | Morphology of blood vessel | **4.95E-10** | 1 |
| 371. | Hypertrophy | **1.47E-09** | 1 |
| 372. | Visceromegaly | **1.96E-09** | 1 |
| 373. | Morphology of muscle | **2.01E-09** | 1 |
| 374. | Development of body axis | **3.82E-09** | 1 |
| 375. | Size of body | **2.54E-08** | 1 |
| 376. | Hypertrophy of heart | **3.04E-08** | 1 |
| 377. | Enlargement of heart | **3.44E-08** | 1 |
| 378. | Morphology of reproductive system | **3.95E-08** | 1 |
| 379. | Development of central nervous system | **4.62E-08** | 1 |
| 380. | Stage II hypertension | **6.30E-08** | 1 |
| 381. | Prehypertension | **6.51E-08** | 1 |
| 382. | Tubulation of cells | **7.76E-08** | 1 |
| 383. | Severe hypertension | **7.81E-08** | 1 |
| 384. | Morphology of head | **8.11E-08** | 1 |
| 385. | Formation of muscle | **8.43E-08** | 1 |
| 386. | Sprouting | **9.06E-08** | 1 |
| 387. | Migration of vascular cells | **1.10E-07** | 1 |
| 388. | Abnormal morphology of blood vessel | **1.21E-07** | 1 |
| 389. | Aneurysm | **1.26E-07** | 1 |
| 390. | Respiratory system development | **1.28E-07** | 1 |
| 391. | Formation of lung | **1.34E-07** | 1 |
| 392. | Perinatal death | **1.53E-07** | 1 |
| 393. | Congenital anomalies of kidney and urinary tract | **1.61E-07** | 1 |
| 394. | Insulin resistance | **1.72E-07** | 1 |
| 395. | Hypertensive renal disease | **2.11E-07** | 1 |
| 396. | Cell-cell contact | **2.38E-07** | 1 |

| | | | |
|---|---|---|---|
| 397. | Proliferation of endothelial cells | **2.38E-07** | 1 |
| 398. | Differentiation of epithelial tissue | **2.68E-07** | 1 |
| 399. | X-linked hereditary disease | **2.74E-07** | 1 |
| 400. | Congenital anomaly of cardiovascular system | **2.75E-07** | 1 |
| 401. | Morphology of nervous system | **2.78E-07** | 1 |
| 402. | Abnormal morphology of nervous system | **2.94E-07** | 1 |
| 403. | Morphology of respiratory system | **3.17E-07** | 1 |
| 404. | Chronic kidney disease | **3.97E-07** | 1 |
| 405. | Blastoma | **4.36E-07** | 1 |
| 406. | Supraventricular arrhythmia | **4.47E-07** | 1 |
| 407. | Embryonal tumor | **4.54E-07** | 1 |
| 408. | Neonatal death | **5.13E-07** | 1 |
| 409. | Overweight disorder | **5.40E-07** | 1 |
| 410. | Development of reproductive system | **5.40E-07** | 1 |
| 411. | Migration of endothelial cells | **5.48E-07** | 1 |
| 412. | Metabolic syndrome X | **5.81E-07** | 1 |
| 413. | Abnormal morphology of head | **5.85E-07** | 1 |
| 414. | Urination disorder | **6.09E-07** | 1 |
| 415. | Hypertension of blood vessel | **6.54E-07** | 1 |
| 416. | Concentration of hormone | **7.75E-07** | 1 |
| 417. | Congenital malformation of genitourinary system | **8.61E-07** | 1 |
| 418. | Proliferation of smooth muscle cells | **9.06E-07** | 1 |
| 419. | Formation of plasma membrane | **9.35E-07** | 1 |
| 420. | Polycystic Kidney Disease | **9.57E-07** | 1 |
| 421. | Cystic kidney disease | **1.05E-06** | 1 |
| 422. | Development of neurons | **1.07E-06** | 1 |
| 423. | Differentiation of bone cells | **1.17E-06** | 1 |
| 424. | Vascularization | **1.34E-06** | 1 |
| 425. | Congenital heart disease | **1.38E-06** | 1 |
| 426. | Growth of muscle tissue | **1.42E-06** | 1 |
| 427. | Morphogenesis of cardiovascular system | **1.44E-06** | 1 |
| 428. | Arrhythmia of atrium | **1.55E-06** | 1 |
| 429. | Organization of extracellular matrix | **1.58E-06** | 1 |
| 430. | Formation of intercellular junctions | **1.78E-06** | 1 |
| 431. | Vasoconstriction | **1.85E-06** | 1 |
| 432. | Hepatorenal fibrocystic disease | **1.96E-06** | 1 |
| 433. | Glioblastoma | **1.98E-06** | 1 |
| 434. | B-cell non-Hodgkin lymphoma | **2.07E-06** | 1 |
| 435. | Differentiation of muscle | **2.10E-06** | 1 |
| 436. | Function of muscle | **2.10E-06** | 1 |
| 437. | Familial congenital anomalies of the kidney and urinary tract | **2.22E-06** | 1 |
| 438. | Differentiation of muscle cells | **2.46E-06** | 1 |
| 439. | Transport of anion | **2.48E-06** | 1 |

| | | | |
|---|---|---|---|
| 440. | Chronic heart failure | **2.49E-06** | 1 |
| 441. | Morphology of artery | **2.51E-06** | 1 |
| 442. | Morphology of vessel component | **2.62E-06** | 1 |
| 443. | Enlargement of left ventricle | **2.69E-06** | 1 |
| 444. | Abnormal morphology of heart ventricle | **2.75E-06** | 1 |
| 445. | Autosomal dominant myopathy | **2.75E-06** | 1 |
| 446. | Mass of genitourinary system | **2.75E-06** | 1 |
| 447. | Abnormal morphology of muscle | **2.90E-06** | 1 |
| 448. | Arterial aneurysm | **3.02E-06** | 1 |
| 449. | Metabolism of cyclic nucleotides | **3.07E-06** | 1 |
| 450. | Familial vascular disease | **3.18E-06** | 1 |
| 451. | Morphology of left ventricle | **3.18E-06** | 1 |
| 452. | Tubulation of epithelial tissue | **3.24E-06** | 1 |
| 453. | Action potential of cells | **3.94E-06** | 1 |
| 454. | Growth of nervous tissue | **3.94E-06** | 1 |
| 455. | Atrial fibrillation | **3.96E-06** | 1 |
| 456. | Ovarian cancer | **4.11E-06** | 1 |
| 457. | Quantity of connective tissue cells | **4.17E-06** | 1 |
| 458. | Familial visceral myopathy | **4.25E-06** | 1 |
| 459. | Hereditary spherocytosis | **4.25E-06** | 1 |
| 460. | Differentiation of progenitor cells | **4.40E-06** | 1 |
| 461. | Organization of cytoskeleton | **4.40E-06** | 1 |
| 462. | Vasoconstriction of blood vessel | **4.56E-06** | 1 |
| 463. | Early stage kidney disease | **4.95E-06** | 1 |
| 464. | Quantity of bone cells | **5.05E-06** | 1 |
| 465. | Cerebral small vessel disease | **5.10E-06** | 1 |
| 466. | Migration of vascular smooth muscle cells | **5.10E-06** | 1 |
| 467. | Tubulation of endothelial cells | **5.16E-06** | 1 |
| 468. | Size of cells | **5.25E-06** | 1 |
| 469. | Abnormal morphology of epithelial tissue | **5.38E-06** | 1 |
| 470. | Morphology of gland | **5.64E-06** | 1 |
| 471. | Extraintestinal functional disorder | **5.91E-06** | 1 |
| 472. | Formation of lymphoid tissue | **6.50E-06** | 1 |
| 473. | Invasive tumor | **6.80E-06** | 1 |
| 474. | Morphology of gonad | **6.84E-06** | 1 |
| 475. | Proteinuria | **6.95E-06** | 1 |
| 476. | Assembly of intercellular junctions | **7.41E-06** | 1 |
| 477. | Morphology of muscle cells | **7.68E-06** | 1 |
| 478. | Protein kinase cascade | **7.79E-06** | 1 |
| 479. | Aortic dilatation | **7.95E-06** | 1 |
| 480. | Action potential of neurons | **8.42E-06** | 1 |
| 481. | Transcription | **8.48E-06** | 1 |
| 482. | Familial polycystic kidney disease | **8.95E-06** | 1 |
| 483. | Abnormal morphology of respiratory system | **9.18E-06** | 1 |

| | | | |
|---|---|---|---|
| 484. | Permeability of vascular system | **1.02E-05** | 1 |
| 485. | Morphology of central nervous system | **1.04E-05** | 1 |
| 486. | Morphology of genital organ | **1.12E-05** | 1 |
| 487. | Proliferation of muscle cells | **1.12E-05** | 1 |
| 488. | Symptomatic stage heart failure | **1.16E-05** | 1 |
| 489. | Severe cardiovascular disorder | **1.19E-05** | 1 |
| 490. | Cardiac fibrillation | **1.20E-05** | 1 |
| 491. | Aortic aneurysm | **1.21E-05** | 1 |
| 492. | Congenital hemolytic anemia | **1.22E-05** | 1 |
| 493. | Abnormal morphology of artery | **1.22E-05** | 1 |
| 494. | Development of urinary tract | **1.22E-05** | 1 |
| 495. | Contraction of cardiac muscle | **1.24E-05** | 1 |
| 496. | Biosynthesis of cyclic nucleotides | **1.26E-05** | 1 |
| 497. | Differentiation of smooth muscle | **1.35E-05** | 1 |
| 498. | Formation of mammary gland | **1.36E-05** | 1 |
| 499. | Development of gap junctions | **1.36E-05** | 1 |
| 500. | Pulmonary vascular disease | **1.39E-05** | 1 |
| 501. | Thoracic neoplasm | 1 | **1.25E-23** |
| 502. | Intrathoracic malignant tumor | 1 | **5.48E-23** |
| 503. | Grade 3-4 glioma | 1 | **2.39E-22** |
| 504. | Grade 3-4 tumor | 1 | **3.46E-22** |
| 505. | Cellular infiltration | 1 | **1.39E-21** |
| 506. | Cellular infiltration by blood cells | 1 | **8.25E-20** |
| 507. | Cellular infiltration by leukocytes | 1 | **1.28E-18** |
| 508. | Inflammation of body cavity | 1 | **5.41E-18** |
| 509. | Cell movement of phagocytes | 1 | **1.32E-17** |
| 510. | Activation of blood cells | 1 | **1.65E-17** |
| 511. | Cell movement of granulocytes | 1 | **1.39E-16** |
| 512. | Arteriosclerosis | 1 | **3.00E-16** |
| 513. | Activation of leukocytes | 1 | **3.16E-16** |
| 514. | Recruitment of phagocytes | 1 | **3.03E-15** |
| 515. | Cellular infiltration by myeloid cells | 1 | **3.34E-15** |
| 516. | Cellular infiltration by granulocytes | 1 | **1.21E-14** |
| 517. | Activation of myeloid cells | 1 | **1.35E-14** |
| 518. | Activation of phagocytes | 1 | **1.46E-14** |
| 519. | Homing of blood cells | 1 | **1.69E-14** |
| 520. | Cell movement of neutrophils | 1 | **1.69E-14** |
| 521. | Cell movement of mononuclear leukocytes | 1 | **2.08E-14** |
| 522. | Homing of leukocytes | 1 | **3.43E-14** |
| 523. | Chemotaxis | 1 | **4.31E-14** |
| 524. | Infection of mammalia | 1 | **5.73E-14** |
| 525. | Quantity of lymphoid cells | 1 | **6.51E-14** |
| 526. | Interaction of blood cells | 1 | **1.24E-13** |
| 527. | Quantity of cytokine | 1 | **1.37E-13** |
| 528. | Binding of blood cells | 1 | **1.75E-13** |

| | | | |
|---|---|---|---|
| 529. | Esophageal lesion | 1 | **1.83E-13** |
| 530. | Pregnancy-induced hypertension | 1 | **4.85E-13** |
| 531. | Chemotaxis of leukocytes | 1 | **1.15E-12** |
| 532. | Cell movement of lymphocytes | 1 | **1.59E-12** |
| 533. | Development of non-small cell lung carcinoma | 1 | **1.89E-12** |
| 534. | Adhesion of blood cells | 1 | **2.01E-12** |
| 535. | Nonsquamous non-small cell lung carcinoma | 1 | **2.77E-12** |
| 536. | Interaction of leukocytes | 1 | **2.79E-12** |
| 537. | Malignant brain glioma | 1 | **2.94E-12** |
| 538. | Esophageal cancer | 1 | **3.03E-12** |
| 539. | Malignant neoplasm of aerodigestive tract | 1 | **3.55E-12** |
| 540. | Migration of lymphatic system cells | 1 | **3.58E-12** |
| 541. | Binding of leukocytes | 1 | **3.84E-12** |
| 542. | Chemotaxis of phagocytes | 1 | **4.30E-12** |
| 543. | Accumulation of cells | 1 | **4.54E-12** |
| 544. | Benign Tumors | 1 | **5.43E-12** |
| 545. | Chemotaxis of myeloid cells | 1 | **5.94E-12** |
| 546. | Quantity of neutrophils | 1 | **6.00E-12** |
| 547. | Abnormality of large intestine | 1 | **6.39E-12** |
| 548. | Upper aerodigestive tract carcinoma | 1 | **6.86E-12** |
| 549. | Function of blood cells | 1 | **7.93E-12** |
| 550. | Adhesion of immune cells | 1 | **9.78E-12** |
| 551. | Infiltration by neutrophils | 1 | **1.01E-11** |
| 552. | Accumulation of granulocytes | 1 | **1.20E-11** |
| 553. | Inflammation of gastrointestinal tract | 1 | **1.49E-11** |
| 554. | Recruitment of mononuclear leukocytes | 1 | **1.57E-11** |
| 555. | Esophageal carcinoma | 1 | **3.52E-11** |
| 556. | Gastroenteritis | 1 | **3.93E-11** |
| 557. | Activation of antigen presenting cells | 1 | **4.42E-11** |
| 558. | Allergy | 1 | **5.01E-11** |
| 559. | Cellular infiltration by phagocytes | 1 | **5.10E-11** |
| 560. | Rheumatoid arthritis | 1 | **6.06E-11** |
| 561. | Migration of mononuclear leukocytes | 1 | **6.22E-11** |
| 562. | Cell movement of antigen presenting cells | 1 | **6.95E-11** |
| 563. | Lymphocyte migration | 1 | **6.96E-11** |
| 564. | Cellular infiltration by mononuclear leukocytes | 1 | **7.94E-11** |
| 565. | Infection by SARS coronavirus | 1 | **7.98E-11** |
| 566. | Function of leukocytes | 1 | **9.38E-11** |
| 567. | Accumulation of leukocytes | 1 | **1.05E-10** |
| 568. | Cell movement of macrophages | 1 | **1.41E-10** |
| 569. | COVID-19 | 1 | **1.73E-10** |
| 570. | Accumulation of myeloid cells | 1 | **2.19E-10** |
| 571. | Morphology of blood cells | 1 | **2.23E-10** |

| | | | |
|---|---|---|---|
| 572. | Cell death of blood cells | 1 | **2.24E-10** |
| 573. | Cell death of immune cells | 1 | **2.32E-10** |
| 574. | Quantity of antigen presenting cells | 1 | **3.37E-10** |
| 575. | Binding of professional phagocytic cells | 1 | **3.45E-10** |
| 576. | Survival of organism | 1 | **3.78E-10** |
| 577. | Quantity of interleukin | 1 | **3.99E-10** |
| 578. | Infection by coronavirus | 1 | **4.02E-10** |
| 579. | Homing of mononuclear leukocytes | 1 | **4.72E-10** |
| 580. | Experimental colitis | 1 | **5.00E-10** |
| 581. | Recruitment of antigen presenting cells | 1 | **5.64E-10** |
| 582. | Interaction of phagocytes | 1 | **6.82E-10** |
| 583. | Binding of tumor cell lines | 1 | **7.53E-10** |
| 584. | Accumulation of blood cells | 1 | **8.25E-10** |
| 585. | Recruitment of lymphocytes | 1 | **8.47E-10** |
| 586. | Function of myeloid cells | 1 | **9.13E-10** |
| 587. | Morphology of trabecular bone | 1 | **9.45E-10** |
| 588. | Binding of myeloid cells | 1 | **1.04E-09** |
| 589. | Inflammation of the large intestine | 1 | **1.23E-09** |
| 590. | Enteritis | 1 | **1.27E-09** |
| 591. | Quantity of steroid | 1 | **1.32E-09** |
| 592. | Inflammatory Bowel Disease | 1 | **1.58E-09** |
| 593. | Dextran sodium sulfate-induced colitis | 1 | **1.59E-09** |
| 594. | Migration of phagocytes | 1 | **1.84E-09** |
| 595. | Concentration of triacylglycerol | 1 | **1.87E-09** |
| 596. | Synthesis of lipid | 1 | **1.87E-09** |
| 597. | Ingestion by rodents | 1 | **1.93E-09** |
| 598. | Inflammation of liver | 1 | **2.00E-09** |
| 599. | Fatty acid metabolism | 1 | **2.09E-09** |
| 600. | Acute enteritis | 1 | **2.17E-09** |
| 601. | Apoptosis of blood cells | 1 | **2.28E-09** |
| 602. | Function of phagocytes | 1 | **2.54E-09** |
| 603. | Amyloidosis | 1 | **2.59E-09** |
| 604. | Systolic pressure | 1 | **2.99E-09** |
| 605. | Acute colitis | 1 | **3.33E-09** |
| 606. | Activation of mononuclear leukocytes | 1 | **3.59E-09** |
| 607. | Apoptosis of leukocytes | 1 | **4.16E-09** |
| 608. | Cellular infiltration by lymphocytes | 1 | **4.21E-09** |
| 609. | Colitis | 1 | **4.25E-09** |
| 610. | Binding of lymphocytes | 1 | **4.58E-09** |
| 611. | Esophageal adenocarcinoma | 1 | **5.18E-09** |
| 612. | Homeostasis of ion | 1 | **5.22E-09** |
| 613. | Interaction of lymphocytes | 1 | **5.35E-09** |
| 614. | Interaction of mononuclear leukocytes | 1 | **6.12E-09** |
| 615. | Quantity of immunoglobulin | 1 | **6.46E-09** |
| 616. | Migration of granulocytes | 1 | **6.65E-09** |

| | | | |
|---|---|---|---|
| 617. | Leukopoiesis | 1 | **7.83E-09** |
| 618. | Necrosis | 1 | **8.17E-09** |
| 619. | Lower respiratory tract disorder | 1 | **8.79E-09** |
| 620. | Inflammation of airway | 1 | **9.28E-09** |
| 621. | Metabolism of carbohydrate | 1 | **9.72E-09** |
| 622. | Activation of macrophages | 1 | **1.00E-08** |
| 623. | Apoptosis | 1 | **1.02E-08** |
| 624. | Experimentally-induced arthritis | 1 | **1.02E-08** |
| 625. | Proliferation of mononuclear leukocytes | 1 | **1.19E-08** |
| 626. | Homing of lymphatic system cells | 1 | **1.26E-08** |
| 627. | Concentration of acylglycerol | 1 | **1.29E-08** |
| 628. | Cell movement of T lymphocytes | 1 | **1.30E-08** |
| 629. | Attraction of cells | 1 | **1.34E-08** |
| 630. | Interaction of endothelial cells | 1 | **1.38E-08** |
| 631. | Extravasation of cells | 1 | **1.43E-08** |
| 632. | Healing of lesion | 1 | **1.49E-08** |
| 633. | Chemotaxis of neutrophils | 1 | **1.61E-08** |
| 634. | Sclerotic lesion | 1 | **1.72E-08** |
| 635. | Acute leukemia | 1 | **1.73E-08** |
| 636. | Proliferation of lymphocytes | 1 | **1.80E-08** |
| 637. | Collagen disease | 1 | **1.85E-08** |

**P*-value: Benjamini Hochberg adjusted *P*-value. Bold values: $P < 0.05$n = 13, 16, 8, and 13 for female-DMSO, male-DMSO, female-ITE, and male-ITE rat placentas, respectively.

Table S4. ITE-dysregulated disease-associated biological functions in rat placentas.*

| | *P*-value [Female Placentas] | *P*-value [Male Placentas] |
|---|---|---|
| Liver tumor | **2.56E-17** | **8.34E-21** |
| Liver carcinoma | **2.56E-17** | **5.84E-20** |
| Liver cancer | **4.22E-17** | **5.84E-20** |
| Infarction of heart | **1.17E-10** | **2.28E-11** |
| Disorder of coronary artery | **1.32E-10** | **7.32E-12** |
| Myocardial infarction | **1.32E-10** | **2.46E-11** |
| Dysfunction of heart | **8.46E-10** | **7.20E-13** |
| Congestive heart failure | **2.32E-09** | **2.98E-10** |
| Ventricular dysfunction | **5.02E-09** | **4.65E-12** |
| Left ventricular dysfunction | **1.42E-08** | **2.46E-11** |
| Failure of heart | **1.42E-08** | **1.33E-10** |
| Enlargement of heart ventricle | **2.15E-08** | **1.84E-09** |
| Hypertrophy of heart | **2.15E-08** | **3.35E-08** |
| Enlargement of heart | **2.28E-08** | **6.37E-08** |
| Supraventricular arrhythmia | **3.30E-07** | **3.35E-08** |
| Congenital heart disease | **1.06E-06** | **7.54E-04** |
| Arrhythmia of atrium | **1.13E-06** | **7.65E-08** |
| Coronary occlusion | **1.46E-06** | **4.80E-09** |
| Chronic heart failure | **1.69E-06** | **4.22E-04** |
| Enlargement of left ventricle | **1.75E-06** | **2.65E-07** |
| Atrial fibrillation | **2.60E-06** | **1.72E-07** |
| Coronary artery disease | **2.97E-06** | **1.08E-08** |
| Cardiac fibrillation | **7.69E-06** | **9.43E-08** |
| Symptomatic stage heart failure | **7.69E-06** | **4.35E-04** |
| Acute heart failure | **1.12E-05** | **1.10E-06** |
| Dilated cardiomyopathy | **1.19E-05** | **1.89E-05** |
| Non-ST elevation myocardial infarction | **2.11E-05** | **3.46E-02** |
| Arrhythmia | **2.63E-05** | **4.05E-08** |
| Stable disease stage heart failure | **4.00E-05** | **1.93E-03** |
| Pulmonary Hypertension | **4.14E-05** | **9.91E-04** |
| Ventricular hypertrophy | **4.39E-05** | **9.28E-06** |
| Cardiorenal syndrome | **4.65E-05** | **2.30E-04** |
| Atrial or ventricular septal defect | **4.65E-05** | **5.45E-03** |
| Carditis | **5.22E-05** | **5.37E-05** |
| Glomerulosclerosis | **7.47E-05** | **1.50E-02** |
| Left ventricular dilation | **1.47E-04** | **1.70E-07** |
| Fibrosis of heart | **1.85E-04** | **1.80E-05** |
| Cardiac death | **2.21E-04** | **5.16E-06** |
| Proximal tubular toxicity | **3.21E-04** | **1.10E-06** |
| ST-elevation myocardial infarction | **6.33E-04** | **1.09E-03** |
| Chronic New York Heart Association class II-IV heart failure | **8.67E-04** | **3.01E-02** |

| | | |
|---|---|---|
| Heart failure with preserved ejection fraction | **8.96E-04** | **1.26E-02** |
| Systolic heart failure | **8.96E-04** | **4.05E-02** |
| Hypertrophy of left ventricle | **9.15E-04** | **2.83E-03** |
| Hypertrophy of cardiac muscle | **1.08E-03** | **1.98E-03** |
| Atrial septal defect | **1.08E-03** | **1.38E-02** |
| Hepatocellular carcinoma | **1.11E-03** | **2.76E-05** |
| Fibrosis of kidney | **1.14E-03** | **1.97E-02** |
| Acute myocardial infarction | **1.23E-03** | **3.65E-02** |
| Atrial flutter | **1.23E-03** | **1.59E-02** |
| Hepatic steatosis | **1.24E-03** | **7.76E-07** |
| Spontaneous coronary artery disease | **1.66E-03** | **2.71E-02** |
| Ventricular septal defect | **1.69E-03** | **2.65E-02** |
| Atrophy of kidney | **1.72E-03** | **1.98E-02** |
| Increased Levels of Hematocrit | **1.75E-03** | **4.25E-03** |
| Activation of hepatic stellate cells | **1.92E-03** | **2.42E-02** |
| Systolic dysfunction | **2.22E-03** | **8.31E-03** |
| Autosomal recessive renal tubular dysgenesis | **2.33E-03** | **4.84E-02** |
| New York Heart Association class IV heart failure | **2.33E-03** | **4.84E-02** |
| Hypertrophy of heart cells | **2.38E-03** | **3.75E-03** |
| Hypertrophy of cardiomyocytes | **2.95E-03** | **3.46E-03** |
| Metastatic liver tumor | **3.15E-03** | **3.01E-02** |
| Injury of kidney | **3.19E-03** | **2.89E-04** |
| Diastolic dysfunction | **3.73E-03** | **3.71E-03** |
| Stenosis of cardiac valve | **4.50E-03** | **1.10E-05** |
| Diastolic dysfunction of heart | **4.50E-03** | **2.30E-04** |
| Myocarditis | **4.50E-03** | **4.72E-04** |
| Acute congestive heart failure | **4.50E-03** | **8.09E-03** |
| Stenosis of mitral valve | **4.50E-03** | **8.09E-03** |
| Liver cholangiocarcinoma | **4.50E-03** | **1.95E-02** |
| Hydronephrosis | **4.50E-03** | **2.17E-02** |
| Left ventricular diastolic dysfunction | **4.89E-03** | **1.07E-04** |
| Chronic renal failure | **4.92E-03** | **8.31E-03** |
| End stage renal disease | **5.16E-03** | **1.20E-02** |
| Bradycardia | **5.34E-03** | **4.64E-02** |
| Decreased Levels of Albumin | **6.44E-03** | **1.41E-03** |
| Tachycardia | **6.96E-03** | **6.11E-04** |
| Fibrosis of liver | **7.40E-03** | **4.06E-06** |
| Cardiotoxicity | **7.40E-03** | **8.31E-03** |
| Ventricular tachycardia | **7.49E-03** | **2.34E-04** |
| Experimental autoimmune myocarditis | **7.49E-03** | **5.99E-04** |
| Failure of kidney | **7.50E-03** | **1.49E-04** |
| Enlargement of right ventricle | **7.70E-03** | **1.98E-03** |
| Nonalcoholic fatty liver disease | **8.00E-03** | **8.43E-03** |
| Arrhythmia of heart ventricle | **8.34E-03** | **6.18E-04** |
| Cirrhosis | **8.75E-03** | **4.66E-05** |

| | | |
|---|---|---|
| Stable disease stage cardiomyopathic heart failure | **8.75E-03** | **1.38E-02** |
| Stable disease stage hypertensive heart failure | **8.75E-03** | **1.38E-02** |
| Stable disease stage ischemic heart failure | **8.75E-03** | **1.38E-02** |
| Symptomatic stage cardiomyopathic heart failure | **8.75E-03** | **1.38E-02** |
| Symptomatic stage hypertensive heart failure | **8.75E-03** | **1.38E-02** |
| Symptomatic stage ischemic heart failure | **8.75E-03** | **1.38E-02** |
| Decreased excretion of albumin | **9.88E-03** | **2.16E-02** |
| Congenital left heart disease | **9.90E-03** | **5.90E-03** |
| Familial arrhythmia | **9.93E-03** | **4.76E-03** |
| Damage of kidney | **1.17E-02** | **2.30E-04** |
| Dysfunction of left ventricle | **1.41E-02** | **2.04E-05** |
| Familial isolated autosomal dominant noncompaction of left ventricular myocardium | **1.53E-02** | **2.71E-02** |
| Chronic hepatitis C | **1.68E-02** | **2.01E-04** |
| Progressive hepatocellular carcinoma | **1.81E-02** | **4.35E-02** |
| Dilation of left ventricle | **1.86E-02** | **4.72E-04** |
| Permanent atrial fibrillation | **1.86E-02** | **8.28E-03** |
| Child-Pugh class A hepatocellular carcinoma | **1.86E-02** | **1.98E-02** |
| Right ventricular dilatation | **1.99E-02** | **3.41E-03** |
| Liver Regeneration | **1.99E-02** | **1.52E-02** |
| Multi vessel coronary artery disease | **1.99E-02** | **3.01E-02** |
| Early stage myocardial infarction | **1.99E-02** | **3.01E-02** |
| Hyperplasia of cardiomyocytes | **1.99E-02** | **3.01E-02** |
| Interauricular communication | **1.99E-02** | **3.01E-02** |
| Perioperative tachycardia | **1.99E-02** | **3.01E-02** |
| Chemotaxis of hepatic stellate cells | **1.99E-02** | **3.40E-02** |
| Stenosis of semilunar valve | **2.14E-02** | **9.86E-05** |
| Recurrent myocardial infarction | **2.50E-02** | **4.41E-02** |
| Inflammation of liver | **2.75E-02** | **1.97E-09** |
| Cardiomyopathy of left ventricle | **2.75E-02** | **2.65E-02** |
| Inherited arrhythmia syndrome | **2.84E-02** | **3.41E-03** |
| Familial dilated cardiomyopathy | **3.01E-02** | **4.64E-02** |
| Cirrhosis of liver | **3.15E-02** | **3.88E-03** |
| Chronic viral hepatitis | **3.20E-02** | **7.13E-04** |
| Hypoplasia of renal medulla | **3.20E-02** | **4.84E-02** |
| Susceptibility to atrioventricular septal defect type 1 | **3.20E-02** | **4.84E-02** |
| Life-threatening sustained ventricular arrhythmia | **3.20E-02** | **4.84E-02** |
| Severe ventricular tachycardia | **3.20E-02** | **4.84E-02** |
| Severely symptomatic disease stage atrial fibrillation | **3.20E-02** | **4.84E-02** |
| Severely symptomatic disease stage atrial flutter | **3.20E-02** | **4.84E-02** |
| Familial atrial fibrillation | **3.61E-02** | **2.12E-02** |
| Stenosis of aortic valve | **3.62E-02** | **1.56E-04** |
| Primary dilated cardiomyopthy | **3.62E-02** | **4.46E-02** |
| Proliferation of liver cells | **4.42E-02** | **1.01E-02** |
| Brugada syndrome | **4.42E-02** | **3.46E-02** |

| | | |
|---|---|---|
| Dilated cardiomyopathy 1A | **4.58E-02** | **7.29E-03** |
| Unresectable hepatocellular carcinoma | **4.58E-02** | **4.63E-02** |
| Ventricular fibrillation | **4.66E-02** | **2.61E-02** |
| Bicuspid aortic valve | 5.19E-02 | **3.75E-03** |
| Familial atrial septal defect | 5.98E-02 | **2.65E-02** |
| Left ventricular systolic dysfunction | 5.98E-02 | **2.65E-02** |
| Acute myocarditis | 6.06E-02 | **9.26E-04** |
| Migration of hepatic stellate cells | 6.06E-02 | **2.79E-03** |
| Right ventricular hypertrophy | 6.06E-02 | **5.75E-03** |
| Child-Pugh class A advanced hepatocellular carcinoma | 6.06E-02 | **1.20E-02** |
| Fibrosis of myocardium | 6.06E-02 | **1.20E-02** |
| Longstanding persistent atrial fibrillation | 6.61E-02 | **3.01E-02** |
| Recurrent ventricular fibrillation | 6.61E-02 | **3.01E-02** |
| Refractory hemodynamically unstable ventricular tachycardia | 6.61E-02 | **3.01E-02** |
| Refractory ventricular fibrillation | 6.61E-02 | **3.01E-02** |
| Acute phase crescentic glomerulonephritis | 8.37E-02 | **2.30E-04** |
| Crescentic glomerulonephritis | 8.37E-02 | **3.22E-04** |
| Chronic hepatitis | 8.37E-02 | **1.86E-03** |
| Pulmonary embolism | 8.37E-02 | **5.74E-03** |
| Growth of liver tumor | 8.37E-02 | **1.62E-02** |
| Familial left ventricular noncompaction | 8.53E-02 | **2.65E-02** |
| Persistent atrial fibrillation | 8.53E-02 | **2.65E-02** |
| Nephritis | 9.06E-02 | **1.29E-02** |
| Hereditary Wolff-Parkinson-White syndrome | 9.13E-02 | **5.47E-03** |
| Primary biliary cirrhosis | 9.46E-02 | **1.45E-02** |
| Necrosis of cardiac muscle | 9.67E-02 | **1.57E-02** |
| Bleeding of liver | 9.81E-02 | **3.12E-02** |
| Dominant dilated cardiomyopathy | 9.81E-02 | **3.12E-02** |
| Viral hepatitis | 1.08E-01 | **2.48E-04** |
| Increased Levels of Creatinine | 1.29E-01 | **2.67E-03** |
| Digenic midventricular hypertrophic cardiomyopathy | 1.29E-01 | **1.38E-02** |
| Ethynylestradiol hepatocellular cholestasis | 1.29E-01 | **1.38E-02** |
| Proliferation of glomerular cells | 1.29E-01 | **4.33E-02** |
| Proliferation of hepatocytes | 1.33E-01 | **4.35E-02** |
| Perimembranous ventricular septal defect | 1.36E-01 | **3.48E-02** |
| Apoptosis of kidney cells | 1.43E-01 | **3.34E-02** |
| Glomerulonephritis | 1.46E-01 | **6.37E-03** |
| Cardiac decompensation | 1.56E-01 | **4.54E-02** |
| Proliferation of mesangial cells | 1.65E-01 | **4.84E-02** |
| Cell death of kidney cells | 1.66E-01 | **2.65E-02** |
| Idiopathic ventricular fibrillation | 1.66E-01 | **3.01E-02** |
| Primary liver cholangiocarcinoma | 1.66E-01 | **3.01E-02** |
| Regeneration of myocardium | 1.66E-01 | **3.01E-02** |
| Cell death of cardiomyocytes | 1.66E-01 | **3.32E-02** |
| Hepatic injury | 1.79E-01 | **4.25E-03** |

| | | |
|---|---|---|
| Liver Damage | 1.92E-01 | **2.30E-04** |
| Alcoholic hepatitis | 1.99E-01 | **4.72E-04** |
| Obstruction of bile duct | 1.99E-01 | **4.84E-02** |
| Decreased secretion of albumin | 1.99E-01 | **4.84E-02** |
| Lipopolysaccharide hepatocellular cholestasis | 1.99E-01 | **4.84E-02** |
| Steatohepatitis | 2.40E-01 | **6.35E-03** |
| Apoptosis of hepatic stellate cells | 3.00E-01 | **2.85E-02** |
| Severe alcoholic hepatitis | 3.43E-01 | **1.20E-02** |
| Alcoholic steatohepatitis | 3.80E-01 | **7.52E-03** |
| Damage of heart | 3.91E-01 | **8.31E-03** |
| Infection by Hepatitis C virus | 1.00E+00 | **1.88E-04** |
| Dilation of heart chamber | 1.00E+00 | **4.46E-04** |
| Dilation of heart ventricle | 1.00E+00 | **6.39E-04** |
| Calcification of aortic valve | 1.00E+00 | **5.74E-03** |
| Severely symptomatic disease stage atrial fibrillation or flutter | 1.00E+00 | **8.09E-03** |
| Hepatic adenocarcinoma | 1.00E+00 | **1.20E-02** |
| Syndromic arrhythmia | 1.00E+00 | **1.26E-02** |
| Advanced stage hepatic steatosis | 1.00E+00 | **3.01E-02** |
| Calcific aortic valve stenosis | 1.00E+00 | **3.40E-02** |
| Sudden cardiac death | 1.00E+00 | **4.32E-02** |
| Pericarditis | **4.43E-04** | 8.76E-02 |
| Single ventricle defect | **5.13E-04** | 7.25E-02 |
| Metastatic liver cholangiocarcinoma | **8.96E-04** | 9.14E-02 |
| New York Heart Association class II heart failure | **4.50E-03** | 7.00E-02 |
| Postmyocardial infarction syndrome | **4.50E-03** | 7.00E-02 |
| Symptomatic stage chronic systolic heart failure | **4.50E-03** | 7.00E-02 |
| Increased Levels of Red Blood Cells | **4.50E-03** | 1.12E-01 |
| Autosomal dominant pulmonary arterial hypertension | **4.89E-03** | 5.22E-02 |
| Interstitial fibrosis of kidney | **5.43E-03** | 1.48E-01 |
| Familial primary pulmonary hypertension | **6.60E-03** | 6.44E-02 |
| Spontaneous coronary artery dissection | **7.40E-03** | 9.02E-02 |
| Common atrium | **7.40E-03** | 9.02E-02 |
| New York Heart Association class III heart failure | **7.40E-03** | 9.02E-02 |
| Congestion of liver | **8.75E-03** | 1.49E-01 |
| Susceptibility to progression to renal failure in IgA nephropathy | **8.75E-03** | 1.49E-01 |
| Advanced liver tumor | **9.30E-03** | 7.35E-02 |
| 5Increased excretion of albumin | **1.02E-02** | 9.83E-02 |
| Pulmonary arterial hypertension associated with congenital heart disease | **1.02E-02** | 9.83E-02 |
| Hypoplasia of kidney | **1.25E-02** | 1.37E-01 |
| BCLC stage B hepatocellular carcinoma | **1.60E-02** | 9.83E-02 |
| Dysplasia of kidney | **1.86E-02** | 2.02E-01 |
| Paroxysmal supraventricular tachycardia | **1.86E-02** | 2.02E-01 |
| Familial pulmonary hypertension | **1.99E-02** | 9.97E-02 |
| Atrophy of renal medulla | **1.99E-02** | 1.31E-01 |

| | | |
|---|---|---|
| Primary pulmonary hypertension 1 | **1.99E-02** | 1.31E-01 |
| Perivascular fibrosis of kidney | **1.99E-02** | 1.89E-01 |
| Hepatoblastoma | **2.04E-02** | 1.89E-01 |
| Increased clearance of creatinine | **2.09E-02** | 1.13E-01 |
| Interstitial nephritis | **2.24E-02** | 2.30E-01 |
| Proliferation of hepatic stellate cells | **2.50E-02** | 1.13E-01 |
| Atrophy of renal glomerulus | **2.50E-02** | 1.48E-01 |
| Unresectable liver cholangiocarcinoma | **2.50E-02** | 2.92E-01 |
| Sustained ventricular tachycardia | **2.84E-02** | 1.37E-01 |
| Symptomatic stage atrial fibrillation | **2.84E-02** | 1.37E-01 |
| Symptomatic stage atrial flutter | **2.84E-02** | 1.37E-01 |
| Ann Arbor stage I-IV hepatosplenic T-cell lymphoma | **2.84E-02** | 3.07E-01 |
| Idiopathic and/or familial pulmonary arterial hypertension | **2.96E-02** | 1.30E-01 |
| Failure of heart looping | **3.11E-02** | 1.37E-01 |
| Type 1 hepatorenal syndrome | **3.11E-02** | 1.54E-01 |
| Stable coronary artery disease | **3.20E-02** | 2.30E-01 |
| Cell death of left ventricle | **3.20E-02** | 1.00E+00 |
| Familial glomerulopathy | **3.31E-02** | 2.76E-01 |
| Atrophy of renal tubule | **3.61E-02** | 1.49E-01 |
| Cell proliferation of kidney cell lines | **3.62E-02** | 9.83E-02 |
| Atrophy of renal cortex | **3.62E-02** | 1.73E-01 |
| Acute kidney injury | **4.11E-02** | 1.09E-01 |
| Small cell undifferentiated hepatoblastoma | **4.11E-02** | 3.63E-01 |
| Thrombosis of atrium | **4.42E-02** | 7.35E-02 |
| Sinus arrhythmia | **4.42E-02** | 7.35E-02 |
| Severe ventricular arrhythmia | **4.42E-02** | 9.14E-02 |
| Stenosis of pulmonary artery | **4.66E-02** | 7.00E-02 |
| Familial sick sinus syndrome | **4.66E-02** | 7.00E-02 |
| Acute-on-chronic liver failure | **4.66E-02** | 2.72E-01 |
| Perivascular fibrosis | **4.77E-02** | 1.70E-01 |

**P*-value: Benjamini Hochberg adjusted *P*-value. Bold values: $P < 0.05$. n = 13, 16, 8, and 13 for female-DMSO, male-DMSO, female-ITE, and male-ITE rat placentas, respectively.

Table S5. ITE-dysregulated canonical pathways in rat placentas.*

| | *P*-value [Female Placentas] | BH *P*-value [Male Placentas] |
|---|---|---|
| Phagosome Formation | **7.13E-04** | **4.72E-08** |
| G-Protein Coupled Receptor Signaling | **1.08E-02** | **4.27E-06** |
| CREB Signaling in Neurons | **5.06E-05** | **5.93E-06** |
| Cellular Effects of Sildenafil (Viagra) | **1.29E-04** | **1.18E-05** |
| Axonal Guidance Signaling | **5.59E-07** | **1.69E-05** |
| cAMP-mediated signaling | **6.55E-03** | **1.69E-05** |
| Cardiac Hypertrophy Signaling (Enhanced) | **6.55E-03** | **2.97E-05** |
| LPS/IL-1 Mediated Inhibition of RXR Function | **2.41E-02** | **3.68E-05** |
| Gap Junction Signaling | **8.93E-05** | **3.89E-05** |
| Leptin Signaling in Obesity | **7.14E-03** | **4.80E-05** |
| Breast Cancer Regulation by Stathmin1 | **2.86E-05** | **5.26E-05** |
| Neuropathic Pain Signaling In Dorsal Horn Neurons | **4.09E-02** | **1.42E-04** |
| Sperm Motility | **3.70E-02** | **1.79E-04** |
| Hepatic Fibrosis Signaling Pathway | **3.19E-02** | **4.10E-04** |
| Adrenomedullin signaling pathway | **7.14E-03** | **4.13E-04** |
| Gαi Signaling | **2.41E-02** | **6.12E-04** |
| Sphingosine-1-phosphate Signaling | **2.18E-02** | **8.50E-04** |
| Gustation Pathway | **6.55E-03** | **1.03E-03** |
| Dilated Cardiomyopathy Signaling Pathway | **1.18E-03** | **1.08E-03** |
| Actin Cytoskeleton Signaling | **2.18E-02** | **1.24E-03** |
| Iron homeostasis signaling pathway | **1.93E-04** | **1.55E-03** |
| Tight Junction Signaling | **3.14E-02** | **1.55E-03** |
| Factors Promoting Cardiogenesis in Vertebrates | **9.67E-03** | **3.11E-03** |
| GP6 Signaling Pathway | **3.33E-02** | **4.06E-03** |
| ILK Signaling | **2.18E-02** | **5.65E-03** |
| Thrombin Signaling | **2.18E-02** | **6.25E-03** |
| Heparan Sulfate Biosynthesis (Late Stages) | **2.24E-02** | **6.25E-03** |
| GPCR-Mediated Integration of Enteroendocrine Signaling Exemplified by an L Cell | **2.32E-02** | **6.79E-03** |
| Sertoli Cell-Sertoli Cell Junction Signaling | **5.90E-03** | **1.12E-02** |
| Heparan Sulfate Biosynthesis | **4.17E-02** | **1.63E-02** |
| GPCR-Mediated Nutrient Sensing in Enteroendocrine Cells | **4.24E-02** | **2.56E-02** |
| Protein Kinase A Signaling | **2.24E-02** | **2.73E-02** |
| Calcium Signaling | **1.49E-02** | **3.11E-02** |
| Human Embryonic Stem Cell Pluripotency | **4.21E-02** | **3.63E-02** |
| VDR/RXR Activation | **4.57E-02** | **3.64E-02** |
| Notch Signaling | **4.21E-02** | 1.46E-01 |
| Coronavirus Replication Pathway | **3.14E-02** | 2.47E-01 |
| Caveolar-mediated Endocytosis Signaling | **4.21E-02** | 2.67E-01 |
| Atherosclerosis Signaling | 1.21E-01 | **1.89E-07** |

| | | |
|---|---|---|
| Role of Pattern Recognition Receptors in Recognition of Bacteria and Viruses | 5.07E-01 | **1.69E-05** |
| Role of Hypercytokinemia/hyperchemokinemia in the Pathogenesis of Influenza | 2.60E-01 | **4.80E-05** |
| FXR/RXR Activation | 1.42E-01 | **1.48E-04** |
| LXR/RXR Activation | 1.88E-01 | **2.03E-04** |
| Maturity Onset Diabetes of Young (MODY) Signaling | 2.11E-01 | **9.72E-04** |
| Insulin Secretion Signaling Pathway | 6.32E-02 | **1.43E-03** |
| Complement System | 7.30E-01 | **1.45E-03** |
| Triacylglycerol Degradation | 6.30E-02 | **1.52E-03** |
| Phospholipases | 1.78E-01 | **1.55E-03** |
| HMGB1 Signaling | 2.77E-01 | **1.64E-03** |
| TREM1 Signaling | 5.47E-01 | **1.91E-03** |
| Neuroinflammation Signaling Pathway | 1.00E+00 | **2.33E-03** |
| Role of Macrophages, Fibroblasts and Endothelial Cells in Rheumatoid Arthritis | 2.14E-01 | **2.41E-03** |
| Acute Phase Response Signaling | 4.18E-01 | **2.76E-03** |
| Glioblastoma Multiforme Signaling | 1.46E-01 | **3.06E-03** |
| IL-17 Signaling | 2.93E-01 | **3.08E-03** |
| Synaptic Long Term Depression | 1.23E-01 | **3.22E-03** |
| Hepatic Cholestasis | 7.02E-02 | **3.38E-03** |
| Glioma Invasiveness Signaling | 8.24E-01 | **3.62E-03** |
| Endothelin-1 Signaling | 1.95E-01 | **3.74E-03** |
| Neuroprotective Role of THOP1 in Alzheimer's Disease | 2.60E-01 | **5.65E-03** |
| Aldosterone Signaling in Epithelial Cells | 5.97E-01 | **5.65E-03** |
| Role of MAPK Signaling in Inhibiting the Pathogenesis of Influenza | 5.57E-01 | **5.96E-03** |
| Relaxin Signaling | 1.10E-01 | **6.20E-03** |
| Apelin Cardiomyocyte Signaling Pathway | 1.44E-01 | **6.80E-03** |
| Eicosanoid Signaling | 1.37E-01 | **9.98E-03** |
| eNOS Signaling | 7.86E-02 | **1.03E-02** |
| HIF1α Signaling | 1.48E-01 | **1.12E-02** |
| Differential Regulation of Cytokine Production in Macrophages and T Helper Cells by IL-17A and IL-17F | 7.75E-01 | **1.28E-02** |
| Aryl Hydrocarbon Receptor Signaling | 1.82E-01 | **1.47E-02** |
| Renin-Angiotensin Signaling | 1.82E-01 | **1.75E-02** |
| Antioxidant Action of Vitamin C | 6.56E-01 | **1.82E-02** |
| Cardiac Hypertrophy Signaling | 1.42E-01 | **1.84E-02** |
| MSP-RON Signaling Pathway | 3.15E-01 | **1.95E-02** |
| Nitric Oxide Signaling in the Cardiovascular System | 6.30E-02 | **2.05E-02** |
| IL-8 Signaling | 2.31E-01 | **2.26E-02** |
| Role of MAPK Signaling in the Pathogenesis of Influenza | 5.87E-01 | **2.33E-02** |
| CXCR4 Signaling | 1.44E-01 | **2.41E-02** |
| IL-15 Production | 2.60E-01 | **2.56E-02** |
| Xenobiotic Metabolism Signaling | 2.14E-01 | **2.57E-02** |
| Inflammasome pathway | 1.00E+00 | **2.70E-02** |

| | | |
|---|---|---|
| Fcγ Receptor-mediated Phagocytosis in Macrophages and Monocytes | 6.69E-01 | **2.97E-02** |
| Retinol Biosynthesis | 1.30E-01 | **2.97E-02** |
| Macropinocytosis Signaling | 1.78E-01 | **2.97E-02** |
| Xenobiotic Metabolism AHR Signaling Pathway | 2.60E-01 | **2.97E-02** |
| NRF2-mediated Oxidative Stress Response | 5.87E-01 | **2.97E-02** |
| Clathrin-mediated Endocytosis Signaling | 3.35E-01 | **3.16E-02** |
| Bladder Cancer Signaling | 1.27E-01 | **3.46E-02** |
| PPARα/RXRα Activation | 1.60E-01 | **3.46E-02** |
| Integrin Signaling | 5.55E-02 | **3.64E-02** |
| Crosstalk between Dendritic Cells and Natural Killer Cells | 4.81E-01 | **3.77E-02** |
| Airway Inflammation in Asthma | 2.30E-01 | **4.09E-02** |
| Ovarian Cancer Signaling | 5.26E-01 | **4.09E-02** |
| Coagulation System | 1.00E+00 | **4.09E-02** |

**P*-value: Benjamini Hochberg adjusted *P*-value. Bold *P*-value: < 0.05. n = 13, 16, 8, and 13 for female-DMSO, male-DMSO, female-ITE, and male-ITE rat placentas, respectively.

Table S6. ITE-dysregulated gene networks in rat placentas.*

| Gene Network | Female Placentas | | Male Placentas | |
|---|---|---|---|---|
| | *P*-value (n) | Target Genes | *P*-value (n) | Target Genes |
| DNMT3B-regulated genes | 5.38E-06 (22) | ADCY5, ADRB1, AGTR1, BIK, CDC25C, CMTM5, CYP1B1, FOXP3, GUCY1A1, HOXA13, LRRC34, MID1, MYH6, MYL3, MYL4, MYL9, PRKCB, RCAN2, RGS4, S1PR1, ST14, TRPC3 | 6.69E-05 (23) | ADCY5, ADRB1, AGTR1, CCL5, CDKN2A, CKM, CYP1B1, DNMT3B, FOXP3, GUCY1A1, HOXA13, MYH6, MYH7, MYL4, MYL9, PAX6, PNPLA3, PPP1R1A, PRKCB, S1PR1, SLC8A2, ST14, TRPC3 |
| CDKN2A-regulated genes | 3.58E-02 (20) | ACTA2, Ccl2, CCL2, CCL3L3, Ccl7, CCNA2, CCND1, CDC25B, CDC25C, CEBPD, CXCL10, E2F2, EBF1, EGR1, EGR2, FST, IL36B, PTGS2, SOX17, TP63 | 3.98E-03 (28) | ACTA2, Ccl2, CCL2, CCL3L3, CCL4, CCL5, Ccl7, Ccl9, CCNA2, CDC25B, CDCA5, CDKN2A, CXCL10, Cxcl9, E2F2, EBF1, EGR1, EGR2, FST, HOXB9, IL1B, IL36B, IL6, MMP9, PSMB9, PTGS2, SOX17, TP63 |
| FOXF2-regulated genes | 5.81E-06 (8) | ACTA2, ACTG2, CCND1, FOXF1, MYH11, MYOCD, PDGFB, TAGLN | 2.60E-04 (7) | ACTA2, ACTG2, FOXF1, MYH11, MYOCD, PDGFB, TAGLN |
| FOXO3-regulated genes | 1.49E-05 (32) | Acan, ACTA2, ANGPT1, BMP2, CAV1, CCND1, CCND2, CLDN1, CXCL10, CXCR4, EGR1, EGR2, ERG, FASLG, FLT4, FOXP3, IGFBP1, IGFBP7, IL1R1, KIF15, MCAM, MMP13, MYOCD, PDE3B, PTPRB, Rcan1, TAL1, TCIM, TIE1, TP63, TRIM63, UBE2C | 2.29E-05 (37) | Acan, ACTA2, ANGPT1, CAV1, CCND2, CLDN1, CXCL10, CXCR4, DOK5, EGR1, EGR2, ERG, FASLG, FLT4, FOXP3, GABRR2, IFNB1, IGFBP7, IL1R1, IL6, KIF15, MCAM, MMP13, MMP9, MYOCD, NCAPD2, PDE3B, PTPRB, Rcan1, TAL1, TCIM, TFF1, TIE1, TNFSF10, TP63, TRIM63, VCAM1 |
| FOXO4-regulated genes | 1.02E-03 (13) | Acan, ANGPT1, CAV1, CCND2, ERG, FASLG, FLT4, IGFBP1, IGFBP7, IL1R1, MAFA, PTPRB, TIE1 | 7.65E-03 (13) | Acan, ANGPT1, CAV1, CCND2, ERG, FASLG, FLT4, IGFBP7, IL1R1, MMP9, PTPRB, TIE1, VCAM1 |
| IFI16-regulated genes | 1.20E-02 (8) | CCL2, CCND1, CD28, CXCL10, CYP1A1, E2F2, IL1RN, IL7R | 1.27E-06 (16) | CCL2, CCL20, CCL5, CDKN2A, CGA, CXCL10, CYP1A1, E2F2, IFI16, IFNB1, IL1RN, IL6, IL7R, ISG15, NKX2-5, VCAM1 |
| SPI1-regulated genes | 7.77E-04 (66) | ACTA2, C3AR1, CCNA2, CCND1, CCND2, CD72, CHIT1, CTSE, | 1.30E-12 (80) | ACTA2, C1QC, C3AR1, CCL4, CCNA2, CCND2, CD14, CD72, |

| | | | | |
|---|---|---|---|---|
| | | CXCL10, E2F2, EBF1, EGR2, FASLG, FLI1, GATA1, ID3, IL1RN, IL24, IL7R, KLF13, MX1, NCF4, NEIL1, PRG2, PTGS2 | | CHI3L1, CHIT1, CTSE, CXCL10, CXCL17, Cxcl9, CYBB, E2F2, EBF1, EGR2, FASLG, FCER1G, FUT7, GATA1, HK3, ID3, IFIT1B, IFIT2, IFIT3, IL1B, IL1RN, IL24, IL7R, ISG15, LILRB3, MX1, NCF1, NCF4, NEIL1, NKX2-5, OASL, OSCAR, PIK3CG, PRG2, PSMB9, PTCRA, PTGS2, SPI1, SPP1, TNFSF10, TREM1 |
| EHF-regulated genes | 4.64E-02 (8) | BMP4, CYP1A1, GATA1, GRHL3, IL1RN, NOTCH3, THY1, TNC | 4.82E-04 (14) | ALOX5, BMP4, CCL20, CDKN2A, COL2A1, CYP1A1, GATA1, GRHL3, IL1RN, NOTCH3, SCEL, THY1, TIMP1, TNC |
| HEY2-regulated genes | 7.47E-07 (11) | ACTA2, BMP2, CNN1, GJA5, IL34, MYH11, MYH6, MYL4, SCN5A, TAGLN, TBX5 | 1.26E-07 (13) | ACTA2, GJA5, IL34, IL6, MYH11, MYH6, MYH7, MYL4, NKX2-5, SCN5A, SLC6A3, TAGLN, TBX5 |
| KDM5B-regulated genes | 5.27E-03 (13) | CAV1, CCND1, CLDN1, ECT2, EGR1, ERBB3, FST, GAL, MCAM, PDE3B, PRPS1, TAL1, TUBB2A | 6.45E-03 (15) | CAV1, CLDN1, ECT2, EGR1, ERBB3, FGFR2, FST, GJB2, ISG15, MCAM, NKX2-5, PDE3B, PRPS1, SCNN1A, TAL1 |
| MYF6-regulated genes | 6.18E-03 (11) | Ccl2, CXCL10, DPP4, EGF, FGF9, FST, IGFBP3, IL1RN, OSM, PDGFB, TNFRSF11B | 6.20E-05 (17) | BDNF, Ccl2, CKM, CXCL10, Cxcl9, DPP4, FGF7, FGF9, FST, GDF11, IGFBP3, IL1RN, IL6, OSM, PDGFB, SPP1, THBS1 |
| NRIP1-regulated genes | 7.28E-03 (8) | AIFM2, CCND1, CYP1A1, FABP3, PTGS2, SLC16A10, SMAD6, SUOX | 2.11E-04 (12) | AIFM2, CDKN2A, CIDEA, CYP1A1, FABP3, IFNB1, IL1B, IL6, PTGS2, SLC16A10, SMAD6, SUOX |
| HEY1-regulated genes | 7.92E-04 (6) | ACTA2, BMP2, CNN1, MYH11, MYH6, TAGLN | | N/A |
| AHR-regulated genes | 1.93E-09 (48) | ACTA2, AHRR, ATOH8, Ccl2, CCNA2, CCND1, CD28, CDC25B, CDC25C, CHST2, COL11A1, COL13A1, CXCL10, CYP1A1, CYP1B1, DCLK3, EBF1, EGR2, FASLG, FBLN5, FBN2, FGF9, FOXP3, GRAP2, HAVCR2, HEY2, | 2.75E-13 (64) | ACTA2, AHRR, ALOX5, ATOH8, C1QB, C1QC, CARD11, CBR1, Ccl2, CCL20, CCL5, CCNA2, CD14, CDC25B, CDKN2A, CHI3L1, CHST2, COL11A1, COL13A1, COL14A1, CXCL10, CXCL11, CXCL2, Cxcl9, CYP1A1, CYP1B1, DCLK3, EBF1, |

| | | | | |
|---|---|---|---|---|
| | | IGFBP1, IL1R1, IL34, IL7R, KLF2, NOTCH3, NPTX1, NQO1, NXPE2, OLFML1, PDGFB, PKLR, PMP22, PTGS2, RBP1, SEMA3A, SERPINA12, SLAMF8, SLC16A10, STEAP4, TGFBI, TUBA8 | | EGR2, FASLG, FBLN5, FBN2, FGF9, FGFR2, FOXP3, GDF15, GRAP2, HAVCR2, HEY2, Hsd3b4, IL1B, IL1R1, IL34, IL6, IL7R, MYH7, NOTCH3, NPTX1, NQO1, OLFML1, PDGFB, PKLR, PMP22, PTGS2, RBP1, SDR9C7, SEMA3A, SERPINA12, SLAMF8, SLC16A10, SPP1, TFF1, TGFBI, THBS1 |
| RARB-regulated genes | 3.90E-02 (13) | ATP6V0A4, BCL2A1, CHGA, EGR1, ENOX1, ID1, ID3, IGFBP3, IGFBP6, KRT1, LYZL1, OXT, PTGS2 | 2.16E-04 (22) | AIM2, ATP6V0A4, BCL2A1, CDKN2A, CHGA, CXCL2, Cyp2c70, EGR1, GBP2, ID1, ID3, IFIT1B, IGFBP3, IL1B, IL6, Ly6g6e, PLA2G1B, PTGS2, SPP1, Tdh, UBA7, ZMAT4 |
| ELANE-regulated genes | | N/A | 1.15E-04 (9) | C5AR1, CCL3L3, CXCL2, ELN, IL1B, IL6, MMP9, OSM, SPI1 |
| USP22-regulated genes | 2.06E-03 (7) | BCL2A1, CCND2, CLEC5A, IL3RA, LYL1, NCF4, OLFM4 | 2.36E-07 (13) | BCL2A1, CCL5, CCND2, CHI3L1, CLEC5A, CYBB, IFNB1, LYL1, MSR1, NCF1, NCF4, OLFM4, SPI1 |
| CBX5-regulated genes | 2.59E-02 (10) | CYP1B1, GRHL3, H2AC18/H2AC19, HBB, HOXA13, ICAM2, PRSS23, TACSTD2, TGFBI, TIMP4 | 1.39E-03 (15) | CKM, CYP1B1, GRHL3, HBB, HOXA13, ICAM2, MAL2, PRR15L, PRSS23, TACSTD2, TGFBI, TIMP4, TM4SF1, TRIM31, TSPAN13 |
| ETS1-regulated genes | 5.86E-03 (18) | BMP4, CAV1, Ccl2, CCL2, CCND1, EGR1, FASLG, FLI1, FOXP3, IL24, IL7R, ITGA2B, ITGB6, Klrk1, MCAM, MMP13, NQO1, THY1 | 1.91E-05 (28) | APOC2, BMP4, CAV1, Ccl2, CCL2, CCL5, CD14, CD69, CDKN2A, COL2A1, EGR1, FASLG, FOXP3, GP9, GZMA, IL24, IL7R, Klrk1, MCAM, MMP13, MMP9, MSR1, NCF1, NQO1, SPI1, SPP1, TFF1, THY1 |
| IRF1-regulated genes | 3.90E-02 (13) | CCL2, CCND1, CXCL10, CXCL3, FASLG, FOXP3, MX1, NFE2, PDCD1LG2, PLAAT1, PTGS2, TBX3, UBD | 1.76E-11 (35) | CCL2, CCL20, CCL5, CD274, CXCL10, CXCL11, CXCL16, CXCL2, CXCL3, Cxcl9, CYBB, FASLG, FOXP3, GBP2, IFIT2, IFIT3, IFNB1, IL1B, IL6, ISG15, MMP9, MX1, NFE2, OAS3, OASL, PLAAT1, PSMB9, |

| | | | | |
|---|---|---|---|---|
| | | | | PTGS2, SAMHD1, SPI1, TBX3, TNFSF10, TNFSF13B, UBD, VCAM1 |
| IRF3-regulated genes | | N/A | 2.07E-08 (32) | Ccl2, CCL2, CCL3L3, CCL5, CD274, CD69, CXCL10, Cxcl3, Cxcl9, FGF1, FST, GBP1, GBP5, IFI16, IFIT1B, IFIT2, IFIT3, IFNB1, IL6, ISG15, Klrk1, MFAP2, MMP9, Mx2, OAS3, OASL, PHF11, PLCG2, TNFSF10, TNFSF15, VCAM1, ZBP1 |
| IRF5-regulated genes | | N/A | 1.11E-04 (14) | CCL4, CCL5, CXCL10, CXCL11, CXCL2, CXCR4, IFIT2, IFIT3, IFNB1, IL6, ISG15, OASL, PTGS2, TNFSF10 |
| IRF7-regulated genes | | N/A | 1.42E-05 (24) | Ccl2, CCL5, CD69, CXCL10, FOXP3, GBP1, GBP4, GBP5, IFI16, IFIT1B, IFIT2, IFIT3, IFNB1, ISG15, MX1, Mx2, OAS3, OASL, PHF11, PSMB9, TNFSF10, TNFSF13B, UBA7, ZBP1 |
| NFKB1-regulated genes | 9.68E-03 (21) | AHRR, BCL2A1, BMP2, Ccl2, CCL2, CCL3L3, CCND1, CCND2, CRMP1, CXCL10, CXCL3, EGR1, ENPP2, FASLG, FOXP3, GBP1, IGFBP3, IL1RN, MMP13, ORAI1, PTGS2 | 2.11E-10 (44) | AHRR, AKR1B1, BCL2A1, Ccl2, CCL2, CCL20, CCL3L3, CCL4, CCL5, CCND2, CD274, CHI3L1, COL2A1, CXCL10, CXCL11, CXCL2, CXCL3, Cxcl9, CYBB, EBI3, EGR1, ENPP2, FABP6, FASLG, FOXP3, GBP1, IFI16, IFIT1B, IFNB1, IGFBP3, IL1B, IL1RN, IL6, ISG15, MMP13, MMP9, MSR1, NOD2, PTGS2, RGS1, SPI1, TNFSF10, TNFSF15, VCAM1 |
| NFKB2-regulated genes | 8.48E-04 (15) | BMP2, BMP4, C1QTNF3, Ccl2, CCND1, CD83, CXCL3, CXCR4, ENPP2, HLA-DQA1, IL34, ORAI1, PTGS2, RRAD, TGM4 | 5.01E-06 (22) | AKR1B1, BMP4, C1QTNF3, Ccl2, CCL20, CCL5, Ccl9, CD83, CRCT1, CXCL3, CXCR4, ENPP2, GDF6, Gvin1, IFNB1, IL1B, IL34, MMP9, NOD2, Ppbp, PTGS2, VCAM1 |
| NFKBIA-regulated genes | 5.40E-09 (48) | AGTR1, ALOX15B, B3GALT5, BCL2A1, BMP2, CAMP, CARD10, Ccl2, CCL2, CCL3L3, Ccl7, | 3.54E-12 (63) | ACOD1, AGTR1, ALOX15B, B3GALT5, BCL2A1, CAMP, CARD10, Ccl2, CCL2, CCL20, CCL3L3, CCL5, |

| | | | | |
|---|---|---|---|---|
| | | CCNA2, CCND1, CCND2, CDC25B, CEBPD, CXCL10, CXCL3, CXCR4, CYP1B1, ECSCR, ENPP2, ESM1, FASLG, HEY1, HLA-DQA1, HOXB2, ID1, ID3, IGFBP6, IL1RN, IL7R, KCNN4, Ly6a, MMP14, NCAN, NDRG2, NOTCH3, OGN, OLFM4, PDHA1, PTGS2, PTPN13, RBP1, RGS3, RGS4, TNC, TNFRSF11B | | Ccl7, Ccl9, CCNA2, CCND2, CD69, CDC25B, CHI3L1, COL2A1, CXCL10, CXCL2, CXCL3, CXCR4, CYP1B1, CYP7B1, ECSCR, ENPP2, ESM1, FASLG, GBP2, HLA-A, ID1, ID3, IFI16, IFIT1B, IFNB1, IL1B, IL1RN, IL6, IL7R, ISG15, KCNN4, LIMA1, MMP9, NCAN, NDRG2, NOD2, NOTCH3, OGN, OLFM4, PDHA1, PTGS2, PTPN13, RBP1, RGS3, SFRP1, TACR1, TFCP2L1, TFRC, TIMP1, TNC, VCAM1 |
| NONO-regulated genes | | N/A | 2.09E-06 (22) | CCL5, CDKN2A, CLEC4F, CXCL10, CXCL11, EPSTI1, GBP1, GBP4, GBP5, IFIT2, IFIT3, ISG15, MX1, OAS3, OASL, PDE2A, PDE3B, PDE7A, PDE8B, TNFSF10, TNFSF13B, TNFSF15 |
| RELA-regulated genes | 4.50E-03 (32) | ACTA2, AGT, AHRR, BCL2A1, BMP2, CAMP, CAV1, Ccl2, CCL2, CCND1, CD28, CXCL10, CXCL3, CXCR4, CYP2C9, EGR1, ELN, FASLG, FOXP3, GBP1, ID1, IL1RN, IL7R, NEFH, ORAI1, PDGFB, PTGES, PTGS2, TNFRSF11B, TP63, TRPC6, UBD | 9.80E-09 (56) | ACTA2, AHRR, ALPK2, BCL2A1, BEX1, CAMP, CAV1, Ccl2, CCL2, CCL20, CCL5, CD14, CD274, CD69, CDKN2A, CHI3L1, COL2A1, CXCL10, CXCL11, CXCL2, CXCL3, CXCR4, CYBB, CYP2C9, DNMT3B, EGR1, ELN, FABP6, FASLG, FOXP3, GBP1, GDF15, ID1, IFIT1B, IFNB1, IL1B, IL1RN, IL6, IL7R, ISG15, MMP9, NEFH, NOD2, PDGFB, PSMB9, PTGES, PTGS2, RGS1, SMOC2, SPI1, TACR1, TP63, TREM1, TWIST2, UBD, VCAM1 |
| STAT1-regulated genes | 1.05E-05 (34) | AGT, ALAS2, ALDH1A3, Ccl2, CCL2, CCND1, CCND2, CEBPD, CXCL10, CXCL3, DPP4, EGR1, FASLG, FOXP3, GBP1, GBP4, GCK, HAVCR2, HEY1, HLA-DQA1, | 2.11E-16 (62) | ACOD1, ALAS2, ALDH1A3, APOC2, C1QTNF7, C1S, C4A/C4B, Ccl2, CCL2, CCL20, CCL5, CCND2, CD14, CD274, CXCL10, CXCL11, CXCL2, CXCL3, Cxcl9, DPP4, EGR1, EPSTI1, |

| | | | | |
|---|---|---|---|---|
| | | IL1R1, INS, Klrk1, MAFA, MX1, Mx2, NEUROD1, NFE2, PDCD1LG2, PDGFC, PTGS2, SLAMF8, UBD, ZBP1 | | FASLG, FCER1G, FGF7, FOXP3, GBP1, GBP2, GBP4, GBP5, GCK, HAVCR2, IFI16, IFIT1B, IFIT2, IFIT3, IFNB1, IL1B, IL1R1, IL6, INS, ISG15, Klrk1, MMP9, MX1, Mx2, NEURL3, NFE2, OAS3, OASL, PAX6, PDGFC, PSMB9, PTGS2, SAMHD1, SLAMF8, SLC2A2, TFF1, TNFSF10, TNFSF13B, UBD, ZBP1 |
| TBX21-regulated genes | 1.70E-02 (7) | CCL3L3, FASLG, HAVCR2, IL23R, IL7R, Klrk1, PTGDR2 | 4.82E-04 (11) | CCL3L3, CCL4, CD244, FASLG, GZMA, HAVCR2, IL7R, Klrk1, PTGDR2, SELPLG, SPP1 |
| TGIF1-regulated genes | | N/A | 3.50E-03 (5) | CXCL2, CXCL3, IL1B, IL6, SOAT2 |
| VAV2-regulated genes | | N/A | 1.14E-02 (5) | BCL2A1, BTC, CD69, FGF7, IL6 |
| ZEB1-regulated genes | | N/A | 8.75E-03 (11) | CD163, CDKN2A, CGA, IL1B, MAL2, MMP9, PTGS2, SFTPB, TGFBI, THBS1, TP63 |
| HNRNPU-regulated genes | 4.45E-02 (3) | CCL2, HBB, NEUROD1 | 3.52E-04 (6) | CCL2, HBB, IL1B, IL6, SPP1, VCAM1 |
| HIPK2-regulated genes | 5.39E-05 (12) | ALAS2, ANK1, CCND2, EGR1, FECH, GATA1, HBA1/HBA2, HBE1, KLF1, NQO1, SLC4A1, SPTB | 4.05E-08 (18) | ALAS2, ANK1, BDNF, CCND2, CIDEA, EGR1, FECH, GATA1, HBA1/HBA2, HBE1, KLF1, NKX2-5, NQO1, RAC2, SLC4A1, SPTB, TRPM5, UPK3B |
| ESR1-regulated genes | 5.65E-05 (90) | ACE, AGT, ALK, BMP2, BMP4, CALML4, CAV1, CAV2, CCL2, CCNA2, CCND1, CD83, CDC42EP2, CEBPD, CHST2, CLDN1, CLEC3A, CNN1, COBL, COL13A1, COL8A1, CTH, CXCL3, CYP1A1, CYP1B1, Cyp4a14, DCDC2, EGF, EGR1, ELN, ENPP2, ERBB3, FGD5, FOXP3, FST, GAL, GHR, GJB1, GPR158, GSTA5, | 2.03E-06 (116) | ACE, ALOX5, BDNF, BMP4, CALML4, CAV1, CAV2, CCL2, CCL20, CCL4, CCNA2, CD69, CD83, CDC42EP2, CDKN2A, CGA, CHST2, CISH, CLDN1, CLDN8, CLEC3A, COBL, COL13A1, COL8A1, CTH, CXCL3, CYP1A1, CYP1B1, Cyp4a14, CYP7B1, DCDC2, EGR1, ELN, ENPP2, ERBB3, FGD5, FGFR2, FOXP3, FST, GABBR2, GDF15, GHR, |

| | | | | |
|---|---|---|---|---|
| | | GUCY1A1, GUCY1B1, HBA1/HBA2, HEY1, HFE, HPGDS, ID1, IGFBP3, IRS1, ITGA2B, KIF20A, KLF1, LAPTM5, LIPC, MCF2L, MX1, NOTCH3, NPY, NQO1, OXT, PADI4, PCBD1, PCDHB14, PHF19, PLCB1, PRSS23, PTGS2, PTPN13, RAMP3, RERG, RGS3, SCARB1, SCNN1B, SHANK3, SLC13A4, SLC16A3, SLC7A2, SMAD6, SOSTDC1, STEAP1, TAC3, TCIM, TEK, TLE5, TNFRSF11B, TNS3, TP53I11, TRPM2, TTR, UNC119 | | GJB1, GJB2, GLRX, GPR158, GSTA5, GUCY1A1, GUCY1B1, HBA1/HBA2, HPGDS, HYAL1, ID1, IGFBP3, IL1B, IL20RA, IL6, INMT, IRS1, KIF20A, KLF1, KRT13, LAPTM5, LIMA1, LIPC, MCF2L, MMP9, MST1, MX1, NOTCH3, NPW, NQO1, NR1D1, PCBD1, PCDHB5, PDE7A, PHF19, PLCB1, PLCE1, PLK4, PRR15L, PRSS23, PTGS2, PTPN13, RAB11FIP4, RAMP3, RBP4, RGS3, RND1, SAMHD1, SCARB1, SCNN1B, SHANK3, SLC13A4, SLC16A3, SLC2A12, SLC6A4, SLC7A2, Slco1a1, SMAD6, SOSTDC1, SPP1, STEAP1, TAC3, TCIM, TEAD4, TEK, TFF1, TFRC, TM4SF1, TNFSF10, TNS3, TRPM2, TTR, UBE2T, UNC119 |
| PPARG-regulated genes | 4.13E-07 (46) | ACADS, ACTA2, ADRB2, AGT, AGTR1, ALDH1A2, ALOX15B, APOA2, CAV1, CAV2, CAVIN2, Ccl2, CCL2, CCND1, Cxcl3, CXCL3, Cyp4a14, FABP3, FST, GCK, GPD1, GPT, IGFBP1, IGFBP3, IGFBP6, IGFBP7, IHH, INS, IRS1, Kctd12b, LEP, LIPE, MMP13, MMP14, PDE3B, PLA2G2A, Prl7d1, PTGES, PTGS2, SCARB1, SFTPB, SLC27A3, TAGLN, TBXA2R, TIMP4, VAMP5 | 1.81E-07 (55) | ACOD1, ACTA2, ADRB2, AGTR1, ALOX15B, APOA2, CAV1, CAV2, CAVIN2, Ccl2, CCL2, CCL5, CD274, CIDEA, CIDEC, CXCL2, Cxcl3, CXCL3, Cyp4a14, FABP3, FGF1, FST, GCK, GPT, IFNB1, IGFBP3, IGFBP7, IL1B, IL6, INS, IRS1, Kctd12b, LEP, LIPE, MMP13, MMP9, MTTP, MYH7, NCF1, NR1D1, PDE3B, PPP2R2B, PTGES, PTGS2, RBP4, SCARB1, SCNN1G, SFTPB, SLC2A2, SPP1, TAGLN, TBXA2R, TIMP4, TNFSF10, VCAM1 |
| THRA-regulated genes | 1.39E-02 (12) | BMP4, Ccl2, CCL2, CCND1, CXCL10, CXCL3, ENG, FRZB, FST, MAFA, MYH6, RCAN2 | 6.53E-02 (12) | BMP4, Ccl2, CCL2, CGA, CXCL10, CXCL2, CXCL3, FST, IL6, MYH6, MYH7, TREM1 |

| | | | | |
|---|---|---|---|---|
| THRB-regulated genes | 1.51E-05 (28) | APOA2, Ccl2, CCL2, CCND1, CCND2, CXCL10, CXCL3, CXCR4, EGF, ENG, ERBB3, FRZB, FZD4, GJA5, ID1, IGFBP3, IL24, MLXIPL, MMP13, MYH6, PLA2G2A, PTGS2, SLAMF8, TCEA3, TGFBI, TLE2, TP63, WNT16 | 6.73E-10 (42) | ANGPTL3, APOA2, APOM, ATP1A3, BCL2L14, Ccl2, CCL2, CCND2, CGA, CXCL10, CXCL2, CXCL3, CXCR4, ERBB3, FZD4, GJA5, ID1, IFIT1B, IGFBP3, IL24, IL6, MLXIPL, MMP13, MMP9, MYH6, MYH7, PTGS2, SFRP1, SH2D4A, SLAMF8, SLC5A11, SPP1, Sult5a1, TCEA3, TGFBI, TLE2, TNFSF10, TP63, TREM1, WNT16, WNT7A, ZNF185 |
| FOXM1-regulated genes | 4.88E-04 (16) | ACE, ACE2, ACTA2, CAV1, CCNA2, CCND1, CCND2, CDC25B, CDC25C, CTNNAL1, FOXF1, FUT4, HEY2, IL1RN, KIF20A, SFTPB | 3.61E-05 (21) | ACE, ACE2, ACTA2, CAV1, CCNA2, CCND2, CDC25B, CDKN2A, CTNNAL1, FOXF1, FUT4, HEY2, IL1RN, IL6, KIF20A, MMP9, PLK4, SFRP1, SFTPB, TWIST2, VCAN |
| GATA1-regulated genes | 2.22E-10 (39) | AHSP, ALAS2, ANK1, CA1, CAVIN1, CCND1, CCND2, CNN3, EGR1, EPB42, F10, FLI1, GATA1, GUCY1B1, HBA1/HBA2, HBB, HBE1, HBG2, IL7R, ITGA2B, KLF1, LYL1, MSX2, NEFH, NFE2, NRARP, POLA2, PRG2, PTGDR2, SLC19A1, SLC4A1, SPOCK2, SPTA1, SPTB, TAC3, TAL1, TEK, TUBB1, ZNF366 | 5.25E-10 (44) | AHSP, ALAS2, ALOX5, ANK1, CA1, CCL5, CCND2, CDKN2A, CNN3, CYBB, EGR1, EPB42, F10, FSD1L, GATA1, GP9, GUCY1B1, HBA1/HBA2, HBB, HBE1, HBG2, IL7R, KLF1, LYL1, MPL, MSX2, NEFH, NFE2, NRARP, PRG2, PTGDR2, RAD54L, SLC19A1, SLC4A1, SPOCK2, SPTA1, SPTB, TAC3, TAL1, TEK, TFRC, TUBB1, VWF, ZNF366 |
| HNF1A-regulated genes | 2.67E-05 (41) | A1CF, ACE2, AGT, Akr1c19, APOA2, B3GALT5, BMP3, C4BPA, CCND1, CPN1, CYP2C9, DPP4, EBF1, GAL, GCK, GJB1, HABP2, HPGDS, IGFBP1, IHH, INS, KIF20A, MELTF, MLXIPL, NOD1, PFKL, PKLR, PLA1A, PLCB1, SLC26A1, SLC4A2, SLC5A1, SLC7A2, SOAT2, SPO11, TMED6, | 1.84E-09 (60) | A1CF, AADAC, ACE2, Akr1c12/Akr1c13, Akr1c19, ALDOB, ANGPTL3, APOA2, APOB, APOH, APOM, B3GALT5, C1S, C4BPA, CDHR2, CLDN8, CPN1, CYP2C9, DPP4, EBF1, F11, FMO1, FUT7, G0S2, GCK, GCKR, GJB1, GLP1R, HPGDS, IGFALS, INS, KIF20A, MELTF, MLXIPL, MTTP, NR1D1, PKLR, PLA1A, PLCB1, PLG, |

| | | | | |
|---|---|---|---|---|
| | | TMPRSS2, TRPM2, TTR, UGT2B11, ZNF365 | | PPP1R1A, PROC, RNF186, SERPINA10, SLC2A2, SLC5A1, SLC7A2, Slco1a1, SLCO1B3, SOAT2, SPO11, TFRC, TMED6, TMPRSS2, TRPM2, TRPM5, TTR, VIL1, VTN, ZNF365 |
| MYOCD-regulated genes | 9.45E-07 (16) | ACE2, ACTA2, CLSTN2, CNN1, GJA5, GJC1, HEY2, LMOD1, MYH11, MYH6, MYL4, MYLK, PLA2G2A, PTPRB, SCN5A, TAGLN | 4.47E-06 (17) | ACE2, ACTA2, CLSTN2, COL2A1, ECM2, GJA5, HEY2, LMOD1, MYH11, MYH6, MYH7, MYL4, MYLK, PTPRB, SCN5A, TAGLN, VCAN |
| MYOD1-regulated genes | 1.67E-02 (18) | ACTA2, AGT, ASB2, BNC1, CADPS2, CCL2, CDH15, E2F2, ENPP2, FAM83D, IRX1, KIF18B, LIPE, MYH6, MYL4, NPTX1, PLEKHO1, Ppfia4 | 3.23E-03 (24) | ACTA2, ASB2, CADPS2, CCL2, CDH15, CKM, E2F2, ENPP2, FAM83D, HSPA4L, IFI16, IFIT2, IGLL1/IGLL5, KIF14, KIF18B, LIPE, MYH6, MYH7, MYL1, MYL4, NPTX1, PRKAR2B, RAC2, UBA7 |
| NCOA3-regulated genes | 4.11E-06 (15) | CCNA2, CCND1, CYP1A1, ERVFRD-1, GCM1, IGFBP1, IGFBP3, IRS1, LECT2, MYH11, OXT, PTGS2, SFTPB, SLC16A3, TAGLN | 1.63E-05 (16) | CCNA2, CYP1A1, ERVFRD-1, FGFR2, GCM1, GJB2, IGFBP3, IRS1, LECT2, MMP9, MYH11, PTGS2, SFTPB, SLC16A3, TAGLN, TFF1 |
| NFE2-regulated genes | 2.06E-07 (9) | ERMAP, GATA1, HBB, ITGA2B, KLF1, NQO1, RAB27B, SPTA1, TUBB1 | 1.61E-05 (8) | ERMAP, GATA1, HBB, KLF1, NQO1, SLC6A4, SPTA1, TUBB1 |
| NOTCH1-regulated genes | 6.08E-10 (36) | ACTA2, ANGPT1, ASB2, BMP2, CCND1, CCND2, CEBPD, CHGA, CNN1, CXCR4, ECT2, ERBB3, ESM1, HBA1/HBA2, HEY1, HEY2, ID1, IGFBP3, IL7R, ITGA2B, LOC100360737/Psg19, Ly6a, LYVE1, MCAM, MYH11, MYLK, NCF4, NEUROD1, NOTCH3, PDGFB, PTGS2, SLC5A5, TAGLN, TEK, TIE1, TP63 | 2.75E-08 (38) | ACTA2, ANGPT1, ASB2, Ccl9, CCND2, CD14, CHGA, CHST1, CKM, CXCR4, ECT2, ERBB3, ESM1, FCER1G, HBA1/HBA2, HEY2, ID1, IFI16, IGFBP3, IL6, IL7R, LYVE1, MCAM, MYH11, MYH7, MYLK, NCF1, NCF4, NOTCH3, PDGFB, PTGS2, SPI1, SPP1, TAGLN, TEK, TFF1, TIE1, TP63 |

| | | | | |
|---|---|---|---|---|
| NOTCH4-regulated genes | 2.07E-08 (10) | ACTA2, ANGPT1, CNN1, FLT4, HEY1, HEY2, MYH11, TAGLN, TEK, TIE1 | 2.30E-05 (8) | ACTA2, ANGPT1, FLT4, HEY2, MYH11, TAGLN, TEK, TIE1 |
| PRDM1-regulated genes | 2.87E-03 (21) | BCL2A1, CADPS2, Ccl2, CLDN1, CXCL3, CXCR4, EGR2, ERG, GSTA5, H2AC18/H2AC19, HLA-DQA1, ID3, IL24, IL7R, LRRN1, Lypd8l, SCARA5, SLC5A8, SLCO2B1, ST14, TTR | 9.00E-07 (34) | AIM2, APOM, BCL2A1, CADPS2, Ccl2, CCL20, CD53, CHI3L1, CLDN1, CXCL2, CXCL3, CXCR4, DBNDD1, EGR2, ERG, FGR, GSTA5, HCK, ID3, IFNB1, IL1B, IL24, IL6, IL7R, Keg1, LRRN1, MAP7D2, SCARA5, SLC5A8, SLCO2B1, ST14, TNFSF10, TNFSF13B, TTR |
| TAL1-regulated genes | 3.10E-02 (16) | ARVCF, C3AR1, EPB42, ERG, GATA1, GUCY1A1, HBB, HBE1, KIF15, KIF20A, KLF1, NDN, P2RY1, PLA2G2A, PLEKHO1, TSPAN7 | 1.00E-02 (21) | C3AR1, CACNG4, CD69, CDKN2A, EPB42, ERG, GATA1, GUCY1A1, HBB, HBE1, HLA-A, KIF14, KIF15, KIF20A, KLF1, LRRN3, PTCRA, RAB33A, SPI1, TNFSF10, TSPAN7 |
| TP73-regulated genes | 8.33E-04 (24) | ALAS2, Ccl2, CCL2, CCNA2, CDC25B, CDC25C, CXCL10, CXCL3, EGR1, ENG, FOXF1, FST, GATA1, HBG2, IGFBP3, MMP13, MMP14, MMP28, PDGFB, PTGES, THY1, TIMP4, TP53I11, UBE2C | 5.10E-04 (29) | ALAS2, Ccl2, CCL2, CCNA2, CDC25B, CXCL10, CXCL2, CXCL3, EGR1, FGFR2, FOXF1, FST, G0S2, GATA1, HBG2, IFNB1, IGFBP3, IL1B, IL6, MMP13, MMP28, NLRC4, PDGFB, PTGES, THBS1, THBS2, THY1, TIMP1, TIMP4 |
| PARK7-regulated genes | | N/A | 4.26E-02 (4) | Ccl2, CCL3L3, IL1B, IL6 |
| ESR2-regulated genes | 2.77E-04 (48) | AGT, AJUBA, ANK1, BMP2, BMP4, CADM3, CAV1, CCL2, CCL3L3, CCNA2, CCND1, CD93, CEBPD, CHST2, CLDN1, CLDN11, CNN1, COL11A1, COL13A1, EGR1, EMB, ENPP2, ENPP3, FBLN5, FST, GATA1, GJB1, GSTA5, IL1R1, IL23R, LAPTM5, MMP13, MMP14, MYH6, MYPN, NDRG2, NPY, OXT, PCBD1, PDGFB, PTGS2, S1PR1, | 1.48E-07 (69) | AJUBA, ANK1, BMP4, CADM3, CAV1, CCL2, CCL20, CCL3L3, CCNA2, CD93, CGA, CHST2, CLDN1, CLDN11, COL11A1, COL13A1, CXCL16, CXCL17, CXCL2, EGR1, ELAVL2, EMB, ENPP2, ENPP3, FBLN5, FST, GABBR2, GATA1, GBP2, GJB1, GLRX, GSTA5, IFIT2, IFIT3, IL1B, IL1R1, IL6, LAPTM5, LRRN3, MEGF10, MMP13, MMP9, MYH6, MYH7, MYPN, Nat8f3, |

| | | | | |
|---|---|---|---|---|
| | | SCARB1, STC1, TLE5, TNFRSF11B, TNS3, TP53I11 | | NDRG2, PCBD1, PDGFB, PLK4, PSMB9, PTCRA, PTGS2, RBP4, S1PR1, SCARB1, SPP1, SRXN1, STC1, TEAD4, TFF1, THBS1, THBS2, TIGD3, TIMP1, TNFSF13B, TNS3, TSPAN13, VCAM1 |
| NR0B2-regulated genes | 1.40E-02 (11) | AGT, Ccl2, CD163, EGR1, GCK, GSTA5, HDC, IL1RN, PIK3C2G, SCARB1, TNFRSF11A | 1.07E-07 (23) | APOM, CARD11, Ccl2, CD163, CIDEC, CYP7B1, EGR1, GCK, GSTA5, HDC, IL1B, IL1RN, IL6, MTTP, PIK3C2G, PIK3CG, PIK3R5, PLCG2, SCARB1, SPP1, TFF1, TIMP1, TNFRSF11A |
| NR1H2-regulated genes | 3.80E-04 (13) | ACE, AGT, Ccl2, Ccl7, CXCL10, GBP4, GCK, IL1RN, INS, LIPE, PTGS2, SCARB1, SLC16A10 | 6.52E-10 (24) | ACE, Ccl2, CCL4, CCL5, Ccl7, CIDEA, CXCL10, CXCL16, CXCL2, GBP4, GCK, IFIT2, IL1B, IL1RN, IL6, INS, ITGAL, LIPE, MMP9, PNPLA2, PTGS2, SCARB1, SLC16A10, SLC2A2 |
| NR1H3-regulated genes | 9.14E-04 (15) | ACTA2, ADRB2, Ccl2, Ccl7, CCNA2, CXCL10, GBP4, GCK, GSTA5, IL1RN, MLXIPL, PKLR, PTGS2, SCARB1, SLC16A10 | 1.39E-09 (28) | ACTA2, ADRB2, ANGPTL3, APOC2, APOM, Ccl2, CCL4, CCL5, Ccl7, CCNA2, CIDEA, CXCL10, CXCL16, CXCL2, GBP4, GCK, GSTA5, IFIT2, IL1B, IL1RN, IL6, ITGAL, MLXIPL, MMP9, PKLR, PTGS2, SCARB1, SLC16A10 |
| NR3C1-regulated genes | 1.31E-05 (56) | ACTA2, AGT, APOA2, BMP2, C1QTNF2, CARD10, CAV1, CAVIN2, Ccl2, CCL2, CCL3L3, CCND1, CD83, CEBPD, CXCL10, CXCL3, CXCR4, CYP2C9, DEAF1, DPP4, EGR1, EPB41L4B, FASLG, GRIA3, GRM5, IGFBP1, IL24, IL3RA, IL7R, ITGB6, KLF13, LEP, LIPC, MCF2, MMP13, MSX2, Mt1m/Mt2A, NOTCH4, NPY, NQO1, NRXN1, PAX3, | 3.67E-10 (81) | ACTA2, ANGPTL7, APOA2, BCL2L14, BDNF, C1QTNF2, C1QTNF7, C4A/C4B, CARD10, CAV1, CAVIN2, CBR1, Ccl2, CCL2, CCL3L3, CCL4, CCL5, CD69, CD83, CGA, CIDEA, CIDEC, CXCL10, CXCL2, CXCL3, Cxcl9, CXCR4, CYP2C9, DPP4, EGR1, EPB41L4B, FASLG, FCER1G, GBP2, GPR65, GRIA3, GRM3, GZMA, Hsd3b4, IFIT1B, IFIT2, IFNB1, IGFALS, IL1B, |

| | | | | |
|---|---|---|---|---|
| | | PDCD1LG2, PLEKHF1, PRKCB, PTGS2, RGS2, SCARB1, SLC7A6, SNCA, STC1, TCIM, TIMP4, TNFRSF11B, TNFRSF21, TUBA4A | | IL24, IL6, IL7R, ISG15, LEP, LIPC, MCF2, MMP13, MMP9, MSX2, Mt1m/Mt2A, NLRC4, NOD2, NOTCH4, NQO1, OASL, PNPLA2, PRKCB, PTGS2, RGS2, SCARB1, SCNN1A, SELENOP, SLC7A6, SNCA, SPP1, STC1, STOM, TCIM, TEAD4, THBS1, TIMP1, TIMP4, TLR1, TNFRSF21, TUBA4A, VCAM1 |
| NR5A2-regulated genes | 1.05E-02 (12) | ACTA2, C3AR1, Ccl7, CCND1, CEBPD, CPA1, CXCL10, IGFBP1, IL1RN, IL24, RGS4, SCARB1 | 8.26E-05 (19) | ACTA2, APOM, C1QB, C3AR1, C5AR1, Ccl7, CPA1, CXCL10, CXCL16, Cxcl9, CYP7B1, FABP6, IL1B, IL1RN, IL24, IL6, MMP9, SCARB1, TNFSF10 |
| BCL3-regulated genes | 4.73E-04 (9) | AGT, CAMP, CCND1, CXCL10, CXCL3, EGR1, EGR2, FABP3, ID1 | 8.51E-07 (14) | CAMP, CCL20, CCL5, CD274, CD69, CXCL10, CXCL2, CXCL3, EGR1, EGR2, FABP3, ID1, IL1B, IL6 |
| BCL6-regulated genes | 2.56E-03 (20) | BCL2A1, BMP2, Ccl2, CCL2, CCL3L3, Ccl7, CCND1, CCND2, CD72, CXCL10, CXCL3, FOXP3, IL24, IL7R, Ly6a, PDCD1LG2, PLA2G2A, PRSS23, RGS4, TACSTD2 | 2.68E-05 (29) | BCL2A1, Ccl2, CCL2, CCL3L3, CCL4, Ccl7, CCND2, CD274, CD69, CD72, CDKN2A, CISH, COL2A1, CXCL10, CXCL3, EBI3, FOXP3, FUT7, IL1B, IL24, IL6, IL7R, PIK3CG, PRSS23, SH2D4A, SLC27A2, TACSTD2, TIMP1, UBA7 |
| CITED2-regulated genes | | N/A | 8.56E-02 (5) | CCL2, CDKN2A, IL1B, IL6, MMP13 |
| FEM1A-regulated genes | 4.25E-02 (2) | Ccl2, CXCL10 | 6.06E-07 (6) | Ccl2, CCL4, CXCL10, CXCL2, IL1B, IL6 |
| FOXA2-regulated genes | 4.64E-07 (29) | ABCC8, ACTA2, ACTG2, CCND1, CNN1, CPN1, CXCR4, CYP2C9, DCX, DYNLRB2, GAL, GCK, IGFBP1, Krt2, LIPE, MAFA, MLXIPL, MUC6, MYH11, MYL9, MYOCD, PPP1R14A, PYY, SCGB1A1, SFTPB, SOX17, TAGLN, TMPRSS2, TTR | 8.45E-08 (35) | ABCC8, ACTA2, ACTG2, ALDOB, ALOX5, APOB, APOM, CHI3L1, CPN1, CXCR4, CYB561, CYP2C9, DYNLRB2, GCK, GSTM3, IL6, KCNJ11, Krt2, LIPE, MLXIPL, MTTP, MYH11, MYL9, MYOCD, PAX6, PYY, SCGB1A1, SFTPB, SLC2A2, |

| | | | | |
|---|---|---|---|---|
| | | | | SLCO1B3, SOX17, TAGLN, TMPRSS2, TTR, WNT7A |
| GPS2-regulated genes | 5.52E-03 (8) | ALAS2, Ccl2, CCL3L3, CCND1, CXCL10, ERMAP, Kctd12b, KLF1 | 2.96E-05 (13) | ALAS2, Ccl2, CCL20, CCL3L3, CCL4, CD14, CXCL10, ERMAP, IL6, Kctd12b, KLF1, TFF1, VCAM1 |
| HLX-regulated genes | 9.25E-03 (5) | BMP2, CLDN1, CYP1B1, EGR1, SEMA3G | 5.50E-03 (6) | CDKN2A, CLDN1, CYP1B1, EGR1, GDF15, SEMA3G |
| IRF4-regulated genes | | N/A | 1.13E-04 (23) | ALPK2, Ccl2, CCL20, CDKN2A, CXCL10, CXCL11, CXCL3, CXCR4, ERBB3, FOXP3, GBP1, IL1B, IL1R1, IL1RN, IL6, IL7R, ISG15, ITGAL, PHF11, RGS1, SPP1, TNFSF10, TNFSF13B |
| KLF2-regulated genes | 2.58E-08 (25) | ACE, BMP4, CAV1, Ccl2, CCL2, Ccl7, CD93, CXCL3, CXCR4, CYP1B1, ELN, FOXP3, HBB, HBE1, HBG2, ID1, ID3, KCNN4, MMP14, NOTCH3, NQO1, PTGS2, RGS3, S1PR1, TBX5 | 5.41E-14 (37) | ACE, BMP4, CAV1, Ccl2, CCL2, CCL4, CCL5, Ccl7, CD93, CXCL2, CXCL3, CXCR4, CYP1B1, ELN, FGR, FOXP3, HBB, HBE1, HBG2, ID1, ID3, IL1B, IL6, KCNN4, NOTCH3, NQO1, PTGS2, RGS3, S1PR1, SELPLG, TBX5, TFRC, THBS1, TIMP1, TNFSF10, VCAM1, VWF |
| LHX1-regulated genes | 5.05E-04 (13) | C7orf57, CADPS2, DPP4, EHD2, EMB, FABP3, GSTA5, LRP2, Ly6a, SLC26A1, SLC5A8, SOSTDC1, SOX17 | 3.26E-08 (22) | AADAT, ALDOB, C7orf57, CADPS2, CUBN, CYB561, DPP4, EHD2, EMB, FABP3, GSTA5, Hsd3b4, Keg1, LRP2, LRRC19, NAPSA, SFRP1, SLC12A1, SLC5A8, SOSTDC1, SOX17, SPP2 |
| NEUROD1-regulated genes | 7.07E-04 (9) | ABCC8, CHGA, DCX, GCK, INS, MAFA, NEUROD1, NTRK2, SCNN1B | 3.58E-03 (9) | ABCC8, CHGA, GCK, INS, KCNJ11, NTRK2, PAX6, SCNN1B, SLC2A2 |
| NFE2L2-regulated genes | 1.84E-02 (29) | ACE, ACE2, AGT, ALAS2, BHMT, Ccl2, CXCL10, CXCL3, Cyp4a14, ELN, EPHB4, F10, FXYD1, GHR, GSTA5, IL24, KRT4, NEUROD1, NQO1, PRKCB, PTGS2, PTPRB, | 1.23E-05 (47) | ACE, ACE2, ALAS2, BDNF, BHMT, CA11, CBR1, Ccl2, CCL20, CCL5, CXCL10, CXCL2, CXCL3, CYBB, Cyp4a14, ELN, EPHB4, F10, FMO1, GHR, GSTA5, IFIT1B, IFNB1, IL1B, |

| | | | | |
|---|---|---|---|---|
| | | RAMP2, S100A13, SCARB1, SYT5, TCN2, TTR, TUB | | IL24, IL6, INMT, ISG15, KRT4, MMP9, NCF1, NQO1, OSCAR, PRKCB, PTGS2, PTPRB, RAMP2, S100A13, SCARB1, SERPIND1, SLCO1B3, SRXN1, TCN2, TTR, TUB, VCAM1, VSNL1 |
| NKX2-3-regulated genes | 1.32E-09 (32) | ACTA2, AOC3, BMP2, BMP4, CAVIN2, CLEC1A, F2RL3, FBN2, FLRT2, GBP1, GHR, H2AC18/H2AC19, HEY1, HEY2, HFE, HNRNPA0, IL7R, KLF2, MFNG, MTURN, MYH11, PLA1A, PLCB1, PLEKHA4, PRSS12, PTGS2, SEMA3A, SNCAIP, SRPX2, TCIM, TMEM100, TRPC6 | 9.81E-07 (31) | ACTA2, AOC3, BMP4, CAVIN2, CLEC1A, CXCL16, F2RL3, FBN2, GBP1, GBP2, GDF15, GHR, HEY2, IFNB1, IL7R, MFNG, MMRN1, MTURN, MYH11, PLA1A, PLCB1, PRSS12, PSMB9, PTGS2, SEMA3A, SNCAIP, SRPX2, TCIM, TMEM100, UBA7, VCAM1 |
| PIAS1-regulated genes | 2.75E-04 (11) | ACTA2, CCL2, CCND1, GUCY1A1, KLF2, MYH11, MYH6, NRXN1, RRAD, SNCA, TAGLN | 6.06E-03 (10) | ACTA2, BDNF, CCL2, GRIK4, GRM4, GUCY1A1, MYH11, MYH6, SNCA, TAGLN |
| PRDM16-regulated genes | | N/A | 7.75E-06 (10) | CCL5, CIDEA, CXCL10, Cxcl9, GBP4, IFIT1B, IFIT2, Mx2, MYH7, OAS3 |
| SIRT1-regulated genes | | N/A | 1.35E-05 (43) | Acan, BDNF, CAV1, Ccl2, CCL5, CCND2, CD72, CD83, CDKN2A, COL2A1, CORO1A, CPA1, ENTPD8, FGF7, FOXP3, GCK, GSTM3, HCK, HLA-A, Hsd3b4, ID1, IFIT1B, IFIT3, IFNB1, IL1B, IL6, LAMA2, LAPTM5, LEP, MMP13, MMP9, NCF1, NR1D1, NTRK2, PNPLA3, PSMB9, SCNN1A, SLC27A6, TACSTD2, TAL1, TRIM31, UBA7, VCAM1 |
| STAT5B-regulated genes | 1.72E-02 (20) | ALAS2, CCL3L3, CCND1, CCND2, CYP2C9, EBF1, FOXP3, GCK, GJB1, GPD1, IL24, LEP, MYH6, MYL4, NQO1, OSM, PCSK5, SCARA5, TNFRSF11A, TULP2 | 5.25E-05 (32) | ACOT4, ALAS2, CBR1, CCL3L3, CCND2, CIDEA, CISH, CYP2C9, CYP39A1, EBF1, FOXP3, G0S2, GCK, GJB1, Hsd3b4, IGFALS, IL24, Keg1, LEP, MYH6, MYH7, MYL1, MYL4, NQO1, OSM, PCSK5, |

| | | | | |
|---|---|---|---|---|
| | | | | SCARA5, Scart1, SLC34A2, Slco1a1, TNFRSF11A, TULP2 |
| TRIM24-regulated genes | | N/A | 1.10E-07 (22) | CCL4, CCL5, CXCL10, EMB, EPSTI1, FLRT3, GBP2, GBP4, IFIT1B, IFIT2, IFIT3, IFNB1, IL1R1, ISG15, OASL, PHF11, PSMB9, SAMHD1, SPP1, TGFBI, TRPV6, UBA7 |
| ZNF281-regulated genes | 7.27E-03 (6) | CCL2, CCL25, CCL3L3, CXCR4, OSM, PTGS2 | 4.70E-05 (10) | CCL2, CCL25, CCL3L3, CCL4, Ccl9, CXCL2, CXCR4, IL6, OSM, PTGS2 |
| BRD4-regulated genes | 1.84E-03 (15) | ACTA2, CCNA2, CCND1, CCND2, CXCL3, CXCR4, ICAM2, IL7R, NQO1, PDGFB, PDGFC, POLR3G, SLC19A1, SLC38A5, SLC7A2 | 5.33E-06 (23) | ACTA2, CCNA2, CCND2, CD274, CDCA5, CXCL2, CXCL3, CXCR4, ICAM2, IGLL1/IGLL5, IL6, IL7R, NQO1, PDGFB, PDGFC, POLR3G, SLC19A1, SLC38A5, SLC7A2, SPP1, THBS1, THBS2, TIMP1 |
| RARA-regulated genes | 1.41E-05 (34) | ALDH1A2, ATP6V0A4, BMP8A, CCN6, CCNA2, CCND1, CD93, CLMN, COBL, EGR1, EGR2, ENOX1, FASLG, FOXP3, FUT4, GCK, GHR, GPR158, IGFBP6, IL1R1, IL23R, IL36B, KIRREL2, OXT, PHF19, PTGS2, RBP1, RBP2, RGS3, SFTPB, SOX17, TG, TNFRSF11B, TP63 | 1.50E-07 (45) | ALOX5, APOM, ATP6V0A4, CCNA2, CD14, CD53, CD93, CLMN, COBL, Cyp2c70, EGR1, EGR2, FASLG, FOXP3, FUT4, GCK, GDF15, GHR, GPR158, IFIT1B, IL1R1, IL36B, IL6, KIRREL2, Ly6g6e, MAOB, PHF19, PIK3CG, PLK4, PTGS2, RBP1, RBP2, RGS3, SAMHD1, SFTPB, SLC2A2, SOX17, SPP1, Tdh, TFF1, TG, TP63, UBA7, UBE2T, ZMAT4 |
| AIRE-regulated genes | 3.92E-04 (11) | BMP2, BMP4, CXCL10, FOXP3, HBG2, INS, Ngp, NPY, OXT, PROKR2, TG | 5.43E-05 (14) | BMP4, C4A/C4B, CCL5, CD274, CXCL10, CXCL11, FOXP3, HBG2, INS, MMP9, Ngp, TG, TNFSF10, VCAM1 |
| CREBBP-regulated genes | 2.21E-07 (86) | AGT, ANGPT1, ARHGAP24, CAMK1G, CCND1, CCND2, CNN3, CRYAA/CRYAA2, CST7, CXCL10, CXCR4, DCX, EGR1, EGR2, FASLG, GIMAP1-GIMAP5, GIMAP4, GRIA3, HBB, IL23R, | 4.42E-12 (108) | ACOXL, ANGPT1, ARHGAP24, CCL5, CCND2, CD53, CGA, CKM, CNN3, COL2A1, CORO2B, CRYAA/CRYAA2, CST7, CXCL10, Cxcl9, CXCR4, CYBB, EGR1, EGR2, EPSTI1, FASLG, FUT7, GIMAP1- |

| | | | | |
|---|---|---|---|---|
| | | IL7R, KLF2, KLHL4, KMO, MCF2L, NEUROD1, PDE3B, PKLR, PLEKHF1, PTGS2, RGS2, SCN4B, SFTPB, SLC6A19, TAGLN, TBX4, TLE2, TMPRSS2, TNC, TUBA8 | | GIMAP5, GIMAP4, GRIA3, HBB, IFIT1B, IFNB1, IL17RE, IL18R1, IL6, IL7R, ISG15, KLHL4, KMO, MCF2L, MMP9, MST1, MYH7, OAS3, PDE3B, PKLR, PSMB9, PTGS2, RGS2, SCN4B, SFTPB, SLC6A19, STOM, TAGLN, TBX4, TFF1, TLE2, TMPRSS2, TNC, TNFSF10, VCAM1 |
| CTNNB1-regulated genes | 3.00E-10 (86) | ACADS, Acan, ACTA2, ADRA2C, ALDH1A2, ALDH1A3, AOC3, BMP2, BMP4, CACNA1G, CBLN1, Ccl2, Ccl7, CCNA2, CCND1, CCND2, Cd52, CHGA, CRIP1, CXCL10, Cxcl3, CXCL3, CYP1A1, CYP1B1, ECRG4, EGR1, ENPP2, EPHB4, ERBB3, FASLG, FGF9, FLI1, FST, GCK, GHR, GJB1, Gulo, HNRNPA0, ID1, ID3, ID4, IHH, IL1R1, IRS1, KRT1, LECT2, LIPE, MMP13, MMP14, MMP16, MPEG1, MSX2, MUC6, MYH6, MYL4, MYLK, NAP1L5, NDRG2, NEUROD1, NOTCH3, NOTUM, NPTX1, OGN, PAX3, PMP22, PROX2, PRX, PTGS2, RPL21, Rpl39l, SCN5A, SCNN1B, SIM2, SLC1A5, SLC38A5, SMAD6, SOX11, SOX17, SOX7, TBX5, TLE5, TNC, TNFRSF11B, TP63, UPB1, WNT16 | 2.11E-12 (108) | Acan, ACTA2, ALDH1A3, AOC3, APCDD1, BCL2L10, BDNF, BMP4, C6, CACNA1G, CBLN1, Ccl2, CCL4, CCL5, Ccl7, Ccl9, CCNA2, CCND2, CD274, Cd52, CDH16, CDKN2A, CHGA, CIDEC, COL2A1, CXCL10, CXCL2, Cxcl3, CXCL3, Cxcl9, CYP1A1, CYP1B1, CYP7B1, DNAJC6, ECRG4, EDIL3, EGR1, ENPP2, EPHB4, ERBB3, FASLG, FCER1G, FGF9, FST, GCK, GDF15, GHR, GJB1, GJB2, Gulo, Hsd3b4, ID1, ID3, ID4, IFI16, IFNB1, IL1B, IL1R1, IL6, INSL6, IRS1, ITGAL, LECT2, LIPE, MMP13, MMP16, MMP9, MPEG1, MSX2, MYH6, MYL4, MYLK, NDRG2, NKX2-5, NOTCH3, NOTUM, NPTX1, OGN, PMP22, PRX, PTCRA, PTGS2, RBP4, SCN5A, SCNN1B, SFRP1, SIM2, SLC1A5, SLC38A5, Slco1a1, SMAD6, SOX11, SOX17, SOX7, SPI1, SPP1, TBX5, TFCP2L1, TFF1, TIMP1, TNC, TNFSF10, TP63, TWIST2, UPB1, VCAM1, VCAN, WNT16 |
| FOXC2-regulated genes | 9.10E-04 (9) | ACTA2, ADRB1, ADRB2, CXCR4, IRS1, LEP, LIPE, PTGS2, TCF15 | 1.25E-03 (10) | ACTA2, ADRB1, ADRB2, CXCR4, IRS1, LEP, LIPE, MMP9, PTGS2, TCF15 |

| | | | | |
|---|---|---|---|---|
| HAND2-regulated genes | 1.11E-04 (11) | ACE2, CCL2, CCL3L3, CLSTN2, CXCR4, GJA5, MYH6, MYL4, PLA2G2A, PTGS2, PTPRB | 2.04E-06 (15) | ACE2, CCL2, CCL3L3, CCL4, Ccl9, CLSTN2, CXCL2, CXCR4, ECM2, GJA5, IL6, MYH6, MYL4, PTGS2, PTPRB |
| HIF1A-regulated genes | 1.70E-05 (42) | ABCC8, ACE, ACTA2, AGT, AGTR1, BMP2, CAV1, CAVIN1, CAVIN2, Ccl2, CCL2, CCL3L3, Ccl7, CCND1, CLDN1, COX4I2, CXCL10, Cxcl3, CXCL3, CXCR4, CYP1A1, ENG, GATA1, GCK, HIF3A, HOXA13, ID1, IGFBP3, IL23R, ITGA2B, LEP, MMP13, NRARP, NTRK2, PDGFB, PDHA1, PFKL, PTGS2, SLC16A3, SLC16A4, TBX5, TNFRSF11B | 1.15E-07 (56) | ABCC8, ACE, ACTA2, AGTR1, CAV1, CAVIN2, Ccl2, CCL2, CCL3L3, CCL4, CCL5, Ccl7, CD274, CD69, CDKN2A, CLDN1, CORO1A, COX4I2, CXCL10, CXCL2, Cxcl3, CXCL3, Cxcl9, CXCR4, CYP1A1, DNAJC5B, FUT7, GATA1, GCK, HIF3A, HOXA13, ID1, IFI16, IGFBP3, IL1B, IL6, LEP, MMP13, MMP9, MYH7, NRARP, NTRK2, PDGFB, PDHA1, PRKAR2B, PTGS2, RAB11FIP4, SFRP1, SLC16A3, SLC16A4, TBX5, TFRC, THBS1, TIMP1, TNFSF10, VCAM1 |
| ID1-regulated genes | 4.99E-04 (10) | CCND1, DTX4, EGR1, FAM110D, Ly6a, MMP13, MMP14, NEUROD1, SLC5A5, TG | 2.93E-03 (10) | CDKN2A, CGA, DTX4, EGR1, FAM110D, IFI16, MMP13, MMP9, TG, THBS1 |
| KAT5-regulated genes | 3.83E-02 (9) | CACNB2, CCND1, CCND2, CXCR4, OGN, PALMD, SOX17, TP63, TTR | 4.84E-03 (13) | CCND2, COL2A1, CXCR4, IL6, OGN, PALMD, PTCRA, RGS1, SOX17, TFF1, TP63, TTR, WNT7A |
| KLF13-regulated genes | 9.64E-04 (5) | BMP2, FECH, GATA1, HBB, HBE1 | 2.74E-03 (5) | CCL5, FECH, GATA1, HBB, HBE1 |
| KMT2D-regulated genes | 1.20E-02 (16) | ADCY5, CA4, CCDC160, CCL2, CRIP1, DAZ2, Eda, FABP3, GRHL3, LRRN4, PRSS16, SCARB1, SMAD6, TCF15, XKRX, ZNF750 | 2.79E-03 (21) | ACOT4, ADCY5, CCDC160, CCL2, EBI3, Eda, FABP3, FBXL7, GJB2, GRHL3, MMP9, PPP2R2B, PRSS16, SCARB1, SMAD6, TCF15, TFF1, VSIG8, VTN, XKRX, ZNF750 |
| NKX2-1-regulated genes | 1.32E-05 (22) | AGER, AGT, AGTR2, BCL2A1, BMP4, CA4, CHIA, DPP4, ENPP2, IGFBP6, LRP2, MYH11, PRRX2, SCGB1A1, SFTPB, SLC5A5, SOX17, TG, THY1, TNC, TP63, TSPAN7 | 5.91E-08 (30) | AGER, AGTR2, BCL2A1, BMP4, CD14, DPP4, EDIL3, ENPP2, HCK, HLA-A, INMT, LRP2, MYH11, NAPSA, PLA2G1B, RAC2, SCGB1A1, SCN7A, SCNN1G, SFTPB, SLC34A2, |

| | | | | |
|---|---|---|---|---|
| | | | | SLC6A14, SLCO1B3, SOX17, TG, THBS1, THY1, TNC, TP63, TSPAN7 |
| NOTCH3-regulated genes | 4.73E-04 (9) | ACTA2, FOXP3, GUCY1B1, HEY1, HEY2, ID1, MYH11, NRARP, TAGLN | 1.39E-04 (11) | ACTA2, FOXP3, GUCY1B1, HEY2, ID1, MMP9, MSR1, MYH11, NRARP, PTCRA, TAGLN |
| PAX8-regulated genes | 4.97E-03 (6) | CCNA2, GSTA5, HBB, LRRN1, SLC5A5, TG | 5.12E-02 (5) | CCNA2, GSTA5, HBB, LRRN1, TG |
| SMARCA4-regulated genes | 1.96E-05 (50) | ABLIM3, ACE, ACE2, ACTA2, AGT, ALDH1A3, BCL2A1, BMP4, C6orf141, CCL2, CCNA2, CCND1, CXCR4, CYP1A1, EBF1, EFNB1, EGR1, FGF9, GPM6A, GPR158, HBB, HBE1, HEPH, HEY1, HFE, ICAM2, ID3, IGFBP7, IL36B, IL7R, KCNS3, LRRC34, MELTF, MYH11, MYL4, MYLK, PLEKHO1, Ppfia4, RBP1, RGS2, SFTPB, SMAD6, SOX17, TACSTD2, TAGLN, TAL1, TCIM, THY1, TTR, UBD | 6.05E-10 (73) | ABLIM3, ACE, ACE2, ACTA2, AIM2, ALDH1A3, ARHGAP30, BCL2A1, BMP4, C6orf141, CCL2, CCNA2, CDH16, CDKN2A, CKM, CORO6, CXCR4, CYP1A1, EBF1, ECM2, EGR1, FABP6, FGF9, FGFR2, GPR158, GSG1, HBB, HBE1, HEPH, ICAM2, ID3, IFI16, IGFBP7, IL1B, IL36B, IL6, IL7R, KCNS3, MAOB, MELTF, MYH11, MYL1, MYL4, MYLK, NCALD, PHEX, PIDD1, PSMB9, RAC2, RBP1, RGS1, RGS2, SELENOP, SERPINB8, SFTPB, SLAMF9, SLC11A1, SMAD6, SOX17, SPP1, TACSTD2, TAGLN, TAL1, TCIM, TFF1, THY1, TNFSF10, TREM1, TSPAN13, TTR, UBD, WNT7A, ZNF185 |
| SP1-regulated genes | 1.62E-13 (66) | ABCC8, ACTA2, ADCY4, AGER, AGTR1, AHRR, ALAS2, ALOX15B, ANK1, ANKH, BMP4, CAMP, CAV1, Ccl2, CCL2, CCNA2, CCND1, CCND2, CD28, CDC25B, CDC25C, CEBPD, CHGA, COL8A1, CTH, CXCL10, CXCL3, CXCR4, CYBRD1, CYP1B1, EGR1, ENG, F10, FASLG, FOXP3, GHR, HBB, HBE1, HBG2, HDC, ID4, | 3.40E-15 (80) | ABCC8, ACTA2, AGER, AGTR1, AHRR, ALAS2, ALOX15B, ALOX5, ANK1, ANKH, BDNF, BMP4, CAMP, CAV1, Ccl2, CCL2, CCL20, CCL5, CCNA2, CCND2, CDC25B, CDKN2A, CHGA, CHI3L1, CHRNA3, COL2A1, COL8A1, CTH, CXCL10, CXCL3, CXCR4, CYBRD1, CYP1B1, Cyp2d1/Cyp2d5, CYP7B1, EGR1, F10, FABP6, FASLG, FOXP3, GDF15, |

| | | | | |
|---|---|---|---|---|
| | | IGFBP1, IGFBP3, KLF1, KRT4, MEOX2, MMP14, MYH11, MYLK, PADI4, PDGFB, PDGFC, PTGS2, PYY, RASGRP2, ROBO4, SCARB1, SLC19A1, SLC4A2, SLC5A1, SNRPN, SPTB, TAGLN, TBXA2R, TIMP4, TNC | | GHR, HBB, HBE1, HBG2, HDC, ID4, IFNB1, IGFBP3, IL1B, KLF1, KRT4, MAOB, MEOX2, MMP9, MPL, MYH11, MYH7, MYLK, PDGFB, PDGFC, PRKAR2B, PTCRA, PTGS2, PYY, SCARB1, SLC11A1, SLC19A1, SLC2A12, SLC5A1, SPP1, SPTB, TAGLN, TBXA2R, TFF1, TIMP1, TIMP4, TNC, TNFSF10, VWF |
| SP3-regulated genes | 4.35E-05 (23) | ACTA2, ADCY4, AHRR, ALOX15B, Ccl2, CCND1, CDC25B, EGR1, F10, FOXP3, GHR, IGFBP3, KLF1, MYH11, MYLK, PADI4, PTGS2, PYY, RASGRP2, SCARB1, SLC19A1, TAGLN, TNC | 1.48E-06 (30) | ACTA2, AHRR, ALOX15B, Ccl2, CCL20, CDC25B, CHI3L1, CHRNA3, COL2A1, CYP7B1, EGR1, F10, FOXP3, GDF15, GHR, IGFBP3, KLF1, MAOB, MYH11, MYH7, MYLK, PTCRA, PTGS2, PYY, SCARB1, SLC19A1, TAGLN, TFF1, TIMP1, TNC |
| SREBF1-regulated genes | 1.55E-02 (19) | ACADS, APOA2, CEBPD, CTH, CXCL10, FABP3, GCK, HFE, INS, KCNA2, LEP, PKLR, PLEKHA4, PMP22, PTGS2, RBP1, S100A13, SCARB1, TRIM63 | 2.83E-05 (31) | APOA2, C5AR1, CD14, CIDEA, CTH, CXCL10, CYP7B1, DBNDD1, FABP3, FABP6, GCK, GLP1R, IL1B, IL6, INS, KCNA2, KCNJ11, LEP, MTTP, MYL1, PKLR, PMP22, PNPLA3, PTGS2, RBP1, RETREG1, S100A13, SCARB1, SELENOP, SLC2A2, TRIM63 |

**P*-value: Benjamini Hochberg adjusted *P*-value; Z Scores > |2| are predicted to be target genes; n = number of target genes. N/A: prediction not available.

Table S7. Comparisons of differential types of immune cells in DMSO-treated rat placentas.*

| Cell subtype (Relative fraction) | Dendritic cells resting | NK cells resting | Macrophages M0 | B cells naive | T cells follicular helper | Macrophages M2 | T cells CD4 memory resting | Eosinophils | Monocytes | Plasma cells | T cells regulatory (Tregs) | T cells CD4 naive | T cells CD4 memory activated | Mast cells resting | Macrophages M1 | Mast cells activated | Neutrophils | NK cells activated | Dendritic cells activated | T cells gamma delta | T cells CD8 | B cells memory |
|---|---|---|---|---|---|---|---|---|---|---|---|---|---|---|---|---|---|---|---|---|---|---|
| Dendritic cells resting (0.298) | | γ | γ | γ | γ | γ | γ | γ | γ | γ | γ | γ | γ | γ | γ | γ | γ | γ | γ | γ | γ | γ |
| NK cells resting (0.155) | | γ | γ | γ | γ | γ | γ | γ | γ | γ | γ | γ | γ | γ | γ | γ | γ | γ | γ | γ | γ | γ |
| Macrophages M0 (0.155) | | γ | γ | γ | γ | γ | γ | γ | γ | γ | γ | γ | γ | γ | γ | γ | γ | γ | γ | γ | γ | γ |
| B cells naïve (0.083) | | γ | γ | γ | γ | γ | γ | γ | γ | γ | γ | γ | γ | γ | γ | γ | γ | γ | γ | γ | γ | γ |
| T cells follicular helper (0.079) | | | | | | ε | γ | γ | γ | γ | γ | γ | γ | γ | γ | γ | γ | γ | γ | γ | γ | γ |
| Macrophages M2 (0.056) | | | | | | | ε | γ | γ | γ | γ | γ | γ | γ | γ | γ | γ | γ | γ | γ | γ | γ |
| T cells CD4 memory resting (0.033) | | | | | | | | γ | γ | γ | γ | γ | γ | γ | γ | γ | γ | γ | γ | γ | γ | γ |
| Eosinophils (0.028) | | | | | | | | | | | | | | | ε | γ | γ | γ | γ | γ | γ | γ |
| Monocytes (0.021) | | | | | | | | | | | | | | | | | | | | | | |
| Plasma cells (0.020) | | | | | | | | | | | | | | | | | | | | | | |
| T cells CD4 naïve (0.019) | | | | | | | | | | | | | | | | | | | | | | |
| T cells regulatory (Tregs) (0.017) | | | | | | | | | | | | | | | | | | | | | | |
| T cells CD4 memory activated (0.017) | | | | | | | | | | | | | | | | | | | | | | |
| Mast cells resting (0.013) | | | | | | | | | | | | | | | | | | | | | | |
| Macrophages M1 (0.004) | | | | | | | | | | | | | | | | | | | | | | |
| Mast cells activated (0.001) | | | | | | | | | | | | | | | | | | | | | | |
| Neutrophils (0.000) | | | | | | | | | | | | | | | | | | | | | | |
| NK cells activated (0.000) | | | | | | | | | | | | | | | | | | | | | | |
| Dendritic cells activated (0.000) | | | | | | | | | | | | | | | | | | | | | | |
| T cells gamma delta (0.000) | | | | | | | | | | | | | | | | | | | | | | |
| T cells CD8 (0.000) | | | | | | | | | | | | | | | | | | | | | | |
| B cells memory (0.000) | | | | | | | | | | | | | | | | | | | | | | |

*The one-way ANOVA with all pairwise multiple comparison was performed using Holm-Sidak tests. $^{\varepsilon}P < 0.01$, $^{\gamma}P < 0.001$ vs corresponding cell type. n = 29 (13 female and 16 male) placentas.

Table S8. ITE-dysregulated immune cell distribution in rat placentas.*

| | Combined Female and Male | | Female | | Male | |
|---|---|---|---|---|---|---|
| | DMSO | ITE | DMSO | ITE | DMSO | ITE |
| B cells naive | 0.083±0.004 | 0.089±0.006 | 0.084±0.007 | 0.102±0.008 | 0.081±0.006 | 0.082±0.009 |
| B cells memory | 0.000±0.000 | 0.000±0.000 | 0.000±0.000 | 0.000±0.000 | 0.000±0.000 | 0.000±0.000 |
| Plasma cells | 0.020±0.003 | 0.042±0.005$^{\varepsilon}$ | 0.016±0.004 | 0.039±0.009 | 0.023±0.004 | 0.044±0.007 |
| T cells CD8 | 0.000±0.000 | 0.000±0.000 | 0.000±0.000 | 0.000±0.000 | 0.000±0.000 | 0.000±0.000 |
| T cells CD4 naive | 0.019±0.004 | 0.029±0.006 | 0.021±0.005 | 0.025±0.010 | 0.018±0.005 | 0.032±0.007 |
| T cells CD4 memory resting | 0.033±0.006 | 0.027±0.007 | 0.035±0.009 | 0.034±0.016 | 0.031±0.009 | 0.023±0.007 |
| T cells CD4 memory activated | 0.017±0.003 | 0.017±0.004 | 0.010±0.004 | 0.017±0.006 | 0.022±0.005 | 0.016±0.005 |
| T cells follicular helper | 0.079±0.004 | 0.046±0.005$^{\gamma}$ | 0.072±0.004 | 0.060±0.008 | 0.085±0.006 | 0.038±0.006$^{\gamma}$ |
| T cells regulatory (Tregs) | 0.017±0.002 | 0.005±0.003 | 0.021±0.003 | 0.008±0.005 | 0.015±0.003 | 0.003±0.003 |
| T cells gamma delta | 0.000±0.000 | 0.000±0.000 | 0.000±0.000 | 0.000±0.000 | 0.000±0.000 | 0.000±0.000 |
| NK cells resting | 0.155±0.006 | 0.117±0.006$^{\gamma}$ | 0.162±0.010 | 0.129±0.013$^{\varepsilon}$ | 0.149±0.007 | 0.109±0.006$^{\varepsilon}$ |
| NK cells activated | 0.000±0.000 | 0.000±0.000 | 0.000±0.000 | 0.000±0.000 | 0.000±0.000 | 0.000±0.000 |
| Monocytes | 0.021±0.005 | 0.056±0.010$^{\gamma}$ | 0.025±0.010 | 0.045±0.020 | 0.018±0.005 | 0.063±0.012$^{\gamma}$ |
| Macrophages M0 | 0.155±0.006 | 0.192±0.017$^{\gamma}$ | 0.160±0.006 | 0.185±0.025$^{\gamma}$ | 0.151±0.009 | 0.196±0.023$^{\gamma}$ |
| Macrophages M1 | 0.004±0.000 | 0.005±0.001 | 0.004±0.001 | 0.006±0.002 | 0.004±0.001 | 0.004±0.002 |
| Macrophages M2 | 0.056±0.008 | 0.126±0.010$^{\gamma}$ | 0.045±0.012 | 0.102±0.014$^{\gamma}$ | 0.065±0.010 | 0.141±0.012$^{\gamma,\delta}$ |
| Dendritic cells resting | 0.298±0.010 | 0.185±0.018$^{\gamma}$ | 0.301±0.014 | 0.190±0.021$^{\gamma}$ | 0.295±0.016$^{\delta}$ | 0.181±0.026$^{\gamma}$ |
| Dendritic cells activated | 0.000±0.000 | 0.000±0.000 | 0.000±0.000 | 0.000±0.000 | 0.000±0.000 | 0.000±0.000 |
| Mast cells resting | 0.013±0.003 | 0.002±0.001 | 0.012±0.004 | 0.001±0.001 | 0.013±0.004 | 0.002±0.002 |
| Mast cells activated | 0.001±0.000 | 0.011±0.004 | 0.001±0.001 | 0.007±0.004 | 0.001±0.001 | 0.014±0.005 |
| Eosinophils | 0.028±0.003 | 0.049±0.003$^{£}$ | 0.031±0.005 | 0.050±0.006 | 0.027±0.003 | 0.049±0.004 |
| Neutrophils | 0.001±0.000 | 0.002±0.002 | 0.001±0.000 | 0.003±0.002 | 0.000±0.000 | 0.002±0.002 |

*The two-way ANOVA with all pairwise multiple comparison was performed using Holm-Sidak tests. Data are expressed as means ± SEM relative cell fractions. $^{\varepsilon}P < 0.03$, $^{\gamma}P < 0.001$ vs DMSO within the same sex group. $^{\delta}P < 0.02$ vs corresponding female. n = 13, 16, 8, and 13 placentas for female DMSO, male DMSO, female ITE, and male ITE, respectively. P < 0.01, P < 0.001

Table S9. Patient demographics for HUVECs used in RNA-seq, RT-qPCR, biofunctional, and phosphoproteomics analyses*.

| | Fetal sex | n | Maternal BMI | Maternal age (years) | Gestational age (weeks) | Fetal weight (gram) | Systolic BP (mmHg) | Diastolic BP (mmHg) |
|---|---|---|---|---|---|---|---|---|
| RNA-seq | Female | 6 | 23.8 ± 0.4 | 33.6 ± 1.9 | 39.6 ± 0.2 | 3495.1 ± 93.2 | 105.9 ± 4.7 | 66.1 ± 5.0 |
| | Male | 5 | 23.7 ± 1.1 | 30.8 ± 1.3 | 39.5 ± 0.6 | 3660.0 ± 154.9 | 112.8 ± 1.9 | 68.8 ± 2.2 |
| | *P*-value | | > 0.05 | > 0.05 | > 0.05 | > 0.05 | > 0.05 | > 0.05 |
| Biofunctional and RT-qPCR assays | Female | 7 | 23.8 ± 0.4 | 32.8 ± 1.8 | 39.7 ± 0.2 | 3611.5 ± 140.6 | 106.3 ± 4.0 | 67.5 ± 4.4 |
| | Male | 7 | 23.8 ± 0.8 | 30.4 ± 1.1 | 39.6 ± 0.5 | 3503.3 ± 148.5 | 115.9 ± 2.4 | 71.7 ± 2.8 |
| | *P*-value | | > 0.05 | > 0.05 | > 0.05 | > 0.05 | > 0.05 | > 0.05 |
| Phosphoproteomics | Female | 2 | 24.7 ± 3.8 | 31.0 ± 4.1 | 39.1 ± 1.3 | 3292.3 ± 186.6 | 124.3 ± 5.4 | 76.5 ± 2.6 |
| | Male | 2 | | | | | | |

*BMI: Body mass index, BP: Blood pressure, RNA-seq: RNA-sequencing, RT-qPCR, Reverse transcription-quantitative PCR. All patients are Caucasians and without current or history of other major complications.

Table S10. ITE-dysregulated genes on HUVECs.

| Gene ID | Gene location on sex chromosome | Gene symbol | Female HUVECs | | Male HUVECs | | Comments |
|---|---|---|---|---|---|---|---|
| | | | log2 (ITE/Ctrl) | FDR-adjusted *P*-Value | log2 (ITE/Ctrl) | FDR-adjusted *P*-Value | |
| ENSG00000140465 | | CYP1A1 | 9.55 | 0.00E+00 | 9.62 | 7.98E-21 | DE in both female and male HUVECs |
| ENSG00000138061 | | CYP1B1 | 7.73 | 0.00E+00 | 6.25 | 6.74E-21 | DE in both female and male HUVECs |
| ENSG00000171246 | | NPTX1 | 7.09 | 0.00E+00 | 2.81 | 1.41E-07 | DE in both female and male HUVECs |
| ENSG00000114812 | | VIPR1 | 3.98 | 9.40E-78 | 4.24 | 4.14E-41 | DE in both female and male HUVECs |
| ENSG00000260349 | | AC087190.2 | 3.78 | 3.50E-04 | 6.72 | 3.36E-04 | DE in both female and male HUVECs |
| ENSG00000127533 | | F2RL3 | 3.63 | 6.55E-32 | 1.88 | 2.32E-02 | DE in both female and male HUVECs |
| ENSG00000137809 | | ITGA11 | 3.34 | 5.53E-214 | 3.56 | 1.36E-65 | DE in both female and male HUVECs |
| ENSG00000196611 | | MMP1 | 2.83 | 0.00E+00 | 3.39 | 1.16E-27 | DE in both female and male HUVECs |
| ENSG00000197632 | | SERPINB2 | 2.71 | 2.14E-105 | 2.78 | 7.76E-32 | DE in both female and male HUVECs |
| ENSG00000180730 | | SHISA2 | 2.66 | 6.73E-77 | 2.66 | 2.20E-35 | DE in both female and male HUVECs |
| ENSG00000117525 | | F3 | 2.65 | 1.37E-53 | 1.97 | 3.39E-03 | DE in both female and male HUVECs |
| ENSG00000205038 | | PKHD1L1 | 2.64 | 5.31E-171 | 1.79 | 3.43E-03 | DE in both female and male HUVECs |
| ENSG00000154864 | | PIEZO2 | 2.61 | 8.58E-193 | 2.97 | 3.98E-82 | DE in both female and male HUVECs |
| ENSG00000126353 | | CCR7 | 2.50 | 1.15E-03 | 3.87 | 4.48E-02 | DE in both female and male HUVECs |
| ENSG00000135914 | | HTR2B | 2.16 | 8.30E-81 | 1.42 | 1.76E-16 | DE in both female and male HUVECs |

| | | | | | | | |
|---|---|---|---|---|---|---|---|
| ENSG00000063438 | | AHRR | 2.08 | 1.03E-80 | 1.98 | 2.56E-23 | DE in both female and male HUVECs |
| ENSG00000125538 | | IL1B | 1.88 | 2.51E-06 | 2.37 | 2.83E-08 | DE in both female and male HUVECs |
| ENSG00000178695 | | KCTD12 | 1.79 | 5.77E-144 | 1.29 | 9.53E-10 | DE in both female and male HUVECs |
| ENSG00000163659 | | TIPARP | 1.69 | 4.41E-113 | 1.17 | 1.43E-15 | DE in both female and male HUVECs |
| ENSG00000172602 | | RND1 | 1.68 | 4.90E-41 | 1.28 | 3.71E-11 | DE in both female and male HUVECs |
| ENSG00000073756 | | PTGS2 | 1.54 | 5.90E-89 | 1.10 | 2.21E-20 | DE in both female and male HUVECs |
| ENSG00000101670 | | LIPG | 1.45 | 9.34E-74 | 1.89 | 8.71E-03 | DE in both female and male HUVECs |
| ENSG00000211445 | | GPX3 | 1.44 | 6.05E-37 | 1.33 | 4.94E-04 | DE in both female and male HUVECs |
| ENSG00000182326 | | C1S | 1.39 | 3.62E-10 | 1.13 | 1.78E-05 | DE in both female and male HUVECs |
| ENSG00000007908 | | SELE | 1.32 | 5.18E-73 | 1.15 | 1.42E-20 | DE in both female and male HUVECs |
| ENSG00000176771 | | NCKAP5 | 1.31 | 1.01E-08 | 1.70 | 7.25E-06 | DE in both female and male HUVECs |
| ENSG00000019991 | | HGF | 1.31 | 4.81E-22 | 1.19 | 1.70E-09 | DE in both female and male HUVECs |
| ENSG00000170891 | | CYTL1 | 1.22 | 1.31E-21 | 1.78 | 9.53E-05 | DE in both female and male HUVECs |
| ENSG00000151789 | | ZNF385D | 1.22 | 5.28E-09 | 1.22 | 1.55E-02 | DE in both female and male HUVECs |
| ENSG00000108342 | | CSF3 | 1.20 | 1.05E-17 | 1.06 | 1.16E-07 | DE in both female and male HUVECs |
| ENSG00000102385 | X | DRP2 | 1.18 | 5.64E-04 | 1.91 | 6.30E-03 | DE in both female and male HUVECs |
| ENSG00000179151 | | EDC3 | 1.15 | 3.05E-40 | 1.00 | 7.31E-15 | DE in both female and male HUVECs |
| ENSG00000166825 | | ANPEP | 1.08 | 5.11E-52 | 1.11 | 7.69E-22 | DE in both female and male HUVECs |

| | | | | | | | |
|---|---|---|---|---|---|---|---|
| ENSG00000151012 | | SLC7A11 | 1.02 | 1.79E-46 | 1.04 | 7.47E-13 | DE in both female and male HUVECs |
| ENSG00000106003 | | LFNG | -1.01 | 6.81E-07 | -1.16 | 8.68E-04 | DE in both female and male HUVECs |
| ENSG00000106852 | | LHX6 | -1.09 | 7.97E-14 | -1.04 | 2.68E-09 | DE in both female and male HUVECs |
| ENSG00000187513 | | GJA4 | -1.10 | 2.65E-18 | -1.26 | 6.50E-18 | DE in both female and male HUVECs |
| ENSG00000164692 | | COL1A2 | -1.11 | 1.55E-37 | -1.38 | 6.48E-30 | DE in both female and male HUVECs |
| ENSG00000271447 | | MMP28 | -1.33 | 2.93E-08 | -1.02 | 1.82E-03 | DE in both female and male HUVECs |
| ENSG00000240583 | | AQP1 | -1.57 | 1.83E-92 | -1.68 | 2.26E-04 | DE in both female and male HUVECs |
| ENSG00000248187 | | AC078850.1 | -1.66 | 1.53E-20 | -1.42 | 3.37E-10 | DE in both female and male HUVECs |
| ENSG00000133110 | | POSTN | -1.88 | 6.71E-54 | -1.80 | 1.70E-23 | DE in both female and male HUVECs |
| ENSG00000276278 | | AC048382.6 | 1.64 | 6.69E-03 | -5.18 | 1.12E-05 | DE in both female and male HUVECs |
| ENSG00000157654 | | PALM2-AKAP2 | -3.77 | 3.99E-08 | 4.50 | 4.37E-02 | DE in both female and male HUVECs |
| ENSG00000262660 | | AC139530.2 | 6.36 | 3.53E-05 | | | DE only in female HUVECs |
| ENSG00000272679 | | AL355987.2 | 5.09 | 8.18E-03 | | | DE only in female HUVECs |
| ENSG00000224533 | X | TMLHE-AS1 | 5.04 | 5.85E-03 | | | DE only in female HUVECs |
| ENSG00000265818 | | EEF1E1-BLOC1S5 | 4.67 | 4.82E-02 | | | DE only in female HUVECs |
| ENSG00000163075 | | CFAP221 | 3.41 | 6.89E-04 | | | DE only in female HUVECs |
| ENSG00000274092 | | AC106739.1 | 3.32 | 1.89E-02 | | | DE only in female HUVECs |
| ENSG00000120251 | | GRIA2 | 3.06 | 1.49E-05 | | | DE only in female HUVECs |
| ENSG00000256861 | | AC048338.1 | 2.85 | 9.13E-03 | | | DE only in female HUVECs |
| ENSG00000265257 | | AP005263.1 | 2.77 | 2.17E-02 | | | DE only in female HUVECs |
| ENSG00000248710 | | AC079594.2 | 2.57 | 7.50E-04 | | | DE only in female HUVECs |
| ENSG00000213860 | | RPL21P75 | 2.50 | 2.32E-07 | | | DE only in female HUVECs |
| ENSG00000275857 | | AC009133.4 | 2.41 | 1.43E-03 | | | DE only in female HUVECs |
| ENSG00000128683 | | GAD1 | 2.31 | 4.31E-02 | | | DE only in female HUVECs |

| | | | | |
|---|---|---|---|---|
| ENSG00000147642 | SYBU | 2.04 | 2.00E-24 | DE only in female HUVECs |
| ENSG00000144785 | AC073896.1 | 1.89 | 4.09E-04 | DE only in female HUVECs |
| ENSG00000168542 | COL3A1 | 1.68 | 2.66E-74 | DE only in female HUVECs |
| ENSG00000178878 | APOLD1 | 1.65 | 3.42E-77 | DE only in female HUVECs |
| ENSG00000138356 | AOX1 | 1.65 | 5.62E-03 | DE only in female HUVECs |
| ENSG00000125965 | GDF5 | 1.64 | 1.16E-02 | DE only in female HUVECs |
| ENSG00000139329 | LUM | 1.60 | 2.90E-13 | DE only in female HUVECs |
| ENSG00000174600 | CMKLR1 | 1.59 | 1.89E-03 | DE only in female HUVECs |
| ENSG00000101680 | LAMA1 | 1.54 | 2.76E-02 | DE only in female HUVECs |
| ENSG00000151715 | TMEM45B | 1.52 | 4.66E-02 | DE only in female HUVECs |
| ENSG00000243509 | TNFRSF6B | 1.52 | 8.25E-08 | DE only in female HUVECs |
| ENSG00000260371 | AC026464.3 | 1.47 | 2.14E-05 | DE only in female HUVECs |
| ENSG00000259460 | AC078909.1 | 1.47 | 1.00E-02 | DE only in female HUVECs |
| ENSG00000171570 | RAB4B-EGLN2 | 1.43 | 2.86E-02 | DE only in female HUVECs |
| ENSG00000116711 | PLA2G4A | 1.30 | 1.36E-65 | DE only in female HUVECs |
| ENSG00000131203 | IDO1 | 1.28 | 7.51E-03 | DE only in female HUVECs |
| ENSG00000219410 | AC125494.1 | 1.25 | 7.15E-03 | DE only in female HUVECs |
| ENSG00000131386 | GALNT15 | 1.25 | 1.05E-55 | DE only in female HUVECs |
| ENSG00000140557 | ST8SIA2 | 1.22 | 2.61E-02 | DE only in female HUVECs |
| ENSG00000165092 | ALDH1A1 | 1.21 | 6.61E-67 | DE only in female HUVECs |
| ENSG00000128342 | LIF | 1.21 | 2.31E-02 | DE only in female HUVECs |
| ENSG00000140873 | ADAMTS18 | 1.20 | 1.03E-63 | DE only in female HUVECs |
| ENSG00000162337 | LRP5 | 1.19 | 2.04E-49 | DE only in female HUVECs |
| ENSG00000184292 | TACSTD2 | 1.19 | 9.49E-54 | DE only in female HUVECs |
| ENSG00000102575 | ACP5 | 1.19 | 1.73E-21 | DE only in female HUVECs |
| ENSG00000111912 | NCOA7 | 1.16 | 8.17E-59 | DE only in female HUVECs |
| ENSG00000159307 | SCUBE1 | 1.13 | 8.51E-08 | DE only in female HUVECs |
| ENSG00000185022 | MAFF | 1.13 | 1.74E-49 | DE only in female HUVECs |
| ENSG00000103888 | CEMIP | 1.12 | 2.18E-06 | DE only in female HUVECs |
| ENSG00000103489 | XYLT1 | 1.12 | 2.55E-13 | DE only in female HUVECs |
| ENSG00000171223 | JUNB | 1.11 | 1.29E-20 | DE only in female HUVECs |
| ENSG00000103241 | FOXF1 | 1.10 | 1.49E-04 | DE only in female HUVECs |
| ENSG00000164484 | TMEM200A | 1.09 | 3.95E-10 | DE only in female HUVECs |
| ENSG00000268861 | AC008878.3 | 1.07 | 2.16E-02 | DE only in female HUVECs |

| | | | | | |
|---|---|---|---|---|---|
| ENSG00000164619 | | BMPER | 1.07 | 1.83E-08 | DE only in female HUVECs |
| ENSG00000158683 | | PKD1L1 | 1.06 | 1.31E-21 | DE only in female HUVECs |
| ENSG00000171227 | | TMEM37 | 1.05 | 1.38E-15 | DE only in female HUVECs |
| ENSG00000254087 | | LYN | 1.05 | 3.84E-40 | DE only in female HUVECs |
| ENSG00000119714 | | GPR68 | 1.04 | 1.24E-02 | DE only in female HUVECs |
| ENSG00000205213 | | LGR4 | 1.03 | 3.12E-23 | DE only in female HUVECs |
| ENSG00000116678 | | LEPR | 1.02 | 6.38E-46 | DE only in female HUVECs |
| ENSG00000183091 | | NEB | 1.01 | 2.41E-12 | DE only in female HUVECs |
| ENSG00000176046 | | NUPR1 | 1.01 | 2.63E-09 | DE only in female HUVECs |
| ENSG00000005471 | | ABCB4 | 1.00 | 1.80E-02 | DE only in female HUVECs |
| ENSG00000179304 | X | FAM156B | 1.00 | 2.99E-07 | DE only in female HUVECs |
| ENSG00000257379 | | AC023509.1 | -1.00 | 3.98E-03 | DE only in female HUVECs |
| ENSG00000188158 | X | NHS | -1.01 | 3.64E-02 | DE only in female HUVECs |
| ENSG00000120669 | | SOHLH2 | -1.02 | 4.95E-02 | DE only in female HUVECs |
| ENSG00000090530 | | P3H2 | -1.03 | 5.68E-05 | DE only in female HUVECs |
| ENSG00000168843 | | FSTL5 | -1.04 | 2.22E-07 | DE only in female HUVECs |
| ENSG00000094755 | | GABRP | -1.05 | 4.81E-03 | DE only in female HUVECs |
| ENSG00000272695 | | GAS6-AS2 | -1.07 | 2.10E-03 | DE only in female HUVECs |
| ENSG00000134198 | | TSPAN2 | -1.08 | 1.53E-04 | DE only in female HUVECs |
| ENSG00000174348 | | PODN | -1.13 | 4.15E-02 | DE only in female HUVECs |
| ENSG00000259305 | | ZHX1-C8orf76 | -1.13 | 5.09E-03 | DE only in female HUVECs |
| ENSG00000265263 | | AC124066.1 | -1.14 | 8.96E-08 | DE only in female HUVECs |
| ENSG00000100336 | | APOL4 | -1.16 | 5.76E-04 | DE only in female HUVECs |
| ENSG00000205670 | | SMIM11A | -1.17 | 3.71E-10 | DE only in female HUVECs |
| ENSG00000183850 | | ZNF730 | -1.20 | 9.22E-03 | DE only in female HUVECs |
| ENSG00000165072 | | MAMDC2 | -1.22 | 1.44E-05 | DE only in female HUVECs |
| ENSG00000256007 | | ARAP1-AS1 | -1.34 | 3.39E-02 | DE only in female HUVECs |
| ENSG00000144278 | | GALNT13 | -1.40 | 3.56E-02 | DE only in female HUVECs |
| ENSG00000204644 | | ZFP57 | -1.46 | 1.55E-02 | DE only in female HUVECs |
| ENSG00000170965 | X | PLAC1 | -1.52 | 4.60E-02 | DE only in female HUVECs |
| ENSG00000175874 | | CREG2 | -1.55 | 3.82E-08 | DE only in female HUVECs |
| ENSG00000233327 | | USP32P2 | -1.67 | 5.67E-03 | DE only in female HUVECs |
| ENSG00000268894 | | PLCE1-AS1 | -1.69 | 1.46E-04 | DE only in female HUVECs |
| ENSG00000277991 | | FP236241.1 | -1.92 | 8.42E-06 | DE only in female HUVECs |

| | | | | | | |
|---|---|---|---|---|---|---|
| ENSG00000255073 | ZFP91-CNTF | -1.96 | 4.67E-03 | | | DE only in female HUVECs |
| ENSG00000141668 | CBLN2 | -2.03 | 4.26E-02 | | | DE only in female HUVECs |
| ENSG00000270898 | GPR75-ASB3 | -2.08 | 4.32E-02 | | | DE only in female HUVECs |
| ENSG00000258588 | TRIM6-TRIM34 | -2.15 | 4.65E-03 | | | DE only in female HUVECs |
| ENSG00000108700 | CCL8 | -2.45 | 5.64E-03 | | | DE only in female HUVECs |
| ENSG00000134817 | APLNR | -2.74 | 8.14E-16 | | | DE only in female HUVECs |
| ENSG00000281453 | TGFB2-OT1 | -2.77 | 4.33E-05 | | | DE only in female HUVECs |
| ENSG00000212643 | ZRSR1 | -3.02 | 3.28E-03 | | | DE only in female HUVECs |
| ENSG00000160683 | CXCR5 | -3.06 | 4.68E-03 | | | DE only in female HUVECs |
| ENSG00000275896 | PRSS2 | -3.21 | 2.11E-02 | | | DE only in female HUVECs |
| ENSG00000180440 | SERTM1 | -3.85 | 2.50E-02 | | | DE only in female HUVECs |
| ENSG00000283528 | AC073264.3 | -4.96 | 2.89E-02 | | | DE only in female HUVECs |
| ENSG00000267261 | AC099811.2 | -5.47 | 1.68E-07 | | | DE only in female HUVECs |
| ENSG00000244491 | AL021707.5 | -6.09 | 3.50E-03 | | | DE only in female HUVECs |
| ENSG00000261684 | AC018362.1 | -7.70 | 2.68E-07 | | | DE only in female HUVECs |
| ENSG00000259607 | AC108449.3 | | | 6.43 | 2.26E-02 | DE only in male HUVECs |
| ENSG00000279592 | AC010653.2 | | | 6.55 | 1.30E-03 | DE only in male HUVECs |
| ENSG00000198121 | LPAR1 | | | 1.65 | 2.55E-02 | DE only in male HUVECs |
| ENSG00000165124 | SVEP1 | | | 1.50 | 2.90E-02 | DE only in male HUVECs |
| ENSG00000136244 | IL6 | | | 1.53 | 3.16E-07 | DE only in male HUVECs |
| ENSG00000181019 | NQO1 | | | 1.07 | 2.56E-20 | DE only in male HUVECs |
| ENSG00000145287 | PLAC8 | | | 1.02 | 1.16E-07 | DE only in male HUVECs |
| ENSG00000108932 | SLC16A6 | | | 1.11 | 8.94E-04 | DE only in male HUVECs |
| ENSG00000159167 | STC1 | | | 1.14 | 2.04E-02 | DE only in male HUVECs |
| ENSG00000229644 | NAMPTP1 | | | 1.23 | 1.31E-16 | DE only in male HUVECs |
| ENSG00000278139 | AL358075.4 | | | 6.46 | 1.93E-02 | DE only in male HUVECs |
| ENSG00000272851 | AC096772.1 | | | 7.79 | 2.48E-06 | DE only in male HUVECs |
| ENSG00000273003 | ARL2-SNX15 | | | 8.77 | 4.17E-07 | DE only in male HUVECs |
| ENSG00000273398 | AC017083.4 | | | 4.06 | 2.27E-03 | DE only in male HUVECs |
| ENSG00000256364 | AC069234.2 | | | 6.79 | 1.96E-03 | DE only in male HUVECs |
| ENSG00000280416 | AC009084.3 | | | 7.23 | 3.46E-05 | DE only in male HUVECs |
| ENSG00000254673 | AC110275.1 | | | 8.36 | 5.23E-11 | DE only in male HUVECs |
| ENSG00000236810 | ELOA-AS1 | | | 1.20 | 1.19E-02 | DE only in male HUVECs |
| ENSG00000196302 | AC146944.1 | | | 4.10 | 3.46E-02 | DE only in male HUVECs |

| | | | | |
|---|---|---|---|---|
| ENSG00000255663 | AP002373.1 | 7.01 | 1.27E-03 | DE only in male HUVECs |
| ENSG00000279033 | AC090984.1 | 8.47 | 1.30E-09 | DE only in male HUVECs |
| ENSG00000282218 | AL132671.2 | 6.88 | 2.28E-02 | DE only in male HUVECs |
| ENSG00000196826 | AC008758.1 | -7.05 | 1.05E-02 | DE only in male HUVECs |
| ENSG00000207008 | SNORA54 | -2.38 | 2.90E-02 | DE only in male HUVECs |
| ENSG00000187608 | ISG15 | -1.05 | 2.52E-06 | DE only in male HUVECs |
| ENSG00000267740 | AC024592.3 | -2.21 | 5.51E-03 | DE only in male HUVECs |
| ENSG00000258644 | SYNJ2BP-COX16 | -1.11 | 1.78E-05 | DE only in male HUVECs |
| ENSG00000152583 | SPARCL1 | -1.59 | 6.99E-06 | DE only in male HUVECs |
| ENSG00000207166 | SNORA68 | -2.18 | 2.65E-02 | DE only in male HUVECs |
| ENSG00000134321 | RSAD2 | -1.13 | 8.68E-14 | DE only in male HUVECs |
| ENSG00000187955 | COL14A1 | -1.49 | 4.60E-03 | DE only in male HUVECs |
| ENSG00000169245 | CXCL10 | -1.06 | 8.39E-06 | DE only in male HUVECs |
| ENSG00000169129 | AFAP1L2 | -1.09 | 1.75E-04 | DE only in male HUVECs |
| ENSG00000049540 | ELN | -1.80 | 6.10E-04 | DE only in male HUVECs |
| ENSG00000163815 | CLEC3B | -2.26 | 2.50E-05 | DE only in male HUVECs |
| ENSG00000249967 | AL355315.1 | -4.30 | 1.34E-02 | DE only in male HUVECs |

*Ctrl: vehicle control (DMSO); *P*-value: Benjamini Hochberg adjusted *P*-value. DE: differentially expressed. $P < 0.05$ is considered significantly differentially expressed. n = 6 and 5 female and male HUVECs preparations.

Table S11. ITE-dysregulated biological function in HUVECs.*

| | *P*-Value [Female HUVECs] | *P*-Value [Male HUVECs] |
|---|---|---|
| Development of vasculature | **1.34E-09** | **1.81E-10** |
| Angiogenesis | **2.66E-09** | **1.81E-10** |
| Anogenital cancer | **5.63E-11** | **8.43E-09** |
| Cell movement | **1.47E-11** | **1.33E-08** |
| Vasculogenesis | **1.59E-08** | **1.33E-08** |
| Fibrosis | **4.63E-08** | **1.33E-08** |
| Migration of cells | **3.67E-11** | **2.09E-08** |
| Genital tract cancer | **6.28E-10** | **2.09E-08** |
| Pelvic cancer | **2.88E-09** | **2.32E-08** |
| Genital tumor | **4.75E-10** | **3.15E-08** |
| Migration of tumor cells | **5.58E-07** | **3.61E-08** |
| Malignant genitourinary solid tumor | **3.52E-10** | **3.92E-08** |
| Aggregation of cells | **3.95E-06** | **7.31E-08** |
| Psoriasis | **6.22E-04** | **8.15E-08** |
| Recruitment of cells | **3.08E-08** | **1.01E-07** |
| Leukocyte migration | **1.58E-07** | **1.18E-07** |
| Benign pelvic disease | **3.52E-10** | **1.28E-07** |
| Periodontal disease | **6.58E-05** | **1.73E-07** |
| Periodontitis | **7.32E-04** | **2.65E-07** |
| Aggregation of blood cells | **1.04E-06** | **3.30E-07** |
| Interaction of mononuclear leukocytes | **2.48E-05** | **3.38E-07** |
| Proliferation of muscle cells | **3.50E-04** | **3.42E-07** |
| Invasive tumor | **1.50E-06** | **3.53E-07** |
| Rheumatic Disease | **1.51E-08** | **4.54E-07** |
| Growth of vessel | **1.08E-03** | **4.54E-07** |
| Survival of organism | **2.55E-05** | **5.38E-07** |
| Female genital tract serous carcinoma | **4.26E-04** | **5.38E-07** |
| Pelvic serous carcinoma | **4.26E-04** | **5.38E-07** |
| Lymphoreticular neoplasm | **1.86E-04** | **6.35E-07** |
| Cell movement of monocytes | **2.30E-04** | **6.35E-07** |
| Cell movement of myeloid cells | **3.98E-05** | **6.58E-07** |
| Development of epithelial tissue | **2.47E-05** | **6.63E-07** |
| Endothelial cell development | **1.60E-04** | **6.63E-07** |
| Vaso-occlusion | **9.57E-04** | **6.63E-07** |
| Morphology of bone | **1.45E-08** | **7.46E-07** |
| Hematologic cancer of cells | **5.21E-04** | **7.46E-07** |
| Breast or gynecological cancer | **2.20E-10** | **7.49E-07** |
| Female genital tract cancer | **7.10E-10** | **8.15E-07** |
| Chemotaxis | **9.82E-06** | **9.38E-07** |
| Infection by coronavirus | **2.26E-04** | **9.38E-07** |
| Neoplasia of blood cells | **5.78E-06** | **9.64E-07** |
| Recruitment of neutrophils | **7.04E-05** | **9.82E-07** |

| | | |
|---|---|---|
| Hematologic cancer | **9.16E-05** | **1.14E-06** |
| Breast or pancreatic cancer | **6.21E-09** | **1.21E-06** |
| Breast or ovarian cancer | **3.36E-07** | **1.21E-06** |
| Recruitment of phagocytes | **6.42E-07** | **1.21E-06** |
| Tumorigenesis of reproductive tract | **6.28E-10** | **1.25E-06** |
| Myeloid or lymphoid neoplasm | **8.94E-06** | **1.26E-06** |
| Binding of mononuclear leukocytes | **9.41E-05** | **1.26E-06** |
| Extrapancreatic malignant tumor | **7.26E-11** | **1.27E-06** |
| Uterine tumor | **9.06E-10** | **1.27E-06** |
| Organismal death | **5.38E-09** | **1.27E-06** |
| Recruitment of leukocytes | **3.36E-07** | **1.27E-06** |
| Migration of vascular cells | **6.50E-04** | **1.27E-06** |
| Diabetes mellitus | **9.47E-04** | **1.34E-06** |
| Hypertrophy | **2.51E-06** | **1.52E-06** |
| COVID-19 | **9.81E-04** | **1.52E-06** |
| Development of genital tumor | **6.28E-10** | **2.16E-06** |
| Recruitment of myeloid cells | **9.70E-06** | **2.20E-06** |
| Quantity of cells | **3.92E-06** | **2.53E-06** |
| Activation of endothelial cells | **4.94E-04** | **2.55E-06** |
| Priming of blood cells | **4.94E-04** | **2.55E-06** |
| Neoplasia of leukocytes | **9.28E-04** | **2.62E-06** |
| Accumulation of blood cells | **3.00E-04** | **2.71E-06** |
| Cell movement of phagocytes | **4.29E-04** | **2.74E-06** |
| Benign oral disorder | **5.58E-06** | **2.74E-06** |
| Binding of blood cells | **1.18E-05** | **2.83E-06** |
| Interaction of phagocytes | **4.34E-05** | **2.98E-06** |
| Migration of granulocytes | **8.22E-04** | **2.98E-06** |
| Breast cancer | **6.92E-07** | **3.04E-06** |
| Pelvic carcinoma | **7.83E-08** | **3.20E-06** |
| Quantity of phagocytes | **1.50E-05** | **3.45E-06** |
| Cell movement of colorectal cancer cell lines | **9.20E-04** | **3.45E-06** |
| Growth of blood vessel | **1.23E-03** | **3.65E-06** |
| Lymphocytic cancer | **1.34E-03** | **3.65E-06** |
| Lymphocytic neoplasm | **1.36E-03** | **3.70E-06** |
| Serous adenocarcinoma | **5.15E-04** | **3.80E-06** |
| Disorder of blood pressure | **1.18E-05** | **4.31E-06** |
| Hypertrophy of cells | **1.25E-04** | **4.44E-06** |
| Function of phagocytes | **2.93E-07** | **4.53E-06** |
| Interaction of lymphocytes | **1.20E-03** | **4.53E-06** |
| Genitourinary carcinoma | **1.54E-08** | **4.64E-06** |
| Cell movement of endothelial cells | **7.92E-04** | **4.90E-06** |
| Fibrogenesis | **6.43E-06** | **5.12E-06** |
| Development of abdomen | **3.41E-06** | **5.14E-06** |
| Quantity of neutrophils | **4.06E-04** | **5.31E-06** |
| Growth of connective tissue | **8.16E-06** | **5.33E-06** |

| | | |
|---|---|---|
| Binding of leukocytes | **1.43E-05** | **5.44E-06** |
| Healing of lesion | **5.05E-05** | **5.44E-06** |
| Migration of cancer cells | **5.38E-05** | **5.86E-06** |
| Extracranial solid tumor | **1.02E-10** | **6.34E-06** |
| Uterine cancer | **3.13E-08** | **6.58E-06** |
| Chronic renal impairment | **3.48E-04** | **6.73E-06** |
| Malignant solid tumor | **1.19E-10** | **7.08E-06** |
| Metastasis | **2.91E-05** | **7.15E-06** |
| Proliferation of endothelial cells | **2.30E-04** | **7.56E-06** |
| Formation of solid tumor | **3.67E-11** | **7.58E-06** |
| Quantity of granulocytes | **2.30E-04** | **7.58E-06** |
| Activation of blood cells | **9.27E-05** | **7.86E-06** |
| Vascularization of absolute anatomical region | **4.93E-06** | **7.86E-06** |
| Disorder of pregnancy | **4.57E-04** | **7.86E-06** |
| Cytokine and chemokine mediated signaling pathway | **4.23E-04** | **8.07E-06** |
| Abdominal cancer | **1.47E-11** | **8.19E-06** |
| Quantity of blood cells | **7.18E-04** | **8.19E-06** |
| Cell movement of mononuclear leukocytes | **6.16E-04** | **8.28E-06** |
| Quantity of leukocytes | **5.14E-04** | **9.54E-06** |
| Immune response of leukocytes | **2.94E-04** | **9.57E-06** |
| Accumulation of leukocytes | **7.55E-04** | **9.57E-06** |
| Proliferation of connective tissue cells | **9.60E-06** | **1.02E-05** |
| Inflammation of joint | **4.38E-08** | **1.06E-05** |
| Hypertension | **2.08E-05** | **1.06E-05** |
| Genitourinary adenocarcinoma | **2.27E-06** | **1.11E-05** |
| Endometriosis | **3.80E-06** | **1.15E-05** |
| Inflammatory response | **1.15E-06** | **1.17E-05** |
| Hypertrophy of tissue | **8.98E-04** | **1.18E-05** |
| Cellular infiltration by myeloid cells | **1.30E-03** | **1.18E-05** |
| Cell surface receptor linked signal transduction | **4.01E-06** | **1.19E-05** |
| Adhesion of mononuclear leukocytes | **6.98E-05** | **1.19E-05** |
| Homing of mononuclear leukocytes | **1.47E-04** | **1.29E-05** |
| Cell movement of squamous cell carcinoma cell lines | **3.53E-06** | **1.31E-05** |
| Adenocarcinoma | **3.52E-10** | **1.32E-05** |
| Invasion of cells | **1.17E-03** | **1.32E-05** |
| Muscular hypertrophy | **7.28E-04** | **1.34E-05** |
| Length of skeleton | **2.26E-04** | **1.39E-05** |
| Mass of bone organ | **2.26E-04** | **1.39E-05** |
| Chemotaxis of monocytes | **3.33E-04** | **1.39E-05** |
| Function of antigen presenting cells | **6.06E-07** | **1.43E-05** |
| Differentiation of connective tissue cells | **1.56E-05** | **1.43E-05** |
| Cell movement of leukocytes | **6.98E-05** | **1.46E-05** |
| Hypertrophy of heart | **2.85E-05** | **1.52E-05** |

| | | |
|---|---|---|
| Chemotaxis of leukocytes | **1.39E-04** | **1.56E-05** |
| Adhesion of immune cells | **3.03E-05** | **1.58E-05** |
| Size of skeleton | **2.55E-04** | **1.61E-05** |
| Abnormal morphology of thoracic cavity | **8.93E-06** | **1.79E-05** |
| Cell viability of phagocytes | **4.26E-04** | **1.79E-05** |
| Aggregation of blood platelets | **9.08E-07** | **1.87E-05** |
| Cellular infiltration by leukocytes | **1.93E-05** | **1.88E-05** |
| Formation of blood vessel | **1.25E-04** | **1.88E-05** |
| Chemotaxis of myeloid cells | **9.13E-05** | **1.90E-05** |
| Development of connective tissue cells | **3.81E-04** | **1.90E-05** |
| Visceromegaly | **1.52E-05** | **1.94E-05** |
| Quantity of myeloid cells | **5.15E-04** | **2.04E-05** |
| Immune response of myeloid cells | **1.10E-03** | **2.09E-05** |
| Wound | **8.16E-06** | **2.12E-05** |
| Chemotaxis of phagocytes | **1.04E-04** | **2.13E-05** |
| Systemic autoimmune syndrome | **9.81E-05** | **2.14E-05** |
| Vascularization | **5.31E-05** | **2.25E-05** |
| Blood clot | **2.79E-04** | **2.28E-05** |
| Localization of cells | **3.58E-04** | **2.28E-05** |
| Development of body trunk | **6.49E-06** | **2.33E-05** |
| Abdominal adenocarcinoma | **3.52E-10** | **2.39E-05** |
| Neutrophilia of blood | **1.59E-04** | **2.40E-05** |
| Malignant neoplasm of large intestine | **1.34E-07** | **2.43E-05** |
| Healing of wound | **1.77E-04** | **2.43E-05** |
| Invasion of tumor cells | **1.77E-04** | **2.43E-05** |
| Incidence of tumor | **1.31E-08** | **2.63E-05** |
| Cancer of secretory structure | **1.46E-06** | **2.71E-05** |
| Pelvic adenocarcinoma | **2.27E-06** | **2.71E-05** |
| Cell movement of carcinoma cell lines | **1.49E-04** | **2.77E-05** |
| Chronic renal failure | **1.00E-03** | **2.77E-05** |
| Osteoarthritis | **3.60E-08** | **2.82E-05** |
| Progression of digestive organ tumor | **2.95E-04** | **2.84E-05** |
| Non-traumatic arthropathy | **1.58E-07** | **2.94E-05** |
| Activation of vascular endothelial cells | **3.23E-04** | **3.13E-05** |
| Cell movement of tumor cells | **2.35E-04** | **3.22E-05** |
| Lower respiratory tract disorder | **1.87E-05** | **3.23E-05** |
| Intraabdominal organ tumor | **3.06E-10** | **3.24E-05** |
| Inflammation of organ | **2.20E-07** | **3.45E-05** |
| Immune response of phagocytes | **4.26E-04** | **3.45E-05** |
| Activation of cells | **1.62E-06** | **3.62E-05** |
| Migration of tumor cell lines | **2.30E-04** | **3.62E-05** |
| Cell movement of granulocytes | **3.58E-04** | **3.71E-05** |
| Induction of cells | **8.78E-04** | **3.71E-05** |
| Function of myeloid cells | **4.66E-05** | **3.78E-05** |
| Gastrointestinal adenocarcinoma | **1.08E-08** | **4.18E-05** |

| | | |
|---|---|---|
| Growth of epithelial tissue | **1.55E-06** | **4.21E-05** |
| Lung cancer | **5.16E-06** | **4.37E-05** |
| Chemotaxis of mononuclear leukocytes | **3.23E-04** | **4.44E-05** |
| Nonsquamous non-small cell lung carcinoma | **2.07E-06** | **4.48E-05** |
| Ovarian tumor | **2.00E-05** | **4.48E-05** |
| Response of myeloid leukocytes | **1.11E-03** | **4.74E-05** |
| Breast or colorectal cancer | **3.36E-07** | **4.82E-05** |
| Ulceration of skin | **7.66E-04** | **4.83E-05** |
| Development of lung tumor | **6.99E-06** | **4.96E-05** |
| Abnormality of left ventricle | **6.41E-04** | **5.03E-05** |
| Cell movement of neutrophils | **1.04E-03** | **5.05E-05** |
| Cell movement of tumor cell lines | **9.70E-06** | **5.07E-05** |
| Invasive carcinoma | **7.21E-05** | **5.17E-05** |
| Abdominal carcinoma | **2.76E-10** | **5.18E-05** |
| Chronic inflammatory disorder | **9.08E-07** | **5.20E-05** |
| Formation of filaments | **3.81E-05** | **5.27E-05** |
| Deterioration of connective tissue | **1.50E-06** | **5.27E-05** |
| Alcohol use disorder | **1.26E-03** | **5.28E-05** |
| Metastatic solid tumor | **5.22E-05** | **5.29E-05** |
| Binding of embryonic cells | **8.38E-04** | **5.29E-05** |
| Malignant neoplasm of retroperitoneum | **1.92E-04** | **5.30E-05** |
| Gastrointestinal carcinoma | **1.59E-08** | **5.52E-05** |
| HER2 negative cancer | **4.33E-04** | **5.84E-05** |
| Invasive breast carcinoma | **1.94E-04** | **5.86E-05** |
| HER2 negative solid tumor | **4.43E-04** | **5.86E-05** |
| Function of leukocytes | **1.11E-05** | **5.96E-05** |
| Malignant neoplasm of respiratory system | **5.24E-07** | **6.02E-05** |
| Mineralization of cells | **1.01E-03** | **6.36E-05** |
| Progression of tumor | **2.26E-04** | **6.69E-05** |
| Non-small cell lung carcinoma | **6.71E-06** | **6.86E-05** |
| Digestive system cancer | **6.28E-10** | **6.87E-05** |
| Immune response of neutrophils | **1.08E-03** | **6.87E-05** |
| Expansion of granulocytes | **4.50E-04** | **7.02E-05** |
| Growth of tumor | **2.34E-05** | **7.07E-05** |
| Migration of carcinoma cell lines | **2.40E-04** | **7.15E-05** |
| Cell movement of cancer cells | **3.58E-04** | **8.45E-05** |
| Demethylation of 2-hydroxyestradiol | **3.47E-04** | **8.52E-05** |
| Metabolism of eicosanoid | **3.00E-04** | **8.75E-05** |
| Migration of squamous cell carcinoma cell lines | **1.93E-05** | **9.03E-05** |
| Peripheral vascular disease | **3.08E-04** | **9.09E-05** |
| Extraadrenal retroperitoneal tumor | **3.12E-05** | **9.25E-05** |
| Endometrial cancer | **6.86E-08** | **9.44E-05** |
| Morphology of body cavity | **5.08E-07** | **9.82E-05** |
| Enlargement of heart | **5.65E-05** | **9.82E-05** |
| Binding of professional phagocytic cells | **7.92E-04** | **9.82E-05** |

| | | |
|---|---|---|
| Inflammation of absolute anatomical region | **7.71E-07** | **1.02E-04** |
| Liver Damage | **2.05E-04** | **1.10E-04** |
| Thrombus | **8.83E-04** | **1.10E-04** |
| Inflammation of body cavity | **4.29E-07** | **1.12E-04** |
| Abnormality of large intestine | **7.97E-06** | **1.19E-04** |
| Advanced extracranial solid tumor | **2.53E-04** | **1.19E-04** |
| Lung adenocarcinoma | **1.43E-05** | **1.20E-04** |
| Adhesion of monocytes | **1.19E-04** | **1.25E-04** |
| Large intestine adenocarcinoma | **1.51E-08** | **1.27E-04** |
| Function of macrophages | **1.02E-04** | **1.27E-04** |
| Adhesion of phagocytes | **5.83E-04** | **1.27E-04** |
| Size of vertebrae | **8.32E-04** | **1.28E-04** |
| Transmigration of dendritic cells | **8.32E-04** | **1.28E-04** |
| Peripheral T-cell lymphoma | **1.06E-03** | **1.29E-04** |
| Ulcer of lower extremity | **1.34E-03** | **1.30E-04** |
| Dermatitis | **2.64E-04** | **1.35E-04** |
| Prostate cancer | **6.19E-05** | **1.37E-04** |
| Size of lesion | **7.53E-05** | **1.45E-04** |
| Acute colitis | **1.47E-04** | **1.45E-04** |
| Breast carcinoma | **2.38E-04** | **1.46E-04** |
| Limb development | **1.23E-03** | **1.46E-04** |
| HER2 negative hormone receptor negative breast cancer | **7.36E-04** | **1.53E-04** |
| Neovascularization | **1.93E-05** | **1.56E-04** |
| Head and neck tumor | **4.10E-07** | **1.59E-04** |
| Aortic valve disease | **3.88E-04** | **1.63E-04** |
| Female genital tract adenocarcinoma | **5.58E-07** | **1.67E-04** |
| Neovascularization of eye | **5.33E-05** | **1.73E-04** |
| Neutrophilia | **1.92E-04** | **1.77E-04** |
| Stimulation of hematopoietic progenitor cells | **1.21E-03** | **1.84E-04** |
| Breast or ovarian carcinoma | **5.90E-05** | **1.85E-04** |
| Neoplasia of cells | **6.51E-10** | **1.92E-04** |
| Head and neck cancer | **7.71E-07** | **1.92E-04** |
| Mature lymphocytic neoplasm | **6.53E-04** | **1.92E-04** |
| Clearance of amodiaquine | **7.92E-04** | **1.92E-04** |
| Growth of brain metastasis | **7.92E-04** | **1.92E-04** |
| Release of docosahexaenoic acid | **7.92E-04** | **1.92E-04** |
| Abnormal morphology of body cavity | **8.67E-07** | **1.93E-04** |
| Adhesion of myeloid cells | **9.57E-04** | **1.97E-04** |
| Familial autoimmune disease | **1.02E-03** | **2.08E-04** |
| Valvulopathy | **1.02E-03** | **2.08E-04** |
| Activation of connective tissue cells | **1.04E-03** | **2.12E-04** |
| Pancreatobiliary tumor | **7.45E-06** | **2.27E-04** |
| Activation of antigen presenting cells | **2.54E-04** | **2.32E-04** |
| Quantity of 12-hydroxyeicosatetraenoic acid | **2.92E-06** | **2.55E-04** |

| | | |
|---|---|---|
| Skin tumor | **3.82E-07** | **2.62E-04** |
| Synthesis of eicosanoid | **5.85E-04** | **2.71E-04** |
| Inflammation of gastrointestinal tract | **3.00E-05** | **2.71E-04** |
| Morphology of cardiovascular system | **1.25E-04** | **2.74E-04** |
| Cancer of cells | **2.60E-09** | **2.77E-04** |
| Carcinoma | **3.41E-09** | **2.77E-04** |
| Ovarian cancer | **2.26E-04** | **2.79E-04** |
| Lipolysis | **6.33E-04** | **2.84E-04** |
| Morphology of heart | **1.93E-04** | **2.85E-04** |
| Hereditary connective tissue disorder | **2.84E-04** | **3.00E-04** |
| Blood pressure | **3.58E-04** | **3.05E-04** |
| Liver lesion | **2.26E-04** | **3.24E-04** |
| Hydroxylation of melatonin | **1.34E-03** | **3.24E-04** |
| Induction of prostaglandin E2 | **1.34E-03** | **3.24E-04** |
| Metabolism of amodiaquine | **1.34E-03** | **3.24E-04** |
| Metabolism of flutamide | **1.34E-03** | **3.24E-04** |
| Recovery of neutrophils | **1.34E-03** | **3.24E-04** |
| Genitourinary tumor | **3.06E-10** | 1.00E+00 |
| Female genital neoplasm | **6.28E-10** | 1.00E+00 |
| Pelvic tumor | **1.87E-09** | 1.00E+00 |
| Nonhematologic malignant neoplasm | **1.92E-09** | 1.00E+00 |
| Morbidity or mortality | **2.36E-09** | 1.00E+00 |
| Endometrium tumor | **2.44E-08** | 1.00E+00 |
| Soft tissue neoplasm | **6.59E-08** | 1.00E+00 |
| Development of malignant tumor | **6.81E-08** | 1.00E+00 |
| Squamous-cell carcinoma | **8.75E-08** | 1.00E+00 |
| Melanoma | **1.36E-07** | 1.00E+00 |
| Malignant soft tissue neoplasm | **1.84E-07** | 1.00E+00 |
| Extrapulmonary squamous cell carcinoma | **2.20E-07** | 1.00E+00 |
| Respiratory system tumor | **2.91E-07** | 1.00E+00 |
| Head and neck squamous cell carcinoma | **2.93E-07** | 1.00E+00 |
| Multiple cancers | **3.64E-07** | 1.00E+00 |
| Cutaneous melanoma | **5.65E-07** | 1.00E+00 |
| Uterine carcinoma | **6.77E-07** | 1.00E+00 |
| Mammary tumor | **8.58E-07** | 1.00E+00 |
| Skin cancer | **8.62E-07** | 1.00E+00 |
| Development of carcinoma | **1.06E-06** | 1.00E+00 |
| Non-colon gastrointestinal cancer | **1.55E-06** | 1.00E+00 |
| Head and neck carcinoma | **2.46E-06** | 1.00E+00 |
| Lung tumor | **2.80E-06** | 1.00E+00 |
| Cellular infiltration | **3.95E-06** | 1.00E+00 |
| Development of adenocarcinoma | **5.55E-06** | 1.00E+00 |
| Upper gastrointestinal tract tumor | **5.92E-06** | 1.00E+00 |
| Endometrial carcinoma | **6.83E-06** | 1.00E+00 |
| Decidualization | **8.94E-06** | 1.00E+00 |

| | | |
|---|---|---|
| Proliferation of osteoblasts | **1.05E-05** | 1.00E+00 |
| Upper gastrointestinal carcinoma | **1.09E-05** | 1.00E+00 |
| Remodeling of bone | **1.12E-05** | 1.00E+00 |
| Size of bone | **1.14E-05** | 1.00E+00 |
| Upper gastrointestinal tract cancer | **1.19E-05** | 1.00E+00 |
| Vascularization of eye | **1.19E-05** | 1.00E+00 |
| Prostatic tumor | **1.31E-05** | 1.00E+00 |
| Decidualization of uterus | **1.33E-05** | 1.00E+00 |
| Damage of bone | **1.37E-05** | 1.00E+00 |
| Organization of extracellular matrix | **1.37E-05** | 1.00E+00 |
| Central nervous system cancer | **1.38E-05** | 1.00E+00 |
| Central nervous system solid tumor | **1.43E-05** | 1.00E+00 |
| Cell movement of stem cells | **1.56E-05** | 1.00E+00 |
| Advanced stage peripheral arterial disease | **1.99E-05** | 1.00E+00 |
| Lung carcinoma | **2.10E-05** | 1.00E+00 |
| Pancreatic tumor | **2.34E-05** | 1.00E+00 |
| Subcutaneous tumor | **2.68E-05** | 1.00E+00 |
| Hypersensitive reaction | **2.76E-05** | 1.00E+00 |
| Abnormal morphology of abdomen | **2.85E-05** | 1.00E+00 |
| Liquid tumor | **3.10E-05** | 1.00E+00 |
| Enteritis | **3.17E-05** | 1.00E+00 |
| Advanced malignant tumor | **3.38E-05** | 1.00E+00 |
| Morphology of digestive system | **3.77E-05** | 1.00E+00 |
| Modification of retinaldehyde | **3.81E-05** | 1.00E+00 |
| Size of body | **4.93E-05** | 1.00E+00 |
| Growth of organism | **5.04E-05** | 1.00E+00 |
| Astrocytoma | **5.30E-05** | 1.00E+00 |
| Migration of stem cells | **5.31E-05** | 1.00E+00 |
| Inflammation of the large intestine | **6.63E-05** | 1.00E+00 |
| Invasion of carcinoma cell lines | **7.31E-05** | 1.00E+00 |
| Brain tumor | **7.56E-05** | 1.00E+00 |
| Oxidation of retinaldehyde | **7.66E-05** | 1.00E+00 |
| Glioma | **8.15E-05** | 1.00E+00 |
| Glioma cancer | **8.50E-05** | 1.00E+00 |
| Benign neoplasm of female genital organ | **8.54E-05** | 1.00E+00 |
| Endometrial adenocarcinoma | **8.62E-05** | 1.00E+00 |
| Quantity of bone | **9.81E-05** | 1.00E+00 |
| Advanced lung cancer | **1.04E-04** | 1.00E+00 |
| Pancreatic cancer | **1.09E-04** | 1.00E+00 |
| Progression of malignant tumor | **1.11E-04** | 1.00E+00 |
| Development of peritoneum | **1.13E-04** | 1.00E+00 |
| Uterine endometrioid carcinoma | **1.32E-04** | 1.00E+00 |
| Brain astrocytoma | **1.36E-04** | 1.00E+00 |
| Synthesis of fatty acid | **1.44E-04** | 1.00E+00 |
| Grade 1-4 astrocytoma | **1.47E-04** | 1.00E+00 |

| | | |
|---|---|---|
| Development of digestive organ tumor | **1.59E-04** | 1.00E+00 |
| Recruitment of antigen presenting cells | **1.64E-04** | 1.00E+00 |
| Endometrioid carcinoma | **1.68E-04** | 1.00E+00 |
| Colitis | **1.75E-04** | 1.00E+00 |
| Morphology of muscle cells | **1.80E-04** | 1.00E+00 |
| Abnormal bone density | **1.86E-04** | 1.00E+00 |
| High grade astrocytoma | **2.03E-04** | 1.00E+00 |
| Development of digestive system | **2.03E-04** | 1.00E+00 |
| Synthesis of prostanoid | **2.12E-04** | 1.00E+00 |
| Brain glioma | **2.26E-04** | 1.00E+00 |
| Peripheral arterial disease | **2.26E-04** | 1.00E+00 |
| Progressive malignant solid tumor | **2.26E-04** | 1.00E+00 |
| Grade 3-4 glioma cancer | **2.30E-04** | 1.00E+00 |
| Stage IIIB-IV non-small cell lung cancer | **2.30E-04** | 1.00E+00 |
| Endocrine gland tumor | **2.30E-04** | 1.00E+00 |
| Endometrioid endometrial adenocarcinoma | **2.38E-04** | 1.00E+00 |
| Abnormal morphology of heart | **2.38E-04** | 1.00E+00 |
| Quantity of connective tissue | **2.38E-04** | 1.00E+00 |
| Leukopoiesis | **2.39E-04** | 1.00E+00 |
| Development of body axis | **2.39E-04** | 1.00E+00 |
| Hyperalgesia | **2.40E-04** | 1.00E+00 |
| Metabolism of retinol | **2.47E-04** | 1.00E+00 |
| Activation of phagocytes | **2.47E-04** | 1.00E+00 |
| Migration of glioma cells | **2.47E-04** | 1.00E+00 |
| Grade 4 astrocytoma | **2.49E-04** | 1.00E+00 |
| Grade 4 high grade glioma | **2.49E-04** | 1.00E+00 |
| Grade 4 malignant glioma | **2.49E-04** | 1.00E+00 |
| Accumulation of cells | **2.49E-04** | 1.00E+00 |
| Idiopathic fibrosis | **2.58E-04** | 1.00E+00 |
| Abnormal morphology of cardiovascular system | **2.72E-04** | 1.00E+00 |
| Colorectal tumor | **2.74E-04** | 1.00E+00 |
| Abnormal morphology of digestive system | **2.79E-04** | 1.00E+00 |
| Function of dendritic cells | **2.80E-04** | 1.00E+00 |
| Abnormality of knee joint | **2.95E-04** | 1.00E+00 |
| Development of head | **3.06E-04** | 1.00E+00 |
| Leukemia | **3.22E-04** | 1.00E+00 |
| Concentration of eicosanoid | **3.23E-04** | 1.00E+00 |
| EGFR expressing lung cancer | **3.23E-04** | 1.00E+00 |
| Morphology of skeleton | **3.23E-04** | 1.00E+00 |
| EGFR mutation negative lung cancer | **3.32E-04** | 1.00E+00 |
| Uterine leiomyoma | **3.32E-04** | 1.00E+00 |
| Development of B-cell follicle | **3.47E-04** | 1.00E+00 |
| Release of linoleic acid | **3.47E-04** | 1.00E+00 |
| Stimulation of colony-forming units granulocytes | **3.47E-04** | 1.00E+00 |
| Release of superoxide | **3.47E-04** | 1.00E+00 |

| | | |
|---|---|---|
| Growth of embryonic tissue | **3.47E-04** | 1.00E+00 |
| Synthesis of thromboxane | **3.47E-04** | 1.00E+00 |
| Benign connective or soft tissue neoplasm | **3.50E-04** | 1.00E+00 |
| Quantity of metal | **3.50E-04** | 1.00E+00 |
| Oral tumor | **3.54E-04** | 1.00E+00 |
| Infection by SARS coronavirus | **3.81E-04** | 1.00E+00 |
| Smooth muscle tumor | **3.86E-04** | 1.00E+00 |
| Rheumatoid arthritis | **3.86E-04** | 1.00E+00 |
| Quantity of metal ion | **3.90E-04** | 1.00E+00 |
| Malignant neoplasm of aerodigestive tract | **3.95E-04** | 1.00E+00 |
| Differentiation of tumor cell lines | **3.98E-04** | 1.00E+00 |
| Sensory disorders | **3.98E-04** | 1.00E+00 |
| Fibrosis of lung | **3.99E-04** | 1.00E+00 |
| Inflammation of genital organ | **4.04E-04** | 1.00E+00 |
| Growth of embryo | **4.14E-04** | 1.00E+00 |
| Oral squamous cell carcinoma | **4.23E-04** | 1.00E+00 |
| Autosomal dominant skeletal dysplasia | **4.29E-04** | 1.00E+00 |
| Release of reactive oxygen species | **4.29E-04** | 1.00E+00 |
| Progression of carcinoma | **4.30E-04** | 1.00E+00 |
| Metabolism of vitamin A | **4.50E-04** | 1.00E+00 |
| Clearance of cells | **4.50E-04** | 1.00E+00 |
| Colon tumor | **4.63E-04** | 1.00E+00 |
| Endocrine carcinoma | **4.87E-04** | 1.00E+00 |
| Morphology of heart cells | **4.94E-04** | 1.00E+00 |
| Connective tissue tumor | **4.96E-04** | 1.00E+00 |
| Quantity of antigen presenting cells | **5.01E-04** | 1.00E+00 |
| Recruitment of macrophages | **5.14E-04** | 1.00E+00 |
| Response of myeloid cells | **5.15E-04** | 1.00E+00 |
| Muscle tumor | **5.26E-04** | 1.00E+00 |
| Dysgenesis | **5.67E-04** | 1.00E+00 |
| Recurrent non-small cell lung cancer | **5.69E-04** | 1.00E+00 |
| Expansion of embryonic tissue | **5.89E-04** | 1.00E+00 |
| Acute inflammation of tissue | **5.89E-04** | 1.00E+00 |
| Stage IIIB-IV advanced non-small cell lung cancer | **5.93E-04** | 1.00E+00 |
| Upper aerodigestive tract carcinoma | **6.05E-04** | 1.00E+00 |
| Fatty acid metabolism | **6.05E-04** | 1.00E+00 |
| Abnormal morphology of sclera | **6.37E-04** | 1.00E+00 |
| Differentiation of leukemia cell lines | **6.37E-04** | 1.00E+00 |
| Colorectal cancer | **6.41E-04** | 1.00E+00 |
| Hepatotoxicity | **6.80E-04** | 1.00E+00 |
| Stomach tumor | **7.01E-04** | 1.00E+00 |
| Quantity of carbohydrate | **7.25E-04** | 1.00E+00 |
| Inflammation of testis | **7.47E-04** | 1.00E+00 |
| Movement of vascular endothelial cells | **7.55E-04** | 1.00E+00 |
| Liver Regeneration | **7.66E-04** | 1.00E+00 |

| | | |
|---|---|---|
| Synthesis of polysaccharide | **7.66E-04** | 1.00E+00 |
| Secretion of prostaglandin | **7.80E-04** | 1.00E+00 |
| Recurrent carcinoma | **7.88E-04** | 1.00E+00 |
| Metabolism of polysaccharide | **7.88E-04** | 1.00E+00 |
| Adhesion of rhabdomyosarcoma cell lines | **7.92E-04** | 1.00E+00 |
| Cell polarity formation by epithelial cells | **7.92E-04** | 1.00E+00 |
| Delay in synthesis of lipid | **7.92E-04** | 1.00E+00 |
| Development of corneal stroma | **7.92E-04** | 1.00E+00 |
| Quantity of peripheral lymph node | **7.92E-04** | 1.00E+00 |
| Nonpituitary endocrine tumor | **7.92E-04** | 1.00E+00 |
| Response of lymphatic system cells | **8.07E-04** | 1.00E+00 |
| Metabolism of arachidonic acid | **8.16E-04** | 1.00E+00 |
| Quantity of protein in blood | **8.25E-04** | 1.00E+00 |
| Colorectal adenocarcinoma | **8.30E-04** | 1.00E+00 |
| Metastatic carcinoma | **8.32E-04** | 1.00E+00 |
| Relapsed malignant solid tumor | **8.32E-04** | 1.00E+00 |
| Proliferation of epithelial cells | **8.32E-04** | 1.00E+00 |
| Limb defect | **8.48E-04** | 1.00E+00 |
| Skin carcinoma | **8.48E-04** | 1.00E+00 |
| Proliferation of stem cells | **8.64E-04** | 1.00E+00 |
| Growth failure or short stature | **8.73E-04** | 1.00E+00 |
| Differentiation of phagocytes | **8.76E-04** | 1.00E+00 |
| Cancer of tumor cell lines | **8.79E-04** | 1.00E+00 |
| Synthesis of leukotriene | **8.98E-04** | 1.00E+00 |
| Activation of liver cells | **8.98E-04** | 1.00E+00 |
| Colony formation | **8.98E-04** | 1.00E+00 |
| Synthesis of thromboxane B2 | **9.47E-04** | 1.00E+00 |
| Pancreatic carcinoma | **9.57E-04** | 1.00E+00 |
| Growth of solid tumor | **9.77E-04** | 1.00E+00 |
| Inflammation of secretory structure | **9.78E-04** | 1.00E+00 |
| Upper aero-digestive squamous cell carcinoma | **1.01E-03** | 1.00E+00 |
| Activation of neurons | **1.02E-03** | 1.00E+00 |
| Chronic kidney disease | **1.03E-03** | 1.00E+00 |
| Recurrent cancer | **1.08E-03** | 1.00E+00 |
| Sarcoma | **1.11E-03** | 1.00E+00 |
| Aplasia or hypoplasia | **1.11E-03** | 1.00E+00 |
| Hand deformity | **1.12E-03** | 1.00E+00 |
| Metastatic non-small cell lung carcinoma | **1.14E-03** | 1.00E+00 |
| Morphology of cardiac muscle | **1.14E-03** | 1.00E+00 |
| Binding of lymphatic system cells | **1.15E-03** | 1.00E+00 |
| Morphology of intestine | **1.17E-03** | 1.00E+00 |
| Proliferation of myoblasts | **1.22E-03** | 1.00E+00 |
| Morphology of head | **1.25E-03** | 1.00E+00 |
| Intermediate disease stage peripheral arterial disease | **1.26E-03** | 1.00E+00 |

| | | |
|---|---|---|
| Recurrent malignant solid tumor | **1.30E-03** | 1.00E+00 |
| Stage III cancer | **1.30E-03** | 1.00E+00 |
| Release of lipid | **1.32E-03** | 1.00E+00 |
| Acne | **1.34E-03** | 1.00E+00 |
| Hypermaturation of dendritic cells | **1.34E-03** | 1.00E+00 |
| Induction of astrocytes | **1.34E-03** | 1.00E+00 |
| Synthesis of 6-keto-prostaglandin F1 alpha | **1.34E-03** | 1.00E+00 |
| Neck neoplasm | **1.35E-03** | 1.00E+00 |
| Metabolism of xenobiotic | **1.36E-03** | 1.00E+00 |
| Morphology of tumor | **1.36E-03** | 1.00E+00 |
| Bleeding | **1.36E-03** | 1.00E+00 |
| Ulcer | **1.37E-03** | 1.00E+00 |
| Cell movement of muscle cells | 1.00E+00 | **1.57E-07** |
| Homing of cells | 1.00E+00 | **3.70E-07** |
| Migration of smooth muscle cells | 1.00E+00 | **4.03E-07** |
| Lymphoma | 1.00E+00 | **5.38E-07** |
| Occlusion of blood vessel | 1.00E+00 | **6.35E-07** |
| Cell movement of connective tissue cells | 1.00E+00 | **6.63E-07** |
| Transmigration of cells | 1.00E+00 | **9.64E-07** |
| Migration of connective tissue cells | 1.00E+00 | **9.71E-07** |
| Glucose metabolism disorder | 1.00E+00 | **1.34E-06** |
| Lymphatic system tumor | 1.00E+00 | **1.52E-06** |
| Severe COVID-19 | 1.00E+00 | **2.07E-06** |
| Occlusion of artery | 1.00E+00 | **2.38E-06** |
| Proliferation of smooth muscle cells | 1.00E+00 | **2.83E-06** |
| Migration of vascular smooth muscle cells | 1.00E+00 | **2.89E-06** |
| Diabetic complication | 1.00E+00 | **3.65E-06** |
| Enlargement of cells | 1.00E+00 | **3.80E-06** |
| Binding of colorectal cancer cell lines | 1.00E+00 | **3.95E-06** |
| Formation of vessel | 1.00E+00 | **4.53E-06** |
| Homing of leukocytes | 1.00E+00 | **4.53E-06** |
| Alcoholic hepatitis | 1.00E+00 | **5.31E-06** |
| Cellular infiltration by granulocytes | 1.00E+00 | **5.33E-06** |
| Cell movement of antigen presenting cells | 1.00E+00 | **6.42E-06** |
| Recruitment of mononuclear leukocytes | 1.00E+00 | **7.08E-06** |
| Alzheimer disease | 1.00E+00 | **7.38E-06** |
| Polarization of blood cells | 1.00E+00 | **7.38E-06** |
| Infiltration by neutrophils | 1.00E+00 | **8.72E-06** |
| Accumulation of myeloid-derived suppressor cells | 1.00E+00 | **9.54E-06** |
| Invasion of myeloid cells | 1.00E+00 | **9.54E-06** |
| Collagen-induced arthritis | 1.00E+00 | **9.57E-06** |
| Migration of dermal cells | 1.00E+00 | **1.08E-05** |
| Orientation of cells | 1.00E+00 | **1.08E-05** |
| Interaction of T lymphocytes | 1.00E+00 | **1.10E-05** |
| Cell proliferation of fibroblasts | 1.00E+00 | **1.17E-05** |

| | | |
|---|---|---|
| Tumorigenesis of lymphocytes | 1.00E+00 | **1.19E-05** |
| Stimulation of hepatoma cell lines | 1.00E+00 | **1.31E-05** |
| Uterine serous carcinoma | 1.00E+00 | **1.32E-05** |
| Barrett syndrome | 1.00E+00 | **1.46E-05** |
| Steroidogenesis of hormone | 1.00E+00 | **1.52E-05** |
| Atherosclerosis | 1.00E+00 | **1.52E-05** |
| Transmigration of leukocytes | 1.00E+00 | **1.52E-05** |
| Recruitment of lymphocytes | 1.00E+00 | **1.56E-05** |
| Metabolism of hydrogen peroxide | 1.00E+00 | **1.67E-05** |
| Progressive neurological disorder | 1.00E+00 | **1.71E-05** |
| Acute myocardial infarction | 1.00E+00 | **1.86E-05** |
| Vascular lesion | 1.00E+00 | **1.88E-05** |
| Infection of mammalia | 1.00E+00 | **1.90E-05** |
| Non-Hodgkin lymphoma | 1.00E+00 | **2.13E-05** |
| Metabolism of hormone | 1.00E+00 | **2.13E-05** |
| Binding of lymphocytes | 1.00E+00 | **2.13E-05** |
| Accumulation of granulocytes | 1.00E+00 | **2.27E-05** |
| Activation of osteoclasts | 1.00E+00 | **2.39E-05** |
| Respiratory distress | 1.00E+00 | **2.55E-05** |
| Invasion of leukocytes | 1.00E+00 | **2.63E-05** |
| Gonadal tumor | 1.00E+00 | **2.71E-05** |
| Proliferation of vascular cells | 1.00E+00 | **2.81E-05** |
| Mature T-cell or NK-cell neoplasm | 1.00E+00 | **2.82E-05** |
| Cell movement of macrophages | 1.00E+00 | **2.94E-05** |
| Priming of leukocytes | 1.00E+00 | **2.98E-05** |
| Cell movement of embryonic cell lines | 1.00E+00 | **3.27E-05** |
| Cellular infiltration by phagocytes | 1.00E+00 | **3.46E-05** |
| Accumulation of myeloid cells | 1.00E+00 | **3.71E-05** |
| Formation of vascular lesion | 1.00E+00 | **3.79E-05** |
| Quantity of lymphoid cells | 1.00E+00 | **4.10E-05** |
| Ischemia of heart | 1.00E+00 | **4.32E-05** |
| Colonization | 1.00E+00 | **4.32E-05** |
| Localization of blood cells | 1.00E+00 | **4.37E-05** |
| Preterm birth | 1.00E+00 | **4.37E-05** |
| Progressive encephalopathy | 1.00E+00 | **4.37E-05** |
| Size of heart | 1.00E+00 | **4.74E-05** |
| Morphology of muscle | 1.00E+00 | **4.77E-05** |
| Uterine serous papillary cancer | 1.00E+00 | **4.87E-05** |
| Mobilization of cells | 1.00E+00 | **5.03E-05** |
| Cell movement of fibroblasts | 1.00E+00 | **5.20E-05** |
| Proliferation of kidney cells | 1.00E+00 | **5.29E-05** |
| Recruitment of T lymphocytes | 1.00E+00 | **5.29E-05** |
| Accumulation of neutrophils | 1.00E+00 | **5.52E-05** |
| Large intestine carcinoma | 1.00E+00 | **5.60E-05** |
| Binding of T lymphocytes | 1.00E+00 | **5.82E-05** |

| | | |
|---|---|---|
| Stimulation of cancer cells | 1.00E+00 | **5.86E-05** |
| Synthesis of cartilage matrix | 1.00E+00 | **5.86E-05** |
| Polarization of leukocytes | 1.00E+00 | **5.86E-05** |
| Attachment of cells | 1.00E+00 | **5.86E-05** |
| Vasodilation of aorta | 1.00E+00 | **6.07E-05** |
| Angiogenesis of lesion | 1.00E+00 | **6.17E-05** |
| Metabolism of reactive oxygen species | 1.00E+00 | **6.24E-05** |
| Inflammation of respiratory system component | 1.00E+00 | **6.55E-05** |
| Migration of endothelial cells | 1.00E+00 | **6.74E-05** |
| Binding of Stat3 binding site | 1.00E+00 | **7.02E-05** |
| Dengue shock syndrome | 1.00E+00 | **7.02E-05** |
| Formation of neovessel | 1.00E+00 | **7.02E-05** |
| Cell proliferation of tumor cell lines | 1.00E+00 | **7.02E-05** |
| Interstitial fibrosis | 1.00E+00 | **7.07E-05** |
| Asthma | 1.00E+00 | **7.38E-05** |
| Diabetic retinopathy | 1.00E+00 | **7.40E-05** |
| Invasion of malignant tumor | 1.00E+00 | **7.42E-05** |
| Steroid metabolism | 1.00E+00 | **7.44E-05** |
| Preeclampsia | 1.00E+00 | **7.50E-05** |
| Migration of keratinocytes | 1.00E+00 | **7.52E-05** |
| T-cell malignant neoplasm | 1.00E+00 | **8.23E-05** |
| Biosynthesis of hydrogen peroxide | 1.00E+00 | **8.23E-05** |
| Binding of SAA activating factor binding site | 1.00E+00 | **8.52E-05** |
| Induction of glucocorticoid | 1.00E+00 | **8.52E-05** |
| Proliferation of Tc17 cells | 1.00E+00 | **8.52E-05** |
| Release of azurophil granules | 1.00E+00 | **8.52E-05** |
| Stimulation of fibroblast-like synoviocytes | 1.00E+00 | **8.52E-05** |
| Migration of colorectal cancer cell lines | 1.00E+00 | **8.52E-05** |
| Demyelination | 1.00E+00 | **8.74E-05** |
| Induction of prostaglandin | 1.00E+00 | **9.82E-05** |
| G0 phase of tumor cell lines | 1.00E+00 | **1.00E-04** |
| G0/G1 phase transition of tumor cell lines | 1.00E+00 | **1.01E-04** |
| Infarction | 1.00E+00 | **1.01E-04** |
| Activation of blood platelets | 1.00E+00 | **1.02E-04** |
| Mature T-cell neoplasm | 1.00E+00 | **1.03E-04** |
| Cell tethering or rolling of leukocytes | 1.00E+00 | **1.07E-04** |
| Injury of mice | 1.00E+00 | **1.07E-04** |
| Renal lesion | 1.00E+00 | **1.07E-04** |
| Fibrosis of liver | 1.00E+00 | **1.08E-04** |
| Accumulation of phagocytes | 1.00E+00 | **1.09E-04** |
| Cell movement of epithelial cell lines | 1.00E+00 | **1.09E-04** |
| Calcinosis | 1.00E+00 | **1.09E-04** |
| T-cell non-Hodgkin lymphoma | 1.00E+00 | **1.09E-04** |
| Damage of cartilage tissue | 1.00E+00 | **1.09E-04** |
| Activation of enzyme | 1.00E+00 | **1.10E-04** |

| | | |
|---|---|---|
| Hypercytokinemia | 1.00E+00 | **1.12E-04** |
| Priming of granulocytes | 1.00E+00 | **1.12E-04** |
| Inflammation of connective tissue | 1.00E+00 | **1.12E-04** |
| Migration of phagocytes | 1.00E+00 | **1.13E-04** |
| Non-melanoma solid tumor | 1.00E+00 | **1.16E-04** |
| Chronic periodontal disease | 1.00E+00 | **1.19E-04** |
| Systemic lupus erythematosus | 1.00E+00 | **1.19E-04** |
| Adhesion of connective tissue cells | 1.00E+00 | **1.25E-04** |
| Mineralization of bone | 1.00E+00 | **1.29E-04** |
| Acute respiratory disorder | 1.00E+00 | **1.29E-04** |
| Function of liver | 1.00E+00 | **1.30E-04** |
| Morphology of malignant tumor | 1.00E+00 | **1.30E-04** |
| Infection by RNA virus | 1.00E+00 | **1.30E-04** |
| Binding of myeloid cells | 1.00E+00 | **1.33E-04** |
| Formation of filopodia | 1.00E+00 | **1.34E-04** |
| Priming of T lymphocytes | 1.00E+00 | **1.37E-04** |
| Cell rolling of lymphocytes | 1.00E+00 | **1.45E-04** |
| Chronic respiratory disorder | 1.00E+00 | **1.45E-04** |
| Recovery of blood cells | 1.00E+00 | **1.45E-04** |
| Quantity of lymphocytes | 1.00E+00 | **1.45E-04** |
| Apoptosis of lung cell lines | 1.00E+00 | **1.45E-04** |
| Formation of actin filaments | 1.00E+00 | **1.46E-04** |
| Chemotaxis of antigen presenting cells | 1.00E+00 | **1.47E-04** |
| Function of kidney | 1.00E+00 | **1.49E-04** |
| Stenosis of aortic valve | 1.00E+00 | **1.52E-04** |
| Cell viability of myeloid cells | 1.00E+00 | **1.53E-04** |
| Apoptosis of granulocytes | 1.00E+00 | **1.53E-04** |
| Development of bone marrow | 1.00E+00 | **1.55E-04** |
| Inflammation of mucosa | 1.00E+00 | **1.59E-04** |
| Formation of lymphoid tissue | 1.00E+00 | **1.66E-04** |
| Proliferation of mesangial cells | 1.00E+00 | **1.67E-04** |
| Sjögren syndrome | 1.00E+00 | **1.67E-04** |
| Myocardial infarction | 1.00E+00 | **1.76E-04** |
| Synthesis of reactive oxygen species | 1.00E+00 | **1.76E-04** |
| Acute respiratory distress syndrome | 1.00E+00 | **1.77E-04** |
| Invasion of cancer cells | 1.00E+00 | **1.77E-04** |
| Cell death of phagocytes | 1.00E+00 | **1.80E-04** |
| Polarization of CD4+ T-lymphocytes | 1.00E+00 | **1.84E-04** |
| Fibrosis of peritoneum | 1.00E+00 | **1.92E-04** |
| Gap junctional intercellular communication of hepatocytes | 1.00E+00 | **1.92E-04** |
| Inhibition of endothelial cells | 1.00E+00 | **1.92E-04** |
| Primary pilocytic astrocytoma | 1.00E+00 | **1.92E-04** |
| Stimulation of acute myeloblastic leukemia cells | 1.00E+00 | **1.92E-04** |
| Transmigration of phagocytes | 1.00E+00 | **1.92E-04** |

| | | |
|---|---|---|
| Resorption of bone | 1.00E+00 | **1.94E-04** |
| Growth of hematological system tumor | 1.00E+00 | **1.95E-04** |
| Immune response of cells | 1.00E+00 | **1.99E-04** |
| Osteoclastogenesis | 1.00E+00 | **2.01E-04** |
| Stimulation of cells | 1.00E+00 | **2.03E-04** |
| Invasion of phagocytes | 1.00E+00 | **2.03E-04** |
| Formation of actin stress fibers | 1.00E+00 | **2.04E-04** |
| Weight gain | 1.00E+00 | **2.07E-04** |
| Mesenchymal tumor | 1.00E+00 | **2.08E-04** |
| Cell viability of cancer cells | 1.00E+00 | **2.11E-04** |
| Growth of lymphoid tissue | 1.00E+00 | **2.16E-04** |
| Apoptosis of bone marrow cell lines | 1.00E+00 | **2.19E-04** |
| Severe pulmonary disease | 1.00E+00 | **2.20E-04** |
| Binding of tumor cell lines | 1.00E+00 | **2.25E-04** |
| Quantity of reactive oxygen species | 1.00E+00 | **2.36E-04** |
| Cell movement of lymphocytes | 1.00E+00 | **2.36E-04** |
| Histiocytosis | 1.00E+00 | **2.43E-04** |
| Cell death of myeloid cells | 1.00E+00 | **2.46E-04** |
| Proliferation of vascular smooth muscle cells | 1.00E+00 | **2.48E-04** |
| Fibrosis of kidney | 1.00E+00 | **2.57E-04** |
| Lymphocytic leukemia | 1.00E+00 | **2.60E-04** |
| Abnormality of cartilage tissue | 1.00E+00 | **2.62E-04** |
| Fever | 1.00E+00 | **2.65E-04** |
| Cell death of immune cells | 1.00E+00 | **2.66E-04** |
| Body temperature | 1.00E+00 | **2.71E-04** |
| Necrosis of liver | 1.00E+00 | **2.71E-04** |
| Morphology of vasculature | 1.00E+00 | **2.74E-04** |
| Cell death of tumor cells | 1.00E+00 | **2.74E-04** |
| Thrombosis of vein | 1.00E+00 | **2.74E-04** |
| Formation of reactive oxygen species | 1.00E+00 | **2.75E-04** |
| Formation of hydrogen peroxide | 1.00E+00 | **2.77E-04** |
| Inflammatory response of respiratory system component | 1.00E+00 | **2.77E-04** |
| Migration of antigen presenting cells | 1.00E+00 | **2.77E-04** |
| Size of infarct | 1.00E+00 | **2.80E-04** |
| Thermoregulation | 1.00E+00 | **2.80E-04** |
| Cell death of antigen presenting cells | 1.00E+00 | **2.80E-04** |
| Activation of Phospholipase C | 1.00E+00 | **2.83E-04** |
| Inflammation of lung | 1.00E+00 | **2.84E-04** |
| Congenital anomaly of skin | 1.00E+00 | **2.84E-04** |
| Severe inflammatory disorder | 1.00E+00 | **2.86E-04** |
| Invasion of epithelial tissue | 1.00E+00 | **2.94E-04** |
| Quantity of bacteria | 1.00E+00 | **2.94E-04** |
| Cell movement of T lymphocytes | 1.00E+00 | **3.03E-04** |
| Morphology of spleen | 1.00E+00 | **3.05E-04** |

| | | |
|---|---|---|
| Permeability of vascular system | 1.00E+00 | **3.05E-04** |
| Proliferation of leukemia cells | 1.00E+00 | **3.21E-04** |
| Decreased secretion of albumin | 1.00E+00 | **3.24E-04** |
| Polarization of T-cell hybrid cells | 1.00E+00 | **3.24E-04** |
| Quantity of polymorphonuclear granulocytic MDSC cells | 1.00E+00 | **3.24E-04** |
| Sensitization of macrophages | 1.00E+00 | **3.24E-04** |
| Shedding of CD34+ progenitor-derived megakaryocyte | 1.00E+00 | **3.24E-04** |
| Adhesion of lymphocytes | 1.00E+00 | **3.24E-04** |

**P*-value: Benjamini Hochberg adjusted *P*-value. Bold *P*-values: < 0.05. n = 6 and 5 female and male HUVECs preparations, respectively.

Table S12. ITE-dysregulated disease-associated biological function in HUVECs.*

| | *P*-Value [Female HUVECs] | *P*-Value [Male HUVECs] |
|---|---|---|
| Hypertrophy of heart | **1.63E-04** | **3.98E-05** |
| Enlargement of heart | **1.84E-04** | **1.70E-04** |
| Liver Damage | **5.78E-04** | **1.70E-04** |
| Liver Regeneration | **2.07E-03** | **6.88E-04** |
| Hepatotoxicity | **2.07E-03** | **9.70E-03** |
| Chronic renal failure | **2.52E-03** | **5.32E-05** |
| Fibrosis of liver | **3.05E-03** | **1.70E-04** |
| Stenosis of aortic valve | **3.05E-03** | **2.31E-04** |
| Inflammation of liver | **3.05E-03** | **4.30E-04** |
| Primary biliary cirrhosis | **3.05E-03** | **6.56E-03** |
| Necrosis of liver | **4.15E-03** | **3.42E-04** |
| Acute myocardial infarction | **4.51E-03** | **3.98E-05** |
| Calcification of aortic valve | **4.51E-03** | **6.44E-03** |
| Drug-induced hepatotoxicity | **4.51E-03** | **2.08E-02** |
| Fibrosis of kidney | **4.78E-03** | **3.42E-04** |
| Damage of kidney | **4.78E-03** | **7.61E-04** |
| Hypertrophy of heart cells | **4.78E-03** | **1.28E-03** |
| Liver tumor | **5.05E-03** | **2.92E-03** |
| Activation of hepatic stellate cells | **5.30E-03** | **4.02E-03** |
| Injury of kidney | **5.81E-03** | **1.61E-03** |
| Myocardial infarction | **6.43E-03** | **2.31E-04** |
| Acute hepatitis | **6.43E-03** | **9.15E-03** |
| Generation of reactive oxygen species in liver | **6.43E-03** | **3.15E-02** |
| Apoptosis of cardiomyocytes | **6.85E-03** | **2.01E-03** |
| Proliferation of liver cells | **7.85E-03** | **2.12E-03** |
| Alcoholic hepatitis | **8.03E-03** | **2.02E-05** |
| End stage renal disease | **8.03E-03** | **7.61E-04** |
| Progression of hepatocellular carcinoma | **8.03E-03** | **2.12E-03** |
| Non-rheumatic aortic stenosis | **8.03E-03** | **2.12E-03** |
| Acute ST-elevation myocardial infarction | **8.03E-03** | **2.12E-03** |
| Cirrhosis | **8.03E-03** | **5.65E-03** |
| Nephrotoxicity | **8.03E-03** | **6.36E-03** |
| Increased activation of alkaline phosphatase | **8.96E-03** | **1.28E-03** |
| Acute liver failure | **1.01E-02** | **2.56E-03** |
| Damage of renal tubule | **1.05E-02** | **1.43E-02** |
| Acute kidney injury | **1.08E-02** | **7.48E-03** |
| Fibrosis of heart | **1.17E-02** | **7.61E-04** |
| Liver cancer | **1.53E-02** | **3.13E-03** |
| Proliferation of hepatic stellate cells | **1.56E-02** | **2.56E-03** |
| Hypertrophy of cardiac muscle | **1.66E-02** | **1.52E-03** |
| ST-elevation myocardial infarction | **1.71E-02** | **3.05E-03** |

| | | |
|---|---|---|
| Cardiac valvular form autosomal recessive Ehlers-Danlos syndrome | **1.71E-02** | **6.44E-03** |
| High grade chronic hepatitis C | **1.71E-02** | **6.44E-03** |
| Inflammation of renal interstitium | **1.71E-02** | **6.44E-03** |
| Development of liver tumor | **1.96E-02** | **2.56E-03** |
| Failure of liver | **1.96E-02** | **3.56E-03** |
| Liver carcinoma | **1.96E-02** | **4.02E-03** |
| Proliferation of kidney cells | **2.21E-02** | **1.02E-04** |
| Left ventricular dysfunction | **2.21E-02** | **2.63E-03** |
| Chronic hepatitis | **2.21E-02** | **3.46E-03** |
| Ventricular hypertrophy | **2.21E-02** | **3.66E-03** |
| Autoimmune glomerulonephritis | **2.21E-02** | **4.31E-03** |
| Apoptosis of mesangial cells | **2.21E-02** | **5.98E-03** |
| Pulmonary embolism | **2.21E-02** | **6.44E-03** |
| Acetaminophen overdose | **2.21E-02** | **9.15E-03** |
| Drug toxicity-related nephropathy | **2.21E-02** | **9.15E-03** |
| Low grade chronic hepatitis C | **2.21E-02** | **9.15E-03** |
| Progressive hepatic fibrosis | **2.21E-02** | **9.15E-03** |
| Rupture of heart ventricle | **2.21E-02** | **9.15E-03** |
| Increased release of albumin | **2.21E-02** | **9.15E-03** |
| Bleeding of liver parenchyma | **2.21E-02** | **9.15E-03** |
| Experimentally-induced heart failure | **2.21E-02** | **9.15E-03** |
| Nephritis | **2.21E-02** | **1.21E-02** |
| Glomerulonephritis | **2.21E-02** | **2.08E-02** |
| Proximal tubular toxicity | **2.21E-02** | **2.81E-02** |
| Cell death of kidney cells | **2.21E-02** | **4.53E-02** |
| Increased Levels of Hematocrit | **2.22E-02** | **5.19E-03** |
| Hepatocellular carcinoma | **2.23E-02** | **1.63E-03** |
| Hypertrophy of cardiomyocytes | **2.32E-02** | **3.94E-03** |
| Hepatomegaly | **2.32E-02** | **3.17E-02** |
| Proliferation of hepatocytes | **2.92E-02** | **4.69E-03** |
| Right ventricular hypertrophy | **2.92E-02** | **9.05E-03** |
| Multi vessel coronary artery disease | **2.92E-02** | **1.30E-02** |
| Apoptosis of liver cell lines | **2.97E-02** | **9.15E-03** |
| Development of hepatocellular carcinoma | **3.22E-02** | **6.86E-03** |
| Benign neoplasm of liver | **3.22E-02** | **9.15E-03** |
| Focal glomerulonephritis | **3.22E-02** | **9.15E-03** |
| Decreased secretion of albumin | **3.40E-02** | **4.14E-04** |
| Hypoplasia of renal medulla | **3.40E-02** | **1.55E-02** |
| Obstruction of bile duct | **3.40E-02** | **1.55E-02** |
| Inflammation of renal glomerulus | **3.40E-02** | **1.55E-02** |
| Fatal myocardial infarction | **3.40E-02** | **1.55E-02** |
| Increased Levels of ALT | **3.79E-02** | **1.52E-03** |
| Apoptosis of kidney cells | **3.80E-02** | **8.78E-03** |
| Apoptosis of oval cells | **4.12E-02** | **1.82E-02** |

| | | |
|---|---|---|
| Syndromic congenital heart disease | **4.71E-02** | **1.43E-02** |
| Acute myocarditis | **4.71E-02** | **2.08E-02** |
| Spontaneous coronary artery dissection | **4.71E-02** | **2.08E-02** |
| Proliferation of mesangial cells | 5.29E-02 | **2.31E-05** |
| Regeneration of kidney | 5.29E-02 | **2.41E-02** |
| Interstitial fibrosis of heart | 5.39E-02 | **1.55E-02** |
| Failure of heart | 5.79E-02 | **6.86E-03** |
| Increased Levels of LDH | 5.79E-02 | **1.67E-02** |
| Hyperplasia of kidney | 5.79E-02 | **2.68E-02** |
| Hypertrophy of renal tubule | 5.79E-02 | **2.68E-02** |
| Calcific aortic valve stenosis | 6.39E-02 | **2.90E-02** |
| Formation of hepatocellular adenoma | 6.39E-02 | **2.90E-02** |
| Interstitial fibrosis of kidney | 6.69E-02 | **3.00E-03** |
| Recurrent myocardial infarction | 6.71E-02 | **3.15E-02** |
| Survival of podocytes | 6.71E-02 | **3.15E-02** |
| Disorder of coronary artery | 7.11E-02 | **8.98E-03** |
| Decreased hematocrit of organism | 7.20E-02 | **3.39E-02** |
| Cortical renal glomerulopathies | 7.20E-02 | **3.39E-02** |
| Increased release of ALT | 7.44E-02 | **2.03E-03** |
| Atrophy of renal cortex | 7.44E-02 | **3.64E-02** |
| Lupus nephritis WHO class III/IV | 7.44E-02 | **3.64E-02** |
| Atrial fibrillation | 7.53E-02 | **1.67E-02** |
| Thrombosis of atrium | 7.82E-02 | **3.88E-02** |
| Fibrosis of renal glomerulus | 7.82E-02 | **3.88E-02** |
| Pyelonephritis | 8.29E-02 | **2.41E-03** |
| Nonfatal myocardial infarction | 8.80E-02 | **4.39E-02** |
| Viral hepatitis | 9.15E-02 | **5.19E-03** |
| Shprintzen syndrome | 9.15E-02 | **4.58E-02** |
| Regeneration of liver cells | 9.15E-02 | **4.58E-02** |
| Apoptosis of liver cells | 1.04E-01 | **1.52E-03** |
| Hypertrophy of left ventricle | 1.12E-01 | **3.88E-02** |
| Arrhythmia | 1.15E-01 | **2.08E-02** |
| Congenital heart disease | 1.16E-01 | **6.44E-03** |
| Cholestasis | 1.19E-01 | **4.39E-02** |
| Severe hepatitis | 1.30E-01 | **4.30E-04** |
| Proliferation of cardiomyocytes | 1.32E-01 | **1.78E-03** |
| Increased localization of ALT | 1.54E-01 | **7.43E-03** |
| Coronary artery disease | 1.58E-01 | **1.03E-02** |
| Congestive heart failure | 1.74E-01 | **1.27E-02** |
| Familial congenital heart disease | 2.04E-01 | **1.55E-02** |
| Apoptosis of hepatocytes | 2.18E-01 | **4.15E-03** |
| Left ventricular dilation | 2.84E-01 | **2.05E-02** |
| Cell proliferation of kidney cell lines | 2.87E-01 | **2.70E-02** |
| Hepatic steatosis | 4.15E-01 | **1.47E-02** |
| Alcoholic steatohepatitis | 1.00E+00 | **1.61E-03** |

| | | |
|---|---|---|
| Chronic hepatitis C | 1.00E+00 | **4.26E-03** |
| Severe alcoholic hepatitis | 1.00E+00 | **5.65E-03** |
| Inflammation of liver lobule | 1.00E+00 | **6.44E-03** |
| Opisthorchiasis periductal fibrosis | 1.00E+00 | **6.44E-03** |
| Autosomal dominant supravalvular aortic stenosis | 1.00E+00 | **6.44E-03** |
| Steatohepatitis | 1.00E+00 | **6.50E-03** |
| Cirrhosis of liver | 1.00E+00 | **8.94E-03** |
| Biliary fibrosis | 1.00E+00 | **9.15E-03** |
| Decreased production of albumin | 1.00E+00 | **9.15E-03** |
| Hypertrophy of ventricular myocardium | 1.00E+00 | **9.15E-03** |
| Increased synthesis of CRP | 1.00E+00 | **9.15E-03** |
| Initiation of regeneration of liver | 1.00E+00 | **9.15E-03** |
| Enlargement of left ventricle | 1.00E+00 | **1.46E-02** |
| Susceptibility to atrioventricular septal defect type 1 | 1.00E+00 | **1.55E-02** |
| Hepatitis subtype infantile hepatitis syndrome | 1.00E+00 | **1.55E-02** |
| Infantile cholestatic hepatitis syndrome | 1.00E+00 | **1.55E-02** |
| Damage of liver tissue | 1.00E+00 | **1.55E-02** |
| Stenosis of pulmonary artery | 1.00E+00 | **1.82E-02** |
| Decreased synthesis of albumin | 1.00E+00 | **1.82E-02** |
| Infection by hepatitis B virus | 1.00E+00 | **2.67E-02** |
| Chemotaxis of hepatic stellate cells | 1.00E+00 | **2.90E-02** |
| Glomerulosclerosis | **4.78E-03** | 1.00E+00 |
| Bleeding of liver | **4.78E-03** | 1.00E+00 |
| Hypertrophy of renal glomerulus | **6.43E-03** | 9.92E-02 |
| Adult idiopathic cirrhosis | **1.71E-02** | 1.00E+00 |
| Autosomal recessive progressive familial intrahepatic cholestasis type 3 | **1.71E-02** | 1.00E+00 |
| Stage III primary biliary cirrhosis | **1.71E-02** | 1.00E+00 |
| Stage IV primary biliary cirrhosis | **1.71E-02** | 1.00E+00 |
| Polycystic liver disease 4 with or without kidney cysts | **1.71E-02** | 1.00E+00 |
| Familial pulmonary hypertension | **2.21E-02** | 1.00E+00 |
| Chronic cholestasis | **2.21E-02** | 1.00E+00 |
| Drug-induced cholestasis | **2.21E-02** | 1.00E+00 |
| Granulomatous hepatitis | **2.21E-02** | 1.00E+00 |
| Intrahepatic cholestasis of pregnancy type 3 | **2.21E-02** | 1.00E+00 |
| Severe hepatic fibrosis | **2.21E-02** | 1.00E+00 |
| Transient idiopathic neonatal cholestasis | **2.21E-02** | 1.00E+00 |
| Acute liver injury | **2.21E-02** | 1.00E+00 |
| Goodpasture syndrome | **2.92E-02** | 1.00E+00 |
| Inflammation of liver portal space | **2.92E-02** | 1.00E+00 |
| Membranous glomerulonephritis | **3.22E-02** | 1.01E-01 |
| Idiopathic and/or familial pulmonary arterial hypertension | **3.22E-02** | 1.00E+00 |
| Liver Hypoplasia | **3.40E-02** | 1.07E-01 |
| Persistent fetal circulation syndrome | **3.40E-02** | 1.00E+00 |
| Bile duct proliferation | **4.71E-02** | 1.00E+00 |

| | | |
|---|---|---|
| Polycystic liver disease 1 | **4.71E-02** | 1.00E+00 |

*$P$-value: Benjamini Hochberg adjusted $P$-value. Bold $P$-values: < 0.05. n = 6 and 5 female and male HUVECs preparations, respectively.

Table 13. ITE-dysregulated canonical pathways in HUVECs.*

| | *P*-value [Female HUVECs] | *P*-value [Male HUVECs] |
|---|---|---|
| Xenobiotic Metabolism AHR Signaling Pathway | **1.58E-03** | **4.77E-06** |
| Atherosclerosis Signaling | **1.31E-04** | **2.55E-05** |
| Aryl Hydrocarbon Receptor Signaling | **2.71E-03** | **6.57E-05** |
| Differential Regulation of Cytokine Production in Macrophages and T Helper Cells by IL-17A and IL-17F | **4.24E-02** | **2.01E-04** |
| Breast Cancer Regulation by Stathmin1 | **5.56E-03** | **1.92E-02** |
| CREB Signaling in Neurons | **6.16E-03** | **1.93E-02** |
| Hepatic Fibrosis Signaling Pathway | **1.03E-02** | **2.04E-02** |
| Phagosome Formation | **5.29E-03** | **2.98E-02** |
| Nicotine Degradation III | **4.22E-02** | **4.62E-02** |
| Apelin Liver Signaling Pathway | **5.29E-03** | 1.47E-01 |
| GP6 Signaling Pathway | **4.22E-02** | 3.42E-01 |
| Role of Hypercytokinemia/hyperchemokinemia in the Pathogenesis of Influenza | 4.72E-01 | **6.36E-05** |
| Xenobiotic Metabolism Signaling | 7.65E-02 | **9.03E-04** |
| Role of IL-17F in Allergic Inflammatory Airway Diseases | 4.01E-01 | **2.47E-03** |
| Role of Cytokines in Mediating Communication between Immune Cells | 1.31E-01 | **3.65E-03** |
| HIF1α Signaling | 2.68E-01 | **1.50E-02** |
| LXR/RXR Activation | 3.40E-01 | **2.25E-02** |
| Role of Macrophages, Fibroblasts and Endothelial Cells in Rheumatoid Arthritis | 9.90E-02 | **3.77E-02** |
| Acute Phase Response Signaling | 4.13E-01 | **4.59E-02** |
| Role of MAPK Signaling in Inhibiting the Pathogenesis of Influenza | 5.65E-02 | **6.30E-04** |
| HGF Signaling | 1.20E-01 | **3.65E-03** |
| HMGB1 Signaling | 6.04E-02 | **7.07E-03** |
| IL-17 Signaling | 7.95E-02 | **1.06E-02** |
| Differential Regulation of Cytokine Production in Intestinal Epithelial Cells by IL-17A and IL-17F | 5.53E-02 | **1.41E-02** |
| Bupropion Degradation | 6.08E-02 | **1.75E-02** |
| Osteoarthritis Pathway | 5.53E-02 | **1.92E-02** |
| Glucocorticoid Receptor Signaling | 2.10E-01 | **1.92E-02** |
| Cholecystokinin/Gastrin-mediated Signaling | 1.08E-01 | **2.04E-02** |
| IL-17A Signaling in Fibroblasts | 3.80E-01 | **2.30E-02** |
| Acetone Degradation I (to Methylglyoxal) | 9.87E-02 | **2.34E-02** |
| HOTAIR Regulatory Pathway | 6.04E-02 | **3.74E-02** |
| Neuroinflammation Signaling Pathway | 9.90E-02 | **3.77E-02** |
| Estrogen Biosynthesis | 1.32E-01 | **3.84E-02** |
| Regulation Of The Epithelial Mesenchymal Transition By Growth Factors Pathway | 4.18E-01 | **4.67E-02** |
| Activation of IRF by Cytosolic Pattern Recognition Receptors | 1.00E+00 | **4.67E-02** |

*$P$-value: Benjamini Hochberg adjusted $P$-value. Bold $P$-value: < 0.05. n = 6 and 5 female and male HUVECs preparations.

Table S14. ITE-dysregulated endothelial function-associated gene networks in HUVECs.*

| | Female HUVECs | | Male HUVECs | |
|---|---|---|---|---|
| Gene network | *P*-value | Target Genes (n) | *P*-value | Target Genes (n) |
| AHR-regulated genes | 8.46E-08 | ACP5, AHRR, CCR7, COL1A2, COL3A1, CYP1A1, CYP1B1, IDO1, IL1B, NPTX1, PTGS2, SERPINB2, TIPARP (13) | 1.15E-12 | AHRR, CCR7, COL14A1, COL1A2, CXCL10, CYP1A1, CYP1B1, IL1B, IL6, NPTX1, NQO1, PTGS2, SERPINB2, TIPARP (14) |
| CREB1-regulated genes | 4.41E-03 | APOLD1, GAD1, IL1B, JUNB, MMP1, PLA2G4A, PTGS2, TIPARP (8) | 2.24E-03 | IL1B, IL6, MMP1, PTGS2, STC1, TIPARP (6) |
| IRF8-regulated genes | 3.38E-04 | AHRR, CCR7, IDO1, IL1B, LIF (5) | 1.52E-05 | AHRR, CCR7, IL1B, IL6, ISG15 (5) |
| JUN-regulated genes | 5.10E-08 | ACP5, CCL8, COL1A2, CSF3, CYP1B1, F3, IL1B, JUNB, LIF, MMP1, PLA2G4A, PTGS2, SERPINB2, XYLT1 (14) | 1.45E-09 | COL1A2, CSF3, CXCL10, CYP1B1, F3, IL1B, IL6, LPAR1, MMP1, NQO1, PTGS2, SERPINB2 (12) |
| RELA-regulated genes | 7.22E-06 | AHRR, CCR7, COL1A2, F3, IL1B, JUNB, LUM, LYN, MMP1, PTGS2, SELE (11) | 6.03E-10 | AHRR, CCR7, COL1A2, CXCL10, ELN, F3, IL1B, IL6, ISG15, MMP1, PTGS2, SELE (12) |
| TP63-regulated genes | 3.27E-04 | F3, FOXF1, IL1B, JUNB, LYN, MAFF, POSTN, SERPINB2, SLC7A11 (9) | 9.12E-04 | F3, IL1B, IL6, POSTN, SERPINB2, SLC7A11 (6) |
| ELK1-regulated genes | 1.59E-05 | JUNB, NUPR1, PTGS2, SLC7A11, TIPARP (5) | 6.00E-04 | PTGS2, SLC7A11, TIPARP (3) |
| EZH2-regulated genes | 6.45E-05 | CEMIP, COL1A2, CSF3, CYP1B1, GPR68, NCOA7, PLA2G4A, PTGS2, SLC7A11 (9) | 3.29E-06 | CLEC3B, COL1A2, CSF3, CXCL10, CYP1B1, IL6, PTGS2, SLC7A11 (8) |
| FOS-regulated genes | 2.78E-04 | CSF3, F3, IL1B, JUNB, MMP1, NPTX1, PTGS2, SERPINB2, TIPARP, XYLT1 (10) | 8.62E-08 | CSF3, CXCL10, F3, IL1B, IL6, MMP1, NPTX1, NQO1, PTGS2, SERPINB2, TIPARP (11) |
| FOXL2-regulated genes | 4.67E-04 | LIF, MAFF, PTGS2, SERPINB2 (4) | 8.85E-04 | IL6, PTGS2, SERPINB2 (3) |
| FOXO1-regulated genes | 5.03E-03 | ABCB4, CCR7, GPX3, IL1B, JUNB, MMP1, SERPINB2 (7) | 1.04E-04 | CCR7, CXCL10, GPX3, IL1B, IL6, MMP1, SERPINB2 (7) |
| HIF1A-regulated genes | 6.93E-03 | CCR7, CEMIP, CYP1A1, IL1B, MAFF, MMP1, PTGS2 (7) | 1.90E-05 | CCR7, CXCL10, CYP1A1, IL1B, IL6, MMP1, PLAC8, PTGS2 (8) |

| | | | | |
|---|---|---|---|---|
| MAFB-regulated genes | 6.41E-04 | CMKLR1, F2RL3, HTR2B, SERPINB2 (4) | 1.13E-03 | F2RL3, HTR2B, SERPINB2 (3) |
| STAT3-regulated genes | 1.76E-07 | ALDH1A1, COL1A2, COL3A1, CSF3, CXCR5, F3, HGF, IL1B, JUNB, LEPR, LGR4, LIF, MMP1, PLA2G4A, PTGS2 (15) | 2.36E-07 | COL1A2, CSF3, CXCL10, F3, HGF, IL1B, IL6, ISG15, MMP1, PTGS2, RSAD2 (11) |
| HMGB1-regulated genes | 2.80E-04 | IL1B, MMP1, PTGS2, SELE (4) | 6.40E-07 | IL1B, IL6, MMP1, PTGS2, SELE (5) |
| MYC-regulated genes | | N/A | 1.83E-03 | AQP1, CLEC3B, COL14A1, COL1A2, CXCL10, F3, NQO1, PTGS2, RSAD2 (9) |
| NKX2-3-regulated genes | 6.37E-05 | F2RL3, GALNT15, IDO1, MAMDC2, POSTN, PTGS2, TIPARP (7) | 1.81E-03 | F2RL3, POSTN, PTGS2, TIPARP (4) |
| CBX5-regulated genes | 1.87E-03 | CEMIP, CYP1B1, TACSTD2, TNFRSF6B (4) | | N/A |
| RBPJ-regulated genes | 7.75E-05 | CSF3, GDF5, HGF, IL1B, LFNG, LIF (6) | 3.38E-05 | CSF3, HGF, IL1B, IL6, LFNG (5) |
| ZFP36-regulated genes | 4.83E-05 | AHRR, COL3A1, IL1B, MMP1, PTGS2 (5) | 5.48E-05 | AHRR, IL1B, MMP1, PTGS2 (4) |

**P*-value: Benjamini Hochberg adjusted *P*-value; Z Scores > |2| are predicted to be target genes; n = number of target genes. N/A: prediction not available.

Table S15. ITE-dysregulated aryl hydrocarbon receptor signaling pathway genes.*

| Gene ID | Gene Name | Cellular Location | Female HUVECs | | Male HUVECs | | Female Rat Placentas | | Male Rat Placentas | |
|---|---|---|---|---|---|---|---|---|---|---|
| | | | Log2 (fold change [ITE/Ctrl]) | *P* -value | Log2 (fold change [ITE/Ctrl]) | *P* -value | Log2 (fold change [ITE/Ctrl]) | *P* -value | Log2 (fold change [ITE/Ctrl]) | *P* -value |
| AHRR | aryl-hydrocarbon receptor repressor | Nucleus | 4.051 | 5.18E-29 | 3.503 | 2.72E-28 | 2.077 | 1.03E-80 | 1.984 | 2.56E-23 |
| ALDH1A1 | aldehyde dehydrogenase 1 family member A1 | Cytoplasm | -1.337 | 7.17E-02 | -0.142 | 1.00E+00 | 1.211 | 6.61E-67 | 0.966 | 1.02E-17 |
| ALDH1A2 | aldehyde dehydrogenase 1 family member A2 | Cytoplasm | -1.375 | 4.78E-02 | -0.969 | 6.48E-02 | -0.373 | 7.06E-05 | -0.141 | 1.00E+00 |
| ALDH1A3 | aldehyde dehydrogenase 1 family member A3 | Cytoplasm | 1.033 | 1.32E-02 | 1.063 | 1.18E-03 | 0.717 | 1.66E-13 | 0.399 | 2.44E-01 |
| ALDH5A1 | aldehyde dehydrogenase 5 family member A1 | Cytoplasm | -0.863 | 3.74E-03 | -1.327 | 2.06E-08 | 0.284 | 9.05E-01 | 0.165 | 1.00E+00 |
| ALDH8A1 | aldehyde dehydrogenase 8 family member A1 | Cytoplasm | -1.270 | 1.82E-01 | -2.239 | 1.33E-04 | -1.058 | 1.00E+00 | 1.612 | 1.00E+00 |
| CCNA2 | cyclin A2 | Nucleus | -1.523 | 3.17E-07 | -1.564 | 3.07E-10 | -0.624 | 1.20E-14 | -0.275 | 9.47E-01 |
| CCND1 | cyclin D1 | Nucleus | -1.010 | 6.20E-09 | -0.944 | 2.99E-09 | 0.030 | 1.00E+00 | 0.157 | 1.00E+00 |
| CCND2 | cyclin D2 | Nucleus | -1.783 | 1.48E-09 | -1.516 | 6.63E-08 | 0.175 | 4.58E-01 | 0.235 | 1.00E+00 |
| CDKN2A | cyclin dependent kinase inhibitor 2A | Nucleus | -1.258 | 1.59E-01 | 1.199 | 9.12E-03 | -0.268 | 1.00E+00 | 0.423 | 1.00E+00 |
| CYP1A1 | cytochrome P450 family 1 subfamily A member 1 | Cytoplasm | 5.255 | 4.75E-14 | 4.709 | 5.98E-14 | 9.554 | 0.00E+00 | 9.618 | 7.98E-21 |
| CYP1B1 | cytochrome P450 family 1 subfamily B member 1 | Cytoplasm | 4.343 | 2.85E-35 | 3.952 | 2.36E-36 | 7.733 | 0.00E+00 | 6.247 | 6.74E-21 |
| Cyp2c70 | cytochrome P450, family 2, subfamily c, polypeptide 70 | Cytoplasm | -1.001 | 4.16E-01 | -1.688 | 6.85E-03 | | | | |
| CYP2C9 | cytochrome P450 family 2 subfamily C member 9 | Cytoplasm | -2.218 | 2.40E-02 | -2.370 | 7.53E-03 | 1.196 | 1.00E+00 | 1.607 | 1.00E+00 |
| FASLG | Fas ligand | Extracellular Space | 1.323 | 1.23E-07 | 1.398 | 7.32E-11 | -1.344 | 1.00E+00 | 1.728 | 1.00E+00 |
| GSTA5 | glutathione S-transferase alpha 5 | Cytoplasm | 2.294 | 2.19E-05 | 2.403 | 1.72E-02 | 0.000 | 1.00E+00 | 0.000 | 1.00E+00 |
| GSTM3 | glutathione S-transferase mu 3 | Cytoplasm | 0.000 | 1.00E+00 | -5.018 | 2.48E-03 | 0.220 | 1.75E-01 | 0.000 | 1.00E+00 |
| IL1B | interleukin 1 beta | Extracellular Space | 0.648 | 8.27E-02 | 1.375 | 3.01E-08 | 1.878 | 2.51E-06 | 2.371 | 2.83E-08 |
| IL6 | interleukin 6 | Extracellular Space | 1.621 | 8.95E-02 | 3.471 | 4.72E-07 | 0.907 | 1.00E-17 | 1.525 | 3.16E-07 |
| NCOA7 | nuclear receptor coactivator 7 | Nucleus | 0.108 | 7.68E-01 | -0.062 | 1.00E+00 | 1.159 | 8.17E-59 | 0.880 | 7.46E-15 |
| NQO1 | NAD(P)H quinone dehydrogenase 1 | Cytoplasm | 1.089 | 2.68E-15 | 1.399 | 3.11E-26 | 0.906 | 3.54E-37 | 1.068 | 2.56E-20 |
| TFF1 | trefoil factor 1 | Extracellular Space | 0.196 | 1.00E+00 | 1.807 | 3.80E-02 | 1.389 | 1.00E+00 | 0.000 | 1.00E+00 |

*Differentially expressed genes with Log2(fold change) >1 or <-1 and Benjamini Hochberg adjusted *P* -value <0.05 are considered significant; Ctrl: vehicle control (DMSO).

Table S16. ITE-dysregulated differential regulation of cytokine production in macrophages and T helper cells by IL-17A and IL-17F pathway genes.*

| Gene ID | Gene Name | Cellular Location | Female HUVECs | | Male HUVECs | | Female Rat Placentas | | Male Rat Placentas | |
|---|---|---|---|---|---|---|---|---|---|---|
| | | | Log2 (fold change [ITE/Ctrl]) | *P* -value | Log2 (fold change [ITE/Ctrl]) | *P* -value | Log2 (fold change [ITE/Ctrl]) | *P* -value | Log2 (fold change [ITE/Ctrl]) | *P* -value |
| CCL2 | C-C motif chemokine ligand 2 | Extracellular Space | 1.975 | 2.65E-05 | 3.414 | 8.32E-15 | 0.462 | 3.58E-09 | 0.649 | 1.96E-02 |
| CCL4 | C-C motif chemokine ligand 4 | Extracellular Space | 1.141 | 8.13E-02 | 2.619 | 6.17E-09 | -0.036 | 1.00E+00 | -0.148 | 1.00E+00 |
| CCL5 | C-C motif chemokine ligand 5 | Extracellular Space | -1.071 | 1.68E-01 | 1.053 | 3.35E-02 | 0.105 | 1.00E+00 | -0.33 | 1.00E+00 |
| CSF3 | colony stimulating factor 3 | Extracellular Space | 0.352 | 9.24E-01 | 0.513 | 5.80E-01 | 1.197 | 1.05E-17 | 1.061 | 1.16E-07 |
| IL1B | interleukin 1 beta | Extracellular Space | 0.648 | 8.27E-02 | 1.375 | 3.01E-08 | 1.878 | 2.51E-06 | 2.371 | 2.83E-08 |
| IL6 | interleukin 6 | Extracellular Space | 1.621 | 8.95E-02 | 3.471 | 4.72E-07 | 0.907 | 1.00E-17 | 1.525 | 3.16E-07 |

*Differentially expressed genes with Log2(fold change) >1 or <-1 and Benjamini Hochberg adjusted P-value <0.05 are considered significant; Ctrl: vehicle control (DMSO).

Table S17. ITE-dysregulated IL17 signalling pathway genes.*

| Gene ID | Gene Name | Cellular Location | Female HUVECs | | Male HUVECs | | Female Rat Placentas | | Male Rat Placentas | |
|---|---|---|---|---|---|---|---|---|---|---|
| | | | Log2 (fold change [ITE/Ctrl]) | *P*-value | Log2 (fold change [ITE/Ctrl]) | *P*-value | Log2 (fold change [ITE/Ctrl]) | *P*-value | Log2 (fold change [ITE/Ctrl]) | *P*-value |
| CSF3 | colony stimulating factor 3 | Extracellular Space | 0.352 | 9.24E-01 | 0.513 | 5.80E-01 | 1.197 | 1.05E-17 | 1.061 | 1.16E-07 |
| IL1B | interleukin 1 beta | Extracellular Space | 0.648 | 8.27E-02 | 1.375 | 3.01E-08 | 1.878 | 2.51E-06 | 2.371 | 2.83E-08 |
| LIF | LIF interleukin 6 family cytokine | Extracellular Space | 0.273 | 4.77E-01 | -0.073 | 1.00E+00 | 1.206 | 2.31E-02 | 1.703 | 1.00E+00 |
| PTGS2 | prostaglandin-endoperoxide synthase 2 | Cytoplasm | -1.613 | 1.87E-04 | -1.997 | 1.61E-07 | 1.542 | 5.90E-89 | 1.098 | 2.21E-20 |
| IL6 | interleukin 6 | Extracellular Space | 1.621 | 8.95E-02 | 3.471 | 4.72E-07 | 0.907 | 1.00E-17 | 1.525 | 3.16E-07 |
| CCL2 | C-C motif chemokine ligand 2 | Extracellular Space | 1.975 | 2.65E-05 | 3.414 | 8.32E-15 | 0.462 | 3.58E-09 | 0.649 | 1.96E-02 |
| CXCL3 | C-X-C motif chemokine ligand 3 | Extracellular Space | 1.174 | 4.67E-02 | 1.449 | 8.35E-04 | 0.615 | 3.01E-04 | 0.438 | 8.31E-01 |
| Eda | ectodysplasin-A | Other | -1.295 | 2.44E-03 | -1.030 | 2.34E-03 | - | - | - | - |
| FASLG | Fas ligand | Extracellular Space | 1.323 | 1.23E-07 | 1.398 | 7.32E-11 | -1.344 | 1.00E+00 | 1.728 | 1.00E+00 |
| IL36B | interleukin 36 beta | Extracellular Space | 1.715 | 2.61E-04 | 2.300 | 3.79E-09 | -0.503 | 1.00E+00 | 4.033 | 1.00E+00 |
| LEP | leptin | Extracellular Space | -1.661 | 4.92E-02 | -2.376 | 1.68E-04 | 0.907 | 1.00E+00 | -0.520 | 1.00E+00 |
| MMP13 | matrix metallopeptidase 13 | Extracellular Space | 1.328 | 4.54E-04 | 2.080 | 2.53E-11 | -0.695 | 1.00E+00 | 0.000 | 1.00E+00 |
| OSM | oncostatin M | Extracellular Space | 1.103 | 6.41E-05 | 1.431 | 1.31E-10 | 1.196 | 1.00E+00 | -0.788 | 1.00E+00 |
| PDGFC | platelet derived growth factor C | Extracellular Space | -1.229 | 8.83E-07 | -1.105 | 8.45E-06 | -0.335 | 4.43E-02 | -0.278 | 1.00E+00 |
| PIK3C2G | phosphatidylinositol-4-phosphate 3-kinase catalytic subunit type 2 gamma | Cytoplasm | 1.345 | 6.49E-09 | 1.243 | 4.63E-10 | -1.344 | 1.00E+00 | -0.947 | 1.00E+00 |
| TNFRSF11B | TNF receptor superfamily member 11b | Plasma Membrane | -1.304 | 9.51E-05 | -0.956 | 6.55E-04 | -0.426 | 1.00E+00 | 0.243 | 1.00E+00 |
| CCL20 | C-C motif chemokine ligand 20 | Extracellular Space | 1.364 | 1.99E-01 | 2.439 | 1.36E-03 | -0.966 | 2.87E-06 | -0.129 | 1.00E+00 |
| IL17RE | interleukin 17 receptor E | Plasma Membrane | -0.895 | 2.06E-05 | -1.002 | 3.16E-09 | -0.288 | 1.00E+00 | -0.294 | 1.00E+00 |
| MMP9 | matrix metallopeptidase 9 | Extracellular Space | 0.601 | 1.75E-02 | 1.066 | 1.24E-08 | 0.876 | 1.00E+00 | 0.196 | 1.00E+00 |
| PIK3CG | phosphatidylinositol-4,5-bisphosphate 3-kinase catalytic subunit gamma | Cytoplasm | 0.948 | 8.23E-05 | 1.027 | 1.82E-07 | 0.193 | 6.94E-01 | 0.171 | 1.00E+00 |
| PIK3R5 | phosphoinositide-3-kinase regulatory subunit 5 | Cytoplasm | 0.322 | 5.86E-01 | 1.151 | 1.71E-06 | 0.139 | 1.00E+00 | -0.390 | 1.00E+00 |
| RRAS2 | RAS related 2 | Plasma Membrane | -0.994 | 1.18E-08 | -1.029 | 1.35E-08 | -0.233 | 5.49E-02 | -0.003 | 1.00E+00 |
| TNFSF10 | TNF superfamily member 10 | Extracellular Space | 0.422 | 6.13E-01 | 1.045 | 2.48E-03 | -0.037 | 1.00E+00 | 0.037 | 1.00E+00 |
| TNFSF13B | TNF superfamily member 13b | Extracellular Space | 0.241 | 1.00E+00 | 1.076 | 9.00E-03 | 0.408 | 1.00E+00 | 0.410 | 1.00E+00 |
| TNFSF15 | TNF superfamily member 15 | Extracellular Space | -0.306 | 1.00E+00 | -1.492 | 1.09E-04 | -0.129 | 8.08E-01 | 0.048 | 1.00E+00 |

*Differentially expressed genes with Log2(fold change) >1 or <-1 and Benjamini Hochberg adjusted P-value <0.05 are considered significant; Ctrl: vehicle control (DMSO).

Table S18. Comparisons of immune cell gene expression signatures in DMSO-treated NT-HUVECs.*

| Cell subtype (Relative fraction) | T cells CD4 memory resting | Dendritic cells activated | T cells CD4 naive | Mast cells resting | NK cells activated | Macrophages M2 | B cells naive | Monocytes | T cells regulatory | Macrophages M1 | Macrophages M0 | Plasma cells | T cells CD8 | NK cells resting | B cells memory | Eosinophils | Dendritic cells resting | T cells gamma delta | T cells follicular helper | Mast cells activated | T cells CD4 memory activated | Neutrophils |
|---|---|---|---|---|---|---|---|---|---|---|---|---|---|---|---|---|---|---|---|---|---|---|
| T cells CD4 memory resting (0.270) | | | | γ | γ | γ | γ | γ | γ | γ | γ | γ | γ | γ | γ | γ | γ | γ | γ | γ | γ | γ |
| Dendritic cells activated (0.263) | | | | γ | γ | γ | γ | γ | γ | γ | γ | γ | γ | γ | γ | γ | γ | γ | γ | γ | γ | γ |
| T cells CD4 naïve (0.220) | | | | γ | γ | γ | γ | γ | γ | γ | γ | γ | γ | γ | γ | γ | γ | γ | γ | γ | γ | γ |
| Mast cells resting (0.086) | | | | | ε | γ | γ | γ | γ | γ | γ | γ | γ | γ | γ | γ | γ | γ | γ | γ | γ | γ |
| NK cells activated (0.069) | | | | | | γ | γ | γ | γ | γ | γ | γ | γ | γ | γ | γ | ε | γ | ε | γ | γ | γ |
| Macrophages M2 (0.037) | | | | | | | | | γ | γ | γ | γ | γ | γ | γ | γ | γ | γ | γ | γ | γ | γ |
| B cells naïve (0.019) | | | | | | | | | ε | γ | γ | γ | γ | γ | γ | γ | γ | γ | ε | γ | γ | γ |
| Monocytes (0.016) | | | | | | | | | γ | γ | γ | γ | γ | γ | γ | γ | γ | γ | γ | γ | γ | γ |
| T cells regulatory (Tregs) (0.005) | | | | | | | | | | | | | | | | | | | | | | |
| Macrophages M1 (0.005) | | | | | | | | | | | | | | | | | | | | | | |
| Macrophages M0 (0.002) | | | | | | | | | | | | | | | | | | | | | | |
| Plasma cells (0.002) | | | | | | | | | | | | | | | | | | | | | | |
| T cells CD8 (0.001) | | | | | | | | | | | | | | | | | | | | | | |
| NK cells resting (0.001) | | | | | | | | | | | | | | | | | | | | | | |
| B cells memory (0.000) | | | | | | | | | | | | | | | | | | | | | | |
| Eosinophils (0.000) | | | | | | | | | | | | | | | | | | | | | | |
| Dendritic cells resting (0.000) | | | | | | | | | | | | | | | | | | | | | | |
| T cells gamma delta (0.000) | | | | | | | | | | | | | | | | | | | | | | |
| T cells follicular helper (0.000) | | | | | | | | | | | | | | | | | | | | | | |
| Mast cells activated (0.000) | | | | | | | | | | | | | | | | | | | | | | |
| T cells CD4 memory activated (0.000) | | | | | | | | | | | | | | | | | | | | | | |
| Neutrophils (0.000) | | | | | | | | | | | | | | | | | | | | | | |

* The one -way ANOVA with all pairwise multiple comparison was performed using Student-Newman-Keuls Method. $^{\varepsilon}P < 0.02$, $^{\gamma}P < 0.001$ vs corresponding cell type. n = 11 (6 female and 5 male) HUVECs preparations.

Table S19. ITE-induced changes in immune cell gene expression signatures in NT-HUVECs.*

| | Combined Female and Male | | Female | | Male | |
|---|---|---|---|---|---|---|
| | DMSO | ITE | DMSO | ITE | DMSO | ITE |
| B cells naive | 0.019±0.003 | 0.027±0.004 | 0.020±0.005 | 0.024±0.008 | 0.017±0.003 | 0.030±0.007 |
| B cells memory | 0.000±0.000 | 0.000±0.000 | 0.000±0.000 | 0.001±0.000 | 0.000±0.000 | 0.000±0.000 |
| Plasma cells | 0.002±0.001 | 0.001±0.001 | 0.002±0.001 | 0.002±0.001 | 0.002±0.002 | 0.000±0.000 |
| T cells CD8 | 0.001±0.001 | 0.006±0.003 | 0.000±0.000 | 0.009±0.007 | 0.002±0.002 | 0.003±0.002 |
| T cells CD4 naive | 0.220±0.024 | 0.200±0.030 | 0.245±0.033 | 0.188±0.051 | 0.190±0.034 | 0.215±0.006 |
| T cells CD4 memory resting | 0.270±0.026 | 0.274±0.030 | 0.240±0.031 | 0.289±0.054 | 0.306±0.042 | 0.257±0.047 |
| T cells CD4 memory activated | 0.000±0.000 | 0.006±0.004 | 0.000±0.000 | 0.000±0.000 | 0.000±0.000 | 0.013±0.013 |
| T cells follicular helper | 0.000±0.000 | 0.002±0.001 | 0.000±0.000 | 0.001±0.001 | 0.000±0.000 | 0.003±0.003 |
| T cells regulatory (Tregs) | 0.005±0.004 | 0.001±0.003 | 0.001±0.001 | 0.000±0.000 | 0.010±0.010 | 0.002±0.002 |
| T cells gamma delta | 0.000±0.000 | 0.000±0.000 | 0.000±0.000 | 0.000±0.000 | 0.000±0.000 | 0.000±0.000 |
| NK cells resting | 0.001±0.001 | 0.007±0.004 | 0.001±0.001 | 0.005±0.004 | 0.000±0.000 | 0.010±0.010 |
| NK cells activated | 0.069±0.010 | 0.057±0.010 | 0.079±0.016 | 0.067±0.018 | 0.057±0.006 | 0.045±0.005 |
| Monocytes | 0.016±0.002 | 0.025±0.004 | 0.013±0.001 | 0.028±0.009 | 0.018±0.005 | 0.022±0.005 |
| Macrophages M0 | 0.005±0.002 | 0.007±0.003 | 0.002±0.002 | 0.009±0.006 | 0.007±0.005 | 0.005±0.005 |
| Macrophages M1 | 0.001±0.001 | 0.000±0.000 | 0.001±0.001 | 0.000±0.000 | 0.000±0.000 | 0.000±0.000 |
| Macrophages M2 | 0.037±0.009 | 0.025±0.010 | 0.041±0.008 | 0.024±0.011 | 0.033±0.017 | 0.026±0.020 |
| Dendritic cells resting | 0.000±0.000 | 0.004±0.002 | 0.000±0.000 | 0.003±0.003 | 0.000±0.000 | 0.005±0.005 |
| Dendritic cells activated | 0.263±0.027 | 0.272±0.031 | 0.258±0.041 | 0.265±0.055 | 0.270±0.039 | 0.282±0.042 |
| Mast cells resting | 0.086±0.012 | 0.082±0.014 | 0.088±0.008 | 0.086±0.022 | 0.085±0.026 | 0.077±0.023 |
| Mast cells activated | 0.000±0.000 | 0.000±0.000 | 0.000±0.000 | 0.000±0.000 | 0.000±0.000 | 0.000±0.000 |
| Eosinophils | 0.005±0.004 | 0.003±0.003 | 0.008±0.007 | 0.002±0.001 | 0.002±0.001 | 0.004±0.002 |
| Neutrophils | 0.000±0.000 | 0.000±0.000 | 0.000±0.000 | 0.000±0.000 | 0.000±0.000 | 0.000±0.000 |

The two-way ANOVA with all pairwise multiple comparison was performed using Holm-Sidak tests. Data are expressed as means ± SEM relative cell fractions. n =11 (6 female and 5 male) HUVECs preparations.

Table S20. Comparisons of immune cell gene expression signatures in P0 NT-HUVECs.*

| Cell subtype (Relative fraction) | T cells CD4 memory resting | Dendritic cells activated | T cells CD4 naive | NK cells resting | Eosinophils | Mast cells activated | Macrophages M0 | Monocytes | B cells naive | Macrophages M2 | Mast cells resting | Plasma cells | T cells CD4 memory | T cells CD8 | B cells memory | Macrophages M1 | Neutrophils | T cells regulatory (Tregs) | Dendritic cells resting | T cells follicular helper | NK cells activated | T cells gamma delta |
|---|---|---|---|---|---|---|---|---|---|---|---|---|---|---|---|---|---|---|---|---|---|---|
| T cells CD4 memory resting (0.312) | | ε | ε | ε | ε | ε | ε | ε | ε | ε | ε | ε | ε | ε | ε | ε | γ | γ | γ | γ | γ | γ |
| Dendritic cells activated (0.192) | | | ε | ε | ε | ε | ε | ε | ε | ε | ε | ε | ε | ε | ε | ε | γ | γ | γ | γ | γ | γ |
| T cells CD4 naïve (0.135) | | | | ε | ε | ε | ε | ε | ε | ε | ε | ε | ε | ε | ε | ε | ε | ε | γ | γ | γ | γ |
| NK cells resting (0.105) | | | | | ε | ε | ε | ε | ε | ε | ε | ε | ε | ε | ε | ε | ε | ε | ε | γ | γ | γ |
| Eosinophils (0.056) | | | | | | ε | ε | ε | ε | ε | ε | ε | ε | ε | ε | ε | ε | ε | ε | ε | γ | γ |
| Mast cells activated (0.050) | | | | | | | ε | ε | ε | ε | ε | ε | ε | ε | ε | ε | ε | ε | ε | ε | ε | γ |
| Macrophages M0 (0.040) | | | | | | | | ε | ε | ε | ε | ε | ε | ε | ε | ε | ε | ε | ε | ε | ε | ε |
| Monocytes (0.030) | | | | | | | | | ε | ε | ε | ε | ε | ε | ε | ε | ε | ε | ε | ε | ε | ε |
| B cells naïve (0.029) | | | | | | | | | | ε | ε | ε | ε | ε | ε | ε | ε | ε | ε | ε | ε | ε |
| Macrophages M2 (0.020) | | | | | | | | | | | | | | | | | | | | | | |
| Mast cells resting (0.013) | | | | | | | | | | | | | | | | | | | | | | |
| Plasma cells (0.001) | | | | | | | | | | | | | | | | | | | | | | |
| T cells CD4 memory activated (0.007) | | | | | | | | | | | | | | | | | | | | | | |
| T cells CD8 (0.004) | | | | | | | | | | | | | | | | | | | | | | |
| B cells memory (0.002) | | | | | | | | | | | | | | | | | | | | | | |
| Macrophages M1 (0.001) | | | | | | | | | | | | | | | | | | | | | | |
| Neutrophils (0.000) | | | | | | | | | | | | | | | | | | | | | | |
| T cells regulatory (Tregs) (0.000) | | | | | | | | | | | | | | | | | | | | | | |
| Dendritic cells resting (0.000) | | | | | | | | | | | | | | | | | | | | | | |
| T cells follicular helper (0.000) | | | | | | | | | | | | | | | | | | | | | | |
| NK cells activated (0.000) | | | | | | | | | | | | | | | | | | | | | | |
| T cells gamma delta (0.000) | | | | | | | | | | | | | | | | | | | | | | |

*The one -way ANOVA with all pairwise multiple comparison was performed using Student-Newman-Keuls Method. Buck RNAseq data from our previous report[1] were subjected to CIBERSORTx. P0: unpassaged cell preparations. NT: normotensive pregnancy. $^{\varepsilon}P < 0.01$, $^{\gamma}P < 0.001$ vs NT within the same sex group. n = 16 (8 female and 8 male) NT-HUVECs.

1. Zhou C, *et al.* Sexual Dimorphisms of Preeclampsia-Dysregulated Transcriptomic Profiles and Cell Function in Fetal Endothelial Cells. *Hypertension* **74**, 154-163 (2019).

0

Table S21. PE-induced changes in immune cell gene expression signatures in P0 HUVECs.*

| | Combined Female and Male | | Female | | Male | |
|---|---|---|---|---|---|---|
| | NT | PE | NT | PE | NT | PE |
| B cells naive | 0.029±0.009 | 0.023±0.005 | 0.027±0.013 | 0.022±0.002 | 0.038±0.015 | 0.023±0.009 |
| B cells memory | 0.002±0.001 | 0.000±0.000 | 0.002±0.002 | 0.000±0.000 | 0.004±0.003 | 0.000±0.000 |
| Plasma cells | 0.001±0.000 | 0.002±0.000 | 0.001±0.001 | 0.001±0.001 | 0.001±0.001 | 0.003±0.001 |
| T cells CD8 | 0.004±0.002 | 0.003±0.001 | 0.005±0.003 | 0.001±0.001 | 0.005±0.005 | 0.005±0.002 |
| T cells CD4 naive | 0.135±0.029 | 0.115±0.028 | 0.176±0.042 | $0.104±0.032^{\varepsilon}$ | $0.103±0.047^{\delta}$ | 0.124±0.047 |
| T cells CD4 memory resting | 0.312±0.024 | 0.281±0.024 | 0.294±0.043 | 0.267±0.048 | 0.332±0.027 | 0.294±0.024 |
| T cells CD4 memory activated | 0.007±0.004 | 0.009±0.005 | 0.004±0.004 | 0.007±0.007 | 0.005±0.003 | 0.010±0.007 |
| T cells follicular helper | 0.000±0.000 | 0.000±0.000 | 0.000±0.000 | 0.000±0.000 | 0.000±0.000 | 0.000±0.000 |
| T cells regulatory (Tregs) | 0.000±0.000 | 0.000±0.000 | 0.000±0.000 | 0.000±0.000 | 0.001±0.001 | 0.000±0.000 |
| T cells gamma delta | 0.000±0.000 | 0.000±0.000 | 0.000±0.000 | 0.000±0.000 | 0.000±0.000 | 0.000±0.000 |
| NK cells resting | 0.105±0.007 | 0.085±0.011 | 0.120±0.008 | 0.067±0.014 | 0.082±0.008 | 0.099±0.014 |
| NK cells activated | 0.000±0.000 | 0.002±0.001 | 0.000±0.000 | 0.000±0.000 | 0.000±0.000 | 0.000±0.000 |
| Monocytes | 0.030±0.005 | 0.025±0.003 | 0.031±0.005 | 0.023±0.005 | 0.033±0.012 | 0.027±0.005 |
| Macrophages M0 | 0.040±0.009 | 0.029±0.008 | 0.039±0.011 | 0.012±0.007 | 0.052±0.019 | 0.043±0.010 |
| Macrophages M1 | 0.002±0.001 | 0.025±0.019 | 0.002±0.002 | 0.049±0.041 | 0.002±0.002 | 0.004±0.002 |
| Macrophages M2 | 0.020±0.008 | 0.002±0.002 | 0.027±0.011 | 0.000±0.000 | 0.017±0.017 | 0.004±0.004 |
| Dendritic cells resting | 0.000±0.000 | 0.000±0.000 | 0.000±0.000 | 0.000±0.000 | 0.001±0.001 | 0.000±0.000 |
| Dendritic cells activated | 0.192±0.025 | 0.227±0.025 | 0.150±0.038 | $0.261±0.016^{\gamma}$ | $0.209±0.027^{\delta}$ | 0.199±0.042 |
| Mast cells resting | 0.013±0.008 | 0.006±0.005 | 0.008±0.004 | 0.004±0.004 | 0.024±0.022 | 0.008±0.008 |
| Mast cells activated | 0.050±0.010 | 0.104±0.022 | 0.039±0.012 | $0.112±0.035^{\varepsilon}$ | 0.057±0.020 | 0.097±0.031 |
| Eosinophils | 0.056±0.010 | 0.059±0.007 | 0.074±0.017 | 0.059±0.009 | 0.034±0.006 | 0.059±0.012 |
| Neutrophils | 0.000±0.000 | 0.004±0.003 | 0.001±0.001 | 0.007±0.006 | 0.000±0.000 | 0.001±0.001 |

*The Two-way ANOVA with all pairwise multiple comparison was performed using Holm-Sidak tests. Buck RNAseq data from our previous report[1] were subjected to CIBERSORTx. Data are expressed as means ± SEM relative cell fractions. P0: unpassaged cell preparations. NT: normotensive pregnancy; PE: preeclampsia. $^{\varepsilon}P < 0.01$, $^{\gamma}P < 0.001$ vs NT within the same sex group. $^{\delta}P < 0.001$ vs. corresponding female. n = 8, 8, 5, and 6 for NT-female, NT-male, PE-female and PE-male cell preparations, respectively.

Table 22. ITE-dysregulated common genes in rat placentas and HUVECs.*

| Rat Placentas | | | | HUVECs | | | |
|---|---|---|---|---|---|---|---|
| Symbol | Gene ID | Log FC (ITE/Ctrl) | FDR *P*-Value | Symbol | Gene ID | Log FC (ITE/Ctrl) | FDR *P*-Value |
| Cyp1b1 | ENSRNOG00000040287 | 4.19 | 0.0000 | CYP1B1 | ENSG00000138061 | 6.96 | 1.30E-13 |
| Ahrr | ENSRNOG00000014721 | 3.80 | 0.0000 | AHRR | ENSG00000063438 | 2.03 | 2.92E-21 |
| Cyp1a1 | ENSRNOG00000019500 | 4.60 | 0.0000 | CYP1A1 | ENSG00000140465 | 9.73 | 1.06E-65 |
| F2rl3 | ENSRNOG00000048186 | 1.21 | 0.0000 | F2RL3 | ENSG00000127533 | 2.69 | 8.45E-11 |
| Rnd1 | ENSRNOG00000059857 | 1.09 | 0.0000 | RND1 | ENSG00000172602 | 1.51 | 2.37E-40 |
| C1s | ENSRNOG00000011971 | 1.01 | 0.0000 | C1S | ENSG00000182326 | 1.25 | 1.67E-14 |
| Nptx1 | ENSRNOG00000003741 | 1.09 | 0.0000 | NPTX1 | ENSG00000171246 | 4.44 | 6.16E-06 |
| Il1b | ENSRNOG00000004649 | 1.16 | 0.0000 | IL1B | ENSG00000125538 | 2.11 | 7.02E-15 |
| Il6 | ENSRNOG00000010278 | 2.84 | 0.0000 | IL6 | ENSG00000136244 | 1.17 | 8.86E-08 |
| Foxf1 | ENSRNOG00000049906 | -1.72 | 0.0000 | FOXF1 | ENSG00000103241 | 1.02 | 8.04E-06 |
| Ptgs2 | ENSRNOG00000002525 | -2.26 | 0.0000 | PTGS2 | ENSG00000073756 | 1.34 | 1.38E-22 |
| Apold1 | ENSRNOG00000007830 | -1.84 | 0.0000 | APOLD1 | ENSG00000178878 | 1.32 | 6.12E-06 |
| Abcb4 | ENSRNOG00000008012 | -1.28 | 0.0000 | ABCB4 | ENSG00000005471 | 1.33 | 0.004743 |
| Tacstd2 | ENSRNOG00000007740 | -1.39 | 0.0000 | TACSTD2 | ENSG00000184292 | 1.05 | 5.68E-19 |
| Cytl1 | ENSRNOG00000028108 | -2.68 | 0.0000 | CYTL1 | ENSG00000170891 | 1.44 | 1.07E-05 |
| Gria2 | ENSRNOG00000054204 | -1.69 | 0.0080 | GRIA2 | ENSG00000120251 | 2.50 | 0.013495 |
| Afap1l2 | ENSRNOG00000017164 | -1.09 | 0.0000 | AFAP1L2 | ENSG00000169129 | -1.03 | 1.64E-12 |
| Sparcl1 | ENSRNOG00000015093 | -1.68 | 0.0000 | SPARCL1 | ENSG00000152583 | -1.20 | 0.000289 |
| Mmp28 | ENSRNOG00000047940 | -2.19 | 0.0000 | MMP28 | ENSG00000271447 | -1.18 | 1.11E-11 |
| Eln | ENSRNOG00000001469 | -1.69 | 0.0000 | ELN | ENSG00000049540 | -1.41 | 8.10E-06 |
| Dhh | ENSRNOG00000053675 | -1.60 | 0.0000 | DHH | ENSG00000139549 | -1.09 | 0.012232 |

*FC: fold change; Ctrl: vehicle control (DMSO); *P*-value: Benjamini Hochberg adjusted *P*-value. $P < 0.05$ is considered significantly differentially expressed.

Table 22B. ITE-dysregulated common genes in female rat placentas and HUVECs.*

| Rat Placentas | | | | HUVECs | | | |
|---|---|---|---|---|---|---|---|
| Symbol | Gene ID | Log FC (ITE/DMSO) | FDR *P*-Value | Symbol | Gene ID | Log FC (ITE/DMSO) | FDR *P*-Value |
| Abcb4 | ENSRNOG00000008012 | -1.07 | 0.000035 | ABCB4 | ENSG00000005471 | 1.00 | 0.018034 |
| Ahrr | ENSRNOG00000014721 | 4.05 | 0.000000 | AHRR | ENSG00000063438 | 2.08 | 1.03E-80 |
| Cyp1a1 | ENSRNOG00000019500 | 5.25 | 0.000000 | CYP1A1 | ENSG00000140465 | 9.55 | 0.000000 |
| Cyp1b1 | ENSRNOG00000040287 | 4.34 | 0.000000 | CYP1B1 | ENSG00000138061 | 7.73 | 0.000000 |
| F2rl3 | ENSRNOG00000048186 | 1.03 | 0.000010 | F2RL3 | ENSG00000127533 | 3.63 | 6.55E-32 |
| Nptx1 | ENSRNOG00000003741 | 1.06 | 0.000047 | NPTX1 | ENSG00000171246 | 7.09 | 0.000000 |
| Apold1 | ENSRNOG00000007830 | -1.79 | 0.000001 | APOLD1 | ENSG00000178878 | 1.65 | 3.42E-77 |
| Foxf1 | ENSRNOG00000049906 | -1.60 | 0.000000 | FOXF1 | ENSG00000103241 | 1.10 | 0.000149 |
| Ptgs2 | ENSRNOG00000002525 | -1.61 | 0.000187 | PTGS2 | ENSG00000073756 | 1.54 | 5.90E-89 |
| Tacstd2 | ENSRNOG00000007740 | -1.30 | 0.000232 | TACSTD2 | ENSG00000184292 | 1.19 | 9.49E-54 |
| Cbln2 | ENSRNOG00000013654 | -1.19 | 0.038098 | CBLN2 | ENSG00000141668 | -2.03 | 0.042592 |
| Mmp28 | ENSRNOG00000047940 | -1.90 | 0.000143 | MMP28 | ENSG00000271447 | -1.33 | 2.93E-08 |

*FC: fold change; Ctrl: vehicle control (DMSO); *P*-value: Benjamini Hochberg adjusted *P*-value. $P < 0.05$ is considered significantly differentially expressed.

Table 22C. ITE-dysregulated common genes in male rat placentas and HUVECs.*

| Rat Placentas | | | | HUVECs | | | |
|---|---|---|---|---|---|---|---|
| Symbol | Gene ID | Log FC (ITE/DMSO) | FDR *P*-Value | Symbol | Gene ID | Log FC (ITE/DMSO) | FDR *P*-Value |
| Cytl1 | ENSRNOG00000028108 | -3.40 | 0.0000 | CYTL1 | ENSG00000170891 | 1.78 | 9.53E-05 |
| Ptgs2 | ENSRNOG00000002525 | -2.73 | 0.0000 | PTGS2 | ENSG00000073756 | 1.10 | 2.21E-20 |
| Stc1 | ENSRNOG00000015075 | -1.15 | 0.0000 | STC1 | ENSG00000159167 | 1.14 | 0.020414 |
| Ahrr | ENSRNOG00000014721 | 3.62 | 0.0000 | AHRR | ENSG00000063438 | 1.98 | 2.56E-23 |
| C1s | ENSRNOG00000011971 | 1.29 | 0.0000 | C1S | ENSG00000182326 | 1.13 | 1.78E-05 |
| Cyp1a1 | ENSRNOG00000019500 | 4.01 | 0.0000 | CYP1A1 | ENSG00000140465 | 9.62 | 7.98E-21 |
| Cyp1b1 | ENSRNOG00000040287 | 4.09 | 0.0000 | CYP1B1 | ENSG00000138061 | 6.25 | 6.74E-21 |
| F2rl3 | ENSRNOG00000048186 | 1.36 | 0.0000 | F2RL3 | ENSG00000127533 | 1.88 | 0.023242 |
| Il1b | ENSRNOG00000004649 | 1.47 | 0.0000 | IL1B | ENSG00000125538 | 2.37 | 2.83E-08 |
| Il6 | ENSRNOG00000010278 | 3.65 | 0.0000 | IL6 | ENSG00000136244 | 1.53 | 3.16E-07 |
| Nptx1 | ENSRNOG00000003741 | 1.17 | 0.0000 | NPTX1 | ENSG00000171246 | 2.81 | 1.41E-07 |
| Nqo1 | ENSRNOG00000012772 | 1.51 | 0.0000 | NQO1 | ENSG00000181019 | 1.07 | 2.56E-20 |
| Rnd1 | ENSRNOG00000059857 | 1.19 | 0.0000 | RND1 | ENSG00000172602 | 1.28 | 3.71E-11 |
| Afap1l2 | ENSRNOG00000017164 | -1.15 | 0.0000 | AFAP1L2 | ENSG00000169129 | -1.09 | 0.000175 |
| Col14a1 | ENSRNOG00000026415 | -1.31 | 0.0000 | COL14A1 | ENSG00000187955 | -1.49 | 0.004600 |
| Eln | ENSRNOG00000001469 | -1.84 | 0.0000 | ELN | ENSG00000049540 | -1.80 | 0.000610 |
| Mmp28 | ENSRNOG00000047940 | -2.40 | 0.0000 | MMP28 | ENSG00000271447 | -1.02 | 0.001815 |
| Sparcl1 | ENSRNOG00000015093 | -1.99 | 0.0000 | SPARCL1 | ENSG00000152583 | -1.59 | 6.99E-06 |
| Cxcl10 | ENSRNOG00000022256 | 3.81 | 0.0000 | CXCL10 | ENSG00000169245 | -1.06 | 8.39E-06 |
| Isg15 | ENSRNOG00000021802 | 1.24 | 0.0000 | ISG15 | ENSG00000187608 | -1.05 | 2.52E-06 |

*FC: fold change; Ctrl: vehicle control (DMSO); *P*-value: Benjamini Hochberg adjusted *P*-value. $P < 0.05$ is considered significantly differentially expressed.

Table S23. ITE-dysregulated common biological function in rat placentas and HUVECs.*

| Biological Function | *P*-Value [Female Placentas] | *P*-Value [Male Placentas] | *P*-Value [Female HUVECs] | *P*-Value [Male HUVECs] |
|---|---|---|---|---|
| Abdominal adenocarcinoma | **9.26E-37** | **3.96E-60** | **3.52E-10** | **2.39E-05** |
| Abdominal cancer | **1.22E-38** | **2.13E-62** | **1.47E-11** | **8.19E-06** |
| Abdominal carcinoma | **6.75E-39** | **1.24E-62** | **2.76E-10** | **5.18E-05** |
| Abnormal morphology of body cavity | **9.77E-19** | **3.43E-18** | **8.67E-07** | **1.93E-04** |
| Abnormal morphology of thoracic cavity | **6.23E-13** | **1.40E-10** | **8.93E-06** | **1.79E-05** |
| Abnormality of left ventricle | **1.08E-10** | **3.72E-12** | **6.41E-04** | **5.03E-05** |
| Activation of cells | **1.37E-08** | **5.84E-24** | **1.62E-06** | **3.62E-05** |
| Adenocarcinoma | **1.33E-37** | **1.22E-61** | **3.52E-10** | **1.32E-05** |
| Angiogenesis | **3.57E-26** | **1.50E-23** | **2.66E-09** | **1.81E-10** |
| Anogenital cancer | **7.51E-19** | **3.12E-30** | **5.63E-11** | **8.43E-09** |
| Blood pressure | **7.09E-09** | **9.27E-17** | **3.58E-04** | **3.05E-04** |
| Breast or colorectal cancer | **1.46E-16** | **1.04E-23** | **3.36E-07** | **4.82E-05** |
| Breast or gynecological cancer | **9.43E-16** | **7.02E-22** | **2.20E-10** | **7.49E-07** |
| Breast or ovarian cancer | **3.74E-07** | **5.61E-11** | **3.36E-07** | **1.21E-06** |
| Breast or ovarian carcinoma | **3.49E-06** | **2.99E-09** | **5.90E-05** | **1.85E-04** |
| Breast or pancreatic cancer | **5.70E-12** | **2.14E-18** | **6.21E-09** | **1.21E-06** |
| Cancer of cells | **1.87E-15** | **2.40E-24** | **2.60E-09** | **2.77E-04** |
| Cancer of secretory structure | **4.58E-38** | **6.05E-52** | **1.46E-06** | **2.71E-05** |
| Carcinoma | **5.72E-49** | **1.39E-70** | **3.41E-09** | **2.77E-04** |
| Cell movement | **6.75E-20** | **1.62E-30** | **1.47E-11** | **1.33E-08** |
| Cell movement of endothelial cells | **2.28E-08** | **1.37E-09** | **7.92E-04** | **4.90E-06** |
| Cell movement of leukocytes | **2.74E-08** | **1.68E-22** | **6.98E-05** | **1.46E-05** |
| Cell movement of myeloid cells | **9.07E-06** | **2.25E-20** | **3.98E-05** | **6.58E-07** |
| Chronic inflammatory disorder | **3.69E-08** | **4.79E-17** | **9.08E-07** | **5.20E-05** |
| Development of abdomen | **5.46E-07** | **9.12E-12** | **3.41E-06** | **5.14E-06** |
| Development of body trunk | **4.31E-21** | **3.34E-19** | **6.49E-06** | **2.33E-05** |
| Development of connective tissue cells | **2.97E-07** | **9.56E-09** | **3.81E-04** | **1.90E-05** |
| Development of epithelial tissue | **4.71E-12** | **1.39E-13** | **2.47E-05** | **6.63E-07** |
| Development of genital tumor | **2.61E-14** | **1.13E-18** | **6.28E-10** | **2.16E-06** |
| Development of lung tumor | **1.21E-08** | **6.90E-13** | **6.99E-06** | **4.96E-05** |
| Development of vasculature | **5.65E-28** | **1.98E-23** | **1.34E-09** | **1.81E-10** |
| Diabetes mellitus | **9.91E-11** | **5.93E-23** | **9.47E-04** | **1.34E-06** |
| Digestive system cancer | **1.49E-34** | **2.03E-57** | **6.28E-10** | **6.87E-05** |
| Disorder of blood pressure | **2.76E-21** | **1.80E-24** | **1.18E-05** | **4.31E-06** |
| Disorder of pregnancy | **1.03E-11** | **4.54E-12** | **4.57E-04** | **7.86E-06** |
| Endometrial cancer | **2.23E-12** | **2.99E-17** | **6.86E-08** | **9.44E-05** |
| Endothelial cell development | **6.89E-12** | **2.79E-12** | **1.60E-04** | **6.63E-07** |
| Extraadrenal retroperitoneal tumor | **4.94E-13** | **1.39E-21** | **3.12E-05** | **9.25E-05** |
| Extracranial solid tumor | **9.86E-45** | **1.67E-65** | **1.02E-10** | **6.34E-06** |
| Extrapancreatic malignant tumor | **9.08E-45** | **1.06E-69** | **7.26E-11** | **1.27E-06** |
| Female genital tract adenocarcinoma | **1.90E-11** | **6.31E-16** | **5.58E-07** | **1.67E-04** |

| | | | | |
|---|---|---|---|---|
| Female genital tract cancer | **2.10E-14** | **7.76E-20** | **7.10E-10** | **8.15E-07** |
| Female genital tract serous carcinoma | **1.19E-12** | **1.28E-09** | **4.26E-04** | **5.38E-07** |
| Fibrosis | **2.54E-08** | **6.67E-13** | **4.63E-08** | **1.33E-08** |
| Formation of solid tumor | **3.38E-40** | **6.27E-60** | **3.67E-11** | **7.58E-06** |
| Gastrointestinal adenocarcinoma | **4.99E-29** | **8.00E-49** | **1.08E-08** | **4.18E-05** |
| Gastrointestinal carcinoma | **4.07E-35** | **2.26E-52** | **1.59E-08** | **5.52E-05** |
| Genital tract cancer | **1.26E-20** | **2.44E-31** | **6.28E-10** | **2.09E-08** |
| Genital tumor | **3.82E-22** | **4.68E-33** | **4.75E-10** | **3.15E-08** |
| Genitourinary adenocarcinoma | **2.02E-15** | **7.09E-26** | **2.27E-06** | **1.11E-05** |
| Genitourinary carcinoma | **1.20E-17** | **3.81E-31** | **1.54E-08** | **4.64E-06** |
| Growth of epithelial tissue | **1.85E-08** | **7.20E-09** | **1.55E-06** | **4.21E-05** |
| Head and neck cancer | **2.35E-38** | **4.64E-52** | **7.71E-07** | **1.92E-04** |
| Head and neck tumor | **3.03E-43** | **1.57E-62** | **4.10E-07** | **1.59E-04** |
| Hematologic cancer | **1.20E-10** | **4.72E-19** | **9.16E-05** | **1.14E-06** |
| Hematologic cancer of cells | **4.14E-10** | **1.90E-16** | **5.21E-04** | **7.46E-07** |
| Hypertension | **6.88E-22** | **1.07E-23** | **2.08E-05** | **1.06E-05** |
| Incidence of tumor | **7.04E-29** | **3.09E-40** | **1.31E-08** | **2.63E-05** |
| Inflammation of absolute anatomical region | **9.91E-06** | **3.94E-17** | **7.71E-07** | **1.02E-04** |
| Inflammation of joint | **6.53E-11** | **1.87E-21** | **4.38E-08** | **1.06E-05** |
| Inflammation of organ | **3.84E-06** | **1.55E-16** | **2.20E-07** | **3.45E-05** |
| Inflammatory response | **3.86E-07** | **1.96E-20** | **1.15E-06** | **1.17E-05** |
| Intraabdominal organ tumor | **1.73E-38** | **5.97E-61** | **3.06E-10** | **3.24E-05** |
| Large intestine adenocarcinoma | **8.67E-31** | **1.86E-46** | **1.51E-08** | **1.27E-04** |
| Leukocyte migration | **4.19E-09** | **7.22E-23** | **1.58E-07** | **1.18E-07** |
| Liver lesion | **3.51E-19** | **4.92E-26** | **2.26E-04** | **3.24E-04** |
| Lung adenocarcinoma | **1.28E-08** | **2.94E-12** | **1.43E-05** | **1.20E-04** |
| Lung cancer | **2.86E-16** | **5.10E-23** | **5.16E-06** | **4.37E-05** |
| Lymphocytic cancer | **1.63E-09** | **5.38E-15** | **1.34E-03** | **3.65E-06** |
| Lymphocytic neoplasm | **1.83E-09** | **6.40E-15** | **1.36E-03** | **3.70E-06** |
| Lymphoreticular neoplasm | **7.89E-08** | **7.91E-15** | **1.86E-04** | **6.35E-07** |
| Malignant genitourinary solid tumor | **7.55E-20** | **1.52E-32** | **3.52E-10** | **3.92E-08** |
| Malignant neoplasm of large intestine | **1.09E-30** | **1.90E-46** | **1.34E-07** | **2.43E-05** |
| Malignant neoplasm of respiratory system | **2.64E-17** | **3.07E-24** | **5.24E-07** | **6.02E-05** |
| Malignant neoplasm of retroperitoneum | **9.88E-12** | **7.18E-17** | **1.92E-04** | **5.30E-05** |
| Malignant solid tumor | **2.41E-46** | **2.11E-66** | **1.19E-10** | **7.08E-06** |
| Mature lymphocytic neoplasm | **1.49E-07** | **6.65E-11** | **6.53E-04** | **1.92E-04** |
| Migration of cells | **1.68E-18** | **1.85E-29** | **3.67E-11** | **2.09E-08** |
| Morphology of body cavity | **3.42E-23** | **2.79E-21** | **5.08E-07** | **9.82E-05** |
| Morphology of bone | **2.77E-08** | **2.06E-12** | **1.45E-08** | **7.46E-07** |
| Morphology of cardiovascular system | **3.02E-16** | **1.26E-11** | **1.25E-04** | **2.74E-04** |
| Morphology of heart | **3.41E-11** | **3.78E-09** | **1.93E-04** | **2.85E-04** |
| Myeloid or lymphoid neoplasm | **5.09E-10** | **3.92E-18** | **8.94E-06** | **1.26E-06** |
| Neoplasia of blood cells | **1.16E-09** | **1.61E-18** | **5.78E-06** | **9.64E-07** |
| Neoplasia of cells | **2.10E-15** | **1.31E-29** | **6.51E-10** | **1.92E-04** |
| Neoplasia of leukocytes | **2.09E-09** | **5.53E-16** | **9.28E-04** | **2.62E-06** |

| | | | | |
|---|---|---|---|---|
| Non-small cell lung carcinoma | **3.38E-10** | **1.98E-14** | **6.71E-06** | **6.86E-05** |
| Non-traumatic arthropathy | **4.97E-08** | **1.62E-15** | **1.58E-07** | **2.94E-05** |
| Organismal death | **4.52E-14** | **8.17E-13** | **5.38E-09** | **1.27E-06** |
| Osteoarthritis | **1.38E-06** | **7.97E-11** | **3.60E-08** | **2.82E-05** |
| Ovarian tumor | **8.19E-07** | **4.21E-09** | **2.00E-05** | **4.48E-05** |
| Pancreatobiliary tumor | **1.48E-11** | **9.53E-20** | **7.45E-06** | **2.27E-04** |
| Pelvic adenocarcinoma | **9.19E-16** | **6.08E-26** | **2.27E-06** | **2.71E-05** |
| Pelvic cancer | **6.37E-23** | **2.28E-33** | **2.88E-09** | **2.32E-08** |
| Pelvic carcinoma | **8.61E-18** | **2.79E-28** | **7.83E-08** | **3.20E-06** |
| Pelvic serous carcinoma | **1.19E-12** | **1.28E-09** | **4.26E-04** | **5.38E-07** |
| Peripheral vascular disease | **3.15E-08** | **3.09E-14** | **3.08E-04** | **9.09E-05** |
| Prostate cancer | **1.00E-14** | **2.34E-22** | **6.19E-05** | **1.37E-04** |
| Quantity of blood cells | **1.78E-15** | **1.62E-24** | **7.18E-04** | **8.19E-06** |
| Quantity of cells | **4.12E-18** | **5.85E-26** | **3.92E-06** | **2.53E-06** |
| Quantity of granulocytes | **1.58E-06** | **2.65E-14** | **2.30E-04** | **7.58E-06** |
| Quantity of leukocytes | **5.63E-13** | **1.38E-23** | **5.14E-04** | **9.54E-06** |
| Quantity of myeloid cells | **4.31E-11** | **4.19E-19** | **5.15E-04** | **2.04E-05** |
| Quantity of phagocytes | **2.37E-08** | **6.44E-17** | **1.50E-05** | **3.45E-06** |
| Recruitment of cells | **1.77E-07** | **1.06E-21** | **3.08E-08** | **1.01E-07** |
| Recruitment of leukocytes | **3.80E-06** | **1.04E-19** | **3.36E-07** | **1.27E-06** |
| Recruitment of myeloid cells | **5.34E-06** | **3.31E-17** | **9.70E-06** | **2.20E-06** |
| Recruitment of neutrophils | **6.36E-06** | **2.10E-17** | **7.04E-05** | **9.82E-07** |
| Rheumatic Disease | **1.04E-09** | **2.08E-22** | **1.51E-08** | **4.54E-07** |
| Serous adenocarcinoma | **1.52E-10** | **5.29E-09** | **5.15E-04** | **3.80E-06** |
| Skin tumor | **4.24E-35** | **2.26E-60** | **3.82E-07** | **2.62E-04** |
| Systemic autoimmune syndrome | **8.92E-08** | **4.60E-21** | **9.81E-05** | **2.14E-05** |
| Tumorigenesis of reproductive tract | **8.89E-15** | **1.08E-20** | **6.28E-10** | **1.25E-06** |
| Uterine cancer | **5.47E-13** | **1.59E-17** | **3.13E-08** | **6.58E-06** |
| Uterine tumor | **8.95E-14** | **1.53E-18** | **9.06E-10** | **1.27E-06** |
| Valvulopathy | **1.77E-06** | **5.64E-10** | **1.02E-03** | **2.08E-04** |
| Vasculogenesis | **1.60E-22** | **1.70E-20** | **1.59E-08** | **1.33E-08** |
| Vaso-occlusion | **4.80E-10** | **4.78E-17** | **9.57E-04** | **6.63E-07** |
| Abnormal morphology of abdomen | **5.98E-09** | **4.40E-12** | **2.85E-05** | 1.00E+00 |
| Abnormal morphology of cardiovascular system | **7.68E-14** | **3.20E-10** | **2.72E-04** | 1.00E+00 |
| Abnormal morphology of digestive system | **3.96E-06** | **4.03E-09** | **2.79E-04** | 1.00E+00 |
| Abnormal morphology of heart | **8.68E-11** | **7.55E-09** | **2.38E-04** | 1.00E+00 |
| Astrocytoma | **2.09E-20** | **7.13E-25** | **5.30E-05** | 1.00E+00 |
| Bleeding | **2.57E-10** | **1.48E-09** | **1.36E-03** | 1.00E+00 |
| Brain astrocytoma | **1.21E-16** | **7.43E-22** | **1.36E-04** | 1.00E+00 |
| Brain glioma | **1.72E-25** | **3.98E-34** | **2.26E-04** | 1.00E+00 |
| Brain tumor | **1.63E-24** | **2.68E-33** | **7.56E-05** | 1.00E+00 |
| Central nervous system cancer | **4.21E-26** | **3.64E-34** | **1.38E-05** | 1.00E+00 |
| Central nervous system solid tumor | **2.31E-26** | **4.62E-36** | **1.43E-05** | 1.00E+00 |
| Colon tumor | **1.67E-17** | **9.60E-26** | **4.63E-04** | 1.00E+00 |

| | | | | |
|---|---|---|---|---|
| Colorectal adenocarcinoma | **4.52E-13** | **6.45E-20** | **8.30E-04** | 1.00E+00 |
| Colorectal cancer | **1.42E-16** | **5.40E-23** | **6.41E-04** | 1.00E+00 |
| Colorectal tumor | **2.79E-18** | **8.46E-24** | **2.74E-04** | 1.00E+00 |
| Cutaneous melanoma | **1.02E-34** | **1.64E-57** | **5.65E-07** | 1.00E+00 |
| Development of adenocarcinoma | **7.92E-11** | **3.28E-17** | **5.55E-06** | 1.00E+00 |
| Development of carcinoma | **2.04E-20** | **2.09E-32** | **1.06E-06** | 1.00E+00 |
| Development of digestive organ tumor | **6.35E-19** | **3.72E-28** | **1.59E-04** | 1.00E+00 |
| Development of head | **1.65E-10** | **1.23E-08** | **3.06E-04** | 1.00E+00 |
| Development of malignant tumor | **3.82E-22** | **3.23E-34** | **6.81E-08** | 1.00E+00 |
| Endocrine carcinoma | **1.68E-33** | **4.94E-45** | **4.87E-04** | 1.00E+00 |
| Endocrine gland tumor | **3.83E-34** | **1.37E-49** | **2.30E-04** | 1.00E+00 |
| Endometrial adenocarcinoma | **5.40E-07** | **6.90E-12** | **8.62E-05** | 1.00E+00 |
| Endometrial carcinoma | **1.50E-07** | **9.36E-14** | **6.83E-06** | 1.00E+00 |
| Endometrioid carcinoma | **3.18E-06** | **7.95E-12** | **1.68E-04** | 1.00E+00 |
| Endometrioid endometrial adenocarcinoma | **6.69E-07** | **1.66E-11** | **2.38E-04** | 1.00E+00 |
| Endometrium tumor | **1.59E-12** | **2.16E-17** | **2.44E-08** | 1.00E+00 |
| Extrapulmonary squamous cell carcinoma | **1.66E-08** | **2.89E-14** | **2.20E-07** | 1.00E+00 |
| Female genital neoplasm | **2.42E-15** | **5.48E-21** | **6.28E-10** | 1.00E+00 |
| Genitourinary tumor | **8.41E-20** | **2.33E-32** | **3.06E-10** | 1.00E+00 |
| Glioma | **2.22E-28** | **2.80E-37** | **8.15E-05** | 1.00E+00 |
| Glioma cancer | **9.04E-28** | **3.21E-35** | **8.50E-05** | 1.00E+00 |
| Grade 1-4 astrocytoma | **4.02E-17** | **3.00E-22** | **1.47E-04** | 1.00E+00 |
| Grade 3-4 glioma cancer | **2.61E-18** | **3.38E-22** | **2.30E-04** | 1.00E+00 |
| Grade 4 astrocytoma | **8.01E-16** | **1.47E-21** | **2.49E-04** | 1.00E+00 |
| Grade 4 high grade glioma | **8.01E-16** | **1.47E-21** | **2.49E-04** | 1.00E+00 |
| Grade 4 malignant glioma | **8.01E-16** | **1.47E-21** | **2.49E-04** | 1.00E+00 |
| Head and neck carcinoma | **3.55E-34** | **4.69E-48** | **2.46E-06** | 1.00E+00 |
| Head and neck squamous cell carcinoma | **2.11E-06** | **2.40E-11** | **2.93E-07** | 1.00E+00 |
| High grade astrocytoma | **3.58E-20** | **1.57E-24** | **2.03E-04** | 1.00E+00 |
| Hypersensitive reaction | **4.27E-07** | **3.98E-11** | **2.76E-05** | 1.00E+00 |
| Leukemia | **1.05E-05** | **9.38E-11** | **3.22E-04** | 1.00E+00 |
| Liquid tumor | **3.04E-06** | **1.24E-11** | **3.10E-05** | 1.00E+00 |
| Lung carcinoma | **6.35E-16** | **5.40E-22** | **2.10E-05** | 1.00E+00 |
| Lung tumor | **9.42E-17** | **1.46E-23** | **2.80E-06** | 1.00E+00 |
| Malignant soft tissue neoplasm | **3.65E-34** | **1.88E-56** | **1.84E-07** | 1.00E+00 |
| Melanoma | **2.87E-35** | **1.42E-61** | **1.36E-07** | 1.00E+00 |
| Morbidity or mortality | **3.06E-14** | **9.67E-14** | **2.36E-09** | 1.00E+00 |
| Morphology of digestive system | **2.11E-08** | **9.90E-13** | **3.77E-05** | 1.00E+00 |
| Multiple cancers | **4.11E-07** | **3.53E-11** | **3.64E-07** | 1.00E+00 |
| Neck neoplasm | **6.26E-33** | **1.92E-42** | **1.35E-03** | 1.00E+00 |
| Non-colon gastrointestinal cancer | **7.60E-16** | **2.14E-20** | **1.55E-06** | 1.00E+00 |
| Nonhematologic malignant neoplasm | **5.75E-50** | **7.88E-71** | **1.92E-09** | 1.00E+00 |
| Nonpituitary endocrine tumor | **2.25E-30** | **9.67E-44** | **7.92E-04** | 1.00E+00 |
| Pancreatic cancer | **7.83E-11** | **1.22E-16** | **1.09E-04** | 1.00E+00 |
| Pancreatic carcinoma | **1.56E-09** | **7.83E-15** | **9.57E-04** | 1.00E+00 |

| | | | | |
|---|---|---|---|---|
| Pancreatic tumor | **1.64E-11** | **1.27E-20** | **2.34E-05** | 1.00E+00 |
| Pelvic tumor | **4.39E-24** | **5.00E-35** | **1.87E-09** | 1.00E+00 |
| Prostatic tumor | **3.99E-15** | **1.14E-22** | **1.31E-05** | 1.00E+00 |
| Quantity of bone | **1.31E-06** | **3.28E-09** | **9.81E-05** | 1.00E+00 |
| Quantity of connective tissue | **5.17E-14** | **6.99E-18** | **2.38E-04** | 1.00E+00 |
| Quantity of metal | **3.76E-10** | **2.67E-15** | **3.50E-04** | 1.00E+00 |
| Quantity of metal ion | **4.62E-08** | **2.09E-13** | **3.90E-04** | 1.00E+00 |
| Quantity of protein in blood | **1.61E-07** | **9.43E-23** | **8.25E-04** | 1.00E+00 |
| Respiratory system tumor | **8.33E-19** | **1.39E-25** | **2.91E-07** | 1.00E+00 |
| Size of bone | **8.48E-06** | **5.23E-10** | **1.14E-05** | 1.00E+00 |
| Skin cancer | **5.65E-35** | **1.32E-59** | **8.62E-07** | 1.00E+00 |
| Skin carcinoma | **9.06E-08** | **1.55E-14** | **8.48E-04** | 1.00E+00 |
| Soft tissue neoplasm | **3.19E-34** | **1.88E-56** | **6.59E-08** | 1.00E+00 |
| Squamous-cell carcinoma | **6.10E-10** | **5.42E-18** | **8.75E-08** | 1.00E+00 |
| Stomach tumor | **1.09E-12** | **1.86E-14** | **7.01E-04** | 1.00E+00 |
| Upper gastrointestinal carcinoma | **1.47E-15** | **3.82E-19** | **1.09E-05** | 1.00E+00 |
| Upper gastrointestinal tract cancer | **9.83E-16** | **2.33E-19** | **1.19E-05** | 1.00E+00 |
| Upper gastrointestinal tract tumor | **1.11E-15** | **1.84E-19** | **5.92E-06** | 1.00E+00 |
| Uterine carcinoma | **7.57E-10** | **1.46E-15** | **6.77E-07** | 1.00E+00 |
| Uterine endometrioid carcinoma | **7.75E-07** | **2.35E-11** | **1.32E-04** | 1.00E+00 |
| Acute respiratory disorder | **7.60E-07** | **1.20E-08** | 1.00E+00 | **1.29E-04** |
| Asthma | **1.30E-08** | **7.30E-10** | 1.00E+00 | **7.38E-05** |
| Atherosclerosis | **3.12E-09** | **3.87E-16** | 1.00E+00 | **1.52E-05** |
| Diabetic complication | **1.37E-08** | **2.24E-14** | 1.00E+00 | **3.65E-06** |
| Glucose metabolism disorder | **9.57E-14** | **3.38E-22** | 1.00E+00 | **1.34E-06** |
| Gonadal tumor | **5.77E-07** | **7.81E-09** | 1.00E+00 | **2.71E-05** |
| Homing of cells | **5.63E-06** | **6.19E-15** | 1.00E+00 | **3.70E-07** |
| Infarction | **1.07E-09** | **3.12E-13** | 1.00E+00 | **1.01E-04** |
| Inflammation of lung | **5.48E-07** | **1.13E-11** | 1.00E+00 | **2.84E-04** |
| Inflammation of respiratory system component | **3.65E-08** | **6.26E-16** | 1.00E+00 | **6.55E-05** |
| Large intestine carcinoma | **2.05E-31** | **6.58E-48** | 1.00E+00 | **5.60E-05** |
| Lymphatic system tumor | **2.11E-09** | **1.33E-14** | 1.00E+00 | **1.52E-06** |
| Lymphoma | **2.21E-08** | **5.47E-13** | 1.00E+00 | **5.38E-07** |
| Morphology of vasculature | **4.36E-13** | **1.17E-09** | 1.00E+00 | **2.74E-04** |
| Myocardial infarction | **1.28E-10** | **2.15E-11** | 1.00E+00 | **1.76E-04** |
| Non-Hodgkin lymphoma | **1.09E-09** | **1.63E-12** | 1.00E+00 | **2.13E-05** |
| Non-melanoma solid tumor | **4.30E-54** | **9.50E-75** | 1.00E+00 | **1.16E-04** |
| Occlusion of artery | **1.72E-09** | **1.25E-16** | 1.00E+00 | **2.38E-06** |
| Occlusion of blood vessel | **2.75E-10** | **5.72E-17** | 1.00E+00 | **6.35E-07** |
| Preeclampsia | **1.69E-12** | **8.71E-13** | 1.00E+00 | **7.50E-05** |
| Quantity of lymphocytes | **2.44E-09** | **2.09E-13** | 1.00E+00 | **1.45E-04** |
| Stimulation of cells | **2.96E-08** | **1.22E-10** | 1.00E+00 | **2.03E-04** |
| Transmigration of cells | **4.15E-09** | **2.94E-11** | 1.00E+00 | **9.64E-07** |
| Transmigration of leukocytes | **4.06E-07** | **8.99E-09** | 1.00E+00 | **1.52E-05** |

| | | | | |
|---|---|---|---|---|
| Tumorigenesis of lymphocytes | **1.49E-09** | **1.82E-15** | 1.00E+00 | **1.19E-05** |
| Uterine serous papillary cancer | **1.03E-09** | **1.37E-08** | 1.00E+00 | **4.87E-05** |
| Vascular lesion | **6.86E-07** | **1.47E-11** | 1.00E+00 | **1.88E-05** |
| Weight gain | **2.38E-07** | **1.80E-11** | 1.00E+00 | **2.07E-04** |
| Enlargement of heart | **3.44E-08** | 1.00E+00 | **5.65E-05** | **9.82E-05** |
| Hypertrophy | **1.47E-09** | 1.00E+00 | **2.51E-06** | **1.52E-06** |
| Hypertrophy of heart | **3.04E-08** | 1.00E+00 | **2.85E-05** | **1.52E-05** |
| Invasive tumor | **6.80E-06** | 1.00E+00 | **1.50E-06** | **3.53E-07** |
| Migration of vascular cells | **1.10E-07** | 1.00E+00 | **6.50E-04** | **1.27E-06** |
| Ovarian cancer | **4.11E-06** | 1.00E+00 | **2.26E-04** | **2.79E-04** |
| Proliferation of endothelial cells | **2.38E-07** | 1.00E+00 | **2.30E-04** | **7.56E-06** |
| Proliferation of muscle cells | **1.12E-05** | 1.00E+00 | **3.50E-04** | **3.42E-07** |
| Vascularization | **1.34E-06** | 1.00E+00 | **5.31E-05** | **2.25E-05** |
| Visceromegaly | **1.96E-09** | 1.00E+00 | **1.52E-05** | **1.94E-05** |
| Chronic kidney disease | **3.97E-07** | 1.00E+00 | **1.03E-03** | 1.00E+00 |
| Development of body axis | **3.82E-09** | 1.00E+00 | **2.39E-04** | 1.00E+00 |
| Morphology of head | **8.11E-08** | 1.00E+00 | **1.25E-03** | 1.00E+00 |
| Morphology of muscle cells | **7.68E-06** | 1.00E+00 | **1.80E-04** | 1.00E+00 |
| Organization of extracellular matrix | **1.58E-06** | 1.00E+00 | **1.37E-05** | 1.00E+00 |
| Size of body | **2.54E-08** | 1.00E+00 | **4.93E-05** | 1.00E+00 |
| Formation of lymphoid tissue | **6.50E-06** | 1.00E+00 | 1.00E+00 | **1.66E-04** |
| Migration of endothelial cells | **5.48E-07** | 1.00E+00 | 1.00E+00 | **6.74E-05** |
| Migration of vascular smooth muscle cells | **5.10E-06** | 1.00E+00 | 1.00E+00 | **2.89E-06** |
| Morphology of muscle | **2.01E-09** | 1.00E+00 | 1.00E+00 | **4.77E-05** |
| Permeability of vascular system | **1.02E-05** | 1.00E+00 | 1.00E+00 | **3.05E-04** |
| Proliferation of smooth muscle cells | **9.06E-07** | 1.00E+00 | 1.00E+00 | **2.83E-06** |
| Uterine serous carcinoma | **6.09E-11** | 1.00E+00 | 1.00E+00 | **1.32E-05** |
| Abnormality of large intestine | 1.00E+00 | **6.39E-12** | **7.97E-06** | **1.19E-04** |
| Accumulation of blood cells | 1.00E+00 | **8.25E-10** | **3.00E-04** | **2.71E-06** |
| Accumulation of leukocytes | 1.00E+00 | **1.05E-10** | **7.55E-04** | **9.57E-06** |
| Activation of antigen presenting cells | 1.00E+00 | **4.42E-11** | **2.54E-04** | **2.32E-04** |
| Activation of blood cells | 1.00E+00 | **1.65E-17** | **9.27E-05** | **7.86E-06** |
| Acute colitis | 1.00E+00 | **3.33E-09** | **1.47E-04** | **1.45E-04** |
| Adhesion of immune cells | 1.00E+00 | **9.78E-12** | **3.03E-05** | **1.58E-05** |
| Binding of blood cells | 1.00E+00 | **1.75E-13** | **1.18E-05** | **2.83E-06** |
| Binding of leukocytes | 1.00E+00 | **3.84E-12** | **1.43E-05** | **5.44E-06** |
| Binding of professional phagocytic cells | 1.00E+00 | **3.45E-10** | **7.92E-04** | **9.82E-05** |
| Cell movement of granulocytes | 1.00E+00 | **1.39E-16** | **3.58E-04** | **3.71E-05** |
| Cell movement of mononuclear leukocytes | 1.00E+00 | **2.08E-14** | **6.16E-04** | **8.28E-06** |
| Cell movement of neutrophils | 1.00E+00 | **1.69E-14** | **1.04E-03** | **5.05E-05** |
| Cell movement of phagocytes | 1.00E+00 | **1.32E-17** | **4.29E-04** | **2.74E-06** |
| Cellular infiltration by leukocytes | 1.00E+00 | **1.28E-18** | **1.93E-05** | **1.88E-05** |
| Cellular infiltration by myeloid cells | 1.00E+00 | **3.34E-15** | **1.30E-03** | **1.18E-05** |
| Chemotaxis | 1.00E+00 | **4.31E-14** | **9.82E-06** | **9.38E-07** |
| Chemotaxis of leukocytes | 1.00E+00 | **1.15E-12** | **1.39E-04** | **1.56E-05** |

| | | | | |
|---|---|---|---|---|
| Chemotaxis of myeloid cells | 1.00E+00 | **5.94E-12** | **9.13E-05** | **1.90E-05** |
| Chemotaxis of phagocytes | 1.00E+00 | **4.30E-12** | **1.04E-04** | **2.13E-05** |
| COVID-19 | 1.00E+00 | **1.73E-10** | **9.81E-04** | **1.52E-06** |
| Function of leukocytes | 1.00E+00 | **9.38E-11** | **1.11E-05** | **5.96E-05** |
| Function of myeloid cells | 1.00E+00 | **9.13E-10** | **4.66E-05** | **3.78E-05** |
| Function of phagocytes | 1.00E+00 | **2.54E-09** | **2.93E-07** | **4.53E-06** |
| Healing of lesion | 1.00E+00 | **1.49E-08** | **5.05E-05** | **5.44E-06** |
| Homing of mononuclear leukocytes | 1.00E+00 | **4.72E-10** | **1.47E-04** | **1.29E-05** |
| Infection by coronavirus | 1.00E+00 | **4.02E-10** | **2.26E-04** | **9.38E-07** |
| Inflammation of body cavity | 1.00E+00 | **5.41E-18** | **4.29E-07** | **1.12E-04** |
| Inflammation of gastrointestinal tract | 1.00E+00 | **1.49E-11** | **3.00E-05** | **2.71E-04** |
| Interaction of lymphocytes | 1.00E+00 | **5.35E-09** | **1.20E-03** | **4.53E-06** |
| Interaction of mononuclear leukocytes | 1.00E+00 | **6.12E-09** | **2.48E-05** | **3.38E-07** |
| Interaction of phagocytes | 1.00E+00 | **6.82E-10** | **4.34E-05** | **2.98E-06** |
| Lower respiratory tract disorder | 1.00E+00 | **8.79E-09** | **1.87E-05** | **3.23E-05** |
| Migration of granulocytes | 1.00E+00 | **6.65E-09** | **8.22E-04** | **2.98E-06** |
| Nonsquamous non-small cell lung carcinoma | 1.00E+00 | **2.77E-12** | **2.07E-06** | **4.48E-05** |
| Quantity of neutrophils | 1.00E+00 | **6.00E-12** | **4.06E-04** | **5.31E-06** |
| Recruitment of phagocytes | 1.00E+00 | **3.03E-15** | **6.42E-07** | **1.21E-06** |
| Survival of organism | 1.00E+00 | **3.78E-10** | **2.55E-05** | **5.38E-07** |
| Accumulation of cells | 1.00E+00 | **4.54E-12** | **2.49E-04** | 1.00E+00 |
| Activation of phagocytes | 1.00E+00 | **1.46E-14** | **2.47E-04** | 1.00E+00 |
| Cellular infiltration | 1.00E+00 | **1.39E-21** | **3.95E-06** | 1.00E+00 |
| Colitis | 1.00E+00 | **4.25E-09** | **1.75E-04** | 1.00E+00 |
| Enteritis | 1.00E+00 | **1.27E-09** | **3.17E-05** | 1.00E+00 |
| Fatty acid metabolism | 1.00E+00 | **2.09E-09** | **6.05E-04** | 1.00E+00 |
| Infection by SARS coronavirus | 1.00E+00 | **7.98E-11** | **3.81E-04** | 1.00E+00 |
| Inflammation of the large intestine | 1.00E+00 | **1.23E-09** | **6.63E-05** | 1.00E+00 |
| Leukopoiesis | 1.00E+00 | **7.83E-09** | **2.39E-04** | 1.00E+00 |
| Malignant neoplasm of aerodigestive tract | 1.00E+00 | **3.55E-12** | **3.95E-04** | 1.00E+00 |
| Quantity of antigen presenting cells | 1.00E+00 | **3.37E-10** | **5.01E-04** | 1.00E+00 |
| Recruitment of antigen presenting cells | 1.00E+00 | **5.64E-10** | **1.64E-04** | 1.00E+00 |
| Rheumatoid arthritis | 1.00E+00 | **6.06E-11** | **3.86E-04** | 1.00E+00 |
| Upper aerodigestive tract carcinoma | 1.00E+00 | **6.86E-12** | **6.05E-04** | 1.00E+00 |
| Accumulation of granulocytes | 1.00E+00 | **1.20E-11** | 1.00E+00 | **2.27E-05** |
| Accumulation of myeloid cells | 1.00E+00 | **2.19E-10** | 1.00E+00 | **3.71E-05** |
| Binding of lymphocytes | 1.00E+00 | **4.58E-09** | 1.00E+00 | **2.13E-05** |
| Binding of myeloid cells | 1.00E+00 | **1.04E-09** | 1.00E+00 | **1.33E-04** |
| Binding of tumor cell lines | 1.00E+00 | **7.53E-10** | 1.00E+00 | **2.25E-04** |
| Cell death of immune cells | 1.00E+00 | **2.32E-10** | 1.00E+00 | **2.66E-04** |
| Cell movement of antigen presenting cells | 1.00E+00 | **6.95E-11** | 1.00E+00 | **6.42E-06** |
| Cell movement of lymphocytes | 1.00E+00 | **1.59E-12** | 1.00E+00 | **2.36E-04** |
| Cell movement of macrophages | 1.00E+00 | **1.41E-10** | 1.00E+00 | **2.94E-05** |
| Cell movement of T lymphocytes | 1.00E+00 | **1.30E-08** | 1.00E+00 | **3.03E-04** |

| | | | | |
|---|---|---|---|---|
| Cellular infiltration by granulocytes | 1.00E+00 | **1.21E-14** | 1.00E+00 | **5.33E-06** |
| Cellular infiltration by phagocytes | 1.00E+00 | **5.10E-11** | 1.00E+00 | **3.46E-05** |
| Homing of leukocytes | 1.00E+00 | **3.43E-14** | 1.00E+00 | **4.53E-06** |
| Infection of mammalia | 1.00E+00 | **5.73E-14** | 1.00E+00 | **1.90E-05** |
| Infiltration by neutrophils | 1.00E+00 | **1.01E-11** | 1.00E+00 | **8.72E-06** |
| Migration of phagocytes | 1.00E+00 | **1.84E-09** | 1.00E+00 | **1.13E-04** |
| Quantity of lymphoid cells | 1.00E+00 | **6.51E-14** | 1.00E+00 | **4.10E-05** |
| Recruitment of lymphocytes | 1.00E+00 | **8.47E-10** | 1.00E+00 | **1.56E-05** |
| Recruitment of mononuclear leukocytes | 1.00E+00 | **1.57E-11** | 1.00E+00 | **7.08E-06** |

**P*-value: Benjamini Hochberg adjusted *P*-value. Bold values: $P < 0.05$.

Table S24. ITE-dysregulated common disease-associated biological function in rat placentas and HUVECs.*

| | *P*-value [Female Placentas] | *P*-value [Male Placentas] | *P*-value [Female HUVECs] | *P*-value [Male HUVECs] |
|---|---|---|---|---|
| Activation of hepatic stellate cells | **1.92E-03** | **2.42E-02** | **5.30E-03** | **4.02E-03** |
| Acute myocardial infarction | **1.23E-03** | **3.65E-02** | **4.51E-03** | **3.98E-05** |
| Chronic renal failure | **4.92E-03** | **8.31E-03** | **2.52E-03** | **5.32E-05** |
| Cirrhosis | **8.75E-03** | **4.66E-05** | **8.03E-03** | **5.65E-03** |
| Damage of kidney | **1.17E-02** | **2.30E-04** | **4.78E-03** | **7.61E-04** |
| End stage renal disease | **5.16E-03** | **1.20E-02** | **8.03E-03** | **7.61E-04** |
| Enlargement of heart | **2.28E-08** | **6.37E-08** | **1.84E-04** | **1.70E-04** |
| Fibrosis of heart | **1.85E-04** | **1.80E-05** | **1.17E-02** | **7.61E-04** |
| Fibrosis of kidney | **1.14E-03** | **1.97E-02** | **4.78E-03** | **3.42E-04** |
| Fibrosis of liver | **7.40E-03** | **4.06E-06** | **3.05E-03** | **1.70E-04** |
| Hepatocellular carcinoma | **1.11E-03** | **2.76E-05** | **2.23E-02** | **1.63E-03** |
| Hypertrophy of cardiac muscle | **1.08E-03** | **1.98E-03** | **1.66E-02** | **1.52E-03** |
| Hypertrophy of cardiomyocytes | **2.95E-03** | **3.46E-03** | **2.32E-02** | **3.94E-03** |
| Hypertrophy of heart | **2.15E-08** | **3.35E-08** | **1.63E-04** | **3.98E-05** |
| Hypertrophy of heart cells | **2.38E-03** | **3.75E-03** | **4.78E-03** | **1.28E-03** |
| Hypoplasia of renal medulla | **3.20E-02** | **4.84E-02** | **3.40E-02** | **1.55E-02** |
| Increased Levels of Hematocrit | **1.75E-03** | **4.25E-03** | **2.22E-02** | **5.19E-03** |
| Inflammation of liver | **2.75E-02** | **1.97E-09** | **3.05E-03** | **4.30E-04** |
| Injury of kidney | **3.19E-03** | **2.89E-04** | **5.81E-03** | **1.61E-03** |
| Left ventricular dysfunction | **1.42E-08** | **2.46E-11** | **2.21E-02** | **2.63E-03** |
| Liver cancer | **4.22E-17** | **5.84E-20** | **1.53E-02** | **3.13E-03** |
| Liver carcinoma | **2.56E-17** | **5.84E-20** | **1.96E-02** | **4.02E-03** |
| Liver Regeneration | **1.99E-02** | **1.52E-02** | **2.07E-03** | **6.88E-04** |
| Liver tumor | **2.56E-17** | **8.34E-21** | **5.05E-03** | **2.92E-03** |
| Multi vessel coronary artery disease | **1.99E-02** | **3.01E-02** | **2.92E-02** | **1.30E-02** |
| Myocardial infarction | **1.32E-10** | **2.46E-11** | **6.43E-03** | **2.31E-04** |
| Proliferation of liver cells | **4.42E-02** | **1.01E-02** | **7.85E-03** | **2.12E-03** |
| Proximal tubular toxicity | **3.21E-04** | **1.10E-06** | **2.21E-02** | **2.81E-02** |
| ST-elevation myocardial infarction | **6.33E-04** | **1.09E-03** | **1.71E-02** | **3.05E-03** |
| Stenosis of aortic valve | **3.62E-02** | **1.56E-04** | **3.05E-03** | **2.31E-04** |
| Ventricular hypertrophy | **4.39E-05** | **9.28E-06** | **2.21E-02** | **3.66E-03** |
| Glomerulosclerosis | **7.47E-05** | **1.50E-02** | **4.78E-03** | 1.00E+00 |
| Arrhythmia | **2.63E-05** | **4.05E-08** | 1.15E-01 | **2.08E-02** |
| Atrial fibrillation | **2.60E-06** | **1.72E-07** | 7.53E-02 | **1.67E-02** |
| Chemotaxis of hepatic stellate cells | **1.99E-02** | **3.40E-02** | 1.00E+00 | **2.90E-02** |
| Chronic hepatitis C | **1.68E-02** | **2.01E-04** | 1.00E+00 | **4.26E-03** |
| Cirrhosis of liver | **3.15E-02** | **3.88E-03** | 1.00E+00 | **8.94E-03** |
| Congenital heart disease | **1.06E-06** | **7.54E-04** | 1.16E-01 | **6.44E-03** |
| Congestive heart failure | **2.32E-09** | **2.98E-10** | 1.74E-01 | **1.27E-02** |
| Coronary artery disease | **2.97E-06** | **1.08E-08** | 1.58E-01 | **1.03E-02** |

| | | | | |
|---|---|---|---|---|
| Disorder of coronary artery | **1.32E-10** | **7.32E-12** | 7.11E-02 | **8.98E-03** |
| Enlargement of left ventricle | **1.75E-06** | **2.65E-07** | 1.00E+00 | **1.46E-02** |
| Failure of heart | **1.42E-08** | **1.33E-10** | 5.79E-02 | **6.86E-03** |
| Hepatic steatosis | **1.24E-03** | **7.76E-07** | 4.15E-01 | **1.47E-02** |
| Hypertrophy of left ventricle | **9.15E-04** | **2.83E-03** | 1.12E-01 | **3.88E-02** |
| Left ventricular dilation | **1.47E-04** | **1.70E-07** | 2.84E-01 | **2.05E-02** |
| Recurrent myocardial infarction | **2.50E-02** | **4.41E-02** | 6.71E-02 | **3.15E-02** |
| Susceptibility to atrioventricular septal defect type 1 | **3.20E-02** | **4.84E-02** | 1.00E+00 | **1.55E-02** |
| Acute kidney injury | **4.11E-02** | 1.09E-01 | **1.08E-02** | **7.48E-03** |
| Proliferation of hepatic stellate cells | **2.50E-02** | 1.13E-01 | **1.56E-02** | **2.56E-03** |
| Spontaneous coronary artery dissection | **7.40E-03** | 9.02E-02 | **4.71E-02** | **2.08E-02** |
| Familial pulmonary hypertension | **1.99E-02** | 9.97E-02 | **2.21E-02** | 1.00E+00 |
| Idiopathic and/or familial pulmonary arterial hypertension | **2.96E-02** | 1.30E-01 | **3.22E-02** | 1.00E+00 |
| Atrophy of renal cortex | **3.62E-02** | 1.73E-01 | 7.44E-02 | **3.64E-02** |
| Cell proliferation of kidney cell lines | **3.62E-02** | 9.83E-02 | 2.87E-01 | **2.70E-02** |
| Interstitial fibrosis of kidney | **5.43E-03** | 1.48E-01 | 6.69E-02 | **3.00E-03** |
| Stenosis of pulmonary artery | **4.66E-02** | 7.00E-02 | 1.00E+00 | **1.82E-02** |
| Thrombosis of atrium | **4.42E-02** | 7.35E-02 | 7.82E-02 | **3.88E-02** |
| Acute myocarditis | 6.06E-02 | **9.26E-04** | **4.71E-02** | **2.08E-02** |
| Alcoholic hepatitis | 1.99E-01 | **4.72E-04** | **8.03E-03** | **2.02E-05** |
| Apoptosis of kidney cells | 1.43E-01 | **3.34E-02** | **3.80E-02** | **8.78E-03** |
| Calcification of aortic valve | 1.00E+00 | **5.74E-03** | **4.51E-03** | **6.44E-03** |
| Cell death of kidney cells | 1.66E-01 | **2.65E-02** | **2.21E-02** | **4.53E-02** |
| Chronic hepatitis | 8.37E-02 | **1.86E-03** | **2.21E-02** | **3.46E-03** |
| Decreased secretion of albumin | 1.99E-01 | **4.84E-02** | **3.40E-02** | **4.14E-04** |
| Glomerulonephritis | 1.46E-01 | **6.37E-03** | **2.21E-02** | **2.08E-02** |
| Liver Damage | 1.92E-01 | **2.30E-04** | **5.78E-04** | **1.70E-04** |
| Nephritis | 9.06E-02 | **1.29E-02** | **2.21E-02** | **1.21E-02** |
| Obstruction of bile duct | 1.99E-01 | **4.84E-02** | **3.40E-02** | **1.55E-02** |
| Primary biliary cirrhosis | 9.46E-02 | **1.45E-02** | **3.05E-03** | **6.56E-03** |
| Proliferation of hepatocytes | 1.33E-01 | **4.35E-02** | **2.92E-02** | **4.69E-03** |
| Pulmonary embolism | 8.37E-02 | **5.74E-03** | **2.21E-02** | **6.44E-03** |
| Right ventricular hypertrophy | 6.06E-02 | **5.75E-03** | **2.92E-02** | **9.05E-03** |
| Bleeding of liver | 9.81E-02 | **3.12E-02** | **4.78E-03** | 1.00E+00 |
| Alcoholic steatohepatitis | 3.80E-01 | **7.52E-03** | 1.00E+00 | **1.61E-03** |
| Calcific aortic valve stenosis | 1.00E+00 | **3.40E-02** | 6.39E-02 | **2.90E-02** |
| Proliferation of mesangial cells | 1.65E-01 | **4.84E-02** | 5.29E-02 | **2.31E-04** |
| Severe alcoholic hepatitis | 3.43E-01 | **1.20E-02** | 1.00E+00 | **5.65E-03** |
| Steatohepatitis | 2.40E-01 | **6.35E-03** | 1.00E+00 | **6.50E-03** |
| Viral hepatitis | 1.08E-01 | **2.48E-04** | 9.15E-02 | **5.19E-03** |

**P*-value: Benjamini Hochberg adjusted *P*-value. Bold *P*-value: < 0.05.

Table S25. ITE-dysregulated common canonical pathways in rat placenta and HUVECs.*

| | *P*-value [Female Rat Placentas] | *P*-value [Male Rat Placentas] | *P*-value [Female HUVECs] | *P*-value [Male HUVECs] |
|---|---|---|---|---|
| Breast Cancer Regulation by Stathmin1 | **2.86E-05** | **5.26E-05** | **5.56E-03** | **1.92E-02** |
| CREB Signaling in Neurons | **5.06E-05** | **5.93E-06** | **6.16E-03** | **1.93E-02** |
| Hepatic Fibrosis Signaling Pathway | **3.19E-02** | **4.10E-04** | **1.03E-02** | **2.04E-02** |
| Phagosome Formation | **7.13E-04** | **4.72E-08** | **5.29E-03** | **2.98E-02** |
| GP6 Signaling Pathway | **3.33E-02** | **4.06E-03** | **4.22E-02** | 3.42E-01 |
| Acute Phase Response Signaling | 4.18E-01 | **2.76E-03** | 4.13E-01 | **4.59E-02** |
| Aryl Hydrocarbon Receptor Signaling | 1.82E-01 | **1.47E-02** | **2.71E-03** | **6.57E-05** |
| Atherosclerosis Signaling | 1.21E-01 | **1.89E-07** | **1.31E-04** | **2.55E-05** |
| Differential Regulation of Cytokine Production in Macrophages and T Helper Cells by IL-17A and IL-17F | 7.75E-01 | **1.28E-02** | **4.24E-02** | **2.01E-04** |
| Xenobiotic Metabolism AHR Signaling Pathway | 2.60E-01 | **2.97E-02** | **1.58E-03** | **4.77E-06** |
| HIF1α Signaling | 1.48E-01 | **1.12E-02** | 2.68E-01 | **1.50E-02** |
| HMGB1 Signaling | 2.77E-01 | **1.64E-03** | 6.04E-02 | **7.07E-03** |
| IL-17 Signaling | 2.93E-01 | **3.08E-03** | 7.95E-02 | **1.06E-02** |
| LXR/RXR Activation | 1.88E-01 | **2.03E-04** | 3.40E-01 | **2.25E-02** |
| Neuroinflammation Signaling Pathway | 1.00E+00 | **2.33E-03** | 9.90E-02 | **3.77E-02** |
| Role of Hypercytokinemia/ hyperchemokinemia in the Pathogenesis of Influenza | 2.60E-01 | **4.80E-05** | 4.72E-01 | **6.36E-05** |
| Role of Macrophages, Fibroblasts and Endothelial Cells in Rheumatoid Arthritis | 2.14E-01 | **2.41E-03** | 9.90E-02 | **3.77E-02** |
| Role of MAPK Signaling in Inhibiting the Pathogenesis of Influenza | 5.57E-01 | **5.96E-03** | 5.65E-02 | **6.30E-04** |
| Xenobiotic Metabolism Signaling | 2.14E-01 | **2.57E-02** | 7.65E-02 | **9.03E-04** |

**P*-value: Benjamini Hochberg adjusted *P*-value. CREB: cAMP response element-binding protein; IL-17: Interleukin-17; GP6: glycoprotein VI platelet; HIF1α: Hypoxia -inducible factor 1-alpha; HMBG1: High mobility group box 1 protein; LXR/RXR, liver X receptor/retinoid X receptor; MAPK: mitogen-activated protein kinase, AHR: acryl hydrocarbon receptor. Bold *P*-value: < 0.05.

Table S26. ITE-dysregulated common gene networks in HUVECs and rat placentas.*

| | Female HUVECs | | Male HUVECs | | Female Rat Placentas | | Male Rat Placentas | |
|---|---|---|---|---|---|---|---|---|
| Gene network | *P*-value (n) | Target Genes | *P*-value (n) | Target Genes | *P*-value (n) | Target Genes | *P*-value (n) | Target Genes |
| AHR-regulated genes | 8.46E-08 (13) | ACP5, AHRR, CCR7, COL1A2, COL3A1, CYP1A1, CYP1B1, IDO1, IL1B, NPTX1, PTGS2, SERPINB2, TIPARP | 1.15E-12 (9) | AHRR, CCR7, COL14A1, COL1A2, CXCL10, CYP1A1, CYP1B1, IL1B, IL6, NPTX1, NQO1, PTGS2, SERPINB2, TIPARP | 1.93E-09 (48) | ACTA2, AHRR, ATOH8, Ccl2, CCNA2, CCND1, CD28, CDC25B, CDC25C, CHST2, COL11A1, COL13A1, CXCL10, CYP1A1, CYP1B1, DCLK3, EBF1, EGR2, FASLG, FBLN5, FBN2, FGF9, FOXP3, GRAP2, HAVCR2, HEY2, IGFBP1, IL1R1, IL34, IL7R, KLF2, NOTCH3, NPTX1, NQO1, NXPE2, OLFML1, PDGFB, PKLR, PMP22, PTGS2, RBP1, SEMA3A, SERPINA12, SLAMF8, SLC16A10, STEAP4, TGFBI, TUBA8 | 2.75E-13 (64) | ACTA2, AHRR, ALOX5, ATOH8, C1QB, C1QC, CARD11, CBR1, Ccl2, CCL20, CCL5, CCNA2, CD14, CDC25B, CDKN2A, CHI3L1, CHST2, COL11A1, COL13A1, COL14A1, CXCL10, CXCL11, CXCL2, Cxcl9, CYP1A1, CYP1B1, DCLK3, EBF1, EGR2, FASLG, FBLN5, FBN2, FGF9, FGFR2, FOXP3, GDF15, GRAP2, HAVCR2, HEY2, Hsd3b4, IL1B, IL1R1, IL34, IL6, IL7R, MYH7, NOTCH3, NPTX1, NQO1, OLFML1, PDGFB, PKLR, PMP22, PTGS2, RBP1, SDR9C7, SEMA3A, SERPINA12, SLAMF8, SLC16A10, SPP1, TFF1, TGFBI, THBS1 |
| CBX5-regulated genes | 1.87E-03 (4) | CEMIP, CYP1B1, TACSTD2, TNFRSF6B | 2.24E-03 (6) | IL1B, IL6, MMP1, PTGS2, STC1, TIPARP | 2.59E-02 (10) | CYP1B1, GRHL3, H2AC18/H2AC19, HBB, HOXA13, ICAM2, PRSS23, TACSTD2, TGFBI, TIMP4 | 1.39E-03 (15) | CKM, CYP1B1, GRHL3, HBB, HOXA13, ICAM2, MAL2, PRR15L, PRSS23, TACSTD2, TGFBI, TIMP4, TM4SF1, TRIM31, TSPAN13 |
| HIF1A-regulated genes | 6.93E-03 (7) | CCR7, CEMIP, CYP1A1, IL1B, MAFF, MMP1, PTGS2 | 1.52E-05 (5) | AHRR, CCR7, IL1B, IL6, ISG15 | 1.70E-05 (42) | ABCC8, ACE, ACTA2, AGT, AGTR1, BMP2, CAV1, CAVIN1, CAVIN2, Ccl2, CCL2, CCL3L3, Ccl7, CCND1, | 1.15E-07 (56) | ABCC8, ACE, ACTA2, AGTR1, CAV1, CAVIN2, Ccl2, CCL2, CCL3L3, CCL4, CCL5, Ccl7, CD274, CD69, CDKN2A, CLDN1, CORO1A, COX4I2, CXCL10, CXCL2, Cxcl3, CXCL3, Cxcl9, |

| | | | | | | | | |
|---|---|---|---|---|---|---|---|---|
| | | | | | | CLDN1, COX4I2, CXCL10, Cxcl3, CXCL3, CXCR4, CYP1A1, ENG, GATA1, GCK, HIF3A, HOXA13, ID1, IGFBP3, IL23R, ITGA2B, LEP, MMP13, NRARP, NTRK2, PDGFB, PDHA1, PFKL, PTGS2, SLC16A3, SLC16A4, TBX5, TNFRSF11B | | CXCR4, CYP1A1, DNAJC5B, FUT7, GATA1, GCK, HIF3A, HOXA13, ID1, IFI16, IGFBP3, IL1B, IL6, LEP, MMP13, MMP9, MYH7, NRARP, NTRK2, PDGFB, PDHA1, PRKAR2B, PTGS2, RAB11FIP4, SFRP1, SLC16A3, SLC16A4, TBX5, TFRC, THBS1, TIMP1, TNFSF10, VCAM1 |
| NKX2-3-regulated genes | 6.37E-05 (7) | F2RL3, GALNT15, IDO1, MAMDC2, POSTN, PTGS2, TIPARP | 1.45E-09 (12) | COL1A2, CSF3, CXCL10, CYP1B1, F3, IL1B, IL6, LPAR1, MMP1, NQO1, PTGS2, SERPINB2 | 1.32E-09 (32) | ACTA2, AOC3, BMP2, BMP4, CAVIN2, CLEC1A, F2RL3, FBN2, FLRT2, GBP1, GHR, H2AC18/H2AC19, HEY1, HEY2, HFE, HNRNPA0, IL7R, KLF2, MFNG, MTURN, MYH11, PLA1A, PLCB1, PLEKHA4, PRSS12, PTGS2, SEMA3A, SNCAIP, SRPX2, TCIM, TMEM100, TRPC6 | 9.81E-07 (31) | ACTA2, AOC3, BMP4, CAVIN2, CLEC1A, CXCL16, F2RL3, FBN2, GBP1, GBP2, GDF15, GHR, HEY2, IFNB1, IL7R, MFNG, MMRN1, MTURN, MYH11, PLA1A, PLCB1, PRSS12, PSMB9, PTGS2, SEMA3A, SNCAIP, SRPX2, TCIM, TMEM100, UBA7, VCAM1 |

| | | | | | | | | |
|---|---|---|---|---|---|---|---|---|
| RELA-regulated genes | 7.22E-06 (11) | AHRR, CCR7, COL1A2, F3, IL1B, JUNB, LUM, LYN, MMP1, PTGS2, SELE | 6.03E-10 (12) | AHRR, CCR7, COL1A2, CXCL10, ELN, F3, IL1B, IL6, ISG15, MMP1, PTGS2, SELE | 4.50E-03 (32) | ACTA2, AGT, AHRR, BCL2A1, BMP2, CAMP, CAV1, Ccl2, CCL2, CCND1, CD28, CXCL10, CXCL3, CXCR4, CYP2C9, EGR1, ELN, FASLG, FOXP3, GBP1, ID1, IL1RN, IL7R, NEFH, ORAI1, PDGFB, PTGES, PTGS2, TNFRSF11B, TP63, TRPC6, UBD | 9.80E-09 (56) | ACTA2, AHRR, ALPK2, BCL2A1, BEX1, CAMP, CAV1, Ccl2, CCL2, CCL20, CCL5, CD14, CD274, CD69, CDKN2A, CHI3L1, COL2A1, CXCL10, CXCL11, CXCL2, CXCL3, CXCR4, CYBB, CYP2C9, DNMT3B, EGR1, ELN, FABP6, FASLG, FOXP3, GBP1, GDF15, ID1, IFIT1B, IFNB1, IL1B, IL1RN, IL6, IL7R, ISG15, MMP9, NEFH, NOD2, PDGFB, PSMB9, PTGES, PTGS2, RGS1, SMOC2, SPI1, TACR1, TP63, TREM1, TWIST2, UBD, VCAM1 |

*A gene network with Z score > |2| is predicted to be altered; *P*-value: Benjamini Hochberg adjusted *P*-value; n = number of target genes.

**Table S27. Primary antibodies used in Western blot analysis.***

A. For rat placentas

| Antibody | Vendor | Cat. # | Dilution |
|---|---|---|---|
| CD31 | R & D systems (Minneapolis, MN, USA) | AF3628 | 1:500 |
| Caspase-3 | Cell Signaling Technology (Beverly, MA, USA) | 9662 | 1:1000 |
| Cleaved Caspase-3 | Cell Signaling Technology | 9661 | 1:1000 |
| β-actin | Cell signaling Technology | 4967 | 1:4000 |
| GAPDH | Cell Signaling Technology | 5174 | 1:3000 |

B. For verification of phosphoproteomics data in HUVECs

| Antibody | Vendor | Cat. # | Dilution |
|---|---|---|---|
| pLMNA a/c (Ser390) | Millipore-Sigma (Burlington, MA, USA) | ABT1388 | 1:1000 |
| tLMNA a/c | Millipore-Sigma | SAB4200236 | 1:5000 |
| pHP1γ/CBX3 (Ser93) | ABcam | ab45270 | 1:1000 |
| tHP1γ/CBX3 | Cell Signaling Technology | 2619 | 1:1000 |
| pEEF2 (Thr 57) | Bioss Antibodies (Woburn, MA, USA) | bs-3486R | 1:500 & 1:1000 |
| Mouse GAPDH | Abnova (Taipei, Taiwan) | H00002597-M01 | 1:10000 |

*p: phospho; t: total.  LMNAa/c: lamin a/c; HP1γ/CBX3:  heterochromatin protein 1 γ/chromobox homolog 3; EEF2: eukaryotic translation elongation factor 2; Ser: serine; Thr: threonine.

**Table S28. ITE-dysregulated phosphosites in HUVECs at 4 hr.***

| Master Protein Accessions | Modifications in Master Proteins | Master Protein Descriptions | Fold change ≥ 1.2 (ITE/DMSO) | Adjusted *P* -Value |
|---|---|---|---|---|
| P35658 | 1xPhospho [T/S] | Nuclear pore complex protein Nup214 GN=NUP214 | 3.0 | 1.06E-05 |
| P51397 | 1xPhospho [S51] | Death-associated protein 1 GN=DAP | 2.6 | **1.16E-14** |
| V9GYM8 | 1xPhospho [S] | Guanine nucleotide exchange factor H1 GN=ARHGEF2 | 2.4 | 6.69E-09 |
| P46821 | 2xPhospho [S1396; S1400] | Microtubule-associated protein 1B GN=MAP1B | 2.0 | **2.73E-11** |
| Q09666 | 1xPhospho [S135] | Neuroblast differentiation-associated protein AHNAK GN=AHNAK | 1.9 | 1.16E-14 |
| E5RGF9 | 1xPhospho [S] | Protein FAM114A2 (Fragment) GN=FAM114A2 | 1.9 | 4.34E-02 |
| Q13283 | 1xPhospho [S] | Ras GTPase-activating protein-binding protein 1 GN=G3BP1 | 1.9 | **2.60E-08** |
| Q6NZI2 | 1xPhospho [S167] | Caveolae-associated protein 1 GN=CAVIN1 | 1.9 | 1.16E-14 |
| Q15149-4 | 1xPhospho [S4269]; | Isoform 4 of Plectin GN=PLEC | 1.8 | 3.32E-04 |
| Q15149 | 1xPhospho [S4406] | | | |
| A0A087WVQ6 | 1xPhospho [T398] | Clathrin heavy chain OS=Homo sapiens GN=CLTC | 0.9 | 9.53E-01 |
| Q96CV9 | 1xPhospho [T/S] | Optineurin GN=OPTN | 1.8 | 9.74E-05 |
| V9GYM8 | 1xPhospho [S/T] | Guanine nucleotide exchange factor H1 GN=ARHGEF2 | 1.8 | 2.39E-04 |
| Q9UDY2 | 1xPhospho [S163] | Tight junction protein ZO-2 GN=TJP2 | 1.7 | **3.17E-06** |
| P62753 | 1xPhospho [S240] | 40S ribosomal protein S6 GN=RPS6 | 1.6 | **1.02E-10** |
| P16989 | 1xPhospho [S134]; | Y-box-binding protein 3 GN=YBX3 | 1.6 | 3.04E-05 |
| P67809 | 1xPhospho [S102] | Y-box-binding protein 1 GN=YBX1 | | |
| Q14204 | 1xPhospho [S4368] | Cytoplasmic dynein 1 heavy chain 1 GN=DYNC1H1 | 1.6 | 5.75E-05 |
| Q15942 | 1xPhospho [S143] | Zyxin GN=ZYX | 1.6 | 2.24E-05 |
| Q96PK6 | 1xPhospho [S627] | RNA-binding protein 14 GN=RBM14 | 1.6 | 1.32E-10 |
| Q9H6F5 | 1xPhospho [S217] | Coiled-coil domain-containing protein 86 GN=CCDC86 | 1.6 | 2.37E-05 |
| B7Z3B9 | 1xPhospho [S448] | GRB2-associated-binding protein 1 GN=GAB1 | 1.6 | 3.24E-05 |
| Q9H2D6 | 1xPhospho [S/T] | TRIO and F-actin-binding protein GN=TRIOBP | 1.6 | 1.71E-02 |
| M0R300 | 1xPhospho [T1348] | Unconventional myosin-IXb (Fragment) GN=MYO9B | 1.6 | 1.45E-04 |
| P48681 | 1xPhospho [T1309] | Nestin GN=NES | 1.6 | 3.50E-08 |
| P61978-2 | 1xPhospho [S/T] | Isoform 2 of Heterogeneous nuclear ribonucleoprotein K GN=HNRNPK | 1.6 | 2.79E-04 |
| P46821 | 1xPhospho [S1265] | Microtubule-associated protein 1B GN=MAP1B | 1.6 | 5.13E-10 |
| P29966 | 1xPhospho [S101] | Myristoylated alanine-rich C-kinase substrate GN=MARCKS | 1.5 | 9.33E-09 |
| Q9UEY8 | 1xPhospho [S423] | Gamma-adducin GN=ADD3 | 1.5 | 3.16E-02 |
| Q13185 | 1xPhospho [S93] | Chromobox protein homolog 3 GN=CBX3 | 1.5 | **2.81E-04** |
| Q7Z417 | 1xPhospho [S629] | Nuclear fragile X mental retardation-interacting protein 2 GN=NUFIP2 | 1.5 | **1.59E-04** |
| Q13409-2 | 1xPhospho [S81] | Isoform 2B of Cytoplasmic dynein 1 intermediate chain 2 GN=DYNC1I2 | 1.5 | 9.44E-04 |
| P19634 | 1xPhospho [S703] | Sodium/hydrogen exchanger 1 GN=SLC9A1 | 1.5 | 1.06E-05 |
| Q09161 | 1xPhospho [T/S] | Nuclear cap-binding protein subunit 1 GN=NCBP1 | 1.5 | 8.45E-05 |
| Q32MZ4-4 | 1xPhospho [S/T] | Isoform 4 of Leucine-rich repeat flightless-interacting protein 1 GN=LRRFIP1 | 1.5 | 5.74E-04 |
| P02545 | 1xPhospho [S390] | Prelamin-A/C GN=LMNA | 1.5 | 4.64E-07 |
| P04792 | 1xPhospho [S] | Heat shock protein beta-1 GN=HSPB1 | 1.5 | 1.01E-05 |

| | | | | |
|---|---|---|---|---|
| Q92609 | 1xPhospho [T/S] | TBC1 domain family member 5 GN=TBC1D5 | 1.5 | 3.04E-06 |
| P48681 | 1xPhospho [S680] | Nestin GN=NES | 1.5 | **1.31E-05** |
| Q3KQU3 | 1xPhospho [S753] | MAP7 domain-containing protein 1 GN=MAP7D1 | 1.5 | **7.96E-06** |
| M0R2K3 | 1xPhospho [T/S] | BRISC and BRCA1-A complex member 1 (Fragment) GN=BABAM1 | 1.5 | 2.09E-03 |
| Q09666 | 1xPhospho [S5110] | Neuroblast differentiation-associated protein AHNAK GN=AHNAK | 1.5 | 4.78E-03 |
| Q9NTI5 | 1xPhospho [S] | Sister chromatid cohesion protein PDS5 homolog B GN=PDS5B | 1.5 | 2.21E-02 |
| E7EX73 | 1xPhospho [S1068] | Eukaryotic translation initiation factor 4 gamma 1 GN=EIF4G1 | 1.5 | **1.08E-05** |
| Q15691 | 1xPhospho [S155] | Microtubule-associated protein RP/EB family member 1 GN=MAPRE1 | 1.5 | **1.94E-05** |
| P27824 | 1xPhospho [T/S] | Calnexin GN=CANX | 1.5 | 2.52E-04 |
| Q9UKV3-5 | 1xPhospho [S991] | Isoform 4 of Apoptotic chromatin condensation inducer in the nucleus GN=ACIN1 | 1.4 | 1.14E-05 |
| Q07352 | 1xPhospho [S331] | mRNA decay activator protein ZFP36L1 GN=ZFP36L1 | 1.4 | 5.47E-04 |
| A0A0A0MQW0 | 1xPhospho [S17] | Myelin expression factor 2 GN=MYEF2 | 1.4 | 1.10E-02 |
| Q5T4S7 | 1xPhospho [S] | E3 ubiquitin-protein ligase UBR4 GN=UBR4 | 1.4 | 2.52E-04 |
| Q8TD19 | 1xPhospho [S] | Serine/threonine-protein kinase Nek9 GN=NEK9 | 1.4 | **5.16E-03** |
| P08670 | 1xPhospho [S72] | Vimentin GN=VIM | 1.4 | 1.14E-05 |
| Q9Y6G9 | 1xPhospho [T/S] | Cytoplasmic dynein 1 light intermediate chain 1 GN=DYNC1LI1 | 1.4 | 1.63E-02 |
| P17812 | 1xPhospho [S] | CTP synthase 1 GN=CTPS1 | 1.4 | 4.64E-03 |
| Q96TA1 | 2xPhospho [S692; S] | Protein Niban 2 GN=NIBAN2 | 1.4 | 5.09E-07 |
| Q3B726 | 1xPhospho [S316] | DNA-directed RNA polymerase I subunit RPA43 GN=POLR1F | 1.4 | 5.68E-03 |
| O00443 | 1xPhospho [S259] | Phosphatidylinositol 4-phosphate 3-kinase C2 domain-containing subunit alpha GN=PIK3C2A | 1.4 | 1.35E-02 |
| P60174-1 | 1xPhospho [S21] | Isoform 2 of Triosephosphate isomerase GN=TPI1 | 1.4 | 6.70E-03 |
| A0A087X0X3 | 1xPhospho [S618] | Heterogeneous nuclear ribonucleoprotein M GN=HNRNPM | 1.4 | 2.73E-02 |
| P05455 | 1xPhospho [S366] | Lupus La protein GN=SSB | 1.4 | 1.12E-02 |
| Q4KMP7 | 1xPhospho [S132] | TBC1 domain family member 10B GN=TBC1D10B | 1.4 | 2.73E-02 |
| Q15149-4 | 1xPhospho [T/S] | Isoform 4 of Plectin GN=PLEC | 1.4 | 1.02E-02 |
| Q02952 | 1xPhospho [S/T] | A-kinase anchor protein 12 GN=AKAP12 | 1.3 | **4.72E-05** |
| Q8TC07-3 | 1xPhospho [S213] | Isoform 3 of TBC1 domain family member 15 GN=TBC1D15 | 1.3 | 2.94E-02 |
| Q02952 | 1xPhospho [S696] | A-kinase anchor protein 12 GN=AKAP12 | 1.3 | **2.57E-02** |
| Q99961 | 1xPhospho [S288] | Endophilin-A2 GN=SH3GL1 | 1.3 | 1.14E-02 |
| O14617-5 | 1xPhospho [S658] | Isoform 5 of AP-3 complex subunit delta-1 GN=AP3D1 | 1.3 | 4.70E-02 |
| Q63ZY3 | 1xPhospho [S540] | KN motif and ankyrin repeat domain-containing protein 2 GN=KANK2 | 1.3 | 5.09E-03 |
| A0A0G2JHC2 | 1xPhospho [S213] | PPP1R18 GN=PPP1R18 | 1.3 | 1.13E-02 |
| A0A1B0GV73 | 1xPhospho [S712] | Rho GTPase-activating protein 21 GN=ARHGAP21 | 1.3 | 2.15E-02 |
| Q8NDI1 | 1xPhospho [S1058] | EH domain-binding protein 1 GN=EHBP1 | 1.3 | 7.57E-04 |
| Q9Y2D5 | 1xPhospho [S152] | A-kinase anchor protein 2 GN=AKAP2 | 1.3 | 1.91E-02 |
| P46821 | 1xPhospho [S] | Microtubule-associated protein 1B GN=MAP1B | 1.3 | **1.99E-02** |
| Q14247 | 2xPhospho [T401; T] | Src substrate cortactin GN=CTTN | 1.3 | 1.25E-02 |
| Q6PKG0 | 1xPhospho [S774] | La-related protein 1 GN=LARP1 | 1.3 | 7.45E-03 |
| Q9H1B7 | 1xPhospho [S547] | Probable E3 ubiquitin-protein ligase IRF2BPL GN=IRF2BPL | 1.3 | 3.29E-02 |

| | | | | |
|---|---|---|---|---|
| J3QRU1 | 1xPhospho [T432] | Tyrosine-protein kinase GN=YES1 | 1.3 | 5.10E-02 |
| Q9UKV3-5 | 1xPhospho [S710] | Isoform 4 of Apoptotic chromatin condensation inducer in the nucleus GN=ACIN1 | 1.3 | 2.43E-02 |
| P02765 | 1xPhospho [S138] | Alpha-2-HS-glycoprotein GN=AHSG | 1.3 | 1.50E-03 |
| Q13263 | 1xPhospho [S473] | Transcription intermediary factor 1-beta GN=TRIM28 | 1.3 | 4.28E-02 |
| Q02952 | 1xPhospho [S1331] | A-kinase anchor protein 12 GN=AKAP12 | 1.3 | **2.49E-02** |
| Q14195-2 | 1xPhospho [S/T] | Isoform LCRMP-4 of Dihydropyrimidinase-related protein 3 GN=DPYSL3 | 1.2 | 3.61E-02 |
| O75369-8 | 1xPhospho [S1474] | Isoform 8 of Filamin-B GN=FLNB | 1.2 | 3.86E-02 |
| H7BXR3 | 1xPhospho [S419] | Sorbin and SH3 domain-containing protein 2 (Fragment) GN=SORBS2 | 1.2 | 5.15E-02 |
| P10644 | 1xPhospho [S83] | cAMP-dependent protein kinase type I-alpha regulatory subunit GN=PRKAR1A | 1.2 | 2.34E-02 |
| P35579 | 1xPhospho [S1943] | Myosin-9 GN=MYH9 | 1.2 | 4.14E-02 |
| A0A0B4J1S8 | 1xPhospho [T292] | Phosphatidylinositol 4-kinase beta GN=PI4KB | 0.8 | 5.41E-02 |
| G3XAC6 | 1xPhospho [S136] | RNA-binding protein 39 GN=RBM39 | 0.8 | 4.28E-02 |
| P06733 | 1xPhospho [T/S] | Alpha-enolase GN=ENO1 | 0.8 | 3.80E-02 |
| A0A0U1RRM6 | 1xPhospho [T721] | Protein enabled homolog GN=ENAH | 0.8 | 3.03E-02 |
| E7EX73 | 1xPhospho [S/T] | Eukaryotic translation initiation factor 4 gamma 1 GN=EIF4G1 | 0.8 | 2.73E-02 |
| Q9UDY2 | 1xPhospho [S441] | Tight junction protein ZO-2 GN=TJP2 | 0.7 | 9.44E-03 |
| A0A0D9SEY1 | 1xPhospho [S793] | Mitogen-activated protein kinase kinase kinase kinase 4 GN=MAP4K4 | 0.7 | 1.91E-02 |
| P07900 | 1xPhospho [S231] | Heat shock protein HSP 90-alpha GN=HSP90AA1 | 0.7 | 3.63E-04 |
| P18669 | 1xPhospho [S118] | Phosphoglycerate mutase 1GN=PGAM1 | 0.7 | 3.93E-02 |
| Q9ULC0 | 1xPhospho [S237] | Endomucin GN=EMCN | 0.7 | 2.67E-02 |
| P13861 | 1xPhospho [S99] | cAMP-dependent protein kinase type II-alpha regulatory subunit GN=PRKAR2A | 0.7 | 2.23E-02 |
| O95810 | 2xPhospho [S288; S293] | Caveolae-associated protein 2 GN=CAVIN2 | 0.7 | 5.87E-05 |
| Q99733 | 1xPhospho [T/S] | Nucleosome assembly protein 1-like 4 GN=NAP1L4 | 0.6 | 2.76E-02 |
| Q53EL6 | 1xPhospho [S94] | Programmed cell death protein 4 GN=PDCD4 | 0.6 | **5.90E-05** |
| Q01082 | 1xPhospho [T] | Spectrin beta chain, non-erythrocytic 1 GN=SPTBN1 | 0.6 | 6.17E-05 |
| A9Z1X7 | 1xPhospho [S722] | Serine/arginine repetitive matrix protein 1 GN=SRRM1 | 0.6 | 7.49E-03 |
| Q13177 | 1xPhospho [S197] | Serine/threonine-protein kinase PAK 2 GN=PAK2 | 0.6 | 4.22E-06 |
| A0A087WTD7 | 1xPhospho [S1325] | A-kinase anchor protein 13 GN=AKAP13 | 0.6 | 3.80E-02 |
| P08670 | 1xPhospho [S430] | Vimentin GN=VIM | 0.6 | **4.85E-06** |
| Q14847 | 1xPhospho [S146] | LIM and SH3 domain protein 1 GN=LASP1 | 0.5 | 3.56E-02 |

*HUVECs were treated with 1 µM ITE or DMSO for 4 hr. Phosphoproteins in lysates were enriched and subjected to the phosphoteomics analysis. S: serine; T: threonine; S/T: serine and threonine. GN: gene name. Bold *P* values: commonly regulated phosphosites at 4 and 24 hr. proteins n = 4 cell preparations (2 female and 2 male cells). *P*- values < 0.05.

**Table S29. ITE-dysregualted phosphosites in HUVECs at 24 hr.***

| Master Protein | Modifications in Master Proteins | Master Protein Descriptions | Fold change ≥ 1.2 (ITE/DMSO) | Adjusted *P* -Value |
|---|---|---|---|---|
| Q12834 | 1xPhospho [S/T] | Cell division cycle protein 20 homolog GN=CDC20 | 3.6 | 1.81E-14 |
| A0A3B3IT92 | 1xPhospho [S120] | DNA helicase GN=MCM4 | 2.5 | 1.81E-14 |
| P52292 | 1xPhospho [T73] | Importin subunit alpha-1 GN=KPNA2 | 2.4 | 1.81E-14 |
| P16401 | 1xPhospho [T138] | Histone H1.5 GN=H1-5 | 2.4 | 1.81E-14 |
| P52292 | 1xPhospho [S62] | Importin subunit alpha-1 GN=KPNA2 | 2.1 | 8.47E-10 |
| Q5H909 | 1xPhospho [S229] | Melanoma-associated antigen D2 GN=MAGED2 | 2.1 | 1.81E-14 |
| P19338 | 1xPhospho [S67] | Nucleolin GN=NCL | 2.1 | 1.21E-12 |
| E9PBY3 | 1xPhospho [S103] | N-acetylgalactosaminyltransferase 7 GN=GALNT7 | 2.0 | 2.51E-08 |
| Q9Y490 | 1xPhospho [S423] | Talin-1 GN=TLN1 | 2.0 | 1.81E-14 |
| A0A096LNZ0 | 1xPhospho [S79] | AP2-associated protein kinase 1 (Fragment) GN=AAK1 | 1.9 | 6.95E-03 |
| A0A0C4DGB5 | 1xPhospho [S289] | Calpain inhibitor GN=CAST | 1.9 | 4.99E-07 |
| P08670 | 1xPhospho [S459] | Vimentin GN=VIM PE=1 SV=4 | 1.8 | 1.81E-14 |
| A0A286YFD6 | 1xPhospho [S20] | Ribonucleoside-diphosphate reductase GN=RRM2 | 1.8 | 8.47E-10 |
| P48681 | 1xPhospho [S325] | Nestin GN=NES PE=1 SV=2 | 1.8 | 3.36E-12 |
| P46821 | 2xPhospho [S1396;S1400] | Microtubule-associated protein 1B GN=MAP1B | 1.8 | **6.56E-10** |
| K7EPZ6 | 1xPhospho [S] | Methyl-CpG-binding domain protein 1 (Fragment) GN=MBD1 PE=1 SV=1 | 1.8 | 9.93E-06 |
| Q15149-4<br>Q15149 | 1xPhospho [S1298];<br>1xPhospho[S1435] | Isoform 4 of Plectin GN=PLEC | 1.8 | 3.34E-12 |
| P16403 | 1xPhospho [T146] | Histone H1.2 GN=H1-2 | 1.8 | 7.16E-09 |
| Q02952 | 1xPhospho [S/T] | A-kinase anchor protein 12 GN=AKAP12 | 1.8 | 9.55E-08 |
| O95997 | 1xPhospho [S165] | Securin OX=9606 GN=PTTG1 | 1.8 | 1.93E-08 |
| Q9UDY2 | 1xPhospho [S163] | Tight junction protein ZO-2 GN=TJP2 | 1.7 | **4.98E-05** |
| F8VQE1 | 1xPhospho [S202] | LIM domain and actin-binding protein 1 GN=LIMA1 | 1.7 | 2.01E-05 |
| P46821 | 1xPhospho [S614] | Microtubule-associated protein 1B GN=MAP1B | 1.7 | 2.05E-07 |
| O15211 | 1xPhospho [S/T] | Ral guanine nucleotide dissociation stimulator-like 2 GN=RGL2 | 1.7 | 1.11E-04 |
| A0A669KAX4 | 1xPhospho [S/T] | Cullin-4B GN=CUL4B | 1.7 | 7.16E-09 |
| P46821 | 1xPhospho [T1067] | Microtubule-associated protein 1B GN=MAP1B | 1.7 | 4.39E-09 |
| E7ETY2 | 1xPhospho [S381];<br>1xPhospho[S446] | Treacle protein GN=TCOF1 | 1.7 | 1.08E-06 |

| | | | | |
|---|---|---|---|---|
| O76021 | 1xPhospho [S427] | Ribosomal L1 domain-containing protein 1 GN=RSL1D1 | 1.7 | 9.08E-08 |
| Q15942 | 1xPhospho [S281] | Zyxin GN=ZYX | 1.7 | 5.02E-07 |
| E7ETY2 | 1xPhospho [S] | Treacle protein GN=TCOF1 | 1.6 | 4.08E-09 |
| P29966 | 1xPhospho [S] | Myristoylated alanine-rich C-kinase substrate GN=MARCKS | 1.6 | 6.16E-06 |
| P13639 | 1xPhospho [T57] | Elongation factor 2 GN=EEF2 | 1.6 | 4.23E-11 |
| Q02952 | 1xPhospho [S627] | A-kinase anchor protein 12 GN=AKAP12 | 1.6 | 8.99E-07 |
| Q02952 | 1xPhospho [S1331] | A-kinase anchor protein 12 GN=AKAP12 | 1.6 | **6.96E-06** |
| P27816 | 1xPhospho [T571] | Microtubule-associated protein 4 GN=MAP4 | 1.6 | 2.24E-06 |
| E7EX73 | 1xPhospho [S1068] | Eukaryotic translation initiation factor 4 gamma 1 GN=EIF4G1 | 1.6 | **5.91E-08** |
| Q7Z417 | 1xPhospho [S629] | Nuclear fragile X mental retardation-interacting protein 2 GN=NUFIP2 | 1.6 | **9.81E-06** |
| I1E4Y6 | 1xPhospho [S258] | GRB10-interacting GYF protein 2 GN=GIGYF2 | 1.6 | 1.05E-03 |
| P27816 | 1xPhospho [S696] | Microtubule-associated protein 4 GN=MAP4 | 1.6 | 1.18E-07 |
| P17096 | 1xPhospho [T53] | High mobility group protein HMG-I/HMG-Y GN=HMGA1 | 1.6 | 6.86E-06 |
| Q13185 | 1xPhospho [S93] | Chromobox protein homolog 3 GN=CBX3 | 1.6 | **2.01E-05** |
| P06748 | 1xPhospho [S254] | Nucleophosmin OX=9606 GN=NPM1 | 1.6 | 6.09E-08 |
| P16989 | 1xPhospho [S293] | Y-box-binding protein 3 GN=YBX3 | 1.6 | 9.62E-09 |
| P46821 | 1xPhospho [T1633] | Microtubule-associated protein 1B GN=MAP1B | 1.6 | 4.10E-03 |
| Q01518-2<br>Q01518 | 1xPhospho [S34];<br>1xPhospho[S34] | Isoform 2 of Adenylyl cyclase-associated protein 1 GN=CAP1<br>Adenylyl cyclase-associated protein 1 GN=CAP1 | 1.5 | 7.57E-05 |
| O75369-8<br>O75369 | 1xPhospho [S1536];<br>1xPhospho[S1505] | Isoform 8 of Filamin-B GN=FLNB; Filamin-B GN=FLNB | 1.5 | 1.51E-04 |
| Q14669 | 1xPhospho [T/S] | E3 ubiquitin-protein ligase TRIP12 GN=TRIP12 | 1.5 | 2.85E-03 |
| Q9H223 | 1xPhospho [T/S] | EH domain-containing protein 4 GN=EHD4 | 1.5 | 3.14E-06 |
| P27816 | 1xPhospho [S358];<br>1xPhospho[S384] | Microtubule-associated protein 4 GN=MAP4 | 1.5 | 2.44E-04 |
| A6NNK5 | 1xPhospho [S1633] | TP53-binding protein 1 GN=TP53BP1 | 1.5 | 1.06E-06 |
| Q9UK59 | 1xPhospho [S514] | Lariat debranching enzyme GN=DBR1 | 1.5 | 5.09E-03 |
| A0A5F9ZHW5 | 1xPhospho [S612] | Ataxin-2 GN=ATXN2 | 1.5 | 6.52E-05 |
| P48681 | 1xPhospho [S905] | Nestin GN=NES | 1.5 | 8.11E-06 |
| Q15555 | 1xPhospho [S228] | Microtubule-associated protein RP/EB family member 2 GN=MAPRE2 | 1.5 | 5.48E-04 |
| A0A2R8YG42 | 1xPhospho [S228] | Rho guanine nucleotide exchange factor 7 GN=ARHGEF7 | 1.5 | 7.86E-03 |
| Q02952 | 1xPhospho [S371] | A-kinase anchor protein 12 GN=AKAP12 | 1.5 | 2.20E-05 |

| | | | | |
|---|---|---|---|---|
| Q02952 | 1xPhospho [S248] | A-kinase anchor protein 12 GN=AKAP12 | 1.5 | 1.78E-02 |
| A0A1C7CYX9 | 1xPhospho [S27] | Dihydropyrimidinase-related protein 2 GN=DPYSL2 | 1.5 | 7.41E-05 |
| Q3KQU3 | 1xPhospho [S753] | MAP7 domain-containing protein 1 GN=MAP7D1 | 1.5 | **2.19E-05** |
| P18887 | 1xPhospho [S241] | DNA repair protein XRCC1 GN=XRCC1 | 1.5 | 5.54E-05 |
| P42695 | 1xPhospho [S1384] | Condensin-2 complex subunit D3 GN=NCAPD3 | 1.5 | 1.37E-02 |
| Q8NC51 | 1xPhospho [S234] | Plasminogen activator inhibitor 1 RNA-binding protein GN=SERBP1 | 1.5 | 3.56E-04 |
| Q6UN15 | 1xPhospho [S/T] | Pre-mRNA 3'-end-processing factor FIP1 GN=FIP1L1 | 1.5 | 2.17E-02 |
| Q9Y6M7-13 | 1xPhospho [S242] | Isoform 13 of Sodium bicarbonate cotransporter 3 GN=SLC4A7 | 1.4 | 1.05E-04 |
| P20700 | 1xPhospho [S23] | Lamin-B1 GN=LMNB1 | 1.4 | 3.26E-04 |
| P22234 | 1xPhospho [S27] | Multifunctional protein ADE2 GN=PAICS | 1.4 | 3.31E-04 |
| E7EX17 | 1xPhospho [S467] | Eukaryotic translation initiation factor 4B GN=EIF4B | 1.4 | 9.17E-04 |
| Q9NR30 | 1xPhospho [S71] | Nucleolar RNA helicase 2 GN=DDX21 | 1.4 | 6.26E-04 |
| P16949 | 2xPhospho [S16;S25] | Stathmin GN=STMN1 | 1.4 | 1.25E-02 |
| Q02952 | 1xPhospho [S1295] | A-kinase anchor protein 12 GN=AKAP12 | 1.4 | 2.06E-04 |
| P78316 | 1xPhospho [S96] | Nucleolar protein 14 GN=NOP14 | 1.4 | 4.21E-03 |
| Q13283 | 1xPhospho [S] | Ras GTPase-activating protein-binding protein 1 GN=G3BP1 | 1.4 | **9.17E-03** |
| Q02952 | 1xPhospho [S696] | A-kinase anchor protein 12 GN=AKAP12 | 1.4 | **9.17E-04** |
| P17096 | 1xPhospho [S36] | High mobility group protein HMG-I/HMG-Y GN=HMGA1 | 1.4 | 6.25E-03 |
| P24941 | 1xPhospho [T14] | Cyclin-dependent kinase 2 GN=CDK2 | 1.4 | 1.48E-03 |
| Q9H1E3 | 1xPhospho [S181] | Nuclear ubiquitous casein and cyclin-dependent kinase substrate 1 GN=NUCKS1 | 1.4 | 1.15E-02 |
| Q96TA1 | 2xPhospho [S665;S681] | Protein Niban 2 GN=NIBAN2 | 1.4 | 4.98E-02 |
| P31323 | 1xPhospho [S114] | cAMP-dependent protein kinase type II-beta regulatory subunit GN=PRKAR2B | 1.4 | 1.26E-04 |
| Q14157-1 | 1xPhospho [S] | Isoform 2 of Ubiquitin-associated protein 2-like GN=UBAP2L | 1.4 | 1.25E-02 |
| A0A5F9UP49 | 1xPhospho [S350] | 45 kDa calcium-binding protein GN=SDF4 | 1.4 | 2.89E-03 |
| Q02952 | 1xPhospho [S651] | A-kinase anchor protein 12 GN=AKAP12 | 1.4 | 3.18E-03 |
| E7EX17 | 1xPhospho [T/S] | Eukaryotic translation initiation factor 4B GN=EIF4B | 1.4 | 5.43E-04 |
| P46821 | 1xPhospho [S2072] | Microtubule-associated protein 1B GN=MAP1B | 1.4 | 6.03E-03 |
| Q9P2E9 | 1xPhospho [T275] | Ribosome-binding protein 1 GN=RRBP1 | 1.4 | 1.01E-03 |
| Q16555 | 2xPhospho [S518;S522]; | Dihydropyrimidinase-related protein 2 GN=DPYSL2 | 1.4 | 7.05E-05 |
| A0A1C7CYX9 | 2xPhospho[S623;S627] | Dihydropyrimidinase-related protein 2 GN=DPYSL2 | | |
| O14578 | 1xPhospho [S1971] | Citron Rho-interacting kinase GN=CIT | 1.4 | 1.75E-03 |

| | | | | |
|---|---|---|---|---|
| A0A3B3IS71 | 1xPhospho [T356] | Retinoblastoma-associated protein GN=RB1 | 1.4 | 3.17E-04 |
| Q02952 | 1xPhospho [S] | A-kinase anchor protein 12 GN=AKAP12 | 1.4 | 8.26E-03 |
| O00429 | 1xPhospho [S616] | Dynamin-1-like protein GN=DNM1L | 1.4 | 8.82E-03 |
| P46013 | 1xPhospho [S357] | Proliferation marker protein Ki-67 GN=MKI67 | 1.4 | 1.39E-02 |
| E7ETY2 | 1xPhospho [S906] | Treacle protein GN=TCOF1 | 1.3 | 4.00E-03 |
| P52926 | 1xPhospho [S44] | High mobility group protein HMGI-C GN=HMGA2 | 1.3 | 4.53E-03 |
| P36507 | 1xPhospho [T394] | Dual specificity mitogen-activated protein kinase kinase 2 GN=MAP2K2 | 1.3 | 1.33E-03 |
| P48681 | 1xPhospho [S680] | Nestin GN=NES | 1.3 | **2.17E-02** |
| Q8TD19 | 1xPhospho [S] | Serine/threonine-protein kinase Nek9 GN=NEK9 | 1.3 | **3.49E-02** |
| P51397 | 1xPhospho [S51] | Death-associated protein 1 GN=DAP | 1.3 | **3.44E-02** |
| A0A0A0MRM9 | 1xPhospho [S547] | Nucleolar and coiled-body phosphoprotein 1 (Fragment) GN=NOLC1 | 1.3 | 2.08E-02 |
| Q15691 | 1xPhospho [S155] | Microtubule-associated protein RP/EB family member 1 GN=MAPRE1 | 1.3 | **3.21E-03** |
| Q9Y2J2-2 | 1xPhospho [S740] | Isoform 2 of Band 4.1-like protein 3 GN=EPB41L3 | 1.3 | 9.13E-03 |
| Q92522 | 1xPhospho [S31] | Histone H1.10 GN=H1-10 | 1.3 | 1.79E-02 |
| P46821 | 1xPhospho [S] | Microtubule-associated protein 1B GN=MAP1B | 1.3 | **1.60E-02** |
| E7ETY2 | 1xPhospho [S1378] | Treacle protein GN=TCOF1 | 1.3 | 1.44E-02 |
| E7ETY2 | 1xPhospho [S381] | Treacle protein GN=TCOF1 | 1.3 | 3.46E-02 |
| P17812 | 1xPhospho [S575] | CTP synthase 1 GN=CTPS1 | 1.3 | 5.03E-02 |
| Q9NRH2 | 1xPhospho [S351] | SNF-related serine/threonine-protein kinase GN=SNRK | 1.3 | 4.26E-02 |
| P13639 | 1xPhospho [T57] | Elongation factor 2 GN=EEF2 | 1.3 | 4.79E-02 |
| P62753 | 1xPhospho [S240] | 40S ribosomal protein S6 GN=RPS6 | 1.3 | **1.81E-02** |
| P20700 | 1xPhospho [S393]; | Lamin-B1 GN=LMNB1 | 1.3 | 2.42E-02 |
| Q03252 | 1xPhospho[S407] | Lamin-B2 GN=LMNB2 | | |
| P46100 | 1xPhospho [S1527] | Transcriptional regulator ATRX GN=ATRX | 1.3 | 4.72E-02 |
| P16284 | 1xPhospho [T731] | Platelet endothelial cell adhesion molecule GN=PECAM1 | 0.8 | 3.27E-02 |
| Q9P266 | 1xPhospho [S1044] | Junctional protein associated with coronary artery disease G N=JCAD | 0.7 | 1.50E-02 |
| J3KTL2 | 1xPhospho [S199] | Serine/arginine-rich splicing factor 1 GN=SRSF1 | 0.7 | 1.26E-02 |
| C9JYJ6 | 1xPhospho [S551] | Filamin A-interacting protein 1-like (Fragment) GN=FILIP1L | 0.7 | 6.37E-03 |
| Q96CW1 | 1xPhospho [T156] | AP-2 complex subunit mu GN=AP2M1 | 0.7 | 4.96E-02 |
| Q8TED9 | 1xPhospho [S329] | Actin filament-associated protein 1-like 1 GN=AFAP1L1 | 0.7 | 1.65E-02 |
| C9JZR2 | 1xPhospho [S] | Catenin delta-1 GN=CTNND1 | 0.7 | 1.06E-02 |

| | | | | |
|---|---|---|---|---|
| Q13177 | 1xPhospho [S55] | Serine/threonine-protein kinase PAK 2 GN=PAK2 | 0.7 | 2.93E-02 |
| C9JZR2 | 1xPhospho [S252] | Catenin delta-1 GN=CTNND1 | 0.7 | 1.24E-02 |
| Q12846 | 1xPhospho [S117]; | Syntaxin-4 GN=STX4 | 0.7 | 1.81E-02 |
| C9JFM5 | 1xPhospho[S169] | Syntaxin-4 (Fragment) GN=STX4 | | |
| P08670 | 1xPhospho [S430] | Vimentin GN=VIM | 0.7 | **9.77E-05** |
| C9JZR2 | 1xPhospho [S268] | Catenin delta-1 GN=CTNND1 | 0.7 | 8.56E-04 |
| P08670 | 1xPhospho [S/T] | Vimentin GN=VIM | 0.7 | 2.77E-03 |
| Q9P266 | 1xPhospho [S885] | Junctional protein associated with coronary artery disease GN=JCAD | 0.7 | 9.97E-03 |
| P04792 | 1xPhospho [S15] | Heat shock protein beta-1 GN=HSPB1 | 0.7 | 1.28E-04 |
| Q16181 | 1xPhospho [T426] | Septin-7 GN=SEPTIN7 | 0.6 | 1.87E-03 |
| C9JJX6 | 1xPhospho [T202] | Armadillo repeat protein deleted in velo-cardio-facial syndrome GN=ARVCF | 0.6 | 2.11E-02 |
| P14923 | 1xPhospho [S665] | Junction plakoglobin GN=JUP | 0.6 | 4.66E-07 |
| Q53EL6 | 1xPhospho [S94] | Programmed cell death protein 4 GN=PDCD4 | 0.6 | **4.99E-07** |
| P17096 | 1xPhospho [S] | High mobility group protein HMG-I/HMG-Y GN=HMGA1 | 0.6 | 1.34E-06 |
| C9JJX6 | 1xPhospho [T769] | Armadillo repeat protein deleted in velo-cardio-facial syndrome GN=ARVCF | 0.6 | 7.05E-07 |
| C9JJX6 | 1xPhospho [S140] | Armadillo repeat protein deleted in velo-cardio-facial syndrome GN=ARVCF | 0.6 | 1.38E-07 |
| P35221 | 1xPhospho [S655] | Catenin alpha-1 GN=CTNNA1 | 0.6 | 1.84E-05 |
| P08670 | 1xPhospho [T426] | Vimentin GN=VIM | 0.5 | 3.60E-05 |
| A0A494C115 | 1xPhospho [S192] | Aquaporin-1 (Fragment) GN=AQP1 | 0.5 | 1.88E-08 |
| I3L1L3 | 1xPhospho [S] | Myb-binding protein 1A (Fragment) GN=MYBBP1A | 0.4 | 1.18E-02 |
| A9Z1X7 | 1xPhospho [S569] | Serine/arginine repetitive matrix protein 1 GN=SRRM1 | 0.4 | 1.02E-05 |

*HUVECs were treated with 1 μM ITE or DMSO for 24 hr. Phosphoproteins in lysates were enriched and subjected to the phosphoteomics analysis. S: serine; T: threonine; S/T: serine and threonine. GN: gene name. Bold P values: commonly regulated phosphosites. S/T: serine and threonine. Bold: commonly regulated phosphosites at 4 and 24 hr. proteins n = 4 cell preparations (2 female and 2 male cells). *P* -values < 0.05.

**Table S30. ITE-dysregulated total protein in HUVECs at 4 and 24 hr.***

| Master Protein Accessions | Master Protein Descriptions | Fold change ≥ 1.2 at 4 hr (ITE/DMSO) | Adjusted *P*-Value at 4 hr | Fold change ≥ 1.2 at 24 hr (ITE/DMSO) | A *P*-Va |
|---|---|---|---|---|---|
| P35579 | Myosin-9 OS=Homo sapiens OX=9606 GN=MYH9 PE=1 SV=4 | 1.014 | 0.904500837 | 0.998 | 0.9 |
| Q09666 | Neuroblast differentiation-associated protein AHNAK OS=Homo sapiens OX=9606 GN=AHNAK PE=1 SV=2 | 1.054 | 0.765235202 | 0.98 | 0.9 |
| O75369 | Filamin-B OS=Homo sapiens OX=9606 GN=FLNB PE=1 SV=2 | 0.982 | 0.750023665 | 0.942 | 0.7 |
| Q9Y490 | Talin-1 OS=Homo sapiens OX=9606 GN=TLN1 PE=1 SV=3 | 0.987 | 0.889482529 | 1.015 | 0.8 |
| P21333 | Filamin-A OS=Homo sapiens OX=9606 GN=FLNA PE=1 SV=4 | 0.994 | 0.866265907 | 0.992 | 0.9 |
| Q15149 | Plectin OS=Homo sapiens OX=9606 GN=PLEC PE=1 SV=3 | 1.042 | 0.858816078 | 0.996 | 0.9 |
| Q15149-3 | Isoform 3 of Plectin OS=Homo sapiens OX=9606 GN=PLEC | 0.912 | 0.835808354 | 0.886 | 0.7 |
| Q14204 | Cytoplasmic dynein 1 heavy chain 1 OS=Homo sapiens OX=9606 GN=DYNC1H1 PE=1 SV=5 | 1.04 | 0.765235202 | 0.98 | 0.7 |
| A0A087WVQ6 | Clathrin heavy chain OS=Homo sapiens OX=9606 GN=CLTC PE=1 SV=1 | 1.004 | 0.953739985 | 0.978 | 0.7 |
| P08670 | Vimentin OS=Homo sapiens OX=9606 GN=VIM PE=1 SV=4 | 1.218 | 0.765235202 | 0.941 | 0.[illegible] |
| P46940 | Ras GTPase-activating-like protein IQGAP1 OS=Homo sapiens OX=9606 GN=IQGAP1 PE=1 SV=1 | 0.973 | 0.765235202 | 0.962 | 0.7 |
| P13639 | Elongation factor 2 OS=Homo sapiens OX=9606 GN=EEF2 PE=1 SV=4 | 1.003 | 0.902016292 | 1.107 | 0.7 |
| P10809 | 60 kDa heat shock protein, mitochondrial OS=Homo sapiens OX=9606 GN=HSPD1 PE=1 SV=2 | 0.981 | 0.766667967 | 0.961 | 0.7 |
| P49327 | Fatty acid synthase OS=Homo sapiens OX=9606 GN=FASN PE=1 SV=3 | 0.98 | 0.827085425 | 1.115 | 0.7 |
| P55072 | Transitional endoplasmic reticulum ATPase OS=Homo sapiens OX=9606 GN=VCP PE=1 SV=4 | 1.006 | 0.977023215 | 1.016 | 0.7 |
| P11142 | Heat shock cognate 71 kDa protein OS=Homo sapiens OX=9606 GN=HSPA8 PE=1 SV=1 | 1.002 | 0.92949421 | 1.133 | 0.7 |
| P22314 | Ubiquitin-like modifier-activating enzyme 1 OS=Homo sapiens OX=9606 GN=UBA1 PE=1 SV=3 | 0.994 | 0.987622091 | 1.022 | 0.8 |
| P04406 | Glyceraldehyde-3-phosphate dehydrogenase OS=Homo sapiens OX=9606 GN=GAPDH PE=1 SV=3 | 0.977 | 0.750023665 | 1.062 | 0.8 |
| P26038 | Moesin OS=Homo sapiens OX=9606 GN=MSN PE=1 SV=3 | 1.002 | 0.942678074 | 1.01 | 0.7 |
| P04275 | von Willebrand factor OS=Homo sapiens OX=9606 GN=VWF PE=1 SV=4 | 0.794 | 0.765235202 | 0.855 | 0.7 |
| O43707 | Alpha-actinin-4 OS=Homo sapiens OX=9606 GN=ACTN4 PE=1 SV=2 | 0.969 | 0.750023665 | 0.887 | 0.7 |
| Q13813-3 | Isoform 3 of Spectrin alpha chain, non-erythrocytic 1 OS=Homo sapiens OX=9606 GN=SPTAN1 | 0.982 | 0.921087329 | 0.881 | 0.7 |
| P08238 | Heat shock protein HSP 90-beta OS=Homo sapiens OX=9606 GN=HSP90AB1 PE=1 SV=4 | 0.999 | 0.924631778 | 1.163 | 0.7 |
| P06733 | Alpha-enolase OS=Homo sapiens OX=9606 GN=ENO1 PE=1 SV=2 | 0.985 | 0.886561233 | 1.021 | 0.8 |
| P14618 | Pyruvate kinase PKM OS=Homo sapiens OX=9606 GN=PKM PE=1 SV=4 | 0.947 | 0.750023665 | 1.042 | 0.7 |
| P21980 | Protein-glutamine gamma-glutamyltransferase 2 OS=Homo sapiens OX=9606 GN=TGM2 PE=1 SV=2 | 0.98 | 0.765825736 | 1.013 | 0.9 |
| P68104 | Elongation factor 1-alpha 1 OS=Homo sapiens OX=9606 GN=EEF1A1 PE=1 SV=1 | 1.061 | 0.765235202 | 1.177 | 0.7 |
| P60709 | Actin, cytoplasmic 1 OS=Homo sapiens OX=9606 GN=ACTB PE=1 SV=1 | 0.991 | 0.925929947 | 0.981 | 0.7 |
| P78527 | DNA-dependent protein kinase catalytic subunit OS=Homo sapiens OX=9606 GN=PRKDC PE=1 SV=3 | 1.026 | 0.886561233 | 0.929 | 0.7 |
| P98160 | Basement membrane-specific heparan sulfate proteoglycan core protein OS=Homo sapiens OX=9606 GN=HSPG2 PE=1 SV=4 | 0.964 | 0.778727018 | 0.946 | 0.8 |
| P06576 | ATP synthase subunit beta, mitochondrial OS=Homo sapiens OX=9606 GN=ATP5F1B PE=1 SV=3 | 0.952 | 0.882454148 | 0.862 | 0.7 |
| P11021 | Endoplasmic reticulum chaperone BiP OS=Homo sapiens OX=9606 GN=HSPA5 PE=1 SV=2 | 1.012 | 0.898725987 | 1.072 | 0.7 |
| P18206 | Vinculin OS=Homo sapiens OX=9606 GN=VCL PE=1 SV=4 | 0.944 | 0.765235202 | 0.952 | 0.7 |
| P07900 | Heat shock protein HSP 90-alpha OS=Homo sapiens OX=9606 GN=HSP90AA1 PE=1 SV=5 | 0.979 | 0.837893552 | 1.069 | 0.7 |
| P12814 | Alpha-actinin-1 OS=Homo sapiens OX=9606 GN=ACTN1 PE=1 SV=2 | 0.978 | 0.869995963 | 0.944 | 0.7 |
| Q71U36 | Tubulin alpha-1A chain OS=Homo sapiens OX=9606 GN=TUBA1A PE=1 SV=1 | 1.003 | 0.882454148 | 1.003 | 0.9 |
| P29401 | Transketolase OS=Homo sapiens OX=9606 GN=TKT PE=1 SV=3 | 0.983 | 0.95980123 | 0.955 | 0.7 |
| P14618-2 | Isoform M1 of Pyruvate kinase PKM OS=Homo sapiens OX=9606 GN=PKM | 0.899 | 0.750023665 | 0.992 | 0.9 |
| F5H5D3 | Tubulin alpha chain OS=Homo sapiens OX=9606 GN=TUBA1C PE=1 SV=1 | 1.023 | 0.983953845 | 1.074 | 0.8 |
| P14625 | Endoplasmin OS=Homo sapiens OX=9606 GN=HSP90B1 PE=1 SV=1 | 1.001 | 0.913279108 | 1.065 | 0.7 |
| Q14764 | Major vault protein OS=Homo sapiens OX=9606 GN=MVP PE=1 SV=4 | 1.023 | 0.917008272 | 1.158 | 0.7 |
| P07437 | Tubulin beta chain OS=Homo sapiens OX=9606 GN=TUBB PE=1 SV=2 | 1.023 | 0.984200071 | 1.058 | 0.8 |
| Q14315 | Filamin-C OS=Homo sapiens OX=9606 GN=FLNC PE=1 SV=3 | 0.998 | 0.944284199 | 0.984 | 0.8 |
| P07355 | Annexin A2 OS=Homo sapiens OX=9606 GN=ANXA2 PE=1 SV=2 | 0.968 | 0.750023665 | 0.979 | 0.7 |
| P53396 | ATP-citrate synthase OS=Homo sapiens OX=9606 GN=ACLY PE=1 SV=3 | 1.002 | 0.860909277 | 1.073 | 0.7 |
| P15144 | Aminopeptidase N OS=Homo sapiens OX=9606 GN=ANPEP PE=1 SV=4 | 0.926 | 0.765235202 | 0.946 | 0.7 |
| P60842 | Eukaryotic initiation factor 4A-I OS=Homo sapiens OX=9606 GN=EIF4A1 PE=1 SV=1 | 0.992 | 0.991211283 | 1.073 | 0.7 |
| P68371 | Tubulin beta-4B chain OS=Homo sapiens OX=9606 GN=TUBB4B PE=1 SV=1 | 1.025 | 0.859267486 | 1.028 | 0.9 |
| P38646 | Stress-70 protein, mitochondrial OS=Homo sapiens OX=9606 GN=HSPA9 PE=1 SV=2 | 0.969 | 0.750023665 | 0.995 | 0.7 |
| P07996 | Thrombospondin-1 OS=Homo sapiens OX=9606 GN=THBS1 PE=1 SV=2 | 1.24 | 0.756935672 | 1.366 | 0.7 |
| Q16881 | Thioredoxin reductase 1, cytoplasmic OS=Homo sapiens OX=9606 GN=TXNRD1 PE=1 SV=3 | 0.994 | 0.956825234 | 1.072 | 0.7 |
| P68366 | Tubulin alpha-4A chain OS=Homo sapiens OX=9606 GN=TUBA4A PE=1 SV=1 | 1.027 | 0.765235202 | 1.026 | 0.[illegible] |
| P48643 | T-complex protein 1 subunit epsilon OS=Homo sapiens OX=9606 GN=CCT5 PE=1 SV=1 | 0.992 | 0.968322408 | 1.035 | 0.[illegible] |
| P13010 | X-ray repair cross-complementing protein 5 OS=Homo sapiens OX=9606 GN=XRCC5 PE=1 SV=3 | 1.034 | 0.889450577 | 0.993 | 0.7 |
| P35580 | Myosin-10 OS=Homo sapiens OX=9606 GN=MYH10 PE=1 SV=3 | 1 | 0.876510992 | 0.89 | 0.7 |
| P78371 | T-complex protein 1 subunit beta OS=Homo sapiens OX=9606 GN=CCT2 PE=1 SV=4 | 0.969 | 0.750023665 | 1.007 | 0.8 |
| P02545 | Prelamin-A/C OS=Homo sapiens OX=9606 GN=LMNA PE=1 SV=1 | 1.007 | 0.917008272 | 0.887 | 0.7 |
| Q01082 | Spectrin beta chain, non-erythrocytic 1 OS=Homo sapiens OX=9606 GN=SPTBN1 PE=1 SV=2 | 0.985 | 0.921087329 | 0.892 | 0.7 |
| Q01813 | ATP-dependent 6-phosphofructokinase, platelet type OS=Homo sapiens OX=9606 GN=PFKP PE=1 SV=2 | 1.004 | 0.886561233 | 1.012 | 0.9 |
| P25705 | ATP synthase subunit alpha, mitochondrial OS=Homo sapiens OX=9606 GN=ATP5F1A PE=1 SV=1 | 0.966 | 0.915804503 | 0.912 | 0.7 |
| Q02952 | A-kinase anchor protein 12 OS=Homo sapiens OX=9606 GN=AKAP12 PE=1 SV=4 | 1.114 | 0.750023665 | 1.352 | 0.7 |
| P13797 | Plastin-3 OS=Homo sapiens OX=9606 GN=PLS3 PE=1 SV=4 | 0.987 | 0.765235202 | 0.938 | 0.7 |
| P52209 | 6-phosphogluconate dehydrogenase, decarboxylating OS=Homo sapiens OX=9606 GN=PGD PE=1 SV=3 | 0.972 | 0.894570507 | 1.045 | 0.[illegible] |
| Q13885 | Tubulin beta-2A chain OS=Homo sapiens OX=9606 GN=TUBB2A PE=1 SV=1 | 0.946 | 0.839845191 | 0.969 | 0.9 |
| Q9NZM1 | Myoferlin OS=Homo sapiens OX=9606 GN=MYOF PE=1 SV=1 | 0.996 | 0.887776944 | 0.934 | 0.7 |
| P08133 | Annexin A6 OS=Homo sapiens OX=9606 GN=ANXA6 PE=1 SV=3 | 0.969 | 0.750023665 | 0.965 | 0.7 |
| P17987 | T-complex protein 1 subunit alpha OS=Homo sapiens OX=9606 GN=TCP1 PE=1 SV=1 | 0.977 | 0.765235202 | 1.007 | 0.7 |
| P50395 | Rab GDP dissociation inhibitor beta OS=Homo sapiens OX=9606 GN=GDI2 PE=1 SV=2 | 0.972 | 0.780884711 | 1.003 | 0.9 |
| P07237 | Protein disulfide-isomerase OS=Homo sapiens OX=9606 GN=P4HB PE=1 SV=3 | 0.977 | 0.904500837 | 1.06 | 0.[illegible] |
| Q08211 | ATP-dependent RNA helicase A OS=Homo sapiens OX=9606 GN=DHX9 PE=1 SV=4 | 0.986 | 0.94475926 | 0.988 | 0.8 |
| P48681 | Nestin OS=Homo sapiens OX=9606 GN=NES PE=1 SV=2 | 1.041 | 0.889482529 | 0.996 | 0.9 |
| P13667 | Protein disulfide-isomerase A4 OS=Homo sapiens OX=9606 GN=PDIA4 PE=1 SV=2 | 0.991 | 0.937953124 | 1.025 | 0.7 |
| P34932 | Heat shock 70 kDa protein 4 OS=Homo sapiens OX=9606 GN=HSPA4 PE=1 SV=4 | 0.999 | 0.948876456 | 1.003 | 0.7 |
| P49588 | Alanine--tRNA ligase, cytoplasmic OS=Homo sapiens OX=9606 GN=AARS1 PE=1 SV=2 | 0.995 | 0.944385799 | 1.053 | 0.7 |
| P12956 | X-ray repair cross-complementing protein 6 OS=Homo sapiens OX=9606 GN=XRCC6 PE=1 SV=2 | 1.018 | 0.816716243 | 1.01 | 0.8 |
| Q14974 | Importin subunit beta-1 OS=Homo sapiens OX=9606 GN=KPNB1 PE=1 SV=2 | 0.962 | 0.764871997 | 1.007 | 0.8 |
| Q9NZN4 | EH domain-containing protein 2 OS=Homo sapiens OX=9606 GN=EHD2 PE=1 SV=2 | 1.058 | 0.840158726 | 0.862 | 0.7 |
| A0A0G2JIW1 | Heat shock 70 kDa protein 1B OS=Homo sapiens OX=9606 GN=HSPA1B PE=1 SV=1 | 0.96 | 0.750023665 | 1.006 | 0.9 |
| P04075 | Fructose-bisphosphate aldolase A OS=Homo sapiens OX=9606 GN=ALDOA PE=1 SV=2 | 1.004 | 0.924992773 | 1.065 | 0.7 |
| Q00839 | Heterogeneous nuclear ribonucleoprotein U OS=Homo sapiens OX=9606 GN=HNRNPU PE=1 SV=6 | 1.041 | 0.750023665 | 1.032 | 0.7 |
| P42704 | Leucine-rich PPR motif-containing protein, mitochondrial OS=Homo sapiens OX=9606 GN=LRPPRC PE=1 SV=3 | 0.988 | 0.766667967 | 1.002 | 0.8 |
| P27797 | Calreticulin OS=Homo sapiens OX=9606 GN=CALR PE=1 SV=1 | 0.935 | 0.765235202 | 0.99 | 0.9 |
| P00558 | Phosphoglycerate kinase 1 OS=Homo sapiens OX=9606 GN=PGK1 PE=1 SV=3 | 0.971 | 0.703015003 | 0.998 | 0.9 |
| A0A1W2PPS1 | Heterogeneous nuclear ribonucleoprotein U OS=Homo sapiens OX=9606 GN=HNRNPU PE=1 SV=1 | 1.278 | 0.750023665 | 1.044 | 0.7 |
| A0A2R8Y5S7 | Radixin OS=Homo sapiens OX=9606 GN=RDX PE=1 SV=1 | 1.002 | 0.985763982 | 0.946 | 0.7 |
| P61978 | Heterogeneous nuclear ribonucleoprotein K OS=Homo sapiens OX=9606 GN=HNRNPK PE=1 SV=1 | 1.036 | 0.918702104 | 0.934 | 0.7 |
| P15311 | Ezrin OS=Homo sapiens OX=9606 GN=EZR PE=1 SV=4 | 0.961 | 0.750023665 | 0.905 | 0.7 |
| P37802 | Transgelin-2 OS=Homo sapiens OX=9606 GN=TAGLN2 PE=1 SV=3 | 0.963 | 0.765235202 | 0.971 | 0.7 |
| H0YFD6 | Enoyl-CoA hydratase OS=Homo sapiens OX=9606 GN=HADHA PE=1 SV=2 | 1 | 0.876510992 | 0.951 | 0.7 |
| O75083 | WD repeat-containing protein 1 OS=Homo sapiens OX=9606 GN=WDR1 PE=1 SV=4 | 0.998 | 0.967779172 | 0.966 | 0.7 |
| P30101 | Protein disulfide-isomerase A3 OS=Homo sapiens OX=9606 GN=PDIA3 PE=1 SV=4 | 0.97 | 0.95980123 | 1.015 | 0.[illegible] |
| P49368 | T-complex protein 1 subunit gamma OS=Homo sapiens OX=9606 GN=CCT3 PE=1 SV=4 | 0.985 | 0.762199181 | 1.033 | 0.7 |
| P27816 | Microtubule-associated protein 4 OS=Homo sapiens OX=9606 GN=MAP4 PE=1 SV=3 | 1.111 | 0.765235202 | 1.058 | 0.9 |

| | | | | | |
|---|---|---|---|---|---|
| P50991 | T-complex protein 1 subunit delta OS=Homo sapiens OX=9606 GN=CCT4 PE=1 SV=4 | 0.99 | 0.886561233 | 1.038 | 0.7 |
| P19338 | Nucleolin OS=Homo sapiens OX=9606 GN=NCL PE=1 SV=3 | 1.044 | 0.750023665 | 1.061 | 0.7 |
| Q16658 | Fascin OS=Homo sapiens OX=9606 GN=FSCN1 PE=1 SV=3 | 0.925 | 0.765235202 | 0.917 | 0.7 |
| P60174-1 | Isoform 2 of Triosephosphate isomerase OS=Homo sapiens OX=9606 GN=TPI1 | 0.993 | 0.978562349 | 1.006 | 0.8 |
| P05121 | Plasminogen activator inhibitor 1 OS=Homo sapiens OX=9606 GN=SERPINE1 PE=1 SV=1 | 1.117 | 0.750023665 | 0.995 | 0.8 |
| Q9Y678 | Coatomer subunit gamma-1 OS=Homo sapiens OX=9606 GN=COPG1 PE=1 SV=1 | 1.028 | 0.779470244 | 1.01 | 0.9 |
| P55060 | Exportin-2 OS=Homo sapiens OX=9606 GN=CSE1L PE=1 SV=3 | 0.977 | 0.750023665 | 0.987 | 0.8 |
| Q14697 | Neutral alpha-glucosidase AB OS=Homo sapiens OX=9606 GN=GANAB PE=1 SV=3 | 0.964 | 0.801245924 | 0.962 | 0.7 |
| H0Y8C6 | Importin-5 (Fragment) OS=Homo sapiens OX=9606 GN=IPO5 PE=1 SV=1 | 0.99 | 0.80706255 | 1.026 | 0.7 |
| P50990 | T-complex protein 1 subunit theta OS=Homo sapiens OX=9606 GN=CCT8 PE=1 SV=4 | 0.978 | 0.762199181 | 1.038 | 0.7 |
| P43121 | Cell surface glycoprotein MUC18 OS=Homo sapiens OX=9606 GN=MCAM PE=1 SV=2 | 0.961 | 0.80706255 | 0.967 | 0.7 |
| P04792 | Heat shock protein beta-1 OS=Homo sapiens OX=9606 GN=HSPB1 PE=1 SV=2 | 0.961 | 0.750023665 | 0.925 | 0.7 |
| P07814 | Bifunctional glutamate/proline--tRNA ligase OS=Homo sapiens OX=9606 GN=EPRS1 PE=1 SV=5 | 1.027 | 0.904802944 | 1.017 | 0.7 |
| Q99832 | T-complex protein 1 subunit eta OS=Homo sapiens OX=9606 GN=CCT7 PE=1 SV=2 | 0.981 | 0.766667967 | 1.022 | 0.7 |
| Q9Y4L1 | Hypoxia up-regulated protein 1 OS=Homo sapiens OX=9606 GN=HYOU1 PE=1 SV=1 | 0.991 | 0.985330338 | 1.056 | 0.7 |
| O14980 | Exportin-1 OS=Homo sapiens OX=9606 GN=XPO1 PE=1 SV=1 | 0.993 | 0.863796857 | 0.954 | 0.7 |
| P22626 | Heterogeneous nuclear ribonucleoproteins A2/B1 OS=Homo sapiens OX=9606 GN=HNRNPA2B1 PE=1 SV=2 | 1.024 | 0.905369216 | 0.906 | 0.7 |
| Q14697-2 | Isoform 2 of Neutral alpha-glucosidase AB OS=Homo sapiens OX=9606 GN=GANAB | 1.046 | 0.765235202 | 0.93 | 0.7 |
| P04083 | Annexin A1 OS=Homo sapiens OX=9606 GN=ANXA1 PE=1 SV=2 | 0.993 | 0.963059765 | 1.001 | 0.8 |
| P26641 | Elongation factor 1-gamma OS=Homo sapiens OX=9606 GN=EEF1G PE=1 SV=3 | 1.014 | 0.885300104 | 1.068 | 0.7 |
| Q13509 | Tubulin beta-3 chain OS=Homo sapiens OX=9606 GN=TUBB3 PE=1 SV=2 | 1.079 | 0.750023665 | 1.026 | 0.9 |
| A0A1C7CYX9 | Dihydropyrimidinase-related protein 2 OS=Homo sapiens OX=9606 GN=DPYSL2 PE=1 SV=1 | 1.008 | 0.895384791 | 1.011 | 0.9 |
| Q14152 | Eukaryotic translation initiation factor 3 subunit A OS=Homo sapiens OX=9606 GN=EIF3A PE=1 SV=1 | 0.999 | 0.879425665 | 1.076 | 0.7 |
| P61978-3 | Isoform 3 of Heterogeneous nuclear ribonucleoprotein K OS=Homo sapiens OX=9606 GN=HNRNPK | 0.97 | 0.887475968 | 0.95 | 0.7 |
| P04632 | Calpain small subunit 1 OS=Homo sapiens OX=9606 GN=CAPNS1 PE=1 SV=1 | 0.995 | 0.924631778 | 0.969 | 0.8 |
| P30153 | Serine/threonine-protein phosphatase 2A 65 kDa regulatory subunit A alpha isoform OS=Homo sapiens OX=9606 GN=PPP2R1A PI | 0.99 | 0.766667967 | 1.005 | 0.8 |
| Q7KZF4 | Staphylococcal nuclease domain-containing protein 1 OS=Homo sapiens OX=9606 GN=SND1 PE=1 SV=1 | 1.021 | 0.790808367 | 1.041 | 0.7 |
| A0A5F9ZHM4 | L-lactate dehydrogenase OS=Homo sapiens OX=9606 GN=LDHB PE=1 SV=1 | 0.965 | 0.750023665 | 1.048 | 0.7 |
| Q9P2E9 | Ribosome-binding protein 1 OS=Homo sapiens OX=9606 GN=RRBP1 PE=1 SV=5 | 1.04 | 0.773977513 | 0.97 | 0.7 |
| P35221 | Catenin alpha-1 OS=Homo sapiens OX=9606 GN=CTNNA1 PE=1 SV=1 | 0.979 | 0.882454148 | 0.887 | 0.7 |
| Q15365 | Poly(rC)-binding protein 1 OS=Homo sapiens OX=9606 GN=PCBP1 PE=1 SV=2 | 0.992 | 0.990981579 | 1.029 | 0.7 |
| Q13200 | 26S proteasome non-ATPase regulatory subunit 2 OS=Homo sapiens OX=9606 GN=PSMD2 PE=1 SV=3 | 0.976 | 0.765235202 | 1.019 | 0.7 |
| P40227 | T-complex protein 1 subunit zeta OS=Homo sapiens OX=9606 GN=CCT6A PE=1 SV=3 | 0.994 | 0.987438982 | 1.052 | 0.7 |
| P00338 | L-lactate dehydrogenase A chain OS=Homo sapiens OX=9606 GN=LDHA PE=1 SV=2 | 1.007 | 0.919690409 | 1.067 | 0.7 |
| Q9BUF5 | Tubulin beta-6 chain OS=Homo sapiens OX=9606 GN=TUBB6 PE=1 SV=1 | 1.032 | 0.765235202 | 1.067 | 0.8 |
| P06744 | Glucose-6-phosphate isomerase OS=Homo sapiens OX=9606 GN=GPI PE=1 SV=4 | 0.982 | 0.750023665 | 1.014 | 0.7 |
| Q9H223 | EH domain-containing protein 4 OS=Homo sapiens OX=9606 GN=EHD4 PE=1 SV=1 | 1.014 | 0.985324977 | 0.941 | 0.7 |
| P06748 | Nucleophosmin OS=Homo sapiens OX=9606 GN=NPM1 PE=1 SV=2 | 1.076 | 0.750023665 | 1.054 | 0.7 |
| P00352 | Retinal dehydrogenase 1 OS=Homo sapiens OX=9606 GN=ALDH1A1 PE=1 SV=2 | 0.996 | 0.989102644 | 1.186 | 0.7 |
| P43490 | Nicotinamide phosphoribosyltransferase OS=Homo sapiens OX=9606 GN=NAMPT PE=1 SV=1 | 1.016 | 0.957701761 | 1.273 | 0.7 |
| P50454 | Serpin H1 OS=Homo sapiens OX=9606 GN=SERPINH1 PE=1 SV=2 | 1 | 0.968322408 | 1.003 | 0.9 |
| Q6YHK3 | CD109 antigen OS=Homo sapiens OX=9606 GN=CD109 PE=1 SV=2 | 0.989 | 0.876510992 | 0.894 | 0.7 |
| O75923-13 | Isoform 13 of Dysferlin OS=Homo sapiens OX=9606 GN=DYSF | 0.977 | 0.9252392 | 0.96 | 0.7 |
| P04264 | Keratin, type II cytoskeletal 1 OS=Homo sapiens OX=9606 GN=KRT1 PE=1 SV=6 | 0.775 | 0.765235202 | 0.629 | 0.7 |
| P68032 | Actin, alpha cardiac muscle 1 OS=Homo sapiens OX=9606 GN=ACTC1 PE=1 SV=1 | 1.008 | 0.902016292 | 0.948 | 0.7 |
| P63104 | 14-3-3 protein zeta/delta OS=Homo sapiens OX=9606 GN=YWHAZ PE=1 SV=1 | 0.942 | 0.750023665 | 0.984 | 0.9 |
| Q8WUM4 | Programmed cell death 6-interacting protein OS=Homo sapiens OX=9606 GN=PDCD6IP PE=1 SV=1 | 0.99 | 0.994361756 | 1.015 | 0.7 |
| Q8NBS9 | Thioredoxin domain-containing protein 5 OS=Homo sapiens OX=9606 GN=TXNDC5 PE=1 SV=2 | 0.948 | 0.765235202 | 0.984 | 0.8 |
| A6NLN1 | Polypyrimidine tract-binding protein 1 OS=Homo sapiens OX=9606 GN=PTBP1 PE=1 SV=4 | 1.031 | 0.985324977 | 0.954 | 0.8 |
| P16284 | Platelet endothelial cell adhesion molecule OS=Homo sapiens OX=9606 GN=PECAM1 PE=1 SV=2 | 0.967 | 0.947009319 | 0.77 | 0.7 |
| P62258 | 14-3-3 protein epsilon OS=Homo sapiens OX=9606 GN=YWHAE PE=1 SV=1 | 0.987 | 0.889450577 | 0.979 | 0.7 |
| P40926 | Malate dehydrogenase, mitochondrial OS=Homo sapiens OX=9606 GN=MDH2 PE=1 SV=3 | 0.97 | 0.809976184 | 0.97 | 0.7 |
| H3BPE1 | Microtubule-actin cross-linking factor 1, isoforms 1/2/3/5 OS=Homo sapiens OX=9606 GN=MACF1 PE=1 SV=1 | 1.01 | 0.806305724 | 0.931 | 0.7 |
| Q86VP6 | Cullin-associated NEDD8-dissociated protein 1 OS=Homo sapiens OX=9606 GN=CAND1 PE=1 SV=2 | 0.967 | 0.765235202 | 0.967 | 0.7 |
| P06753-5 | Isoform 5 of Tropomyosin alpha-3 chain OS=Homo sapiens OX=9606 GN=TPM3 | 0.963 | 0.750023665 | 0.985 | 0.8 |
| A0A087X0X3 | Heterogeneous nuclear ribonucleoprotein M OS=Homo sapiens OX=9606 GN=HNRNPM PE=1 SV=1 | 1.09 | 0.845656075 | 0.974 | 0.7 |
| O75643 | U5 small nuclear ribonucleoprotein 200 kDa helicase OS=Homo sapiens OX=9606 GN=SNRNP200 PE=1 SV=2 | 1.02 | 0.889450577 | 0.96 | 0.7 |
| P00491 | Purine nucleoside phosphorylase OS=Homo sapiens OX=9606 GN=PNP PE=1 SV=2 | 0.961 | 0.642276967 | 0.979 | 0.7 |
| P33176 | Kinesin-1 heavy chain OS=Homo sapiens OX=9606 GN=KIF5B PE=1 SV=1 | 1.024 | 0.765235202 | 1.027 | 0.7 |
| J3QRS3 | Myosin regulatory light chain 12A OS=Homo sapiens OX=9606 GN=MYL12A PE=1 SV=1 | 0.963 | 0.845656075 | 0.949 | 0.7 |
| P23381 | Tryptophan--tRNA ligase, cytoplasmic OS=Homo sapiens OX=9606 GN=WARS1 PE=1 SV=2 | 0.985 | 0.987438982 | 0.921 | 0.8 |
| O14950 | Myosin regulatory light chain 12B OS=Homo sapiens OX=9606 GN=MYL12B PE=1 SV=2 | 0.923 | 0.770292944 | 0.933 | 0.7 |
| P27824 | Calnexin OS=Homo sapiens OX=9606 GN=CANX PE=1 SV=2 | 0.986 | 0.972616875 | 0.991 | 0.7 |
| P49411 | Elongation factor Tu, mitochondrial OS=Homo sapiens OX=9606 GN=TUFM PE=1 SV=2 | 0.958 | 0.750023665 | 0.963 | 0.7 |
| F8VZX2 | Poly(rC)-binding protein 2 OS=Homo sapiens OX=9606 GN=PCBP2 PE=1 SV=1 | 1.004 | 0.951537478 | 0.985 | 0.7 |
| P28838 | Cytosol aminopeptidase OS=Homo sapiens OX=9606 GN=LAP3 PE=1 SV=3 | 0.968 | 0.774484855 | 0.837 | 0.7 |
| P05556 | Integrin beta-1 OS=Homo sapiens OX=9606 GN=ITGB1 PE=1 SV=2 | 0.956 | 0.808147351 | 1.034 | 0.7 |
| P53618 | Coatomer subunit beta OS=Homo sapiens OX=9606 GN=COPB1 PE=1 SV=3 | 1.017 | 0.76858622 | 1.057 | 0.7 |
| P31948 | Stress-induced-phosphoprotein 1 OS=Homo sapiens OX=9606 GN=STIP1 PE=1 SV=1 | 0.98 | 0.974919191 | 1.026 | 0.7 |
| P17655 | Calpain-2 catalytic subunit OS=Homo sapiens OX=9606 GN=CAPN2 PE=1 SV=6 | 0.986 | 0.792509319 | 0.998 | 0.9 |
| O00429 | Dynamin-1-like protein OS=Homo sapiens OX=9606 GN=DNM1L PE=1 SV=2 | 1.008 | 0.904500837 | 0.972 | 0.7 |
| Q14240 | Eukaryotic initiation factor 4A-II OS=Homo sapiens OX=9606 GN=EIF4A2 PE=1 SV=2 | 0.984 | 0.859846322 | 0.939 | 0.7 |
| P08865 | 40S ribosomal protein SA OS=Homo sapiens OX=9606 GN=RPSA PE=1 SV=4 | 0.979 | 0.816703245 | 1.029 | 0.7 |
| P26640 | Valine--tRNA ligase OS=Homo sapiens OX=9606 GN=VARS1 PE=1 SV=4 | 0.965 | 0.750023665 | 0.987 | 0.7 |
| Q96CX2 | BTB/POZ domain-containing protein KCTD12 OS=Homo sapiens OX=9606 GN=KCTD12 PE=1 SV=1 | 0.964 | 0.913279108 | 1.125 | 0.7 |
| Q12906-7 | Isoform 7 of Interleukin enhancer-binding factor 3 OS=Homo sapiens OX=9606 GN=ILF3 | 0.988 | 0.970974607 | 0.969 | 0.7 |
| P41250 | Glycine--tRNA ligase OS=Homo sapiens OX=9606 GN=GARS1 PE=1 SV=3 | 0.982 | 0.829390484 | 1 | 0.7 |
| Q92616 | eIF-2-alpha kinase activator GCN1 OS=Homo sapiens OX=9606 GN=GCN1 PE=1 SV=6 | 1.027 | 0.762971614 | 1.001 | 0.8 |
| O95373 | Importin-7 OS=Homo sapiens OX=9606 GN=IPO7 PE=1 SV=1 | 0.947 | 0.766667967 | 1.008 | 0.8 |
| P53621 | Coatomer subunit alpha OS=Homo sapiens OX=9606 GN=COPA PE=1 SV=2 | 1.004 | 0.906091542 | 0.985 | 0.7 |
| P43686 | 26S proteasome regulatory subunit 6B OS=Homo sapiens OX=9606 GN=PSMC4 PE=1 SV=2 | 1.002 | 0.984200071 | 0.988 | 0.7 |
| P17980 | 26S proteasome regulatory subunit 6A OS=Homo sapiens OX=9606 GN=PSMC3 PE=1 SV=3 | 0.998 | 0.983953845 | 1.017 | 0.7 |
| P61158 | Actin-related protein 3 OS=Homo sapiens OX=9606 GN=ACTR3 PE=1 SV=3 | 0.996 | 0.939024443 | 0.922 | 0.7 |
| Q07065 | Cytoskeleton-associated protein 4 OS=Homo sapiens OX=9606 GN=CKAP4 PE=1 SV=2 | 1.016 | 0.98478736 | 0.993 | 0.9 |
| Q9BSJ8 | Extended synaptotagmin-1 OS=Homo sapiens OX=9606 GN=ESYT1 PE=1 SV=1 | 1.029 | 0.766667967 | 0.945 | 0.7 |
| O60506 | Heterogeneous nuclear ribonucleoprotein Q OS=Homo sapiens OX=9606 GN=SYNCRIP PE=1 SV=2 | 1.02 | 0.913279108 | 1.015 | 0.7 |
| Q15436 | Protein transport protein Sec23A OS=Homo sapiens OX=9606 GN=SEC23A PE=1 SV=2 | 0.991 | 0.990333809 | 1.015 | 0.7 |
| P09936 | Ubiquitin carboxyl-terminal hydrolase isozyme L1 OS=Homo sapiens OX=9606 GN=UCHL1 PE=1 SV=2 | 0.972 | 0.709661036 | 1.021 | 0.9 |
| Q99536 | Synaptic vesicle membrane protein VAT-1 homolog OS=Homo sapiens OX=9606 GN=VAT1 PE=1 SV=2 | 1.015 | 0.833700751 | 0.953 | 0.7 |
| P05091 | Aldehyde dehydrogenase, mitochondrial OS=Homo sapiens OX=9606 GN=ALDH2 PE=1 SV=2 | 0.976 | 0.750023665 | 0.918 | 0.7 |
| Q99460 | 26S proteasome non-ATPase regulatory subunit 1 OS=Homo sapiens OX=9606 GN=PSMD1 PE=1 SV=2 | 1.027 | 0.917008272 | 0.982 | 0.9 |
| P31946 | 14-3-3 protein beta/alpha OS=Homo sapiens OX=9606 GN=YWHAB PE=1 SV=3 | 0.96 | 0.750023665 | 0.985 | 0.7 |
| Q9UH65 | Switch-associated protein 70 OS=Homo sapiens OX=9606 GN=SWAP70 PE=1 SV=1 | 1.013 | 0.942678074 | 0.966 | 0.7 |
| Q86UX7 | Fermitin family homolog 3 OS=Homo sapiens OX=9606 GN=FERMT3 PE=1 SV=1 | 1.002 | 0.984805107 | 1.008 | 0.7 |
| O00299 | Chloride intracellular channel protein 1 OS=Homo sapiens OX=9606 GN=CLIC1 PE=1 SV=4 | 0.974 | 0.774484855 | 1.023 | 0.7 |
| P39023 | 60S ribosomal protein L3 OS=Homo sapiens OX=9606 GN=RPL3 PE=1 SV=2 | 1.029 | 0.756287988 | 1.126 | 0.7 |

| | | | | | |
|---|---|---|---|---|---|
| P61981 | 14-3-3 protein gamma OS=Homo sapiens OX=9606 GN=YWHAG PE=1 SV=2 | 0.991 | 0.915968895 | 1.01 | 0.7 |
| P07737 | Profilin-1 OS=Homo sapiens OX=9606 GN=PFN1 PE=1 SV=2 | 0.96 | 0.750023665 | 0.988 | 0.8 |
| P63244 | Receptor of activated protein C kinase 1 OS=Homo sapiens OX=9606 GN=RACK1 PE=1 SV=3 | 1.031 | 0.765235202 | 1.041 | 0.7 |
| P62081 | 40S ribosomal protein S7 OS=Homo sapiens OX=9606 GN=RPS7 PE=1 SV=1 | 1.126 | 0.750023665 | 1.1 | 0.7 |
| Q96QK1 | Vacuolar protein sorting-associated protein 35 OS=Homo sapiens OX=9606 GN=VPS35 PE=1 SV=2 | 0.948 | 0.750023665 | 0.939 | 0.7 |
| P54136 | Arginine--tRNA ligase, cytoplasmic OS=Homo sapiens OX=9606 GN=RARS1 PE=1 SV=2 | 1.031 | 0.839959734 | 1.026 | 0.7 |
| Q6NZI2 | Caveolae-associated protein 1 OS=Homo sapiens OX=9606 GN=CAVIN1 PE=1 SV=1 | 1.026 | 0.946791131 | 0.934 | 0.7 |
| P11586 | C-1-tetrahydrofolate synthase, cytoplasmic OS=Homo sapiens OX=9606 GN=MTHFD1 PE=1 SV=3 | 0.969 | 0.750023665 | 1.017 | 0.7 |
| Q15293 | Reticulocalbin-1 OS=Homo sapiens OX=9606 GN=RCN1 PE=1 SV=1 | 0.991 | 0.904500837 | 1.018 | 0.7 |
| Q01518 | Adenylyl cyclase-associated protein 1 OS=Homo sapiens OX=9606 GN=CAP1 PE=1 SV=5 | 0.874 | 0.750023665 | 1.056 | 0.8 |
| E9PLK3 | Aminopeptidase OS=Homo sapiens OX=9606 GN=NPEPPS PE=1 SV=1 | 0.992 | 0.827871327 | 0.981 | 0.7 |
| Q13263 | Transcription intermediary factor 1-beta OS=Homo sapiens OX=9606 GN=TRIM28 PE=1 SV=5 | 0.997 | 0.978562349 | 0.959 | 0.7 |
| P67936 | Tropomyosin alpha-4 chain OS=Homo sapiens OX=9606 GN=TPM4 PE=1 SV=3 | 0.961 | 0.835336742 | 0.962 | 0.7 |
| P52597 | Heterogeneous nuclear ribonucleoprotein F OS=Homo sapiens OX=9606 GN=HNRNPF PE=1 SV=3 | 0.957 | 0.874689721 | 0.955 | 0.7 |
| Q01518-2 | Isoform 2 of Adenylyl cyclase-associated protein 1 OS=Homo sapiens OX=9606 GN=CAP1 | 1.001 | 0.956825234 | 1.045 | 0.7 |
| P24534 | Elongation factor 1-beta OS=Homo sapiens OX=9606 GN=EEF1B2 PE=1 SV=3 | 0.994 | 0.917008272 | 1.054 | 0.7 |
| Q32Q12 | Nucleoside diphosphate kinase OS=Homo sapiens OX=9606 GN=NME1-NME2 PE=1 SV=1 | 0.967 | 0.765235202 | 1.098 | 0.7 |
| E9PK25 | Cofilin, non-muscle isoform OS=Homo sapiens OX=9606 GN=CFL1 PE=1 SV=1 | 0.995 | 0.958664425 | 1.013 | 0.7 |
| P04899 | Guanine nucleotide-binding protein G(i) subunit alpha-2 OS=Homo sapiens OX=9606 GN=GNAI2 PE=1 SV=3 | 0.932 | 0.642276967 | 0.899 | 0.7 |
| P23396 | 40S ribosomal protein S3 OS=Homo sapiens OX=9606 GN=RPS3 PE=1 SV=2 | 1.068 | 0.765235202 | 1.122 | 0.7 |
| H0YIV4 | Nucleosome assembly protein 1-like 1 (Fragment) OS=Homo sapiens OX=9606 GN=NAP1L1 PE=1 SV=1 | 0.927 | 0.765235202 | 1.048 | 0.7 |
| P00505 | Aspartate aminotransferase, mitochondrial OS=Homo sapiens OX=9606 GN=GOT2 PE=1 SV=3 | 1.017 | 0.924992773 | 0.93 | 0.7 |
| O75390 | Citrate synthase, mitochondrial OS=Homo sapiens OX=9606 GN=CS PE=1 SV=2 | 0.995 | 0.95980123 | 0.992 | 0.7 |
| P27708 | CAD protein OS=Homo sapiens OX=9606 GN=CAD PE=1 SV=3 | 1.009 | 0.791905807 | 1.02 | 0.7 |
| P35609-2 | Isoform 2 of Alpha-actinin-2 OS=Homo sapiens OX=9606 GN=ACTN2 | 1.001 | 0.930855729 | 0.937 | 0.7 |
| P36578 | 60S ribosomal protein L4 OS=Homo sapiens OX=9606 GN=RPL4 PE=1 SV=5 | 1.087 | 0.750023665 | 1.103 | 0.7 |
| P09651 | Heterogeneous nuclear ribonucleoprotein A1 OS=Homo sapiens OX=9606 GN=HNRNPA1 PE=1 SV=5 | 1.076 | 0.842027256 | 0.987 | 0.8 |
| O43852 | Calumenin OS=Homo sapiens OX=9606 GN=CALU PE=1 SV=2 | 0.967 | 0.750023665 | 0.945 | 0.7 |
| P61160 | Actin-related protein 2 OS=Homo sapiens OX=9606 GN=ACTR2 PE=1 SV=1 | 0.993 | 0.918702104 | 0.98 | 0.9 |
| Q13201 | Multimerin-1 OS=Homo sapiens OX=9606 GN=MMRN1 PE=1 SV=3 | 0.99 | 0.990981579 | 1.05 | 0.8 |
| O00159 | Unconventional myosin-Ic OS=Homo sapiens OX=9606 GN=MYO1C PE=1 SV=4 | 1.012 | 0.939024443 | 0.923 | 0.7 |
| P09382 | Galectin-1 OS=Homo sapiens OX=9606 GN=LGALS1 PE=1 SV=2 | 0.995 | 0.901934028 | 1.011 | 0.7 |
| P31150 | Rab GDP dissociation inhibitor alpha OS=Homo sapiens OX=9606 GN=GDI1 PE=1 SV=2 | 0.969 | 0.761475165 | 0.944 | 0.7 |
| P46777 | 60S ribosomal protein L5 OS=Homo sapiens OX=9606 GN=RPL5 PE=1 SV=3 | 1.045 | 0.790808367 | 1.068 | 0.7 |
| Q12905 | Interleukin enhancer-binding factor 2 OS=Homo sapiens OX=9606 GN=ILF2 PE=1 SV=2 | 1.003 | 0.905369216 | 0.984 | 0.7 |
| A0A087X253 | AP complex subunit beta OS=Homo sapiens OX=9606 GN=AP2B1 PE=1 SV=1 | 0.978 | 0.76635507 | 0.983 | 0.7 |
| Q15084 | Protein disulfide-isomerase A6 OS=Homo sapiens OX=9606 GN=PDIA6 PE=1 SV=1 | 0.913 | 0.765235202 | 1.052 | 0.7 |
| P14866 | Heterogeneous nuclear ribonucleoprotein L OS=Homo sapiens OX=9606 GN=HNRNPL PE=1 SV=2 | 1.026 | 0.971097726 | 0.919 | 0.7 |
| P11413 | Glucose-6-phosphate 1-dehydrogenase OS=Homo sapiens OX=9606 GN=G6PD PE=1 SV=4 | 0.932 | 0.761475165 | 0.994 | 0.9 |
| P05023 | Sodium/potassium-transporting ATPase subunit alpha-1 OS=Homo sapiens OX=9606 GN=ATP1A1 PE=1 SV=1 | 1.038 | 0.750023665 | 1.033 | 0.7 |
| Q92598 | Heat shock protein 105 kDa OS=Homo sapiens OX=9606 GN=HSPH1 PE=1 SV=1 | 1 | 0.917008272 | 1.105 | 0.7 |
| J3KN67 | Tropomyosin alpha-3 chain OS=Homo sapiens OX=9606 GN=TPM3 PE=1 SV=1 | 0.938 | 0.765235202 | 0.935 | 0.7 |
| D6REX3 | Protein transport protein Sec31A OS=Homo sapiens OX=9606 GN=SEC31A PE=1 SV=1 | 1.038 | 0.750023665 | 1.039 | 0.7 |
| P49321 | Nuclear autoantigenic sperm protein OS=Homo sapiens OX=9606 GN=NASP PE=1 SV=2 | 0.976 | 0.766667967 | 0.997 | 0.8 |
| A0A087WTT1 | Polyadenylate-binding protein OS=Homo sapiens OX=9606 GN=PABPC1 PE=1 SV=1 | 1.031 | 0.996637279 | 1.053 | 0.7 |
| G3V4C1 | Heterogeneous nuclear ribonucleoproteins C1/C2 OS=Homo sapiens OX=9606 GN=HNRNPC PE=1 SV=1 | 1.062 | 0.837941467 | 0.927 | 0.7 |
| P62424 | 60S ribosomal protein L7a OS=Homo sapiens OX=9606 GN=RPL7A PE=1 SV=2 | 1.032 | 0.765235202 | 1.057 | 0.7 |
| E9PCY7 | Heterogeneous nuclear ribonucleoprotein H OS=Homo sapiens OX=9606 GN=HNRNPH1 PE=1 SV=1 | 1.031 | 0.996637279 | 0.934 | 0.7 |
| P62937 | Peptidyl-prolyl cis-trans isomerase A OS=Homo sapiens OX=9606 GN=PPIA PE=1 SV=2 | 1.009 | 0.997835552 | 0.894 | 0.7 |
| O60701 | UDP-glucose 6-dehydrogenase OS=Homo sapiens OX=9606 GN=UGDH PE=1 SV=1 | 0.997 | 0.952913364 | 0.987 | 0.8 |
| P35237 | Serpin B6 OS=Homo sapiens OX=9606 GN=SERPINB6 PE=1 SV=3 | 1.031 | 0.829390484 | 0.953 | 0.7 |
| Q9NYU2 | UDP-glucose:glycoprotein glucosyltransferase 1 OS=Homo sapiens OX=9606 GN=UGGT1 PE=1 SV=3 | 0.973 | 0.750023665 | 0.99 | 0.9 |
| O15144 | Actin-related protein 2/3 complex subunit 2 OS=Homo sapiens OX=9606 GN=ARPC2 PE=1 SV=1 | 0.989 | 0.849231872 | 0.98 | 0.7 |
| Q10567 | AP-1 complex subunit beta-1 OS=Homo sapiens OX=9606 GN=AP1B1 PE=1 SV=2 | 0.971 | 0.980966594 | 0.981 | 0.7 |
| Q12931 | Heat shock protein 75 kDa, mitochondrial OS=Homo sapiens OX=9606 GN=TRAP1 PE=1 SV=3 | 1 | 0.984200071 | 1.026 | 0.8 |
| P15880 | 40S ribosomal protein S2 OS=Homo sapiens OX=9606 GN=RPS2 PE=1 SV=2 | 1.042 | 0.750023665 | 1.09 | 0.7 |
| O43852-4 | Isoform 4 of Calumenin OS=Homo sapiens OX=9606 GN=CALU | 0.972 | 0.750023665 | 1.029 | 0.9 |
| Q16531 | DNA damage-binding protein 1 OS=Homo sapiens OX=9606 GN=DDB1 PE=1 SV=1 | 1.016 | 0.765235202 | 0.978 | 0.8 |
| A0A669KAX4 | Cullin-4B OS=Homo sapiens OX=9606 GN=CUL4B PE=1 SV=1 | 0.973 | 0.765235202 | 1.029 | 0.7 |
| P04843 | Dolichyl-diphosphooligosaccharide--protein glycosyltransferase subunit 1 OS=Homo sapiens OX=9606 GN=RPN1 PE=1 SV=1 | 1.013 | 0.994361756 | 0.94 | 0.7 |
| P04844 | Dolichyl-diphosphooligosaccharide--protein glycosyltransferase subunit 2 OS=Homo sapiens OX=9606 GN=RPN2 PE=1 SV=3 | 1.012 | 0.968693062 | 0.937 | 0.8 |
| Q9H0U4 | Ras-related protein Rab-1B OS=Homo sapiens OX=9606 GN=RAB1B PE=1 SV=1 | 0.98 | 0.889450577 | 1.001 | 0.7 |
| O60664 | Perilipin-3 OS=Homo sapiens OX=9606 GN=PLIN3 PE=1 SV=3 | 0.984 | 0.765235202 | 0.991 | 0.8 |
| P05455 | Lupus La protein OS=Homo sapiens OX=9606 GN=SSB PE=1 SV=2 | 0.988 | 0.882454148 | 1.013 | 0.7 |
| P30041 | Peroxiredoxin-6 OS=Homo sapiens OX=9606 GN=PRDX6 PE=1 SV=3 | 0.94 | 0.750023665 | 0.982 | 0.7 |
| P55884 | Eukaryotic translation initiation factor 3 subunit B OS=Homo sapiens OX=9606 GN=EIF3B PE=1 SV=3 | 1.021 | 0.750023665 | 1.06 | 0.7 |
| Q9BWD1 | Acetyl-CoA acetyltransferase, cytosolic OS=Homo sapiens OX=9606 GN=ACAT2 PE=1 SV=2 | 1.023 | 0.863796857 | 1.02 | 0.9 |
| P31939 | Bifunctional purine biosynthesis protein ATIC OS=Homo sapiens OX=9606 GN=ATIC PE=1 SV=3 | 0.996 | 0.883033668 | 1.011 | 0.9 |
| Q05682 | Caldesmon OS=Homo sapiens OX=9606 GN=CALD1 PE=1 SV=3 | 1.134 | 0.750023665 | 0.888 | 0.7 |
| A0A024R571 | EH domain-containing protein 1 OS=Homo sapiens OX=9606 GN=EHD1 PE=1 SV=1 | 1.028 | 0.889450577 | 0.934 | 0.7 |
| P54652 | Heat shock-related 70 kDa protein 2 OS=Homo sapiens OX=9606 GN=HSPA2 PE=1 SV=1 | 1.017 | 0.913279108 | 1.082 | 0.7 |
| P22102 | Trifunctional purine biosynthetic protein adenosine-3 OS=Homo sapiens OX=9606 GN=GART PE=1 SV=1 | 0.995 | 0.95980123 | 1.034 | 0.7 |
| Q96AC1 | Fermitin family homolog 2 OS=Homo sapiens OX=9606 GN=FERMT2 PE=1 SV=1 | 0.985 | 0.793453909 | 0.925 | 0.7 |
| O43390 | Heterogeneous nuclear ribonucleoprotein R OS=Homo sapiens OX=9606 GN=HNRNPR PE=1 SV=1 | 1.053 | 0.761475165 | 0.956 | 0.7 |
| P49593 | Protein phosphatase 1F OS=Homo sapiens OX=9606 GN=PPM1F PE=1 SV=3 | 0.966 | 0.856817632 | 0.939 | 0.7 |
| P27105 | Erythrocyte band 7 integral membrane protein OS=Homo sapiens OX=9606 GN=STOM PE=1 SV=3 | 0.943 | 0.642276967 | 0.908 | 0.7 |
| H0Y4R1 | Inosine-5'-monophosphate dehydrogenase 2 (Fragment) OS=Homo sapiens OX=9606 GN=IMPDH2 PE=1 SV=1 | 0.961 | 0.750023665 | 1.067 | 0.7 |
| Q13561 | Dynactin subunit 2 OS=Homo sapiens OX=9606 GN=DCTN2 PE=1 SV=4 | 1.028 | 0.765235202 | 1.021 | 0.7 |
| Q7L1Q6 | Basic leucine zipper and W2 domain-containing protein 1 OS=Homo sapiens OX=9606 GN=BZW1 PE=1 SV=1 | 1.056 | 0.750023665 | 1.06 | 0.7 |
| Q9UQ80 | Proliferation-associated protein 2G4 OS=Homo sapiens OX=9606 GN=PA2G4 PE=1 SV=3 | 1.026 | 0.829390484 | 1.033 | 0.7 |
| P37837 | Transaldolase OS=Homo sapiens OX=9606 GN=TALDO1 PE=1 SV=2 | 0.963 | 0.773977513 | 0.998 | 0.8 |
| P27348 | 14-3-3 protein theta OS=Homo sapiens OX=9606 GN=YWHAQ PE=1 SV=1 | 0.979 | 0.809976184 | 0.99 | 0.9 |
| Q92973 | Transportin-1 OS=Homo sapiens OX=9606 GN=TNPO1 PE=1 SV=2 | 0.972 | 0.750023665 | 1.093 | 0.7 |
| Q04637 | Eukaryotic translation initiation factor 4 gamma 1 OS=Homo sapiens OX=9606 GN=EIF4G1 PE=1 SV=4 | 1.042 | 0.750023665 | 1.092 | 0.7 |
| P41252 | Isoleucine--tRNA ligase, cytoplasmic OS=Homo sapiens OX=9606 GN=IARS1 PE=1 SV=2 | 1.015 | 0.765937763 | 1.011 | 0.7 |
| P07954 | Fumarate hydratase, mitochondrial OS=Homo sapiens OX=9606 GN=FH PE=1 SV=3 | 0.939 | 0.703015003 | 0.993 | 0.7 |
| Q99497 | Parkinson disease protein 7 OS=Homo sapiens OX=9606 GN=PARK7 PE=1 SV=2 | 0.951 | 0.762971614 | 0.982 | 0.7 |
| P55795 | Heterogeneous nuclear ribonucleoprotein H2 OS=Homo sapiens OX=9606 GN=HNRNPH2 PE=1 SV=1 | 0.985 | 0.765235202 | 0.935 | 0.7 |
| A0A0C4DGA2 | Enoyl-CoA delta isomerase 2, mitochondrial OS=Homo sapiens OX=9606 GN=ECI2 PE=1 SV=1 | 0.986 | 0.827137513 | 0.977 | 0.8 |
| P15121 | Aldo-keto reductase family 1 member B1 OS=Homo sapiens OX=9606 GN=AKR1B1 PE=1 SV=3 | 0.974 | 0.896335635 | 0.998 | 0.7 |
| Q6P2Q9 | Pre-mRNA-processing-splicing factor 8 OS=Homo sapiens OX=9606 GN=PRPF8 PE=1 SV=2 | 1.046 | 0.870355966 | 0.987 | 0.7 |
| P00367 | Glutamate dehydrogenase 1, mitochondrial OS=Homo sapiens OX=9606 GN=GLUD1 PE=1 SV=2 | 0.984 | 0.765235202 | 0.968 | 0.7 |
| Q9Y696 | Chloride intracellular channel protein 4 OS=Homo sapiens OX=9606 GN=CLIC4 PE=1 SV=4 | 0.967 | 0.889450577 | 0.913 | 0.7 |
| Q13838 | Spliceosome RNA helicase DDX39B OS=Homo sapiens OX=9606 GN=DDX39B PE=1 SV=1 | 0.982 | 0.766667967 | 0.955 | 0.8 |
| O00571 | ATP-dependent RNA helicase DDX3X OS=Homo sapiens OX=9606 GN=DDX3X PE=1 SV=3 | 1.037 | 0.750023665 | 1.071 | 0.7 |
| Q92499 | ATP-dependent RNA helicase DDX1 OS=Homo sapiens OX=9606 GN=DDX1 PE=1 SV=2 | 1.003 | 0.891352629 | 0.991 | 0.7 |

| | | | | | |
|---|---|---|---|---|---|
| P35527 | Keratin, type I cytoskeletal 9 OS=Homo sapiens OX=9606 GN=KRT9 PE=1 SV=3 | 0.747 | 0.765235202 | 0.654 | 0.7 |
| J3KTA4 | DEAD box protein 5 OS=Homo sapiens OX=9606 GN=DDX5 PE=1 SV=1 | 1.12 | 0.750023665 | 1.023 | 0.7 |
| P05388 | 60S acidic ribosomal protein P0 OS=Homo sapiens OX=9606 GN=RPLP0 PE=1 SV=1 | 1.024 | 0.840158726 | 1.128 | 0.7 |
| P13489 | Ribonuclease inhibitor OS=Homo sapiens OX=9606 GN=RNH1 PE=1 SV=2 | 0.947 | 0.849561288 | 0.998 | 0.9 |
| P50453 | Serpin B9 OS=Homo sapiens OX=9606 GN=SERPINB9 PE=1 SV=1 | 1.006 | 0.916568161 | 1.016 | 0.7 |
| A0A6I8PL42 | Leucine--tRNA ligase, cytoplasmic OS=Homo sapiens OX=9606 GN=LARS1 PE=4 SV=1 | 1.039 | 0.840158726 | 1.037 | 0.7 |
| P06737 | Glycogen phosphorylase, liver form OS=Homo sapiens OX=9606 GN=PYGL PE=1 SV=4 | 0.976 | 0.750023665 | 0.967 | 0.7 |
| P23246 | Splicing factor, proline- and glutamine-rich OS=Homo sapiens OX=9606 GN=SFPQ PE=1 SV=2 | 1.156 | 0.805136244 | 1.029 | 0.7 |
| O75874 | Isocitrate dehydrogenase [NADP] cytoplasmic OS=Homo sapiens OX=9606 GN=IDH1 PE=1 SV=2 | 0.96 | 0.750023665 | 0.98 | 0.8 |
| P54578 | Ubiquitin carboxyl-terminal hydrolase 14 OS=Homo sapiens OX=9606 GN=USP14 PE=1 SV=3 | 0.969 | 0.762971614 | 0.963 | 0.7 |
| P67936-2 | Isoform 2 of Tropomyosin alpha-4 chain OS=Homo sapiens OX=9606 GN=TPM4 | 0.976 | 0.765235202 | 0.901 | 0.7 |
| P61586 | Transforming protein RhoA OS=Homo sapiens OX=9606 GN=RHOA PE=1 SV=1 | 1.004 | 0.866300162 | 1.025 | 0.7 |
| P62701 | 40S ribosomal protein S4, X isoform OS=Homo sapiens OX=9606 GN=RPS4X PE=1 SV=2 | 1.046 | 0.750023665 | 1.108 | 0.7 |
| P22234 | Multifunctional protein ADE2 OS=Homo sapiens OX=9606 GN=PAICS PE=1 SV=3 | 0.969 | 0.765235202 | 1.054 | 0.7 |
| E9PGM4 | 1,4-alpha-glucan branching enzyme OS=Homo sapiens OX=9606 GN=GBE1 PE=1 SV=1 | 0.971 | 0.783600149 | 1.051 | 0.7 |
| K7EKE6 | Lon protease homolog, mitochondrial OS=Homo sapiens OX=9606 GN=LONP1 PE=1 SV=1 | 0.997 | 0.961750547 | 0.96 | 0.7 |
| P56192 | Methionine--tRNA ligase, cytoplasmic OS=Homo sapiens OX=9606 GN=MARS1 PE=1 SV=2 | 0.963 | 0.792509319 | 0.993 | 0.7 |
| Q14914 | Prostaglandin reductase 1 OS=Homo sapiens OX=9606 GN=PTGR1 PE=1 SV=2 | 0.968 | 0.927148669 | 0.98 | 0.7 |
| E9PEX6 | Dihydrolipoyl dehydrogenase OS=Homo sapiens OX=9606 GN=DLD PE=1 SV=1 | 0.967 | 0.750023665 | 0.963 | 0.7 |
| P18669 | Phosphoglycerate mutase 1 OS=Homo sapiens OX=9606 GN=PGAM1 PE=1 SV=2 | 1.008 | 0.892311206 | 0.956 | 0.9 |
| A2A274 | Aconitate hydratase, mitochondrial OS=Homo sapiens OX=9606 GN=ACO2 PE=1 SV=1 | 1.001 | 0.984442117 | 0.949 | 0.7 |
| Q99714 | 3-hydroxyacyl-CoA dehydrogenase type-2 OS=Homo sapiens OX=9606 GN=HSD17B10 PE=1 SV=3 | 0.989 | 0.829390484 | 0.994 | 0.8 |
| Q9NYL9 | Tropomodulin-3 OS=Homo sapiens OX=9606 GN=TMOD3 PE=1 SV=1 | 0.994 | 0.949254122 | 0.982 | 0.7 |
| P45974 | Ubiquitin carboxyl-terminal hydrolase 5 OS=Homo sapiens OX=9606 GN=USP5 PE=1 SV=2 | 0.97 | 0.765235202 | 0.963 | 0.7 |
| P78417 | Glutathione S-transferase omega-1 OS=Homo sapiens OX=9606 GN=GSTO1 PE=1 SV=2 | 0.994 | 0.973388826 | 0.999 | 0.9 |
| P09525 | Annexin A4 OS=Homo sapiens OX=9606 GN=ANXA4 PE=1 SV=4 | 0.996 | 0.956825234 | 0.964 | 0.7 |
| Q5T5C7 | Seryl-tRNA synthetase OS=Homo sapiens OX=9606 GN=SARS1 PE=1 SV=1 | 0.988 | 0.766667967 | 1.038 | 0.7 |
| Q15019 | Septin-2 OS=Homo sapiens OX=9606 GN=SEPTIN2 PE=1 SV=1 | 0.995 | 0.990017996 | 0.953 | 0.7 |
| Q04917 | 14-3-3 protein eta OS=Homo sapiens OX=9606 GN=YWHAH PE=1 SV=4 | 0.984 | 0.750023665 | 1.033 | 0.9 |
| P46939 | Utrophin OS=Homo sapiens OX=9606 GN=UTRN PE=1 SV=2 | 1.017 | 0.908555656 | 0.938 | 0.7 |
| Q9UBT2 | SUMO-activating enzyme subunit 2 OS=Homo sapiens OX=9606 GN=UBA2 PE=1 SV=2 | 0.969 | 0.876510992 | 0.977 | 0.7 |
| P25786 | Proteasome subunit alpha type-1 OS=Homo sapiens OX=9606 GN=PSMA1 PE=1 SV=1 | 0.968 | 0.766667967 | 1.016 | 0.7 |
| H0Y8G5 | Heterogeneous nuclear ribonucleoprotein D0 (Fragment) OS=Homo sapiens OX=9606 GN=HNRNPD PE=1 SV=8 | 0.996 | 0.860909277 | 0.927 | 0.7 |
| Q92945 | Far upstream element-binding protein 2 OS=Homo sapiens OX=9606 GN=KHSRP PE=1 SV=4 | 0.973 | 0.947009319 | 0.969 | 0.7 |
| P18124 | 60S ribosomal protein L7 OS=Homo sapiens OX=9606 GN=RPL7 PE=1 SV=1 | 1.038 | 0.803837966 | 1.076 | 0.7 |
| Q06323 | Proteasome activator complex subunit 1 OS=Homo sapiens OX=9606 GN=PSME1 PE=1 SV=1 | 0.975 | 0.95980123 | 1.034 | 0.7 |
| Q562R1 | Beta-actin-like protein 2 OS=Homo sapiens OX=9606 GN=ACTBL2 PE=1 SV=2 | 0.963 | 0.869995963 | 0.901 | 0.7 |
| J3KN16 | Proteasome adapter and scaffold protein ECM29 OS=Homo sapiens OX=9606 GN=ECPAS PE=1 SV=1 | 0.992 | 0.996578046 | 0.971 | 0.7 |
| P0DP25 | Calmodulin-3 OS=Homo sapiens OX=9606 GN=CALM3 PE=1 SV=1 | 0.934 | 0.956825234 | 1.005 | 0.8 |
| P60981 | Destrin OS=Homo sapiens OX=9606 GN=DSTN PE=1 SV=3 | 0.982 | 0.934711762 | 1.013 | 0.7 |
| P68036 | Ubiquitin-conjugating enzyme E2 L3 OS=Homo sapiens OX=9606 GN=UBE2L3 PE=1 SV=1 | 1.003 | 0.905369216 | 1.046 | 0.7 |
| Q96TA1 | Protein Niban 2 OS=Homo sapiens OX=9606 GN=NIBAN2 PE=1 SV=3 | 1.016 | 0.765235202 | 0.974 | 0.7 |
| P21399 | Cytoplasmic aconitate hydratase OS=Homo sapiens OX=9606 GN=ACO1 PE=1 SV=3 | 0.998 | 0.904500837 | 0.946 | 0.7 |
| O43242 | 26S proteasome non-ATPase regulatory subunit 3 OS=Homo sapiens OX=9606 GN=PSMD3 PE=1 SV=2 | 1.016 | 0.873855294 | 1.021 | 0.8 |
| P52306 | Rap1 GTPase-GDP dissociation stimulator 1 OS=Homo sapiens OX=9606 GN=RAP1GDS1 PE=1 SV=3 | 0.98 | 0.750023665 | 0.997 | 0.8 |
| P05387 | 60S acidic ribosomal protein P2 OS=Homo sapiens OX=9606 GN=RPLP2 PE=1 SV=1 | 0.965 | 0.978562349 | 1.051 | 0.7 |
| A0A5F9ZHL1 | Acetyl-CoA acetyltransferase, mitochondrial OS=Homo sapiens OX=9606 GN=ACAT1 PE=1 SV=1 | 0.983 | 0.829390484 | 0.943 | 0.7 |
| P62820 | Ras-related protein Rab-1A OS=Homo sapiens OX=9606 GN=RAB1A PE=1 SV=3 | 1.022 | 0.901460271 | 1.036 | 0.7 |
| G3V1V0 | Myosin light polypeptide 6 OS=Homo sapiens OX=9606 GN=MYL6 PE=1 SV=1 | 0.998 | 0.922498596 | 0.974 | 0.9 |
| Q5JR08 | Rho-related GTP-binding protein RhoC (Fragment) OS=Homo sapiens OX=9606 GN=RHOC PE=1 SV=8 | 0.962 | 0.784922527 | 1.053 | 0.7 |
| P51659 | Peroxisomal multifunctional enzyme type 2 OS=Homo sapiens OX=9606 GN=HSD17B4 PE=1 SV=3 | 1.007 | 0.89814168 | 0.995 | 0.8 |
| P63241 | Eukaryotic translation initiation factor 5A-1 OS=Homo sapiens OX=9606 GN=EIF5A PE=1 SV=2 | 0.959 | 0.869094069 | 1.064 | 0.7 |
| P23284 | Peptidyl-prolyl cis-trans isomerase B OS=Homo sapiens OX=9606 GN=PPIB PE=1 SV=2 | 0.994 | 0.949678359 | 0.986 | 0.8 |
| O95810 | Caveolae-associated protein 2 OS=Homo sapiens OX=9606 GN=CAVIN2 PE=1 SV=3 | 1.004 | 0.99130786 | 0.884 | 0.7 |
| P62140 | Serine/threonine-protein phosphatase PP1-beta catalytic subunit OS=Homo sapiens OX=9606 GN=PPP1CB PE=1 SV=3 | 1.056 | 0.873607677 | 0.896 | 0.7 |
| P29966 | Myristoylated alanine-rich C-kinase substrate OS=Homo sapiens OX=9606 GN=MARCKS PE=1 SV=4 | 0.952 | 0.985324977 | 0.866 | 0.7 |
| Q15417 | Calponin-3 OS=Homo sapiens OX=9606 GN=CNN3 PE=1 SV=1 | 1.018 | 0.765235202 | 0.931 | 0.7 |
| O14579 | Coatomer subunit epsilon OS=Homo sapiens OX=9606 GN=COPE PE=1 SV=3 | 1.051 | 0.765235202 | 1 | 0.9 |
| P61247 | 40S ribosomal protein S3a OS=Homo sapiens OX=9606 GN=RPS3A PE=1 SV=2 | 1.031 | 0.805488634 | 1.088 | 0.7 |
| Q13409-2 | Isoform 2B of Cytoplasmic dynein 1 intermediate chain 2 OS=Homo sapiens OX=9606 GN=DYNC1I2 | 0.999 | 0.9962822 | 0.981 | 0.7 |
| E9PKG1 | Protein arginine N-methyltransferase 1 OS=Homo sapiens OX=9606 GN=PRMT1 PE=1 SV=1 | 0.968 | 0.750023665 | 0.942 | 0.7 |
| P38606 | V-type proton ATPase catalytic subunit A OS=Homo sapiens OX=9606 GN=ATP6V1A PE=1 SV=2 | 0.989 | 0.994361756 | 1.011 | 0.9 |
| P30044 | Peroxiredoxin-5, mitochondrial OS=Homo sapiens OX=9606 GN=PRDX5 PE=1 SV=4 | 0.987 | 0.902016292 | 0.966 | 0.7 |
| P60953 | Cell division control protein 42 homolog OS=Homo sapiens OX=9606 GN=CDC42 PE=1 SV=2 | 0.999 | 0.991801854 | 0.946 | 0.7 |
| P62841 | 40S ribosomal protein S15 OS=Homo sapiens OX=9606 GN=RPS15 PE=1 SV=2 | 1.075 | 0.750023665 | 1.049 | 0.7 |
| P62136 | Serine/threonine-protein phosphatase PP1-alpha catalytic subunit OS=Homo sapiens OX=9606 GN=PPP1CA PE=1 SV=1 | 1.027 | 0.839845191 | 0.93 | 0.7 |
| P35998 | 26S proteasome regulatory subunit 7 OS=Homo sapiens OX=9606 GN=PSMC2 PE=1 SV=3 | 0.98 | 0.886561233 | 0.993 | 0.8 |
| P61204 | ADP-ribosylation factor 3 OS=Homo sapiens OX=9606 GN=ARF3 PE=1 SV=2 | 0.981 | 0.809976184 | 1.008 | 0.8 |
| P48444 | Coatomer subunit delta OS=Homo sapiens OX=9606 GN=ARCN1 PE=1 SV=1 | 1.015 | 0.756935672 | 0.991 | 0.8 |
| O75533 | Splicing factor 3B subunit 1 OS=Homo sapiens OX=9606 GN=SF3B1 PE=1 SV=3 | 1.03 | 0.882454148 | 0.969 | 0.7 |
| P62805 | Histone H4 OS=Homo sapiens OX=9606 GN=H4C1 PE=1 SV=2 | 0.64 | 0.765235202 | 0.891 | 0.7 |
| P27635 | 60S ribosomal protein L10 OS=Homo sapiens OX=9606 GN=RPL10 PE=1 SV=4 | 1.092 | 0.642276967 | 1.128 | 0.7 |
| M0R0F0 | 40S ribosomal protein S5 (Fragment) OS=Homo sapiens OX=9606 GN=RPS5 PE=1 SV=1 | 1.073 | 0.765235202 | 1.171 | 0.7 |
| A0A087WUQ6 | Glutathione peroxidase OS=Homo sapiens OX=9606 GN=GPX1 PE=1 SV=1 | 0.972 | 0.824559577 | 1.018 | 0.7 |
| Q14195-2 | Isoform LCRMP-4 of Dihydropyrimidinase-related protein 3 OS=Homo sapiens OX=9606 GN=DPYSL3 | 0.996 | 0.962240823 | 0.94 | 0.7 |
| E7ETU9 | Procollagen-lysine 5-dioxygenase OS=Homo sapiens OX=9606 GN=PLOD2 PE=1 SV=1 | 0.963 | 0.863796857 | 0.977 | 0.7 |
| Q92841 | Probable ATP-dependent RNA helicase DDX17 OS=Homo sapiens OX=9606 GN=DDX17 PE=1 SV=2 | 1.044 | 0.835336742 | 0.955 | 0.7 |
| Q99880 | Histone H2B type 1-L OS=Homo sapiens OX=9606 GN=H2BC13 PE=1 SV=3 | | | | |
| P55263 | Adenosine kinase OS=Homo sapiens OX=9606 GN=ADK PE=1 SV=2 | 0.989 | 0.790808367 | 0.97 | 0.7 |
| P25787 | Proteasome subunit alpha type-2 OS=Homo sapiens OX=9606 GN=PSMA2 PE=1 SV=2 | 1.004 | 0.95980123 | 0.991 | 0.9 |
| P67809 | Y-box-binding protein 1 OS=Homo sapiens OX=9606 GN=YBX1 PE=1 SV=3 | 1.037 | 0.780884711 | 1.128 | 0.7 |
| Q02878 | 60S ribosomal protein L6 OS=Homo sapiens OX=9606 GN=RPL6 PE=1 SV=3 | 1.05 | 0.765235202 | 1.096 | 0.7 |
| Q99879 | Histone H2B type 1-M OS=Homo sapiens OX=9606 GN=H2BC14 PE=1 SV=3 | 0.731 | 0.766667967 | 0.94 | 0.9 |
| P26639 | Threonine--tRNA ligase 1, cytoplasmic OS=Homo sapiens OX=9606 GN=TARS1 PE=1 SV=3 | 0.988 | 0.917008272 | 1.019 | 0.7 |
| O15067 | Phosphoribosylformylglycinamidine synthase OS=Homo sapiens OX=9606 GN=PFAS PE=1 SV=4 | 1.048 | 0.882454148 | 0.978 | 0.7 |
| P34897 | Serine hydroxymethyltransferase, mitochondrial OS=Homo sapiens OX=9606 GN=SHMT2 PE=1 SV=3 | 1.013 | 0.905369216 | 0.99 | 0.8 |
| P11216 | Glycogen phosphorylase, brain form OS=Homo sapiens OX=9606 GN=PYGB PE=1 SV=5 | 0.972 | 0.980741283 | 0.998 | 0.9 |
| P48735 | Isocitrate dehydrogenase [NADP], mitochondrial OS=Homo sapiens OX=9606 GN=IDH2 PE=1 SV=2 | 1.006 | 0.900915273 | 1.012 | 0.9 |
| P09429 | High mobility group protein B1 OS=Homo sapiens OX=9606 GN=HMGB1 PE=1 SV=3 | 1.072 | 0.765235202 | 0.998 | 0.8 |
| Q9Y230 | RuvB-like 2 OS=Homo sapiens OX=9606 GN=RUVBL2 PE=1 SV=3 | 0.964 | 0.750023665 | 0.99 | 0.9 |
| Q16643 | Drebrin OS=Homo sapiens OX=9606 GN=DBN1 PE=1 SV=4 | 1.024 | 0.886561233 | 0.954 | 0.9 |
| Q02790 | Peptidyl-prolyl cis-trans isomerase FKBP4 OS=Homo sapiens OX=9606 GN=FKBP4 PE=1 SV=3 | 0.979 | 0.702711636 | 1.05 | 0.7 |
| P51148 | Ras-related protein Rab-5C OS=Homo sapiens OX=9606 GN=RAB5C PE=1 SV=2 | 0.977 | 0.854598401 | 0.991 | 0.8 |
| P46821 | Microtubule-associated protein 1B OS=Homo sapiens OX=9606 GN=MAP1B PE=1 SV=2 | 1.139 | 0.827871327 | 1.065 | 0.8 |
| Q16778 | Histone H2B type 2-E OS=Homo sapiens OX=9606 GN=H2BC21 PE=1 SV=3 | 0.915 | 0.869473756 | 0.959 | 0.9 |
| Q00341 | Vigilin OS=Homo sapiens OX=9606 GN=HDLBP PE=1 SV=2 | 1.035 | 0.765937763 | 1.019 | 0.7 |

| | | | | | |
|---|---|---|---|---|---|
| Q15942 | Zyxin OS=Homo sapiens OX=9606 GN=ZYX PE=1 SV=1 | 1.044 | 0.766667967 | 0.994 | 0.9 |
| E7EUC7 | UTP--glucose-1-phosphate uridylyltransferase OS=Homo sapiens OX=9606 GN=UGP2 PE=1 SV=1 | 0.962 | 0.901934028 | 0.996 | 0.7 |
| A0A2R8Y7Z0 | Catenin beta-1 OS=Homo sapiens OX=9606 GN=CTNNB1 PE=1 SV=1 | 0.944 | 0.750023665 | 0.851 | 0.7 |
| Q5T4S7 | E3 ubiquitin-protein ligase UBR4 OS=Homo sapiens OX=9606 GN=UBR4 PE=1 SV=1 | 1.072 | 0.750023665 | 0.94 | 0.7 |
| P30154 | Serine/threonine-protein phosphatase 2A 65 kDa regulatory subunit A beta isoform OS=Homo sapiens OX=9606 GN=PPP2R1B PE= | 1.007 | 0.973388826 | 0.931 | 0.7 |
| B7Z7P8 | Eukaryotic peptide chain release factor subunit 1 OS=Homo sapiens OX=9606 GN=ETF1 PE=1 SV=1 | 1.027 | 0.750023665 | 1.045 | 0.7 |
| P20290 | Transcription factor BTF3 OS=Homo sapiens OX=9606 GN=BTF3 PE=1 SV=1 | 1.04 | 0.750023665 | 1.127 | 0.7 |
| P00387 | NADH-cytochrome b5 reductase 3 OS=Homo sapiens OX=9606 GN=CYB5R3 PE=1 SV=3 | 1.032 | 0.909956852 | 0.972 | 0.7 |
| P12429 | Annexin A3 OS=Homo sapiens OX=9606 GN=ANXA3 PE=1 SV=3 | 0.964 | 0.750023665 | 0.957 | 0.7 |
| P09104 | Gamma-enolase OS=Homo sapiens OX=9606 GN=ENO2 PE=1 SV=3 | 0.996 | 0.989102644 | 0.899 | 0.7 |
| Q9UDY2 | Tight junction protein ZO-2 OS=Homo sapiens OX=9606 GN=TJP2 PE=1 SV=2 | 1.025 | 0.770292944 | 0.978 | 0.7 |
| P49915 | GMP synthase [glutamine-hydrolyzing] OS=Homo sapiens OX=9606 GN=GMPS PE=1 SV=1 | 0.977 | 0.750023665 | 0.952 | 0.7 |
| P15531 | Nucleoside diphosphate kinase A OS=Homo sapiens OX=9606 GN=NME1 PE=1 SV=1 | 0.895 | 0.870355966 | 0.965 | 0.8 |
| P07741 | Adenine phosphoribosyltransferase OS=Homo sapiens OX=9606 GN=APRT PE=1 SV=2 | 0.958 | 0.750023665 | 0.994 | 0.8 |
| P51665 | 26S proteasome non-ATPase regulatory subunit 7 OS=Homo sapiens OX=9606 GN=PSMD7 PE=1 SV=2 | 0.991 | 0.996637279 | 0.99 | 0.9 |
| Q15181 | Inorganic pyrophosphatase OS=Homo sapiens OX=9606 GN=PPA1 PE=1 SV=2 | 0.978 | 0.765235202 | 1.018 | 0.7 |
| P84077 | ADP-ribosylation factor 1 OS=Homo sapiens OX=9606 GN=ARF1 PE=1 SV=2 | 1.072 | 0.803600946 | 0.924 | 0.7 |
| Q07021 | Complement component 1 Q subcomponent-binding protein, mitochondrial OS=Homo sapiens OX=9606 GN=C1QBP PE=1 SV=1 | 0.955 | 0.762971614 | 0.982 | 0.7 |
| Q9Y265 | RuvB-like 1 OS=Homo sapiens OX=9606 GN=RUVBL1 PE=1 SV=1 | 0.98 | 0.766667967 | 0.983 | 0.9 |
| E7EX90 | Dynactin subunit 1 OS=Homo sapiens OX=9606 GN=DCTN1 PE=1 SV=1 | 1.03 | 0.830305674 | 0.979 | 0.7 |
| H3BNC9 | 40S ribosomal protein S17 OS=Homo sapiens OX=9606 PE=3 SV=2 | 1.029 | 0.916842788 | 1.048 | 0.7 |
| Q96P70 | Importin-9 OS=Homo sapiens OX=9606 GN=IPO9 PE=1 SV=3 | 1.002 | 0.890382159 | 0.969 | 0.7 |
| O43175 | D-3-phosphoglycerate dehydrogenase OS=Homo sapiens OX=9606 GN=PHGDH PE=1 SV=4 | 0.988 | 0.898315154 | 0.946 | 0.7 |
| P17858 | ATP-dependent 6-phosphofructokinase, liver type OS=Homo sapiens OX=9606 GN=PFKL PE=1 SV=6 | 1.007 | 0.860909277 | 0.94 | 0.7 |
| P10768 | S-formylglutathione hydrolase OS=Homo sapiens OX=9606 GN=ESD PE=1 SV=2 | 0.952 | 0.765235202 | 1.018 | 0.9 |
| H9KV28 | Protein diaphanous homolog 1 OS=Homo sapiens OX=9606 GN=DIAPH1 PE=1 SV=2 | 0.995 | 0.905868507 | 0.981 | 0.9 |
| Q1KMD3 | Heterogeneous nuclear ribonucleoprotein U-like protein 2 OS=Homo sapiens OX=9606 GN=HNRNPUL2 PE=1 SV=1 | 1.028 | 0.876510992 | 0.918 | 0.7 |
| G3V180 | Dipeptidyl peptidase 3 OS=Homo sapiens OX=9606 GN=DPP3 PE=1 SV=1 | 1.015 | 0.963059765 | 0.94 | 0.7 |
| Q99729-3 | Isoform 3 of Heterogeneous nuclear ribonucleoprotein A/B OS=Homo sapiens OX=9606 GN=HNRNPAB | 1.056 | 0.780884711 | 0.97 | 0.9 |
| P14923 | Junction plakoglobin OS=Homo sapiens OX=9606 GN=JUP PE=1 SV=3 | 0.986 | 0.869094069 | 0.808 | 0.7 |
| P62195 | 26S proteasome regulatory subunit 8 OS=Homo sapiens OX=9606 GN=PSMC5 PE=1 SV=1 | 0.999 | 0.984200071 | 0.996 | 0.9 |
| Q14103-2 | Isoform 2 of Heterogeneous nuclear ribonucleoprotein D0 OS=Homo sapiens OX=9606 GN=HNRNPD | 1.158 | 0.765235202 | 0.963 | 0.8 |
| P62753 | 40S ribosomal protein S6 OS=Homo sapiens OX=9606 GN=RPS6 PE=1 SV=1 | 1.106 | 0.750023665 | 1.146 | 0.7 |
| P08648 | Integrin alpha-5 OS=Homo sapiens OX=9606 GN=ITGA5 PE=1 SV=2 | 0.934 | 0.765235202 | 1.085 | 0.7 |
| P52907 | F-actin-capping protein subunit alpha-1 OS=Homo sapiens OX=9606 GN=CAPZA1 PE=1 SV=3 | 1.009 | 0.889482529 | 1.009 | 0.9 |
| Q9NQC3-2 | Isoform B of Reticulon-4 OS=Homo sapiens OX=9606 GN=RTN4 | 0.976 | 0.902766821 | 0.962 | 0.8 |
| Q7L576 | Cytoplasmic FMR1-interacting protein 1 OS=Homo sapiens OX=9606 GN=CYFIP1 PE=1 SV=1 | 1.02 | 0.766667967 | 0.958 | 0.7 |
| Q9Y224 | RNA transcription, translation and transport factor protein OS=Homo sapiens OX=9606 GN=RTRAF PE=1 SV=1 | 0.988 | 0.951324063 | 0.999 | 0.9 |
| C9JZR2 | Catenin delta-1 OS=Homo sapiens OX=9606 GN=CTNND1 PE=1 SV=2 | 0.987 | 0.765235202 | 0.922 | 0.7 |
| A5A3E0 | POTE ankyrin domain family member F OS=Homo sapiens OX=9606 GN=POTEF PE=1 SV=2 | 1.014 | 0.906273459 | 0.956 | 0.7 |
| P47755 | F-actin-capping protein subunit alpha-2 OS=Homo sapiens OX=9606 GN=CAPZA2 PE=1 SV=3 | 0.972 | 0.765235202 | 0.967 | 0.7 |
| P52566 | Rho GDP-dissociation inhibitor 2 OS=Homo sapiens OX=9606 GN=ARHGDIB PE=1 SV=3 | 0.979 | 0.873607677 | 1.026 | 0.7 |
| P23921 | Ribonucleoside-diphosphate reductase large subunit OS=Homo sapiens OX=9606 GN=RRM1 PE=1 SV=1 | 0.969 | 0.80706255 | 1.029 | 0.7 |
| P52788 | Spermine synthase OS=Homo sapiens OX=9606 GN=SMS PE=1 SV=2 | 0.958 | 0.750023665 | 0.94 | 0.7 |
| P48163 | NADP-dependent malic enzyme OS=Homo sapiens OX=9606 GN=ME1 PE=1 SV=1 | 1.003 | 0.957217311 | 0.984 | 0.7 |
| O95782 | AP-2 complex subunit alpha-1 OS=Homo sapiens OX=9606 GN=AP2A1 PE=1 SV=3 | 1.008 | 0.766667967 | 0.982 | 0.7 |
| Q9BQG0 | Myb-binding protein 1A OS=Homo sapiens OX=9606 GN=MYBBP1A PE=1 SV=2 | 0.998 | 0.972946678 | 0.761 | 0.7 |
| Q9Y262 | Eukaryotic translation initiation factor 3 subunit L OS=Homo sapiens OX=9606 GN=EIF3L PE=1 SV=1 | 0.992 | 0.891750829 | 0.991 | 0.9 |
| P30520 | Adenylosuccinate synthetase isozyme 2 OS=Homo sapiens OX=9606 GN=ADSS2 PE=1 SV=3 | 0.999 | 0.889450577 | 1.121 | 0.7 |
| B5MDF5 | GTP-binding nuclear protein Ran OS=Homo sapiens OX=9606 GN=RAN PE=1 SV=1 | 1.022 | 0.954224541 | 1.008 | 0.9 |
| Q16543 | Hsp90 co-chaperone Cdc37 OS=Homo sapiens OX=9606 GN=CDC37 PE=1 SV=1 | 0.975 | 0.792509319 | 1.037 | 0.7 |
| Q15029 | 116 kDa U5 small nuclear ribonucleoprotein component OS=Homo sapiens OX=9606 GN=EFTUD2 PE=1 SV=1 | 1.007 | 0.997448818 | 0.981 | 0.7 |
| Q99733 | Nucleosome assembly protein 1-like 4 OS=Homo sapiens OX=9606 GN=NAP1L4 PE=1 SV=1 | 1.011 | 0.889482529 | 1.006 | 0.9 |
| P12004 | Proliferating cell nuclear antigen OS=Homo sapiens OX=9606 GN=PCNA PE=1 SV=1 | 0.959 | 0.845656075 | 1.128 | 0.7 |
| P62879 | Guanine nucleotide-binding protein G(I)/G(S)/G(T) subunit beta-2 OS=Homo sapiens OX=9606 GN=GNB2 PE=1 SV=3 | 0.968 | 0.80706255 | 0.913 | 0.7 |
| Q5TCU3 | Tropomyosin beta chain OS=Homo sapiens OX=9606 GN=TPM2 PE=1 SV=1 | 0.881 | 0.754126086 | 1.196 | 0.7 |
| P20700 | Lamin-B1 OS=Homo sapiens OX=9606 GN=LMNB1 PE=1 SV=2 | 1.076 | 0.916035389 | 0.926 | 0.7 |
| P28482 | Mitogen-activated protein kinase 1 OS=Homo sapiens OX=9606 GN=MAPK1 PE=1 SV=3 | 0.97 | 0.765235202 | 1.004 | 0.9 |
| Q7Z6Z7 | E3 ubiquitin-protein ligase HUWE1 OS=Homo sapiens OX=9606 GN=HUWE1 PE=1 SV=3 | 0.988 | 0.983953845 | 0.97 | 0.7 |
| P55084 | Trifunctional enzyme subunit beta, mitochondrial OS=Homo sapiens OX=9606 GN=HADHB PE=1 SV=3 | 0.994 | 0.989102644 | 0.94 | 0.7 |
| A0AVT1 | Ubiquitin-like modifier-activating enzyme 6 OS=Homo sapiens OX=9606 GN=UBA6 PE=1 SV=1 | 1.001 | 0.911695794 | 0.959 | 0.7 |
| Q9ULV4 | Coronin-1C OS=Homo sapiens OX=9606 GN=CORO1C PE=1 SV=1 | 0.997 | 0.972073425 | 0.901 | 0.7 |
| P51149 | Ras-related protein Rab-7a OS=Homo sapiens OX=9606 GN=RAB7A PE=1 SV=1 | 0.969 | 0.825252681 | 1.009 | 0.9 |
| B5ME19 | Eukaryotic translation initiation factor 3 subunit C-like protein OS=Homo sapiens OX=9606 GN=EIF3CL PE=1 SV=1 | 1.019 | 0.774484855 | 1.047 | 0.7 |
| P23526 | Adenosylhomocysteinase OS=Homo sapiens OX=9606 GN=AHCY PE=1 SV=4 | 0.997 | 0.932716491 | 1.002 | 0.7 |
| P46060 | Ran GTPase-activating protein 1 OS=Homo sapiens OX=9606 GN=RANGAP1 PE=1 SV=1 | 1.041 | 0.750023665 | 0.963 | 0.9 |
| Q8N163 | Cell cycle and apoptosis regulator protein 2 OS=Homo sapiens OX=9606 GN=CCAR2 PE=1 SV=2 | 1 | 0.876510992 | 0.963 | 0.7 |
| P30084 | Enoyl-CoA hydratase, mitochondrial OS=Homo sapiens OX=9606 GN=ECHS1 PE=1 SV=4 | 0.965 | 0.80706255 | 0.962 | 0.7 |
| Q15459 | Splicing factor 3A subunit 1 OS=Homo sapiens OX=9606 GN=SF3A1 PE=1 SV=1 | 0.977 | 0.977023215 | 0.96 | 0.9 |
| P28066 | Proteasome subunit alpha type-5 OS=Homo sapiens OX=9606 GN=PSMA5 PE=1 SV=3 | 0.964 | 0.765235202 | 1 | 0.7 |
| P62873 | Guanine nucleotide-binding protein G(I)/G(S)/G(T) subunit beta-1 OS=Homo sapiens OX=9606 GN=GNB1 PE=1 SV=3 | 0.985 | 0.96247978 | 0.971 | 0.7 |
| P11279 | Lysosome-associated membrane glycoprotein 1 OS=Homo sapiens OX=9606 GN=LAMP1 PE=1 SV=3 | 0.987 | 0.80706255 | 0.884 | 0.7 |
| Q96KP4 | Cytosolic non-specific dipeptidase OS=Homo sapiens OX=9606 GN=CNDP2 PE=1 SV=2 | 0.983 | 0.765235202 | 0.983 | 0.7 |
| P55809 | Succinyl-CoA:3-ketoacid coenzyme A transferase 1, mitochondrial OS=Homo sapiens OX=9606 GN=OXCT1 PE=1 SV=1 | 1.109 | 0.96647353 | 0.959 | 0.7 |
| P50502 | Hsc70-interacting protein OS=Homo sapiens OX=9606 GN=ST13 PE=1 SV=2 | 0.982 | 0.820675725 | 1.007 | 0.7 |
| P26368 | Splicing factor U2AF 65 kDa subunit OS=Homo sapiens OX=9606 GN=U2AF2 PE=1 SV=4 | 0.974 | 0.939024443 | 0.967 | 0.7 |
| O14818 | Proteasome subunit alpha type-7 OS=Homo sapiens OX=9606 GN=PSMA7 PE=1 SV=1 | 0.954 | 0.866300162 | 1.03 | 0.8 |
| P09493-3 | Isoform 3 of Tropomyosin alpha-1 chain OS=Homo sapiens OX=9606 GN=TPM1 | 0.944 | 0.750023665 | 0.834 | 0.7 |
| P07686 | Beta-hexosaminidase subunit beta OS=Homo sapiens OX=9606 GN=HEXB PE=1 SV=3 | 0.97 | 0.889482529 | 0.926 | 0.7 |
| Q15075 | Early endosome antigen 1 OS=Homo sapiens OX=9606 GN=EEA1 PE=1 SV=2 | 1.017 | 0.983953845 | 0.902 | 0.7 |
| P15170 | Eukaryotic peptide chain release factor GTP-binding subunit ERF3A OS=Homo sapiens OX=9606 GN=GSPT1 PE=1 SV=1 | 1.015 | 0.866300162 | 1.047 | 0.7 |
| O43237 | Cytoplasmic dynein 1 light intermediate chain 2 OS=Homo sapiens OX=9606 GN=DYNC1LI2 PE=1 SV=1 | 1.053 | 0.858815413 | 0.914 | 0.7 |
| A0A087X1Z3 | Proteasome activator complex subunit 2 OS=Homo sapiens OX=9606 GN=PSME2 PE=1 SV=1 | 0.962 | 0.750023665 | 1.008 | 0.9 |
| O95865 | N(G),N(G)-dimethylarginine dimethylaminohydrolase 2 OS=Homo sapiens OX=9606 GN=DDAH2 PE=1 SV=1 | 0.883 | 0.765235202 | 0.918 | 0.7 |
| P18085 | ADP-ribosylation factor 4 OS=Homo sapiens OX=9606 GN=ARF4 PE=1 SV=3 | 0.965 | 0.750023665 | 1.044 | 0.7 |
| P11233 | Ras-related protein Ral-A OS=Homo sapiens OX=9606 GN=RALA PE=1 SV=1 | 0.969 | 0.853363211 | 0.841 | 0.7 |
| P61221 | ATP-binding cassette sub-family E member 1 OS=Homo sapiens OX=9606 GN=ABCE1 PE=1 SV=1 | 0.983 | 0.863796857 | 1.031 | 0.7 |
| O43143 | Pre-mRNA-splicing factor ATP-dependent RNA helicase DHX15 OS=Homo sapiens OX=9606 GN=DHX15 PE=1 SV=2 | 1.026 | 0.882454148 | 0.978 | 0.7 |
| P35606 | Coatomer subunit beta' OS=Homo sapiens OX=9606 GN=COPB2 PE=1 SV=2 | 1.002 | 0.869995963 | 0.984 | 0.9 |
| P09972 | Fructose-bisphosphate aldolase C OS=Homo sapiens OX=9606 GN=ALDOC PE=1 SV=2 | 0.963 | 0.765235202 | 1.067 | 0.7 |
| Q13310-3 | Isoform 3 of Polyadenylate-binding protein 4 OS=Homo sapiens OX=9606 GN=PABPC4 | 1.015 | 0.829390484 | 1.024 | 0.7 |
| O00154 | Cytosolic acyl coenzyme A thioester hydrolase OS=Homo sapiens OX=9606 GN=ACOT7 PE=1 SV=3 | 0.995 | 0.882968647 | 0.985 | 0.9 |
| O00231 | 26S proteasome non-ATPase regulatory subunit 11 OS=Homo sapiens OX=9606 GN=PSMD11 PE=1 SV=3 | 0.984 | 0.799266635 | 1.056 | 0.7 |
| Q02218 | 2-oxoglutarate dehydrogenase, mitochondrial OS=Homo sapiens OX=9606 GN=OGDH PE=1 SV=3 | 0.986 | 0.869094069 | 0.97 | 0.7 |
| P62906 | 60S ribosomal protein L10a OS=Homo sapiens OX=9606 GN=RPL10A PE=1 SV=2 | 1.023 | 0.868906519 | 1.086 | 0.7 |
| Q15404 | Ras suppressor protein 1 OS=Homo sapiens OX=9606 GN=RSU1 PE=1 SV=3 | 1.002 | 0.903226585 | 0.941 | 0.7 |

| | | | | | |
|---|---|---|---|---|---|
| A8MXP9 | Matrin-3 OS=Homo sapiens OX=9606 GN=MATR3 PE=1 SV=1 | 1.12 | 0.829390484 | 0.972 | 0.8 |
| Q15691 | Microtubule-associated protein RP/EB family member 1 OS=Homo sapiens OX=9606 GN=MAPRE1 PE=1 SV=3 | 1.028 | 0.765235202 | 1.046 | 0.9 |
| P29692 | Elongation factor 1-delta OS=Homo sapiens OX=9606 GN=EEF1D PE=1 SV=5 | 0.977 | 0.902016292 | 1.029 | 0.7 |
| E9PB61 | THO complex subunit 4 OS=Homo sapiens OX=9606 GN=ALYREF PE=1 SV=1 | 1.042 | 0.796408135 | 0.946 | 0.7 |
| P61088 | Ubiquitin-conjugating enzyme E2 N OS=Homo sapiens OX=9606 GN=UBE2N PE=1 SV=1 | 0.962 | 0.750023665 | 1.008 | 0.7 |
| Q5JRX3 | Presequence protease, mitochondrial OS=Homo sapiens OX=9606 GN=PITRM1 PE=1 SV=3 | 1.013 | 0.905369216 | 0.967 | 0.7 |
| O00151 | PDZ and LIM domain protein 1 OS=Homo sapiens OX=9606 GN=PDLIM1 PE=1 SV=4 | 1.024 | 0.835307991 | 0.932 | 0.7 |
| P19367 | Hexokinase-1 OS=Homo sapiens OX=9606 GN=HK1 PE=1 SV=3 | 0.96 | 0.765235202 | 0.951 | 0.9 |
| A0A0C4DGB5 | Calpain inhibitor OS=Homo sapiens OX=9606 GN=CAST PE=1 SV=1 | 1.022 | 0.953521381 | 1.023 | 0.9 |
| Q9BS26 | Endoplasmic reticulum resident protein 44 OS=Homo sapiens OX=9606 GN=ERP44 PE=1 SV=1 | 0.956 | 0.765235202 | 0.928 | 0.7 |
| O00148 | ATP-dependent RNA helicase DDX39A OS=Homo sapiens OX=9606 GN=DDX39A PE=1 SV=2 | 1 | 0.995582922 | 0.938 | 0.7 |
| P20618 | Proteasome subunit beta type-1 OS=Homo sapiens OX=9606 GN=PSMB1 PE=1 SV=2 | 1.001 | 0.916363276 | 0.938 | 0.7 |
| P21281 | V-type proton ATPase subunit B, brain isoform OS=Homo sapiens OX=9606 GN=ATP6V1B2 PE=1 SV=3 | 1.009 | 0.95980123 | 0.975 | 0.7 |
| P14324 | Farnesyl pyrophosphate synthase OS=Homo sapiens OX=9606 GN=FDPS PE=1 SV=4 | 0.99 | 0.809976184 | 1.033 | 0.9 |
| O00425 | Insulin-like growth factor 2 mRNA-binding protein 3 OS=Homo sapiens OX=9606 GN=IGF2BP3 PE=1 SV=2 | 0.999 | 0.925929947 | 1.013 | 0.8 |
| Q9Y266 | Nuclear migration protein nudC OS=Homo sapiens OX=9606 GN=NUDC PE=1 SV=1 | 1.029 | 0.887776944 | 1.07 | 0.7 |
| Q7Z406 | Myosin-14 OS=Homo sapiens OX=9606 GN=MYH14 PE=1 SV=2 | 1.263 | 0.750023665 | 0.993 | 0.9 |
| O95433 | Activator of 90 kDa heat shock protein ATPase homolog 1 OS=Homo sapiens OX=9606 GN=AHSA1 PE=1 SV=1 | 0.984 | 0.750023665 | 1.029 | 0.7 |
| Q13185 | Chromobox protein homolog 3 OS=Homo sapiens OX=9606 GN=CBX3 PE=1 SV=4 | 0.943 | 0.904500837 | 1 | 0.7 |
| P39687 | Acidic leucine-rich nuclear phosphoprotein 32 family member A OS=Homo sapiens OX=9606 GN=ANP32A PE=1 SV=1 | 0.969 | 0.835336742 | 0.943 | 0.7 |
| Q8NC51-3 | Isoform 3 of Plasminogen activator inhibitor 1 RNA-binding protein OS=Homo sapiens OX=9606 GN=SERBP1 | 1.106 | 0.996637279 | 1.08 | 0.8 |
| Q9Y281 | Cofilin-2 OS=Homo sapiens OX=9606 GN=CFL2 PE=1 SV=1 | 0.964 | 0.765235202 | 1.012 | 0.9 |
| P07384 | Calpain-1 catalytic subunit OS=Homo sapiens OX=9606 GN=CAPN1 PE=1 SV=1 | 0.964 | 0.902016292 | 0.994 | 0.7 |
| J3KR97 | Tubulin-specific chaperone D OS=Homo sapiens OX=9606 GN=TBCD PE=1 SV=1 | 1.095 | 0.798032881 | 0.915 | 0.7 |
| P49748 | Very long-chain specific acyl-CoA dehydrogenase, mitochondrial OS=Homo sapiens OX=9606 GN=ACADVL PE=1 SV=1 | 1.024 | 0.932424612 | 0.957 | 0.7 |
| P32969 | 60S ribosomal protein L9 OS=Homo sapiens OX=9606 GN=RPL9 PE=1 SV=1 | 1.107 | 0.750023665 | 1.13 | 0.7 |
| P25789 | Proteasome subunit alpha type-4 OS=Homo sapiens OX=9606 GN=PSMA4 PE=1 SV=1 | 0.994 | 0.765235202 | 1.001 | 0.8 |
| P17066 | Heat shock 70 kDa protein 6 OS=Homo sapiens OX=9606 GN=HSPA6 PE=1 SV=2 | 0.925 | 0.876510992 | 0.933 | 0.7 |
| Q16181 | Septin-7 OS=Homo sapiens OX=9606 GN=SEPTIN7 PE=1 SV=2 | 1.04 | 0.882454148 | 0.926 | 0.7 |
| P60900 | Proteasome subunit alpha type-6 OS=Homo sapiens OX=9606 GN=PSMA6 PE=1 SV=1 | 0.983 | 0.913279108 | 1.002 | 0.8 |
| P02751 | Fibronectin OS=Homo sapiens OX=9606 GN=FN1 PE=1 SV=5 | 0.978 | 0.882454148 | 0.912 | 0.8 |
| Q92621 | Nuclear pore complex protein Nup205 OS=Homo sapiens OX=9606 GN=NUP205 PE=1 SV=3 | 1.002 | 0.870355966 | 0.86 | 0.7 |
| P30040 | Endoplasmic reticulum resident protein 29 OS=Homo sapiens OX=9606 GN=ERP29 PE=1 SV=4 | 0.965 | 0.750023665 | 1.009 | 0.7 |
| Q9H4A4 | Aminopeptidase B OS=Homo sapiens OX=9606 GN=RNPEP PE=1 SV=2 | 0.991 | 0.95980123 | 0.981 | 0.7 |
| A0A087X2I1 | 26S proteasome regulatory subunit 10B OS=Homo sapiens OX=9606 GN=PSMC6 PE=1 SV=1 | 1.01 | 0.703015003 | 1 | 0.9 |
| Q9HC38 | Glyoxalase domain-containing protein 4 OS=Homo sapiens OX=9606 GN=GLOD4 PE=1 SV=1 | 0.968 | 0.895384791 | 0.947 | 0.7 |
| Q01105-2 | Isoform 2 of Protein SET OS=Homo sapiens OX=9606 GN=SET | 0.996 | 0.869756587 | 0.973 | 0.7 |
| P27695 | DNA-(apurinic or apyrimidinic site) endonuclease OS=Homo sapiens OX=9606 GN=APEX1 PE=1 SV=2 | 1.02 | 0.925981427 | 0.938 | 0.7 |
| P60228 | Eukaryotic translation initiation factor 3 subunit E OS=Homo sapiens OX=9606 GN=EIF3E PE=1 SV=1 | 1.009 | 0.972946678 | 0.988 | 0.8 |
| P40925 | Malate dehydrogenase, cytoplasmic OS=Homo sapiens OX=9606 GN=MDH1 PE=1 SV=4 | 0.969 | 0.750023665 | 1.015 | 0.7 |
| O15143 | Actin-related protein 2/3 complex subunit 1B OS=Homo sapiens OX=9606 GN=ARPC1B PE=1 SV=3 | 0.971 | 0.849526028 | 0.976 | 0.7 |
| Q15008 | 26S proteasome non-ATPase regulatory subunit 6 OS=Homo sapiens OX=9606 GN=PSMD6 PE=1 SV=1 | 0.997 | 0.889450577 | 1.007 | 0.9 |
| Q9UBE0 | SUMO-activating enzyme subunit 1 OS=Homo sapiens OX=9606 GN=SAE1 PE=1 SV=1 | 0.968 | 0.750023665 | 0.916 | 0.7 |
| P07942 | Laminin subunit beta-1 OS=Homo sapiens OX=9606 GN=LAMB1 PE=1 SV=2 | 0.995 | 0.984805107 | 0.994 | 0.9 |
| Q9Y3I0 | RNA-splicing ligase RtcB homolog OS=Homo sapiens OX=9606 GN=RTCB PE=1 SV=1 | 1.045 | 0.750023665 | 0.994 | 0.7 |
| Q9BXJ9 | N-alpha-acetyltransferase 15, NatA auxiliary subunit OS=Homo sapiens OX=9606 GN=NAA15 PE=1 SV=1 | 0.987 | 0.951693691 | 1.008 | 0.8 |
| Q8NC51 | Plasminogen activator inhibitor 1 RNA-binding protein OS=Homo sapiens OX=9606 GN=SERBP1 PE=1 SV=2 | 1.137 | 0.805488634 | 1.099 | 0.7 |
| P40763 | Signal transducer and activator of transcription 3 OS=Homo sapiens OX=9606 GN=STAT3 PE=1 SV=2 | 0.989 | 0.774484855 | 1.015 | 0.8 |
| P62241 | 40S ribosomal protein S8 OS=Homo sapiens OX=9606 GN=RPS8 PE=1 SV=2 | 1.032 | 0.99634713 | 1.114 | 0.7 |
| Q71DI3 | Histone H3.2 OS=Homo sapiens OX=9606 GN=H3C15 PE=1 SV=3 | 0.759 | 0.761475165 | 0.817 | 0.7 |
| P11234 | Ras-related protein Ral-B OS=Homo sapiens OX=9606 GN=RALB PE=1 SV=1 | 0.986 | 0.927756817 | 0.89 | 0.7 |
| Q9BZZ5 | Apoptosis inhibitor 5 OS=Homo sapiens OX=9606 GN=API5 PE=1 SV=3 | 0.996 | 0.963059765 | 0.967 | 0.7 |
| O76003 | Glutaredoxin-3 OS=Homo sapiens OX=9606 GN=GLRX3 PE=1 SV=2 | 0.976 | 0.826213663 | 1.037 | 0.7 |
| Q15056 | Eukaryotic translation initiation factor 4H OS=Homo sapiens OX=9606 GN=EIF4H PE=1 SV=5 | 0.957 | 0.902766821 | 0.957 | 0.7 |
| Q15233 | Non-POU domain-containing octamer-binding protein OS=Homo sapiens OX=9606 GN=NONO PE=1 SV=4 | 1.151 | 0.765235202 | 1.034 | 0.7 |
| Q9NVD7 | Alpha-parvin OS=Homo sapiens OX=9606 GN=PARVA PE=1 SV=1 | 0.988 | 0.994361756 | 0.934 | 0.7 |
| P30086 | Phosphatidylethanolamine-binding protein 1 OS=Homo sapiens OX=9606 GN=PEBP1 PE=1 SV=3 | 0.974 | 0.750023665 | 0.974 | 0.7 |
| P40261 | Nicotinamide N-methyltransferase OS=Homo sapiens OX=9606 GN=NNMT PE=1 SV=1 | 0.987 | 0.996637279 | 0.967 | 0.8 |
| Q7L2H7 | Eukaryotic translation initiation factor 3 subunit M OS=Homo sapiens OX=9606 GN=EIF3M PE=1 SV=1 | 1.021 | 0.765235202 | 1.049 | 0.7 |
| O00629 | Importin subunit alpha-3 OS=Homo sapiens OX=9606 GN=KPNA4 PE=1 SV=1 | 1.019 | 0.766667967 | 0.987 | 0.9 |
| Q16777 | Histone H2A type 2-C OS=Homo sapiens OX=9606 GN=H2AC20 PE=1 SV=4 | 0.867 | 0.889482529 | 0.939 | 0.7 |
| G5EA09 | Syndecan binding protein (Syntenin), isoform CRA_a OS=Homo sapiens OX=9606 GN=SDCBP PE=1 SV=1 | 1.045 | 0.80706255 | 1.041 | 0.7 |
| P31153 | S-adenosylmethionine synthase isoform type-2 OS=Homo sapiens OX=9606 GN=MAT2A PE=1 SV=1 | 0.96 | 0.765235202 | 1.044 | 0.7 |
| Q9NTK5 | Obg-like ATPase 1 OS=Homo sapiens OX=9606 GN=OLA1 PE=1 SV=2 | 0.994 | 0.924111368 | 1.026 | 0.7 |
| B1AK88 | F-actin-capping protein subunit beta OS=Homo sapiens OX=9606 GN=CAPZB PE=1 SV=1 | 0.974 | 0.829390484 | 0.99 | 0.9 |
| O14979 | Heterogeneous nuclear ribonucleoprotein D-like OS=Homo sapiens OX=9606 GN=HNRNPDL PE=1 SV=3 | 1.025 | 0.945142068 | 1.008 | 0.9 |
| P31949 | Protein S100-A11 OS=Homo sapiens OX=9606 GN=S100A11 PE=1 SV=2 | 0.958 | 0.765235202 | 0.992 | 0.9 |
| Q13724 | Mannosyl-oligosaccharide glucosidase OS=Homo sapiens OX=9606 GN=MOGS PE=1 SV=5 | 1.037 | 0.754126086 | 0.956 | 0.7 |
| P61163 | Alpha-centractin OS=Homo sapiens OX=9606 GN=ACTR1A PE=1 SV=1 | 0.99 | 0.869995963 | 1.001 | 0.8 |
| O15355 | Protein phosphatase 1G OS=Homo sapiens OX=9606 GN=PPM1G PE=1 SV=1 | 1.004 | 0.983953845 | 0.995 | 0.9 |
| P62917 | 60S ribosomal protein L8 OS=Homo sapiens OX=9606 GN=RPL8 PE=1 SV=2 | 1.064 | 0.765235202 | 1.099 | 0.7 |
| P42785 | Lysosomal Pro-X carboxypeptidase OS=Homo sapiens OX=9606 GN=PRCP PE=1 SV=1 | 0.936 | 0.750023665 | 0.884 | 0.7 |
| O95340 | Bifunctional 3'-phosphoadenosine 5'-phosphosulfate synthase 2 OS=Homo sapiens OX=9606 GN=PAPSS2 PE=1 SV=2 | 1.021 | 0.703015003 | 1.124 | 0.7 |
| A0A0J9YYL3 | 60 kDa poly(U)-binding-splicing factor (Fragment) OS=Homo sapiens OX=9606 GN=PUF60 PE=1 SV=1 | 1.014 | 0.983953845 | 0.97 | 0.8 |
| Q9UNM6 | 26S proteasome non-ATPase regulatory subunit 13 OS=Homo sapiens OX=9606 GN=PSMD13 PE=1 SV=2 | 0.966 | 0.815069882 | 1.012 | 0.7 |
| O00232 | 26S proteasome non-ATPase regulatory subunit 12 OS=Homo sapiens OX=9606 GN=PSMD12 PE=1 SV=3 | 0.99 | 0.937953124 | 0.978 | 0.8 |
| Q86UP2 | Kinectin OS=Homo sapiens OX=9606 GN=KTN1 PE=1 SV=1 | 1.017 | 0.996637279 | 0.897 | 0.7 |
| J3KRX5 | 60S ribosomal protein L17 (Fragment) OS=Homo sapiens OX=9606 GN=RPL17 PE=1 SV=1 | 1.051 | 0.779783772 | 1.097 | 0.7 |
| P30740 | Leukocyte elastase inhibitor OS=Homo sapiens OX=9606 GN=SERPINB1 PE=1 SV=1 | 1.006 | 0.983953845 | 0.987 | 0. |
| P62191 | 26S proteasome regulatory subunit 4 OS=Homo sapiens OX=9606 GN=PSMC1 PE=1 SV=1 | 0.978 | 0.765235202 | 1.033 | 0.7 |
| P52292 | Importin subunit alpha-1 OS=Homo sapiens OX=9606 GN=KPNA2 PE=1 SV=1 | 1.017 | 0.95980123 | 1.233 | 0.7 |
| Q9Y3F4 | Serine-threonine kinase receptor-associated protein OS=Homo sapiens OX=9606 GN=STRAP PE=1 SV=1 | 1.021 | 0.756935672 | 1.095 | 0.7 |
| P32119 | Peroxiredoxin-2 OS=Homo sapiens OX=9606 GN=PRDX2 PE=1 SV=5 | 1.02 | 0.860909277 | 0.999 | 0.9 |
| Q14019 | Coactosin-like protein OS=Homo sapiens OX=9606 GN=COTL1 PE=1 SV=3 | 0.976 | 0.901934028 | 0.998 | 0.9 |
| P62829 | 60S ribosomal protein L23 OS=Homo sapiens OX=9606 GN=RPL23 PE=1 SV=1 | 1.103 | 0.703015003 | 1.034 | 0.7 |
| P62280 | 40S ribosomal protein S11 OS=Homo sapiens OX=9606 GN=RPS11 PE=1 SV=3 | 1.07 | 0.750023665 | 1.111 | 0.7 |
| J3KR44 | Ubiquitin thioesterase OS=Homo sapiens OX=9606 GN=OTUB1 PE=1 SV=2 | 1.045 | 0.924111368 | 0.967 | 0.7 |
| P04181 | Ornithine aminotransferase, mitochondrial OS=Homo sapiens OX=9606 GN=OAT PE=1 SV=1 | 1.03 | 0.758950588 | 1.068 | 0.7 |
| P54577 | Tyrosine--tRNA ligase, cytoplasmic OS=Homo sapiens OX=9606 GN=YARS1 PE=1 SV=4 | 1.011 | 0.984200071 | 1.014 | 0.7 |
| P17301 | Integrin alpha-2 OS=Homo sapiens OX=9606 GN=ITGA2 PE=1 SV=1 | 0.963 | 0.829390484 | 0.992 | 0.9 |
| O94973 | AP-2 complex subunit alpha-2 OS=Homo sapiens OX=9606 GN=AP2A2 PE=1 SV=2 | 0.973 | 0.827871327 | 0.94 | 0.7 |
| A0A2R8Y811 | 40S ribosomal protein S14 (Fragment) OS=Homo sapiens OX=9606 GN=RPS14 PE=1 SV=1 | 1.033 | 0.750023665 | 1.079 | 0.7 |
| P21589 | 5'-nucleotidase OS=Homo sapiens OX=9606 GN=NT5E PE=1 SV=1 | 0.931 | 0.776314646 | 0.841 | 0.7 |
| P00568 | Adenylate kinase isoenzyme 1 OS=Homo sapiens OX=9606 GN=AK1 PE=1 SV=3 | 0.981 | 0.824559577 | 0.969 | 0.7 |
| P62888 | 60S ribosomal protein L30 OS=Homo sapiens OX=9606 GN=RPL30 PE=1 SV=2 | 1.063 | 0.750023665 | 1.07 | 0.7 |
| P06703 | Protein S100-A6 OS=Homo sapiens OX=9606 GN=S100A6 PE=1 SV=1 | 0.978 | 0.985324977 | 1.025 | 0.9 |

| | | | | | |
|---|---|---|---|---|---|
| P50570 | Dynamin-2 OS=Homo sapiens OX=9606 GN=DNM2 PE=1 SV=2 | 1.02 | 0.859846322 | 0.931 | 0.7 |
| P21291 | Cysteine and glycine-rich protein 1 OS=Homo sapiens OX=9606 GN=CSRP1 PE=1 SV=3 | 0.969 | 0.942843201 | 0.997 | 0.8 |
| P14550 | Aldo-keto reductase family 1 member A1 OS=Homo sapiens OX=9606 GN=AKR1A1 PE=1 SV=3 | 0.995 | 0.991211283 | 1.004 | 0.9 |
| H7BYY1 | Tropomyosin 1 (Alpha), isoform CRA_m OS=Homo sapiens OX=9606 GN=TPM1 PE=1 SV=1 | 0.985 | 0.983953845 | 0.904 | 0.7 |
| P54727 | UV excision repair protein RAD23 homolog B OS=Homo sapiens OX=9606 GN=RAD23B PE=1 SV=1 | 0.933 | 0.840158726 | 1.017 | 0.7 |
| P68431 | Histone H3.1 OS=Homo sapiens OX=9606 GN=H3C1 PE=1 SV=2 | 0.87 | 0.766667967 | 0.858 | 0.7 |
| Q9BRX8 | Peroxiredoxin-like 2A OS=Homo sapiens OX=9606 GN=PRXL2A PE=1 SV=3 | 1.052 | 0.849526028 | 1.164 | 0.7 |
| Q9Y617 | Phosphoserine aminotransferase OS=Homo sapiens OX=9606 GN=PSAT1 PE=1 SV=2 | 1.026 | 0.942678074 | 1.054 | 0.7 |
| P52565 | Rho GDP-dissociation inhibitor 1 OS=Homo sapiens OX=9606 GN=ARHGDIA PE=1 SV=3 | 0.98 | 0.978562349 | 1.05 | 0.8 |
| Q5SRN7 | HLA class I histocompatibility antigen, A alpha chain OS=Homo sapiens OX=9606 GN=HLA-A PE=1 SV=2 | 0.988 | 0.916363276 | 0.885 | 0.8 |
| J3QRU1 | Tyrosine-protein kinase OS=Homo sapiens OX=9606 GN=YES1 PE=1 SV=1 | 0.949 | 0.794788863 | 0.956 | 0.7 |
| Q9H8L6 | Multimerin-2 OS=Homo sapiens OX=9606 GN=MMRN2 PE=1 SV=2 | 0.96 | 0.889482529 | 0.915 | 0.7 |
| P84243 | Histone H3.3 OS=Homo sapiens OX=9606 GN=H3-3A PE=1 SV=2 | 0.773 | 0.790136102 | 0.957 | 0.8 |
| Q6IBS0 | Twinfilin-2 OS=Homo sapiens OX=9606 GN=TWF2 PE=1 SV=2 | 0.976 | 0.809976184 | 0.983 | 0.8 |
| P11766 | Alcohol dehydrogenase class-3 OS=Homo sapiens OX=9606 GN=ADH5 PE=1 SV=4 | 1.006 | 0.994361756 | 1.007 | 0.9 |
| P09960 | Leukotriene A-4 hydrolase OS=Homo sapiens OX=9606 GN=LTA4H PE=1 SV=2 | 1.007 | 0.985324977 | 0.934 | 0.7 |
| Q53Z42 | HLA class I antigen OS=Homo sapiens OX=9606 GN=HLA-A PE=1 SV=1 | 0.941 | 0.80706255 | 0.88 | 0.7 |
| P38159 | RNA-binding motif protein, X chromosome OS=Homo sapiens OX=9606 GN=RBMX PE=1 SV=3 | 1.097 | 0.939024443 | 0.888 | 0.7 |
| P17813 | Endoglin OS=Homo sapiens OX=9606 GN=ENG PE=1 SV=2 | 0.963 | 0.765235202 | 1.128 | 0.7 |
| Q9NR45 | Sialic acid synthase OS=Homo sapiens OX=9606 GN=NANS PE=1 SV=2 | 1.008 | 0.913279108 | 0.984 | 0.8 |
| P06132 | Uroporphyrinogen decarboxylase OS=Homo sapiens OX=9606 GN=UROD PE=1 SV=2 | 0.925 | 0.703015003 | 0.961 | 0.7 |
| O00487 | 26S proteasome non-ATPase regulatory subunit 14 OS=Homo sapiens OX=9606 GN=PSMD14 PE=1 SV=1 | 0.963 | 0.750023665 | 1.032 | 0.7 |
| P19623 | Spermidine synthase OS=Homo sapiens OX=9606 GN=SRM PE=1 SV=1 | 0.976 | 0.776314646 | 1.043 | 0.7 |
| A0A087WV22 | 60S ribosomal protein L10 OS=Homo sapiens OX=9606 GN=RPL10 PE=1 SV=1 | 1.25 | 0.765235202 | 1.209 | 0.7 |
| P10412 | Histone H1.4 OS=Homo sapiens OX=9606 GN=H1-4 PE=1 SV=2 | 1.168 | 0.825275231 | 0.985 | 0.7 |
| P51991 | Heterogeneous nuclear ribonucleoprotein A3 OS=Homo sapiens OX=9606 GN=HNRNPA3 PE=1 SV=2 | 1.046 | 0.925929947 | 0.945 | 0.7 |
| Q15393 | Splicing factor 3B subunit 3 OS=Homo sapiens OX=9606 GN=SF3B3 PE=1 SV=4 | 1.056 | 0.765235202 | 0.977 | 0.8 |
| Q16401 | 26S proteasome non-ATPase regulatory subunit 5 OS=Homo sapiens OX=9606 GN=PSMD5 PE=1 SV=3 | 0.999 | 0.972946678 | 0.937 | 0.7 |
| B4DUT8 | Calponin OS=Homo sapiens OX=9606 GN=CNN2 PE=1 SV=1 | 1.025 | 0.750023665 | 0.91 | 0.7 |
| P61224 | Ras-related protein Rap-1b OS=Homo sapiens OX=9606 GN=RAP1B PE=1 SV=1 | 1.006 | 0.963059765 | 0.968 | 0.7 |
| O00625 | Pirin OS=Homo sapiens OX=9606 GN=PIR PE=1 SV=1 | 1.009 | 0.876510992 | 1.01 | 0.8 |
| Q96M27 | Protein PRRC1 OS=Homo sapiens OX=9606 GN=PRRC1 PE=1 SV=1 | 1.045 | 0.750023665 | 0.972 | 0.9 |
| E7EX17 | Eukaryotic translation initiation factor 4B OS=Homo sapiens OX=9606 GN=EIF4B PE=1 SV=1 | 1.03 | 0.765235202 | 1.077 | 0.7 |
| Q9UHD8 | Septin-9 OS=Homo sapiens OX=9606 GN=SEPTIN9 PE=1 SV=2 | 1.074 | 0.765235202 | 0.974 | 0.8 |
| P13861 | cAMP-dependent protein kinase type II-alpha regulatory subunit OS=Homo sapiens OX=9606 GN=PRKAR2A PE=1 SV=2 | 1.023 | 0.961735044 | 0.952 | 0.7 |
| P61026 | Ras-related protein Rab-10 OS=Homo sapiens OX=9606 GN=RAB10 PE=1 SV=1 | 0.968 | 0.750023665 | 0.973 | 0.7 |
| A0A0G2JR96 | Cytoplasmic FMR1-interacting protein 1 (Fragment) OS=Homo sapiens OX=9606 GN=CYFIP1 PE=1 SV=4 | 0.992 | 0.931793839 | 0.715 | 0.8 |
| P39019 | 40S ribosomal protein S19 OS=Homo sapiens OX=9606 GN=RPS19 PE=1 SV=2 | 1.087 | 0.761475165 | 1.181 | 0.7 |
| O00303 | Eukaryotic translation initiation factor 3 subunit F OS=Homo sapiens OX=9606 GN=EIF3F PE=1 SV=1 | 0.981 | 0.921087329 | 1.016 | 0.7 |
| Q9UDY4 | DnaJ homolog subfamily B member 4 OS=Homo sapiens OX=9606 GN=DNAJB4 PE=1 SV=1 | 1.016 | 0.899793601 | 1.032 | 0.8 |
| O75131 | Copine-3 OS=Homo sapiens OX=9606 GN=CPNE3 PE=1 SV=1 | 0.946 | 0.756287988 | 0.983 | 0.7 |
| Q9Y2A7 | Nck-associated protein 1 OS=Homo sapiens OX=9606 GN=NCKAP1 PE=1 SV=1 | 1.033 | 0.785314375 | 0.964 | 0.7 |
| P62834 | Ras-related protein Rap-1A OS=Homo sapiens OX=9606 GN=RAP1A PE=1 SV=1 | 1.02 | 0.986297807 | 0.999 | 0.8 |
| Q07960 | Rho GTPase-activating protein 1 OS=Homo sapiens OX=9606 GN=ARHGAP1 PE=1 SV=1 | 1.008 | 0.939784289 | 1.025 | 0.9 |
| P42224 | Signal transducer and activator of transcription 1-alpha/beta OS=Homo sapiens OX=9606 GN=STAT1 PE=1 SV=2 | 1.021 | 0.765937763 | 0.915 | 0.7 |
| P37840 | Alpha-synuclein OS=Homo sapiens OX=9606 GN=SNCA PE=1 SV=1 | 1.082 | 0.754126086 | 1.006 | 0.7 |
| Q9Y305 | Acyl-coenzyme A thioesterase 9, mitochondrial OS=Homo sapiens OX=9606 GN=ACOT9 PE=1 SV=2 | 0.956 | 0.908555656 | 0.968 | 0.7 |
| G3V203 | 60S ribosomal protein L18 OS=Homo sapiens OX=9606 GN=RPL18 PE=1 SV=1 | 1.009 | 0.985330338 | 1.051 | 0.8 |
| P50570-4 | Isoform 4 of Dynamin-2 OS=Homo sapiens OX=9606 GN=DNM2 | 1.067 | 0.840158726 | 0.96 | 0.7 |
| P21964 | Catechol O-methyltransferase OS=Homo sapiens OX=9606 GN=COMT PE=1 SV=2 | 0.999 | 0.954224541 | 0.916 | 0.7 |
| P61758 | Prefoldin subunit 3 OS=Homo sapiens OX=9606 GN=VBP1 PE=1 SV=4 | 0.96 | 0.765235202 | 0.998 | 0.8 |
| Q9NUV9 | GTPase IMAP family member 4 OS=Homo sapiens OX=9606 GN=GIMAP4 PE=1 SV=1 | 0.927 | 0.765235202 | 0.966 | 0.7 |
| P16949 | Stathmin OS=Homo sapiens OX=9606 GN=STMN1 PE=1 SV=3 | 0.942 | 0.827871327 | 0.962 | 0.7 |
| D6RER5 | Septin-11 OS=Homo sapiens OX=9606 GN=SEPTIN11 PE=1 SV=1 | 0.996 | 0.978562349 | 0.897 | 0.7 |
| H7BZJ3 | Protein disulfide-isomerase A3 (Fragment) OS=Homo sapiens OX=9606 GN=PDIA3 PE=1 SV=1 | 0.814 | 0.780884711 | 0.911 | 0.7 |
| P23368 | NAD-dependent malic enzyme, mitochondrial OS=Homo sapiens OX=9606 GN=ME2 PE=1 SV=1 | 1.013 | 0.970601862 | 0.957 | 0.7 |
| Q9NQW7 | Xaa-Pro aminopeptidase 1 OS=Homo sapiens OX=9606 GN=XPNPEP1 PE=1 SV=3 | 0.966 | 0.750023665 | 0.96 | 0.7 |
| P16402 | Histone H1.3 OS=Homo sapiens OX=9606 GN=H1-3 PE=1 SV=2 | 1.083 | 0.848368819 | 0.98 | 0.7 |
| P30626 | Sorcin OS=Homo sapiens OX=9606 GN=SRI PE=1 SV=1 | 0.978 | 0.765235202 | 0.975 | 0.7 |
| P30048 | Thioredoxin-dependent peroxide reductase, mitochondrial OS=Homo sapiens OX=9606 GN=PRDX3 PE=1 SV=3 | 0.97 | 0.750023665 | 0.967 | 0.7 |
| Q9H2U2 | Inorganic pyrophosphatase 2, mitochondrial OS=Homo sapiens OX=9606 GN=PPA2 PE=1 SV=2 | 0.97 | 0.809976184 | 0.946 | 0.7 |
| O15523 | ATP-dependent RNA helicase DDX3Y OS=Homo sapiens OX=9606 GN=DDX3Y PE=1 SV=2 | 1.004 | 0.860909277 | 0.787 | 0.8 |
| P59998 | Actin-related protein 2/3 complex subunit 4 OS=Homo sapiens OX=9606 GN=ARPC4 PE=1 SV=3 | 0.96 | 0.750023665 | 0.969 | 0.8 |
| P16403 | Histone H1.2 OS=Homo sapiens OX=9606 GN=H1-2 PE=1 SV=2 | 1.04 | 0.899987172 | 0.948 | 0.7 |
| Q13148 | TAR DNA-binding protein 43 OS=Homo sapiens OX=9606 GN=TARDBP PE=1 SV=1 | 1 | 0.88308648 | 0.904 | 0.7 |
| P11047 | Laminin subunit gamma-1 OS=Homo sapiens OX=9606 GN=LAMC1 PE=1 SV=3 | 0.979 | 0.996578046 | 0.945 | 0.7 |
| P11177 | Pyruvate dehydrogenase E1 component subunit beta, mitochondrial OS=Homo sapiens OX=9606 GN=PDHB PE=1 SV=3 | 0.98 | 0.900915273 | 0.952 | 0.7 |
| P25788 | Proteasome subunit alpha type-3 OS=Homo sapiens OX=9606 GN=PSMA3 PE=1 SV=2 | 0.946 | 0.750023665 | 0.986 | 0.7 |
| Q96JJ7 | Protein disulfide-isomerase TMX3 OS=Homo sapiens OX=9606 GN=TMX3 PE=1 SV=2 | 1.012 | 0.985324977 | 0.992 | 0.7 |
| O15371 | Eukaryotic translation initiation factor 3 subunit D OS=Homo sapiens OX=9606 GN=EIF3D PE=1 SV=1 | 1.026 | 0.765235202 | 1.002 | 0.8 |
| Q12769 | Nuclear pore complex protein Nup160 OS=Homo sapiens OX=9606 GN=NUP160 PE=1 SV=3 | 1.035 | 0.902016292 | 0.898 | 0.7 |
| P12236 | ADP/ATP translocase 3 OS=Homo sapiens OX=9606 GN=SLC25A6 PE=1 SV=4 | 0.975 | 0.916936955 | 0.883 | 0.7 |
| P12270 | Nucleoprotein TPR OS=Homo sapiens OX=9606 GN=TPR PE=1 SV=3 | 1.025 | 0.830179531 | 0.942 | 0.7 |
| I3L3H2 | RNA helicase OS=Homo sapiens OX=9606 GN=EIF4A3 PE=1 SV=2 | 0.978 | 0.924631778 | 1.001 | 0.8 |
| P07858 | Cathepsin B OS=Homo sapiens OX=9606 GN=CTSB PE=1 SV=3 | 0.947 | 0.756935672 | 0.907 | 0.7 |
| Q9UJZ1 | Stomatin-like protein 2, mitochondrial OS=Homo sapiens OX=9606 GN=STOML2 PE=1 SV=1 | 0.978 | 0.983953845 | 0.893 | 0.7 |
| P28070 | Proteasome subunit beta type-4 OS=Homo sapiens OX=9606 GN=PSMB4 PE=1 SV=4 | 0.901 | 0.765235202 | 0.978 | 0.7 |
| P54725 | UV excision repair protein RAD23 homolog A OS=Homo sapiens OX=9606 GN=RAD23A PE=1 SV=1 | 1.023 | 0.829390484 | 0.948 | 0.7 |
| P62491 | Ras-related protein Rab-11A OS=Homo sapiens OX=9606 GN=RAB11A PE=1 SV=3 | 1.132 | 0.754126086 | 0.786 | 0.7 |
| Q5SSJ5 | Heterochromatin protein 1-binding protein 3 OS=Homo sapiens OX=9606 GN=HP1BP3 PE=1 SV=1 | 1.017 | 0.912083887 | 0.947 | 0.9 |
| P62266 | 40S ribosomal protein S23 OS=Homo sapiens OX=9606 GN=RPS23 PE=1 SV=3 | 1.06 | 0.750023665 | 1.138 | 0.7 |
| Q14847 | LIM and SH3 domain protein 1 OS=Homo sapiens OX=9606 GN=LASP1 PE=1 SV=2 | 0.986 | 0.996637279 | 0.982 | 0.8 |
| P61201 | COP9 signalosome complex subunit 2 OS=Homo sapiens OX=9606 GN=COPS2 PE=1 SV=1 | 1.001 | 0.916363276 | 0.988 | 0.7 |
| P46783 | 40S ribosomal protein S10 OS=Homo sapiens OX=9606 GN=RPS10 PE=1 SV=1 | 1.077 | 0.765235202 | 1.18 | 0.7 |
| P53634 | Dipeptidyl peptidase 1 OS=Homo sapiens OX=9606 GN=CTSC PE=1 SV=2 | 0.933 | 0.750023665 | 0.871 | 0.7 |
| P0CG38 | POTE ankyrin domain family member I OS=Homo sapiens OX=9606 GN=POTEI PE=3 SV=1 | | | | |
| Q14376 | UDP-glucose 4-epimerase OS=Homo sapiens OX=9606 GN=GALE PE=1 SV=2 | 1.078 | 0.765235202 | 0.954 | 0.7 |
| P08237 | ATP-dependent 6-phosphofructokinase, muscle type OS=Homo sapiens OX=9606 GN=PFKM PE=1 SV=2 | 0.996 | 0.983953845 | 0.919 | 0.7 |
| P46778 | 60S ribosomal protein L21 OS=Homo sapiens OX=9606 GN=RPL21 PE=1 SV=2 | 1.129 | 0.750023665 | 1.139 | 0.7 |
| Q6XQN6 | Nicotinate phosphoribosyltransferase OS=Homo sapiens OX=9606 GN=NAPRT PE=1 SV=2 | 1.028 | 0.905369216 | 0.831 | 0.7 |
| P09661 | U2 small nuclear ribonucleoprotein A' OS=Homo sapiens OX=9606 GN=SNRPA1 PE=1 SV=2 | 0.934 | 0.840158726 | 0.938 | 0.7 |
| E9PND2 | Cysteine and glycine-rich protein 1 (Fragment) OS=Homo sapiens OX=9606 GN=CSRP1 PE=1 SV=1 | 1.055 | 0.964209379 | 0.878 | 0.7 |
| O75368 | SH3 domain-binding glutamic acid-rich-like protein OS=Homo sapiens OX=9606 GN=SH3BGRL PE=1 SV=1 | 0.98 | 0.955325044 | 0.945 | 0.7 |
| Q7L7L0 | Histone H2A type 3 OS=Homo sapiens OX=9606 GN=H2AW PE=1 SV=3 | 0.81 | 0.80706255 | 0.98 | 0.8 |
| Q15046 | Lysine--tRNA ligase OS=Homo sapiens OX=9606 GN=KARS1 PE=1 SV=3 | 1.032 | 0.765235202 | 1.029 | 0.7 |
| P54687 | Branched-chain-amino-acid aminotransferase, cytosolic OS=Homo sapiens OX=9606 GN=BCAT1 PE=1 SV=3 | 0.951 | 0.765235202 | 0.976 | 0.9 |

| | | | | | |
|---|---|---|---|---|---|
| H0YLU7 | Electron transfer flavoprotein subunit alpha, mitochondrial (Fragment) OS=Homo sapiens OX=9606 GN=ETFA PE=1 SV=1 | 1.023 | 0.983953845 | 0.95 | 0.7 |
| P43304 | Glycerol-3-phosphate dehydrogenase, mitochondrial OS=Homo sapiens OX=9606 GN=GPD2 PE=1 SV=3 | 0.987 | 0.882454148 | 0.936 | 0.7 |
| P83731 | 60S ribosomal protein L24 OS=Homo sapiens OX=9606 GN=RPL24 PE=1 SV=1 | 1.123 | 0.750023665 | 1.14 | 0.7 |
| Q13045 | Protein flightless-1 homolog OS=Homo sapiens OX=9606 GN=FLII PE=1 SV=2 | 0.944 | 0.770292944 | 0.96 | 0.7 |
| P25398 | 40S ribosomal protein S12 OS=Homo sapiens OX=9606 GN=RPS12 PE=1 SV=3 | 1.017 | 0.80706255 | 1.101 | 0.7 |
| O00264 | Membrane-associated progesterone receptor component 1 OS=Homo sapiens OX=9606 GN=PGRMC1 PE=1 SV=3 | 1.029 | 0.902016292 | 0.975 | 0.7 |
| P46926 | Glucosamine-6-phosphate isomerase 1 OS=Homo sapiens OX=9606 GN=GNPDA1 PE=1 SV=1 | 0.959 | 0.750023665 | 0.934 | 0.7 |
| P49755 | Transmembrane emp24 domain-containing protein 10 OS=Homo sapiens OX=9606 GN=TMED10 PE=1 SV=2 | 1.101 | 0.782074241 | 1.01 | 0.8 |
| P62269 | 40S ribosomal protein S18 OS=Homo sapiens OX=9606 GN=RPS18 PE=1 SV=3 | 1.053 | 0.766667967 | 1.129 | 0.7 |
| P46781 | 40S ribosomal protein S9 OS=Homo sapiens OX=9606 GN=RPS9 PE=1 SV=3 | 1.062 | 0.908928234 | 1.06 | 0.7 |
| Q92820 | Gamma-glutamyl hydrolase OS=Homo sapiens OX=9606 GN=GGH PE=1 SV=2 | 0.979 | 0.959346973 | 0.925 | 0.7 |
| Q5T4U5 | Medium-chain specific acyl-CoA dehydrogenase, mitochondrial OS=Homo sapiens OX=9606 GN=ACADM PE=1 SV=1 | 1.022 | 0.876510992 | 0.958 | 0.7 |
| P36957 | Dihydrolipoyllysine-residue succinyltransferase component of 2-oxoglutarate dehydrogenase complex, mitochondrial OS=Homo s | 1.018 | 0.985324977 | 0.939 | 0.7 |
| Q14257 | Reticulocalbin-2 OS=Homo sapiens OX=9606 GN=RCN2 PE=1 SV=1 | 0.986 | 0.803355963 | 0.919 | 0.7 |
| Q16363 | Laminin subunit alpha-4 OS=Homo sapiens OX=9606 GN=LAMA4 PE=1 SV=4 | 0.951 | 0.750023665 | 0.911 | 0.7 |
| P62750 | 60S ribosomal protein L23a OS=Homo sapiens OX=9606 GN=RPL23A PE=1 SV=1 | 1.064 | 0.80706255 | 1.084 | 0.7 |
| P30085 | UMP-CMP kinase OS=Homo sapiens OX=9606 GN=CMPK1 PE=1 SV=3 | 0.966 | 0.750023665 | 1.027 | 0.9 |
| Q96G03 | Phosphoglucomutase-2 OS=Homo sapiens OX=9606 GN=PGM2 PE=1 SV=4 | 0.97 | 0.750023665 | 0.972 | 0.7 |
| H0Y512 | Adipocyte plasma membrane-associated protein (Fragment) OS=Homo sapiens OX=9606 GN=APMAP PE=1 SV=1 | 1.006 | 0.918702104 | 0.966 | 0.8 |
| P10644 | cAMP-dependent protein kinase type I-alpha regulatory subunit OS=Homo sapiens OX=9606 GN=PRKAR1A PE=1 SV=1 | 0.972 | 0.765235202 | 1.047 | 0.7 |
| Q53GQ0 | Very-long-chain 3-oxoacyl-CoA reductase OS=Homo sapiens OX=9606 GN=HSD17B12 PE=1 SV=2 | 1.029 | 0.908696432 | 1.059 | 0.9 |
| Q9Y6N5 | Sulfide:quinone oxidoreductase, mitochondrial OS=Homo sapiens OX=9606 GN=SQOR PE=1 SV=1 | 0.979 | 0.830179531 | 0.94 | 0.7 |
| Q02543 | 60S ribosomal protein L18a OS=Homo sapiens OX=9606 GN=RPL18A PE=1 SV=2 | 1.09 | 0.765235202 | 1.136 | 0.7 |
| O60763 | General vesicular transport factor p115 OS=Homo sapiens OX=9606 GN=USO1 PE=1 SV=2 | 1.011 | 0.895589243 | 0.949 | 0.7 |
| K7ER96 | Thioredoxin-like protein 1 (Fragment) OS=Homo sapiens OX=9606 GN=TXNL1 PE=1 SV=1 | 1.011 | 0.790136102 | 1.01 | 0.7 |
| P62249 | 40S ribosomal protein S16 OS=Homo sapiens OX=9606 GN=RPS16 PE=1 SV=2 | 1.064 | 0.785314375 | 1.091 | 0.7 |
| Q9UMS4 | Pre-mRNA-processing factor 19 OS=Homo sapiens OX=9606 GN=PRPF19 PE=1 SV=1 | 1.082 | 0.924992773 | 0.965 | 0.7 |
| Q9Y3C6 | Peptidyl-prolyl cis-trans isomerase-like 1 OS=Homo sapiens OX=9606 GN=PPIL1 PE=1 SV=1 | 0.986 | 0.96882036 | 0.948 | 0.7 |
| Q5W0H4 | Translationally-controlled tumor protein OS=Homo sapiens OX=9606 GN=TPT1 PE=1 SV=1 | 1.089 | 0.774484855 | 1.164 | 0.7 |
| Q01469 | Fatty acid-binding protein 5 OS=Homo sapiens OX=9606 GN=FABP5 PE=1 SV=3 | 0.947 | 0.902016292 | 1.016 | 0.8 |
| A0A3B3IUA7 | E3 ubiquitin/ISG15 ligase TRIM25 OS=Homo sapiens OX=9606 GN=TRIM25 PE=1 SV=1 | 1.018 | 0.955325044 | 0.957 | 0.8 |
| O60841 | Eukaryotic translation initiation factor 5B OS=Homo sapiens OX=9606 GN=EIF5B PE=1 SV=4 | 1.055 | 0.750023665 | 0.998 | 0.9 |
| B3KS98 | Eukaryotic translation initiation factor 3 subunit H OS=Homo sapiens OX=9606 GN=EIF3H PE=1 SV=1 | 1.028 | 0.750023665 | 1.048 | 0.7 |
| Q9NSD9 | Phenylalanine--tRNA ligase beta subunit OS=Homo sapiens OX=9606 GN=FARSB PE=1 SV=3 | 1.002 | 0.994361756 | 1.056 | 0.7 |
| P30050 | 60S ribosomal protein L12 OS=Homo sapiens OX=9606 GN=RPL12 PE=1 SV=1 | 1.053 | 0.780884711 | 1.08 | 0.7 |
| Q9NZI8 | Insulin-like growth factor 2 mRNA-binding protein 1 OS=Homo sapiens OX=9606 GN=IGF2BP1 PE=1 SV=2 | 0.98 | 0.937888222 | 0.957 | 0.7 |
| O43776 | Asparagine--tRNA ligase, cytoplasmic OS=Homo sapiens OX=9606 GN=NARS1 PE=1 SV=1 | 0.972 | 0.908555656 | 0.976 | 0.8 |
| P16401 | Histone H1.5 OS=Homo sapiens OX=9606 GN=H1-5 PE=1 SV=3 | 1.163 | 0.837893552 | 0.952 | 0.7 |
| Q9UMX0 | Ubiquilin-1 OS=Homo sapiens OX=9606 GN=UBQLN1 PE=1 SV=2 | 1.008 | 0.982301221 | 1.002 | 0.9 |
| O60884 | DnaJ homolog subfamily A member 2 OS=Homo sapiens OX=9606 GN=DNAJA2 PE=1 SV=1 | 0.964 | 0.829390484 | 1.03 | 0.7 |
| A0A0G2JMZ6 | Kinesin-like protein (Fragment) OS=Homo sapiens OX=9606 GN=KIF5C PE=1 SV=1 | 1.091 | 0.765235202 | 1.159 | 0.7 |
| A0A087WT44 | Heme oxygenase (biliverdin-producing) OS=Homo sapiens OX=9606 GN=HMOX2 PE=1 SV=1 | 1.013 | 0.912604818 | 0.991 | 0.9 |
| P61604 | 10 kDa heat shock protein, mitochondrial OS=Homo sapiens OX=9606 GN=HSPE1 PE=1 SV=2 | 0.972 | 0.963059765 | 0.989 | 0.9 |
| P54886 | Delta-1-pyrroline-5-carboxylate synthase OS=Homo sapiens OX=9606 GN=ALDH18A1 PE=1 SV=2 | 1.018 | 0.805488634 | 0.939 | 0.7 |
| P47897 | Glutamine--tRNA ligase OS=Homo sapiens OX=9606 GN=QARS1 PE=1 SV=1 | 1.063 | 0.765235202 | 1.022 | 0.9 |
| P63208 | S-phase kinase-associated protein 1 OS=Homo sapiens OX=9606 GN=SKP1 PE=1 SV=2 | 1.051 | 0.95980123 | 1.088 | 0.7 |
| Q14166 | Tubulin--tyrosine ligase-like protein 12 OS=Homo sapiens OX=9606 GN=TTLL12 PE=1 SV=2 | 0.99 | 0.978562349 | 0.957 | 0.7 |
| Q9BXP5 | Serrate RNA effector molecule homolog OS=Homo sapiens OX=9606 GN=SRRT PE=1 SV=1 | 0.963 | 0.765235202 | 0.973 | 0.7 |
| Q13283 | Ras GTPase-activating protein-binding protein 1 OS=Homo sapiens OX=9606 GN=G3BP1 PE=1 SV=1 | 1.039 | 0.989102644 | 1.123 | 0.7 |
| O43592 | Exportin-T OS=Homo sapiens OX=9606 GN=XPOT PE=1 SV=2 | 0.967 | 0.750023665 | 1.012 | 0.9 |
| P23229-6 | Isoform Alpha-6X1X2A of Integrin alpha-6 OS=Homo sapiens OX=9606 GN=ITGA6 | 0.963 | 0.750023665 | 0.966 | 0.7 |
| P06865 | Beta-hexosaminidase subunit alpha OS=Homo sapiens OX=9606 GN=HEXA PE=1 SV=2 | 0.982 | 0.827871327 | 0.856 | 0.7 |
| Q9Y2X3 | Nucleolar protein 58 OS=Homo sapiens OX=9606 GN=NOP58 PE=1 SV=1 | 1.024 | 0.882968647 | 0.925 | 0.8 |
| P11908 | Ribose-phosphate pyrophosphokinase 2 OS=Homo sapiens OX=9606 GN=PRPS2 PE=1 SV=2 | 0.961 | 0.765235202 | 1.016 | 0.9 |
| P36507 | Dual specificity mitogen-activated protein kinase kinase 2 OS=Homo sapiens OX=9606 GN=MAP2K2 PE=1 SV=1 | 0.978 | 0.750023665 | 0.988 | 0.7 |
| Q12874 | Splicing factor 3A subunit 3 OS=Homo sapiens OX=9606 GN=SF3A3 PE=1 SV=1 | 1.022 | 0.765235202 | 0.953 | 0.7 |
| O76094 | Signal recognition particle subunit SRP72 OS=Homo sapiens OX=9606 GN=SRP72 PE=1 SV=3 | 0.997 | 0.990981579 | 1.014 | 0.9 |
| O75947 | ATP synthase subunit d, mitochondrial OS=Homo sapiens OX=9606 GN=ATP5PD PE=1 SV=3 | 1.027 | 0.973458115 | 0.81 | 0.7 |
| P04179 | Superoxide dismutase [Mn], mitochondrial OS=Homo sapiens OX=9606 GN=SOD2 PE=1 SV=3 | 0.947 | 0.750023665 | 0.953 | 0.7 |
| P55036 | 26S proteasome non-ATPase regulatory subunit 4 OS=Homo sapiens OX=9606 GN=PSMD4 PE=1 SV=1 | 0.989 | 0.957217311 | 1.004 | 0.8 |
| E9PAV3-2 | Isoform skNAC-2 of Nascent polypeptide-associated complex subunit alpha, muscle-specific form OS=Homo sapiens OX=9606 GN= | 1.024 | 0.765235202 | 1.109 | 0.7 |
| Q15363 | Transmembrane emp24 domain-containing protein 2 OS=Homo sapiens OX=9606 GN=TMED2 PE=1 SV=1 | 1.041 | 0.978562349 | 0.93 | 0.8 |
| P50552 | Vasodilator-stimulated phosphoprotein OS=Homo sapiens OX=9606 GN=VASP PE=1 SV=3 | 1.009 | 0.99319413 | 0.963 | 0.7 |
| Q13347 | Eukaryotic translation initiation factor 3 subunit I OS=Homo sapiens OX=9606 GN=EIF3I PE=1 SV=1 | 1.044 | 0.792509319 | 1.01 | 0.8 |
| P21926 | CD9 antigen OS=Homo sapiens OX=9606 GN=CD9 PE=1 SV=4 | 0.926 | 0.765235202 | 0.978 | 0.9 |
| O00505 | Importin subunit alpha-4 OS=Homo sapiens OX=9606 GN=KPNA3 PE=1 SV=2 | 1.002 | 0.889482529 | 1.027 | 0.7 |
| P49720 | Proteasome subunit beta type-3 OS=Homo sapiens OX=9606 GN=PSMB3 PE=1 SV=2 | 0.943 | 0.913279108 | 0.97 | 0.7 |
| E9PPJ0 | Splicing factor 3B subunit 2 OS=Homo sapiens OX=9606 GN=SF3B2 PE=1 SV=1 | 1.062 | 0.892311206 | 0.964 | 0.7 |
| P42892 | Endothelin-converting enzyme 1 OS=Homo sapiens OX=9606 GN=ECE1 PE=1 SV=2 | 0.946 | 0.765235202 | 0.974 | 0.7 |
| P60891 | Ribose-phosphate pyrophosphokinase 1 OS=Homo sapiens OX=9606 GN=PRPS1 PE=1 SV=2 | 1.02 | 0.859846322 | 1.026 | 0.7 |
| P16615 | Sarcoplasmic/endoplasmic reticulum calcium ATPase 2 OS=Homo sapiens OX=9606 GN=ATP2A2 PE=1 SV=1 | 1.027 | 0.750023665 | 1.005 | 0.9 |
| Q07666 | KH domain-containing, RNA-binding, signal transduction-associated protein 1 OS=Homo sapiens OX=9606 GN=KHDRBS1 PE=1 SV= | 1.018 | 0.860909277 | 0.908 | 0.7 |
| F5H6I7 | Atlastin-3 OS=Homo sapiens OX=9606 GN=ATL3 PE=1 SV=1 | 1.013 | 0.99214485 | 1.021 | 0.7 |
| O94925-3 | Isoform 3 of Glutaminase kidney isoform, mitochondrial OS=Homo sapiens OX=9606 GN=GLS | 0.988 | 0.924992773 | 0.97 | 0.7 |
| Q9UKM9 | RNA-binding protein Raly OS=Homo sapiens OX=9606 GN=RALY PE=1 SV=1 | 1.053 | 0.87038391 | 0.922 | 0.7 |
| Q93009 | Ubiquitin carboxyl-terminal hydrolase 7 OS=Homo sapiens OX=9606 GN=USP7 PE=1 SV=2 | 0.998 | 0.985324977 | 1.004 | 0.7 |
| K7ELL7 | Glucosidase 2 subunit beta OS=Homo sapiens OX=9606 GN=PRKCSH PE=1 SV=1 | 1.001 | 0.968322408 | 0.999 | 0.9 |
| P51858 | Hepatoma-derived growth factor OS=Homo sapiens OX=9606 GN=HDGF PE=1 SV=1 | 0.939 | 0.829390484 | 0.915 | 0.7 |
| Q27J81 | Inverted formin-2 OS=Homo sapiens OX=9606 GN=INF2 PE=1 SV=2 | 1.006 | 0.837893552 | 1.014 | 0.9 |
| Q96AG4 | Leucine-rich repeat-containing protein 59 OS=Homo sapiens OX=9606 GN=LRRC59 PE=1 SV=1 | 1.028 | 0.904500837 | 1.07 | 0.7 |
| P25325 | 3-mercaptopyruvate sulfurtransferase OS=Homo sapiens OX=9606 GN=MPST PE=1 SV=3 | 0.934 | 0.765235202 | 0.926 | 0.7 |
| P48556 | 26S proteasome non-ATPase regulatory subunit 8 OS=Homo sapiens OX=9606 GN=PSMD8 PE=1 SV=2 | 1.016 | 0.930855729 | 0.996 | 0.9 |
| Q15907 | Ras-related protein Rab-11B OS=Homo sapiens OX=9606 GN=RAB11B PE=1 SV=4 | 0.974 | 0.770292944 | 1.025 | 0.7 |
| A0A087WY61 | Nuclear mitotic apparatus protein 1 OS=Homo sapiens OX=9606 GN=NUMA1 PE=1 SV=1 | 1.075 | 0.908555656 | 0.946 | 0.9 |
| A0A1B0GVU9 | Glutaminyl-tRNA synthetase (Fragment) OS=Homo sapiens OX=9606 GN=QARS1 PE=1 SV=1 | 0.811 | 0.828363915 | 0.914 | 0.7 |
| Q9H299 | SH3 domain-binding glutamic acid-rich-like protein 3 OS=Homo sapiens OX=9606 GN=SH3BGRL3 PE=1 SV=1 | 0.994 | 0.961750547 | 1.062 | 0.7 |
| Q9NUJ1 | Mycophenolic acid acyl-glucuronide esterase, mitochondrial OS=Homo sapiens OX=9606 GN=ABHD10 PE=1 SV=1 | 0.972 | 0.765235202 | 0.927 | 0.7 |
| B4DKY1 | Cysteinyl-tRNA synthetase OS=Homo sapiens OX=9606 GN=CARS1 PE=1 SV=1 | 0.995 | 0.916363276 | 0.996 | 0.7 |
| P55290 | Cadherin-13 OS=Homo sapiens OX=9606 GN=CDH13 PE=1 SV=1 | 0.965 | 0.750023665 | 0.941 | 0.7 |
| Q15185 | Prostaglandin E synthase 3 OS=Homo sapiens OX=9606 GN=PTGES3 PE=1 SV=1 | 0.969 | 0.904500837 | 1.031 | 0.7 |
| Q92688 | Acidic leucine-rich nuclear phosphoprotein 32 family member B OS=Homo sapiens OX=9606 GN=ANP32B PE=1 SV=1 | 0.932 | 0.750023665 | 0.934 | 0.7 |
| P10253 | Lysosomal alpha-glucosidase OS=Homo sapiens OX=9606 GN=GAA PE=1 SV=4 | 0.956 | 0.750023665 | 0.943 | 0.7 |
| M0R1B5 | 2-hydroxyacyl-CoA lyase 2 (Fragment) OS=Homo sapiens OX=9606 GN=ILVBL PE=1 SV=8 | 1.088 | 0.798529197 | 0.919 | 0.7 |
| Q96AE4 | Far upstream element-binding protein 1 OS=Homo sapiens OX=9606 GN=FUBP1 PE=1 SV=3 | 0.962 | 0.9252392 | 0.944 | 0.7 |
| P16104 | Histone H2AX OS=Homo sapiens OX=9606 GN=H2AX PE=1 SV=2 | 0.686 | 0.765235202 | 0.931 | 0.9 |
| Q14157-1 | Isoform 2 of Ubiquitin-associated protein 2-like OS=Homo sapiens OX=9606 GN=UBAP2L | 1.016 | 0.87017509 | 1.017 | 0.7 |

| | | | | | |
|---|---|---|---|---|---|
| P31942 | Heterogeneous nuclear ribonucleoprotein H3 OS=Homo sapiens OX=9606 GN=HNRNPH3 PE=1 SV=2 | 1.034 | 0.933097749 | 0.901 | 0.7 |
| Q9Y4D7 | Plexin-D1 OS=Homo sapiens OX=9606 GN=PLXND1 PE=1 SV=3 | 0.965 | 0.750023665 | 0.891 | 0.7 |
| Q99961 | Endophilin-A2 OS=Homo sapiens OX=9606 GN=SH3GL1 PE=1 SV=1 | 1.001 | 0.983953845 | 1.018 | 0.7 |
| O76021 | Ribosomal L1 domain-containing protein 1 OS=Homo sapiens OX=9606 GN=RSL1D1 PE=1 SV=3 | 1.007 | 0.973643049 | 0.93 | 0.9 |
| Q8TEX9 | Importin-4 OS=Homo sapiens OX=9606 GN=IPO4 PE=1 SV=2 | 0.998 | 0.882968647 | 1.031 | 0.8 |
| P62277 | 40S ribosomal protein S13 OS=Homo sapiens OX=9606 GN=RPS13 PE=1 SV=2 | 1.056 | 0.883217803 | 1.097 | 0.7 |
| Q02809 | Procollagen-lysine,2-oxoglutarate 5-dioxygenase 1 OS=Homo sapiens OX=9606 GN=PLOD1 PE=1 SV=2 | 1.012 | 0.779470244 | 0.96 | 0.7 |
| O14787-2 | Isoform 2 of Transportin-2 OS=Homo sapiens OX=9606 GN=TNPO2 | 1 | 0.886561233 | 0.993 | 0.7 |
| Q08J23 | RNA cytosine C(5)-methyltransferase NSUN2 OS=Homo sapiens OX=9606 GN=NSUN2 PE=1 SV=2 | 1.041 | 0.876510992 | 0.985 | 0.7 |
| P49189 | 4-trimethylaminobutyraldehyde dehydrogenase OS=Homo sapiens OX=9606 GN=ALDH9A1 PE=1 SV=3 | 0.941 | 0.829390484 | 0.998 | 0.8 |
| Q92905 | COP9 signalosome complex subunit 5 OS=Homo sapiens OX=9606 GN=COPS5 PE=1 SV=4 | 0.986 | 0.895384791 | 0.934 | 0.7 |
| P39060 | Collagen alpha-1(XVIII) chain OS=Homo sapiens OX=9606 GN=COL18A1 PE=1 SV=5 | 1.061 | 0.765235202 | 0.898 | 0.9 |
| Q6P179 | Endoplasmic reticulum aminopeptidase 2 OS=Homo sapiens OX=9606 GN=ERAP2 PE=1 SV=2 | 0.98 | 0.766667967 | 0.958 | 0.9 |
| Q14247 | Src substrate cortactin OS=Homo sapiens OX=9606 GN=CTTN PE=1 SV=2 | 1.069 | 0.750023665 | 1.045 | 0.7 |
| Q1JUQ5 | Peptidylprolyl isomerase OS=Homo sapiens OX=9606 GN=FKBP1A PE=1 SV=1 | 0.943 | 0.80706255 | 0.969 | 0.7 |
| Q9UIJ7 | GTP:AMP phosphotransferase AK3, mitochondrial OS=Homo sapiens OX=9606 GN=AK3 PE=1 SV=4 | 0.975 | 0.829390484 | 0.953 | 0.7 |
| Q13151 | Heterogeneous nuclear ribonucleoprotein A0 OS=Homo sapiens OX=9606 GN=HNRNPA0 PE=1 SV=1 | 1.021 | 0.814079429 | 1.025 | 0.7 |
| Q9UBV8 | Peflin OS=Homo sapiens OX=9606 GN=PEF1 PE=1 SV=1 | 0.971 | 0.795797569 | 0.937 | 0.7 |
| P0CG39 | POTE ankyrin domain family member J OS=Homo sapiens OX=9606 GN=POTEJ PE=3 SV=1 | | | | |
| P41091 | Eukaryotic translation initiation factor 2 subunit 3 OS=Homo sapiens OX=9606 GN=EIF2S3 PE=1 SV=3 | 1.071 | 0.960293674 | 1.002 | 0.9 |
| P55010 | Eukaryotic translation initiation factor 5 OS=Homo sapiens OX=9606 GN=EIF5 PE=1 SV=2 | 1.002 | 0.978003638 | 1.022 | 0.7 |
| P43487 | Ran-specific GTPase-activating protein OS=Homo sapiens OX=9606 GN=RANBP1 PE=1 SV=1 | 0.98 | 0.765235202 | 1.054 | 0.7 |
| P13598 | Intercellular adhesion molecule 2 OS=Homo sapiens OX=9606 GN=ICAM2 PE=1 SV=2 | 0.885 | 0.765235202 | 0.901 | 0.7 |
| P84085 | ADP-ribosylation factor 5 OS=Homo sapiens OX=9606 GN=ARF5 PE=1 SV=2 | 0.935 | 0.766521608 | 0.949 | 0.7 |
| P63000 | Ras-related C3 botulinum toxin substrate 1 OS=Homo sapiens OX=9606 GN=RAC1 PE=1 SV=1 | 1.049 | 0.904500837 | 1 | 0.8 |
| J3KQ69 | DNA helicase OS=Homo sapiens OX=9606 GN=MCM3 PE=1 SV=3 | 0.944 | 0.765235202 | 1.089 | 0.7 |
| Q96I99 | Succinate--CoA ligase [GDP-forming] subunit beta, mitochondrial OS=Homo sapiens OX=9606 GN=SUCLG2 PE=1 SV=2 | 0.98 | 0.766667967 | 0.947 | 0.7 |
| B4DR61 | Protein transport protein Sec61 subunit alpha isoform 1 OS=Homo sapiens OX=9606 GN=SEC61A1 PE=1 SV=1 | 1.022 | 0.996637279 | 0.973 | 0.9 |
| P48047 | ATP synthase subunit O, mitochondrial OS=Homo sapiens OX=9606 GN=ATP5PO PE=1 SV=1 | 0.998 | 0.983953845 | 0.86 | 0.7 |
| A0A286YFF7 | Palmitoyl-protein hydrolase 1 OS=Homo sapiens OX=9606 GN=PPT1 PE=1 SV=1 | 1.006 | 0.983953845 | 0.911 | 0.7 |
| P84103 | Serine/arginine-rich splicing factor 3 OS=Homo sapiens OX=9606 GN=SRSF3 PE=1 SV=1 | 0.985 | 0.910777517 | 0.966 | 0.7 |
| A0A2R8Y849 | 40S ribosomal protein S24 OS=Homo sapiens OX=9606 GN=RPS24 PE=1 SV=1 | 1.071 | 0.750023665 | 1.15 | 0.7 |
| P52888 | Thimet oligopeptidase OS=Homo sapiens OX=9606 GN=THOP1 PE=1 SV=2 | 0.968 | 0.765235202 | 0.952 | 0.7 |
| P20073 | Annexin A7 OS=Homo sapiens OX=9606 GN=ANXA7 PE=1 SV=3 | 0.983 | 0.869995963 | 0.995 | 0.9 |
| P51570 | Galactokinase OS=Homo sapiens OX=9606 GN=GALK1 PE=1 SV=1 | 0.975 | 0.765235202 | 0.954 | 0.7 |
| D6RFN0 | COP9 signalosome complex subunit 4 OS=Homo sapiens OX=9606 GN=COPS4 PE=1 SV=1 | 1.012 | 0.905369216 | 0.974 | 0.7 |
| P05120 | Plasminogen activator inhibitor 2 OS=Homo sapiens OX=9606 GN=SERPINB2 PE=1 SV=2 | 0.989 | 0.905369216 | 1.117 | 0.9 |
| Q02750 | Dual specificity mitogen-activated protein kinase kinase 1 OS=Homo sapiens OX=9606 GN=MAP2K1 PE=1 SV=2 | 1.031 | 0.766667967 | 1.048 | 0.7 |
| P28074 | Proteasome subunit beta type-5 OS=Homo sapiens OX=9606 GN=PSMB5 PE=1 SV=3 | 0.955 | 0.750023665 | 0.958 | 0.7 |
| Q13492 | Phosphatidylinositol-binding clathrin assembly protein OS=Homo sapiens OX=9606 GN=PICALM PE=1 SV=2 | 1.025 | 0.750023665 | 0.97 | 0.7 |
| P27694 | Replication protein A 70 kDa DNA-binding subunit OS=Homo sapiens OX=9606 GN=RPA1 PE=1 SV=2 | 0.988 | 0.978562349 | 0.986 | 0.7 |
| Q15637 | Splicing factor 1 OS=Homo sapiens OX=9606 GN=SF1 PE=1 SV=4 | 1.019 | 0.840110974 | 0.945 | 0.7 |
| A6NDG6 | Glycerol-3-phosphate phosphatase OS=Homo sapiens OX=9606 GN=PGP PE=1 SV=1 | 0.912 | 0.750023665 | 0.952 | 0.8 |
| Q8IYB3 | Serine/arginine repetitive matrix protein 1 OS=Homo sapiens OX=9606 GN=SRRM1 PE=1 SV=2 | 1.07 | 0.827871327 | 0.945 | 0.7 |
| J3KS05 | Chromobox protein homolog 1 (Fragment) OS=Homo sapiens OX=9606 GN=CBX1 PE=1 SV=8 | 1.027 | 0.765235202 | 0.906 | 0.7 |
| P51572 | B-cell receptor-associated protein 31 OS=Homo sapiens OX=9606 GN=BCAP31 PE=1 SV=3 | 1.048 | 0.750023665 | 1.048 | 0.7 |
| Q92743 | Serine protease HTRA1 OS=Homo sapiens OX=9606 GN=HTRA1 PE=1 SV=1 | 0.987 | 0.985324977 | 0.803 | 0.7 |
| P06756 | Integrin alpha-V OS=Homo sapiens OX=9606 GN=ITGAV PE=1 SV=2 | 0.973 | 0.809976184 | 0.872 | 0.7 |
| E5RGR0 | Acyl-protein thioesterase 1 (Fragment) OS=Homo sapiens OX=9606 GN=LYPLA1 PE=1 SV=1 | 0.934 | 0.765235202 | 0.987 | 0.7 |
| A0A1B0GUS4 | Ubiquitin-conjugating enzyme E2 L5 OS=Homo sapiens OX=9606 GN=UBE2L5 PE=2 SV=1 | 1.035 | 0.827871327 | 0.952 | 0.7 |
| P54819 | Adenylate kinase 2, mitochondrial OS=Homo sapiens OX=9606 GN=AK2 PE=1 SV=2 | 0.982 | 0.917008272 | 0.953 | 0.7 |
| Q04837 | Single-stranded DNA-binding protein, mitochondrial OS=Homo sapiens OX=9606 GN=SSBP1 PE=1 SV=1 | 0.97 | 0.750023665 | 0.925 | 0.7 |
| P13798 | Acylamino-acid-releasing enzyme OS=Homo sapiens OX=9606 GN=APEH PE=1 SV=4 | 1.005 | 0.994361756 | 1.016 | 0.7 |
| P62318 | Small nuclear ribonucleoprotein Sm D3 OS=Homo sapiens OX=9606 GN=SNRPD3 PE=1 SV=1 | 0.945 | 0.906273459 | 0.916 | 0.7 |
| Q99436 | Proteasome subunit beta type-7 OS=Homo sapiens OX=9606 GN=PSMB7 PE=1 SV=1 | 0.964 | 0.750023665 | 1.013 | 0.7 |
| Q16222 | UDP-N-acetylhexosamine pyrophosphorylase OS=Homo sapiens OX=9606 GN=UAP1 PE=1 SV=3 | 1.027 | 0.88409663 | 1.083 | 0.7 |
| P48147 | Prolyl endopeptidase OS=Homo sapiens OX=9606 GN=PREP PE=1 SV=2 | 0.999 | 0.985324977 | 0.912 | 0.7 |
| Q03135 | Caveolin-1 OS=Homo sapiens OX=9606 GN=CAV1 PE=1 SV=4 | 1.015 | 0.995175368 | 0.853 | 0.7 |
| O75937 | DnaJ homolog subfamily C member 8 OS=Homo sapiens OX=9606 GN=DNAJC8 PE=1 SV=2 | 1.022 | 0.765235202 | 0.956 | 0.7 |
| P22307 | Non-specific lipid-transfer protein OS=Homo sapiens OX=9606 GN=SCP2 PE=1 SV=2 | 0.958 | 0.815069882 | 0.972 | 0.9 |
| P50995 | Annexin A11 OS=Homo sapiens OX=9606 GN=ANXA11 PE=1 SV=1 | 0.984 | 0.794788863 | 1.038 | 0.7 |
| Q9Y6E2 | Basic leucine zipper and W2 domain-containing protein 2 OS=Homo sapiens OX=9606 GN=BZW2 PE=1 SV=1 | 1.054 | 0.765235202 | 1.133 | 0.7 |
| Q9NR31 | GTP-binding protein SAR1a OS=Homo sapiens OX=9606 GN=SAR1A PE=1 SV=1 | 1.017 | 0.903226585 | 1.034 | 0.7 |
| P84098 | 60S ribosomal protein L19 OS=Homo sapiens OX=9606 GN=RPL19 PE=1 SV=1 | 1.019 | 0.765235202 | 1.101 | 0.7 |
| Q66K74 | Microtubule-associated protein 1S OS=Homo sapiens OX=9606 GN=MAP1S PE=1 SV=2 | 1.047 | 0.779783772 | 0.894 | 0.7 |
| Q14444 | Caprin-1 OS=Homo sapiens OX=9606 GN=CAPRIN1 PE=1 SV=2 | 1.06 | 0.917008272 | 1.104 | 0.7 |
| P31689 | DnaJ homolog subfamily A member 1 OS=Homo sapiens OX=9606 GN=DNAJA1 PE=1 SV=2 | 1.057 | 0.765235202 | 1.109 | 0.7 |
| P61353 | 60S ribosomal protein L27 OS=Homo sapiens OX=9606 GN=RPL27 PE=1 SV=2 | 1.028 | 0.969565732 | 1.082 | 0.7 |
| P17612 | cAMP-dependent protein kinase catalytic subunit alpha OS=Homo sapiens OX=9606 GN=PRKACA PE=1 SV=2 | 0.966 | 0.700264056 | 0.96 | 0.7 |
| Q9UFN0 | Protein NipSnap homolog 3A OS=Homo sapiens OX=9606 GN=NIPSNAP3A PE=1 SV=2 | 0.928 | 0.750023665 | 0.933 | 0.7 |
| Q9Y3C4 | EKC/KEOPS complex subunit TPRKB OS=Homo sapiens OX=9606 GN=TPRKB PE=1 SV=1 | 1.003 | 0.985324977 | 0.917 | 0.7 |
| P61201-2 | Isoform 2 of COP9 signalosome complex subunit 2 OS=Homo sapiens OX=9606 GN=COPS2 | 1.026 | 0.994361756 | 0.784 | 0.7 |
| P39656 | Dolichyl-diphosphooligosaccharide--protein glycosyltransferase 48 kDa subunit OS=Homo sapiens OX=9606 GN=DDOST PE=1 SV=4 | 1.007 | 0.973370361 | 0.953 | 0.8 |
| Q9Y5S9 | RNA-binding protein 8A OS=Homo sapiens OX=9606 GN=RBM8A PE=1 SV=1 | 0.993 | 0.876510992 | 0.985 | 0.7 |
| P61019 | Ras-related protein Rab-2A OS=Homo sapiens OX=9606 GN=RAB2A PE=1 SV=1 | 0.995 | 0.939024443 | 1.018 | 0.8 |
| B1AHB1 | DNA helicase OS=Homo sapiens OX=9606 GN=MCM5 PE=1 SV=1 | 0.997 | 0.984805107 | 1.105 | 0.7 |
| P36542 | ATP synthase subunit gamma, mitochondrial OS=Homo sapiens OX=9606 GN=ATP5F1C PE=1 SV=1 | 0.962 | 0.978138288 | 0.851 | 0.7 |
| P61313 | 60S ribosomal protein L15 OS=Homo sapiens OX=9606 GN=RPL15 PE=1 SV=2 | 1.055 | 0.750023665 | 1.047 | 0.7 |
| Q9NSE4 | Isoleucine--tRNA ligase, mitochondrial OS=Homo sapiens OX=9606 GN=IARS2 PE=1 SV=2 | 1.012 | 0.895384791 | 0.896 | 0.7 |
| Q99627 | COP9 signalosome complex subunit 8 OS=Homo sapiens OX=9606 GN=COPS8 PE=1 SV=1 | 1.028 | 0.902016292 | 0.958 | 0.7 |
| P30837 | Aldehyde dehydrogenase X, mitochondrial OS=Homo sapiens OX=9606 GN=ALDH1B1 PE=1 SV=3 | 0.948 | 0.750023665 | 0.876 | 0.7 |
| Q68EM7 | Rho GTPase-activating protein 17 OS=Homo sapiens OX=9606 GN=ARHGAP17 PE=1 SV=1 | 0.974 | 0.765235202 | 0.987 | 0.8 |
| Q14254 | Flotillin-2 OS=Homo sapiens OX=9606 GN=FLOT2 PE=1 SV=2 | 0.987 | 0.917008272 | 0.908 | 0.7 |
| P31323 | cAMP-dependent protein kinase type II-beta regulatory subunit OS=Homo sapiens OX=9606 GN=PRKAR2B PE=1 SV=3 | 0.952 | 0.937888222 | 0.935 | 0.7 |
| Q13155 | Aminoacyl tRNA synthase complex-interacting multifunctional protein 2 OS=Homo sapiens OX=9606 GN=AIMP2 PE=1 SV=2 | 1.065 | 0.792509319 | 1.005 | 0.9 |
| Q9BUJ2 | Heterogeneous nuclear ribonucleoprotein U-like protein 1 OS=Homo sapiens OX=9606 GN=HNRNPUL1 PE=1 SV=2 | 0.867 | 0.829390484 | 0.887 | 0.7 |
| Q8WUM0 | Nuclear pore complex protein Nup133 OS=Homo sapiens OX=9606 GN=NUP133 PE=1 SV=2 | 1.037 | 0.765235202 | 0.931 | 0.8 |
| P16152 | Carbonyl reductase [NADPH] 1 OS=Homo sapiens OX=9606 GN=CBR1 PE=1 SV=3 | 0.979 | 0.96863015 | 0.964 | 0.7 |
| P40121 | Macrophage-capping protein OS=Homo sapiens OX=9606 GN=CAPG PE=1 SV=2 | 0.961 | 0.776314646 | 1.006 | 0.9 |
| Q93008-1 | Isoform 2 of Probable ubiquitin carboxyl-terminal hydrolase FAF-X OS=Homo sapiens OX=9606 GN=USP9X | 1.02 | 0.765235202 | 0.923 | 0.7 |
| O14737 | Programmed cell death protein 5 OS=Homo sapiens OX=9606 GN=PDCD5 PE=1 SV=3 | 0.985 | 0.876510992 | 1.049 | 0.7 |
| P62995 | Transformer-2 protein homolog beta OS=Homo sapiens OX=9606 GN=TRA2B PE=1 SV=1 | 0.995 | 0.905369216 | 0.943 | 0.7 |
| O94760 | N(G),N(G)-dimethylarginine dimethylaminohydrolase 1 OS=Homo sapiens OX=9606 GN=DDAH1 PE=1 SV=3 | 0.969 | 0.750023665 | 0.92 | 0.7 |
| X6RFL8 | Ras-related protein Rab-14 (Fragment) OS=Homo sapiens OX=9606 GN=RAB14 PE=1 SV=1 | 1.006 | 0.985324977 | 0.975 | 0.7 |
| P62244 | 40S ribosomal protein S15a OS=Homo sapiens OX=9606 GN=RPS15A PE=1 SV=1 | 1.086 | 0.794788863 | 1.076 | 0.7 |
| P40429 | 60S ribosomal protein L13a OS=Homo sapiens OX=9606 GN=RPL13A PE=1 SV=2 | 1.049 | 0.830305674 | 1.106 | 0.7 |

| | | | | | |
|---|---|---|---|---|---|
| Q13867 | Bleomycin hydrolase OS=Homo sapiens OX=9606 GN=BLMH PE=1 SV=1 | 0.975 | 0.898725987 | 0.936 | 0.7 |
| O95394 | Phosphoacetylglucosamine mutase OS=Homo sapiens OX=9606 GN=PGM3 PE=1 SV=1 | 0.979 | 0.765235202 | 0.973 | 0.7 |
| P33993 | DNA replication licensing factor MCM7 OS=Homo sapiens OX=9606 GN=MCM7 PE=1 SV=4 | 0.999 | 0.889482529 | 1.02 | 0.7 |
| P52815 | 39S ribosomal protein L12, mitochondrial OS=Homo sapiens OX=9606 GN=MRPL12 PE=1 SV=2 | 1.002 | 0.925929947 | 0.907 | 0.7 |
| P21266 | Glutathione S-transferase Mu 3 OS=Homo sapiens OX=9606 GN=GSTM3 PE=1 SV=3 | 0.982 | 0.973458115 | 0.921 | 0.8 |
| A0A1W2PNX8 | Protein unc-45 homolog A OS=Homo sapiens OX=9606 GN=UNC45A PE=1 SV=1 | 0.986 | 0.957217311 | 0.964 | 0.7 |
| P61006 | Ras-related protein Rab-8A OS=Homo sapiens OX=9606 GN=RAB8A PE=1 SV=1 | 0.945 | 0.866566453 | 0.926 | 0.7 |
| P61960 | Ubiquitin-fold modifier 1 OS=Homo sapiens OX=9606 GN=UFM1 PE=1 SV=1 | 1.062 | 0.899342793 | 1.066 | 0.8 |
| Q52LJ0-1 | Isoform 1 of Protein FAM98B OS=Homo sapiens OX=9606 GN=FAM98B | 0.955 | 0.766667967 | 1.023 | 0.8 |
| Q9P0V9 | Septin-10 OS=Homo sapiens OX=9606 GN=SEPTIN10 PE=1 SV=2 | 0.966 | 0.906273459 | 0.933 | 0.7 |
| Q9NR28 | Diablo homolog, mitochondrial OS=Homo sapiens OX=9606 GN=DIABLO PE=1 SV=1 | 1.068 | 0.913279108 | 0.937 | 0.7 |
| P13796 | Plastin-2 OS=Homo sapiens OX=9606 GN=LCP1 PE=1 SV=6 | 1.059 | 0.879331928 | 0.746 | 0.7 |
| P00492 | Hypoxanthine-guanine phosphoribosyltransferase OS=Homo sapiens OX=9606 GN=HPRT1 PE=1 SV=2 | 1.016 | 0.886561233 | 0.977 | 0.7 |
| O95573 | Long-chain-fatty-acid--CoA ligase 3 OS=Homo sapiens OX=9606 GN=ACSL3 PE=1 SV=3 | 1.052 | 0.765235202 | 1.001 | 0.9 |
| Q5VVC8 | 60S ribosomal protein L11 OS=Homo sapiens OX=9606 GN=RPL11 PE=1 SV=2 | 1.062 | 0.765235202 | 1.138 | 0.7 |
| F6WIT2 | Serine/threonine-protein phosphatase 2A activator OS=Homo sapiens OX=9606 GN=PTPA PE=1 SV=1 | 0.942 | 0.750023665 | 0.958 | 0.7 |
| G5E9Q6 | Profilin OS=Homo sapiens OX=9606 GN=PFN2 PE=1 SV=1 | 0.997 | 0.96882036 | 0.976 | 0.8 |
| P05386 | 60S acidic ribosomal protein P1 OS=Homo sapiens OX=9606 GN=RPLP1 PE=1 SV=1 | 1.015 | 0.939024443 | 1.063 | 0.7 |
| J3KTL2 | Serine/arginine-rich splicing factor 1 OS=Homo sapiens OX=9606 GN=SRSF1 PE=1 SV=1 | 1.022 | 0.983953845 | 0.956 | 0.7 |
| G3V0E4 | Beta-MPP OS=Homo sapiens OX=9606 GN=PMPCB PE=1 SV=1 | 0.997 | 0.996637279 | 0.938 | 0.7 |
| P63092 | Guanine nucleotide-binding protein G(s) subunit alpha isoforms short OS=Homo sapiens OX=9606 GN=GNAS PE=1 SV=1 | 0.964 | 0.902016292 | 0.891 | 0.7 |
| O15212 | Prefoldin subunit 6 OS=Homo sapiens OX=9606 GN=PFDN6 PE=1 SV=1 | 0.995 | 0.983953845 | 1.016 | 0.7 |
| Q13011 | Delta(3,5)-Delta(2,4)-dienoyl-CoA isomerase, mitochondrial OS=Homo sapiens OX=9606 GN=ECH1 PE=1 SV=2 | 0.953 | 0.786409349 | 0.923 | 0.7 |
| P35268 | 60S ribosomal protein L22 OS=Homo sapiens OX=9606 GN=RPL22 PE=1 SV=2 | 1.066 | 0.839845191 | 1.087 | 0.7 |
| Q6NVY1 | 3-hydroxyisobutyryl-CoA hydrolase, mitochondrial OS=Homo sapiens OX=9606 GN=HIBCH PE=1 SV=2 | 0.961 | 0.765235202 | 0.912 | 0.7 |
| A0A087X0R6 | Sorting nexin-12 OS=Homo sapiens OX=9606 GN=SNX12 PE=1 SV=1 | 1.02 | 0.773977513 | 0.998 | 0.9 |
| Q96PK6 | RNA-binding protein 14 OS=Homo sapiens OX=9606 GN=RBM14 PE=1 SV=2 | 1.115 | 0.765235202 | 0.966 | 0.7 |
| Q8IZ83 | Aldehyde dehydrogenase family 16 member A1 OS=Homo sapiens OX=9606 GN=ALDH16A1 PE=1 SV=2 | 1.002 | 0.985763982 | 0.988 | 0.8 |
| O14617 | AP-3 complex subunit delta-1 OS=Homo sapiens OX=9606 GN=AP3D1 PE=1 SV=1 | 0.98 | 0.886561233 | 0.96 | 0.7 |
| P49207 | 60S ribosomal protein L34 OS=Homo sapiens OX=9606 GN=RPL34 PE=1 SV=3 | 1.077 | 0.754126086 | 1.104 | 0.7 |
| P46063 | ATP-dependent DNA helicase Q1 OS=Homo sapiens OX=9606 GN=RECQL PE=1 SV=3 | 1.078 | 0.750023665 | 0.947 | 0.8 |
| P62899 | 60S ribosomal protein L31 OS=Homo sapiens OX=9606 GN=RPL31 PE=1 SV=1 | 1.014 | 0.840158726 | 1.056 | 0.7 |
| Q92900 | Regulator of nonsense transcripts 1 OS=Homo sapiens OX=9606 GN=UPF1 PE=1 SV=2 | 1.019 | 0.750023665 | 1.014 | 0.9 |
| P27361 | Mitogen-activated protein kinase 3 OS=Homo sapiens OX=9606 GN=MAPK3 PE=1 SV=4 | 0.992 | 0.996578046 | 0.956 | 0.7 |
| Q5VW36 | Focadhesin OS=Homo sapiens OX=9606 GN=FOCAD PE=1 SV=1 | 0.964 | 0.765235202 | 0.98 | 0.8 |
| P49407 | Beta-arrestin-1 OS=Homo sapiens OX=9606 GN=ARRB1 PE=1 SV=2 | 0.945 | 0.811788015 | 0.895 | 0.7 |
| P36542-2 | Isoform Heart of ATP synthase subunit gamma, mitochondrial OS=Homo sapiens OX=9606 GN=ATP5F1C | 0.978 | 0.996847952 | 0.839 | 0.7 |
| Q9NPQ8 | Synembryn-A OS=Homo sapiens OX=9606 GN=RIC8A PE=1 SV=3 | 1.058 | 0.750023665 | 1.041 | 0.7 |
| Q15286 | Ras-related protein Rab-35 OS=Homo sapiens OX=9606 GN=RAB35 PE=1 SV=1 | 0.927 | 0.765235202 | 0.86 | 0.7 |
| O60684 | Importin subunit alpha-7 OS=Homo sapiens OX=9606 GN=KPNA6 PE=1 SV=1 | 0.95 | 0.780884711 | 1.016 | 0.8 |
| P49419 | Alpha-aminoadipic semialdehyde dehydrogenase OS=Homo sapiens OX=9606 GN=ALDH7A1 PE=1 SV=5 | 0.89 | 0.765235202 | 0.905 | 0.7 |
| P47813 | Eukaryotic translation initiation factor 1A, X-chromosomal OS=Homo sapiens OX=9606 GN=EIF1AX PE=1 SV=2 | 0.937 | 0.765235202 | 1.102 | 0.7 |
| O15511 | Actin-related protein 2/3 complex subunit 5 OS=Homo sapiens OX=9606 GN=ARPC5 PE=1 SV=3 | 0.977 | 0.933097749 | 1.001 | 0.7 |
| P50914 | 60S ribosomal protein L14 OS=Homo sapiens OX=9606 GN=RPL14 PE=1 SV=4 | 1.047 | 0.765235202 | 1.121 | 0.7 |
| G3V3E8 | Epididymal secretory protein E1 OS=Homo sapiens OX=9606 GN=NPC2 PE=1 SV=1 | 0.954 | 0.765235202 | 0.878 | 0.7 |
| Q96HP0 | Dedicator of cytokinesis protein 6 OS=Homo sapiens OX=9606 GN=DOCK6 PE=1 SV=3 | 0.981 | 0.937953124 | 0.888 | 0.7 |
| B4E0K5 | Mitogen-activated protein kinase OS=Homo sapiens OX=9606 GN=MAPK14 PE=1 SV=1 | 0.964 | 0.750023665 | 0.974 | 0.9 |
| Q8IYD1 | Eukaryotic peptide chain release factor GTP-binding subunit ERF3B OS=Homo sapiens OX=9606 GN=GSPT2 PE=1 SV=2 | 0.984 | 0.774484855 | 0.915 | 0.7 |
| Q9NQR4 | Omega-amidase NIT2 OS=Homo sapiens OX=9606 GN=NIT2 PE=1 SV=1 | 0.995 | 0.97688166 | 0.939 | 0.7 |
| Q12805 | EGF-containing fibulin-like extracellular matrix protein 1 OS=Homo sapiens OX=9606 GN=EFEMP1 PE=1 SV=2 | 0.984 | 0.779783772 | 1.11 | 0.8 |
| O95336 | 6-phosphogluconolactonase OS=Homo sapiens OX=9606 GN=PGLS PE=1 SV=2 | 0.992 | 0.985324977 | 0.99 | 0.7 |
| P42025 | Beta-centractin OS=Homo sapiens OX=9606 GN=ACTR1B PE=1 SV=1 | 0.866 | 0.750023665 | 1.042 | 0.7 |
| Q8NCW5 | NAD(P)H-hydrate epimerase OS=Homo sapiens OX=9606 GN=NAXE PE=1 SV=2 | 1 | 0.972073425 | 0.909 | 0.7 |
| Q12792 | Twinfilin-1 OS=Homo sapiens OX=9606 GN=TWF1 PE=1 SV=3 | 0.983 | 0.775827921 | 1.005 | 0.9 |
| O75915 | PRA1 family protein 3 OS=Homo sapiens OX=9606 GN=ARL6IP5 PE=1 SV=1 | 0.883 | 0.815614436 | 0.954 | 0.8 |
| O43809 | Cleavage and polyadenylation specificity factor subunit 5 OS=Homo sapiens OX=9606 GN=NUDT21 PE=1 SV=1 | 0.98 | 0.985330338 | 0.958 | 0.7 |
| Q08623 | Pseudouridine-5'-phosphatase OS=Homo sapiens OX=9606 GN=PUDP PE=1 SV=3 | 0.902 | 0.889450577 | 0.88 | 0.7 |
| F8WJN3 | Cleavage and polyadenylation specificity factor subunit 6 OS=Homo sapiens OX=9606 GN=CPSF6 PE=1 SV=1 | 0.96 | 0.765235202 | 0.962 | 0.7 |
| F2Z2Y4 | Pyridoxal kinase OS=Homo sapiens OX=9606 GN=PDXK PE=1 SV=1 | 0.998 | 0.991801854 | 0.996 | 0.8 |
| F8W9J4 | Dystonin OS=Homo sapiens OX=9606 GN=DST PE=1 SV=1 | 0.97 | 0.882454148 | 0.909 | 0.7 |
| Q15427 | Splicing factor 3B subunit 4 OS=Homo sapiens OX=9606 GN=SF3B4 PE=1 SV=1 | 1.027 | 0.841691633 | 0.953 | 0.7 |
| Q9H845 | Complex I assembly factor ACAD9, mitochondrial OS=Homo sapiens OX=9606 GN=ACAD9 PE=1 SV=1 | 0.949 | 0.750023665 | 0.948 | 0.7 |
| Q9P0L0 | Vesicle-associated membrane protein-associated protein A OS=Homo sapiens OX=9606 GN=VAPA PE=1 SV=3 | 1.103 | 0.770292944 | 0.976 | 0.8 |
| E7ESD2 | WASH complex subunit 2A OS=Homo sapiens OX=9606 GN=WASHC2A PE=1 SV=2 | 0.982 | 0.96882036 | 0.952 | 0.7 |
| P35232 | Prohibitin OS=Homo sapiens OX=9606 GN=PHB PE=1 SV=1 | 1.042 | 0.942145703 | 0.846 | 0.7 |
| A0A0A0MS54 | cAMP-dependent protein kinase catalytic subunit beta OS=Homo sapiens OX=9606 GN=PRKACB PE=1 SV=1 | 0.963 | 0.955325044 | 1.031 | 0.7 |
| O60551 | Glycylpeptide N-tetradecanoyltransferase 2 OS=Homo sapiens OX=9606 GN=NMT2 PE=1 SV=1 | 0.991 | 0.953792095 | 0.954 | 0.9 |
| P20042 | Eukaryotic translation initiation factor 2 subunit 2 OS=Homo sapiens OX=9606 GN=EIF2S2 PE=1 SV=2 | 1.101 | 0.761475165 | 1.047 | 0.9 |
| F5GZS6 | 4F2 cell-surface antigen heavy chain OS=Homo sapiens OX=9606 GN=SLC3A2 PE=1 SV=1 | 0.98 | 0.937953124 | 0.994 | 0.7 |
| P26373 | 60S ribosomal protein L13 OS=Homo sapiens OX=9606 GN=RPL13 PE=1 SV=4 | 1.054 | 0.765235202 | 1.14 | 0.7 |
| Q15041 | ADP-ribosylation factor-like protein 6-interacting protein 1 OS=Homo sapiens OX=9606 GN=ARL6IP1 PE=1 SV=2 | 1.11 | 0.878144903 | 1.074 | 0.8 |
| H7C2N1 | Prothymosin alpha (Fragment) OS=Homo sapiens OX=9606 GN=PTMA PE=1 SV=1 | 1.028 | 0.916363276 | 1.146 | 0.8 |
| O43747 | AP-1 complex subunit gamma-1 OS=Homo sapiens OX=9606 GN=AP1G1 PE=1 SV=5 | 1.002 | 0.792509319 | 1.03 | 0.7 |
| Q9NRV9 | Heme-binding protein 1 OS=Homo sapiens OX=9606 GN=HEBP1 PE=1 SV=1 | 0.914 | 0.766667967 | 0.954 | 0.7 |
| H0Y3P2 | Eukaryotic translation initiation factor 4 gamma 2 OS=Homo sapiens OX=9606 GN=EIF4G2 PE=1 SV=1 | 1.026 | 0.750023665 | 0.983 | 0.7 |
| P46977 | Dolichyl-diphosphooligosaccharide--protein glycosyltransferase subunit STT3A OS=Homo sapiens OX=9606 GN=STT3A PE=1 SV=2 | 1.014 | 0.997835552 | 0.91 | 0.7 |
| Q9UNH7 | Sorting nexin-6 OS=Homo sapiens OX=9606 GN=SNX6 PE=1 SV=1 | 0.948 | 0.750023665 | 0.984 | 0.7 |
| A0A494C0U1 | Tripeptidyl-peptidase 2 (Fragment) OS=Homo sapiens OX=9606 GN=TPP2 PE=1 SV=1 | 0.999 | 0.991801854 | 0.99 | 0.7 |
| P22090 | 40S ribosomal protein S4, Y isoform 1 OS=Homo sapiens OX=9606 GN=RPS4Y1 PE=1 SV=2 | 1.063 | 0.806305724 | 1.063 | 0.9 |
| Q96T76 | MMS19 nucleotide excision repair protein homolog OS=Homo sapiens OX=9606 GN=MMS19 PE=1 SV=2 | 1.001 | 0.990981579 | 1 | 0.7 |
| P20340 | Ras-related protein Rab-6A OS=Homo sapiens OX=9606 GN=RAB6A PE=1 SV=3 | 0.991 | 0.976601777 | 0.981 | 0.7 |
| Q9BTT0 | Acidic leucine-rich nuclear phosphoprotein 32 family member E OS=Homo sapiens OX=9606 GN=ANP32E PE=1 SV=1 | 0.972 | 0.895384791 | 0.945 | 0.5 |
| Q9C0B1 | Alpha-ketoglutarate-dependent dioxygenase FTO OS=Homo sapiens OX=9606 GN=FTO PE=1 SV=3 | 0.934 | 0.750023665 | 0.904 | 0.7 |
| P53004 | Biliverdin reductase A OS=Homo sapiens OX=9606 GN=BLVRA PE=1 SV=2 | 0.965 | 0.765235202 | 1.009 | 0.9 |
| Q13084 | 39S ribosomal protein L28, mitochondrial OS=Homo sapiens OX=9606 GN=MRPL28 PE=1 SV=4 | 1.029 | 0.886561233 | 0.896 | 0.7 |
| P61923 | Coatomer subunit zeta-1 OS=Homo sapiens OX=9606 GN=COPZ1 PE=1 SV=1 | 1.029 | 0.765235202 | 1.005 | 0.7 |
| P62857 | 40S ribosomal protein S28 OS=Homo sapiens OX=9606 GN=RPS28 PE=1 SV=1 | 0.93 | 0.754126086 | 1.052 | 0.7 |
| Q9H444 | Charged multivesicular body protein 4b OS=Homo sapiens OX=9606 GN=CHMP4B PE=1 SV=1 | 0.96 | 0.765235202 | 0.973 | 0.7 |
| A0A1W2PQR6 | Lysosome membrane protein 2 OS=Homo sapiens OX=9606 GN=SCARB2 PE=1 SV=1 | 0.947 | 0.779783772 | 0.868 | 0.7 |
| Q8TBC4 | NEDD8-activating enzyme E1 catalytic subunit OS=Homo sapiens OX=9606 GN=UBA3 PE=1 SV=2 | 0.989 | 0.944284199 | 0.961 | 0.7 |
| Q9UBF2 | Coatomer subunit gamma-2 OS=Homo sapiens OX=9606 GN=COPG2 PE=1 SV=1 | 1.01 | 0.985324977 | 0.879 | 0.7 |
| Q9HAV4 | Exportin-5 OS=Homo sapiens OX=9606 GN=XPO5 PE=1 SV=1 | 0.992 | 0.994361756 | 0.968 | 0.7 |
| A0A0G2JLB3 | Glucosylceramidase OS=Homo sapiens OX=9606 GN=GBA PE=1 SV=1 | 1.003 | 0.971097726 | 0.94 | 0.7 |
| B0QZ18 | Copine-1 OS=Homo sapiens OX=9606 GN=CPNE1 PE=1 SV=1 | 1.023 | 0.893277754 | 0.958 | 0.7 |
| P05141 | ADP/ATP translocase 2 OS=Homo sapiens OX=9606 GN=SLC25A5 PE=1 SV=7 | 0.959 | 0.869503832 | 0.868 | 0.7 |
| Q9Y5B9 | FACT complex subunit SPT16 OS=Homo sapiens OX=9606 GN=SUPT16H PE=1 SV=1 | 1.027 | 0.96647353 | 0.896 | 0.7 |

| | | | | | |
|---|---|---|---|---|---|
| Q9BR76 | Coronin-1B OS=Homo sapiens OX=9606 GN=CORO1B PE=1 SV=1 | 1.024 | 0.809976184 | 0.911 | 0.7 |
| P62316 | Small nuclear ribonucleoprotein Sm D2 OS=Homo sapiens OX=9606 GN=SNRPD2 PE=1 SV=1 | 0.987 | 0.984200071 | 0.978 | 0.7 |
| A0A3B3IRQ8 | CTP synthase OS=Homo sapiens OX=9606 GN=CTPS1 PE=1 SV=1 | 1.005 | 0.969984528 | 0.996 | 0.9 |
| I1E4Y6 | GRB10-interacting GYF protein 2 OS=Homo sapiens OX=9606 GN=GIGYF2 PE=1 SV=1 | 1.038 | 0.863796857 | 0.984 | 0.7 |
| Q9Y371 | Endophilin-B1 OS=Homo sapiens OX=9606 GN=SH3GLB1 PE=1 SV=1 | 1.003 | 0.996637279 | 0.995 | 0.8 |
| P36871 | Phosphoglucomutase-1 OS=Homo sapiens OX=9606 GN=PGM1 PE=1 SV=3 | 0.989 | 0.944284199 | 0.984 | 0.8 |
| P61086 | Ubiquitin-conjugating enzyme E2 K OS=Homo sapiens OX=9606 GN=UBE2K PE=1 SV=3 | 1 | 0.994361756 | 0.987 | 0.7 |
| P16989 | Y-box-binding protein 3 OS=Homo sapiens OX=9606 GN=YBX3 PE=1 SV=4 | 1.074 | 0.766667967 | 1.073 | 0.7 |
| Q9UHL4 | Dipeptidyl peptidase 2 OS=Homo sapiens OX=9606 GN=DPP7 PE=1 SV=3 | 1.021 | 0.956912276 | 0.88 | 0.7 |
| Q09028 | Histone-binding protein RBBP4 OS=Homo sapiens OX=9606 GN=RBBP4 PE=1 SV=3 | 1.008 | 0.985324977 | 1.033 | 0.7 |
| Q08945 | FACT complex subunit SSRP1 OS=Homo sapiens OX=9606 GN=SSRP1 PE=1 SV=1 | 0.972 | 0.929783322 | 0.989 | 0.9 |
| Q96CW1 | AP-2 complex subunit mu OS=Homo sapiens OX=9606 GN=AP2M1 PE=1 SV=2 | 0.975 | 0.91574315 | 0.992 | 0.7 |
| Q8TDQ7 | Glucosamine-6-phosphate isomerase 2 OS=Homo sapiens OX=9606 GN=GNPDA2 PE=1 SV=1 | 1.054 | 0.859267486 | 0.921 | 0.7 |
| P62851 | 40S ribosomal protein S25 OS=Homo sapiens OX=9606 GN=RPS25 PE=1 SV=1 | 1.165 | 0.754126086 | 1.148 | 0.7 |
| H0YDU8 | Serine/threonine-protein phosphatase (Fragment) OS=Homo sapiens OX=9606 GN=PPP5C PE=1 SV=1 | 0.994 | 0.904500837 | 1.006 | 0.7 |
| Q9UKY7 | Protein CDV3 homolog OS=Homo sapiens OX=9606 GN=CDV3 PE=1 SV=1 | 1.013 | 0.927148669 | 0.997 | 0.7 |
| P10155 | 60 kDa SS-A/Ro ribonucleoprotein OS=Homo sapiens OX=9606 GN=RO60 PE=1 SV=2 | 1.016 | 0.881113553 | 0.963 | 0.7 |
| P09874 | Poly [ADP-ribose] polymerase 1 OS=Homo sapiens OX=9606 GN=PARP1 PE=1 SV=4 | 1.078 | 0.900915273 | 0.929 | 0.8 |
| Q9UJ70 | N-acetyl-D-glucosamine kinase OS=Homo sapiens OX=9606 GN=NAGK PE=1 SV=4 | 0.99 | 0.92320619 | 0.983 | 0.7 |
| Q96C19 | EF-hand domain-containing protein D2 OS=Homo sapiens OX=9606 GN=EFHD2 PE=1 SV=1 | 1.026 | 0.765937763 | 1.005 | 0.8 |
| A0A2R8YFH0 | DNA mismatch repair protein OS=Homo sapiens OX=9606 GN=MSH2 PE=1 SV=1 | 1.065 | 0.835808354 | 0.954 | 0.7 |
| P35590 | Tyrosine-protein kinase receptor Tie-1 OS=Homo sapiens OX=9606 GN=TIE1 PE=1 SV=1 | 0.95 | 0.943791335 | 0.823 | 0.7 |
| P29317 | Ephrin type-A receptor 2 OS=Homo sapiens OX=9606 GN=EPHA2 PE=1 SV=2 | 0.979 | 0.750023665 | 0.969 | 0.7 |
| P23786 | Carnitine O-palmitoyltransferase 2, mitochondrial OS=Homo sapiens OX=9606 GN=CPT2 PE=1 SV=2 | 1.023 | 0.811788015 | 0.961 | 0.7 |
| H0YNE9 | Ras-related protein Rab-8B (Fragment) OS=Homo sapiens OX=9606 GN=RAB8B PE=1 SV=1 | 1.044 | 0.854598401 | 0.923 | 0.7 |
| H0YMG7 | Retinal dehydrogenase 2 OS=Homo sapiens OX=9606 GN=ALDH1A2 PE=1 SV=1 | 0.987 | 0.809976184 | 0.986 | 0.7 |
| P10619 | Lysosomal protective protein OS=Homo sapiens OX=9606 GN=CTSA PE=1 SV=2 | 0.962 | 0.829390484 | 0.913 | 0.7 |
| P51114 | Fragile X mental retardation syndrome-related protein 1 OS=Homo sapiens OX=9606 GN=FXR1 PE=1 SV=3 | 1.045 | 0.765235202 | 1.026 | 0.9 |
| K7ES25 | Xaa-Pro dipeptidase (Fragment) OS=Homo sapiens OX=9606 GN=PEPD PE=1 SV=3 | 1.002 | 0.956825234 | 0.948 | 0.7 |
| O43399 | Tumor protein D54 OS=Homo sapiens OX=9606 GN=TPD52L2 PE=1 SV=2 | 1.03 | 0.774484855 | 1.036 | 0.7 |
| Q8WUP2 | Filamin-binding LIM protein 1 OS=Homo sapiens OX=9606 GN=FBLIM1 PE=1 SV=2 | 0.951 | 0.780884711 | 0.949 | 0.7 |
| Q8TAT6 | Nuclear protein localization protein 4 homolog OS=Homo sapiens OX=9606 GN=NPLOC4 PE=1 SV=3 | 0.989 | 0.905868507 | 0.991 | 0.7 |
| P29992 | Guanine nucleotide-binding protein subunit alpha-11 OS=Homo sapiens OX=9606 GN=GNA11 PE=1 SV=2 | 1.111 | 0.806327385 | 0.968 | 0.7 |
| P13473 | Lysosome-associated membrane glycoprotein 2 OS=Homo sapiens OX=9606 GN=LAMP2 PE=1 SV=2 | 0.806 | 0.765235202 | 0.895 | 0.7 |
| O60518 | Ran-binding protein 6 OS=Homo sapiens OX=9606 GN=RANBP6 PE=1 SV=2 | 1.003 | 0.996637279 | 0.883 | 0.7 |
| Q13564 | NEDD8-activating enzyme E1 regulatory subunit OS=Homo sapiens OX=9606 GN=NAE1 PE=1 SV=1 | 0.975 | 0.895384791 | 0.936 | 0.7 |
| E9PEP6 | Neuropilin OS=Homo sapiens OX=9606 GN=NRP1 PE=1 SV=2 | 0.985 | 0.866300162 | 1.005 | 0.8 |
| O60313 | Dynamin-like 120 kDa protein, mitochondrial OS=Homo sapiens OX=9606 GN=OPA1 PE=1 SV=3 | 0.998 | 0.872474259 | 0.901 | 0.7 |
| O00592 | Podocalyxin OS=Homo sapiens OX=9606 GN=PODXL PE=1 SV=2 | 0.958 | 0.750023665 | 0.989 | 0.7 |
| Q9GZS3 | WD repeat-containing protein 61 OS=Homo sapiens OX=9606 GN=WDR61 PE=1 SV=1 | 1.068 | 0.860909277 | 0.966 | 0.7 |
| Q9HB71 | Calcyclin-binding protein OS=Homo sapiens OX=9606 GN=CACYBP PE=1 SV=2 | 0.963 | 0.765235202 | 1.03 | 0.7 |
| Q10713 | Mitochondrial-processing peptidase subunit alpha OS=Homo sapiens OX=9606 GN=PMPCA PE=1 SV=2 | 0.997 | 0.980966594 | 0.938 | 0.7 |
| O00203 | AP-3 complex subunit beta-1 OS=Homo sapiens OX=9606 GN=AP3B1 PE=1 SV=3 | 1.005 | 0.792509319 | 0.991 | 0.8 |
| Q99623 | Prohibitin-2 OS=Homo sapiens OX=9606 GN=PHB2 PE=1 SV=2 | 1.002 | 0.913279108 | 0.791 | 0.7 |
| Q9UHG3 | Prenylcysteine oxidase 1 OS=Homo sapiens OX=9606 GN=PCYOX1 PE=1 SV=3 | 0.936 | 0.750023665 | 0.903 | 0.7 |
| P62854 | 40S ribosomal protein S26 OS=Homo sapiens OX=9606 GN=RPS26 PE=1 SV=3 | 1.103 | 0.750023665 | 1.158 | 0.7 |
| K7ENA8 | Eukaryotic translation initiation factor 3 subunit G (Fragment) OS=Homo sapiens OX=9606 GN=EIF3G PE=1 SV=1 | 0.995 | 0.902225702 | 1.04 | 0.7 |
| P40616 | ADP-ribosylation factor-like protein 1 OS=Homo sapiens OX=9606 GN=ARL1 PE=1 SV=1 | 0.955 | 0.750023665 | 1.01 | 0.8 |
| Q8IUE6 | Histone H2A type 2-B OS=Homo sapiens OX=9606 GN=H2AC21 PE=1 SV=3 | 1.032 | 0.944284199 | 1.003 | 0.9 |
| Q9ULC4 | Malignant T-cell-amplified sequence 1 OS=Homo sapiens OX=9606 GN=MCTS1 PE=1 SV=1 | 1.071 | 0.750023665 | 0.96 | 0.7 |
| Q13162 | Peroxiredoxin-4 OS=Homo sapiens OX=9606 GN=PRDX4 PE=1 SV=1 | 1.007 | 0.939024443 | 1.066 | 0.7 |
| P35080 | Profilin-2 OS=Homo sapiens OX=9606 GN=PFN2 PE=1 SV=3 | 1.004 | 0.978693882 | 0.972 | 0.7 |
| Q13177 | Serine/threonine-protein kinase PAK 2 OS=Homo sapiens OX=9606 GN=PAK2 PE=1 SV=3 | 1.03 | 0.835336742 | 0.997 | 0.8 |
| Q9Y6M1 | Insulin-like growth factor 2 mRNA-binding protein 2 OS=Homo sapiens OX=9606 GN=IGF2BP2 PE=1 SV=2 | 0.999 | 0.955325044 | 0.936 | 0.7 |
| Q969G6 | Riboflavin kinase OS=Homo sapiens OX=9606 GN=RFK PE=1 SV=2 | 0.918 | 0.766667967 | 0.826 | 0.7 |
| Q16629 | Serine/arginine-rich splicing factor 7 OS=Homo sapiens OX=9606 GN=SRSF7 PE=1 SV=1 | 0.969 | 0.765235202 | 0.966 | 0.7 |
| Q9UHB9 | Signal recognition particle subunit SRP68 OS=Homo sapiens OX=9606 GN=SRP68 PE=1 SV=2 | 0.99 | 0.985324977 | 0.993 | 0.9 |
| A0A087X0K9 | Tight junction protein ZO-1 OS=Homo sapiens OX=9606 GN=TJP1 PE=1 SV=1 | 1 | 0.957217311 | 0.95 | 0.7 |
| P33151 | Cadherin-5 OS=Homo sapiens OX=9606 GN=CDH5 PE=1 SV=5 | 0.977 | 0.827085425 | 0.811 | 0.7 |
| O00161 | Synaptosomal-associated protein 23 OS=Homo sapiens OX=9606 GN=SNAP23 PE=1 SV=1 | 0.969 | 0.750023665 | 0.904 | 0.7 |
| O75165 | DnaJ homolog subfamily C member 13 OS=Homo sapiens OX=9606 GN=DNAJC13 PE=1 SV=5 | 0.925 | 0.750023665 | 0.948 | 0.7 |
| Q92626 | Peroxidasin homolog OS=Homo sapiens OX=9606 GN=PXDN PE=1 SV=2 | 1.013 | 0.905369216 | 0.896 | 0.7 |
| Q12768 | WASH complex subunit 5 OS=Homo sapiens OX=9606 GN=WASHC5 PE=1 SV=1 | 0.944 | 0.882454148 | 0.914 | 0.7 |
| Q96FQ6 | Protein S100-A16 OS=Homo sapiens OX=9606 GN=S100A16 PE=1 SV=1 | 0.928 | 0.765235202 | 0.942 | 0.7 |
| Q03252 | Lamin-B2 OS=Homo sapiens OX=9606 GN=LMNB2 PE=1 SV=4 | 1.097 | 0.984200071 | 0.932 | 0.7 |
| O15145 | Actin-related protein 2/3 complex subunit 3 OS=Homo sapiens OX=9606 GN=ARPC3 PE=1 SV=3 | 0.962 | 0.765235202 | 0.955 | 0.7 |
| Q9Y5L0-1 | Isoform 1 of Transportin-3 OS=Homo sapiens OX=9606 GN=TNPO3 | 1.043 | 0.765235202 | 1.007 | 0.9 |
| P48739 | Phosphatidylinositol transfer protein beta isoform OS=Homo sapiens OX=9606 GN=PITPNB PE=1 SV=2 | 0.974 | 0.80706255 | 0.968 | 0.7 |
| Q92769 | Histone deacetylase 2 OS=Homo sapiens OX=9606 GN=HDAC2 PE=1 SV=2 | 1.059 | 0.766667967 | 0.944 | 0.7 |
| P54920 | Alpha-soluble NSF attachment protein OS=Homo sapiens OX=9606 GN=NAPA PE=1 SV=3 | 0.938 | 0.765235202 | 0.95 | 0.7 |
| Q12904 | Aminoacyl tRNA synthase complex-interacting multifunctional protein 1 OS=Homo sapiens OX=9606 GN=AIMP1 PE=1 SV=2 | 1.019 | 0.904500837 | 1.01 | 0.9 |
| P15153 | Ras-related C3 botulinum toxin substrate 2 OS=Homo sapiens OX=9606 GN=RAC2 PE=1 SV=1 | 0.991 | 0.883217803 | 1.073 | 0.7 |
| P14868 | Aspartate--tRNA ligase, cytoplasmic OS=Homo sapiens OX=9606 GN=DARS1 PE=1 SV=2 | 1.012 | 0.967779172 | 0.985 | 0.8 |
| A0A087X0K1 | Calcium-binding protein 39 OS=Homo sapiens OX=9606 GN=CAB39 PE=1 SV=1 | 0.98 | 0.878144903 | 1.038 | 0.7 |
| E9PL57 | NEDD8 (Fragment) OS=Homo sapiens OX=9606 GN=NEDD8-MDP1 PE=3 SV=1 | 1.023 | 0.951324063 | 0.842 | 0.7 |
| Q96HC4 | PDZ and LIM domain protein 5 OS=Homo sapiens OX=9606 GN=PDLIM5 PE=1 SV=5 | 1.021 | 0.840158726 | 0.995 | 0.9 |
| P29474 | Nitric oxide synthase, endothelial OS=Homo sapiens OX=9606 GN=NOS3 PE=1 SV=4 | 1.004 | 0.984200071 | 1.029 | 0.8 |
| Q9UDR5 | Alpha-aminoadipic semialdehyde synthase, mitochondrial OS=Homo sapiens OX=9606 GN=AASS PE=1 SV=1 | 1.003 | 0.924631778 | 0.941 | 0.7 |
| Q9NVP1 | ATP-dependent RNA helicase DDX18 OS=Homo sapiens OX=9606 GN=DDX18 PE=1 SV=2 | 1.007 | 0.841965383 | 0.907 | 0.7 |
| Q9H0A0 | RNA cytidine acetyltransferase OS=Homo sapiens OX=9606 GN=NAT10 PE=1 SV=2 | 1.028 | 0.902016292 | 0.937 | 0.7 |
| O60488 | Long-chain-fatty-acid--CoA ligase 4 OS=Homo sapiens OX=9606 GN=ACSL4 PE=1 SV=2 | 0.978 | 0.996578046 | 1.003 | 0.9 |
| Q13596 | Sorting nexin-1 OS=Homo sapiens OX=9606 GN=SNX1 PE=1 SV=3 | 0.982 | 0.882454148 | 0.948 | 0.7 |
| P32455 | Guanylate-binding protein 1 OS=Homo sapiens OX=9606 GN=GBP1 PE=1 SV=2 | 0.992 | 0.930855729 | 0.943 | 0.7 |
| O43598 | 2'-deoxynucleoside 5'-phosphate N-hydrolase 1 OS=Homo sapiens OX=9606 GN=DNPH1 PE=1 SV=1 | 1.014 | 0.829390484 | 1 | 0.9 |
| P98082 | Disabled homolog 2 OS=Homo sapiens OX=9606 GN=DAB2 PE=1 SV=3 | 0.978 | 0.935312766 | 0.866 | 0.7 |
| O75146 | Huntingtin-interacting protein 1-related protein OS=Homo sapiens OX=9606 GN=HIP1R PE=1 SV=2 | 0.988 | 0.889450577 | 0.897 | 0.7 |
| P18077 | 60S ribosomal protein L35a OS=Homo sapiens OX=9606 GN=RPL35A PE=1 SV=2 | 1.027 | 0.83113959 | 1.095 | 0.7 |
| Q9Y3B8 | Oligoribonuclease, mitochondrial OS=Homo sapiens OX=9606 GN=REXO2 PE=1 SV=3 | 1.07 | 0.803463851 | 0.942 | 0.7 |
| Q6FI81 | Anamorsin OS=Homo sapiens OX=9606 GN=CIAPIN1 PE=1 SV=2 | 0.974 | 0.765235202 | 1.018 | 0.7 |
| Q96HE7 | ERO1-like protein alpha OS=Homo sapiens OX=9606 GN=ERO1A PE=1 SV=2 | 0.974 | 0.766667967 | 0.981 | 0.7 |
| Q9UNZ2 | NSFL1 cofactor p47 OS=Homo sapiens OX=9606 GN=NSFL1C PE=1 SV=2 | 1.034 | 0.80706255 | 0.949 | 0.7 |
| Q9NZZ3 | Charged multivesicular body protein 5 OS=Homo sapiens OX=9606 GN=CHMP5 PE=1 SV=1 | 1.026 | 0.902016292 | 0.947 | 0.7 |
| P17174 | Aspartate aminotransferase, cytoplasmic OS=Homo sapiens OX=9606 GN=GOT1 PE=1 SV=3 | 0.982 | 0.765235202 | 0.97 | 0.7 |
| Q9BVK6 | Transmembrane emp24 domain-containing protein 9 OS=Homo sapiens OX=9606 GN=TMED9 PE=1 SV=2 | 1.046 | 0.905868507 | 1.016 | 0.8 |
| O00567 | Nucleolar protein 56 OS=Homo sapiens OX=9606 GN=NOP56 PE=1 SV=4 | 1.013 | 0.967795554 | 0.887 | 0.7 |
| P36405 | ADP-ribosylation factor-like protein 3 OS=Homo sapiens OX=9606 GN=ARL3 PE=1 SV=2 | 0.963 | 0.902016292 | 0.985 | 0.7 |

| | | | | | |
|---|---|---|---|---|---|
| Q16798 | NADP-dependent malic enzyme, mitochondrial OS=Homo sapiens OX=9606 GN=ME3 PE=2 SV=2 | 1.028 | 0.750023665 | 0.903 | 0.7 |
| Q32P28 | Prolyl 3-hydroxylase 1 OS=Homo sapiens OX=9606 GN=P3H1 PE=1 SV=2 | 0.908 | 0.76858622 | 1.014 | 0.9 |
| A0A1B0GTJ7 | Adenylosuccinate lyase OS=Homo sapiens OX=9606 GN=ADSL PE=1 SV=1 | 0.988 | 0.809976184 | 0.982 | 0.7 |
| Q6UWE0 | E3 ubiquitin-protein ligase LRSAM1 OS=Homo sapiens OX=9606 GN=LRSAM1 PE=1 SV=1 | 1.016 | 0.998901726 | 0.933 | 0.7 |
| A0A0C4DG89 | RNA helicase OS=Homo sapiens OX=9606 GN=DDX46 PE=1 SV=1 | 1.032 | 0.765235202 | 0.975 | 0.7 |
| Q08209 | Serine/threonine-protein phosphatase 2B catalytic subunit alpha isoform OS=Homo sapiens OX=9606 GN=PPP3CA PE=1 SV=1 | 1.052 | 0.765235202 | 0.971 | 0.7 |
| Q13619 | Cullin-4A OS=Homo sapiens OX=9606 GN=CUL4A PE=1 SV=3 | 1.003 | 0.942853463 | 0.99 | 0.8 |
| Q8N1G4 | Leucine-rich repeat-containing protein 47 OS=Homo sapiens OX=9606 GN=LRRC47 PE=1 SV=1 | 1.015 | 0.902016292 | 0.981 | 0.7 |
| Q9NR30 | Nucleolar RNA helicase 2 OS=Homo sapiens OX=9606 GN=DDX21 PE=1 SV=5 | 1.184 | 0.762199181 | 0.984 | 0.9 |
| P61020 | Ras-related protein Rab-5B OS=Homo sapiens OX=9606 GN=RAB5B PE=1 SV=1 | 1.065 | 0.766667967 | 0.968 | 0.8 |
| P06241 | Tyrosine-protein kinase Fyn OS=Homo sapiens OX=9606 GN=FYN PE=1 SV=3 | 0.93 | 0.801301898 | 0.929 | 0.7 |
| P20339 | Ras-related protein Rab-5A OS=Homo sapiens OX=9606 GN=RAB5A PE=1 SV=2 | 1.052 | 0.811788015 | 0.907 | 0.7 |
| Q96RP9 | Elongation factor G, mitochondrial OS=Homo sapiens OX=9606 GN=GFM1 PE=1 SV=2 | 0.967 | 0.750023665 | 0.984 | 0.7 |
| P12931 | Proto-oncogene tyrosine-protein kinase Src OS=Homo sapiens OX=9606 GN=SRC PE=1 SV=3 | 1.002 | 0.985324977 | 0.906 | 0.7 |
| C9JIZ6 | Prosaposin OS=Homo sapiens OX=9606 GN=PSAP PE=1 SV=2 | 0.962 | 0.855041013 | 0.885 | 0.7 |
| Q9Y5X1 | Sorting nexin-9 OS=Homo sapiens OX=9606 GN=SNX9 PE=1 SV=1 | 1.02 | 0.913279108 | 0.902 | 0.7 |
| Q92804 | TATA-binding protein-associated factor 2N OS=Homo sapiens OX=9606 GN=TAF15 PE=1 SV=1 | 0.995 | 0.978562349 | 0.978 | 0.9 |
| Q96AY3 | Peptidyl-prolyl cis-trans isomerase FKBP10 OS=Homo sapiens OX=9606 GN=FKBP10 PE=1 SV=1 | 1.072 | 0.905369216 | 1.048 | 0.7 |
| A0A3B3IT92 | DNA helicase OS=Homo sapiens OX=9606 GN=MCM4 PE=1 SV=1 | 1.018 | 0.840158726 | 1.056 | 0.7 |
| P09012 | U1 small nuclear ribonucleoprotein A OS=Homo sapiens OX=9606 GN=SNRPA PE=1 SV=3 | 0.964 | 0.909475037 | 0.945 | 0.7 |
| A0A590UK80 | Uncharacterized protein (Fragment) OS=Homo sapiens OX=9606 PE=3 SV=1 | 1.006 | 0.978562349 | 0.982 | 0.8 |
| O94855 | Protein transport protein Sec24D OS=Homo sapiens OX=9606 GN=SEC24D PE=1 SV=2 | 0.967 | 0.956825234 | 1.05 | 0.7 |
| O95861 | 3'(2'),5'-bisphosphate nucleotidase 1 OS=Homo sapiens OX=9606 GN=BPNT1 PE=1 SV=1 | 0.947 | 0.765235202 | 0.944 | 0.7 |
| Q14566 | DNA replication licensing factor MCM6 OS=Homo sapiens OX=9606 GN=MCM6 PE=1 SV=1 | 1.139 | 0.951537478 | 1.004 | 0.9 |
| Q6PKG0 | La-related protein 1 OS=Homo sapiens OX=9606 GN=LARP1 PE=1 SV=2 | 1.01 | 0.934553308 | 0.953 | 0.7 |
| O75534 | Cold shock domain-containing protein E1 OS=Homo sapiens OX=9606 GN=CSDE1 PE=1 SV=2 | 1.015 | 0.929783322 | 1.038 | 0.7 |
| Q8N392 | Rho GTPase-activating protein 18 OS=Homo sapiens OX=9606 GN=ARHGAP18 PE=1 SV=3 | 1.011 | 0.994361756 | 0.892 | 0.7 |
| A0A6I8PTT9 | Glutamine--fructose-6-phosphate aminotransferase [isomerizing] 1 OS=Homo sapiens OX=9606 GN=GFPT1 PE=4 SV=1 | 0.994 | 0.921641483 | 1.045 | 0.7 |
| Q9HAV7 | GrpE protein homolog 1, mitochondrial OS=Homo sapiens OX=9606 GN=GRPEL1 PE=1 SV=2 | 0.975 | 0.912604818 | 1.011 | 0.7 |
| P62304 | Small nuclear ribonucleoprotein E OS=Homo sapiens OX=9606 GN=SNRPE PE=1 SV=1 | 0.97 | 0.863796857 | 0.94 | 0.7 |
| P63167 | Dynein light chain 1, cytoplasmic OS=Homo sapiens OX=9606 GN=DYNLL1 PE=1 SV=1 | 1.014 | 0.824101611 | 1.019 | 0.9 |
| P61289 | Proteasome activator complex subunit 3 OS=Homo sapiens OX=9606 GN=PSME3 PE=1 SV=1 | 0.95 | 0.780884711 | 0.998 | 0.9 |
| Q7Z2Z2 | Elongation factor-like GTPase 1 OS=Homo sapiens OX=9606 GN=EFL1 PE=1 SV=2 | 0.998 | 0.983479158 | 0.956 | 0.7 |
| P26196 | Probable ATP-dependent RNA helicase DDX6 OS=Homo sapiens OX=9606 GN=DDX6 PE=1 SV=2 | 1.048 | 0.750023665 | 1.015 | 0.7 |
| G5EA31 | Protein transport protein Sec24C OS=Homo sapiens OX=9606 GN=SEC24C PE=1 SV=1 | 1.012 | 0.96647353 | 1.011 | [illegible] |
| P23919 | Thymidylate kinase OS=Homo sapiens OX=9606 GN=DTYMK PE=1 SV=4 | 1.016 | 0.989102644 | 0.98 | 0.7 |
| P52943 | Cysteine-rich protein 2 OS=Homo sapiens OX=9606 GN=CRIP2 PE=1 SV=1 | 1.028 | 0.761475165 | 0.976 | 0.7 |
| O75116 | Rho-associated protein kinase 2 OS=Homo sapiens OX=9606 GN=ROCK2 PE=1 SV=4 | 1.034 | 0.8215945 | 0.991 | 0.7 |
| P09471-2 | Isoform Alpha-2 of Guanine nucleotide-binding protein G(o) subunit alpha OS=Homo sapiens OX=9606 GN=GNAO1 | 0.896 | 0.765235202 | 0.927 | 0.7 |
| Q9H2G2 | STE20-like serine/threonine-protein kinase OS=Homo sapiens OX=9606 GN=SLK PE=1 SV=1 | 1.045 | 0.750023665 | 0.971 | 0.7 |
| Q9UI26 | Importin-11 OS=Homo sapiens OX=9606 GN=IPO11 PE=1 SV=1 | 0.971 | 0.902016292 | 0.973 | 0.7 |
| Q8NBF2 | NHL repeat-containing protein 2 OS=Homo sapiens OX=9606 GN=NHLRC2 PE=1 SV=1 | 1.077 | 0.688026711 | 0.991 | 0.7 |
| Q9HB07 | MYG1 exonuclease OS=Homo sapiens OX=9606 GN=MYG1 PE=1 SV=3 | 1.02 | 0.961998268 | 0.923 | 0.7 |
| Q9BY32 | Inosine triphosphate pyrophosphatase OS=Homo sapiens OX=9606 GN=ITPA PE=1 SV=2 | 0.999 | 0.885528205 | 0.979 | 0.8 |
| O14744 | Protein arginine N-methyltransferase 5 OS=Homo sapiens OX=9606 GN=PRMT5 PE=1 SV=4 | 0.958 | 0.829390484 | 0.97 | 0.9 |
| Q9BV20 | Methylthioribose-1-phosphate isomerase OS=Homo sapiens OX=9606 GN=MRI1 PE=1 SV=1 | 0.957 | 0.750023665 | 0.953 | 0.7 |
| Q9UNF1 | Melanoma-associated antigen D2 OS=Homo sapiens OX=9606 GN=MAGED2 PE=1 SV=2 | 0.999 | 0.939024443 | 0.948 | 0.9 |
| Q12907 | Vesicular integral-membrane protein VIP36 OS=Homo sapiens OX=9606 GN=LMAN2 PE=1 SV=1 | 1.046 | 0.880337807 | 0.972 | 0.7 |
| D3YTB1 | 60S ribosomal protein L32 (Fragment) OS=Homo sapiens OX=9606 GN=RPL32 PE=1 SV=1 | 1.071 | 0.790808367 | 1.12 | 0.7 |
| Q99614 | Tetratricopeptide repeat protein 1 OS=Homo sapiens OX=9606 GN=TTC1 PE=1 SV=1 | 1.011 | 0.961457894 | 1.026 | 0.8 |
| P26232 | Catenin alpha-2 OS=Homo sapiens OX=9606 GN=CTNNA2 PE=1 SV=5 | 1.184 | 0.902016292 | 0.829 | 0.7 |
| P49006 | MARCKS-related protein OS=Homo sapiens OX=9606 GN=MARCKSL1 PE=1 SV=2 | 0.967 | 0.902016292 | 0.87 | 0.7 |
| Q9H8Y8 | Golgi reassembly-stacking protein 2 OS=Homo sapiens OX=9606 GN=GORASP2 PE=1 SV=3 | 1.03 | 0.765235202 | 0.99 | 0.9 |
| Q9Y295 | Developmentally-regulated GTP-binding protein 1 OS=Homo sapiens OX=9606 GN=DRG1 PE=1 SV=1 | 1.03 | 0.886561233 | 0.981 | 0.7 |
| P78347 | General transcription factor II-I OS=Homo sapiens OX=9606 GN=GTF2I PE=1 SV=2 | 1.004 | 0.989102644 | 0.947 | 0.7 |
| O43681 | ATPase GET3 OS=Homo sapiens OX=9606 GN=GET3 PE=1 SV=2 | 0.971 | 0.908928234 | 0.909 | 0.7 |
| P49721 | Proteasome subunit beta type-2 OS=Homo sapiens OX=9606 GN=PSMB2 PE=1 SV=1 | 0.893 | 0.750023665 | 0.969 | 0.7 |
| A0A140T9R1 | Flotillin (Fragment) OS=Homo sapiens OX=9606 GN=FLOT1 PE=1 SV=1 | 0.971 | 0.868906519 | 0.901 | 0.7 |
| P23634 | Plasma membrane calcium-transporting ATPase 4 OS=Homo sapiens OX=9606 GN=ATP2B4 PE=1 SV=2 | 0.961 | 0.845656075 | 0.899 | 0.7 |
| Q96BM9 | ADP-ribosylation factor-like protein 8A OS=Homo sapiens OX=9606 GN=ARL8A PE=1 SV=1 | 1.016 | 0.806303314 | 0.947 | 0.7 |
| D6RIR6 | 3-hydroxybutyrate dehydrogenase type 2 (Fragment) OS=Homo sapiens OX=9606 GN=BDH2 PE=1 SV=1 | 0.939 | 0.837893552 | 0.924 | 0.7 |
| Q5JPE7 | Nodal modulator 2 OS=Homo sapiens OX=9606 GN=NOMO2 PE=1 SV=1 | 0.969 | 0.985324977 | 0.952 | 0.9 |
| A0A2R8Y5Q8 | Tubulin-folding cofactor E OS=Homo sapiens OX=9606 GN=TBCE PE=1 SV=1 | 0.956 | 0.765235202 | 0.915 | 0.7 |
| Q6DKJ4 | Nucleoredoxin OS=Homo sapiens OX=9606 GN=NXN PE=1 SV=2 | 1.035 | 0.829390484 | 0.866 | 0.7 |
| Q92734 | Protein TFG OS=Homo sapiens OX=9606 GN=TFG PE=1 SV=2 | 1.028 | 0.843246212 | 1.006 | 0.9 |
| O95202 | Mitochondrial proton/calcium exchanger protein OS=Homo sapiens OX=9606 GN=LETM1 PE=1 SV=1 | 1.019 | 0.994361756 | 0.912 | 0.7 |
| A0A0A0MSB8 | Exocyst complex component 7 OS=Homo sapiens OX=9606 GN=EXOC7 PE=1 SV=1 | 1.038 | 0.893277754 | 0.981 | 0.7 |
| P21283 | V-type proton ATPase subunit C 1 OS=Homo sapiens OX=9606 GN=ATP6V1C1 PE=1 SV=4 | 0.964 | 0.815069882 | 0.962 | 0.7 |
| Q9Y3A5 | Ribosome maturation protein SBDS OS=Homo sapiens OX=9606 GN=SBDS PE=1 SV=4 | 1.034 | 0.886561233 | 1.013 | 0.7 |
| Q15370 | Elongin-B OS=Homo sapiens OX=9606 GN=ELOB PE=1 SV=1 | 0.938 | 0.829390484 | 0.922 | 0.7 |
| X6RGP5 | Protein transport protein sec16 (Fragment) OS=Homo sapiens OX=9606 GN=SEC16A PE=1 SV=1 | 1.028 | 0.869995963 | 0.944 | 0.7 |
| P22061 | Protein-L-isoaspartate(D-aspartate) O-methyltransferase OS=Homo sapiens OX=9606 GN=PCMT1 PE=1 SV=4 | 0.982 | 0.863882725 | 0.985 | 0.7 |
| H0YKD8 | 60S ribosomal protein L28 OS=Homo sapiens OX=9606 GN=RPL28 PE=1 SV=1 | 1.076 | 0.750023665 | 1.109 | 0.7 |
| P62330 | ADP-ribosylation factor 6 OS=Homo sapiens OX=9606 GN=ARF6 PE=1 SV=2 | 1.109 | 0.810840988 | 0.938 | 0.7 |
| P63151 | Serine/threonine-protein phosphatase 2A 55 kDa regulatory subunit B alpha isoform OS=Homo sapiens OX=9606 GN=PPP2R2A PE | 0.964 | 0.889450577 | 0.965 | 0.7 |
| A0A0J9YXF2 | Paraoxonase 2, isoform CRA_a OS=Homo sapiens OX=9606 GN=PON2 PE=1 SV=1 | 0.999 | 0.917150381 | 0.888 | 0.7 |
| D9J307 | HLA class I antigen OS=Homo sapiens OX=9606 GN=HLA-B PE=1 SV=1 | 0.891 | 0.765235202 | 0.861 | 0.7 |
| Q2M389 | WASH complex subunit 4 OS=Homo sapiens OX=9606 GN=WASHC4 PE=1 SV=2 | 0.955 | 0.750023665 | 0.988 | 0.7 |
| Q8NBX0 | Saccharopine dehydrogenase-like oxidoreductase OS=Homo sapiens OX=9606 GN=SCCPDH PE=1 SV=1 | 0.999 | 0.985324977 | 0.972 | 0.7 |
| P00390 | Glutathione reductase, mitochondrial OS=Homo sapiens OX=9606 GN=GSR PE=1 SV=2 | 0.974 | 0.882454148 | 0.985 | 0.8 |
| P50148 | Guanine nucleotide-binding protein G(q) subunit alpha OS=Homo sapiens OX=9606 GN=GNAQ PE=1 SV=4 | 1.007 | 0.994361756 | 0.985 | 0.7 |
| Q09161 | Nuclear cap-binding protein subunit 1 OS=Homo sapiens OX=9606 GN=NCBP1 PE=1 SV=1 | 0.963 | 0.845656075 | 0.977 | 0.7 |
| P54709 | Sodium/potassium-transporting ATPase subunit beta-3 OS=Homo sapiens OX=9606 GN=ATP1B3 PE=1 SV=1 | 1.023 | 0.889482529 | 0.947 | 0.7 |
| Q14141 | Septin-6 OS=Homo sapiens OX=9606 GN=SEPTIN6 PE=1 SV=4 | 0.988 | 0.994361756 | 0.877 | 0.7 |
| P55735 | Protein SEC13 homolog OS=Homo sapiens OX=9606 GN=SEC13 PE=1 SV=3 | 0.97 | 0.984805107 | 1.041 | 0. |
| P35914 | Hydroxymethylglutaryl-CoA lyase, mitochondrial OS=Homo sapiens OX=9606 GN=HMGCL PE=1 SV=2 | 1.016 | 0.829390484 | 0.93 | 0.7 |
| O00442 | RNA 3'-terminal phosphate cyclase OS=Homo sapiens OX=9606 GN=RTCA PE=1 SV=1 | 1.049 | 0.765235202 | 0.914 | 0.7 |
| P14543 | Nidogen-1 OS=Homo sapiens OX=9606 GN=NID1 PE=1 SV=3 | 0.991 | 0.95071661 | 0.887 | 0.7 |
| Q9UJU6 | Drebrin-like protein OS=Homo sapiens OX=9606 GN=DBNL PE=1 SV=1 | 0.812 | 0.937978988 | 0.98 | 0.8 |
| Q5JTH9 | RRP12-like protein OS=Homo sapiens OX=9606 GN=RRP12 PE=1 SV=2 | 1.008 | 0.953792095 | 0.898 | 0.7 |
| F8VXU5 | Vacuolar protein sorting-associated protein 29 OS=Homo sapiens OX=9606 GN=VPS29 PE=1 SV=1 | 0.967 | 0.765235202 | 0.942 | 0.7 |
| A0A3B3IUA2 | Ribonucleoprotein OS=Homo sapiens OX=9606 GN=SNU13 PE=1 SV=1 | 1.011 | 0.918114651 | 0.894 | 0.9 |
| Q15645 | Pachytene checkpoint protein 2 homolog OS=Homo sapiens OX=9606 GN=TRIP13 PE=1 SV=2 | 0.988 | 0.900868056 | 0.966 | 0.7 |
| Q71UI9 | Histone H2A.V OS=Homo sapiens OX=9606 GN=H2AZ2 PE=1 SV=3 | 0.913 | 0.765235202 | 0.98 | 0.7 |
| Q9UBQ5 | Eukaryotic translation initiation factor 3 subunit K OS=Homo sapiens OX=9606 GN=EIF3K PE=1 SV=1 | 1.021 | 0.825672127 | 1.004 | 0.8 |
| Q9BWS9 | Chitinase domain-containing protein 1 OS=Homo sapiens OX=9606 GN=CHID1 PE=1 SV=1 | 1.055 | 0.765235202 | 0.939 | 0.7 |

| | | | | | |
|---|---|---|---|---|---|
| P17096 | High mobility group protein HMG-I/HMG-Y OS=Homo sapiens OX=9606 GN=HMGA1 PE=1 SV=3 | 1.28 | 0.863796857 | 0.966 | 0.8 |
| P10301 | Ras-related protein R-Ras OS=Homo sapiens OX=9606 GN=RRAS PE=1 SV=1 | 0.96 | 0.834564551 | 0.95 | 0.7 |
| P30043 | Flavin reductase (NADPH) OS=Homo sapiens OX=9606 GN=BLVRB PE=1 SV=3 | 0.967 | 0.876510992 | 0.993 | 0.7 |
| C9J177 | Protein phosphatase 1 regulatory subunit 7 (Fragment) OS=Homo sapiens OX=9606 GN=PPP1R7 PE=1 SV=1 | 0.999 | 0.889450577 | 1.036 | 0.7 |
| P61081 | NEDD8-conjugating enzyme Ubc12 OS=Homo sapiens OX=9606 GN=UBE2M PE=1 SV=1 | 0.979 | 0.789400225 | 0.963 | 0.7 |
| Q969H8 | Myeloid-derived growth factor OS=Homo sapiens OX=9606 GN=MYDGF PE=1 SV=1 | 0.944 | 0.750023665 | 1.018 | 0.7 |
| H7C4T5 | Muscleblind-like protein 1 (Fragment) OS=Homo sapiens OX=9606 GN=MBNL1 PE=1 SV=1 | 1.046 | 0.765235202 | 0.998 | 0.9 |
| Q9NVZ3 | Adaptin ear-binding coat-associated protein 2 OS=Homo sapiens OX=9606 GN=NECAP2 PE=1 SV=1 | 0.996 | 0.992258896 | 0.961 | 0.7 |
| P61254 | 60S ribosomal protein L26 OS=Homo sapiens OX=9606 GN=RPL26 PE=1 SV=1 | 1.041 | 0.76858622 | 1.111 | 0.7 |
| P49257 | Protein ERGIC-53 OS=Homo sapiens OX=9606 GN=LMAN1 PE=1 SV=2 | 1.027 | 0.930855729 | 1.002 | 0.8 |
| P51813 | Cytoplasmic tyrosine-protein kinase BMX OS=Homo sapiens OX=9606 GN=BMX PE=1 SV=1 | 1.007 | 0.829390484 | 0.913 | 0.7 |
| P16278 | Beta-galactosidase OS=Homo sapiens OX=9606 GN=GLB1 PE=1 SV=2 | 0.964 | 0.750023665 | 0.894 | 0.7 |
| Q9Y6W5 | Wiskott-Aldrich syndrome protein family member 2 OS=Homo sapiens OX=9606 GN=WASF2 PE=1 SV=3 | 1.02 | 0.765235202 | 0.948 | 0.7 |
| P05198 | Eukaryotic translation initiation factor 2 subunit 1 OS=Homo sapiens OX=9606 GN=EIF2S1 PE=1 SV=3 | 1.065 | 0.884127064 | 1.07 | 0. |
| Q16836 | Hydroxyacyl-coenzyme A dehydrogenase, mitochondrial OS=Homo sapiens OX=9606 GN=HADH PE=1 SV=3 | 0.985 | 0.765235202 | 0.92 | 0.7 |
| G3XAC6 | RNA-binding protein 39 OS=Homo sapiens OX=9606 GN=RBM39 PE=1 SV=1 | 1.022 | 0.886561233 | 0.965 | 0.8 |
| Q5U651 | Ras-interacting protein 1 OS=Homo sapiens OX=9606 GN=RASIP1 PE=1 SV=1 | 1.177 | 0.642276967 | 0.935 | 0.7 |
| Q13907 | Isopentenyl-diphosphate Delta-isomerase 1 OS=Homo sapiens OX=9606 GN=IDI1 PE=1 SV=2 | 1.124 | 0.940631922 | 1.093 | 0.9 |
| Q92896 | Golgi apparatus protein 1 OS=Homo sapiens OX=9606 GN=GLG1 PE=1 SV=2 | 0.982 | 0.889482529 | 0.999 | 0.9 |
| Q99584 | Protein S100-A13 OS=Homo sapiens OX=9606 GN=S100A13 PE=1 SV=1 | 1.008 | 0.904500837 | 1.091 | 0.7 |
| P30419 | Glycylpeptide N-tetradecanoyltransferase 1 OS=Homo sapiens OX=9606 GN=NMT1 PE=1 SV=2 | 0.956 | 0.920144076 | 0.96 | 0.7 |
| P28072 | Proteasome subunit beta type-6 OS=Homo sapiens OX=9606 GN=PSMB6 PE=1 SV=4 | 0.952 | 0.750023665 | 1.119 | 0.8 |
| P43034 | Platelet-activating factor acetylhydrolase IB subunit alpha OS=Homo sapiens OX=9606 GN=PAFAH1B1 PE=1 SV=2 | 1.036 | 0.912636218 | 0.971 | 0.7 |
| Q99471 | Prefoldin subunit 5 OS=Homo sapiens OX=9606 GN=PFDN5 PE=1 SV=2 | 1.008 | 0.876510992 | 1.023 | 0.8 |
| F5GZ78 | Paxillin OS=Homo sapiens OX=9606 GN=PXN PE=1 SV=1 | 1.051 | 0.80706255 | 1.02 | 0.7 |
| P17931 | Galectin-3 OS=Homo sapiens OX=9606 GN=LGALS3 PE=1 SV=5 | 1.037 | 0.809976184 | 1.002 | 0.8 |
| Q9UMR2 | ATP-dependent RNA helicase DDX19B OS=Homo sapiens OX=9606 GN=DDX19B PE=1 SV=1 | 1.092 | 0.784693444 | 0.997 | 0.9 |
| Q9NUU7 | ATP-dependent RNA helicase DDX19A OS=Homo sapiens OX=9606 GN=DDX19A PE=1 SV=1 | 0.991 | 0.946055791 | 1.024 | 0.9 |
| A0A590UK22 | Unconventional myosin-VI (Fragment) OS=Homo sapiens OX=9606 GN=MYO6 PE=1 SV=1 | 0.909 | 0.750023665 | 0.965 | 0.8 |
| Q9ULZ3 | Apoptosis-associated speck-like protein containing a CARD OS=Homo sapiens OX=9606 GN=PYCARD PE=1 SV=2 | 0.971 | 0.829390484 | 0.955 | 0.7 |
| O15020 | Spectrin beta chain, non-erythrocytic 2 OS=Homo sapiens OX=9606 GN=SPTBN2 PE=1 SV=3 | 0.978 | 0.902777383 | 1.461 | 0.7 |
| Q12996 | Cleavage stimulation factor subunit 3 OS=Homo sapiens OX=9606 GN=CSTF3 PE=1 SV=1 | 1.036 | 0.854598401 | 0.964 | 0.7 |
| Q9Y624 | Junctional adhesion molecule A OS=Homo sapiens OX=9606 GN=F11R PE=1 SV=1 | 0.986 | 0.904500837 | 0.939 | 0.7 |
| O75400 | Pre-mRNA-processing factor 40 homolog A OS=Homo sapiens OX=9606 GN=PRPF40A PE=1 SV=2 | 1.024 | 0.865970839 | 0.925 | 0.7 |
| O60271 | C-Jun-amino-terminal kinase-interacting protein 4 OS=Homo sapiens OX=9606 GN=SPAG9 PE=1 SV=4 | 1.067 | 0.829390484 | 1.044 | 0.9 |
| P60866 | 40S ribosomal protein S20 OS=Homo sapiens OX=9606 GN=RPS20 PE=1 SV=1 | 1.113 | 0.761475165 | 1.181 | 0.7 |
| Q7LG56 | Ribonucleoside-diphosphate reductase subunit M2 B OS=Homo sapiens OX=9606 GN=RRM2B PE=1 SV=1 | 1.008 | 0.946055791 | 0.913 | 0.7 |
| P19174 | 1-phosphatidylinositol 4,5-bisphosphate phosphodiesterase gamma-1 OS=Homo sapiens OX=9606 GN=PLCG1 PE=1 SV=1 | 1.038 | 0.886561233 | 0.953 | 0.7 |
| Q96FS4 | Signal-induced proliferation-associated protein 1 OS=Homo sapiens OX=9606 GN=SIPA1 PE=1 SV=1 | 1.016 | 0.989102644 | 0.933 | 0.9 |
| Q9NVJ2 | ADP-ribosylation factor-like protein 8B OS=Homo sapiens OX=9606 GN=ARL8B PE=1 SV=1 | 1.121 | 0.765235202 | 0.963 | 0.8 |
| Q9NR12 | PDZ and LIM domain protein 7 OS=Homo sapiens OX=9606 GN=PDLIM7 PE=1 SV=1 | 0.976 | 0.961750547 | 0.926 | 0.7 |
| H7BY94 | Bystin (Fragment) OS=Homo sapiens OX=9606 GN=BYSL PE=1 SV=1 | 0.992 | 0.902016292 | 0.95 | 0.8 |
| Q9UK45 | U6 snRNA-associated Sm-like protein LSm7 OS=Homo sapiens OX=9606 GN=LSM7 PE=1 SV=1 | 0.952 | 0.750023665 | 0.954 | 0.7 |
| P33240 | Cleavage stimulation factor subunit 2 OS=Homo sapiens OX=9606 GN=CSTF2 PE=1 SV=1 | 0.969 | 0.935867662 | 0.919 | 0.7 |
| Q5T3Q7 | HEAT repeat-containing protein 1 OS=Homo sapiens OX=9606 GN=HEATR1 PE=1 SV=1 | 0.956 | 0.889450577 | 0.912 | 0.7 |
| Q9NUQ9 | CYFIP-related Rac1 interactor B OS=Homo sapiens OX=9606 GN=CYRIB PE=1 SV=1 | 0.905 | 0.750023665 | 0.964 | 0.7 |
| Q16666 | Gamma-interferon-inducible protein 16 OS=Homo sapiens OX=9606 GN=IFI16 PE=1 SV=3 | 1.001 | 0.974919191 | 1.001 | 0.9 |
| Q9H4A6 | Golgi phosphoprotein 3 OS=Homo sapiens OX=9606 GN=GOLPH3 PE=1 SV=1 | 1.038 | 0.765235202 | 0.957 | 0.7 |
| P60903 | Protein S100-A10 OS=Homo sapiens OX=9606 GN=S100A10 PE=1 SV=2 | 1.081 | 0.813779274 | 0.887 | 0.7 |
| P42574 | Caspase-3 OS=Homo sapiens OX=9606 GN=CASP3 PE=1 SV=2 | 0.951 | 0.770292944 | 0.945 | 0.7 |
| Q9NRR5 | Ubiquilin-4 OS=Homo sapiens OX=9606 GN=UBQLN4 PE=1 SV=2 | 1.072 | 0.765235202 | 0.966 | 0.7 |
| O43615 | Mitochondrial import inner membrane translocase subunit TIM44 OS=Homo sapiens OX=9606 GN=TIMM44 PE=1 SV=2 | 1.095 | 0.765235202 | 0.906 | 0.7 |
| E9PC52 | Histone-binding protein RBBP7 OS=Homo sapiens OX=9606 GN=RBBP7 PE=1 SV=1 | 1.077 | 0.750023665 | 0.993 | 0.9 |
| O95881 | Thioredoxin domain-containing protein 12 OS=Homo sapiens OX=9606 GN=TXNDC12 PE=1 SV=1 | 0.995 | 0.983953845 | 1.046 | 0.7 |
| P49792 | E3 SUMO-protein ligase RanBP2 OS=Homo sapiens OX=9606 GN=RANBP2 PE=1 SV=2 | 1.019 | 0.965904544 | 1.032 | 0.9 |
| P29279 | CCN family member 2 OS=Homo sapiens OX=9606 GN=CCN2 PE=1 SV=2 | 0.935 | 0.759321117 | 0.972 | 0.7 |
| O95487 | Protein transport protein Sec24B OS=Homo sapiens OX=9606 GN=SEC24B PE=1 SV=2 | 1.002 | 0.95980123 | 0.929 | 0.7 |
| F8VQE1 | LIM domain and actin-binding protein 1 OS=Homo sapiens OX=9606 GN=LIMA1 PE=1 SV=1 | 1.062 | 0.766667967 | 1.018 | 0.8 |
| J3QRN6 | Unconventional myosin-Id OS=Homo sapiens OX=9606 GN=MYO1D PE=1 SV=1 | 0.979 | 0.951537478 | 0.855 | 0.7 |
| Q5VW32 | BRO1 domain-containing protein BROX OS=Homo sapiens OX=9606 GN=BROX PE=1 SV=1 | 0.974 | 0.840158726 | 0.949 | 0.7 |
| F8W031 | DUF3456 domain-containing protein (Fragment) OS=Homo sapiens OX=9606 PE=1 SV=1 | 1.031 | 0.983953845 | 1.089 | 0.7 |
| Q16698 | 2,4-dienoyl-CoA reductase, mitochondrial OS=Homo sapiens OX=9606 GN=DECR1 PE=1 SV=1 | 1.003 | 0.972616875 | 0.935 | 0.7 |
| Q9BTY7 | Protein HGH1 homolog OS=Homo sapiens OX=9606 GN=HGH1 PE=1 SV=1 | 0.981 | 0.929783322 | 0.931 | 0.8 |
| Q9UHV9 | Prefoldin subunit 2 OS=Homo sapiens OX=9606 GN=PFDN2 PE=1 SV=1 | 0.966 | 0.99634713 | 1.01 | 0.7 |
| P09496-2 | Isoform Non-brain of Clathrin light chain A OS=Homo sapiens OX=9606 GN=CLTA | 0.924 | 0.878144903 | 1.006 | 0.9 |
| Q12765 | Secernin-1 OS=Homo sapiens OX=9606 GN=SCRN1 PE=1 SV=2 | 0.98 | 0.983752935 | 0.955 | 0.7 |
| Q9H0D6 | 5'-3' exoribonuclease 2 OS=Homo sapiens OX=9606 GN=XRN2 PE=1 SV=1 | 0.997 | 0.897516642 | 0.96 | 0.7 |
| Q9Y2Z0 | Protein SGT1 homolog OS=Homo sapiens OX=9606 GN=SUGT1 PE=1 SV=3 | 1.021 | 0.750023665 | 1.04 | 0.7 |
| A0A0A0MRA8 | Band 4.1-like protein 3 OS=Homo sapiens OX=9606 GN=EPB41L3 PE=1 SV=1 | 1.003 | 0.983953845 | 0.972 | 0.7 |
| G3V3H3 | Kinesin light chain OS=Homo sapiens OX=9606 GN=KLC1 PE=1 SV=1 | 1.037 | 0.830179531 | 1.033 | 0.7 |
| Q92597 | Protein NDRG1 OS=Homo sapiens OX=9606 GN=NDRG1 PE=1 SV=1 | 1.077 | 0.765235202 | 1.068 | 0.7 |
| P13647 | Keratin, type II cytoskeletal 5 OS=Homo sapiens OX=9606 GN=KRT5 PE=1 SV=3 | 0.867 | 0.765235202 | 0.73 | 0.7 |
| Q9P258 | Protein RCC2 OS=Homo sapiens OX=9606 GN=RCC2 PE=1 SV=2 | 1.028 | 0.811611688 | 0.965 | 0.7 |
| P36543 | V-type proton ATPase subunit E 1 OS=Homo sapiens OX=9606 GN=ATP6V1E1 PE=1 SV=1 | 0.984 | 0.917008272 | 0.97 | 0.7 |
| O75694 | Nuclear pore complex protein Nup155 OS=Homo sapiens OX=9606 GN=NUP155 PE=1 SV=1 | 0.966 | 0.792509319 | 0.917 | 0.7 |
| Q00169 | Phosphatidylinositol transfer protein alpha isoform OS=Homo sapiens OX=9606 GN=PITPNA PE=1 SV=2 | 0.989 | 0.765235202 | 1.013 | 0.8 |
| Q7L9L4 | MOB kinase activator 1B OS=Homo sapiens OX=9606 GN=MOB1B PE=1 SV=3 | 0.955 | 0.750023665 | 0.946 | 0.7 |
| Q9UJU6-3 | Isoform 3 of Drebrin-like protein OS=Homo sapiens OX=9606 GN=DBNL | 1.04 | 0.765235202 | 1.046 | 0.7 |
| O00622 | CCN family member 1 OS=Homo sapiens OX=9606 GN=CCN1 PE=1 SV=1 | 0.929 | 0.889482529 | 0.691 | 0.7 |
| Q9UK76 | Jupiter microtubule associated homolog 1 OS=Homo sapiens OX=9606 GN=JPT1 PE=1 SV=3 | 1.029 | 0.954224541 | 1.18 | 0.7 |
| Q9NZ08 | Endoplasmic reticulum aminopeptidase 1 OS=Homo sapiens OX=9606 GN=ERAP1 PE=1 SV=3 | 0.959 | 0.765235202 | 0.98 | 0.7 |
| Q8NHP8 | Putative phospholipase B-like 2 OS=Homo sapiens OX=9606 GN=PLBD2 PE=1 SV=2 | 0.946 | 0.824101611 | 0.9 | 0.7 |
| O15269 | Serine palmitoyltransferase 1 OS=Homo sapiens OX=9606 GN=SPTLC1 PE=1 SV=1 | 0.988 | 0.978562349 | 0.962 | 0.7 |
| A0A2R8Y7U1 | Tripeptidyl aminopeptidase (Fragment) OS=Homo sapiens OX=9606 GN=TPP1 PE=1 SV=1 | 1.008 | 0.978562349 | 0.935 | 0.7 |
| Q9Y5Z4 | Heme-binding protein 2 OS=Homo sapiens OX=9606 GN=HEBP2 PE=1 SV=1 | 0.958 | 0.829390484 | 1.003 | 0.9 |
| P42356 | Phosphatidylinositol 4-kinase alpha OS=Homo sapiens OX=9606 GN=PI4KA PE=1 SV=4 | 0.953 | 0.750023665 | 0.895 | 0.7 |
| P61011 | Signal recognition particle 54 kDa protein OS=Homo sapiens OX=9606 GN=SRP54 PE=1 SV=1 | 0.968 | 0.993362873 | 1.025 | 0.7 |
| P08240 | Signal recognition particle receptor subunit alpha OS=Homo sapiens OX=9606 GN=SRPRA PE=1 SV=2 | 1.056 | 0.765235202 | 0.987 | 0.8 |
| Q92542 | Nicastrin OS=Homo sapiens OX=9606 GN=NCSTN PE=1 SV=2 | 0.734 | 0.905369216 | 0.891 | 0.7 |
| A0A087WVM4 | Formyltetrahydrofolate synthetase OS=Homo sapiens OX=9606 GN=MTHFD1L PE=1 SV=1 | 0.81 | 0.965945681 | 1.024 | 0.8 |
| Q6L8Q7 | 2',5'-phosphodiesterase 12 OS=Homo sapiens OX=9606 GN=PDE12 PE=1 SV=2 | 1.005 | 0.961457894 | 0.925 | 0.7 |
| E5RFJ3 | Cell cycle and apoptosis regulator protein 2 (Fragment) OS=Homo sapiens OX=9606 GN=CCAR2 PE=1 SV=1 | 0.944 | 0.905369216 | 0.906 | 0.7 |
| P61326 | Protein mago nashi homolog OS=Homo sapiens OX=9606 GN=MAGOH PE=1 SV=1 | 0.968 | 0.902016292 | 0.987 | 0.7 |
| Q96B26 | Exosome complex component RRP43 OS=Homo sapiens OX=9606 GN=EXOSC8 PE=1 SV=1 | 0.971 | 0.902016292 | 0.944 | 0.7 |
| Q04760 | Lactoylglutathione lyase OS=Homo sapiens OX=9606 GN=GLO1 PE=1 SV=4 | 0.907 | 0.978562349 | 0.948 | 0.7 |
| O15260 | Surfeit locus protein 4 OS=Homo sapiens OX=9606 GN=SURF4 PE=1 SV=3 | 0.885 | 0.824559577 | 0.886 | 0.7 |

| | | | | | |
|---|---|---|---|---|---|
| Q9Y5P6 | Mannose-1-phosphate guanyltransferase beta OS=Homo sapiens OX=9606 GN=GMPPB PE=1 SV=2 | 0.988 | 0.996637279 | 1.01 | 0.9 |
| Q13557 | Calcium/calmodulin-dependent protein kinase type II subunit delta OS=Homo sapiens OX=9606 GN=CAMK2D PE=1 SV=3 | 1.005 | 0.995582922 | 0.909 | 0.7 |
| Q9Y2W1 | Thyroid hormone receptor-associated protein 3 OS=Homo sapiens OX=9606 GN=THRAP3 PE=1 SV=2 | 0.884 | 0.949407053 | 0.943 | 0.7 |
| M0R299 | rRNA 2'-O-methyltransferase fibrillarin (Fragment) OS=Homo sapiens OX=9606 GN=FBL PE=1 SV=1 | 0.964 | 0.904500837 | 0.908 | 0.8 |
| Q13126 | S-methyl-5'-thioadenosine phosphorylase OS=Homo sapiens OX=9606 GN=MTAP PE=1 SV=2 | 1.009 | 0.9252392 | 0.964 | 0.7 |
| A0A0A0MSN4 | Angiotensin-converting enzyme OS=Homo sapiens OX=9606 GN=ACE PE=1 SV=1 | 1.002 | 0.949934729 | 0.863 | 0.7 |
| Q13451 | Peptidyl-prolyl cis-trans isomerase FKBP5 OS=Homo sapiens OX=9606 GN=FKBP5 PE=1 SV=2 | 0.966 | 0.902016292 | 0.973 | 0.9 |
| Q96AX1 | Vacuolar protein sorting-associated protein 33A OS=Homo sapiens OX=9606 GN=VPS33A PE=1 SV=1 | 0.996 | 0.893277754 | 0.961 | 0.7 |
| Q13098 | COP9 signalosome complex subunit 1 OS=Homo sapiens OX=9606 GN=GPS1 PE=1 SV=4 | 1.022 | 0.827871327 | 0.966 | 0.7 |
| Q6P996 | Pyridoxal-dependent decarboxylase domain-containing protein 1 OS=Homo sapiens OX=9606 GN=PDXDC1 PE=1 SV=2 | 1.012 | 0.962117424 | 0.972 | 0.7 |
| Q14232 | Translation initiation factor eIF-2B subunit alpha OS=Homo sapiens OX=9606 GN=EIF2B1 PE=1 SV=1 | 1.023 | 0.902016292 | 1.004 | 0.9 |
| J3KQN4 | 60S ribosomal protein L36a OS=Homo sapiens OX=9606 GN=RPL36A PE=1 SV=1 | 1.099 | 0.750023665 | 1.12 | 0.7 |
| A0A087X142 | Septin-8 OS=Homo sapiens OX=9606 GN=SEPTIN8 PE=1 SV=1 | 1.119 | 0.857755811 | 0.965 | 0.7 |
| Q92879 | CUGBP Elav-like family member 1 OS=Homo sapiens OX=9606 GN=CELF1 PE=1 SV=2 | 0.993 | 0.983953845 | 0.86 | 0.7 |
| P46459 | Vesicle-fusing ATPase OS=Homo sapiens OX=9606 GN=NSF PE=1 SV=3 | 0.965 | 0.860909277 | 0.956 | 0.7 |
| Q12965 | Unconventional myosin-Ie OS=Homo sapiens OX=9606 GN=MYO1E PE=1 SV=2 | 0.989 | 0.955325044 | 0.959 | 0.7 |
| O60343 | TBC1 domain family member 4 OS=Homo sapiens OX=9606 GN=TBC1D4 PE=1 SV=2 | 0.992 | 0.906273459 | 1.005 | 0.9 |
| G3V1U3 | Methionine aminopeptidase 2 OS=Homo sapiens OX=9606 GN=METAP2 PE=1 SV=1 | 1.022 | 0.765235202 | 1.055 | 0.7 |
| Q5RKV6 | Exosome complex component MTR3 OS=Homo sapiens OX=9606 GN=EXOSC6 PE=1 SV=1 | 1.249 | 0.765235202 | 1.004 | 0.8 |
| Q9Y2T2 | AP-3 complex subunit mu-1 OS=Homo sapiens OX=9606 GN=AP3M1 PE=1 SV=1 | 1.026 | 0.904568496 | 1.022 | 0.7 |
| Q9BYT8 | Neurolysin, mitochondrial OS=Homo sapiens OX=9606 GN=NLN PE=1 SV=1 | 0.988 | 0.902016292 | 0.96 | 0.7 |
| P21796 | Voltage-dependent anion-selective channel protein 1 OS=Homo sapiens OX=9606 GN=VDAC1 PE=1 SV=2 | 0.913 | 0.793453909 | 0.739 | 0.7 |
| Q9Y263 | Phospholipase A-2-activating protein OS=Homo sapiens OX=9606 GN=PLAA PE=1 SV=2 | 0.924 | 0.750023665 | 0.957 | 0.7 |
| O60749 | Sorting nexin-2 OS=Homo sapiens OX=9606 GN=SNX2 PE=1 SV=2 | 1.004 | 0.927756817 | 0.976 | 0.8 |
| A0A3B3ITT5 | 60S ribosomal protein L29 OS=Homo sapiens OX=9606 GN=RPL29 PE=1 SV=1 | 1.082 | 0.750023665 | 1.136 | 0.7 |
| O75381 | Peroxisomal membrane protein PEX14 OS=Homo sapiens OX=9606 GN=PEX14 PE=1 SV=1 | 0.978 | 0.898725987 | 0.939 | 0.7 |
| P61970 | Nuclear transport factor 2 OS=Homo sapiens OX=9606 GN=NUTF2 PE=1 SV=1 | 1.01 | 0.904500837 | 1.015 | 0.9 |
| E7EU96 | Casein kinase II subunit alpha OS=Homo sapiens OX=9606 GN=CSNK2A1 PE=1 SV=1 | 0.984 | 0.765235202 | 0.955 | 0.8 |
| Q9Y2Q5 | Ragulator complex protein LAMTOR2 OS=Homo sapiens OX=9606 GN=LAMTOR2 PE=1 SV=1 | 0.951 | 0.882968647 | 0.926 | 0.7 |
| Q14CX7 | N-alpha-acetyltransferase 25, NatB auxiliary subunit OS=Homo sapiens OX=9606 GN=NAA25 PE=1 SV=1 | 0.994 | 0.904500837 | 0.941 | 0.7 |
| O75718 | Cartilage-associated protein OS=Homo sapiens OX=9606 GN=CRTAP PE=1 SV=1 | 0.974 | 0.964013256 | 0.987 | 0.9 |
| P43897 | Elongation factor Ts, mitochondrial OS=Homo sapiens OX=9606 GN=TSFM PE=1 SV=2 | 1.022 | 0.962711234 | 0.886 | 0.7 |
| Q9H9H4 | Vacuolar protein sorting-associated protein 37B OS=Homo sapiens OX=9606 GN=VPS37B PE=1 SV=1 | 1.028 | 0.827871327 | 0.962 | 0.9 |
| E7EVX8 | Pre-mRNA-processing factor 31 OS=Homo sapiens OX=9606 GN=PRPF31 PE=1 SV=1 | 1.023 | 0.902016292 | 0.94 | 0.7 |
| Q9H9B4 | Sideroflexin-1 OS=Homo sapiens OX=9606 GN=SFXN1 PE=1 SV=4 | 0.941 | 0.875967645 | 0.877 | 0.7 |
| P52294 | Importin subunit alpha-5 OS=Homo sapiens OX=9606 GN=KPNA1 PE=1 SV=3 | 1.265 | 0.829390484 | 0.771 | 0.7 |
| K7EJL1 | AP-1 complex subunit mu-1 OS=Homo sapiens OX=9606 GN=AP1M1 PE=1 SV=1 | 0.949 | 0.895384791 | 0.987 | 0.7 |
| P48637 | Glutathione synthetase OS=Homo sapiens OX=9606 GN=GSS PE=1 SV=1 | 0.975 | 0.860909277 | 1 | 0.9 |
| A0A0D9SEM4 | Serine/arginine-rich-splicing factor 4 (Fragment) OS=Homo sapiens OX=9606 GN=SRSF4 PE=1 SV=1 | 0.882 | 0.882454148 | 0.901 | 0.7 |
| P61225 | Ras-related protein Rap-2b OS=Homo sapiens OX=9606 GN=RAP2B PE=1 SV=1 | 0.972 | 0.829390484 | 0.945 | 0.7 |
| Q15021 | Condensin complex subunit 1 OS=Homo sapiens OX=9606 GN=NCAPD2 PE=1 SV=3 | 1.009 | 0.902016292 | 0.955 | 0.7 |
| Q92522 | Histone H1.10 OS=Homo sapiens OX=9606 GN=H1-10 PE=1 SV=1 | 1.015 | 0.986048646 | 0.85 | 0.7 |
| O75396 | Vesicle-trafficking protein SEC22b OS=Homo sapiens OX=9606 GN=SEC22B PE=1 SV=4 | 1.106 | 0.765235202 | 1.02 | 0.9 |
| O43252 | Bifunctional 3'-phosphoadenosine 5'-phosphosulfate synthase 1 OS=Homo sapiens OX=9606 GN=PAPSS1 PE=1 SV=2 | 0.986 | 0.996637279 | 1.07 | 0.7 |
| A6NP24 | Quinone oxidoreductase (Fragment) OS=Homo sapiens OX=9606 GN=CRYZ PE=1 SV=1 | 0.992 | 0.935556205 | 0.974 | 0.9 |
| P67870 | Casein kinase II subunit beta OS=Homo sapiens OX=9606 GN=CSNK2B PE=1 SV=1 | 0.998 | 0.985324977 | 1.029 | 0.7 |
| Q9NQ88 | Fructose-2,6-bisphosphatase TIGAR OS=Homo sapiens OX=9606 GN=TIGAR PE=1 SV=1 | 0.955 | 0.765235202 | 0.934 | 0.7 |
| P29590 | Protein PML OS=Homo sapiens OX=9606 GN=PML PE=1 SV=3 | 1.015 | 0.963059765 | 0.961 | 0.7 |
| P41240 | Tyrosine-protein kinase CSK OS=Homo sapiens OX=9606 GN=CSK PE=1 SV=1 | 0.982 | 0.991801854 | 0.996 | 0.7 |
| Q15424 | Scaffold attachment factor B1 OS=Homo sapiens OX=9606 GN=SAFB PE=1 SV=4 | 0.928 | 0.750023665 | 0.981 | 0.8 |
| Q15738 | Sterol-4-alpha-carboxylate 3-dehydrogenase, decarboxylating OS=Homo sapiens OX=9606 GN=NSDHL PE=1 SV=2 | 1.017 | 0.956104752 | 1.016 | 0.9 |
| Q8TF05 | Serine/threonine-protein phosphatase 4 regulatory subunit 1 OS=Homo sapiens OX=9606 GN=PPP4R1 PE=1 SV=1 | 0.997 | 0.886561233 | 0.933 | 0.7 |
| A0A2R8Y5P7 | Histidine--tRNA ligase OS=Homo sapiens OX=9606 GN=HARS2 PE=1 SV=1 | 1.085 | 0.765235202 | 0.79 | 0.7 |
| Q96F85 | CB1 cannabinoid receptor-interacting protein 1 OS=Homo sapiens OX=9606 GN=CNRIP1 PE=1 SV=1 | 1.012 | 0.902016292 | 0.959 | 0.7 |
| Q96EP5 | DAZ-associated protein 1 OS=Homo sapiens OX=9606 GN=DAZAP1 PE=1 SV=1 | 1.071 | 0.785314375 | 0.941 | 0.7 |
| Q9Y613 | FH1/FH2 domain-containing protein 1 OS=Homo sapiens OX=9606 GN=FHOD1 PE=1 SV=3 | 1.012 | 0.985324977 | 0.927 | 0.7 |
| Q9Y3Z3 | Deoxynucleoside triphosphate triphosphohydrolase SAMHD1 OS=Homo sapiens OX=9606 GN=SAMHD1 PE=1 SV=2 | 0.999 | 0.996637279 | 0.985 | 0.8 |
| P37108 | Signal recognition particle 14 kDa protein OS=Homo sapiens OX=9606 GN=SRP14 PE=1 SV=2 | 1.01 | 0.984805107 | 0.939 | 0.7 |
| Q9UBQ7 | Glyoxylate reductase/hydroxypyruvate reductase OS=Homo sapiens OX=9606 GN=GRHPR PE=1 SV=1 | 0.939 | 0.765235202 | 0.935 | 0.7 |
| Q13501 | Sequestosome-1 OS=Homo sapiens OX=9606 GN=SQSTM1 PE=1 SV=1 | 1.01 | 0.885528205 | 0.939 | 0.7 |
| B1AJY7 | 26S proteasome non-ATPase regulatory subunit 10 OS=Homo sapiens OX=9606 GN=PSMD10 PE=1 SV=1 | 0.988 | 0.95980123 | 0.971 | 0.7 |
| P67775 | Serine/threonine-protein phosphatase 2A catalytic subunit alpha isoform OS=Homo sapiens OX=9606 GN=PPP2CA PE=1 SV=1 | 1.003 | 0.989102644 | 1.033 | 0.7 |
| P35637 | RNA-binding protein FUS OS=Homo sapiens OX=9606 GN=FUS PE=1 SV=1 | 1.12 | 0.765235202 | 0.995 | 0.9 |
| Q9UNX4 | WD repeat-containing protein 3 OS=Homo sapiens OX=9606 GN=WDR3 PE=1 SV=1 | 1.003 | 0.985330338 | 0.895 | 0.7 |
| K7ER00 | Phenylalanine--tRNA ligase OS=Homo sapiens OX=9606 GN=FARSA PE=1 SV=1 | 1.01 | 0.968024925 | 0.971 | 0.7 |
| Q9BY44 | Eukaryotic translation initiation factor 2A OS=Homo sapiens OX=9606 GN=EIF2A PE=1 SV=3 | 1.051 | 0.765235202 | 1.03 | 0.7 |
| H0YIC9 | 5'-AMP-activated protein kinase subunit gamma-1 (Fragment) OS=Homo sapiens OX=9606 GN=PRKAG1 PE=1 SV=1 | 1.052 | 0.952913364 | 0.871 | 0.7 |
| Q92882 | Osteoclast-stimulating factor 1 OS=Homo sapiens OX=9606 GN=OSTF1 PE=1 SV=2 | 0.934 | 0.642276967 | 0.941 | 0.7 |
| A0A088AWN3 | Dedicator of cytokinesis protein 9 OS=Homo sapiens OX=9606 GN=DOCK9 PE=1 SV=2 | 0.959 | 0.834564551 | 0.85 | 0.7 |
| Q16186 | Proteasomal ubiquitin receptor ADRM1 OS=Homo sapiens OX=9606 GN=ADRM1 PE=1 SV=2 | 0.991 | 0.997835552 | 1.01 | 0.7 |
| Q00688 | Peptidyl-prolyl cis-trans isomerase FKBP3 OS=Homo sapiens OX=9606 GN=FKBP3 PE=1 SV=1 | 1.011 | 0.882454148 | 1.012 | 0.9 |
| Q9H993 | Damage-control phosphatase ARMT1 OS=Homo sapiens OX=9606 GN=ARMT1 PE=1 SV=1 | 0.995 | 0.965475754 | 0.924 | 0.7 |
| I3L0A0 | HCG2044781 OS=Homo sapiens OX=9606 GN=TMEM189-UBE2V1 PE=3 SV=1 | 0.992 | 0.904500837 | 0.983 | 0.7 |
| P36959 | GMP reductase 1 OS=Homo sapiens OX=9606 GN=GMPR PE=1 SV=1 | 0.974 | 0.999880031 | 0.996 | 0.8 |
| O95347 | Structural maintenance of chromosomes protein 2 OS=Homo sapiens OX=9606 GN=SMC2 PE=1 SV=2 | 1.034 | 0.948891344 | 1.022 | 0.7 |
| Q15555 | Microtubule-associated protein RP/EB family member 2 OS=Homo sapiens OX=9606 GN=MAPRE2 PE=1 SV=1 | 1.053 | 0.750023665 | 1.017 | 0.7 |
| P08559 | Pyruvate dehydrogenase E1 component subunit alpha, somatic form, mitochondrial OS=Homo sapiens OX=9606 GN=PDHA1 PE=1 | 1.009 | 0.902016292 | 0.942 | 0.7 |
| Q9UHY7 | Enolase-phosphatase E1 OS=Homo sapiens OX=9606 GN=ENOPH1 PE=1 SV=1 | 1.01 | 0.889450577 | 0.913 | 0.7 |
| O00116 | Alkyldihydroxyacetonephosphate synthase, peroxisomal OS=Homo sapiens OX=9606 GN=AGPS PE=1 SV=1 | 1.109 | 0.765235202 | 0.936 | 0.8 |
| Q9UII2 | ATPase inhibitor, mitochondrial OS=Homo sapiens OX=9606 GN=ATP5IF1 PE=1 SV=1 | 0.947 | 0.775827921 | 0.963 | 0.7 |
| Q96JB5 | CDK5 regulatory subunit-associated protein 3 OS=Homo sapiens OX=9606 GN=CDK5RAP3 PE=1 SV=2 | 1.014 | 0.985324977 | 0.986 | 0.7 |
| P09601 | Heme oxygenase 1 OS=Homo sapiens OX=9606 GN=HMOX1 PE=1 SV=1 | 1.088 | 0.750023665 | 0.927 | 0.7 |
| A0A2U3TZL5 | CD59 glycoprotein (Fragment) OS=Homo sapiens OX=9606 GN=CD59 PE=1 SV=1 | 0.896 | 0.754126086 | 0.943 | 0.7 |
| A0A1W2PQV2 | Glycine cleavage system H protein, mitochondrial (Fragment) OS=Homo sapiens OX=9606 GN=GCSH PE=1 SV=1 | 1.104 | 0.873607677 | 0.911 | 0.7 |
| P09110 | 3-ketoacyl-CoA thiolase, peroxisomal OS=Homo sapiens OX=9606 GN=ACAA1 PE=1 SV=2 | 0.922 | 0.827871327 | 0.96 | 0.7 |
| O95479 | GDH/6PGL endoplasmic bifunctional protein OS=Homo sapiens OX=9606 GN=H6PD PE=1 SV=2 | 1.03 | 0.750023665 | 1.009 | 0.9 |
| Q86T13 | C-type lectin domain family 14 member A OS=Homo sapiens OX=9606 GN=CLEC14A PE=1 SV=1 | 0.928 | 0.765235202 | 0.894 | 0.7 |
| Q05209 | Tyrosine-protein phosphatase non-receptor type 12 OS=Homo sapiens OX=9606 GN=PTPN12 PE=1 SV=3 | 0.974 | 0.900915273 | 0.915 | 0.7 |
| P14174 | Macrophage migration inhibitory factor OS=Homo sapiens OX=9606 GN=MIF PE=1 SV=4 | 0.909 | 0.765235202 | 0.962 | 0.7 |
| H3BRG4 | Cytochrome b-c1 complex subunit 2, mitochondrial OS=Homo sapiens OX=9606 GN=UQCRC2 PE=1 SV=1 | 0.99 | 0.900868056 | 0.997 | 0.8 |
| H0Y7H7 | Dedicator of cytokinesis protein 4 (Fragment) OS=Homo sapiens OX=9606 GN=DOCK4 PE=1 SV=1 | 1.114 | 0.765235202 | 0.911 | 0.7 |
| O00499 | Myc box-dependent-interacting protein 1 OS=Homo sapiens OX=9606 GN=BIN1 PE=1 SV=1 | 0.998 | 0.942678074 | 0.925 | 0.7 |
| P58546 | Myotrophin OS=Homo sapiens OX=9606 GN=MTPN PE=1 SV=2 | 0.974 | 0.944317006 | 0.977 | 0.8 |
| P35754 | Glutaredoxin-1 OS=Homo sapiens OX=9606 GN=GLRX PE=1 SV=2 | 0.944 | 0.829390484 | 0.98 | 0.7 |
| G5E948 | Mitogen-activated protein kinase kinase kinase kinase 4 OS=Homo sapiens OX=9606 GN=MAP4K4 PE=1 SV=2 | 1.031 | 0.761475165 | 0.955 | 0.7 |
| Q9Y333 | U6 snRNA-associated Sm-like protein LSm2 OS=Homo sapiens OX=9606 GN=LSM2 PE=1 SV=1 | 0.997 | 0.949934729 | 0.928 | 0.7 |

| | | | | | |
|---|---|---|---|---|---|
| P04080 | Cystatin-B OS=Homo sapiens OX=9606 GN=CSTB PE=1 SV=2 | 0.916 | 0.837254987 | 1 | 0.9 |
| Q16775 | Hydroxyacylglutathione hydrolase, mitochondrial OS=Homo sapiens OX=9606 GN=HAGH PE=1 SV=2 | 0.926 | 0.765235202 | 0.92 | 0.7 |
| Q9BRA2 | Thioredoxin domain-containing protein 17 OS=Homo sapiens OX=9606 GN=TXNDC17 PE=1 SV=1 | 0.974 | 0.80706255 | 1.012 | 0.7 |
| Q7Z4H8 | Protein O-glucosyltransferase 3 OS=Homo sapiens OX=9606 GN=POGLUT3 PE=1 SV=2 | 0.975 | 0.765235202 | 0.994 | 0.7 |
| P14735 | Insulin-degrading enzyme OS=Homo sapiens OX=9606 GN=IDE PE=1 SV=4 | 0.974 | 0.750023665 | 1.026 | 0.8 |
| Q9Y5K5 | Ubiquitin carboxyl-terminal hydrolase isozyme L5 OS=Homo sapiens OX=9606 GN=UCHL5 PE=1 SV=3 | 0.987 | 0.80706255 | 0.969 | 0.7 |
| P53597 | Succinate--CoA ligase [ADP/GDP-forming] subunit alpha, mitochondrial OS=Homo sapiens OX=9606 GN=SUCLG1 PE=1 SV=4 | 1.041 | 0.861425366 | 0.92 | 0.7 |
| Q96DB5 | Regulator of microtubule dynamics protein 1 OS=Homo sapiens OX=9606 GN=RMDN1 PE=1 SV=1 | 0.996 | 0.972586641 | 0.94 | 0.7 |
| H0YHS6 | Tyrosine--tRNA ligase (Fragment) OS=Homo sapiens OX=9606 GN=YARS2 PE=1 SV=1 | 0.93 | 0.750023665 | 0.902 | 0.7 |
| Q9BVG4 | Protein PBDC1 OS=Homo sapiens OX=9606 GN=PBDC1 PE=1 SV=1 | 1.001 | 0.904500837 | 1.031 | 0.7 |
| P53985 | Monocarboxylate transporter 1 OS=Homo sapiens OX=9606 GN=SLC16A1 PE=1 SV=3 | 1 | 0.983074461 | 0.928 | 0.8 |
| Q8WWN8 | Arf-GAP with Rho-GAP domain, ANK repeat and PH domain-containing protein 3 OS=Homo sapiens OX=9606 GN=ARAP3 PE=1 SV= | 1.047 | 0.765235202 | 1.048 | 0.9 |
| C9J1V9 | EEF1E1-BLOC1S5 readthrough (NMD candidate) OS=Homo sapiens OX=9606 GN=EEF1E1-BLOC1S5 PE=4 SV=2 | 1.009 | 0.902016292 | 0.996 | 0.8 |
| A0A087WZF1 | Lipoma-preferred partner OS=Homo sapiens OX=9606 GN=LPP PE=1 SV=1 | 1.048 | 0.750023665 | 0.951 | 0.7 |
| O95716 | Ras-related protein Rab-3D OS=Homo sapiens OX=9606 GN=RAB3D PE=1 SV=1 | 0.93 | 0.750023665 | 0.909 | 0.7 |
| Q9UBB6 | Neurochondrin OS=Homo sapiens OX=9606 GN=NCDN PE=1 SV=1 | 0.981 | 0.916363276 | 0.913 | 0.7 |
| Q02818 | Nucleobindin-1 OS=Homo sapiens OX=9606 GN=NUCB1 PE=1 SV=4 | 1 | 0.987438982 | 0.951 | 0.7 |
| P32456 | Guanylate-binding protein 2 OS=Homo sapiens OX=9606 GN=GBP2 PE=1 SV=3 | 1.067 | 0.779470244 | 1.121 | 0.7 |
| P37235 | Hippocalcin-like protein 1 OS=Homo sapiens OX=9606 GN=HPCAL1 PE=1 SV=3 | 0.952 | 0.932272096 | 1.006 | 0.9 |
| Q14318 | Peptidyl-prolyl cis-trans isomerase FKBP8 OS=Homo sapiens OX=9606 GN=FKBP8 PE=1 SV=2 | 0.981 | 0.879425665 | 1.056 | 0.7 |
| P49736 | DNA replication licensing factor MCM2 OS=Homo sapiens OX=9606 GN=MCM2 PE=1 SV=4 | 0.996 | 0.917008272 | 1.024 | 0.7 |
| Q96DG6 | Carboxymethylenebutenolidase homolog OS=Homo sapiens OX=9606 GN=CMBL PE=1 SV=1 | 0.967 | 0.905369216 | 0.954 | 0.7 |
| E7ET15 | U2 snRNP-associated SURP motif-containing protein OS=Homo sapiens OX=9606 GN=U2SURP PE=1 SV=1 | 1.041 | 0.840133488 | 0.946 | 0.7 |
| Q8NHV1 | GTPase IMAP family member 7 OS=Homo sapiens OX=9606 GN=GIMAP7 PE=1 SV=1 | 1.038 | 0.984200071 | 0.983 | 0.7 |
| P57740 | Nuclear pore complex protein Nup107 OS=Homo sapiens OX=9606 GN=NUP107 PE=1 SV=1 | 1.133 | 0.702711636 | 0.91 | 0.9 |
| Q86X83 | COMM domain-containing protein 2 OS=Homo sapiens OX=9606 GN=COMMD2 PE=1 SV=2 | 1.157 | 0.750023665 | 0.996 | 0.8 |
| P31930 | Cytochrome b-c1 complex subunit 1, mitochondrial OS=Homo sapiens OX=9606 GN=UQCRC1 PE=1 SV=3 | 0.96 | 0.805488634 | 0.981 | 0.8 |
| A0A0A0MTH3 | Integrin-linked protein kinase OS=Homo sapiens OX=9606 GN=ILK PE=1 SV=1 | 1.008 | 0.845656075 | 0.974 | 0.7 |
| Q8TC12 | Retinol dehydrogenase 11 OS=Homo sapiens OX=9606 GN=RDH11 PE=1 SV=2 | 1.026 | 0.78623955 | 0.986 | 0.7 |
| P50452 | Serpin B8 OS=Homo sapiens OX=9606 GN=SERPINB8 PE=1 SV=2 | 0.996 | 0.879425665 | 0.904 | 0.7 |
| Q05397 | Focal adhesion kinase 1 OS=Homo sapiens OX=9606 GN=PTK2 PE=1 SV=2 | 1.1 | 0.849526028 | 0.937 | 0.7 |
| Q9P2A4 | ABI gene family member 3 OS=Homo sapiens OX=9606 GN=ABI3 PE=1 SV=2 | 0.882 | 0.876510992 | 0.93 | 0.7 |
| Q08AM6 | Protein VAC14 homolog OS=Homo sapiens OX=9606 GN=VAC14 PE=1 SV=1 | 1.043 | 0.765235202 | 0.912 | 0.7 |
| Q16795 | NADH dehydrogenase [ubiquinone] 1 alpha subcomplex subunit 9, mitochondrial OS=Homo sapiens OX=9606 GN=NDUFA9 PE=1 | 0.916 | 0.765235202 | 0.948 | 0.8 |
| Q96AP7 | Endothelial cell-selective adhesion molecule OS=Homo sapiens OX=9606 GN=ESAM PE=1 SV=1 | 0.983 | 0.866054955 | 0.89 | 0.7 |
| Q5GLZ8 | Probable E3 ubiquitin-protein ligase HERC4 OS=Homo sapiens OX=9606 GN=HERC4 PE=1 SV=1 | 0.975 | 0.829390484 | 0.913 | 0.7 |
| J3QKW7 | Developmentally-regulated GTP-binding protein 2 OS=Homo sapiens OX=9606 GN=DRG2 PE=1 SV=1 | 0.999 | 0.985324977 | 0.951 | 0.7 |
| P35270 | Sepiapterin reductase OS=Homo sapiens OX=9606 GN=SPR PE=1 SV=1 | 0.988 | 0.902016292 | 0.959 | 0.7 |
| P30049 | ATP synthase subunit delta, mitochondrial OS=Homo sapiens OX=9606 GN=ATP5F1D PE=1 SV=2 | 1.012 | 0.99634713 | 0.882 | 0.7 |
| A0A2R8Y6Y7 | Succinate--CoA ligase [ADP-forming] subunit beta, mitochondrial OS=Homo sapiens OX=9606 GN=SUCLA2 PE=1 SV=1 | 0.976 | 0.978464867 | 0.924 | 0.7 |
| O75822 | Eukaryotic translation initiation factor 3 subunit J OS=Homo sapiens OX=9606 GN=EIF3J PE=1 SV=2 | 1.048 | 0.829390484 | 0.988 | 0.9 |
| P50213 | Isocitrate dehydrogenase [NAD] subunit alpha, mitochondrial OS=Homo sapiens OX=9606 GN=IDH3A PE=1 SV=1 | 0.953 | 0.766667967 | 0.961 | 0.7 |
| Q8TCS8 | Polyribonucleotide nucleotidyltransferase 1, mitochondrial OS=Homo sapiens OX=9606 GN=PNPT1 PE=1 SV=2 | 1.053 | 0.750023665 | 0.935 | 0.7 |
| P56537 | Eukaryotic translation initiation factor 6 OS=Homo sapiens OX=9606 GN=EIF6 PE=1 SV=1 | 0.978 | 0.845656075 | 1.086 | 0.7 |
| Q9UKK3 | Protein mono-ADP-ribosyltransferase PARP4 OS=Homo sapiens OX=9606 GN=PARP4 PE=1 SV=3 | 0.972 | 0.906273459 | 1.13 | 0.7 |
| P19525 | Interferon-induced, double-stranded RNA-activated protein kinase OS=Homo sapiens OX=9606 GN=EIF2AK2 PE=1 SV=2 | 1.018 | 0.766667967 | 0.968 | 0.8 |
| P26583 | High mobility group protein B2 OS=Homo sapiens OX=9606 GN=HMGB2 PE=1 SV=2 | 1.077 | 0.765937763 | 0.961 | 0.8 |
| Q9UBB4 | Ataxin-10 OS=Homo sapiens OX=9606 GN=ATXN10 PE=1 SV=1 | 0.984 | 0.876510992 | 1.11 | 0.7 |
| P38117 | Electron transfer flavoprotein subunit beta OS=Homo sapiens OX=9606 GN=ETFB PE=1 SV=3 | 1.033 | 0.939024443 | 0.911 | 0.7 |
| Q9UI15 | Transgelin-3 OS=Homo sapiens OX=9606 GN=TAGLN3 PE=1 SV=2 | 1.133 | 0.765235202 | 1.075 | 0.7 |
| P78318 | Immunoglobulin-binding protein 1 OS=Homo sapiens OX=9606 GN=IGBP1 PE=1 SV=1 | 1.164 | 0.889450577 | 0.914 | 0.7 |
| Q14669 | E3 ubiquitin-protein ligase TRIP12 OS=Homo sapiens OX=9606 GN=TRIP12 PE=1 SV=1 | 0.996 | 0.765235202 | 0.977 | 0.9 |
| Q53FA7 | Quinone oxidoreductase PIG3 OS=Homo sapiens OX=9606 GN=TP53I3 PE=1 SV=2 | 1.026 | 0.889482529 | 0.955 | 0.8 |
| P15586 | N-acetylglucosamine-6-sulfatase OS=Homo sapiens OX=9606 GN=GNS PE=1 SV=3 | 0.926 | 0.766667967 | 0.887 | 0.7 |
| P55327 | Tumor protein D52 OS=Homo sapiens OX=9606 GN=TPD52 PE=1 SV=2 | 0.978 | 0.889450577 | 0.921 | 0.7 |
| Q969Q0 | 60S ribosomal protein L36a-like OS=Homo sapiens OX=9606 GN=RPL36AL PE=1 SV=3 | 1.032 | 0.996578046 | 1.082 | 0.7 |
| Q9UNW9 | RNA-binding protein Nova-2 OS=Homo sapiens OX=9606 GN=NOVA2 PE=1 SV=1 | 0.95 | 0.917008272 | 0.938 | 0.7 |
| O00401 | Neural Wiskott-Aldrich syndrome protein OS=Homo sapiens OX=9606 GN=WASL PE=1 SV=2 | 1.008 | 0.80706255 | 0.972 | 0.7 |
| Q15785 | Mitochondrial import receptor subunit TOM34 OS=Homo sapiens OX=9606 GN=TOMM34 PE=1 SV=2 | 1.02 | 0.80706255 | 1.005 | 0.8 |
| A0A087X2B5 | Basigin (Fragment) OS=Homo sapiens OX=9606 GN=BSG PE=1 SV=1 | 0.997 | 0.938907165 | 0.946 | 0.8 |
| Q14008 | Cytoskeleton-associated protein 5 OS=Homo sapiens OX=9606 GN=CKAP5 PE=1 SV=3 | 1.085 | 0.765235202 | 0.842 | 0.7 |
| Q96C90 | Protein phosphatase 1 regulatory subunit 14B OS=Homo sapiens OX=9606 GN=PPP1R14B PE=1 SV=3 | 1.005 | 0.983953845 | 0.942 | 0.7 |
| F5GYF7 | COP9 signalosome complex subunit 7a (Fragment) OS=Homo sapiens OX=9606 GN=COPS7A PE=1 SV=8 | 0.979 | 0.905868507 | 0.973 | 0.7 |
| A0A0C4DFL7 | Lanosterol 14-alpha demethylase OS=Homo sapiens OX=9606 GN=CYP51A1 PE=1 SV=1 | 1.018 | 0.765235202 | 1.003 | 0.7 |
| Q9H4A5 | Golgi phosphoprotein 3-like OS=Homo sapiens OX=9606 GN=GOLPH3L PE=1 SV=1 | 1.101 | 0.765235202 | 0.964 | 0.8 |
| Q96KG9 | N-terminal kinase-like protein OS=Homo sapiens OX=9606 GN=SCYL1 PE=1 SV=1 | 0.983 | 0.956825234 | 0.954 | 0.7 |
| Q8WWP7 | GTPase IMAP family member 1 OS=Homo sapiens OX=9606 GN=GIMAP1 PE=1 SV=1 | 1.03 | 0.901934028 | 0.917 | 0.7 |
| C9JVE2 | DCN1-like protein OS=Homo sapiens OX=9606 GN=DCUN1D1 PE=1 SV=1 | 0.945 | 0.765235202 | 0.973 | 0.7 |
| Q9Y2D5 | A-kinase anchor protein 2 OS=Homo sapiens OX=9606 GN=AKAP2 PE=1 SV=3 | 1.083 | 0.750023665 | 0.982 | 0.9 |
| O75340 | Programmed cell death protein 6 OS=Homo sapiens OX=9606 GN=PDCD6 PE=1 SV=1 | 0.963 | 0.765235202 | 0.976 | 0.7 |
| O60568 | Multifunctional procollagen lysine hydroxylase and glycosyltransferase LH3 OS=Homo sapiens OX=9606 GN=PLOD3 PE=1 SV=1 | 0.972 | 0.834564551 | 1.035 | 0.7 |
| Q9Y383 | Putative RNA-binding protein Luc7-like 2 OS=Homo sapiens OX=9606 GN=LUC7L2 PE=1 SV=2 | 1.032 | 0.829390484 | 1.05 | 0.7 |
| P35573 | Glycogen debranching enzyme OS=Homo sapiens OX=9606 GN=AGL PE=1 SV=3 | 1.054 | 0.750023665 | 0.956 | 0.7 |
| P42765 | 3-ketoacyl-CoA thiolase, mitochondrial OS=Homo sapiens OX=9606 GN=ACAA2 PE=1 SV=2 | 0.962 | 0.851637841 | 0.969 | 0.7 |
| Q13442 | 28 kDa heat- and acid-stable phosphoprotein OS=Homo sapiens OX=9606 GN=PDAP1 PE=1 SV=1 | 1.091 | 0.750023665 | 1.054 | 0.7 |
| Q6DN03 | Putative histone H2B type 2-C OS=Homo sapiens OX=9606 GN=H2BC20P PE=5 SV=3 | 1.027 | 0.984805107 | 0.892 | 0.7 |
| P11172 | Uridine 5'-monophosphate synthase OS=Homo sapiens OX=9606 GN=UMPS PE=1 SV=1 | 0.994 | 0.997835552 | 0.957 | 0.9 |
| Q66K14 | TBC1 domain family member 9B OS=Homo sapiens OX=9606 GN=TBC1D9B PE=1 SV=3 | 1.075 | 0.765235202 | 0.978 | 0.7 |
| P49756 | RNA-binding protein 25 OS=Homo sapiens OX=9606 GN=RBM25 PE=1 SV=3 | 1.005 | 0.925091317 | 0.965 | 0.7 |
| Q16537 | Serine/threonine-protein phosphatase 2A 56 kDa regulatory subunit epsilon isoform OS=Homo sapiens OX=9606 GN=PPP2R5E PE | 0.988 | 0.983953845 | 0.92 | 0.7 |
| O43488 | Aflatoxin B1 aldehyde reductase member 2 OS=Homo sapiens OX=9606 GN=AKR7A2 PE=1 SV=3 | 1.003 | 0.917008272 | 0.956 | 0.7 |
| P17050 | Alpha-N-acetylgalactosaminidase OS=Homo sapiens OX=9606 GN=NAGA PE=1 SV=2 | 1.035 | 0.994585949 | 0.923 | 0.7 |
| Q6ICL3 | Transport and Golgi organization protein 2 homolog OS=Homo sapiens OX=9606 GN=TANGO2 PE=1 SV=1 | 0.949 | 0.905396375 | 0.927 | 0.7 |
| P23258 | Tubulin gamma-1 chain OS=Homo sapiens OX=9606 GN=TUBG1 PE=1 SV=2 | 1.174 | 0.750023665 | 0.98 | 0.7 |
| H0YDD4 | Acetyltransferase component of pyruvate dehydrogenase complex (Fragment) OS=Homo sapiens OX=9606 GN=DLAT PE=1 SV=1 | 1.064 | 0.824551728 | 0.875 | 0.7 |
| P49458 | Signal recognition particle 9 kDa protein OS=Homo sapiens OX=9606 GN=SRP9 PE=1 SV=2 | 1.08 | 0.854598401 | 0.967 | 0.7 |
| Q8N0X7 | Spartin OS=Homo sapiens OX=9606 GN=SPART PE=1 SV=1 | 1.045 | 0.812755903 | 1.066 | 0.8 |
| P51153 | Ras-related protein Rab-13 OS=Homo sapiens OX=9606 GN=RAB13 PE=1 SV=1 | 1.077 | 0.785314375 | 1.071 | 0.9 |
| O15400 | Syntaxin-7 OS=Homo sapiens OX=9606 GN=STX7 PE=1 SV=4 | 0.99 | 0.883217803 | 0.973 | 0.7 |
| Q96HY6 | DDRGK domain-containing protein 1 OS=Homo sapiens OX=9606 GN=DDRGK1 PE=1 SV=2 | 1.019 | 0.983953845 | 1.03 | 0.9 |
| D6RBW1 | Eukaryotic translation initiation factor 4E OS=Homo sapiens OX=9606 GN=EIF4E PE=1 SV=1 | 1.055 | 0.785314375 | 0.94 | 0.7 |
| Q8NEZ5 | F-box only protein 22 OS=Homo sapiens OX=9606 GN=FBXO22 PE=1 SV=1 | 0.885 | 0.750023665 | 0.914 | 0.7 |
| P62308 | Small nuclear ribonucleoprotein G OS=Homo sapiens OX=9606 GN=SNRPG PE=1 SV=1 | 1.026 | 0.886561233 | 1.003 | 0.8 |
| A0A1B0GTK8 | Tumor necrosis factor alpha-induced protein 8-like protein 3 OS=Homo sapiens OX=9606 GN=TNFAIP8L3 PE=1 SV=1 | 1.025 | 0.835336742 | 1.1 | 0.7 |
| E9PB90 | Hexokinase OS=Homo sapiens OX=9606 GN=HK2 PE=1 SV=1 | 0.997 | 0.930855729 | 0.965 | 0.9 |
| A0A494C0X0 | Tubulin-folding cofactor B OS=Homo sapiens OX=9606 GN=TBCB PE=1 SV=1 | 0.952 | 0.77467523 | 1.047 | 0.7 |

| | | | | | |
|---|---|---|---|---|---|
| P42766 | 60S ribosomal protein L35 OS=Homo sapiens OX=9606 GN=RPL35 PE=1 SV=2 | 1.081 | 0.765235202 | 1.109 | 0.7 |
| Q9UQ35 | Serine/arginine repetitive matrix protein 2 OS=Homo sapiens OX=9606 GN=SRRM2 PE=1 SV=2 | 1.039 | 0.954224541 | 0.975 | 0.8 |
| Q8IXM3 | 39S ribosomal protein L41, mitochondrial OS=Homo sapiens OX=9606 GN=MRPL41 PE=1 SV=1 | 0.993 | 0.902777383 | 0.907 | 0.7 |
| P46776 | 60S ribosomal protein L27a OS=Homo sapiens OX=9606 GN=RPL27A PE=1 SV=2 | 0.954 | 0.985324977 | 1.131 | 0.9 |
| Q13158 | FAS-associated death domain protein OS=Homo sapiens OX=9606 GN=FADD PE=1 SV=1 | 1.048 | 0.902016292 | 0.937 | 0.7 |
| O95671 | Probable bifunctional dTTP/UTP pyrophosphatase/methyltransferase protein OS=Homo sapiens OX=9606 GN=ASMTL PE=1 SV=3 | 0.944 | 0.765235202 | 1.018 | 0.9 |
| Q9NQT5 | Exosome complex component RRP40 OS=Homo sapiens OX=9606 GN=EXOSC3 PE=1 SV=3 | 1.088 | 0.765235202 | 0.943 | 0.7 |
| Q96CN7 | Isochorismatase domain-containing protein 1 OS=Homo sapiens OX=9606 GN=ISOC1 PE=1 SV=3 | 0.993 | 0.837893552 | 0.933 | 0.7 |
| C9JXA7 | Late endosomal/lysosomal adaptor and MAPK and MTOR activator 4 OS=Homo sapiens OX=9606 GN=LAMTOR4 PE=1 SV=1 | 16.299 | 0.985330338 | 1.519 | 0.8 |
| F8W9B8 | Exocyst complex component 5 OS=Homo sapiens OX=9606 GN=EXOC5 PE=1 SV=1 | 0.956 | 0.859846322 | 0.939 | 0.7 |
| Q9NPH2 | Inositol-3-phosphate synthase 1 OS=Homo sapiens OX=9606 GN=ISYNA1 PE=1 SV=1 | 1.059 | 0.754126086 | 0.974 | 0.7 |
| G3V0E5 | Transferrin receptor protein 1 OS=Homo sapiens OX=9606 GN=TFRC PE=1 SV=1 | 0.959 | 0.765235202 | 1.038 | 0.7 |
| Q96RS6 | NudC domain-containing protein 1 OS=Homo sapiens OX=9606 GN=NUDCD1 PE=1 SV=2 | 0.986 | 0.9252392 | 0.973 | 0.9 |
| Q8IVF7 | Formin-like protein 3 OS=Homo sapiens OX=9606 GN=FMNL3 PE=1 SV=3 | 0.965 | 0.863796857 | 0.834 | 0.7 |
| Q13363 | C-terminal-binding protein 1 OS=Homo sapiens OX=9606 GN=CTBP1 PE=1 SV=2 | 0.947 | 0.765235202 | 0.953 | 0.7 |
| F5H5I6 | G-rich sequence factor 1 OS=Homo sapiens OX=9606 GN=GRSF1 PE=1 SV=2 | 1.046 | 0.780884711 | 1.016 | 0.7 |
| Q8N684 | Cleavage and polyadenylation specificity factor subunit 7 OS=Homo sapiens OX=9606 GN=CPSF7 PE=1 SV=1 | 1.085 | 0.968712192 | 0.804 | 0.7 |
| O60502 | Protein O-GlcNAcase OS=Homo sapiens OX=9606 GN=OGA PE=1 SV=2 | 1.053 | 0.750023665 | 1.01 | 0.8 |
| Q9P2R3 | Rabankyrin-5 OS=Homo sapiens OX=9606 GN=ANKFY1 PE=1 SV=2 | 0.983 | 0.809976184 | 0.947 | 0.9 |
| P62633 | Cellular nucleic acid-binding protein OS=Homo sapiens OX=9606 GN=CNBP PE=1 SV=1 | 0.952 | 0.985324977 | 1.09 | 0.8 |
| K7ERU7 | Telomerase RNA component-interacting RNase OS=Homo sapiens OX=9606 GN=TRIR PE=1 SV=1 | 1.104 | 0.882454148 | 1.067 | 0.8 |
| Q8WUX9 | Charged multivesicular body protein 7 OS=Homo sapiens OX=9606 GN=CHMP7 PE=1 SV=1 | 0.962 | 0.905868507 | 0.89 | 0.7 |
| Q9NX58 | Cell growth-regulating nucleolar protein OS=Homo sapiens OX=9606 GN=LYAR PE=1 SV=2 | 1.022 | 0.827451813 | 1.026 | 0.9 |
| E9PQP6 | Protein farnesyltransferase/geranylgeranyltransferase type-1 subunit alpha (Fragment) OS=Homo sapiens OX=9606 GN=FNTA PE= | 1.009 | 0.925836304 | 0.924 | 0.7 |
| E7EMD0 | NADPH--cytochrome P450 reductase OS=Homo sapiens OX=9606 GN=POR PE=1 SV=1 | 1.016 | 0.965945681 | 0.896 | 0.7 |
| K7ERP4 | Glutathione peroxidase (Fragment) OS=Homo sapiens OX=9606 GN=GPX4 PE=1 SV=1 | 0.996 | 0.889482529 | 0.956 | 0.7 |
| P50479 | PDZ and LIM domain protein 4 OS=Homo sapiens OX=9606 GN=PDLIM4 PE=1 SV=2 | 0.994 | 0.956825234 | 0.932 | 0.7 |
| A0A499FIX8 | Extended synaptotagmin-2 OS=Homo sapiens OX=9606 GN=ESYT2 PE=1 SV=1 | 0.962 | 0.859327485 | 0.97 | 0.7 |
| Q15819 | Ubiquitin-conjugating enzyme E2 variant 2 OS=Homo sapiens OX=9606 GN=UBE2V2 PE=1 SV=4 | 0.978 | 0.810054304 | 1.009 | 0.9 |
| Q15836 | Vesicle-associated membrane protein 3 OS=Homo sapiens OX=9606 GN=VAMP3 PE=1 SV=3 | 1.024 | 0.939024443 | 0.938 | 0.7 |
| Q9GZL7 | Ribosome biogenesis protein WDR12 OS=Homo sapiens OX=9606 GN=WDR12 PE=1 SV=2 | 1.026 | 0.765235202 | 0.867 | 0.8 |
| F8WE88 | Unconventional myosin-Va OS=Homo sapiens OX=9606 GN=MYO5A PE=1 SV=2 | 0.981 | 0.918702104 | 0.914 | 0.7 |
| Q5EBL4 | RILP-like protein 1 OS=Homo sapiens OX=9606 GN=RILPL1 PE=1 SV=1 | 0.959 | 0.824101611 | 0.989 | 0.7 |
| P52594 | Arf-GAP domain and FG repeat-containing protein 1 OS=Homo sapiens OX=9606 GN=AGFG1 PE=1 SV=2 | 0.917 | 0.865970839 | 0.93 | 0.7 |
| P01116 | GTPase KRas OS=Homo sapiens OX=9606 GN=KRAS PE=1 SV=1 | 0.999 | 0.947161221 | 0.906 | 0.7 |
| P62714 | Serine/threonine-protein phosphatase 2A catalytic subunit beta isoform OS=Homo sapiens OX=9606 GN=PPP2CB PE=1 SV=1 | 0.999 | 0.989102644 | 0.994 | 0.7 |
| A0A3B3IRQ9 | Double-stranded RNA-specific adenosine deaminase (Fragment) OS=Homo sapiens OX=9606 GN=ADAR PE=1 SV=1 | 1.015 | 0.862309708 | 1.001 | 0 |
| Q14155-1 | Isoform 1 of Rho guanine nucleotide exchange factor 7 OS=Homo sapiens OX=9606 GN=ARHGEF7 | 1.01 | 0.96244577 | 0.882 | 0.7 |
| Q15642 | Cdc42-interacting protein 4 OS=Homo sapiens OX=9606 GN=TRIP10 PE=1 SV=3 | 0.972 | 0.642276967 | 0.961 | 0.7 |
| P41567 | Eukaryotic translation initiation factor 1 OS=Homo sapiens OX=9606 GN=EIF1 PE=1 SV=1 | 1.065 | 0.901934028 | 0.99 | 0.8 |
| P51648 | Aldehyde dehydrogenase family 3 member A2 OS=Homo sapiens OX=9606 GN=ALDH3A2 PE=1 SV=1 | 1.051 | 0.780884711 | 0.871 | 0.7 |
| Q92878 | DNA repair protein RAD50 OS=Homo sapiens OX=9606 GN=RAD50 PE=1 SV=1 | 1.039 | 0.827871327 | 0.943 | 0.7 |
| O94906 | Pre-mRNA-processing factor 6 OS=Homo sapiens OX=9606 GN=PRPF6 PE=1 SV=1 | 1.046 | 0.765235202 | 0.961 | 0.7 |
| P62993 | Growth factor receptor-bound protein 2 OS=Homo sapiens OX=9606 GN=GRB2 PE=1 SV=1 | 0.982 | 0.889450577 | 0.968 | 0.7 |
| Q9H9J2 | 39S ribosomal protein L44, mitochondrial OS=Homo sapiens OX=9606 GN=MRPL44 PE=1 SV=1 | 1.012 | 0.938907165 | 0.889 | 0.7 |
| Q9UN86 | Ras GTPase-activating protein-binding protein 2 OS=Homo sapiens OX=9606 GN=G3BP2 PE=1 SV=2 | 0.981 | 0.927756817 | 1.034 | 0.7 |
| Q9H3N1 | Thioredoxin-related transmembrane protein 1 OS=Homo sapiens OX=9606 GN=TMX1 PE=1 SV=1 | 0.995 | 0.978003638 | 1.047 | 0.7 |
| P14209 | CD99 antigen OS=Homo sapiens OX=9606 GN=CD99 PE=1 SV=1 | 0.865 | 0.776314646 | 1.022 | 0.7 |
| A0A087X1I3 | Flavoprotein subunit of complex II OS=Homo sapiens OX=9606 GN=SDHA PE=1 SV=1 | 1.064 | 0.766667967 | 0.885 | 0.7 |
| P53999 | Activated RNA polymerase II transcriptional coactivator p15 OS=Homo sapiens OX=9606 GN=SUB1 PE=1 SV=3 | 1.017 | 0.968293464 | 0.901 | 0.7 |
| Q9Y3D6 | Mitochondrial fission 1 protein OS=Homo sapiens OX=9606 GN=FIS1 PE=1 SV=2 | 0.988 | 0.95980123 | 0.967 | 0.7 |
| P46734 | Dual specificity mitogen-activated protein kinase kinase 3 OS=Homo sapiens OX=9606 GN=MAP2K3 PE=1 SV=2 | 1.026 | 0.809976184 | 0.916 | 0.7 |
| O95352 | Ubiquitin-like modifier-activating enzyme ATG7 OS=Homo sapiens OX=9606 GN=ATG7 PE=1 SV=1 | 0.977 | 0.765235202 | 0.945 | 0.7 |
| Q14344 | Guanine nucleotide-binding protein subunit alpha-13 OS=Homo sapiens OX=9606 GN=GNA13 PE=1 SV=2 | 1.035 | 0.837941467 | 0.916 | 0.7 |
| Q8NDH3 | Probable aminopeptidase NPEPL1 OS=Homo sapiens OX=9606 GN=NPEPL1 PE=1 SV=3 | 1.038 | 0.980556319 | 0.969 | 0.8 |
| P57737 | Coronin-7 OS=Homo sapiens OX=9606 GN=CORO7 PE=1 SV=2 | 1.017 | 0.904500837 | 0.976 | 0.7 |
| Q00325 | Phosphate carrier protein, mitochondrial OS=Homo sapiens OX=9606 GN=SLC25A3 PE=1 SV=2 | 0.909 | 0.766667967 | 0.855 | 0.7 |
| A6NHJ8 | Quinone oxidoreductase-like protein 1 OS=Homo sapiens OX=9606 GN=CRYZL1 PE=1 SV=2 | 1.001 | 0.954224541 | 0.921 | 0.7 |
| A0A140TA73 | Beta-2-syntrophin (Fragment) OS=Homo sapiens OX=9606 GN=SNTB2 PE=1 SV=1 | 0.918 | 0.933097749 | 0.922 | 0.7 |
| P55145 | Mesencephalic astrocyte-derived neurotrophic factor OS=Homo sapiens OX=9606 GN=MANF PE=1 SV=3 | 1.028 | 0.765235202 | 0.89 | 0.7 |
| P62875 | DNA-directed RNA polymerases I, II, and III subunit RPABC5 OS=Homo sapiens OX=9606 GN=POLR2L PE=1 SV=1 | 1.011 | 0.951537478 | 0.974 | 0.7 |
| O00273 | DNA fragmentation factor subunit alpha OS=Homo sapiens OX=9606 GN=DFFA PE=1 SV=1 | 0.855 | 0.765235202 | 0.92 | 0.7 |
| Q9NTZ6 | RNA-binding protein 12 OS=Homo sapiens OX=9606 GN=RBM12 PE=1 SV=1 | 0.929 | 0.750023665 | 0.968 | 0.7 |
| O94919 | Endonuclease domain-containing 1 protein OS=Homo sapiens OX=9606 GN=ENDOD1 PE=1 SV=2 | 0.976 | 0.95980123 | 0.897 | 0.7 |
| P63220 | 40S ribosomal protein S21 OS=Homo sapiens OX=9606 GN=RPS21 PE=1 SV=1 | 1.022 | 0.886561233 | 1.053 | 0.7 |
| P98179 | RNA-binding protein 3 OS=Homo sapiens OX=9606 GN=RBM3 PE=1 SV=1 | 1 | 0.955325044 | 1.053 | 0.7 |
| Q9Y6E0 | Serine/threonine-protein kinase 24 OS=Homo sapiens OX=9606 GN=STK24 PE=1 SV=1 | 0.988 | 0.809976184 | 1 | 0.8 |
| Q6PI48 | Aspartate--tRNA ligase, mitochondrial OS=Homo sapiens OX=9606 GN=DARS2 PE=1 SV=1 | 0.939 | 0.776314646 | 0.923 | 0.7 |
| P42126 | Enoyl-CoA delta isomerase 1, mitochondrial OS=Homo sapiens OX=9606 GN=ECI1 PE=1 SV=1 | 1.003 | 0.918702104 | 0.986 | 0.7 |
| Q9HB40 | Retinoid-inducible serine carboxypeptidase OS=Homo sapiens OX=9606 GN=SCPEP1 PE=1 SV=1 | 1.001 | 0.902016292 | 0.906 | 0.7 |
| Q6XZF7 | Dynamin-binding protein OS=Homo sapiens OX=9606 GN=DNMBP PE=1 SV=1 | 0.994 | 0.996637279 | 1.056 | 0.7 |
| P35658 | Nuclear pore complex protein Nup214 OS=Homo sapiens OX=9606 GN=NUP214 PE=1 SV=2 | 1.032 | 0.859846322 | 0.989 | 0.9 |
| V9GYX7 | Actin-related protein 10 (Fragment) OS=Homo sapiens OX=9606 GN=ACTR10 PE=1 SV=1 | 1.009 | 0.893277754 | 0.936 | 0.7 |
| P51398 | 28S ribosomal protein S29, mitochondrial OS=Homo sapiens OX=9606 GN=DAP3 PE=1 SV=1 | 0.98 | 0.947526471 | 1.47 | 0.7 |
| Q06124 | Tyrosine-protein phosphatase non-receptor type 11 OS=Homo sapiens OX=9606 GN=PTPN11 PE=1 SV=3 | 1.005 | 0.994361756 | 1.021 | 0.8 |
| P01889 | HLA class I histocompatibility antigen, B alpha chain OS=Homo sapiens OX=9606 GN=HLA-B PE=1 SV=3 | 0.791 | 0.765235202 | 1.387 | 0.7 |
| Q86W92 | Liprin-beta-1 OS=Homo sapiens OX=9606 GN=PPFIBP1 PE=1 SV=2 | 1.043 | 0.750023665 | 0.993 | 0.7 |
| P08621 | U1 small nuclear ribonucleoprotein 70 kDa OS=Homo sapiens OX=9606 GN=SNRNP70 PE=1 SV=2 | 1.007 | 0.977023215 | 0.986 | 0.7 |
| P46109 | Crk-like protein OS=Homo sapiens OX=9606 GN=CRKL PE=1 SV=1 | 0.985 | 0.827871327 | 0.941 | 0.7 |
| Q14192 | Four and a half LIM domains protein 2 OS=Homo sapiens OX=9606 GN=FHL2 PE=1 SV=3 | 0.966 | 0.937467594 | 0.805 | 0.7 |
| Q6IQ42 | FUSIP1 protein OS=Homo sapiens OX=9606 GN=SRSF10 PE=1 SV=1 | 1.008 | 0.984805107 | 0.944 | 0.7 |
| E7ESC6 | Exportin-7 OS=Homo sapiens OX=9606 GN=XPO7 PE=1 SV=1 | 0.966 | 0.765235202 | 0.968 | 0.7 |
| Q8N1B4 | Vacuolar protein sorting-associated protein 52 homolog OS=Homo sapiens OX=9606 GN=VPS52 PE=1 SV=1 | 1.171 | 0.889482529 | 0.934 | 0.7 |
| F5H442 | Tumor susceptibility gene 101 protein OS=Homo sapiens OX=9606 GN=TSG101 PE=1 SV=1 | 1.013 | 0.809976184 | 0.977 | 0.7 |
| O95793 | Double-stranded RNA-binding protein Staufen homolog 1 OS=Homo sapiens OX=9606 GN=STAU1 PE=1 SV=2 | 1.104 | 0.765235202 | 0.949 | 0.7 |
| J3QQV6 | Pyridoxal 5'-phosphate synthase (Fragment) OS=Homo sapiens OX=9606 GN=PNPO PE=1 SV=2 | 1.046 | 0.765235202 | 0.881 | 0.7 |
| P08579 | U2 small nuclear ribonucleoprotein B'' OS=Homo sapiens OX=9606 GN=SNRPB2 PE=1 SV=1 | 1.081 | 0.791040593 | 0.966 | 0.8 |
| O00193 | Small acidic protein OS=Homo sapiens OX=9606 GN=SMAP PE=1 SV=1 | 0.979 | 0.996637279 | 0.99 | 0.9 |
| A0A3B3ITX4 | Uncharacterized protein OS=Homo sapiens OX=9606 PE=3 SV=1 | 1.003 | 0.994361756 | 0.795 | 0.7 |
| P49459 | Ubiquitin-conjugating enzyme E2 A OS=Homo sapiens OX=9606 GN=UBE2A PE=1 SV=2 | 1.068 | 0.902016292 | 0.97 | 0.8 |
| Q9HA77 | Probable cysteine--tRNA ligase, mitochondrial OS=Homo sapiens OX=9606 GN=CARS2 PE=1 SV=1 | 1.026 | 0.983479158 | 0.934 | 0.8 |
| Q15437 | Protein transport protein Sec23B OS=Homo sapiens OX=9606 GN=SEC23B PE=1 SV=2 | 1.039 | 0.765235202 | 0.993 | 0.7 |
| F8W8I6 | Nucleolysin TIA-1 isoform p40 OS=Homo sapiens OX=9606 GN=TIA1 PE=1 SV=1 | 0.814 | 0.87017509 | 0.936 | 0.7 |
| P09486 | SPARC OS=Homo sapiens OX=9606 GN=SPARC PE=1 SV=1 | 0.916 | 0.865970839 | 0.84 | 0.7 |
| O15270 | Serine palmitoyltransferase 2 OS=Homo sapiens OX=9606 GN=SPTLC2 PE=1 SV=1 | 1.011 | 0.994361756 | 0.957 | 0.7 |
| Q86Y82 | Syntaxin-12 OS=Homo sapiens OX=9606 GN=STX12 PE=1 SV=1 | 0.921 | 0.750023665 | 0.982 | 0.8 |

| | | | | | |
|---|---|---|---|---|---|
| Q92556 | Engulfment and cell motility protein 1 OS=Homo sapiens OX=9606 GN=ELMO1 PE=1 SV=2 | 0.967 | 0.80706255 | 0.94 | 0.7 |
| Q01970 | 1-phosphatidylinositol 4,5-bisphosphate phosphodiesterase beta-3 OS=Homo sapiens OX=9606 GN=PLCB3 PE=1 SV=2 | 1.056 | 0.750023665 | 0.956 | 0.7 |
| H7C0M1 | H/ACA ribonucleoprotein complex subunit DKC1 (Fragment) OS=Homo sapiens OX=9606 GN=DKC1 PE=1 SV=1 | 0.951 | 0.889482529 | 0.925 | 0.8 |
| J3KS22 | L-xylulose reductase (Fragment) OS=Homo sapiens OX=9606 GN=DCXR PE=1 SV=8 | 0.959 | 0.963059765 | 0.935 | 0.7 |
| A0A5F9ZH07 | Tyrosine-protein kinase (Fragment) OS=Homo sapiens OX=9606 GN=JAK1 PE=1 SV=1 | 1.125 | 0.750023665 | 1.209 | 0.7 |
| Q9UHD1 | Cysteine and histidine-rich domain-containing protein 1 OS=Homo sapiens OX=9606 GN=CHORDC1 PE=1 SV=2 | 0.982 | 0.927148669 | 1.043 | 0.7 |
| P54802 | Alpha-N-acetylglucosaminidase OS=Homo sapiens OX=9606 GN=NAGLU PE=1 SV=2 | 0.948 | 0.765235202 | 0.867 | 0.7 |
| E9PFR3 | Serine/threonine-protein phosphatase 2A 56 kDa regulatory subunit OS=Homo sapiens OX=9606 GN=PPP2R5D PE=1 SV=1 | 1.006 | 0.974919191 | 0.934 | 0.7 |
| Q8WXF1 | Paraspeckle component 1 OS=Homo sapiens OX=9606 GN=PSPC1 PE=1 SV=1 | 1.076 | 0.830179531 | 0.934 | 0.9 |
| Q9BPX5 | Actin-related protein 2/3 complex subunit 5-like protein OS=Homo sapiens OX=9606 GN=ARPC5L PE=1 SV=1 | 0.998 | 0.860909277 | 1.02 | 0.7 |
| P46108 | Adapter molecule crk OS=Homo sapiens OX=9606 GN=CRK PE=1 SV=2 | 0.957 | 0.924631778 | 1.059 | 0.7 |
| J3KRC4 | 5'(3')-deoxyribonucleotidase, cytosolic type OS=Homo sapiens OX=9606 GN=NT5C PE=1 SV=1 | 0.865 | 0.803463851 | 0.931 | 0.7 |
| Q9Y5M8 | Signal recognition particle receptor subunit beta OS=Homo sapiens OX=9606 GN=SRPRB PE=1 SV=3 | 0.988 | 0.872869468 | 1.073 | 0.8 |
| Q14696 | LRP chaperone MESD OS=Homo sapiens OX=9606 GN=MESD PE=1 SV=2 | 0.98 | 0.869995963 | 0.991 | 0.7 |
| A0A0A0MT83 | Butyryl-CoA dehydrogenase OS=Homo sapiens OX=9606 GN=IVD PE=1 SV=2 | 1.003 | 0.996578046 | 1.157 | 0.7 |
| Q6EEV6 | Small ubiquitin-related modifier 4 OS=Homo sapiens OX=9606 GN=SUMO4 PE=1 SV=2 | 0.936 | 0.89188144 | 1.035 | 0.9 |
| Q9Y394 | Dehydrogenase/reductase SDR family member 7 OS=Homo sapiens OX=9606 GN=DHRS7 PE=1 SV=1 | 1.037 | 0.937888222 | 0.942 | 0.7 |
| O14929 | Histone acetyltransferase type B catalytic subunit OS=Homo sapiens OX=9606 GN=HAT1 PE=1 SV=1 | 0.986 | 0.882454148 | 1.058 | 0.7 |
| Q53GS9 | U4/U6.U5 tri-snRNP-associated protein 2 OS=Homo sapiens OX=9606 GN=USP39 PE=1 SV=2 | 0.938 | 0.99319413 | 0.981 | 0.7 |
| A0A0A0MTJ9 | Neutral cholesterol ester hydrolase 1 OS=Homo sapiens OX=9606 GN=NCEH1 PE=1 SV=1 | 0.908 | 0.750023665 | 0.927 | 0.7 |
| Q6PCE3 | Glucose 1,6-bisphosphate synthase OS=Homo sapiens OX=9606 GN=PGM2L1 PE=1 SV=3 | 1.045 | 0.872474259 | 0.957 | 0.7 |
| O75844 | CAAX prenyl protease 1 homolog OS=Homo sapiens OX=9606 GN=ZMPSTE24 PE=1 SV=2 | 1.031 | 0.829390484 | 0.907 | 0.7 |
| Q9UPU5 | Ubiquitin carboxyl-terminal hydrolase 24 OS=Homo sapiens OX=9606 GN=USP24 PE=1 SV=3 | 0.995 | 0.865970839 | 0.931 | 0.7 |
| F8VXI9 | ARF GTPase-activating protein GIT2 OS=Homo sapiens OX=9606 GN=GIT2 PE=1 SV=1 | 1.062 | 0.765235202 | 0.934 | 0.7 |
| Q14558 | Phosphoribosyl pyrophosphate synthase-associated protein 1 OS=Homo sapiens OX=9606 GN=PRPSAP1 PE=1 SV=2 | 0.941 | 0.750023665 | 0.925 | 0.7 |
| P01111 | GTPase NRas OS=Homo sapiens OX=9606 GN=NRAS PE=1 SV=1 | 0.977 | 0.996637279 | 0.971 | 0.7 |
| A0A0A0MRM9 | Nucleolar and coiled-body phosphoprotein 1 (Fragment) OS=Homo sapiens OX=9606 GN=NOLC1 PE=1 SV=1 | 0.967 | 0.766667967 | 0.934 | 0.8 |
| O60443 | Gasdermin-E OS=Homo sapiens OX=9606 GN=GSDME PE=1 SV=2 | 1.077 | 0.932716491 | 0.934 | 0.7 |
| O14907 | Tax1-binding protein 3 OS=Homo sapiens OX=9606 GN=TAX1BP3 PE=1 SV=2 | 1.018 | 0.754126086 | 0.964 | 0.7 |
| Q96EY8 | Corrinoid adenosyltransferase OS=Homo sapiens OX=9606 GN=MMAB PE=1 SV=1 | 0.963 | 0.952913364 | 0.935 | 0.7 |
| J3KNP2 | Transducin beta-like protein 3 (Fragment) OS=Homo sapiens OX=9606 GN=TBL3 PE=1 SV=1 | 0.981 | 0.840158726 | 0.932 | 0.8 |
| E7EWX8 | Monoglyceride lipase OS=Homo sapiens OX=9606 GN=MGLL PE=1 SV=1 | 0.976 | 0.900868056 | 0.971 | 0.7 |
| O60287 | Nucleolar pre-ribosomal-associated protein 1 OS=Homo sapiens OX=9606 GN=URB1 PE=1 SV=4 | 1.005 | 0.994361756 | 0.928 | 0.9 |
| E9PK91 | Bcl-2-associated transcription factor 1 OS=Homo sapiens OX=9606 GN=BCLAF1 PE=1 SV=1 | 0.987 | 0.990981579 | 1.029 | 0.9 |
| Q32MZ4 | Leucine-rich repeat flightless-interacting protein 1 OS=Homo sapiens OX=9606 GN=LRRFIP1 PE=1 SV=2 | 1.182 | 0.780884711 | 0.981 | 0.9 |
| E9PK21 | Peptidyl-prolyl cis-trans isomerase E OS=Homo sapiens OX=9606 GN=PPIE PE=1 SV=1 | 0.983 | 0.996637279 | 0.934 | 0.7 |
| C9JA28 | Signal sequence receptor subunit gamma OS=Homo sapiens OX=9606 GN=SSR3 PE=1 SV=1 | 1.069 | 0.865970839 | 1.052 | 0.8 |
| Q9NP92 | 39S ribosomal protein S30, mitochondrial OS=Homo sapiens OX=9606 GN=MRPS30 PE=1 SV=2 | 1.038 | 0.889482529 | 0.9 | 0.7 |
| Q9H008 | Phospholysine phosphohistidine inorganic pyrophosphate phosphatase OS=Homo sapiens OX=9606 GN=LHPP PE=1 SV=2 | 1.035 | 0.921087329 | 0.875 | 0.7 |
| Q9BX68 | Histidine triad nucleotide-binding protein 2, mitochondrial OS=Homo sapiens OX=9606 GN=HINT2 PE=1 SV=1 | 1.056 | 0.750023665 | 0.975 | 0.7 |
| P62314 | Small nuclear ribonucleoprotein Sm D1 OS=Homo sapiens OX=9606 GN=SNRPD1 PE=1 SV=1 | 1.015 | 0.994361756 | 0.962 | 0.7 |
| Q15631 | Translin OS=Homo sapiens OX=9606 GN=TSN PE=1 SV=1 | 0.95 | 0.765235202 | 0.995 | 0.7 |
| P49441 | Inositol polyphosphate 1-phosphatase OS=Homo sapiens OX=9606 GN=INPP1 PE=1 SV=1 | 0.981 | 0.939929386 | 0.965 | 0.7 |
| Q8IVL6 | Prolyl 3-hydroxylase 3 OS=Homo sapiens OX=9606 GN=P3H3 PE=1 SV=1 | 1.011 | 0.891161036 | 0.989 | 0.8 |
| Q15042 | Rab3 GTPase-activating protein catalytic subunit OS=Homo sapiens OX=9606 GN=RAB3GAP1 PE=1 SV=3 | 1.065 | 0.882454148 | 0.918 | 0.7 |
| Q92851-4 | Isoform D of Caspase-10 OS=Homo sapiens OX=9606 GN=CASP10 | 0.964 | 0.889450577 | 0.903 | 0.7 |
| Q15036 | Sorting nexin-17 OS=Homo sapiens OX=9606 GN=SNX17 PE=1 SV=1 | 0.914 | 0.835336742 | 0.92 | 0.7 |
| G5EA06 | 28S ribosomal protein S27, mitochondrial OS=Homo sapiens OX=9606 GN=MRPS27 PE=1 SV=1 | 1.057 | 0.827451813 | 0.897 | 0.7 |
| A0A0C4DGN6 | ARF GTPase-activating protein GIT1 OS=Homo sapiens OX=9606 GN=GIT1 PE=1 SV=1 | 0.995 | 0.97688166 | 0.933 | 0.7 |
| J3QS63 | Nuclear pore complex protein Nup85 (Fragment) OS=Homo sapiens OX=9606 GN=NUP85 PE=1 SV=1 | 1.044 | 0.766667967 | 0.813 | 0.7 |
| G3V2S9 | SRA stem-loop-interacting RNA-binding protein, mitochondrial OS=Homo sapiens OX=9606 GN=SLIRP PE=1 SV=1 | 1.001 | 0.978562349 | 0.99 | 0.9 |
| E2QRB3 | Pyrroline-5-carboxylate reductase OS=Homo sapiens OX=9606 GN=PYCR1 PE=1 SV=2 | 1.056 | 0.766667967 | 1.011 | 0.9 |
| P41226 | Ubiquitin-like modifier-activating enzyme 7 OS=Homo sapiens OX=9606 GN=UBA7 PE=1 SV=2 | 1.145 | 0.750023665 | 1.011 | 0.9 |
| Q9H0A8 | COMM domain-containing protein 4 OS=Homo sapiens OX=9606 GN=COMMD4 PE=1 SV=1 | 0.944 | 0.765235202 | 1.026 | 0.9 |
| P17096-2 | Isoform HMG-Y of High mobility group protein HMG-I/HMG-Y OS=Homo sapiens OX=9606 GN=HMGA1 | 1.473 | 0.765235202 | 1.004 | 0.9 |
| O75351 | Vacuolar protein sorting-associated protein 4B OS=Homo sapiens OX=9606 GN=VPS4B PE=1 SV=2 | 0.985 | 0.971955334 | 0.995 | 0.9 |
| Q7L523 | Ras-related GTP-binding protein A OS=Homo sapiens OX=9606 GN=RRAGA PE=1 SV=1 | 0.977 | 0.942678074 | 0.915 | 0.7 |
| Q15796 | Mothers against decapentaplegic homolog 2 OS=Homo sapiens OX=9606 GN=SMAD2 PE=1 SV=1 | 1.098 | 0.910115867 | 1.021 | 0.9 |
| P56545-2 | Isoform 2 of C-terminal-binding protein 2 OS=Homo sapiens OX=9606 GN=CTBP2 | 1.018 | 0.901934028 | 0.972 | 0.7 |
| Q96K17 | Transcription factor BTF3 homolog 4 OS=Homo sapiens OX=9606 GN=BTF3L4 PE=1 SV=1 | 1.087 | 0.750023665 | 0.928 | 0.7 |
| A0A494C0A8 | Serine/threonine-protein kinase TBK1 OS=Homo sapiens OX=9606 GN=TBK1 PE=1 SV=1 | 0.976 | 0.953792095 | 1.016 | 0.8 |
| Q86UE4 | Protein LYRIC OS=Homo sapiens OX=9606 GN=MTDH PE=1 SV=2 | 0.97 | 0.886561233 | 0.943 | 0.8 |
| O95302 | Peptidyl-prolyl cis-trans isomerase FKBP9 OS=Homo sapiens OX=9606 GN=FKBP9 PE=1 SV=2 | 1.036 | 0.986297807 | 0.922 | 0.7 |
| Q9NV70 | Exocyst complex component 1 OS=Homo sapiens OX=9606 GN=EXOC1 PE=1 SV=4 | 1.002 | 0.985324977 | 1.008 | 0.8 |
| Q9BYB0 | SH3 and multiple ankyrin repeat domains protein 3 OS=Homo sapiens OX=9606 GN=SHANK3 PE=1 SV=3 | 1.13 | 0.830179531 | 0.852 | 0.7 |
| O43149 | Zinc finger ZZ-type and EF-hand domain-containing protein 1 OS=Homo sapiens OX=9606 GN=ZZEF1 PE=1 SV=6 | 1.034 | 0.994361756 | 0.769 | 0.7 |
| Q9P1F3 | Costars family protein ABRACL OS=Homo sapiens OX=9606 GN=ABRACL PE=1 SV=1 | 1.066 | 0.750023665 | 1.09 | 0.7 |
| O94874-3 | Isoform 3 of E3 UFM1-protein ligase 1 OS=Homo sapiens OX=9606 GN=UFL1 | 0.944 | 0.865165562 | 0.953 | 0.9 |
| Q04323 | UBX domain-containing protein 1 OS=Homo sapiens OX=9606 GN=UBXN1 PE=1 SV=2 | 0.981 | 0.956825234 | 1.002 | 0.8 |
| P20962 | Parathymosin OS=Homo sapiens OX=9606 GN=PTMS PE=1 SV=2 | 0.939 | 0.925981427 | 0.957 | 0.7 |
| P14136 | Glial fibrillary acidic protein OS=Homo sapiens OX=9606 GN=GFAP PE=1 SV=1 | 1.063 | 0.680740328 | 0.964 | 0.9 |
| Q8IY67 | Ribonucleoprotein PTB-binding 1 OS=Homo sapiens OX=9606 GN=RAVER1 PE=1 SV=1 | 0.982 | 0.889482529 | 0.868 | 0.7 |
| Q3KQV9 | UDP-N-acetylhexosamine pyrophosphorylase-like protein 1 OS=Homo sapiens OX=9606 GN=UAP1L1 PE=1 SV=2 | 1.003 | 0.961445805 | 0.916 | 0.7 |
| P24666 | Low molecular weight phosphotyrosine protein phosphatase OS=Homo sapiens OX=9606 GN=ACP1 PE=1 SV=3 | 1.091 | 0.750023665 | 0.975 | 0.7 |
| Q92922 | SWI/SNF complex subunit SMARCC1 OS=Homo sapiens OX=9606 GN=SMARCC1 PE=1 SV=3 | 1.01 | 0.997276249 | 0.859 | 0.7 |
| O95372 | Acyl-protein thioesterase 2 OS=Homo sapiens OX=9606 GN=LYPLA2 PE=1 SV=1 | 0.804 | 0.750023665 | 0.958 | 0.7 |
| Q15020 | Squamous cell carcinoma antigen recognized by T-cells 3 OS=Homo sapiens OX=9606 GN=SART3 PE=1 SV=1 | 0.935 | 0.849526028 | 0.968 | 0.7 |
| Q96IZ0 | PRKC apoptosis WT1 regulator protein OS=Homo sapiens OX=9606 GN=PAWR PE=1 SV=1 | 0.923 | 0.978562349 | 0.866 | 0.7 |
| E9PDF6 | Unconventional myosin-Ib OS=Homo sapiens OX=9606 GN=MYO1B PE=1 SV=1 | 0.925 | 0.750023665 | 0.835 | 0.7 |
| O14976 | Cyclin-G-associated kinase OS=Homo sapiens OX=9606 GN=GAK PE=1 SV=2 | 0.997 | 0.95980123 | 1.004 | 0.8 |
| Q9UBI6 | Guanine nucleotide-binding protein G(I)/G(S)/G(O) subunit gamma-12 OS=Homo sapiens OX=9606 GN=GNG12 PE=1 SV=3 | 0.891 | 0.750023665 | 0.928 | 0.7 |
| O75348 | V-type proton ATPase subunit G 1 OS=Homo sapiens OX=9606 GN=ATP6V1G1 PE=1 SV=3 | 0.966 | 0.897516642 | 0.94 | 0.7 |
| Q9BQE5 | Apolipoprotein L2 OS=Homo sapiens OX=9606 GN=APOL2 PE=1 SV=1 | 0.815 | 0.765235202 | 1.015 | 0.7 |
| P13674 | Prolyl 4-hydroxylase subunit alpha-1 OS=Homo sapiens OX=9606 GN=P4HA1 PE=1 SV=2 | 0.945 | 0.750023665 | 0.902 | 0.7 |
| Q96PU8 | Protein quaking OS=Homo sapiens OX=9606 GN=QKI PE=1 SV=1 | 0.956 | 0.809976184 | 0.936 | 0.7 |
| Q9BZH6 | WD repeat-containing protein 11 OS=Homo sapiens OX=9606 GN=WDR11 PE=1 SV=1 | 0.847 | 0.774484855 | 0.894 | 0.7 |
| G3V4W4 | Glucosamine 6-phosphate N-acetyltransferase (Fragment) OS=Homo sapiens OX=9606 GN=GNPNAT1 PE=1 SV=1 | 1.026 | 0.876510992 | 0.999 | 0.7 |
| Q9NQZ2 | Something about silencing protein 10 OS=Homo sapiens OX=9606 GN=UTP3 PE=1 SV=1 | 0.971 | 0.990418431 | 0.888 | 0.7 |
| Q96DE0 | U8 snoRNA-decapping enzyme OS=Homo sapiens OX=9606 GN=NUDT16 PE=1 SV=2 | 1.005 | 0.902016292 | 0.993 | 0.8 |
| B1AH58 | Intraflagellar transport protein 27 homolog (Fragment) OS=Homo sapiens OX=9606 GN=IFT27 PE=1 SV=1 | 0.969 | 0.876510992 | 0.973 | 0.7 |
| P23497 | Nuclear autoantigen Sp-100 OS=Homo sapiens OX=9606 GN=SP100 PE=1 SV=3 | 1.033 | 0.889450577 | 1.011 | 0.8 |
| Q16512 | Serine/threonine-protein kinase N1 OS=Homo sapiens OX=9606 GN=PKN1 PE=1 SV=2 | 0.968 | 0.876510992 | 0.975 | 0.7 |
| Q8NF37 | Lysophosphatidylcholine acyltransferase 1 OS=Homo sapiens OX=9606 GN=LPCAT1 PE=1 SV=2 | 1.008 | 0.987438982 | 1.08 | 0.7 |
| Q9UBM7 | 7-dehydrocholesterol reductase OS=Homo sapiens OX=9606 GN=DHCR7 PE=1 SV=1 | 0.965 | 0.809976184 | 1.007 | 0.9 |
| Q9Y3A3 | MOB-like protein phocein OS=Homo sapiens OX=9606 GN=MOB4 PE=1 SV=1 | 0.961 | 0.829390484 | 0.973 | 0.7 |
| Q96BJ3 | Axin interactor, dorsalization-associated protein OS=Homo sapiens OX=9606 GN=AIDA PE=1 SV=1 | 0.98 | 0.924631778 | 0.941 | 0.7 |

| | | | | | |
|---|---|---|---|---|---|
| P28331 | NADH-ubiquinone oxidoreductase 75 kDa subunit, mitochondrial OS=Homo sapiens OX=9606 GN=NDUFS1 PE=1 SV=3 | 0.946 | 0.765235202 | 0.923 | 0.7 |
| Q86U42 | Polyadenylate-binding protein 2 OS=Homo sapiens OX=9606 GN=PABPN1 PE=1 SV=3 | 1.028 | 0.985324977 | 0.937 | 0.7 |
| Q8WVM8 | Sec1 family domain-containing protein 1 OS=Homo sapiens OX=9606 GN=SCFD1 PE=1 SV=4 | 0.976 | 0.750023665 | 0.978 | 0.7 |
| Q9BQA1 | Methylosome protein 50 OS=Homo sapiens OX=9606 GN=WDR77 PE=1 SV=1 | 0.955 | 0.750023665 | 0.962 | 0.8 |
| P09543 | 2',3'-cyclic-nucleotide 3'-phosphodiesterase OS=Homo sapiens OX=9606 GN=CNP PE=1 SV=2 | 1.017 | 0.985324977 | 0.944 | 0.7 |
| Q9UKX7 | Nuclear pore complex protein Nup50 OS=Homo sapiens OX=9606 GN=NUP50 PE=1 SV=2 | 0.963 | 0.984200071 | 0.97 | 0.7 |
| Q9BZV1 | UBX domain-containing protein 6 OS=Homo sapiens OX=9606 GN=UBXN6 PE=1 SV=1 | 1.016 | 0.889450577 | 0.914 | 0.7 |
| P19823 | Inter-alpha-trypsin inhibitor heavy chain H2 OS=Homo sapiens OX=9606 GN=ITIH2 PE=1 SV=2 | 1.086 | 0.750023665 | 0.963 | 0.7 |
| Q92538 | Golgi-specific brefeldin A-resistance guanine nucleotide exchange factor 1 OS=Homo sapiens OX=9606 GN=GBF1 PE=1 SV=2 | 0.981 | 0.978562349 | 1.095 | 0.7 |
| Q99575 | Ribonucleases P/MRP protein subunit POP1 OS=Homo sapiens OX=9606 GN=POP1 PE=1 SV=2 | 1.032 | 0.832436816 | 0.981 | 0.9 |
| Q9NVT9 | Armadillo repeat-containing protein 1 OS=Homo sapiens OX=9606 GN=ARMC1 PE=1 SV=1 | 0.998 | 0.920144076 | 0.9 | 0.7 |
| O15254 | Peroxisomal acyl-coenzyme A oxidase 3 OS=Homo sapiens OX=9606 GN=ACOX3 PE=1 SV=2 | 0.956 | 0.840158726 | 0.915 | 0.7 |
| J3KQ18 | D-dopachrome decarboxylase OS=Homo sapiens OX=9606 GN=DDT PE=1 SV=1 | 1.028 | 0.917008272 | 0.972 | 0.8 |
| B5MC59 | Replication protein A 14 kDa subunit OS=Homo sapiens OX=9606 GN=RPA3 PE=1 SV=1 | 1.01 | 0.994361756 | 0.842 | 0.7 |
| Q8N6T3 | ADP-ribosylation factor GTPase-activating protein 1 OS=Homo sapiens OX=9606 GN=ARFGAP1 PE=1 SV=2 | 1.082 | 0.837893552 | 0.993 | 0.9 |
| Q8ND71 | GTPase IMAP family member 8 OS=Homo sapiens OX=9606 GN=GIMAP8 PE=2 SV=2 | 0.955 | 0.750023665 | 0.939 | 0.7 |
| F8VZQ9 | SAP domain-containing ribonucleoprotein OS=Homo sapiens OX=9606 GN=SARNP PE=1 SV=1 | 1.012 | 0.971280121 | 0.992 | 0.9 |
| A0A6I8PRZ0 | DnaJ homolog subfamily C member 7 (Fragment) OS=Homo sapiens OX=9606 GN=DNAJC7 PE=4 SV=1 | 1.012 | 0.881099595 | 0.959 | 0.7 |
| P42285 | Exosome RNA helicase MTR4 OS=Homo sapiens OX=9606 GN=MTREX PE=1 SV=3 | 1.014 | 0.969984528 | 0.966 | 0.7 |
| Q9BVC6 | Transmembrane protein 109 OS=Homo sapiens OX=9606 GN=TMEM109 PE=1 SV=1 | 0.953 | 0.750023665 | 1.019 | 0.9 |
| P07099 | Epoxide hydrolase 1 OS=Homo sapiens OX=9606 GN=EPHX1 PE=1 SV=1 | 0.972 | 0.765235202 | 0.905 | 0.7 |
| Q01085 | Nucleolysin TIAR OS=Homo sapiens OX=9606 GN=TIAL1 PE=1 SV=1 | 0.991 | 0.943791335 | 0.999 | 0.8 |
| Q8IXB1 | DnaJ homolog subfamily C member 10 OS=Homo sapiens OX=9606 GN=DNAJC10 PE=1 SV=2 | 1.031 | 0.913279108 | 1.01 | 0.9 |
| Q5JTZ9 | Alanine--tRNA ligase, mitochondrial OS=Homo sapiens OX=9606 GN=AARS2 PE=1 SV=1 | 1.04 | 0.883217803 | 0.761 | 0.7 |
| B4E1B9 | Cell surface hyaluronidase OS=Homo sapiens OX=9606 GN=CEMIP2 PE=1 SV=1 | 1.174 | 0.80706255 | 0.883 | 0.7 |
| Q02252 | Methylmalonate-semialdehyde dehydrogenase [acylating], mitochondrial OS=Homo sapiens OX=9606 GN=ALDH6A1 PE=1 SV=2 | 0.963 | 0.917008272 | 0.917 | 0.7 |
| Q92747 | Actin-related protein 2/3 complex subunit 1A OS=Homo sapiens OX=9606 GN=ARPC1A PE=2 SV=2 | 0.974 | 0.87017509 | 0.869 | 0.7 |
| M0R165 | Epidermal growth factor receptor substrate 15-like 1 OS=Homo sapiens OX=9606 GN=EPS15L1 PE=1 SV=1 | 0.884 | 0.889450577 | 0.973 | 0.7 |
| C9J5C3 | Programmed cell death protein 10 (Fragment) OS=Homo sapiens OX=9606 GN=PDCD10 PE=1 SV=1 | 1.051 | 0.750023665 | 0.991 | 0.7 |
| Q9UJX2 | Cell division cycle protein 23 homolog OS=Homo sapiens OX=9606 GN=CDC23 PE=1 SV=3 | 1.145 | 0.792509319 | 0.795 | 0.7 |
| A0A0A0MRV4 | Prostamide/prostaglandin F synthase OS=Homo sapiens OX=9606 GN=PRXL2B PE=1 SV=1 | 0.916 | 0.766667967 | 0.836 | 0.7 |
| O75347 | Tubulin-specific chaperone A OS=Homo sapiens OX=9606 GN=TBCA PE=1 SV=3 | 0.969 | 0.765235202 | 1.037 | 0.7 |
| Q9Y3B3 | Transmembrane emp24 domain-containing protein 7 OS=Homo sapiens OX=9606 GN=TMED7 PE=1 SV=2 | 1.047 | 0.700264056 | 0.948 | 0.7 |
| Q5TH30 | N-myc downstream-regulated gene 3 protein OS=Homo sapiens OX=9606 GN=NDRG3 PE=1 SV=1 | 0.967 | 0.820396985 | 1.013 | 0.9 |
| Q9Y2Q9 | 28S ribosomal protein S28, mitochondrial OS=Homo sapiens OX=9606 GN=MRPS28 PE=1 SV=1 | 1.037 | 0.985324977 | 0.895 | 0.7 |
| Q96A65 | Exocyst complex component 4 OS=Homo sapiens OX=9606 GN=EXOC4 PE=1 SV=1 | 1.019 | 0.97688166 | 0.886 | 0.7 |
| Q9H7Z7 | Prostaglandin E synthase 2 OS=Homo sapiens OX=9606 GN=PTGES2 PE=1 SV=1 | 0.947 | 0.820451399 | 0.923 | 0.7 |
| Q9UID3 | Vacuolar protein sorting-associated protein 51 homolog OS=Homo sapiens OX=9606 GN=VPS51 PE=1 SV=2 | 0.922 | 0.889482529 | 0.883 | 0.7 |
| Q5VSL9 | Striatin-interacting protein 1 OS=Homo sapiens OX=9606 GN=STRIP1 PE=1 SV=1 | 1.007 | 0.954224541 | 0.963 | 0.8 |
| O43815 | Striatin OS=Homo sapiens OX=9606 GN=STRN PE=1 SV=4 | 1.018 | 0.876510992 | 0.891 | 0.7 |
| Q9UK41 | Vacuolar protein sorting-associated protein 28 homolog OS=Homo sapiens OX=9606 GN=VPS28 PE=1 SV=1 | 0.998 | 0.957217311 | 0.925 | 0.7 |
| P62256 | Ubiquitin-conjugating enzyme E2 H OS=Homo sapiens OX=9606 GN=UBE2H PE=1 SV=1 | 1.008 | 0.973388826 | 0.95 | 0.7 |
| P19784 | Casein kinase II subunit alpha' OS=Homo sapiens OX=9606 GN=CSNK2A2 PE=1 SV=1 | 0.933 | 0.750023665 | 0.929 | 0.7 |
| P04004 | Vitronectin OS=Homo sapiens OX=9606 GN=VTN PE=1 SV=1 | 0.974 | 0.901934028 | 0.659 | 0.7 |
| O75531 | Barrier-to-autointegration factor OS=Homo sapiens OX=9606 GN=BANF1 PE=1 SV=1 | 0.91 | 0.765235202 | 0.71 | 0.7 |
| A0A087X1G7 | Selenoprotein F OS=Homo sapiens OX=9606 GN=SELENOF PE=1 SV=1 | 0.942 | 0.765235202 | 1.007 | 0.9 |
| D6RA70 | RNA helicase OS=Homo sapiens OX=9606 GN=YTHDC2 PE=1 SV=1 | 1.043 | 0.750023665 | 0.933 | 0.7 |
| Q9BRF8 | Serine/threonine-protein phosphatase CPPED1 OS=Homo sapiens OX=9606 GN=CPPED1 PE=1 SV=3 | 0.985 | 0.827871327 | 0.906 | 0.7 |
| H3BST1 | Biogenesis of lysosome-related organelles complex 1 subunit 6 OS=Homo sapiens OX=9606 GN=BLOC1S6 PE=1 SV=1 | 1.12 | 0.750023665 | 0.949 | 0.7 |
| O95486 | Protein transport protein Sec24A OS=Homo sapiens OX=9606 GN=SEC24A PE=1 SV=2 | 0.991 | 0.984805107 | 0.837 | 0.7 |
| P10909 | Clusterin OS=Homo sapiens OX=9606 GN=CLU PE=1 SV=1 | 0.968 | 0.889482529 | 0.826 | 0.7 |
| O43837 | Isocitrate dehydrogenase [NAD] subunit beta, mitochondrial OS=Homo sapiens OX=9606 GN=IDH3B PE=1 SV=2 | 0.969 | 0.765235202 | 0.983 | 0.8 |
| Q9H074 | Polyadenylate-binding protein-interacting protein 1 OS=Homo sapiens OX=9606 GN=PAIP1 PE=1 SV=1 | 1.022 | 0.750023665 | 0.98 | 0.7 |
| A0A2U3TZM0 | DNA helicase OS=Homo sapiens OX=9606 GN=CHD4 PE=1 SV=1 | 1.122 | 0.765235202 | 1.026 | 0.7 |
| O75828 | Carbonyl reductase [NADPH] 3 OS=Homo sapiens OX=9606 GN=CBR3 PE=1 SV=3 | 1.116 | 0.750023665 | 0.945 | 0.7 |
| Q99598 | Translin-associated protein X OS=Homo sapiens OX=9606 GN=TSNAX PE=1 SV=1 | 0.987 | 0.972581915 | 0.935 | 0.7 |
| D6RAN1 | PDZ and LIM domain protein 7 (Fragment) OS=Homo sapiens OX=9606 GN=PDLIM7 PE=1 SV=1 | | | | |
| Q8TDN6 | Ribosome biogenesis protein BRX1 homolog OS=Homo sapiens OX=9606 GN=BRIX1 PE=1 SV=2 | 0.995 | 0.917008272 | 0.906 | 0.8 |
| I3L4J1 | Vesicle-fusing ATPase (Fragment) OS=Homo sapiens OX=9606 PE=1 SV=3 | 0.947 | 0.983196715 | 1.029 | 0.8 |
| Q9NRX4 | 14 kDa phosphohistidine phosphatase OS=Homo sapiens OX=9606 GN=PHPT1 PE=1 SV=1 | 1.016 | 0.756287988 | 0.946 | 0.7 |
| Q9UI12 | V-type proton ATPase subunit H OS=Homo sapiens OX=9606 GN=ATP6V1H PE=1 SV=1 | 0.97 | 0.794788863 | 0.915 | 0.7 |
| Q9NT62 | Ubiquitin-like-conjugating enzyme ATG3 OS=Homo sapiens OX=9606 GN=ATG3 PE=1 SV=1 | 0.962 | 0.750023665 | 0.965 | 0.7 |
| E9PHM2 | Leucine--tRNA ligase OS=Homo sapiens OX=9606 GN=LARS2 PE=1 SV=1 | 0.957 | 0.765235202 | 0.918 | 0.7 |
| Q9UHD9 | Ubiquilin-2 OS=Homo sapiens OX=9606 GN=UBQLN2 PE=1 SV=2 | 0.95 | 0.765235202 | 1.003 | 0.8 |
| A0A3B3IUD2 | Protein misato homolog 1 OS=Homo sapiens OX=9606 GN=MSTO1 PE=1 SV=1 | 1.022 | 0.9962822 | 0.917 | 0.7 |
| Q9UKV3 | Apoptotic chromatin condensation inducer in the nucleus OS=Homo sapiens OX=9606 GN=ACIN1 PE=1 SV=2 | 1.065 | 0.750023665 | 0.996 | 0.8 |
| P49662 | Caspase-4 OS=Homo sapiens OX=9606 GN=CASP4 PE=1 SV=1 | 1.004 | 0.994361756 | 0.957 | 0.8 |
| A8MU27 | Small ubiquitin-related modifier 3 OS=Homo sapiens OX=9606 GN=SUMO3 PE=1 SV=1 | 1.043 | 0.860909277 | 0.992 | 0.9 |
| O96019 | Actin-like protein 6A OS=Homo sapiens OX=9606 GN=ACTL6A PE=1 SV=1 | 1.05 | 0.754126086 | 1.035 | 0.7 |
| Q08752 | Peptidyl-prolyl cis-trans isomerase D OS=Homo sapiens OX=9606 GN=PPID PE=1 SV=3 | 0.95 | 0.829390484 | 0.848 | 0.7 |
| Q99570 | Phosphoinositide 3-kinase regulatory subunit 4 OS=Homo sapiens OX=9606 GN=PIK3R4 PE=1 SV=3 | 0.914 | 0.765235202 | 0.9 | 0.7 |
| P68402 | Platelet-activating factor acetylhydrolase IB subunit beta OS=Homo sapiens OX=9606 GN=PAFAH1B2 PE=1 SV=1 | 0.985 | 0.869995963 | 1.002 | 0.7 |
| P12035 | Keratin, type II cytoskeletal 3 OS=Homo sapiens OX=9606 GN=KRT3 PE=1 SV=3 | 0.973 | 0.904500837 | 0.745 | 0.7 |
| Q86VS8 | Protein Hook homolog 3 OS=Homo sapiens OX=9606 GN=HOOK3 PE=1 SV=2 | 1.039 | 0.830179531 | 1.052 | 0.8 |
| Q15388 | Mitochondrial import receptor subunit TOM20 homolog OS=Homo sapiens OX=9606 GN=TOMM20 PE=1 SV=1 | 0.956 | 0.765235202 | 0.908 | 0.7 |
| E9PHI6 | Dynein light intermediate chain OS=Homo sapiens OX=9606 GN=DYNC1LI1 PE=1 SV=1 | 1.009 | 0.886561233 | 0.986 | 0.7 |
| P14678 | Small nuclear ribonucleoprotein-associated proteins B and B' OS=Homo sapiens OX=9606 GN=SNRPB PE=1 SV=2 | 0.981 | 0.912604818 | 0.925 | 0.7 |
| Q01081 | Splicing factor U2AF 35 kDa subunit OS=Homo sapiens OX=9606 GN=U2AF1 PE=1 SV=3 | 1.003 | 0.95980123 | 0.98 | 0.9 |
| Q9HBI1 | Beta-parvin OS=Homo sapiens OX=9606 GN=PARVB PE=1 SV=1 | 0.886 | 0.765235202 | 0.945 | 0.7 |
| Q15061 | WD repeat-containing protein 43 OS=Homo sapiens OX=9606 GN=WDR43 PE=1 SV=3 | 0.977 | 0.990981579 | 0.912 | 0.8 |
| Q6P2E9 | Enhancer of mRNA-decapping protein 4 OS=Homo sapiens OX=9606 GN=EDC4 PE=1 SV=1 | 1.021 | 0.845656075 | 1.023 | 0.9 |
| Q9Y4E8 | Ubiquitin carboxyl-terminal hydrolase 15 OS=Homo sapiens OX=9606 GN=USP15 PE=1 SV=3 | 0.949 | 0.803390775 | 0.981 | 0.7 |
| P49773 | Histidine triad nucleotide-binding protein 1 OS=Homo sapiens OX=9606 GN=HINT1 PE=1 SV=2 | 1.009 | 0.95980123 | 0.981 | 0.9 |
| Q92791 | Endoplasmic reticulum protein SC65 OS=Homo sapiens OX=9606 GN=P3H4 PE=1 SV=1 | 0.881 | 0.750023665 | 0.986 | 0.7 |
| O75323 | Protein NipSnap homolog 2 OS=Homo sapiens OX=9606 GN=NIPSNAP2 PE=1 SV=1 | | | | |
| O94903 | Pyridoxal phosphate homeostasis protein OS=Homo sapiens OX=9606 GN=PLPBP PE=1 SV=1 | 0.978 | 0.765235202 | 1.024 | 0.8 |
| Q9Y6B6 | GTP-binding protein SAR1b OS=Homo sapiens OX=9606 GN=SAR1B PE=1 SV=1 | 0.974 | 0.809976184 | 0.883 | 0.7 |
| Q5TEC6 | Histone H3 OS=Homo sapiens OX=9606 GN=H3-2 PE=1 SV=1 | 0.791 | 0.765235202 | 0.826 | 0.7 |
| P51610 | Host cell factor 1 OS=Homo sapiens OX=9606 GN=HCFC1 PE=1 SV=2 | 1.119 | 0.750023665 | 0.937 | 0.7 |
| Q6P1J9 | Parafibromin OS=Homo sapiens OX=9606 GN=CDC73 PE=1 SV=1 | 1.044 | 0.750023665 | 0.969 | 0.8 |
| K7ENV7 | Isochorismatase domain-containing protein 2 (Fragment) OS=Homo sapiens OX=9606 GN=ISOC2 PE=1 SV=8 | 1.109 | 0.750023665 | 0.988 | 0.8 |
| P17900 | Ganglioside GM2 activator OS=Homo sapiens OX=9606 GN=GM2A PE=1 SV=4 | 0.961 | 0.798032881 | 0.952 | 0.7 |
| Q15428 | Splicing factor 3A subunit 2 OS=Homo sapiens OX=9606 GN=SF3A2 PE=1 SV=2 | 1.01 | 0.892311206 | 1.028 | 0.7 |
| Q92990 | Glomulin OS=Homo sapiens OX=9606 GN=GLMN PE=1 SV=2 | 1.057 | 0.896983349 | 0.794 | 0.7 |
| Q15386 | Ubiquitin-protein ligase E3C OS=Homo sapiens OX=9606 GN=UBE3C PE=1 SV=3 | 1.012 | 0.901934028 | 0.907 | 0.7 |
| P50583 | Bis(5'-nucleosyl)-tetraphosphatase [asymmetrical] OS=Homo sapiens OX=9606 GN=NUDT2 PE=1 SV=3 | 0.958 | 0.830179531 | 0.961 | 0.7 |

| | | | | | |
|---|---|---|---|---|---|
| Q53EP0 | Fibronectin type III domain-containing protein 3B OS=Homo sapiens OX=9606 GN=FNDC3B PE=1 SV=2 | 0.99 | 0.968322408 | 0.907 | 0.7 |
| P08473 | Neprilysin OS=Homo sapiens OX=9606 GN=MME PE=1 SV=2 | 1.017 | 0.983953845 | 0.975 | 0.7 |
| Q9C0E2 | Exportin-4 OS=Homo sapiens OX=9606 GN=XPO4 PE=1 SV=2 | 0.956 | 0.703015003 | 0.854 | 0.7 |
| Q06481 | Amyloid-like protein 2 OS=Homo sapiens OX=9606 GN=APLP2 PE=1 SV=2 | 1.043 | 0.952140784 | 1.001 | 0.9 |
| Q9Y2D4 | Exocyst complex component 6B OS=Homo sapiens OX=9606 GN=EXOC6B PE=1 SV=3 | 1.096 | 0.765235202 | 0.765 | 0.7 |
| A0A1B0GUE3 | Acid ceramidase OS=Homo sapiens OX=9606 GN=ASAH1 PE=1 SV=1 | 1.005 | 0.985324977 | 0.917 | 0.7 |
| Q9NZL9 | Methionine adenosyltransferase 2 subunit beta OS=Homo sapiens OX=9606 GN=MAT2B PE=1 SV=1 | 1.038 | 0.765235202 | 0.988 | 0.9 |
| P45877 | Peptidyl-prolyl cis-trans isomerase C OS=Homo sapiens OX=9606 GN=PPIC PE=1 SV=1 | 0.951 | 0.883924281 | 0.976 | 0.9 |
| Q12979 | Active breakpoint cluster region-related protein OS=Homo sapiens OX=9606 GN=ABR PE=1 SV=2 | 1.058 | 0.765235202 | 0.881 | 0.7 |
| P26885 | Peptidyl-prolyl cis-trans isomerase FKBP2 OS=Homo sapiens OX=9606 GN=FKBP2 PE=1 SV=2 | 0.996 | 0.930093779 | 0.952 | 0.7 |
| Q9H1I8 | Activating signal cointegrator 1 complex subunit 2 OS=Homo sapiens OX=9606 GN=ASCC2 PE=1 SV=3 | 1.068 | 0.986048646 | 0.748 | 0.7 |
| J3QRG0 | U3 small nucleolar RNA-associated protein 18 homolog (Fragment) OS=Homo sapiens OX=9606 GN=UTP18 PE=1 SV=2 | 1.163 | 0.829390484 | 0.87 | 0.8 |
| A0A0A0MTC1 | E3 ubiquitin-protein ligase RNF213 OS=Homo sapiens OX=9606 GN=RNF213 PE=1 SV=1 | 0.941 | 0.765235202 | 0.819 | 0.7 |
| Q5TFE4 | 5'-nucleotidase domain-containing protein 1 OS=Homo sapiens OX=9606 GN=NT5DC1 PE=1 SV=1 | 0.952 | 0.830958723 | 1.006 | 0.8 |
| Q96BN8 | Ubiquitin thioesterase otulin OS=Homo sapiens OX=9606 GN=OTULIN PE=1 SV=3 | 1.025 | 0.882454148 | 0.821 | 0.7 |
| Q9NP72 | Ras-related protein Rab-18 OS=Homo sapiens OX=9606 GN=RAB18 PE=1 SV=1 | 0.975 | 0.889450577 | 1.024 | 0.9 |
| P61457 | Pterin-4-alpha-carbinolamine dehydratase OS=Homo sapiens OX=9606 GN=PCBD1 PE=1 SV=2 | 1.023 | 0.921087329 | 0.987 | 0.8 |
| Q8WZ82 | Esterase OVCA2 OS=Homo sapiens OX=9606 GN=OVCA2 PE=1 SV=1 | 1.096 | 0.765235202 | 0.99 | 0.7 |
| Q5NDL2 | EGF domain-specific O-linked N-acetylglucosamine transferase OS=Homo sapiens OX=9606 GN=EOGT PE=1 SV=1 | 0.971 | 0.921087329 | 0.939 | 0.7 |
| O95155 | Ubiquitin conjugation factor E4 B OS=Homo sapiens OX=9606 GN=UBE4B PE=1 SV=1 | 1.06 | 0.863796857 | 0.982 | 0.7 |
| Q9P2I0 | Cleavage and polyadenylation specificity factor subunit 2 OS=Homo sapiens OX=9606 GN=CPSF2 PE=1 SV=2 | 1.027 | 0.750023665 | 0.985 | 0.7 |
| O15231 | Zinc finger protein 185 OS=Homo sapiens OX=9606 GN=ZNF185 PE=1 SV=3 | 0.995 | 0.946055791 | 1 | 0.8 |
| M0R0X1 | Ras-related protein Rab-4B (Fragment) OS=Homo sapiens OX=9606 GN=RAB4B PE=1 SV=1 | | | | |
| Q9BU89 | Deoxyhypusine hydroxylase OS=Homo sapiens OX=9606 GN=DOHH PE=1 SV=1 | 0.904 | 0.958664425 | 0.99 | 0.8 |
| A0A0G2JHC2 | PPP1R18 OS=Homo sapiens OX=9606 GN=PPP1R18 PE=1 SV=1 | 1.04 | 0.985763982 | 0.897 | 0.7 |
| E9PK52 | Band 4.1-like protein 2 OS=Homo sapiens OX=9606 GN=EPB41L2 PE=1 SV=1 | 0.968 | 0.827871327 | 0.942 | 0.7 |
| P60604 | Ubiquitin-conjugating enzyme E2 G2 OS=Homo sapiens OX=9606 GN=UBE2G2 PE=1 SV=1 | 0.896 | 0.765235202 | 0.984 | 0.9 |
| Q5T2E6 | Armadillo-like helical domain-containing protein 3 OS=Homo sapiens OX=9606 GN=ARMH3 PE=1 SV=1 | 1.009 | 0.948132568 | 0.745 | 0.7 |
| Q16864 | V-type proton ATPase subunit F OS=Homo sapiens OX=9606 GN=ATP6V1F PE=1 SV=2 | 0.967 | 0.968952473 | 0.811 | 0.7 |
| Q9BPW8 | Protein NipSnap homolog 1 OS=Homo sapiens OX=9606 GN=NIPSNAP1 PE=1 SV=1 | 1.069 | 0.765235202 | 0.947 | 0.7 |
| B4DSN5 | Tyrosine-protein phosphatase non-receptor type OS=Homo sapiens OX=9606 GN=PTPN1 PE=1 SV=1 | 0.983 | 0.983953845 | 0.973 | 0.8 |
| Q8NEU8 | DCC-interacting protein 13-beta OS=Homo sapiens OX=9606 GN=APPL2 PE=1 SV=3 | 0.982 | 0.983953845 | 0.975 | 0.7 |
| A0A0A0MTL6 | Rho family-interacting cell polarization regulator 1 OS=Homo sapiens OX=9606 GN=RIPOR1 PE=1 SV=1 | 0.96 | 0.983953845 | 0.933 | 0.7 |
| Q5TDF0 | Cancer-related nucleoside-triphosphatase OS=Homo sapiens OX=9606 GN=NTPCR PE=1 SV=1 | 0.985 | 0.913279108 | 0.88 | 0.7 |
| P02533 | Keratin, type I cytoskeletal 14 OS=Homo sapiens OX=9606 GN=KRT14 PE=1 SV=4 | 0.752 | 0.765235202 | 0.597 | 0.7 |
| G3V1R5 | Nardilysin OS=Homo sapiens OX=9606 GN=NRDC PE=1 SV=1 | 1.008 | 0.889450577 | 0.97 | 0.8 |
| F8VWK8 | Tetraspanin (Fragment) OS=Homo sapiens OX=9606 GN=CD63 PE=1 SV=1 | 0.975 | 0.886561233 | 0.913 | 0.7 |
| Q8NE71 | ATP-binding cassette sub-family F member 1 OS=Homo sapiens OX=9606 GN=ABCF1 PE=1 SV=2 | 1.021 | 0.829390484 | 1.017 | 0.7 |
| P04066 | Tissue alpha-L-fucosidase OS=Homo sapiens OX=9606 GN=FUCA1 PE=1 SV=4 | 0.933 | 0.935391021 | 1.002 | 0.8 |
| P11274 | Breakpoint cluster region protein OS=Homo sapiens OX=9606 GN=BCR PE=1 SV=2 | 1.007 | 0.98333918 | 0.924 | 0.7 |
| I3L2C7 | Gem-associated protein 4 OS=Homo sapiens OX=9606 GN=GEMIN4 PE=1 SV=1 | 0.978 | 0.824101611 | 0.913 | 0.9 |
| P16930 | Fumarylacetoacetase OS=Homo sapiens OX=9606 GN=FAH PE=1 SV=2 | 0.934 | 0.750023665 | 0.96 | 0.7 |
| Q96S52 | GPI transamidase component PIG-S OS=Homo sapiens OX=9606 GN=PIGS PE=1 SV=3 | 0.933 | 0.814337508 | 0.976 | 0.8 |
| P35250 | Replication factor C subunit 2 OS=Homo sapiens OX=9606 GN=RFC2 PE=1 SV=3 | 0.902 | 0.703015003 | 1.001 | 0.9 |
| Q9NTJ3 | Structural maintenance of chromosomes protein 4 OS=Homo sapiens OX=9606 GN=SMC4 PE=1 SV=2 | 0.936 | 0.765235202 | 1.01 | 0.9 |
| Q00537 | Cyclin-dependent kinase 17 OS=Homo sapiens OX=9606 GN=CDK17 PE=1 SV=2 | 0.968 | 0.827091161 | 0.984 | 0.7 |
| P84090 | Enhancer of rudimentary homolog OS=Homo sapiens OX=9606 GN=ERH PE=1 SV=1 | 0.923 | 0.882455549 | 0.933 | 0.8 |
| Q9NX62 | Golgi-resident adenosine 3',5'-bisphosphate 3'-phosphatase OS=Homo sapiens OX=9606 GN=BPNT2 PE=1 SV=1 | 1.022 | 0.95980123 | 1.046 | 0.7 |
| Q9Y276 | Mitochondrial chaperone BCS1 OS=Homo sapiens OX=9606 GN=BCS1L PE=1 SV=1 | 1.041 | 0.972946678 | 0.85 | 0.7 |
| O43617 | Trafficking protein particle complex subunit 3 OS=Homo sapiens OX=9606 GN=TRAPPC3 PE=1 SV=1 | 1 | 0.873607677 | 1.029 | 0.7 |
| Q8WUI4 | Histone deacetylase 7 OS=Homo sapiens OX=9606 GN=HDAC7 PE=1 SV=2 | 1.155 | 0.680740328 | 0.897 | 0.7 |
| P39748 | Flap endonuclease 1 OS=Homo sapiens OX=9606 GN=FEN1 PE=1 SV=1 | 1.077 | 0.750023665 | 1.009 | 0.7 |
| A0A0U1RR22 | Protein kinase C and casein kinase substrate in neurons protein 2 (Fragment) OS=Homo sapiens OX=9606 GN=PACSIN2 PE=1 SV=1 | 1.087 | 0.785314375 | 1.002 | 0.7 |
| O75884 | Serine hydrolase RBBP9 OS=Homo sapiens OX=9606 GN=RBBP9 PE=1 SV=2 | 1 | 0.94960029 | 0.972 | 0.7 |
| A0A087WUC6 | Microsomal signal peptidase 25 kDa subunit OS=Homo sapiens OX=9606 GN=SPCS2 PE=1 SV=1 | 1.016 | 0.863796857 | 0.973 | 0.8 |
| Q5VU11 | Ribonuclease P protein subunit p30 (Fragment) OS=Homo sapiens OX=9606 GN=RPP30 PE=1 SV=1 | 0.988 | 0.913279108 | 0.934 | 0.7 |
| B8ZZK2 | Gamma-glutamylcyclotransferase OS=Homo sapiens OX=9606 GN=GGCT PE=1 SV=1 | 0.974 | 0.750023665 | 0.99 | 0.7 |
| Q7Z4Q2 | HEAT repeat-containing protein 3 OS=Homo sapiens OX=9606 GN=HEATR3 PE=1 SV=2 | 1.059 | 0.924631778 | 0.913 | 0.7 |
| A0A286YFF8 | Protein MON2 homolog OS=Homo sapiens OX=9606 GN=MON2 PE=1 SV=1 | 1.018 | 0.9962822 | 0.896 | 0.7 |
| Q9H488 | GDP-fucose protein O-fucosyltransferase 1 OS=Homo sapiens OX=9606 GN=POFUT1 PE=1 SV=1 | 1.02 | 0.866054955 | 0.95 | 0.7 |
| Q14997 | Proteasome activator complex subunit 4 OS=Homo sapiens OX=9606 GN=PSME4 PE=1 SV=2 | 1.011 | 0.904500837 | 0.924 | 0.7 |
| Q9Y6A4 | Cilia- and flagella-associated protein 20 OS=Homo sapiens OX=9606 GN=CFAP20 PE=1 SV=1 | 0.99 | 0.959800973 | 0.9 | 0.7 |
| Q3ZCQ8 | Mitochondrial import inner membrane translocase subunit TIM50 OS=Homo sapiens OX=9606 GN=TIMM50 PE=1 SV=2 | 1.056 | 0.869995963 | 0.981 | 0.8 |
| Q8NBJ5 | Procollagen galactosyltransferase 1 OS=Homo sapiens OX=9606 GN=COLGALT1 PE=1 SV=1 | 0.977 | 0.774484855 | 0.861 | 0.7 |
| A0A3B3ITW3 | ATP-binding cassette sub-family D member 3 OS=Homo sapiens OX=9606 GN=ABCD3 PE=1 SV=2 | 1.139 | 0.765235202 | 0.963 | 0.7 |
| H0YK99 | DnaJ homolog subfamily C member 17 OS=Homo sapiens OX=9606 GN=DNAJC17 PE=1 SV=1 | 1.144 | 0.815069882 | 0.992 | 0.9 |
| P02765 | Alpha-2-HS-glycoprotein OS=Homo sapiens OX=9606 GN=AHSG PE=1 SV=2 | 1.367 | 0.920074608 | 0.971 | 0.8 |
| A8MQ02 | Afadin OS=Homo sapiens OX=9606 GN=AFDN PE=1 SV=2 | 1.143 | 0.766667967 | 0.87 | 0.7 |
| A0A140T998 | Proteasome endopeptidase complex OS=Homo sapiens OX=9606 GN=PSMB8 PE=1 SV=1 | 0.965 | 0.750023665 | 1.021 | 0.7 |
| P63173 | 60S ribosomal protein L38 OS=Homo sapiens OX=9606 GN=RPL38 PE=1 SV=2 | 1.216 | 0.750023665 | 1.05 | 0.8 |
| Q8N183 | NADH dehydrogenase [ubiquinone] 1 alpha subcomplex assembly factor 2 OS=Homo sapiens OX=9606 GN=NDUFAF2 PE=1 SV=1 | 1.027 | 0.860909277 | 0.934 | 0.7 |
| Q9UBR2 | Cathepsin Z OS=Homo sapiens OX=9606 GN=CTSZ PE=1 SV=1 | 1.012 | 0.991211283 | 1.172 | 0.8 |
| Q9GZP4 | PITH domain-containing protein 1 OS=Homo sapiens OX=9606 GN=PITHD1 PE=1 SV=1 | 0.936 | 0.839954644 | 0.85 | 0.7 |
| P51688 | N-sulphoglucosamine sulphohydrolase OS=Homo sapiens OX=9606 GN=SGSH PE=1 SV=1 | 1.008 | 0.900630465 | 0.953 | 0.7 |
| Q14011 | Cold-inducible RNA-binding protein OS=Homo sapiens OX=9606 GN=CIRBP PE=1 SV=1 | 1.014 | 0.925693085 | 0.929 | 0.7 |
| Q9H3P7 | Golgi resident protein GCP60 OS=Homo sapiens OX=9606 GN=ACBD3 PE=1 SV=4 | 1 | 0.920357062 | 1.049 | 0.7 |
| Q2TAL8 | Glutamine-rich protein 1 OS=Homo sapiens OX=9606 GN=QRICH1 PE=1 SV=1 | 1.021 | 0.912604818 | 0.926 | 0.7 |
| Q6ZVM7 | TOM1-like protein 2 OS=Homo sapiens OX=9606 GN=TOM1L2 PE=1 SV=1 | 0.997 | 0.87017509 | 0.973 | 0.7 |
| Q9P287 | BRCA2 and CDKN1A-interacting protein OS=Homo sapiens OX=9606 GN=BCCIP PE=1 SV=1 | 0.91 | 0.765235202 | 1.018 | 0.8 |
| Q71UM5 | 40S ribosomal protein S27-like OS=Homo sapiens OX=9606 GN=RPS27L PE=1 SV=3 | 1.068 | 0.765235202 | 1.051 | 0.7 |
| A0A0C4DGW6 | UPF0600 protein C5orf51 OS=Homo sapiens OX=9606 GN=C5orf51 PE=1 SV=1 | 0.857 | 0.765235202 | 0.808 | 0.7 |
| Q9Y5L4 | Mitochondrial import inner membrane translocase subunit Tim13 OS=Homo sapiens OX=9606 GN=TIMM13 PE=1 SV=1 | 1.047 | 0.765235202 | 0.916 | 0.7 |
| Q969T9 | WW domain-binding protein 2 OS=Homo sapiens OX=9606 GN=WBP2 PE=1 SV=1 | 0.939 | 0.765235202 | 0.965 | 0.7 |
| Q9H3S7 | Tyrosine-protein phosphatase non-receptor type 23 OS=Homo sapiens OX=9606 GN=PTPN23 PE=1 SV=1 | 0.921 | 0.762750886 | 0.928 | 0.7 |
| E5RGS4 | Prefoldin subunit 1 OS=Homo sapiens OX=9606 GN=PFDN1 PE=1 SV=1 | 1.018 | 0.845656075 | 1.005 | 0.7 |
| A0A2R8Y5V8 | Protein furry homolog OS=Homo sapiens OX=9606 GN=FRY PE=1 SV=1 | 1.006 | 0.991801854 | 0.886 | 0.7 |
| Q9NZJ4 | Sacsin OS=Homo sapiens OX=9606 GN=SACS PE=1 SV=2 | 1.003 | 0.983086621 | 0.888 | 0.7 |
| A0A087WYW7 | Protein Hikeshi OS=Homo sapiens OX=9606 GN=HIKESHI PE=1 SV=1 | 1.101 | 0.766667967 | 0.906 | 0.8 |
| A6NFX8 | ADP-sugar pyrophosphatase OS=Homo sapiens OX=9606 GN=NUDT5 PE=1 SV=1 | 0.977 | 0.853774607 | 0.95 | 0.7 |
| Q6PJG6 | BRCA1-associated ATM activator 1 OS=Homo sapiens OX=9606 GN=BRAT1 PE=1 SV=2 | 0.978 | 0.994361756 | 1.02 | 0.8 |
| Q9P0J1 | [Pyruvate dehydrogenase [acetyl-transferring]]-phosphatase 1, mitochondrial OS=Homo sapiens OX=9606 GN=PDP1 PE=1 SV=3 | 1.034 | 0.888073112 | 0.953 | 0.7 |
| F6U1T9 | Calcineurin subunit B type 1 OS=Homo sapiens OX=9606 GN=PPP3R1 PE=1 SV=1 | 1.039 | 0.801467784 | 0.934 | 0.7 |
| H0YIQ2 | YLP motif-containing protein 1 (Fragment) OS=Homo sapiens OX=9606 GN=YLPM1 PE=1 SV=1 | 1.136 | 0.902016292 | 0.927 | 0.7 |
| E7EVW7 | Hematopoietic lineage cell-specific protein OS=Homo sapiens OX=9606 GN=HCLS1 PE=1 SV=1 | 1.032 | 0.827871327 | 1.11 | 0.7 |
| Q7Z2W4 | Zinc finger CCCH-type antiviral protein 1 OS=Homo sapiens OX=9606 GN=ZC3HAV1 PE=1 SV=3 | 1.005 | 0.969565732 | 1.034 | 0.7 |
| Q86XP3 | ATP-dependent RNA helicase DDX42 OS=Homo sapiens OX=9606 GN=DDX42 PE=1 SV=1 | 1.028 | 0.902766821 | 0.909 | 0.7 |

| | | | | | |
|---|---|---|---|---|---|
| P80297 | Metallothionein-1X OS=Homo sapiens OX=9606 GN=MT1X PE=1 SV=1 | 0.986 | 0.908555656 | 1.621 | 0.7 |
| P42345 | Serine/threonine-protein kinase mTOR OS=Homo sapiens OX=9606 GN=MTOR PE=1 SV=1 | 0.937 | 0.834564551 | 0.945 | 0.7 |
| Q92888 | Rho guanine nucleotide exchange factor 1 OS=Homo sapiens OX=9606 GN=ARHGEF1 PE=1 SV=2 | 1.014 | 0.839845191 | 0.963 | 0.7 |
| P55212 | Caspase-6 OS=Homo sapiens OX=9606 GN=CASP6 PE=1 SV=2 | 0.954 | 0.765235202 | 0.92 | 0.7 |
| Q9UJY4 | ADP-ribosylation factor-binding protein GGA2 OS=Homo sapiens OX=9606 GN=GGA2 PE=1 SV=3 | 0.98 | 0.992258896 | 1.028 | 0. |
| Q8N668 | COMM domain-containing protein 1 OS=Homo sapiens OX=9606 GN=COMMD1 PE=1 SV=1 | 1.138 | 0.750023665 | 1.015 | 0.9 |
| A6NLM8 | Signal sequence receptor subunit delta OS=Homo sapiens OX=9606 GN=SSR4 PE=1 SV=1 | 1.05 | 0.860909277 | 1.005 | 0.9 |
| P61513 | 60S ribosomal protein L37a OS=Homo sapiens OX=9606 GN=RPL37A PE=1 SV=2 | 1.109 | 0.763877559 | 1.116 | 0.7 |
| Q13242 | Serine/arginine-rich splicing factor 9 OS=Homo sapiens OX=9606 GN=SRSF9 PE=1 SV=1 | 1.097 | 0.830179531 | 0.948 | 0.7 |
| P49903 | Selenide, water dikinase 1 OS=Homo sapiens OX=9606 GN=SEPHS1 PE=1 SV=2 | 1.028 | 0.765235202 | 0.942 | 0.7 |
| H0YJ92 | CDK-activating kinase assembly factor MAT1 (Fragment) OS=Homo sapiens OX=9606 GN=MNAT1 PE=1 SV=1 | 1.088 | 0.722091132 | 0.991 | 0.9 |
| O00186 | Syntaxin-binding protein 3 OS=Homo sapiens OX=9606 GN=STXBP3 PE=1 SV=2 | 1.062 | 0.765235202 | 0.996 | 0.7 |
| Q9NP74 | Palmdelphin OS=Homo sapiens OX=9606 GN=PALMD PE=1 SV=1 | 0.965 | 0.902016292 | 0.902 | 0.7 |
| O43765 | Small glutamine-rich tetratricopeptide repeat-containing protein alpha OS=Homo sapiens OX=9606 GN=SGTA PE=1 SV=1 | 1.012 | 0.863796857 | 1.037 | 0.7 |
| P13716 | Delta-aminolevulinic acid dehydratase OS=Homo sapiens OX=9606 GN=ALAD PE=1 SV=1 | 0.995 | 0.984805107 | 0.865 | 0.7 |
| O60493 | Sorting nexin-3 OS=Homo sapiens OX=9606 GN=SNX3 PE=1 SV=3 | 0.942 | 0.766667967 | 1.006 | 0.9 |
| Q9Y2Q3 | Glutathione S-transferase kappa 1 OS=Homo sapiens OX=9606 GN=GSTK1 PE=1 SV=3 | 0.865 | 0.765235202 | 0.923 | 0.7 |
| E5RJ09 | Erlin-2 (Fragment) OS=Homo sapiens OX=9606 GN=ERLIN2 PE=1 SV=1 | 1.019 | 0.992258896 | 1.018 | 0.9 |
| Q9BYG3 | MKI67 FHA domain-interacting nucleolar phosphoprotein OS=Homo sapiens OX=9606 GN=NIFK PE=1 SV=1 | 0.957 | 0.93503857 | 0.957 | 0.9 |
| O95292 | Vesicle-associated membrane protein-associated protein B/C OS=Homo sapiens OX=9606 GN=VAPB PE=1 SV=3 | 1.018 | 0.868906519 | 0.994 | 0.7 |
| O60234 | Glia maturation factor gamma OS=Homo sapiens OX=9606 GN=GMFG PE=1 SV=1 | 1 | 0.904500837 | 1.024 | 0.8 |
| Q9P0K7 | Ankycorbin OS=Homo sapiens OX=9606 GN=RAI14 PE=1 SV=2 | 1.009 | 0.939784289 | 1 | 0.9 |
| Q9BRX2 | Protein pelota homolog OS=Homo sapiens OX=9606 GN=PELO PE=1 SV=2 | 1.008 | 0.994361756 | 0.955 | 0.7 |
| Q5T013 | Putative hydroxypyruvate isomerase OS=Homo sapiens OX=9606 GN=HYI PE=1 SV=2 | 0.99 | 0.991801854 | 0.94 | 0.7 |
| Q8N1F7 | Nuclear pore complex protein Nup93 OS=Homo sapiens OX=9606 GN=NUP93 PE=1 SV=2 | 1.034 | 0.830415523 | 1.004 | 0.9 |
| Q13393 | Phospholipase D1 OS=Homo sapiens OX=9606 GN=PLD1 PE=1 SV=1 | 0.883 | 0.815069882 | 0.911 | 0.7 |
| P53367 | Arfaptin-1 OS=Homo sapiens OX=9606 GN=ARFIP1 PE=1 SV=2 | 1.091 | 0.750023665 | 1.002 | 0.8 |
| Q8WVY7 | Ubiquitin-like domain-containing CTD phosphatase 1 OS=Homo sapiens OX=9606 GN=UBLCP1 PE=1 SV=2 | 1.01 | 0.95980123 | 0.951 | 0.8 |
| Q9UNI6 | Dual specificity protein phosphatase 12 OS=Homo sapiens OX=9606 GN=DUSP12 PE=1 SV=1 | 0.978 | 0.970974607 | 0.937 | 0.7 |
| Q93034 | Cullin-5 OS=Homo sapiens OX=9606 GN=CUL5 PE=1 SV=4 | 1.003 | 0.984805107 | 0.926 | 0.7 |
| H7C3P9 | COP9 signalosome complex subunit 3 OS=Homo sapiens OX=9606 GN=COPS3 PE=1 SV=2 | 1.017 | 0.902016292 | 0.934 | 0.7 |
| Q9Y3U8 | 60S ribosomal protein L36 OS=Homo sapiens OX=9606 GN=RPL36 PE=1 SV=3 | 1.019 | 0.920144076 | 1.032 | 0.8 |
| O95747 | Serine/threonine-protein kinase OSR1 OS=Homo sapiens OX=9606 GN=OXSR1 PE=1 SV=1 | 1.002 | 0.901934028 | 0.977 | 0.7 |
| Q9UEY8 | Gamma-adducin OS=Homo sapiens OX=9606 GN=ADD3 PE=1 SV=1 | 1.026 | 0.902016292 | 0.94 | 0.7 |
| Q92665 | 28S ribosomal protein S31, mitochondrial OS=Homo sapiens OX=9606 GN=MRPS31 PE=1 SV=3 | 1.085 | 0.766667967 | 0.924 | 0.7 |
| F2Z2Y6 | U6 snRNA-associated Sm-like protein LSm8 OS=Homo sapiens OX=9606 GN=LSM8 PE=1 SV=1 | 1.042 | 0.765235202 | 0.961 | 0.7 |
| A8MQB8 | Synaptic functional regulator FMR1 OS=Homo sapiens OX=9606 GN=FMR1 PE=1 SV=1 | 0.966 | 0.996578046 | 0.957 | 0.7 |
| A2AB27 | Guanine nucleotide-binding protein-like 1 (Fragment) OS=Homo sapiens OX=9606 GN=GNL1 PE=1 SV=1 | 1.07 | 0.901934028 | 0.917 | 0.7 |
| Q86X55 | Histone-arginine methyltransferase CARM1 OS=Homo sapiens OX=9606 GN=CARM1 PE=1 SV=3 | 0.907 | 0.750023665 | 0.926 | 0.7 |
| G3V4P8 | Glia maturation factor beta (Fragment) OS=Homo sapiens OX=9606 GN=GMFB PE=1 SV=1 | 0.977 | 0.881623512 | 1.001 | 0.9 |
| P45973 | Chromobox protein homolog 5 OS=Homo sapiens OX=9606 GN=CBX5 PE=1 SV=1 | 0.958 | 0.845656075 | 0.896 | 0.7 |
| P23610 | 40-kDa huntingtin-associated protein OS=Homo sapiens OX=9606 GN=F8A1 PE=1 SV=2 | 1.094 | 0.873607677 | 0.932 | 0.7 |
| Q13630 | GDP-L-fucose synthase OS=Homo sapiens OX=9606 GN=TSTA3 PE=1 SV=1 | 1.01 | 0.766667967 | 1.015 | 0.8 |
| Q9BPZ3 | Polyadenylate-binding protein-interacting protein 2 OS=Homo sapiens OX=9606 GN=PAIP2 PE=1 SV=1 | 0.983 | 0.956825234 | 0.869 | 0.7 |
| M0R0M7 | Syntaxin-binding protein 2 OS=Homo sapiens OX=9606 GN=STXBP2 PE=1 SV=1 | 1.19 | 0.776314646 | 0.974 | 0.7 |
| A0A0B4J2D5 | Glutamine amidotransferase-like class 1 domain-containing protein 3B, mitochondrial OS=Homo sapiens OX=9606 GN=GATD3B P | 1.239 | 0.765235202 | 0.801 | 0.7 |
| O00443 | Phosphatidylinositol 4-phosphate 3-kinase C2 domain-containing subunit alpha OS=Homo sapiens OX=9606 GN=PIK3C2A PE=1 SV | 0.921 | 0.750023665 | 1.044 | 0.9 |
| Q14966 | Zinc finger protein 638 OS=Homo sapiens OX=9606 GN=ZNF638 PE=1 SV=2 | 1.017 | 0.840158726 | 0.884 | 0.7 |
| Q15397 | Pumilio homolog 3 OS=Homo sapiens OX=9606 GN=PUM3 PE=1 SV=3 | 0.918 | 0.996138267 | 0.871 | 0.7 |
| P48059-3 | Isoform 3 of LIM and senescent cell antigen-like-containing domain protein 1 OS=Homo sapiens OX=9606 GN=LIMS1 | 1.039 | 0.849526028 | 0.966 | 0.7 |
| P45954 | Short/branched chain specific acyl-CoA dehydrogenase, mitochondrial OS=Homo sapiens OX=9606 GN=ACADSB PE=1 SV=1 | 1.051 | 0.939024443 | 0.976 | 0.8 |
| Q96D15 | Reticulocalbin-3 OS=Homo sapiens OX=9606 GN=RCN3 PE=1 SV=1 | 0.924 | 0.754126086 | 1.02 | 0.9 |
| D6RFH4 | Cytochrome b5 type B OS=Homo sapiens OX=9606 GN=CYB5B PE=1 SV=2 | 0.976 | 0.889482529 | 0.947 | 0.7 |
| Q9NPY3 | Complement component C1q receptor OS=Homo sapiens OX=9606 GN=CD93 PE=1 SV=3 | 0.98 | 0.931038555 | 0.95 | 0.7 |
| E9PF16 | Medium-chain acyl-CoA ligase ACSF2, mitochondrial OS=Homo sapiens OX=9606 GN=ACSF2 PE=1 SV=1 | 1.097 | 0.77554521 | 0.839 | 0.7 |
| P40222 | Alpha-taxilin OS=Homo sapiens OX=9606 GN=TXLNA PE=1 SV=3 | 1.066 | 0.750023665 | 0.995 | 0.8 |
| A0A3B3IUB9 | Baculoviral IAP repeat-containing protein 6 (Fragment) OS=Homo sapiens OX=9606 GN=BIRC6 PE=1 SV=1 | 1.024 | 0.913279108 | 0.978 | 0.8 |
| Q86TX2 | Acyl-coenzyme A thioesterase 1 OS=Homo sapiens OX=9606 GN=ACOT1 PE=1 SV=1 | 1.011 | 0.96258468 | 0.918 | 0.7 |
| P80303 | Nucleobindin-2 OS=Homo sapiens OX=9606 GN=NUCB2 PE=1 SV=3 | 1.005 | 0.985763982 | 1.008 | 0.9 |
| E9PSH3 | Tetraspanin-4 (Fragment) OS=Homo sapiens OX=9606 GN=TSPAN4 PE=1 SV=1 | 0.928 | 0.785314375 | 0.939 | 0.7 |
| Q6P587 | Acylpyruvase FAHD1, mitochondrial OS=Homo sapiens OX=9606 GN=FAHD1 PE=1 SV=2 | 0.955 | 0.95980123 | 0.888 | 0.7 |
| Q92520 | Protein FAM3C OS=Homo sapiens OX=9606 GN=FAM3C PE=1 SV=1 | 0.927 | 0.765235202 | 0.918 | 0.7 |
| Q9NQ29 | Putative RNA-binding protein Luc7-like 1 OS=Homo sapiens OX=9606 GN=LUC7L PE=1 SV=1 | 0.988 | 0.956825234 | 0.963 | 0.7 |
| Q9UKN8 | General transcription factor 3C polypeptide 4 OS=Homo sapiens OX=9606 GN=GTF3C4 PE=1 SV=2 | 1.163 | 0.837893552 | 0.793 | 0.7 |
| F5H479 | Late endosomal/lysosomal adaptor and MAPK and MTOR activator 1 OS=Homo sapiens OX=9606 GN=LAMTOR1 PE=1 SV=1 | 0.949 | 0.94475926 | 0.908 | 0.7 |
| Q8NBY1 | Serine/threonine-protein kinase 26 OS=Homo sapiens OX=9606 GN=STK26 PE=1 SV=1 | 1.067 | 0.908555656 | 0.775 | 0.7 |
| O15460 | Prolyl 4-hydroxylase subunit alpha-2 OS=Homo sapiens OX=9606 GN=P4HA2 PE=1 SV=1 | 0.992 | 0.978693882 | 0.946 | 0.7 |
| Q13123 | Protein Red OS=Homo sapiens OX=9606 GN=IK PE=1 SV=3 | 1.025 | 0.904500837 | 0.967 | 0.9 |
| Q5T1J5 | Putative coiled-coil-helix-coiled-coil-helix domain-containing protein CHCHD2P9, mitochondrial OS=Homo sapiens OX=9606 GN=( | 1.054 | 0.938907165 | 0.943 | 0.7 |
| Q9Y277 | Voltage-dependent anion-selective channel protein 3 OS=Homo sapiens OX=9606 GN=VDAC3 PE=1 SV=1 | 0.772 | 0.750023665 | 0.741 | 0.7 |
| O60739 | Eukaryotic translation initiation factor 1b OS=Homo sapiens OX=9606 GN=EIF1B PE=1 SV=2 | 1.108 | 0.750023665 | 1.033 | 0.7 |
| O76070 | Gamma-synuclein OS=Homo sapiens OX=9606 GN=SNCG PE=1 SV=2 | 1.007 | 0.953792095 | 0.969 | 0.7 |
| Q9Y2S6 | Translation machinery-associated protein 7 OS=Homo sapiens OX=9606 GN=TMA7 PE=1 SV=1 | 0.997 | 0.901934028 | 1.042 | 0.7 |
| Q13325 | Interferon-induced protein with tetratricopeptide repeats 5 OS=Homo sapiens OX=9606 GN=IFIT5 PE=1 SV=1 | 0.93 | 0.903226585 | 0.898 | 0.7 |
| C9J2Q2 | ATP synthase mitochondrial F1 complex assembly factor 2 (Fragment) OS=Homo sapiens OX=9606 GN=ATPAF2 PE=1 SV=1 | 0.964 | 0.916936955 | 0.927 | 0.7 |
| Q6P9B6 | MTOR-associated protein MEAK7 OS=Homo sapiens OX=9606 GN=MEAK7 PE=1 SV=2 | 0.996 | 0.996578046 | 1.014 | 0.8 |
| Q96II8 | DISP complex protein LRCH3 OS=Homo sapiens OX=9606 GN=LRCH3 PE=1 SV=2 | 1.093 | 0.942843201 | 0.819 | 0.7 |
| E7EM64 | COP9 signalosome complex subunit 6 OS=Homo sapiens OX=9606 GN=COPS6 PE=1 SV=1 | 1.065 | 0.883924281 | 0.871 | 0.7 |
| Q9Y3I1 | F-box only protein 7 OS=Homo sapiens OX=9606 GN=FBXO7 PE=1 SV=1 | 1.03 | 0.985763982 | 1.027 | 0.8 |
| P51668 | Ubiquitin-conjugating enzyme E2 D1 OS=Homo sapiens OX=9606 GN=UBE2D1 PE=1 SV=1 | 0.856 | 0.765235202 | 0.893 | 0.7 |
| C9IZG4 | Protein CutA OS=Homo sapiens OX=9606 GN=CUTA PE=1 SV=1 | 0.937 | 0.750023665 | 0.931 | 0.7 |
| P00374 | Dihydrofolate reductase OS=Homo sapiens OX=9606 GN=DHFR PE=1 SV=2 | 0.928 | 0.771965148 | 0.968 | 0.7 |
| Q9Y5Y2 | Cytosolic Fe-S cluster assembly factor NUBP2 OS=Homo sapiens OX=9606 GN=NUBP2 PE=1 SV=1 | 1.023 | 0.917008272 | 0.95 | 0.8 |
| C9JFA7 | Glycerol-3-phosphate dehydrogenase 1-like protein (Fragment) OS=Homo sapiens OX=9606 GN=GPD1L PE=1 SV=1 | 1.189 | 0.829390484 | 0.906 | 0.7 |
| E5RIA4 | Aspartyl aminopeptidase OS=Homo sapiens OX=9606 GN=DNPEP PE=1 SV=1 | 1.089 | 0.750023665 | 0.848 | 0.7 |
| D6RG18 | Cyclin-H OS=Homo sapiens OX=9606 GN=CCNH PE=1 SV=1 | 0.997 | 0.974919191 | 0.811 | 0.7 |
| Q9NSC5 | Homer protein homolog 3 OS=Homo sapiens OX=9606 GN=HOMER3 PE=1 SV=2 | 0.999 | 0.963059765 | 0.838 | 0.7 |
| Q9Y6Y8 | SEC23-interacting protein OS=Homo sapiens OX=9606 GN=SEC23IP PE=1 SV=1 | 1.009 | 0.97688166 | 0.878 | 0.7 |
| Q7KZ85 | Transcription elongation factor SPT6 OS=Homo sapiens OX=9606 GN=SUPT6H PE=1 SV=2 | 0.911 | 0.908928234 | 0.817 | 0.8 |
| Q8TAQ2 | SWI/SNF complex subunit SMARCC2 OS=Homo sapiens OX=9606 GN=SMARCC2 PE=1 SV=1 | 1.032 | 0.881113553 | 0.898 | 0.7 |
| A6NHR9 | Structural maintenance of chromosomes flexible hinge domain-containing protein 1 OS=Homo sapiens OX=9606 GN=SMCHD1 PE | 0.985 | 0.953828078 | 0.972 | 0.9 |
| A0A1B0GUF2 | SH3 and PX domain-containing protein 2B (Fragment) OS=Homo sapiens OX=9606 GN=SH3PXD2B PE=1 SV=1 | 1.008 | 0.997276249 | 0.911 | 0.7 |
| O75110 | Probable phospholipid-transporting ATPase IIA OS=Homo sapiens OX=9606 GN=ATP9A PE=1 SV=3 | 0.941 | 0.766667967 | 1.155 | 0.8 |
| Q9C0C2 | 182 kDa tankyrase-1-binding protein OS=Homo sapiens OX=9606 GN=TNKS1BP1 PE=1 SV=4 | 1.096 | 0.750023665 | 0.992 | 0.8 |
| A0A0C4DFM1 | Transmembrane 9 superfamily member OS=Homo sapiens OX=9606 GN=TM9SF4 PE=1 SV=1 | 1.006 | 0.917008272 | 0.969 | 0.7 |
| P24539 | ATP synthase F(0) complex subunit B1, mitochondrial OS=Homo sapiens OX=9606 GN=ATP5PB PE=1 SV=2 | 0.951 | 0.95274708 | 0.849 | 0.7 |

| | | | | | |
|---|---|---|---|---|---|
| E9PH82 | Protein FAM98A OS=Homo sapiens OX=9606 GN=FAM98A PE=1 SV=1 | 1.019 | 0.957217311 | 1.012 | 0.9 |
| A0A590UJ50 | Cullin-1 OS=Homo sapiens OX=9606 GN=CUL1 PE=1 SV=1 | 0.995 | 0.895199053 | 0.9 | 0.8 |
| Q9NW15 | Anoctamin-10 OS=Homo sapiens OX=9606 GN=ANO10 PE=1 SV=2 | 0.977 | 0.985324977 | 0.925 | 0.8 |
| Q8IWE2 | Protein NOXP20 OS=Homo sapiens OX=9606 GN=FAM114A1 PE=1 SV=2 | 0.894 | 0.766667967 | 0.976 | 0.8 |
| H0YAC6 | Lipopolysaccharide-responsive and beige-like anchor protein (Fragment) OS=Homo sapiens OX=9606 GN=LRBA PE=1 SV=1 | 0.985 | 0.904500837 | 0.849 | 0.7 |
| Q5T440 | Putative transferase CAF17, mitochondrial OS=Homo sapiens OX=9606 GN=IBA57 PE=1 SV=1 | 1.094 | 0.766667967 | 0.895 | 0.7 |
| B4DIP2 | Erbin OS=Homo sapiens OX=9606 GN=ERBIN PE=1 SV=1 | 1.065 | 0.765235202 | 0.867 | 0.7 |
| O75691 | Small subunit processome component 20 homolog OS=Homo sapiens OX=9606 GN=UTP20 PE=1 SV=3 | 1.166 | 0.904500837 | 0.833 | 0.8 |
| Q9UBN7 | Histone deacetylase 6 OS=Homo sapiens OX=9606 GN=HDAC6 PE=1 SV=2 | 1.096 | 0.973388826 | 0.92 | 0.8 |
| P05783 | Keratin, type I cytoskeletal 18 OS=Homo sapiens OX=9606 GN=KRT18 PE=1 SV=2 | 0.968 | 0.983953845 | 1.14 | 0.7 |
| Q9H446 | RWD domain-containing protein 1 OS=Homo sapiens OX=9606 GN=RWDD1 PE=1 SV=1 | 1.064 | 0.805711413 | 0.981 | 0.7 |
| Q9H490 | Phosphatidylinositol glycan anchor biosynthesis class U protein OS=Homo sapiens OX=9606 GN=PIGU PE=1 SV=3 | 1.091 | 0.983953845 | 0.93 | 0.7 |
| Q86TI2 | Dipeptidyl peptidase 9 OS=Homo sapiens OX=9606 GN=DPP9 PE=1 SV=3 | 1.017 | 0.978477275 | 0.808 | 0.7 |
| M0QZR8 | DNA polymerase OS=Homo sapiens OX=9606 GN=POLD1 PE=1 SV=1 | 1.022 | 0.829390484 | 0.868 | 0.7 |
| O75880 | Protein SCO1 homolog, mitochondrial OS=Homo sapiens OX=9606 GN=SCO1 PE=1 SV=1 | 1.117 | 0.829390484 | 0.655 | 0.7 |
| Q12972 | Nuclear inhibitor of protein phosphatase 1 OS=Homo sapiens OX=9606 GN=PPP1R8 PE=1 SV=2 | 1.089 | 0.840158726 | 1.008 | 0.8 |
| Q9Y237 | Peptidyl-prolyl cis-trans isomerase NIMA-interacting 4 OS=Homo sapiens OX=9606 GN=PIN4 PE=1 SV=1 | 1.039 | 0.765235202 | 0.946 | 0.7 |
| Q15067 | Peroxisomal acyl-coenzyme A oxidase 1 OS=Homo sapiens OX=9606 GN=ACOX1 PE=1 SV=3 | 1.041 | 0.765235202 | 0.959 | 0.7 |
| Q15477 | Helicase SKI2W OS=Homo sapiens OX=9606 GN=SKIV2L PE=1 SV=3 | 0.931 | 0.80706255 | 0.983 | 0.7 |
| P11717 | Cation-independent mannose-6-phosphate receptor OS=Homo sapiens OX=9606 GN=IGF2R PE=1 SV=3 | 0.918 | 0.754250616 | 0.928 | 0.7 |
| Q9BS40 | Latexin OS=Homo sapiens OX=9606 GN=LXN PE=1 SV=2 | 1.015 | 0.994361756 | 0.919 | 0.7 |
| Q3SXM5 | Inactive hydroxysteroid dehydrogenase-like protein 1 OS=Homo sapiens OX=9606 GN=HSDL1 PE=1 SV=3 | 0.84 | 0.765235202 | 0.772 | 0.7 |
| U3KPZ7 | Uncharacterized protein OS=Homo sapiens OX=9606 PE=1 SV=1 | 1.013 | 0.889482529 | 0.799 | 0.7 |
| Q8WZA0 | Protein LZIC OS=Homo sapiens OX=9606 GN=LZIC PE=1 SV=1 | 0.979 | 0.884192327 | 0.958 | 0.7 |
| Q14657 | EKC/KEOPS complex subunit LAGE3 OS=Homo sapiens OX=9606 GN=LAGE3 PE=1 SV=2 | 0.93 | 0.886561233 | 0.937 | 0.9 |
| Q9UJY5 | ADP-ribosylation factor-binding protein GGA1 OS=Homo sapiens OX=9606 GN=GGA1 PE=1 SV=1 | 0.946 | 0.80706255 | 0.847 | 0.7 |
| P31937 | 3-hydroxyisobutyrate dehydrogenase, mitochondrial OS=Homo sapiens OX=9606 GN=HIBADH PE=1 SV=2 | 1.024 | 0.994361756 | 0.919 | 0.7 |
| Q92620 | Pre-mRNA-splicing factor ATP-dependent RNA helicase PRP16 OS=Homo sapiens OX=9606 GN=DHX38 PE=1 SV=2 | 1.026 | 0.984637593 | 1.01 | 0.9 |
| O15397 | Importin-8 OS=Homo sapiens OX=9606 GN=IPO8 PE=1 SV=2 | 0.97 | 0.983953845 | 1.008 | 0.8 |
| Q9H0W9 | Ester hydrolase C11orf54 OS=Homo sapiens OX=9606 GN=C11orf54 PE=1 SV=1 | 0.962 | 0.790136102 | 0.951 | 0.8 |
| Q86W42 | THO complex subunit 6 homolog OS=Homo sapiens OX=9606 GN=THOC6 PE=1 SV=1 | 1.064 | 0.752950049 | 0.933 | 0.7 |
| Q9Y570 | Protein phosphatase methylesterase 1 OS=Homo sapiens OX=9606 GN=PPME1 PE=1 SV=3 | 1.018 | 0.861425366 | 0.957 | 0.9 |
| Q9GZR7 | ATP-dependent RNA helicase DDX24 OS=Homo sapiens OX=9606 GN=DDX24 PE=1 SV=1 | 1.016 | 0.939024443 | 0.848 | 0.7 |
| C9J6A7 | Ribulose-phosphate 3-epimerase (Fragment) OS=Homo sapiens OX=9606 GN=RPE PE=1 SV=1 | 1.072 | 0.765235202 | 0.953 | 0.8 |
| Q9NWY4 | Histone PARylation factor 1 OS=Homo sapiens OX=9606 GN=HPF1 PE=1 SV=2 | 1.006 | 0.905369216 | 0.975 | 0.9 |
| O75155 | Cullin-associated NEDD8-dissociated protein 2 OS=Homo sapiens OX=9606 GN=CAND2 PE=1 SV=3 | 0.74 | 0.750023665 | 0.937 | 0.7 |
| O75489 | NADH dehydrogenase [ubiquinone] iron-sulfur protein 3, mitochondrial OS=Homo sapiens OX=9606 GN=NDUFS3 PE=1 SV=1 | 0.873 | 0.816716243 | 0.937 | 0.7 |
| Q5EG05 | Caspase recruitment domain-containing protein 16 OS=Homo sapiens OX=9606 GN=CARD16 PE=1 SV=1 | 0.961 | 0.882454148 | 0.861 | 0.7 |
| P49585 | Choline-phosphate cytidylyltransferase A OS=Homo sapiens OX=9606 GN=PCYT1A PE=1 SV=2 | 1.006 | 0.981259097 | 0.985 | 0.7 |
| Q9BVL4 | Protein adenylyltransferase SelO, mitochondrial OS=Homo sapiens OX=9606 GN=SELENOO PE=1 SV=3 | 1.251 | 0.766667967 | 0.822 | 0.7 |
| Q2TAA2 | Isoamyl acetate-hydrolyzing esterase 1 homolog OS=Homo sapiens OX=9606 GN=IAH1 PE=1 SV=1 | 0.899 | 0.924992773 | 0.817 | 0.7 |
| Q93100 | Phosphorylase b kinase regulatory subunit beta OS=Homo sapiens OX=9606 GN=PHKB PE=1 SV=3 | 1.052 | 0.905369216 | 0.943 | 0.7 |
| Q13526 | Peptidyl-prolyl cis-trans isomerase NIMA-interacting 1 OS=Homo sapiens OX=9606 GN=PIN1 PE=1 SV=1 | 0.945 | 0.750023665 | 0.943 | 0.7 |
| H3BRM1 | Abscission/NoCut checkpoint regulator (Fragment) OS=Homo sapiens OX=9606 GN=ZFYVE19 PE=1 SV=1 | | | | |
| Q5VZS7 | Programmed cell death protein 4 (Fragment) OS=Homo sapiens OX=9606 GN=PDCD4 PE=1 SV=1 | 0.883 | 0.765235202 | 0.708 | 0.7 |
| O75695 | Protein XRP2 OS=Homo sapiens OX=9606 GN=RP2 PE=1 SV=4 | 0.884 | 0.754126086 | 0.841 | 0.7 |
| Q9UHY1 | Nuclear receptor-binding protein OS=Homo sapiens OX=9606 GN=NRBP1 PE=1 SV=1 | 0.997 | 0.990166412 | 0.942 | 0.7 |
| Q92797 | Symplekin OS=Homo sapiens OX=9606 GN=SYMPK PE=1 SV=2 | 1.014 | 0.860909277 | 0.893 | 0.7 |
| O00178 | GTP-binding protein 1 OS=Homo sapiens OX=9606 GN=GTPBP1 PE=1 SV=3 | 1.008 | 0.921087329 | 0.901 | 0.8 |
| Q92572 | AP-3 complex subunit sigma-1 OS=Homo sapiens OX=9606 GN=AP3S1 PE=1 SV=1 | 0.982 | 0.823903628 | 1.008 | 0.8 |
| Q9BPX3 | Condensin complex subunit 3 OS=Homo sapiens OX=9606 GN=NCAPG PE=1 SV=1 | 0.97 | 0.765235202 | 0.984 | 0.7 |
| I3L1R7 | Ethanolamine-phosphate cytidylyltransferase OS=Homo sapiens OX=9606 GN=PCYT2 PE=1 SV=1 | 0.909 | 0.765235202 | 0.923 | 0.7 |
| O14964 | Hepatocyte growth factor-regulated tyrosine kinase substrate OS=Homo sapiens OX=9606 GN=HGS PE=1 SV=1 | 1.045 | 0.765235202 | 0.97 | 0.8 |
| A0A3B3IUB5 | Minor histocompatibility antigen H13 OS=Homo sapiens OX=9606 GN=HM13 PE=1 SV=1 | 0.951 | 0.816673047 | 1.036 | 0.9 |
| D6REM4 | Casein kinase I isoform alpha OS=Homo sapiens OX=9606 GN=CSNK1A1 PE=1 SV=1 | 1.125 | 0.750023665 | 1.045 | 0.9 |
| Q6DKI1 | 60S ribosomal protein L7-like 1 OS=Homo sapiens OX=9606 GN=RPL7L1 PE=1 SV=2 | 1.069 | 0.750023665 | 0.893 | 0.7 |
| Q4KMQ2 | Anoctamin-6 OS=Homo sapiens OX=9606 GN=ANO6 PE=1 SV=2 | 1.003 | 0.997835552 | 0.885 | 0.7 |
| A0A2R8Y566 | RAB11-binding protein RELCH OS=Homo sapiens OX=9606 GN=RELCH PE=1 SV=1 | 0.997 | 0.913279108 | 0.923 | 0.7 |
| M0R0P8 | Unconventional myosin-IXb OS=Homo sapiens OX=9606 GN=MYO9B PE=1 SV=1 | 1.046 | 0.766667967 | 0.988 | 0.9 |
| Q9C005 | Protein dpy-30 homolog OS=Homo sapiens OX=9606 GN=DPY30 PE=1 SV=1 | 0.997 | 0.95980123 | 0.872 | 0.7 |
| Q9NXR7-1 | Isoform 1 of BRISC and BRCA1-A complex member 2 OS=Homo sapiens OX=9606 GN=BABAM2 | 1.036 | 0.79828785 | 0.887 | 0.7 |
| P84095 | Rho-related GTP-binding protein RhoG OS=Homo sapiens OX=9606 GN=RHOG PE=1 SV=1 | 1.064 | 0.750023665 | 0.942 | 0.7 |
| I3L4X2 | Multidrug resistance-associated protein 1 (Fragment) OS=Homo sapiens OX=9606 GN=ABCC1 PE=1 SV=1 | 0.938 | 0.750023665 | 0.928 | 0.7 |
| F2Z2S2 | Protein furry homolog-like OS=Homo sapiens OX=9606 GN=FRYL PE=1 SV=1 | 1.025 | 0.883217803 | 0.911 | 0.7 |
| Q9Y680-1 | Isoform 1 of Peptidyl-prolyl cis-trans isomerase FKBP7 OS=Homo sapiens OX=9606 GN=FKBP7 | 0.988 | 0.882454148 | 0.913 | 0.7 |
| Q96NY7 | Chloride intracellular channel protein 6 OS=Homo sapiens OX=9606 GN=CLIC6 PE=2 SV=3 | 0.984 | 0.997738183 | 0.912 | 0.7 |
| Q9H0C8 | Integrin-linked kinase-associated serine/threonine phosphatase 2C OS=Homo sapiens OX=9606 GN=ILKAP PE=1 SV=1 | 0.945 | 0.869473756 | 1.061 | 0.8 |
| Q96A33 | Coiled-coil domain-containing protein 47 OS=Homo sapiens OX=9606 GN=CCDC47 PE=1 SV=1 | 1.041 | 0.780884711 | 0.937 | 0.8 |
| P42566 | Epidermal growth factor receptor substrate 15 OS=Homo sapiens OX=9606 GN=EPS15 PE=1 SV=2 | 0.983 | 0.905369216 | 0.861 | 0.7 |
| P40337 | von Hippel-Lindau disease tumor suppressor OS=Homo sapiens OX=9606 GN=VHL PE=1 SV=2 | 1.067 | 0.996138267 | 0.951 | 0.7 |
| P21912 | Succinate dehydrogenase [ubiquinone] iron-sulfur subunit, mitochondrial OS=Homo sapiens OX=9606 GN=SDHB PE=1 SV=3 | 1.042 | 0.905369216 | 0.834 | 0.7 |
| C9J3L8 | Signal sequence receptor subunit alpha OS=Homo sapiens OX=9606 GN=SSR1 PE=1 SV=1 | 1.033 | 0.994361756 | 1.017 | 0.9 |
| O75179 | Ankyrin repeat domain-containing protein 17 OS=Homo sapiens OX=9606 GN=ANKRD17 PE=1 SV=3 | 1.1 | 0.827871327 | 1 | 0.8 |
| Q96GA3 | Protein LTV1 homolog OS=Homo sapiens OX=9606 GN=LTV1 PE=1 SV=1 | 0.982 | 0.928138909 | 0.981 | 0.9 |
| C9JQD4 | Peptidyl-prolyl cis-trans isomerase (Fragment) OS=Homo sapiens OX=9606 GN=PPIH PE=1 SV=1 | 0.933 | 0.756287988 | 0.96 | 0.7 |
| Q8TEQ6 | Gem-associated protein 5 OS=Homo sapiens OX=9606 GN=GEMIN5 PE=1 SV=3 | 1.003 | 0.989102644 | 1.036 | 0.8 |
| P07902 | Galactose-1-phosphate uridylyltransferase OS=Homo sapiens OX=9606 GN=GALT PE=1 SV=3 | 0.893 | 0.882454148 | 0.79 | 0.7 |
| G5E9D5 | ElaC homolog protein 2 OS=Homo sapiens OX=9606 GN=ELAC2 PE=1 SV=1 | 1.002 | 0.961750547 | 0.933 | 0.7 |
| P48507 | Glutamate--cysteine ligase regulatory subunit OS=Homo sapiens OX=9606 GN=GCLM PE=1 SV=1 | 0.968 | 0.916363276 | 0.992 | 0.7 |
| Q9UBU9 | Nuclear RNA export factor 1 OS=Homo sapiens OX=9606 GN=NXF1 PE=1 SV=1 | 1.008 | 0.96647353 | 0.924 | 0.7 |
| O95456 | Proteasome assembly chaperone 1 OS=Homo sapiens OX=9606 GN=PSMG1 PE=1 SV=1 | 0.839 | 0.750023665 | 0.914 | 0.7 |
| Q96EH3 | Mitochondrial assembly of ribosomal large subunit protein 1 OS=Homo sapiens OX=9606 GN=MALSU1 PE=1 SV=1 | 0.867 | 0.824559577 | 0.815 | 0.7 |
| A0A087X211 | Protein CIP2A OS=Homo sapiens OX=9606 GN=CIP2A PE=1 SV=1 | 1.071 | 0.750023665 | 1.493 | 0.7 |
| O00743 | Serine/threonine-protein phosphatase 6 catalytic subunit OS=Homo sapiens OX=9606 GN=PPP6C PE=1 SV=1 | 1.033 | 0.866300162 | 0.984 | 0.8 |
| E9PHS0 | Glutathione S-transferase LANCL1 (Fragment) OS=Homo sapiens OX=9606 GN=LANCL1 PE=1 SV=1 | 0.899 | 0.829390484 | 0.954 | 0.7 |
| O95831 | Apoptosis-inducing factor 1, mitochondrial OS=Homo sapiens OX=9606 GN=AIFM1 PE=1 SV=1 | 1.002 | 0.990981579 | 0.952 | 0.7 |
| H0Y650 | Dynein assembly factor 5, axonemal (Fragment) OS=Homo sapiens OX=9606 GN=DNAAF5 PE=1 SV=1 | 0.969 | 0.95980123 | 0.98 | 0.7 |
| P25685 | DnaJ homolog subfamily B member 1 OS=Homo sapiens OX=9606 GN=DNAJB1 PE=1 SV=4 | 1.066 | 0.845356491 | 0.921 | 0.7 |
| Q9BSL1 | Ubiquitin-associated domain-containing protein 1 OS=Homo sapiens OX=9606 GN=UBAC1 PE=1 SV=1 | 1.022 | 0.921641483 | 1.03 | 0.9 |
| Q9UEW8 | STE20/SPS1-related proline-alanine-rich protein kinase OS=Homo sapiens OX=9606 GN=STK39 PE=1 SV=3 | 0.912 | 0.876510992 | 0.862 | 0.7 |
| P62306 | Small nuclear ribonucleoprotein F OS=Homo sapiens OX=9606 GN=SNRPF PE=1 SV=1 | 1.056 | 0.902016292 | 1.028 | 0.8 |
| O75251 | NADH dehydrogenase [ubiquinone] iron-sulfur protein 7, mitochondrial OS=Homo sapiens OX=9606 GN=NDUFS7 PE=1 SV=3 | 0.938 | 0.994361756 | 0.907 | 0.7 |
| P10620 | Microsomal glutathione S-transferase 1 OS=Homo sapiens OX=9606 GN=MGST1 PE=1 SV=1 | 0.925 | 0.876510992 | 1.04 | 0.8 |
| A0A087X2B3 | Neuronal cell adhesion molecule OS=Homo sapiens OX=9606 GN=NRCAM PE=1 SV=1 | 1.013 | 0.886561233 | 1.018 | 0.8 |
| G3V445 | N6-L-threonylcarbamoyladenine synthase (Fragment) OS=Homo sapiens OX=9606 GN=OSGEP PE=1 SV=8 | 1.067 | 0.927148669 | 0.826 | 0.7 |
| Q15582 | Transforming growth factor-beta-induced protein ig-h3 OS=Homo sapiens OX=9606 GN=TGFBI PE=1 SV=1 | 1.008 | 0.996138267 | 0.759 | 0.7 |

| | | | | | |
|---|---|---|---|---|---|
| Q6NXT2 | Histone H3.3C OS=Homo sapiens OX=9606 GN=H3-5 PE=1 SV=3 | 0.649 | 0.765235202 | 0.827 | 0.7 |
| Q96L92 | Sorting nexin-27 OS=Homo sapiens OX=9606 GN=SNX27 PE=1 SV=2 | 1.024 | 0.782074241 | 0.918 | 0.7 |
| O43294 | Transforming growth factor beta-1-induced transcript 1 protein OS=Homo sapiens OX=9606 GN=TGFB1I1 PE=1 SV=2 | 1.05 | 0.994361756 | 0.907 | 0.7 |
| R4GMW6 | COMMD3-BMI1 readthrough (Fragment) OS=Homo sapiens OX=9606 GN=COMMD3-BMI1 PE=4 SV=1 | 0.987 | 0.996637279 | 0.898 | 0.7 |
| Q96N66 | Lysophospholipid acyltransferase 7 OS=Homo sapiens OX=9606 GN=MBOAT7 PE=1 SV=2 | 1.002 | 0.921087329 | 0.866 | 0.7 |
| P30038 | Delta-1-pyrroline-5-carboxylate dehydrogenase, mitochondrial OS=Homo sapiens OX=9606 GN=ALDH4A1 PE=1 SV=3 | 1.031 | 0.956825234 | 0.93 | 0.7 |
| O14562 | Ubiquitin domain-containing protein UBFD1 OS=Homo sapiens OX=9606 GN=UBFD1 PE=1 SV=2 | 1.026 | 0.953792095 | 0.925 | 0.7 |
| O15173 | Membrane-associated progesterone receptor component 2 OS=Homo sapiens OX=9606 GN=PGRMC2 PE=1 SV=1 | 1.027 | 0.964351664 | 1.008 | 0.8 |
| Q02880 | DNA topoisomerase 2-beta OS=Homo sapiens OX=9606 GN=TOP2B PE=1 SV=3 | 0.955 | 0.886561233 | 0.948 | 0.9 |
| Q9BQ04 | RNA-binding protein 4B OS=Homo sapiens OX=9606 GN=RBM4B PE=1 SV=1 | 1.042 | 0.766667967 | 0.969 | 0.8 |
| Q14690 | Protein RRP5 homolog OS=Homo sapiens OX=9606 GN=PDCD11 PE=1 SV=3 | 1.062 | 0.750023665 | 0.936 | 0.8 |
| Q99567 | Nuclear pore complex protein Nup88 OS=Homo sapiens OX=9606 GN=NUP88 PE=1 SV=2 | 1.055 | 0.779783772 | 1.006 | 0.7 |
| Q7LBR1 | Charged multivesicular body protein 1b OS=Homo sapiens OX=9606 GN=CHMP1B PE=1 SV=1 | 1.014 | 0.929783322 | 0.903 | 0.7 |
| Q9BUQ8 | Probable ATP-dependent RNA helicase DDX23 OS=Homo sapiens OX=9606 GN=DDX23 PE=1 SV=3 | 1.038 | 0.87017509 | 0.923 | 0.7 |
| Q9BZQ8 | Protein Niban 1 OS=Homo sapiens OX=9606 GN=NIBAN1 PE=1 SV=1 | 1.041 | 0.845656075 | 0.972 | 0.7 |
| A0A3B3ISK9 | Malonate--CoA ligase ACSF3, mitochondrial OS=Homo sapiens OX=9606 GN=ACSF3 PE=1 SV=1 | 1.089 | 0.902016292 | 0.908 | 0.7 |
| Q9BZE4 | Nucleolar GTP-binding protein 1 OS=Homo sapiens OX=9606 GN=GTPBP4 PE=1 SV=3 | 1.221 | 0.750023665 | 1.084 | 0.7 |
| H0YLH3 | Protein geranylgeranyltransferase type II OS=Homo sapiens OX=9606 GN=RABGGTA PE=1 SV=1 | 1.132 | 0.869995963 | 1.025 | 0.9 |
| B5MDL5 | Mitogen-activated protein kinase OS=Homo sapiens OX=9606 GN=MAPK12 PE=1 SV=1 | 0.868 | 0.750023665 | 0.865 | 0.7 |
| Q9HCC0 | Methylcrotonoyl-CoA carboxylase beta chain, mitochondrial OS=Homo sapiens OX=9606 GN=MCCC2 PE=1 SV=1 | 0.937 | 0.889482529 | 0.869 | 0.7 |
| Q6ZXV5 | Protein O-mannosyl-transferase TMTC3 OS=Homo sapiens OX=9606 GN=TMTC3 PE=1 SV=2 | 1.021 | 0.997835552 | 0.884 | 0.7 |
| B1AKR6 | Dynein light chain roadblock-type 1 OS=Homo sapiens OX=9606 GN=DYNLRB1 PE=1 SV=1 | 0.913 | 0.790136102 | 0.89 | 0.7 |
| Q9UQE7 | Structural maintenance of chromosomes protein 3 OS=Homo sapiens OX=9606 GN=SMC3 PE=1 SV=2 | 1.004 | 0.996637279 | 0.899 | 0.7 |
| P49770 | Translation initiation factor eIF-2B subunit beta OS=Homo sapiens OX=9606 GN=EIF2B2 PE=1 SV=3 | 1.075 | 0.781370192 | 0.923 | 0.7 |
| Q13637 | Ras-related protein Rab-32 OS=Homo sapiens OX=9606 GN=RAB32 PE=1 SV=3 | 0.965 | 0.703015003 | 1.036 | 0.7 |
| Q8NHH9 | Atlastin-2 OS=Homo sapiens OX=9606 GN=ATL2 PE=1 SV=2 | 0.953 | 0.902016292 | 0.979 | 0.9 |
| O75934 | Pre-mRNA-splicing factor SPF27 OS=Homo sapiens OX=9606 GN=BCAS2 PE=1 SV=1 | 1.089 | 0.750023665 | 0.924 | 0.7 |
| Q9Y5Q0 | Fatty acid desaturase 3 OS=Homo sapiens OX=9606 GN=FADS3 PE=1 SV=1 | 1.048 | 0.927756817 | 0.831 | 0.7 |
| O15498 | Synaptobrevin homolog YKT6 OS=Homo sapiens OX=9606 GN=YKT6 PE=1 SV=1 | 0.996 | 0.884192327 | 1.014 | 0.7 |
| Q8WVJ2 | NudC domain-containing protein 2 OS=Homo sapiens OX=9606 GN=NUDCD2 PE=1 SV=1 | 1.028 | 0.889450577 | 0.939 | 0.7 |
| Q969T4 | Ubiquitin-conjugating enzyme E2 E3 OS=Homo sapiens OX=9606 GN=UBE2E3 PE=1 SV=1 | 0.973 | 0.917134289 | 0.838 | 0.7 |
| Q8NB90 | ATPase family protein 2 homolog OS=Homo sapiens OX=9606 GN=SPATA5 PE=1 SV=3 | 1.146 | 0.750023665 | 1.062 | 0.7 |
| O43464 | Serine protease HTRA2, mitochondrial OS=Homo sapiens OX=9606 GN=HTRA2 PE=1 SV=2 | 0.888 | 0.829390484 | 0.897 | 0.7 |
| Q96JG6 | Syndetin OS=Homo sapiens OX=9606 GN=VPS50 PE=1 SV=3 | 0.953 | 0.834946606 | 0.997 | 0.8 |
| Q9NWV4 | CXXC motif containing zinc binding protein OS=Homo sapiens OX=9606 GN=CZIB PE=1 SV=1 | 1.106 | 0.752950049 | 1.029 | 0.7 |
| Q96H79 | Zinc finger CCCH-type antiviral protein 1-like OS=Homo sapiens OX=9606 GN=ZC3HAV1L PE=1 SV=2 | | | | |
| E7ER68 | Protein FAM91A1 OS=Homo sapiens OX=9606 GN=FAM91A1 PE=1 SV=1 | 1.025 | 0.9962822 | 0.954 | 0.8 |
| A0A0A0MRG2 | ABPP OS=Homo sapiens OX=9606 GN=APP PE=1 SV=1 | 0.992 | 0.889482529 | 1.055 | 0.7 |
| Q07812 | Apoptosis regulator BAX OS=Homo sapiens OX=9606 GN=BAX PE=1 SV=1 | 0.934 | 0.758551152 | 0.981 | 0.7 |
| O95905 | Protein ecdysoneless homolog OS=Homo sapiens OX=9606 GN=ECD PE=1 SV=1 | 0.989 | 0.984925478 | 0.986 | 0.8 |
| P21810 | Biglycan OS=Homo sapiens OX=9606 GN=BGN PE=1 SV=2 | | | | |
| Q15811 | Intersectin-1 OS=Homo sapiens OX=9606 GN=ITSN1 PE=1 SV=3 | 0.99 | 0.985324977 | 1.092 | 0.7 |
| E7EQ69 | N-alpha-acetyltransferase 50 OS=Homo sapiens OX=9606 GN=NAA50 PE=1 SV=1 | 1.017 | 0.902373639 | 1.006 | 0.8 |
| M0QXD6 | Transcription initiation factor IIF subunit alpha (Fragment) OS=Homo sapiens OX=9606 GN=GTF2F1 PE=1 SV=1 | 1.005 | 0.996637279 | 0.999 | 0.9 |
| O75051 | Plexin-A2 OS=Homo sapiens OX=9606 GN=PLXNA2 PE=1 SV=4 | 1.042 | 0.878541488 | 1 | 0.9 |
| Q92947 | Glutaryl-CoA dehydrogenase, mitochondrial OS=Homo sapiens OX=9606 GN=GCDH PE=1 SV=1 | 0.987 | 0.985733613 | 1.007 | 0.9 |
| Q9Y3E7 | Charged multivesicular body protein 3 OS=Homo sapiens OX=9606 GN=CHMP3 PE=1 SV=3 | 0.907 | 0.750023665 | 1.043 | 0.8 |
| E9PQN9 | Interferon-induced transmembrane protein 2 OS=Homo sapiens OX=9606 GN=IFITM2 PE=1 SV=1 | 1.015 | 0.942678074 | 1.06 | 0.9 |
| Q9Y3D5 | 28S ribosomal protein S18c, mitochondrial OS=Homo sapiens OX=9606 GN=MRPS18C PE=1 SV=1 | 0.977 | 0.957217311 | 0.998 | 0.9 |
| Q8N201 | Integrator complex subunit 1 OS=Homo sapiens OX=9606 GN=INTS1 PE=1 SV=2 | 1.244 | 0.829390484 | 0.789 | 0.9 |
| Q15126 | Phosphomevalonate kinase OS=Homo sapiens OX=9606 GN=PMVK PE=1 SV=3 | 1.054 | 0.889450577 | 0.953 | 0.7 |
| A0A075B7C8 | SLIT-ROBO Rho GTPase-activating protein 2 (Fragment) OS=Homo sapiens OX=9606 GN=SRGAP2 PE=4 SV=1 | 0.952 | 0.765235202 | 1.05 | 0.9 |
| X6R390 | AP complex subunit sigma OS=Homo sapiens OX=9606 GN=AP2S1 PE=1 SV=1 | 0.94 | 0.779783772 | 0.922 | 0.7 |
| Q9Y6M7 | Sodium bicarbonate cotransporter 3 OS=Homo sapiens OX=9606 GN=SLC4A7 PE=1 SV=2 | 0.995 | 0.978562349 | 1.052 | 0.9 |
| P61764 | Syntaxin-binding protein 1 OS=Homo sapiens OX=9606 GN=STXBP1 PE=1 SV=1 | 1.009 | 0.973388826 | 0.959 | 0.8 |
| Q9Y399 | 28S ribosomal protein S2, mitochondrial OS=Homo sapiens OX=9606 GN=MRPS2 PE=1 SV=1 | 0.869 | 0.774484855 | 0.828 | 0.7 |
| Q16134 | Electron transfer flavoprotein-ubiquinone oxidoreductase, mitochondrial OS=Homo sapiens OX=9606 GN=ETFDH PE=1 SV=2 | 1.037 | 0.829390484 | 0.951 | 0.7 |
| Q9H1E3 | Nuclear ubiquitous casein and cyclin-dependent kinase substrate 1 OS=Homo sapiens OX=9606 GN=NUCKS1 PE=1 SV=1 | 1.009 | 0.983953845 | 0.982 | 0.7 |
| Q9BYD6 | 39S ribosomal protein L1, mitochondrial OS=Homo sapiens OX=9606 GN=MRPL1 PE=1 SV=2 | 1.062 | 0.809976184 | 0.938 | 0.7 |
| F8W954 | Negative elongation factor A OS=Homo sapiens OX=9606 GN=NELFA PE=1 SV=1 | 0.939 | 0.995582922 | 0.921 | 0.7 |
| F5GYQ1 | V-type proton ATPase subunit OS=Homo sapiens OX=9606 GN=ATP6V0D1 PE=1 SV=1 | 1.022 | 0.902016292 | 0.944 | 0.7 |
| Q00765 | Receptor expression-enhancing protein 5 OS=Homo sapiens OX=9606 GN=REEP5 PE=1 SV=3 | 0.974 | 0.886561233 | 1.038 | 0.7 |
| A0A0C4DGL3 | Deoxyuridine 5'-triphosphate nucleotidohydrolase OS=Homo sapiens OX=9606 GN=DUT PE=1 SV=1 | 0.973 | 0.916363276 | 1.05 | 0.7 |
| Q70UQ0 | Inhibitor of nuclear factor kappa-B kinase-interacting protein OS=Homo sapiens OX=9606 GN=IKBIP PE=1 SV=1 | 1.111 | 0.778727018 | 0.859 | 0.7 |
| E9PNS3 | Toll-interacting protein OS=Homo sapiens OX=9606 GN=TOLLIP PE=1 SV=1 | 1.011 | 0.956825234 | 0.931 | 0.7 |
| F8WFC6 | Large subunit GTPase 1 homolog OS=Homo sapiens OX=9606 GN=LSG1 PE=1 SV=1 | 0.907 | 0.80706255 | 1.121 | 0.9 |
| Q9Y4K0 | Lysyl oxidase homolog 2 OS=Homo sapiens OX=9606 GN=LOXL2 PE=1 SV=1 | 0.91 | 0.889450577 | 0.9 | 0.7 |
| F2Z2E2 | Ras GTPase-activating-like protein IQGAP3 OS=Homo sapiens OX=9606 GN=IQGAP3 PE=1 SV=1 | 0.906 | 0.902597392 | 0.944 | 0.7 |
| P02792 | Ferritin light chain OS=Homo sapiens OX=9606 GN=FTL PE=1 SV=2 | 1.023 | 0.879425665 | 0.911 | 0.7 |
| Q8WVV9 | Heterogeneous nuclear ribonucleoprotein L-like OS=Homo sapiens OX=9606 GN=HNRNPLL PE=1 SV=1 | 0.987 | 0.978562349 | 0.975 | 0.7 |
| M0QXN5 | Nuclear pore glycoprotein p62 OS=Homo sapiens OX=9606 GN=NUP62 PE=1 SV=1 | 0.999 | 0.966113364 | 0.996 | 0.9 |
| Q7Z460 | CLIP-associating protein 1 OS=Homo sapiens OX=9606 GN=CLASP1 PE=1 SV=1 | 1.033 | 0.765235202 | 0.906 | 0.7 |
| A0A494C0R8 | Clustered mitochondria protein homolog OS=Homo sapiens OX=9606 GN=CLUH PE=1 SV=1 | 0.945 | 0.892311206 | 1.055 | 0.9 |
| Q9Y2H6 | Fibronectin type-III domain-containing protein 3A OS=Homo sapiens OX=9606 GN=FNDC3A PE=1 SV=4 | 0.944 | 0.80706255 | 0.962 | 0.7 |
| A0A0B4J1V8 | HCG2039996 OS=Homo sapiens OX=9606 GN=PPAN-P2RY11 PE=3 SV=1 | 1.128 | 0.908555656 | 0.926 | 0.9 |
| Q9BSJ2 | Gamma-tubulin complex component 2 OS=Homo sapiens OX=9606 GN=TUBGCP2 PE=1 SV=2 | 1.022 | 0.902777383 | 0.922 | 0.7 |
| Q53T59 | HCLS1-binding protein 3 OS=Homo sapiens OX=9606 GN=HS1BP3 PE=1 SV=1 | 1.018 | 0.908696432 | 0.935 | 0.7 |
| P98175 | RNA-binding protein 10 OS=Homo sapiens OX=9606 GN=RBM10 PE=1 SV=3 | 1.177 | 0.865970839 | 0.973 | 0.7 |
| A0A087WUD3 | Oligosaccharyltransferase complex subunit OS=Homo sapiens OX=9606 GN=OSTC PE=1 SV=1 | 1.024 | 0.949934729 | 0.871 | 0.7 |
| E9PE17 | 28S ribosomal protein S17, mitochondrial (Fragment) OS=Homo sapiens OX=9606 GN=MRPS17 PE=1 SV=1 | 1.031 | 0.818672523 | 0.963 | 0.7 |
| C9J4M6 | DNA-directed RNA polymerase subunit beta OS=Homo sapiens OX=9606 GN=POLR2B PE=1 SV=3 | 0.953 | 0.913279108 | 0.915 | 0.7 |
| X6R3N0 | Solute carrier family 27 (Fatty acid transporter), member 3, isoform CRA_d OS=Homo sapiens OX=9606 GN=SLC27A3 PE=1 SV=1 | 1.086 | 0.996578046 | 0.984 | 0.9 |
| Q5R3I4 | Tetratricopeptide repeat protein 38 OS=Homo sapiens OX=9606 GN=TTC38 PE=1 SV=1 | 1.018 | 0.914316836 | 0.882 | 0.7 |
| A0A3B3IUC0 | Integral membrane protein 2 OS=Homo sapiens OX=9606 GN=ITM2B PE=1 SV=1 | 1.061 | 0.901934028 | 0.964 | 0.7 |
| Q9Y496 | Kinesin-like protein KIF3A OS=Homo sapiens OX=9606 GN=KIF3A PE=1 SV=4 | 1.071 | 0.908555656 | 0.75 | 0.7 |
| A0A0J9YXE1 | Protein canopy homolog 4 (Fragment) OS=Homo sapiens OX=9606 GN=CNPY4 PE=1 SV=1 | 0.967 | 0.91999213 | 0.864 | 0.7 |
| Q14156 | Protein EFR3 homolog A OS=Homo sapiens OX=9606 GN=EFR3A PE=1 SV=2 | 1.063 | 0.889450577 | 0.878 | 0.7 |
| P63218 | Guanine nucleotide-binding protein G(I)/G(S)/G(O) subunit gamma-5 OS=Homo sapiens OX=9606 GN=GNG5 PE=1 SV=3 | 0.989 | 0.905369216 | 0.969 | 0.7 |
| Q8IWE4 | DCN1-like protein 3 OS=Homo sapiens OX=9606 GN=DCUN1D3 PE=1 SV=1 | 1.002 | 0.963059765 | 0.918 | 0.7 |
| Q9H1B7 | Probable E3 ubiquitin-protein ligase IRF2BPL OS=Homo sapiens OX=9606 GN=IRF2BPL PE=1 SV=1 | 1.06 | 0.886561233 | 0.998 | 0.9 |
| Q9HC36 | rRNA methyltransferase 3, mitochondrial OS=Homo sapiens OX=9606 GN=MRM3 PE=1 SV=2 | 0.982 | 0.978562349 | 0.853 | 0.7 |
| Q9NZR1 | Tropomodulin-2 OS=Homo sapiens OX=9606 GN=TMOD2 PE=1 SV=1 | 1.048 | 0.997835552 | 0.9 | 0.9 |
| J3KMZ7 | Integrator complex subunit 2 OS=Homo sapiens OX=9606 GN=INTS2 PE=1 SV=1 | 0.943 | 0.976558546 | 1.011 | 0.7 |
| O14657 | Torsin-1B OS=Homo sapiens OX=9606 GN=TOR1B PE=1 SV=2 | 0.959 | 0.944419315 | 0.908 | 0.7 |
| Q8WTT2 | Nucleolar complex protein 3 homolog OS=Homo sapiens OX=9606 GN=NOC3L PE=1 SV=1 | 0.964 | 0.642276967 | 0.93 | 0.7 |
| E9PNS0 | Tubulin-specific chaperone cofactor E-like protein (Fragment) OS=Homo sapiens OX=9606 GN=TBCEL PE=1 SV=1 | 0.993 | 0.900915273 | 0.918 | 0.7 |

| | | | | | |
|---|---|---|---|---|---|
| P06493 | Cyclin-dependent kinase 1 OS=Homo sapiens OX=9606 GN=CDK1 PE=1 SV=3 | 1.054 | 0.765235202 | 1.18 | 0.8 |
| Q9UNN5 | FAS-associated factor 1 OS=Homo sapiens OX=9606 GN=FAF1 PE=1 SV=2 | 1.021 | 0.920144076 | 0.968 | 0.8 |
| O60869 | Endothelial differentiation-related factor 1 OS=Homo sapiens OX=9606 GN=EDF1 PE=1 SV=1 | 1.123 | 0.765235202 | 0.958 | 0.7 |
| Q96DI7 | U5 small nuclear ribonucleoprotein 40 kDa protein OS=Homo sapiens OX=9606 GN=SNRNP40 PE=1 SV=1 | 1.072 | 0.902016292 | 0.95 | 0.7 |
| F5H345 | Hydroxymethylbilane synthase OS=Homo sapiens OX=9606 GN=HMBS PE=1 SV=1 | 0.856 | 0.886561233 | 0.942 | 0.7 |
| Q14694 | Ubiquitin carboxyl-terminal hydrolase 10 OS=Homo sapiens OX=9606 GN=USP10 PE=1 SV=2 | 0.986 | 0.886561233 | 1.07 | 0.7 |
| Q9P0V3 | SH3 domain-binding protein 4 OS=Homo sapiens OX=9606 GN=SH3BP4 PE=1 SV=1 | | | | |
| O14974 | Protein phosphatase 1 regulatory subunit 12A OS=Homo sapiens OX=9606 GN=PPP1R12A PE=1 SV=1 | 1.096 | 0.750023665 | 0.877 | 0.7 |
| Q7Z4V5 | Hepatoma-derived growth factor-related protein 2 OS=Homo sapiens OX=9606 GN=HDGFL2 PE=1 SV=1 | 0.947 | 0.750023665 | 0.985 | 0.7 |
| Q9UBS4 | DnaJ homolog subfamily B member 11 OS=Homo sapiens OX=9606 GN=DNAJB11 PE=1 SV=1 | 0.994 | 0.978562349 | 1.097 | 0.7 |
| Q9Y3C8 | Ubiquitin-fold modifier-conjugating enzyme 1 OS=Homo sapiens OX=9606 GN=UFC1 PE=1 SV=3 | 0.973 | 0.765235202 | 0.985 | 0.8 |
| G3V4K3 | Spermatogenesis-defective protein 39 homolog OS=Homo sapiens OX=9606 GN=VIPAS39 PE=1 SV=1 | 0.904 | 0.750023665 | 1.03 | 0.8 |
| P46087 | Probable 28S rRNA (cytosine(4447)-C(5))-methyltransferase OS=Homo sapiens OX=9606 GN=NOP2 PE=1 SV=2 | 1.007 | 0.902016292 | 0.97 | 0.7 |
| C9JG87 | 39S ribosomal protein L39, mitochondrial (Fragment) OS=Homo sapiens OX=9606 GN=MRPL39 PE=1 SV=1 | 0.993 | 0.95980123 | 1.105 | 0.7 |
| P20936 | Ras GTPase-activating protein 1 OS=Homo sapiens OX=9606 GN=RASA1 PE=1 SV=1 | 0.992 | 0.968359197 | 1.003 | 0.7 |
| Q9HB90 | Ras-related GTP-binding protein C OS=Homo sapiens OX=9606 GN=RRAGC PE=1 SV=1 | 0.965 | 0.765235202 | 0.841 | 0.7 |
| P14406 | Cytochrome c oxidase subunit 7A2, mitochondrial OS=Homo sapiens OX=9606 GN=COX7A2 PE=1 SV=1 | 0.932 | 0.883924281 | 0.754 | 0.7 |
| Q99459 | Cell division cycle 5-like protein OS=Homo sapiens OX=9606 GN=CDC5L PE=1 SV=2 | 1.029 | 0.771520178 | 1.008 | 0.8 |
| H3BUF6 | Ataxin-2-like protein OS=Homo sapiens OX=9606 GN=ATXN2L PE=1 SV=1 | 1.06 | 0.984805107 | 1.044 | 0.8 |
| Q15276 | Rab GTPase-binding effector protein 1 OS=Homo sapiens OX=9606 GN=RABEP1 PE=1 SV=2 | 0.884 | 0.750023665 | 0.995 | 0.9 |
| Q9BYD1 | 39S ribosomal protein L13, mitochondrial OS=Homo sapiens OX=9606 GN=MRPL13 PE=1 SV=1 | 1.053 | 0.807858213 | 0.945 | 0.7 |
| Q04828 | Aldo-keto reductase family 1 member C1 OS=Homo sapiens OX=9606 GN=AKR1C1 PE=1 SV=1 | 1.024 | 0.889450577 | 1.021 | 0.9 |
| Q5T447 | E3 ubiquitin-protein ligase HECTD3 OS=Homo sapiens OX=9606 GN=HECTD3 PE=1 SV=1 | 1.029 | 0.957493616 | 0.99 | 0.9 |
| Q5RGS4 | Pleckstrin homology domain-containing family A member 1 OS=Homo sapiens OX=9606 GN=PLEKHA1 PE=1 SV=1 | 1.011 | 0.994361756 | 0.929 | 0.7 |
| P61927 | 60S ribosomal protein L37 OS=Homo sapiens OX=9606 GN=RPL37 PE=1 SV=2 | 0.997 | 0.945523581 | 1.224 | 0.8 |
| Q6UXN9 | WD repeat-containing protein 82 OS=Homo sapiens OX=9606 GN=WDR82 PE=1 SV=1 | 1.043 | 0.798032881 | 0.912 | 0.7 |
| H3BV55 | Phosphomannomutase OS=Homo sapiens OX=9606 GN=PMM2 PE=1 SV=1 | 0.951 | 0.765235202 | 1.052 | 0.8 |
| Q32MZ4-4 | Isoform 4 of Leucine-rich repeat flightless-interacting protein 1 OS=Homo sapiens OX=9606 GN=LRRFIP1 | 0.659 | 0.750023665 | 0.861 | 0.7 |
| A0A087WTD7 | A-kinase anchor protein 13 OS=Homo sapiens OX=9606 GN=AKAP13 PE=1 SV=1 | 0.836 | 0.876510992 | 0.819 | 0.7 |
| P48449 | Lanosterol synthase OS=Homo sapiens OX=9606 GN=LSS PE=1 SV=1 | 1.059 | 0.954224541 | 0.912 | 0.7 |
| B7ZC39 | Endophilin-B2 OS=Homo sapiens OX=9606 GN=SH3GLB2 PE=1 SV=1 | 0.99 | 0.985324977 | 1.009 | 0.7 |
| P21953 | 2-oxoisovalerate dehydrogenase subunit beta, mitochondrial OS=Homo sapiens OX=9606 GN=BCKDHB PE=1 SV=2 | 0.948 | 0.750023665 | 0.908 | 0.7 |
| Q8NGA1 | Olfactory receptor 1M1 OS=Homo sapiens OX=9606 GN=OR1M1 PE=2 SV=1 | 0.921 | 0.750023665 | 0.946 | 0.7 |
| P80723 | Brain acid soluble protein 1 OS=Homo sapiens OX=9606 GN=BASP1 PE=1 SV=2 | 0.987 | 0.842765836 | 0.931 | 0.7 |
| X6RI37 | Leucine-rich repeat protein SHOC-2 (Fragment) OS=Homo sapiens OX=9606 GN=SHOC2 PE=1 SV=1 | 1.009 | 0.995582922 | 0.855 | 0.[illegible] |
| H0YJA0 | Putative E3 ubiquitin-protein ligase UBR7 (Fragment) OS=Homo sapiens OX=9606 GN=UBR7 PE=1 SV=1 | 0.957 | 0.957217311 | 0.914 | 0.7 |
| P36551 | Oxygen-dependent coproporphyrinogen-III oxidase, mitochondrial OS=Homo sapiens OX=9606 GN=CPOX PE=1 SV=3 | 1.094 | 0.750023665 | 0.97 | 0.7 |
| Q13144 | Translation initiation factor eIF-2B subunit epsilon OS=Homo sapiens OX=9606 GN=EIF2B5 PE=1 SV=3 | 0.974 | 0.980966594 | 0.969 | 0.8 |
| A8MTY9 | Vacuolar protein sorting-associated protein 26C OS=Homo sapiens OX=9606 GN=VPS26C PE=1 SV=1 | 0.969 | 0.924631778 | 0.928 | 0.7 |
| P53801 | Pituitary tumor-transforming gene 1 protein-interacting protein OS=Homo sapiens OX=9606 GN=PTTG1IP PE=1 SV=1 | 0.987 | 0.905369216 | 0.956 | 0.7 |
| A0A087WY55 | Chromosome 6 open reading frame 55, isoform CRA_b OS=Homo sapiens OX=9606 GN=VTA1 PE=1 SV=1 | 1.031 | 0.984805107 | 1.071 | 0.7 |
| Q9H9Y4 | GPN-loop GTPase 2 OS=Homo sapiens OX=9606 GN=GPN2 PE=1 SV=2 | 0.973 | 0.990981579 | 0.888 | 0.7 |
| P40189 | Interleukin-6 receptor subunit beta OS=Homo sapiens OX=9606 GN=IL6ST PE=1 SV=2 | 1.115 | 0.750023665 | 1.112 | 0.7 |
| E9PRZ1 | Protein SAAL1 OS=Homo sapiens OX=9606 GN=SAAL1 PE=1 SV=1 | 0.804 | 0.750023665 | 0.898 | 0.7 |
| Q9Y2V7 | Conserved oligomeric Golgi complex subunit 6 OS=Homo sapiens OX=9606 GN=COG6 PE=1 SV=2 | 0.987 | 0.992258896 | 0.942 | 0.7 |
| A6NM71 | PRA1 family protein OS=Homo sapiens OX=9606 PE=3 SV=3 | 0.995 | 0.942843201 | 0.831 | 0.7 |
| Q9BTV4 | Transmembrane protein 43 OS=Homo sapiens OX=9606 GN=TMEM43 PE=1 SV=1 | 0.985 | 0.904500837 | 1.017 | 0.9 |
| O76031 | ATP-dependent Clp protease ATP-binding subunit clpX-like, mitochondrial OS=Homo sapiens OX=9606 GN=CLPX PE=1 SV=2 | 1.056 | 0.935312766 | 0.931 | 0.7 |
| Q9HAT2 | Sialate O-acetylesterase OS=Homo sapiens OX=9606 GN=SIAE PE=1 SV=1 | 0.978 | 0.994833431 | 0.959 | 0.7 |
| C9JQS9 | Propionyl-CoA carboxylase beta chain, mitochondrial OS=Homo sapiens OX=9606 GN=PCCB PE=1 SV=1 | 0.83 | 0.762971614 | 0.771 | 0.7 |
| Q8NFH3 | Nucleoporin Nup43 OS=Homo sapiens OX=9606 GN=NUP43 PE=1 SV=1 | 1.126 | 0.765235202 | 1.068 | 0.9 |
| Q96B36 | Proline-rich AKT1 substrate 1 OS=Homo sapiens OX=9606 GN=AKT1S1 PE=1 SV=1 | 1.025 | 0.979006266 | 1.017 | 0.[illegible] |
| Q3MHD2 | Protein LSM12 homolog OS=Homo sapiens OX=9606 GN=LSM12 PE=1 SV=2 | 1.093 | 0.766667967 | 1.014 | 0.8 |
| E5RIZ4 | 39S ribosomal protein L15, mitochondrial (Fragment) OS=Homo sapiens OX=9606 GN=MRPL15 PE=1 SV=2 | 1.02 | 0.904500837 | 0.863 | 0.7 |
| Q5TA50 | Ceramide-1-phosphate transfer protein OS=Homo sapiens OX=9606 GN=CPTP PE=1 SV=1 | 0.958 | 0.876510992 | 1.044 | 0.8 |
| Q13618 | Cullin-3 OS=Homo sapiens OX=9606 GN=CUL3 PE=1 SV=2 | 1.006 | 0.886561233 | 0.901 | 0.7 |
| Q9P2C4 | Transmembrane protein 181 OS=Homo sapiens OX=9606 GN=TMEM181 PE=1 SV=2 | 0.952 | 0.876510992 | 0.804 | 0.7 |
| Q969S9 | Ribosome-releasing factor 2, mitochondrial OS=Homo sapiens OX=9606 GN=GFM2 PE=1 SV=1 | 0.986 | 0.899593149 | 0.788 | 0.7 |
| P49959 | Double-strand break repair protein MRE11 OS=Homo sapiens OX=9606 GN=MRE11 PE=1 SV=3 | 1.04 | 0.820396985 | 0.963 | 0.7 |
| P36639 | 7,8-dihydro-8-oxoguanine triphosphatase OS=Homo sapiens OX=9606 GN=NUDT1 PE=1 SV=3 | 0.951 | 0.763877559 | 0.912 | 0.7 |
| Q9Y478 | 5'-AMP-activated protein kinase subunit beta-1 OS=Homo sapiens OX=9606 GN=PRKAB1 PE=1 SV=4 | 1.064 | 0.907894335 | 0.927 | 0.7 |
| Q9NV88 | Integrator complex subunit 9 OS=Homo sapiens OX=9606 GN=INTS9 PE=1 SV=2 | 0.906 | 0.829390484 | 0.872 | 0.7 |
| J3KQ48 | Aminoacyl-tRNA hydrolase OS=Homo sapiens OX=9606 GN=PTRH2 PE=1 SV=1 | 0.929 | 0.977023215 | 0.843 | 0.7 |
| O00754 | Lysosomal alpha-mannosidase OS=Homo sapiens OX=9606 GN=MAN2B1 PE=1 SV=3 | 1.02 | 0.956825234 | 0.942 | 0.7 |
| Q8NHV4 | Protein NEDD1 OS=Homo sapiens OX=9606 GN=NEDD1 PE=1 SV=1 | 0.969 | 0.958996004 | 0.965 | 0.7 |
| F6XZQ7 | Glutathione S-transferase OS=Homo sapiens OX=9606 GN=GSTM2 PE=1 SV=1 | 1.068 | 0.845656075 | 0.948 | 0.7 |
| A0A494C1P1 | Actin-binding LIM protein 1 (Fragment) OS=Homo sapiens OX=9606 GN=ABLIM1 PE=1 SV=1 | 1.036 | 0.827871327 | 0.792 | 0.7 |
| Q16822 | Phosphoenolpyruvate carboxykinase [GTP], mitochondrial OS=Homo sapiens OX=9606 GN=PCK2 PE=1 SV=4 | 0.915 | 0.750023665 | 0.963 | 0.8 |
| A5YKK6 | CCR4-NOT transcription complex subunit 1 OS=Homo sapiens OX=9606 GN=CNOT1 PE=1 SV=2 | 0.93 | 0.85315742 | 0.932 | 0.7 |
| H0YIY8 | Serine/threonine-protein phosphatase 4 regulatory subunit 3A (Fragment) OS=Homo sapiens OX=9606 GN=PPP4R3A PE=1 SV=1 | 0.965 | 0.765235202 | 0.85 | 0.7 |
| Q9Y2V2 | Calcium-regulated heat-stable protein 1 OS=Homo sapiens OX=9606 GN=CARHSP1 PE=1 SV=2 | 0.917 | 0.827451813 | 0.921 | 0.8 |
| H0Y764 | Neurobeachin-like protein 2 (Fragment) OS=Homo sapiens OX=9606 GN=NBEAL2 PE=1 SV=1 | 0.98 | 0.895384791 | 0.886 | 0.7 |
| Q96CN9 | GRIP and coiled-coil domain-containing protein 1 OS=Homo sapiens OX=9606 GN=GCC1 PE=1 SV=1 | 0.827 | 0.750023665 | 1.016 | 0.[illegible] |
| Q9P035 | Very-long-chain (3R)-3-hydroxyacyl-CoA dehydratase 3 OS=Homo sapiens OX=9606 GN=HACD3 PE=1 SV=2 | 0.959 | 0.93219119 | 0.912 | 0.7 |
| P05114 | Non-histone chromosomal protein HMG-14 OS=Homo sapiens OX=9606 GN=HMGN1 PE=1 SV=3 | 1.039 | 0.809620887 | 0.942 | 0.8 |
| Q7L5N7 | Lysophosphatidylcholine acyltransferase 2 OS=Homo sapiens OX=9606 GN=LPCAT2 PE=1 SV=1 | 1.072 | 0.984200071 | 0.923 | 0.7 |
| B1AXG1 | Non-specific serine/threonine protein kinase OS=Homo sapiens OX=9606 GN=RPS6KA3 PE=1 SV=2 | 0.999 | 0.859846322 | 0.949 | 0.7 |
| G8JLA1 | Retinol dehydrogenase 13 OS=Homo sapiens OX=9606 GN=RDH13 PE=1 SV=1 | 1.063 | 0.91571242 | 0.806 | 0.7 |
| H7BXY3 | RNA helicase OS=Homo sapiens OX=9606 GN=DHX30 PE=1 SV=1 | 0.877 | 0.750023665 | 0.918 | 0.7 |
| B5MC98 | Prolactin regulatory element-binding protein OS=Homo sapiens OX=9606 GN=PREB PE=1 SV=1 | 1.098 | 0.766667967 | 0.937 | 0.8 |
| Q92890-1 | Isoform Long of Ubiquitin recognition factor in ER-associated degradation protein 1 OS=Homo sapiens OX=9606 GN=UFD1 | 1.048 | 0.765235202 | 1.01 | 0.[illegible] |
| O43704 | Sulfotransferase family cytosolic 1B member 1 OS=Homo sapiens OX=9606 GN=SULT1B1 PE=1 SV=2 | 1.153 | 0.750023665 | 0.962 | 0.7 |
| Q96HW7 | Integrator complex subunit 4 OS=Homo sapiens OX=9606 GN=INTS4 PE=1 SV=2 | 0.954 | 0.927148669 | 1.042 | 0.8 |
| E7EWW0 | VPS35 endosomal protein sorting factor-like OS=Homo sapiens OX=9606 GN=VPS35L PE=1 SV=2 | 1.011 | 0.932424612 | 0.845 | 0.7 |
| P11308 | Transcriptional regulator ERG OS=Homo sapiens OX=9606 GN=ERG PE=1 SV=3 | 1.033 | 0.750023665 | 1.023 | 0.[illegible] |
| F8VU90 | Peptidylprolyl isomerase OS=Homo sapiens OX=9606 GN=FKBP11 PE=1 SV=1 | | | | |
| H0YAN8 | Rho guanine nucleotide exchange factor 10 (Fragment) OS=Homo sapiens OX=9606 GN=ARHGEF10 PE=1 SV=1 | 0.946 | 0.829390484 | 0.908 | 0.7 |
| Q8WTW3 | Conserved oligomeric Golgi complex subunit 1 OS=Homo sapiens OX=9606 GN=COG1 PE=1 SV=1 | 0.997 | 0.95980123 | 0.89 | 0.7 |
| P53990 | IST1 homolog OS=Homo sapiens OX=9606 GN=IST1 PE=1 SV=1 | 1.048 | 0.979678753 | 0.974 | 0.7 |
| Q460N5-1 | Isoform 1 of Protein mono-ADP-ribosyltransferase PARP14 OS=Homo sapiens OX=9606 GN=PARP14 | 1.354 | 0.820396985 | 0.857 | 0.7 |
| Q9BU23 | Lipase maturation factor 2 OS=Homo sapiens OX=9606 GN=LMF2 PE=1 SV=2 | 1.018 | 0.876510992 | 0.869 | 0.[illegible] |
| Q13405 | 39S ribosomal protein L49, mitochondrial OS=Homo sapiens OX=9606 GN=MRPL49 PE=1 SV=1 | 1.086 | 0.750023665 | 0.934 | 0.7 |
| Q9ULE6 | Paladin OS=Homo sapiens OX=9606 GN=PALD1 PE=1 SV=3 | 0.875 | 0.779241609 | 0.717 | 0.7 |
| Q8NFH4 | Nucleoporin Nup37 OS=Homo sapiens OX=9606 GN=NUP37 PE=1 SV=1 | 0.863 | 0.766667967 | 0.938 | 0.7 |
| P36404 | ADP-ribosylation factor-like protein 2 OS=Homo sapiens OX=9606 GN=ARL2 PE=1 SV=4 | 0.973 | 0.765235202 | 0.957 | 0.7 |
| P52948 | Nuclear pore complex protein Nup98-Nup96 OS=Homo sapiens OX=9606 GN=NUP98 PE=1 SV=4 | 0.9 | 0.765235202 | 0.925 | 0.7 |

| | | | | | |
|---|---|---|---|---|---|
| Q9NXS2 | Glutaminyl-peptide cyclotransferase-like protein OS=Homo sapiens OX=9606 GN=QPCTL PE=1 SV=2 | 0.98 | 0.978562349 | 0.881 | 0.7 |
| Q16762 | Thiosulfate sulfurtransferase OS=Homo sapiens OX=9606 GN=TST PE=1 SV=4 | 1.043 | 0.9962822 | 0.879 | 0.7 |
| P62861 | 40S ribosomal protein S30 OS=Homo sapiens OX=9606 GN=FAU PE=1 SV=1 | 1.193 | 0.765235202 | 1.108 | 0.8 |
| E5RGN3 | Copper transport protein ATOX1 OS=Homo sapiens OX=9606 GN=ATOX1 PE=1 SV=1 | 0.972 | 0.816240759 | 1.039 | 0.7 |
| O14908 | PDZ domain-containing protein GIPC1 OS=Homo sapiens OX=9606 GN=GIPC1 PE=1 SV=2 | 1.054 | 0.876510992 | 0.869 | 0.7 |
| Q9NQT8 | Kinesin-like protein KIF13B OS=Homo sapiens OX=9606 GN=KIF13B PE=1 SV=2 | 0.93 | 0.889482529 | 0.96 | 0.7 |
| H3BLV9 | SRSF protein kinase 1 (Fragment) OS=Homo sapiens OX=9606 GN=SRPK1 PE=1 SV=1 | 1.066 | 0.754126086 | 1.128 | 0.7 |
| O60547 | GDP-mannose 4,6 dehydratase OS=Homo sapiens OX=9606 GN=GMDS PE=1 SV=1 | 0.974 | 0.845656075 | 1.06 | 0.7 |
| Q96P16 | Regulation of nuclear pre-mRNA domain-containing protein 1A OS=Homo sapiens OX=9606 GN=RPRD1A PE=1 SV=1 | 1.004 | 0.983953845 | 0.861 | 0.7 |
| O00506 | Serine/threonine-protein kinase 25 OS=Homo sapiens OX=9606 GN=STK25 PE=1 SV=1 | 1.075 | 0.881099595 | 1.161 | 0.7 |
| P51397 | Death-associated protein 1 OS=Homo sapiens OX=9606 GN=DAP PE=1 SV=3 | 1.083 | 0.951324063 | 0.995 | 0.9 |
| P29353 | SHC-transforming protein 1 OS=Homo sapiens OX=9606 GN=SHC1 PE=1 SV=4 | 1.012 | 0.988027887 | 0.993 | 0.9 |
| Q99988 | Growth/differentiation factor 15 OS=Homo sapiens OX=9606 GN=GDF15 PE=1 SV=3 | 0.94 | 0.991321414 | 0.901 | 0.7 |
| Q8NFF5 | FAD synthase OS=Homo sapiens OX=9606 GN=FLAD1 PE=1 SV=1 | 0.992 | 0.996637279 | 1.038 | 0.8 |
| C9JJX6 | Armadillo repeat protein deleted in velo-cardio-facial syndrome OS=Homo sapiens OX=9606 GN=ARVCF PE=1 SV=1 | 0.951 | 0.750023665 | 0.704 | 0.7 |
| O95232 | Luc7-like protein 3 OS=Homo sapiens OX=9606 GN=LUC7L3 PE=1 SV=2 | 0.988 | 0.860909277 | 1.001 | 0.9 |
| Q12797 | Aspartyl/asparaginyl beta-hydroxylase OS=Homo sapiens OX=9606 GN=ASPH PE=1 SV=3 | 1.033 | 0.889450577 | 0.925 | 0.7 |
| Q8WUA2 | Peptidyl-prolyl cis-trans isomerase-like 4 OS=Homo sapiens OX=9606 GN=PPIL4 PE=1 SV=1 | 0.97 | 0.995582922 | 0.975 | 0.9 |
| Q92979 | Ribosomal RNA small subunit methyltransferase NEP1 OS=Homo sapiens OX=9606 GN=EMG1 PE=1 SV=4 | 0.984 | 0.953792095 | 1.11 | 0.7 |
| P09234 | U1 small nuclear ribonucleoprotein C OS=Homo sapiens OX=9606 GN=SNRPC PE=1 SV=1 | 1.106 | 0.765235202 | 0.967 | 0.7 |
| Q9UKF6 | Cleavage and polyadenylation specificity factor subunit 3 OS=Homo sapiens OX=9606 GN=CPSF3 PE=1 SV=1 | 0.966 | 0.983953845 | 0.96 | 0.8 |
| A0A3B3ITR4 | Uncharacterized protein (Fragment) OS=Homo sapiens OX=9606 PE=4 SV=1 | 1.006 | 0.91999213 | 0.985 | 0.8 |
| P0DMN0 | Sulfotransferase 1A4 OS=Homo sapiens OX=9606 GN=SULT1A4 PE=1 SV=1 | 0.995 | 0.883217803 | 0.928 | 0.7 |
| Q9Y2E5 | Epididymis-specific alpha-mannosidase OS=Homo sapiens OX=9606 GN=MAN2B2 PE=1 SV=4 | 1.01 | 0.996578046 | 0.972 | 0.7 |
| P08243 | Asparagine synthetase [glutamine-hydrolyzing] OS=Homo sapiens OX=9606 GN=ASNS PE=1 SV=4 | 0.973 | 0.984805107 | 1.154 | 0.7 |
| O15347 | High mobility group protein B3 OS=Homo sapiens OX=9606 GN=HMGB3 PE=1 SV=4 | 1.208 | 0.750023665 | 1.014 | 0.8 |
| Q6P2I3 | Fumarylacetoacetate hydrolase domain-containing protein 2B OS=Homo sapiens OX=9606 GN=FAHD2B PE=1 SV=1 | 0.961 | 0.901934028 | 1.099 | 0.7 |
| Q8WYA6 | Beta-catenin-like protein 1 OS=Homo sapiens OX=9606 GN=CTNNBL1 PE=1 SV=1 | 1.031 | 0.779783772 | 0.943 | 0.7 |
| H7C0W7 | Hyccin (Fragment) OS=Homo sapiens OX=9606 GN=FAM126A PE=1 SV=1 | 1.01 | 0.994361756 | 0.964 | 0.8 |
| Q9UNW1 | Multiple inositol polyphosphate phosphatase 1 OS=Homo sapiens OX=9606 GN=MINPP1 PE=1 SV=1 | 0.938 | 0.750023665 | 1.005 | 0.7 |
| Q9H2M9 | Rab3 GTPase-activating protein non-catalytic subunit OS=Homo sapiens OX=9606 GN=RAB3GAP2 PE=1 SV=1 | 1.026 | 0.951699549 | 0.906 | 0.7 |
| Q9BXK5 | Bcl-2-like protein 13 OS=Homo sapiens OX=9606 GN=BCL2L13 PE=1 SV=1 | 0.954 | 0.81199948 | 0.883 | 0.7 |
| Q969V3 | Nicalin OS=Homo sapiens OX=9606 GN=NCLN PE=1 SV=2 | 1.014 | 0.901934028 | 1.015 | 0.9 |
| F5GZY7 | Gamma-aminobutyric acid receptor-associated protein-like 1 (Fragment) OS=Homo sapiens OX=9606 GN=GABARAPL1 PE=1 SV=1 | 0.886 | 0.845356491 | 0.905 | 0.7 |
| Q9UKG1 | DCC-interacting protein 13-alpha OS=Homo sapiens OX=9606 GN=APPL1 PE=1 SV=1 | 0.889 | 0.913279108 | 1.17 | 0.9 |
| O95183 | Vesicle-associated membrane protein 5 OS=Homo sapiens OX=9606 GN=VAMP5 PE=1 SV=1 | 1.063 | 0.765235202 | 1.077 | 0.8 |
| Q9BV79 | Enoyl-[acyl-carrier-protein] reductase, mitochondrial OS=Homo sapiens OX=9606 GN=MECR PE=1 SV=2 | 1.166 | 0.859846322 | 1.274 | 0.7 |
| A0A286YEW0 | ER membrane protein complex subunit 1 (Fragment) OS=Homo sapiens OX=9606 GN=EMC1 PE=1 SV=1 | 0.88 | 0.750023665 | 0.849 | 0.7 |
| P41208 | Centrin-2 OS=Homo sapiens OX=9606 GN=CETN2 PE=1 SV=1 | 1.045 | 0.87017509 | 0.992 | 0.8 |
| I3L3X5 | Phospholipid scramblase OS=Homo sapiens OX=9606 GN=PLSCR3 PE=1 SV=1 | 0.998 | 0.978562349 | 0.882 | 0.7 |
| Q8WYP3 | Ras and Rab interactor 2 OS=Homo sapiens OX=9606 GN=RIN2 PE=1 SV=1 | 0.95 | 0.754126086 | 1.023 | 0.8 |
| Q13243 | Serine/arginine-rich splicing factor 5 OS=Homo sapiens OX=9606 GN=SRSF5 PE=1 SV=1 | 1.018 | 0.952913364 | 1.018 | 0.8 |
| Q8WUH2 | Transforming growth factor-beta receptor-associated protein 1 OS=Homo sapiens OX=9606 GN=TGFBRAP1 PE=1 SV=1 | 0.954 | 0.750023665 | 0.857 | 0.7 |
| P15090 | Fatty acid-binding protein, adipocyte OS=Homo sapiens OX=9606 GN=FABP4 PE=1 SV=3 | 1.012 | 0.983953845 | 0.751 | 0.7 |
| Q9NZ45 | CDGSH iron-sulfur domain-containing protein 1 OS=Homo sapiens OX=9606 GN=CISD1 PE=1 SV=1 | | | | |
| Q9NSY1 | BMP-2-inducible protein kinase OS=Homo sapiens OX=9606 GN=BMP2K PE=1 SV=2 | 0.936 | 0.765235202 | 0.972 | 0.8 |
| Q9UNN8 | Endothelial protein C receptor OS=Homo sapiens OX=9606 GN=PROCR PE=1 SV=1 | 0.922 | 0.77554521 | 1.049 | 0.7 |
| A0A3B3IRL5 | Inosine-5'-monophosphate dehydrogenase (Fragment) OS=Homo sapiens OX=9606 GN=IMPDH1 PE=1 SV=1 | 1.016 | 0.87017509 | 0.999 | 0.9 |
| P41214 | Eukaryotic translation initiation factor 2D OS=Homo sapiens OX=9606 GN=EIF2D PE=1 SV=3 | 0.915 | 0.951693691 | 1.017 | 0.8 |
| Q9NUQ8 | ATP-binding cassette sub-family F member 3 OS=Homo sapiens OX=9606 GN=ABCF3 PE=1 SV=2 | 1.047 | 0.765235202 | 1.021 | 0.9 |
| Q15717 | ELAV-like protein 1 OS=Homo sapiens OX=9606 GN=ELAVL1 PE=1 SV=2 | 0.985 | 0.985324977 | 1.02 | 0.9 |
| A0A0A0MRW6 | Nucleolar protein 6 OS=Homo sapiens OX=9606 GN=NOL6 PE=1 SV=1 | 1.333 | 0.680740328 | 0.688 | 0.7 |
| E9PQX9 | Dr1-associated corepressor OS=Homo sapiens OX=9606 GN=DRAP1 PE=1 SV=1 | 1.005 | 0.994361756 | 0.766 | 0.7 |
| P63172 | Dynein light chain Tctex-type 1 OS=Homo sapiens OX=9606 GN=DYNLT1 PE=1 SV=1 | 1.048 | 0.902016292 | 0.902 | 0.7 |
| F5GZ49 | Glycolipid transfer protein OS=Homo sapiens OX=9606 GN=GLTP PE=1 SV=1 | 0.83 | 0.889482529 | 0.892 | 0.8 |
| P29218 | Inositol monophosphatase 1 OS=Homo sapiens OX=9606 GN=IMPA1 PE=1 SV=1 | 0.962 | 0.829390484 | 0.941 | 0.7 |
| E9PC62 | CUGBP Elav-like family member 2 OS=Homo sapiens OX=9606 GN=CELF2 PE=1 SV=1 | 0.86 | 0.765235202 | 0.967 | 0.9 |
| A0A1W2PRL9 | Transcription elongation factor A protein 1 OS=Homo sapiens OX=9606 GN=TCEA1 PE=1 SV=1 | 0.894 | 0.930855729 | 1.066 | 0.7 |
| P19022 | Cadherin-2 OS=Homo sapiens OX=9606 GN=CDH2 PE=1 SV=4 | 0.985 | 0.886561233 | 0.979 | 0.7 |
| Q712K3 | Ubiquitin-conjugating enzyme E2 R2 OS=Homo sapiens OX=9606 GN=UBE2R2 PE=1 SV=1 | 0.986 | 0.889482529 | 1.008 | 0.9 |
| Q8WWB7 | Glycosylated lysosomal membrane protein OS=Homo sapiens OX=9606 GN=GLMP PE=1 SV=1 | | | | |
| P05106 | Integrin beta-3 OS=Homo sapiens OX=9606 GN=ITGB3 PE=1 SV=2 | 1.036 | 0.765235202 | 0.94 | 0.9 |
| Q8TBQ9 | Protein kish-A OS=Homo sapiens OX=9606 GN=TMEM167A PE=1 SV=1 | 0.98 | 0.766667967 | 0.92 | 0.7 |
| X6RLX0 | ELKS/Rab6-interacting/CAST family member 1 OS=Homo sapiens OX=9606 GN=ERC1 PE=1 SV=1 | 1.038 | 0.703015003 | 1.009 | 0.8 |
| Q12891 | Hyaluronidase-2 OS=Homo sapiens OX=9606 GN=HYAL2 PE=1 SV=4 | 0.988 | 0.860909277 | 1.004 | 0.8 |
| J3KSL8 | Trafficking protein particle complex subunit 8 OS=Homo sapiens OX=9606 GN=TRAPPC8 PE=1 SV=1 | 1.07 | 0.884192327 | 0.921 | 0.7 |
| Q9NQ50 | 39S ribosomal protein L40, mitochondrial OS=Homo sapiens OX=9606 GN=MRPL40 PE=1 SV=1 | 1.004 | 0.925659097 | 0.885 | 0.7 |
| D6RBG2 | Protocadherin-1 (Fragment) OS=Homo sapiens OX=9606 GN=PCDH1 PE=1 SV=1 | 1.056 | 0.964351664 | 0.855 | 0.7 |
| A0A6I8PIX7 | Supervillin OS=Homo sapiens OX=9606 GN=SVIL PE=4 SV=1 | 0.961 | 0.766667967 | 0.963 | 0.7 |
| A0A590UJL2 | Disks large homolog 1 OS=Homo sapiens OX=9606 GN=DLG1 PE=1 SV=1 | 1.069 | 0.994361756 | 0.837 | 0.7 |
| A0A0A0MSS8 | Aldo-keto reductase family 1 member C3 OS=Homo sapiens OX=9606 GN=AKR1C3 PE=1 SV=1 | 1.022 | 0.963059765 | 0.997 | 0.9 |
| O75475 | PC4 and SFRS1-interacting protein OS=Homo sapiens OX=9606 GN=PSIP1 PE=1 SV=1 | 1.109 | 0.786409349 | 1.002 | 0.9 |
| O94826 | Mitochondrial import receptor subunit TOM70 OS=Homo sapiens OX=9606 GN=TOMM70 PE=1 SV=1 | 0.993 | 0.973083765 | 0.969 | 0.7 |
| O95816 | BAG family molecular chaperone regulator 2 OS=Homo sapiens OX=9606 GN=BAG2 PE=1 SV=1 | 1.012 | 0.95071661 | 1.01 | 0.9 |
| A0A096LP25 | AP2-associated protein kinase 1 (Fragment) OS=Homo sapiens OX=9606 GN=AAK1 PE=1 SV=1 | 1.025 | 0.909996844 | 1.007 | 0.9 |
| Q69YN4 | Protein virilizer homolog OS=Homo sapiens OX=9606 GN=VIRMA PE=1 SV=2 | 0.886 | 0.95980123 | 0.878 | 0.7 |
| A0A1W2PQ12 | Golgi SNAP receptor complex member 2 (Fragment) OS=Homo sapiens OX=9606 GN=GOSR2 PE=4 SV=1 | 0.941 | 0.95980123 | 0.968 | 0.7 |
| A0A0D9SFD6 | Bifunctional polynucleotide phosphatase/kinase (Fragment) OS=Homo sapiens OX=9606 GN=PNKP PE=1 SV=1 | 1.002 | 0.990981579 | 0.974 | 0.7 |
| C9J4U8 | Ventricular zone-expressed PH domain-containing protein homolog 1 (Fragment) OS=Homo sapiens OX=9606 GN=VEPH1 PE=1 SV | 0.968 | 0.956825234 | 0.935 | 0.7 |
| Q9Y4Y9 | U6 snRNA-associated Sm-like protein LSm5 OS=Homo sapiens OX=9606 GN=LSM5 PE=1 SV=3 | 0.983 | 0.942843201 | 0.991 | 0.8 |
| B7Z2L0 | B-cell receptor-associated protein OS=Homo sapiens OX=9606 GN=BCAP29 PE=1 SV=1 | 0.97 | 0.882454148 | 1.002 | 0.9 |
| P50402 | Emerin OS=Homo sapiens OX=9606 GN=EMD PE=1 SV=1 | 0.929 | 0.829390484 | 0.967 | 0.9 |
| Q5JS79 | Peripheral plasma membrane protein CASK (Fragment) OS=Homo sapiens OX=9606 GN=CASK PE=1 SV=2 | 1.15 | 0.765235202 | 0.936 | 0.9 |
| O60784 | Target of Myb protein 1 OS=Homo sapiens OX=9606 GN=TOM1 PE=1 SV=2 | 0.953 | 0.992258896 | 0.88 | 0.7 |
| Q9Y3E0 | Vesicle transport protein GOT1B OS=Homo sapiens OX=9606 GN=GOLT1B PE=1 SV=1 | 0.83 | 0.750023665 | 1.045 | 0.9 |
| Q9P0J7 | E3 ubiquitin-protein ligase KCMF1 OS=Homo sapiens OX=9606 GN=KCMF1 PE=1 SV=2 | 0.931 | 0.756287988 | 0.945 | 0.7 |
| C9JTT7 | Replication factor C subunit 4 (Fragment) OS=Homo sapiens OX=9606 GN=RFC4 PE=1 SV=1 | 1.034 | 0.839954644 | 0.887 | 0.7 |
| P23469 | Receptor-type tyrosine-protein phosphatase epsilon OS=Homo sapiens OX=9606 GN=PTPRE PE=1 SV=1 | 1.017 | 0.929331324 | 0.888 | 0.9 |
| O75970 | Multiple PDZ domain protein OS=Homo sapiens OX=9606 GN=MPDZ PE=1 SV=2 | 0.922 | 0.883924281 | 0.816 | 0.7 |
| A0A087WYN9 | ATP-dependent RNA helicase DHX29 OS=Homo sapiens OX=9606 GN=DHX29 PE=1 SV=1 | 1.062 | 0.882454148 | 0.931 | 0.7 |
| H0YE25 | Glutamine amidotransferase-like class 1 domain-containing protein 1 (Fragment) OS=Homo sapiens OX=9606 GN=GATD1 PE=1 SV | 0.954 | 0.750023665 | 0.92 | 0.7 |
| Q99653 | Calcineurin B homologous protein 1 OS=Homo sapiens OX=9606 GN=CHP1 PE=1 SV=3 | 1.013 | 0.905369216 | 0.943 | 0.7 |
| P47985 | Cytochrome b-c1 complex subunit Rieske, mitochondrial OS=Homo sapiens OX=9606 GN=UQCRFS1 PE=1 SV=2 | 0.986 | 0.85673819 | 1.009 | 0.9 |
| E9PHA2 | Condensin complex subunit 2 OS=Homo sapiens OX=9606 GN=NCAPH PE=1 SV=1 | 0.951 | 0.884192327 | 1.062 | 0.8 |
| Q3HKR1 | RAP1 short isoform OS=Homo sapiens OX=9606 GN=RABGEF1 PE=1 SV=1 | 0.995 | 0.983953845 | 0.818 | 0.7 |
| X6RLR1 | Dynactin subunit 3 OS=Homo sapiens OX=9606 GN=DCTN3 PE=1 SV=1 | 1.003 | 0.942843201 | 1.033 | 0.9 |

| | | | | | |
|---|---|---|---|---|---|
| Q9BYN0 | Sulfiredoxin-1 OS=Homo sapiens OX=9606 GN=SRXN1 PE=1 SV=2 | 1.071 | 0.750023665 | 1.105 | 0.9 |
| H7C5I5 | Phospholipid scramblase (Fragment) OS=Homo sapiens OX=9606 GN=PLSCR1 PE=1 SV=1 | 0.983 | 0.750023665 | 1.069 | 0.9 |
| Q8WU90 | Zinc finger CCCH domain-containing protein 15 OS=Homo sapiens OX=9606 GN=ZC3H15 PE=1 SV=1 | 1.051 | 0.765235202 | 1.046 | 0.7 |
| Q16740 | ATP-dependent Clp protease proteolytic subunit, mitochondrial OS=Homo sapiens OX=9606 GN=CLPP PE=1 SV=1 | 1.045 | 0.809976184 | 0.995 | 0.9 |
| F8VSC5 | SCY1-like protein 2 (Fragment) OS=Homo sapiens OX=9606 GN=SCYL2 PE=1 SV=1 | 1.031 | 0.750023665 | 0.939 | 0.7 |
| P08574 | Cytochrome c1, heme protein, mitochondrial OS=Homo sapiens OX=9606 GN=CYC1 PE=1 SV=3 | 1.034 | 0.949927239 | 0.79 | 0.7 |
| Q9Y3A6 | Transmembrane emp24 domain-containing protein 5 OS=Homo sapiens OX=9606 GN=TMED5 PE=1 SV=1 | 0.976 | 0.909944495 | 0.875 | 0.7 |
| H7C440 | DIS3-like exonuclease 2 (Fragment) OS=Homo sapiens OX=9606 GN=DIS3L2 PE=1 SV=1 | | | | |
| K7EIU8 | Mothers against decapentaplegic homolog OS=Homo sapiens OX=9606 GN=SMAD4 PE=1 SV=1 | 0.938 | 0.917008272 | 0.985 | 0.7 |
| P53701 | Cytochrome c-type heme lyase OS=Homo sapiens OX=9606 GN=HCCS PE=1 SV=1 | 1.034 | 0.913279108 | 0.914 | 0.7 |
| P05362 | Intercellular adhesion molecule 1 OS=Homo sapiens OX=9606 GN=ICAM1 PE=1 SV=2 | 1.017 | 0.963059765 | 1.041 | 0.7 |
| Q01581 | Hydroxymethylglutaryl-CoA synthase, cytoplasmic OS=Homo sapiens OX=9606 GN=HMGCS1 PE=1 SV=2 | 0.995 | 0.974919191 | 0.902 | 0.8 |
| O60645 | Exocyst complex component 3 OS=Homo sapiens OX=9606 GN=EXOC3 PE=1 SV=3 | 1.033 | 0.994361756 | 0.862 | 0.7 |
| P57081 | tRNA (guanine-N(7)-)-methyltransferase non-catalytic subunit WDR4 OS=Homo sapiens OX=9606 GN=WDR4 PE=1 SV=2 | 1.036 | 0.9962822 | 0.974 | 0.7 |
| P03956 | Interstitial collagenase OS=Homo sapiens OX=9606 GN=MMP1 PE=1 SV=3 | 1.134 | 0.750023665 | 1.253 | 0.7 |
| Q9P266 | Junctional protein associated with coronary artery disease OS=Homo sapiens OX=9606 GN=JCAD PE=1 SV=3 | 0.831 | 0.765235202 | 0.847 | 0.7 |
| I3L2L5 | Mapk-regulated corepressor-interacting protein 1 OS=Homo sapiens OX=9606 GN=MCRIP1 PE=1 SV=1 | 0.996 | 0.908555656 | 0.97 | 0.8 |
| P49790 | Nuclear pore complex protein Nup153 OS=Homo sapiens OX=9606 GN=NUP153 PE=1 SV=2 | | | | |
| A2A376 | RNA helicase OS=Homo sapiens OX=9606 GN=DDX58 PE=1 SV=1 | 0.914 | 0.806305724 | 0.796 | 0.7 |
| K4DIA7 | Tetraspanin (Fragment) OS=Homo sapiens OX=9606 GN=CD151 PE=1 SV=1 | 0.93 | 0.859846322 | 1.025 | 0.7 |
| S4R369 | 39S ribosomal protein L37, mitochondrial OS=Homo sapiens OX=9606 GN=MRPL37 PE=1 SV=1 | 1 | 0.985763982 | 0.907 | 0.7 |
| Q8IV08 | 5'-3' exonuclease PLD3 OS=Homo sapiens OX=9606 GN=PLD3 PE=1 SV=1 | 0.953 | 0.752728302 | 0.95 | 0.7 |
| A6NNK5 | TP53-binding protein 1 OS=Homo sapiens OX=9606 GN=TP53BP1 PE=1 SV=2 | 1.014 | 0.918702104 | 0.793 | 0.7 |
| A0A1B0GUA8 | Kinesin-like protein KIF13A OS=Homo sapiens OX=9606 GN=KIF13A PE=1 SV=1 | 0.973 | 0.866300162 | 1.043 | 0.9 |
| Q13085 | Acetyl-CoA carboxylase 1 OS=Homo sapiens OX=9606 GN=ACACA PE=1 SV=2 | 1.117 | 0.849526028 | 0.933 | 0.7 |
| Q9Y3B4 | Splicing factor 3B subunit 6 OS=Homo sapiens OX=9606 GN=SF3B6 PE=1 SV=1 | 0.959 | 0.987438982 | 0.987 | 0.7 |
| O43823 | A-kinase anchor protein 8 OS=Homo sapiens OX=9606 GN=AKAP8 PE=1 SV=1 | 0.971 | 0.900915273 | 0.656 | 0.7 |
| Q9UKD2 | mRNA turnover protein 4 homolog OS=Homo sapiens OX=9606 GN=MRTO4 PE=1 SV=2 | 1.002 | 0.880494956 | 1.026 | 0.8 |
| Q01780 | Exosome component 10 OS=Homo sapiens OX=9606 GN=EXOSC10 PE=1 SV=2 | 1.041 | 0.750023665 | 0.916 | 0.8 |
| Q9NY15 | Stabilin-1 OS=Homo sapiens OX=9606 GN=STAB1 PE=1 SV=3 | 0.938 | 0.873607677 | 1.096 | 0.7 |
| Q3LXA3 | Triokinase/FMN cyclase OS=Homo sapiens OX=9606 GN=TKFC PE=1 SV=2 | 1.026 | 0.938907165 | 0.904 | 0.7 |
| A6NML8 | Diaphanous homolog 2 (Drosophila), isoform CRA_c OS=Homo sapiens OX=9606 GN=DIAPH2 PE=1 SV=3 | 0.991 | 0.901934028 | 0.812 | 0.7 |
| Q9NVH6 | Trimethyllysine dioxygenase, mitochondrial OS=Homo sapiens OX=9606 GN=TMLHE PE=1 SV=1 | 1.107 | 0.750023665 | 0.882 | 0.7 |
| Q9Y2C4 | Nuclease EXOG, mitochondrial OS=Homo sapiens OX=9606 GN=EXOG PE=1 SV=2 | 0.84 | 0.914805529 | 0.723 | 0.7 |
| Q8WX93 | Palladin OS=Homo sapiens OX=9606 GN=PALLD PE=1 SV=3 | 1.039 | 0.750023665 | 0.865 | 0.7 |
| O43865 | S-adenosylhomocysteine hydrolase-like protein 1 OS=Homo sapiens OX=9606 GN=AHCYL1 PE=1 SV=2 | 0.865 | 0.703015003 | 0.923 | 0.7 |
| A0A5F9ZHW5 | Ataxin-2 OS=Homo sapiens OX=9606 GN=ATXN2 PE=1 SV=1 | 1.054 | 0.96647353 | 1.013 | 0.9 |
| Q8TC07 | TBC1 domain family member 15 OS=Homo sapiens OX=9606 GN=TBC1D15 PE=1 SV=2 | 1.232 | 0.703015003 | 0.963 | 0.7 |
| Q70CQ2 | Ubiquitin carboxyl-terminal hydrolase 34 OS=Homo sapiens OX=9606 GN=USP34 PE=1 SV=2 | 0.831 | 0.816716243 | 0.828 | 0.7 |
| Q8WXA9 | Splicing regulatory glutamine/lysine-rich protein 1 OS=Homo sapiens OX=9606 GN=SREK1 PE=1 SV=1 | 0.969 | 0.918702104 | 0.91 | 0.7 |
| Q9NZT2 | Opioid growth factor receptor OS=Homo sapiens OX=9606 GN=OGFR PE=1 SV=3 | 0.87 | 0.876510992 | 0.981 | 0.7 |
| H0YHU4 | tRNA (guanine-N(7)-)-methyltransferase (Fragment) OS=Homo sapiens OX=9606 GN=METTL1 PE=1 SV=1 | 0.982 | 0.985324977 | 0.889 | 0.7 |
| H0YB09 | D-glutamate cyclase, mitochondrial (Fragment) OS=Homo sapiens OX=9606 GN=DGLUCY PE=1 SV=1 | 0.991 | 0.989102644 | 0.979 | 0.9 |
| Q9HA64 | Ketosamine-3-kinase OS=Homo sapiens OX=9606 GN=FN3KRP PE=1 SV=2 | 0.922 | 0.703015003 | 1.011 | 0.9 |
| Q9NYU1 | UDP-glucose:glycoprotein glucosyltransferase 2 OS=Homo sapiens OX=9606 GN=UGGT2 PE=1 SV=4 | 0.84 | 0.820396985 | 0.824 | 0.7 |
| Q13444 | Disintegrin and metalloproteinase domain-containing protein 15 OS=Homo sapiens OX=9606 GN=ADAM15 PE=1 SV=4 | 1.041 | 0.902016292 | 1.005 | 0.8 |
| A0A6I8PRA0 | Treacle protein (Fragment) OS=Homo sapiens OX=9606 GN=TCOF1 PE=4 SV=1 | 1.031 | 0.903226585 | 1.149 | 0.7 |
| Q9Y6K5 | 2'-5'-oligoadenylate synthase 3 OS=Homo sapiens OX=9606 GN=OAS3 PE=1 SV=3 | 0.95 | 0.946055791 | 0.88 | 0.7 |
| Q13614 | Myotubularin-related protein 2 OS=Homo sapiens OX=9606 GN=MTMR2 PE=1 SV=4 | | | | |
| Q8WXX5 | DnaJ homolog subfamily C member 9 OS=Homo sapiens OX=9606 GN=DNAJC9 PE=1 SV=1 | 1.012 | 0.952913364 | 0.931 | 0.7 |
| Q9BW92 | Threonine--tRNA ligase, mitochondrial OS=Homo sapiens OX=9606 GN=TARS2 PE=1 SV=1 | 1.02 | 0.921087329 | 0.92 | 0.7 |
| O15066 | Kinesin-like protein KIF3B OS=Homo sapiens OX=9606 GN=KIF3B PE=1 SV=1 | 0.845 | 0.924563362 | 0.933 | 0.8 |
| P50416 | Carnitine O-palmitoyltransferase 1, liver isoform OS=Homo sapiens OX=9606 GN=CPT1A PE=1 SV=2 | 1.015 | 0.930855729 | 0.829 | 0.7 |
| Q96T51 | RUN and FYVE domain-containing protein 1 OS=Homo sapiens OX=9606 GN=RUFY1 PE=1 SV=2 | 0.963 | 0.766667967 | 0.913 | 0.7 |
| Q9BTE6 | Alanyl-tRNA editing protein Aarsd1 OS=Homo sapiens OX=9606 GN=AARSD1 PE=1 SV=2 | 1.027 | 0.930855729 | 0.93 | 0.7 |
| P49761-1 | Isoform 1 of Dual specificity protein kinase CLK3 OS=Homo sapiens OX=9606 GN=CLK3 | 1.183 | 0.849526028 | 0.96 | 0.8 |
| Q9NPJ3 | Acyl-coenzyme A thioesterase 13 OS=Homo sapiens OX=9606 GN=ACOT13 PE=1 SV=1 | 0.974 | 0.792509319 | 0.964 | 0.8 |
| F8W9U3 | Protein ABHD14B OS=Homo sapiens OX=9606 GN=ABHD14B PE=1 SV=1 | 1.101 | 0.765235202 | 0.908 | 0.7 |
| H0Y9U7 | Long-chain-fatty-acid--CoA ligase 1 (Fragment) OS=Homo sapiens OX=9606 GN=ACSL1 PE=1 SV=1 | 1.037 | 0.910192239 | 0.969 | 0.7 |
| O95817 | BAG family molecular chaperone regulator 3 OS=Homo sapiens OX=9606 GN=BAG3 PE=1 SV=3 | 1.126 | 0.766667967 | 1.054 | 0.7 |
| Q15121 | Astrocytic phosphoprotein PEA-15 OS=Homo sapiens OX=9606 GN=PEA15 PE=1 SV=2 | 0.915 | 0.930427413 | 0.976 | 0.9 |
| F5GX71 | Mannose-6-phosphate isomerase OS=Homo sapiens OX=9606 GN=MPI PE=1 SV=1 | 1.042 | 0.863796857 | 0.941 | 0.7 |
| E9PIB2 | Gasdermin-D (Fragment) OS=Homo sapiens OX=9606 GN=GSDMD PE=1 SV=1 | 0.941 | 0.994361756 | 0.999 | 0.9 |
| Q9BVL2 | Nucleoporin p58/p45 OS=Homo sapiens OX=9606 GN=NUP58 PE=1 SV=1 | 1.004 | 0.902016292 | 0.96 | 0.8 |
| Q9H307 | Pinin OS=Homo sapiens OX=9606 GN=PNN PE=1 SV=5 | 1.069 | 0.80706255 | 0.836 | 0.7 |
| Q9H773 | dCTP pyrophosphatase 1 OS=Homo sapiens OX=9606 GN=DCTPP1 PE=1 SV=1 | 1.126 | 0.980966594 | 1.547 | 0.7 |
| Q7Z6K5 | Arpin OS=Homo sapiens OX=9606 GN=ARPIN PE=1 SV=1 | 0.961 | 0.889482529 | 1.057 | 0.7 |
| E5RGJ2 | ER membrane protein complex subunit 2 (Fragment) OS=Homo sapiens OX=9606 GN=EMC2 PE=1 SV=1 | 1.131 | 0.765235202 | 1 | 0.9 |
| O14920 | Inhibitor of nuclear factor kappa-B kinase subunit beta OS=Homo sapiens OX=9606 GN=IKBKB PE=1 SV=1 | 1.016 | 0.889450577 | 0.999 | 0.7 |
| Q9UHA4 | Ragulator complex protein LAMTOR3 OS=Homo sapiens OX=9606 GN=LAMTOR3 PE=1 SV=1 | 0.93 | 0.766667967 | 0.944 | 0.7 |
| P15927 | Replication protein A 32 kDa subunit OS=Homo sapiens OX=9606 GN=RPA2 PE=1 SV=1 | 1.066 | 0.790136102 | 0.918 | 0.7 |
| O14734 | Acyl-coenzyme A thioesterase 8 OS=Homo sapiens OX=9606 GN=ACOT8 PE=1 SV=1 | 0.973 | 0.996637279 | 1.118 | 0.9 |
| Q9Y450 | HBS1-like protein OS=Homo sapiens OX=9606 GN=HBS1L PE=1 SV=1 | 0.924 | 0.98478736 | 0.883 | 0.7 |
| Q9H009 | Nascent polypeptide-associated complex subunit alpha-2 OS=Homo sapiens OX=9606 GN=NACA2 PE=1 SV=1 | 0.958 | 0.873607677 | 1.109 | 0.7 |
| Q96CV9 | Optineurin OS=Homo sapiens OX=9606 GN=OPTN PE=1 SV=3 | 1.079 | 0.765235202 | 0.981 | 0.8 |
| P52732 | Kinesin-like protein KIF11 OS=Homo sapiens OX=9606 GN=KIF11 PE=1 SV=2 | 0.958 | 0.95980123 | 0.945 | 0.7 |
| X6R4W8 | BUB3-interacting and GLEBS motif-containing protein ZNF207 OS=Homo sapiens OX=9606 GN=ZNF207 PE=1 SV=1 | 0.961 | 0.991801854 | 0.936 | 0.7 |
| X6R674 | Influenza virus NS1A-binding protein OS=Homo sapiens OX=9606 GN=IVNS1ABP PE=1 SV=1 | 1.156 | 0.750023665 | 1.322 | 0.7 |
| O95674 | Phosphatidate cytidylyltransferase 2 OS=Homo sapiens OX=9606 GN=CDS2 PE=1 SV=1 | 1.066 | 0.765235202 | 1.023 | 0.7 |
| Q8TDW0 | Volume-regulated anion channel subunit LRRC8C OS=Homo sapiens OX=9606 GN=LRRC8C PE=1 SV=2 | 0.992 | 0.9962822 | 0.926 | 0.7 |
| Q6YN16 | Hydroxysteroid dehydrogenase-like protein 2 OS=Homo sapiens OX=9606 GN=HSDL2 PE=1 SV=1 | 0.88 | 0.765235202 | 0.935 | 0.7 |
| Q9NTX5 | Ethylmalonyl-CoA decarboxylase OS=Homo sapiens OX=9606 GN=ECHDC1 PE=1 SV=2 | 0.995 | 0.997835552 | 0.925 | 0.8 |
| Q9UPY6 | Wiskott-Aldrich syndrome protein family member 3 OS=Homo sapiens OX=9606 GN=WASF3 PE=1 SV=2 | 0.867 | 0.761475165 | 1.193 | 0.9 |
| E9PHH9 | RNA polymerase III subunit C3 OS=Homo sapiens OX=9606 GN=POLR3C PE=1 SV=1 | 1.008 | 0.984200071 | 0.967 | 0.8 |
| Q9NYH9 | U3 small nucleolar RNA-associated protein 6 homolog OS=Homo sapiens OX=9606 GN=UTP6 PE=2 SV=2 | 1.066 | 0.819646566 | 0.901 | 0.7 |
| Q92835 | Phosphatidylinositol 3,4,5-trisphosphate 5-phosphatase 1 OS=Homo sapiens OX=9606 GN=INPP5D PE=1 SV=2 | 1.065 | 0.876510992 | 0.877 | 0.7 |
| Q8IZL8 | Proline-, glutamic acid- and leucine-rich protein 1 OS=Homo sapiens OX=9606 GN=PELP1 PE=1 SV=2 | 0.994 | 0.984200071 | 0.961 | 0.7 |
| Q9NXG2 | THUMP domain-containing protein 1 OS=Homo sapiens OX=9606 GN=THUMPD1 PE=1 SV=2 | 0.974 | 0.906273459 | 0.842 | 0.7 |
| Q8IWA0 | WD repeat-containing protein 75 OS=Homo sapiens OX=9606 GN=WDR75 PE=1 SV=1 | 0.958 | 0.985324977 | 0.92 | 0.8 |
| Q5VT25 | Serine/threonine-protein kinase MRCK alpha OS=Homo sapiens OX=9606 GN=CDC42BPA PE=1 SV=1 | 0.992 | 0.985324977 | 0.859 | 0.7 |
| Q5VZ89 | DENN domain-containing protein 4C OS=Homo sapiens OX=9606 GN=DENND4C PE=1 SV=3 | 1.123 | 0.954224541 | 0.992 | 0.9 |
| P33947 | ER lumen protein-retaining receptor 2 OS=Homo sapiens OX=9606 GN=KDELR2 PE=1 SV=1 | 0.84 | 0.869995963 | 0.842 | 0.8 |
| P78539 | Sushi repeat-containing protein SRPX OS=Homo sapiens OX=9606 GN=SRPX PE=1 SV=1 | 1.004 | 0.978562349 | 0.932 | 0.8 |
| Q9UJW0 | Dynactin subunit 4 OS=Homo sapiens OX=9606 GN=DCTN4 PE=1 SV=1 | 1.042 | 0.809976184 | 0.98 | 0.7 |
| J3QRU7 | Biogenesis of lysosome-related organelles complex 1 subunit 2 (Fragment) OS=Homo sapiens OX=9606 GN=BLOC1S2 PE=1 SV=1 | 1.015 | 0.924563362 | 0.829 | 0.7 |
| K7EML4 | WD repeat-containing protein 18 (Fragment) OS=Homo sapiens OX=9606 GN=WDR18 PE=1 SV=1 | 1.058 | 0.642276967 | 0.867 | 0.7 |

| | | | | | |
|---|---|---|---|---|---|
| P07204 | Thrombomodulin OS=Homo sapiens OX=9606 GN=THBD PE=1 SV=2 | 1.053 | 0.896983349 | 0.95 | 0.9 |
| Q9Y6D6 | Brefeldin A-inhibited guanine nucleotide-exchange protein 1 OS=Homo sapiens OX=9606 GN=ARFGEF1 PE=1 SV=2 | 1.134 | 0.848975923 | 0.881 | 0.7 |
| Q9P253 | Vacuolar protein sorting-associated protein 18 homolog OS=Homo sapiens OX=9606 GN=VPS18 PE=1 SV=2 | 1.075 | 0.916842788 | 0.996 | 0.7 |
| K7EPJ1 | Ubiquitin-conjugating enzyme E2 S (Fragment) OS=Homo sapiens OX=9606 GN=UBE2S PE=1 SV=1 | 1.071 | 0.994361756 | 1.048 | 0.9 |
| O43314 | Inositol hexakisphosphate and diphosphoinositol-pentakisphosphate kinase 2 OS=Homo sapiens OX=9606 GN=PPIP5K2 PE=1 SV= | 1.406 | 0.703015003 | 1.03 | 0.9 |
| O43684 | Mitotic checkpoint protein BUB3 OS=Homo sapiens OX=9606 GN=BUB3 PE=1 SV=1 | 1.056 | 0.816673047 | 1.027 | 0. |
| B4DJV5 | Periodic tryptophan protein 1 homolog OS=Homo sapiens OX=9606 GN=PWP1 PE=1 SV=1 | 0.972 | 0.968322408 | 0.989 | 0.9 |
| P29084 | Transcription initiation factor IIE subunit beta OS=Homo sapiens OX=9606 GN=GTF2E2 PE=1 SV=1 | 1.181 | 0.765235202 | 0.945 | 0.9 |
| O96013 | Serine/threonine-protein kinase PAK 4 OS=Homo sapiens OX=9606 GN=PAK4 PE=1 SV=1 | 0.876 | 0.766667967 | 0.867 | 0.9 |
| Q5W011 | Splicing factor 45 (Fragment) OS=Homo sapiens OX=9606 GN=RBM17 PE=1 SV=1 | 1.009 | 0.9962822 | 0.865 | 0.7 |
| Q9H2D6 | TRIO and F-actin-binding protein OS=Homo sapiens OX=9606 GN=TRIOBP PE=1 SV=3 | 1.049 | 0.815614436 | 0.937 | 0.7 |
| P41227 | N-alpha-acetyltransferase 10 OS=Homo sapiens OX=9606 GN=NAA10 PE=1 SV=1 | 0.982 | 0.925929947 | 1.133 | 0.7 |
| X6R700 | Chromatin target of PRMT1 protein OS=Homo sapiens OX=9606 GN=CHTOP PE=1 SV=1 | 1.311 | 0.750023665 | 0.879 | 0.7 |
| H0YIW3 | HEAT repeat-containing protein 5A (Fragment) OS=Homo sapiens OX=9606 GN=HEATR5A PE=1 SV=1 | 1.06 | 0.920074608 | 0.827 | 0.7 |
| P29466 | Caspase-1 OS=Homo sapiens OX=9606 GN=CASP1 PE=1 SV=1 | 0.983 | 0.913279108 | 0.846 | 0.7 |
| A0A0D9SF70 | ADP-ribosylation factor GTPase-activating protein 2 OS=Homo sapiens OX=9606 GN=ARFGAP2 PE=1 SV=1 | 1.118 | 0.761475165 | 0.986 | 0.9 |
| Q6IQ49 | Replication stress response regulator SDE2 OS=Homo sapiens OX=9606 GN=SDE2 PE=1 SV=1 | 1.105 | 0.80706255 | 0.709 | 0.7 |
| E9PQY2 | Prefoldin subunit 4 OS=Homo sapiens OX=9606 GN=PFDN4 PE=1 SV=1 | 0.995 | 0.886561233 | 1.028 | 0.7 |
| E9PFL3 | Dehydrogenase/reductase SDR family member 4-like 2 OS=Homo sapiens OX=9606 GN=DHRS4L2 PE=1 SV=1 | 0.973 | 0.904500837 | 1 | 0.9 |
| O43583 | Density-regulated protein OS=Homo sapiens OX=9606 GN=DENR PE=1 SV=2 | 0.994 | 0.984805107 | 1.03 | 0.8 |
| B7WP74 | Pre-mRNA-splicing factor CWC22 homolog (Fragment) OS=Homo sapiens OX=9606 GN=CWC22 PE=1 SV=1 | | | | |
| F5GZ97 | Coiled-coil domain-containing protein 53 OS=Homo sapiens OX=9606 GN=WASHC3 PE=1 SV=1 | 1.035 | 0.750023665 | 0.998 | 0.9 |
| F8WEE7 | SEC14-like protein 2 OS=Homo sapiens OX=9606 GN=SEC14L2 PE=1 SV=1 | 1.136 | 0.754126086 | 1.22 | 0.7 |
| A0A1B0GUS7 | Protein unc-13 homolog B OS=Homo sapiens OX=9606 GN=UNC13B PE=1 SV=1 | 1.065 | 0.972073425 | 1.135 | 0. |
| A0A2Y9D025 | Complex I-B14 OS=Homo sapiens OX=9606 GN=NDUFA6 PE=1 SV=1 | 0.954 | 0.750023665 | 1.062 | 0.7 |
| Q0VDF9 | Heat shock 70 kDa protein 14 OS=Homo sapiens OX=9606 GN=HSPA14 PE=1 SV=1 | 1.097 | 0.882454148 | 0.959 | 0.8 |
| H3BQQ2 | E3 ubiquitin-protein ligase ZNF598 OS=Homo sapiens OX=9606 GN=ZNF598 PE=1 SV=1 | 1.055 | 0.904500837 | 1.168 | 0.7 |
| A0A0A0MR02 | Outer mitochondrial membrane protein porin 2 (Fragment) OS=Homo sapiens OX=9606 GN=VDAC2 PE=1 SV=1 | 0.956 | 0.766667967 | 0.866 | 0.7 |
| H7BZ28 | TRAF family member-associated NF-kappa-B activator (Fragment) OS=Homo sapiens OX=9606 GN=TANK PE=1 SV=1 | 1.239 | 0.765235202 | 0.901 | 0.7 |
| Q96AY4 | Tetratricopeptide repeat protein 28 OS=Homo sapiens OX=9606 GN=TTC28 PE=1 SV=4 | 1.183 | 0.765235202 | 0.822 | 0.7 |
| O95236 | Apolipoprotein L3 OS=Homo sapiens OX=9606 GN=APOL3 PE=1 SV=3 | 0.868 | 0.983953845 | 0.952 | 0.7 |
| Q96JM3 | Chromosome alignment-maintaining phosphoprotein 1 OS=Homo sapiens OX=9606 GN=CHAMP1 PE=1 SV=2 | 0.993 | 0.926688896 | 0.918 | 0.7 |
| Q9H2P9 | Diphthine methyl ester synthase OS=Homo sapiens OX=9606 GN=DPH5 PE=1 SV=2 | 0.933 | 0.954224541 | 0.942 | 0.9 |
| Q8IVG5 | Sterile alpha motif domain-containing protein 9-like OS=Homo sapiens OX=9606 GN=SAMD9L PE=1 SV=2 | 1.108 | 0.9252392 | 0.877 | 0.7 |
| B0QYK0 | RNA-binding protein EWS OS=Homo sapiens OX=9606 GN=EWSR1 PE=1 SV=1 | 1.1 | 0.765235202 | 1.007 | 0. |
| Q9Y2L1 | Exosome complex exonuclease RRP44 OS=Homo sapiens OX=9606 GN=DIS3 PE=1 SV=2 | 1.026 | 0.86207051 | 1.068 | 0.7 |
| Q96I24 | Far upstream element-binding protein 3 OS=Homo sapiens OX=9606 GN=FUBP3 PE=1 SV=2 | | | | |
| M0QWZ7 | Seryl-tRNA synthetase OS=Homo sapiens OX=9606 GN=SARS2 PE=1 SV=1 | 0.929 | 0.750023665 | 0.904 | 0.7 |
| Q9UL25 | Ras-related protein Rab-21 OS=Homo sapiens OX=9606 GN=RAB21 PE=1 SV=3 | 1.04 | 0.765235202 | 0.984 | 0.7 |
| Q96ND0 | Protein FAM210A OS=Homo sapiens OX=9606 GN=FAM210A PE=1 SV=2 | 1 | 0.975477573 | 0.912 | 0.7 |
| Q9P2D3 | HEAT repeat-containing protein 5B OS=Homo sapiens OX=9606 GN=HEATR5B PE=1 SV=2 | 1.011 | 0.985324977 | 0.823 | 0.7 |
| E9PJ93 | Ubiquitin carboxyl-terminal hydrolase 48 (Fragment) OS=Homo sapiens OX=9606 GN=USP48 PE=1 SV=8 | 0.983 | 0.827451813 | 0.888 | 0.7 |
| C9JAW5 | HIG1 domain-containing protein OS=Homo sapiens OX=9606 PE=4 SV=1 | 0.832 | 0.756935672 | 0.821 | 0.7 |
| A0A0A0MSK5 | Torsin-1A-interacting protein 1 OS=Homo sapiens OX=9606 GN=TOR1AIP1 PE=1 SV=1 | 0.929 | 0.826889527 | 0.854 | 0.7 |
| H0YJA4 | Chromodomain-helicase-DNA-binding protein 8 (Fragment) OS=Homo sapiens OX=9606 GN=CHD8 PE=1 SV=2 | 0.852 | 0.766667967 | 0.78 | 0. |
| D6REA0 | Glutamyl-tRNA(Gln) amidotransferase subunit B, mitochondrial OS=Homo sapiens OX=9606 GN=GATB PE=1 SV=1 | 1.032 | 0.991211283 | 0.804 | 0.9 |
| M0QY80 | Persulfide dioxygenase ETHE1, mitochondrial OS=Homo sapiens OX=9606 GN=ETHE1 PE=1 SV=1 | 1.023 | 0.948413966 | 0.969 | 0.7 |
| B1ANE3 | Methionine synthase (Fragment) OS=Homo sapiens OX=9606 GN=MTR PE=1 SV=1 | 1.191 | 0.765235202 | 0.994 | 0.9 |
| Q13217 | DnaJ homolog subfamily C member 3 OS=Homo sapiens OX=9606 GN=DNAJC3 PE=1 SV=1 | 0.984 | 0.869995963 | 0.899 | 0.7 |
| Q13547 | Histone deacetylase 1 OS=Homo sapiens OX=9606 GN=HDAC1 PE=1 SV=1 | 1.022 | 0.80706255 | 0.868 | 0.7 |
| Q56VL3 | OCIA domain-containing protein 2 OS=Homo sapiens OX=9606 GN=OCIAD2 PE=1 SV=1 | 0.925 | 0.765235202 | 0.787 | 0.7 |
| Q96PP9 | Guanylate-binding protein 4 OS=Homo sapiens OX=9606 GN=GBP4 PE=2 SV=2 | 0.932 | 0.961249803 | 0.952 | 0.9 |
| P52630 | Signal transducer and activator of transcription 2 OS=Homo sapiens OX=9606 GN=STAT2 PE=1 SV=1 | 0.888 | 0.86414952 | 0.958 | 0.8 |
| P01137 | Transforming growth factor beta-1 proprotein OS=Homo sapiens OX=9606 GN=TGFB1 PE=1 SV=2 | 1.037 | 0.985324977 | 0.959 | 0.7 |
| F5H658 | RNA helicase OS=Homo sapiens OX=9606 GN=DHX8 PE=1 SV=1 | 1.073 | 0.9962822 | 0.981 | 0.8 |
| Q96QU8 | Exportin-6 OS=Homo sapiens OX=9606 GN=XPO6 PE=1 SV=1 | | | | |
| O43818 | U3 small nucleolar RNA-interacting protein 2 OS=Homo sapiens OX=9606 GN=RRP9 PE=1 SV=1 | 0.779 | 0.750023665 | 0.998 | 0.9 |
| A0A087WZT0 | Protein Smaug homolog 2 OS=Homo sapiens OX=9606 GN=SAMD4B PE=1 SV=1 | 0.904 | 0.809976184 | 0.87 | 0.7 |
| E7EW35 | Phosphoribosyl pyrophosphate synthase-associated protein 2 OS=Homo sapiens OX=9606 GN=PRPSAP2 PE=1 SV=1 | 0.939 | 0.785314375 | 1.042 | 0.7 |
| Q2VPB7 | AP-5 complex subunit beta-1 OS=Homo sapiens OX=9606 GN=AP5B1 PE=1 SV=4 | 0.807 | 0.750023665 | 1.171 | 0.9 |
| P53365 | Arfaptin-2 OS=Homo sapiens OX=9606 GN=ARFIP2 PE=1 SV=1 | 1.177 | 0.80706255 | 1.104 | 0.7 |
| A0A087WZK0 | Deoxyhypusine synthase OS=Homo sapiens OX=9606 GN=DHPS PE=1 SV=1 | 1.047 | 0.798032881 | 1.003 | 0.8 |
| A0A2R8YFD4 | E3 ubiquitin-protein ligase CBL OS=Homo sapiens OX=9606 GN=CBLB PE=1 SV=1 | 1.089 | 0.869995963 | 0.873 | 0.7 |
| Q16540 | 39S ribosomal protein L23, mitochondrial OS=Homo sapiens OX=9606 GN=MRPL23 PE=1 SV=1 | 0.995 | 0.916363276 | 0.937 | 0.7 |
| Q05519 | Serine/arginine-rich splicing factor 11 OS=Homo sapiens OX=9606 GN=SRSF11 PE=1 SV=1 | 1.104 | 0.765235202 | 0.962 | 0.7 |
| G3V5V3 | Nuclear export mediator factor NEMF (Fragment) OS=Homo sapiens OX=9606 GN=NEMF PE=1 SV=8 | 1.151 | 0.765235202 | 1.049 | 0.9 |
| Q13057 | Bifunctional coenzyme A synthase OS=Homo sapiens OX=9606 GN=COASY PE=1 SV=4 | 0.968 | 0.848746927 | 0.992 | 0.9 |
| Q15172 | Serine/threonine-protein phosphatase 2A 56 kDa regulatory subunit alpha isoform OS=Homo sapiens OX=9606 GN=PPP2R5A PE= | 0.781 | 0.750023665 | 0.9 | 0.7 |
| P16144 | Integrin beta-4 OS=Homo sapiens OX=9606 GN=ITGB4 PE=1 SV=5 | 1.025 | 0.905369216 | 0.963 | 0.7 |
| Q96EY7 | Pentatricopeptide repeat domain-containing protein 3, mitochondrial OS=Homo sapiens OX=9606 GN=PTCD3 PE=1 SV=3 | 0.998 | 0.904500837 | 0.946 | 0.9 |
| Q6P3X3 | Tetratricopeptide repeat protein 27 OS=Homo sapiens OX=9606 GN=TTC27 PE=1 SV=1 | 1.481 | 0.766667967 | 0.799 | 0.7 |
| Q9UQ88 | Cyclin-dependent kinase 11A OS=Homo sapiens OX=9606 GN=CDK11A PE=1 SV=4 | | | | |
| O00170 | AH receptor-interacting protein OS=Homo sapiens OX=9606 GN=AIP PE=1 SV=2 | 1.008 | 0.996578046 | 0.908 | 0.7 |
| G3V3A4 | SNW domain-containing protein 1 OS=Homo sapiens OX=9606 GN=SNW1 PE=1 SV=1 | 1.089 | 0.765235202 | 0.929 | 0.7 |
| P56385 | ATP synthase subunit e, mitochondrial OS=Homo sapiens OX=9606 GN=ATP5ME PE=1 SV=2 | 0.872 | 0.786635798 | 0.764 | 0.7 |
| Q8N2R8 | Protein FAM43A OS=Homo sapiens OX=9606 GN=FAM43A PE=1 SV=2 | 0.967 | 0.978562349 | 1.052 | 0.8 |
| O75600 | 2-amino-3-ketobutyrate coenzyme A ligase, mitochondrial OS=Homo sapiens OX=9606 GN=GCAT PE=1 SV=1 | 1.004 | 0.913279108 | 0.931 | 0.9 |
| Q9NZM3 | Intersectin-2 OS=Homo sapiens OX=9606 GN=ITSN2 PE=1 SV=3 | 0.919 | 0.892311206 | 0.928 | 0.9 |
| Q8IY81 | pre-rRNA 2'-O-ribose RNA methyltransferase FTSJ3 OS=Homo sapiens OX=9606 GN=FTSJ3 PE=1 SV=2 | 0.986 | 0.899593149 | 0.927 | 0.7 |
| Q9BSY9 | Deubiquitinase DESI2 OS=Homo sapiens OX=9606 GN=DESI2 PE=1 SV=1 | 0.913 | 0.765235202 | 0.989 | 0.8 |
| Q14165 | Malectin OS=Homo sapiens OX=9606 GN=MLEC PE=1 SV=1 | 1.041 | 0.966586493 | 0.924 | 0.7 |
| P08493 | Matrix Gla protein OS=Homo sapiens OX=9606 GN=MGP PE=1 SV=2 | 0.998 | 0.908555656 | 1.121 | 0.8 |
| Q9C0H2 | Protein tweety homolog 3 OS=Homo sapiens OX=9606 GN=TTYH3 PE=1 SV=3 | 1.169 | 0.765235202 | 0.926 | 0.9 |
| O43493 | Trans-Golgi network integral membrane protein 2 OS=Homo sapiens OX=9606 GN=TGOLN2 PE=1 SV=4 | 0.956 | 0.944284199 | 1.277 | 0.7 |
| Q99717 | Mothers against decapentaplegic homolog 5 OS=Homo sapiens OX=9606 GN=SMAD5 PE=1 SV=1 | 0.965 | 0.912604818 | 0.978 | 0. |
| P17302 | Gap junction alpha-1 protein OS=Homo sapiens OX=9606 GN=GJA1 PE=1 SV=2 | 1.236 | 0.750023665 | 1.267 | 0.7 |
| Q9NXJ5 | Pyroglutamyl-peptidase 1 OS=Homo sapiens OX=9606 GN=PGPEP1 PE=1 SV=1 | 1.057 | 0.999280989 | 0.895 | 0.7 |
| A2A2F0 | Ral GTPase-activating protein subunit beta (Fragment) OS=Homo sapiens OX=9606 GN=RALGAPB PE=1 SV=1 | 0.936 | 0.924992773 | 1.12 | 0.8 |
| Q9Y5Q9 | General transcription factor 3C polypeptide 3 OS=Homo sapiens OX=9606 GN=GTF3C3 PE=1 SV=1 | 1.028 | 0.985763982 | 0.958 | 0.8 |
| B3KSI3 | Branched-chain-amino-acid aminotransferase OS=Homo sapiens OX=9606 GN=BCAT2 PE=1 SV=1 | 0.959 | 0.908555656 | 0.962 | 0.7 |
| Q96QD8 | Sodium-coupled neutral amino acid transporter 2 OS=Homo sapiens OX=9606 GN=SLC38A2 PE=1 SV=2 | 1.088 | 0.950763163 | 0.755 | 0.7 |
| O95197 | Reticulon-3 OS=Homo sapiens OX=9606 GN=RTN3 PE=1 SV=2 | 1.021 | 0.984200071 | 1.014 | 0.9 |
| Q8IYS2 | Uncharacterized protein KIAA2013 OS=Homo sapiens OX=9606 GN=KIAA2013 PE=1 SV=1 | 1.056 | 0.984805107 | 0.92 | 0.8 |
| H7C1X9 | CCR4-NOT transcription complex subunit 10 (Fragment) OS=Homo sapiens OX=9606 GN=CNOT10 PE=1 SV=1 | 0.961 | 0.985324977 | 0.96 | 0. |
| F5GWI4 | Adenosine deaminase OS=Homo sapiens OX=9606 GN=ADA PE=1 SV=1 | 1.018 | 0.985324977 | 0.966 | 0.7 |
| Q86TU7 | Actin-histidine N-methyltransferase OS=Homo sapiens OX=9606 GN=SETD3 PE=1 SV=1 | 0.939 | 0.85315742 | 0.996 | 0.9 |

| | | | | | |
|---|---|---|---|---|---|
| Q92575 | UBX domain-containing protein 4 OS=Homo sapiens OX=9606 GN=UBXN4 PE=1 SV=2 | 1.021 | 0.773977513 | 0.964 | 0.7 |
| Q8NFQ8 | Torsin-1A-interacting protein 2 OS=Homo sapiens OX=9606 GN=TOR1AIP2 PE=1 SV=1 | | | | |
| A0A087WVU6 | E3 ubiquitin-protein ligase HECTD1 (Fragment) OS=Homo sapiens OX=9606 GN=HECTD1 PE=1 SV=1 | 1.014 | 0.9962822 | 1.017 | 0.9 |
| P27707 | Deoxycytidine kinase OS=Homo sapiens OX=9606 GN=DCK PE=1 SV=1 | | | | |
| P13807 | Glycogen [starch] synthase, muscle OS=Homo sapiens OX=9606 GN=GYS1 PE=1 SV=2 | 0.96 | 0.916363276 | 1.023 | 0.9 |
| Q15392 | Delta(24)-sterol reductase OS=Homo sapiens OX=9606 GN=DHCR24 PE=1 SV=2 | 0.865 | 0.937953124 | 0.881 | 0.7 |
| Q9H1E5 | Thioredoxin-related transmembrane protein 4 OS=Homo sapiens OX=9606 GN=TMX4 PE=1 SV=1 | 1.023 | 0.869995963 | 0.936 | 0.7 |
| A8K0Z3 | WASH complex subunit 1 OS=Homo sapiens OX=9606 GN=WASHC1 PE=1 SV=2 | 1.106 | 0.830179531 | 0.964 | 0.8 |
| J3KNF4 | Copper chaperone for superoxide dismutase OS=Homo sapiens OX=9606 GN=CCS PE=1 SV=1 | 1.011 | 0.983953845 | 1 | 0.9 |
| Q9GZV5 | WW domain-containing transcription regulator protein 1 OS=Homo sapiens OX=9606 GN=WWTR1 PE=1 SV=1 | 1.058 | 0.855041013 | 0.815 | 0.8 |
| Q16204 | Coiled-coil domain-containing protein 6 OS=Homo sapiens OX=9606 GN=CCDC6 PE=1 SV=2 | 1.026 | 0.902766821 | 0.948 | 0.7 |
| F5H538 | Mitogen-activated protein kinase kinase kinase 4 OS=Homo sapiens OX=9606 GN=MAP3K4 PE=1 SV=1 | 0.956 | 0.87017509 | 0.885 | 0.7 |
| K7ES07 | Thioredoxin-like protein 4A OS=Homo sapiens OX=9606 GN=TXNL4A PE=1 SV=1 | 0.891 | 0.750023665 | 0.941 | 0.7 |
| E9PC15 | Acylglycerol kinase, mitochondrial OS=Homo sapiens OX=9606 GN=AGK PE=1 SV=1 | 1.123 | 0.865165562 | 0.783 | 0.7 |
| Q9BVP2 | Guanine nucleotide-binding protein-like 3 OS=Homo sapiens OX=9606 GN=GNL3 PE=1 SV=2 | 1.065 | 0.785314375 | 0.902 | 0.9 |
| G3V4E4 | Legumain OS=Homo sapiens OX=9606 GN=LGMN PE=1 SV=1 | 1.056 | 0.876510992 | 1.036 | 0.9 |
| A0A0B4J286 | Trafficking protein particle complex subunit 2-like protein OS=Homo sapiens OX=9606 GN=TRAPPC2L PE=1 SV=1 | 1.078 | 0.859846322 | 0.997 | 0.8 |
| Q9Y6I3 | Epsin-1 OS=Homo sapiens OX=9606 GN=EPN1 PE=1 SV=2 | 0.994 | 0.901934028 | 0.961 | 0.7 |
| A8MYR4 | Transcription initiation factor IIA gamma chain OS=Homo sapiens OX=9606 GN=GTF2A2 PE=1 SV=1 | 1.116 | 0.902016292 | 0.958 | 0.9 |
| A2ACR1 | Proteasome subunit beta OS=Homo sapiens OX=9606 GN=PSMB9 PE=1 SV=1 | 0.964 | 0.889482529 | 0.577 | 0.7 |
| Q8IVD9 | NudC domain-containing protein 3 OS=Homo sapiens OX=9606 GN=NUDCD3 PE=1 SV=3 | 0.948 | 0.994833431 | 0.824 | 0.7 |
| Q7Z3B4 | Nucleoporin p54 OS=Homo sapiens OX=9606 GN=NUP54 PE=1 SV=2 | 1.028 | 0.982290035 | 1.004 | 0.9 |
| P51452 | Dual specificity protein phosphatase 3 OS=Homo sapiens OX=9606 GN=DUSP3 PE=1 SV=1 | 1.015 | 0.985330338 | 0.916 | 0.7 |
| Q03154 | Aminoacylase-1 OS=Homo sapiens OX=9606 GN=ACY1 PE=1 SV=1 | 0.929 | 0.93716325 | 0.914 | 0.7 |
| A0A5H1ZRP4 | Negative elongation factor B OS=Homo sapiens OX=9606 GN=NELFB PE=1 SV=1 | 1.078 | 0.840158726 | 0.976 | 0.7 |
| A0A087WXZ7 | HEAT repeat-containing protein 6 OS=Homo sapiens OX=9606 GN=HEATR6 PE=1 SV=1 | 1.063 | 0.765235202 | 0.996 | 0.8 |
| Q8N3F8 | MICAL-like protein 1 OS=Homo sapiens OX=9606 GN=MICALL1 PE=1 SV=2 | 0.994 | 0.949254122 | 0.94 | 0.7 |
| H7C173 | Mitotic-spindle organizing protein 2B (Fragment) OS=Homo sapiens OX=9606 GN=MZT2B PE=1 SV=1 | 1.04 | 0.961249803 | 0.976 | 0.9 |
| Q9H496 | Torsin-1A-interacting protein 2, isoform IFRG15 OS=Homo sapiens OX=9606 GN=TOR1AIP2 PE=1 SV=1 | 0.93 | 0.95980123 | 1.053 | 0.8 |
| C9JW51 | Protein FAM107B (Fragment) OS=Homo sapiens OX=9606 GN=FAM107B PE=1 SV=1 | 1.071 | 0.905369216 | 1.152 | 0.7 |
| A0A2R8YFQ7 | Lambda-crystallin homolog OS=Homo sapiens OX=9606 GN=CRYL1 PE=1 SV=1 | 1.004 | 0.961249803 | 0.897 | 0.7 |
| P22059 | Oxysterol-binding protein 1 OS=Homo sapiens OX=9606 GN=OSBP PE=1 SV=1 | 1.038 | 0.837941467 | 1.014 | 0.9 |
| O15118 | NPC intracellular cholesterol transporter 1 OS=Homo sapiens OX=9606 GN=NPC1 PE=1 SV=2 | 1.03 | 0.95980123 | 0.962 | 0.8 |
| Q16706 | Alpha-mannosidase 2 OS=Homo sapiens OX=9606 GN=MAN2A1 PE=1 SV=2 | 0.929 | 0.750023665 | 0.922 | 0.7 |
| F8WAN9 | GMP reductase OS=Homo sapiens OX=9606 GN=GMPR2 PE=1 SV=1 | 1.039 | 0.942678074 | 1 | 0.8 |
| H7BXQ8 | Armadillo repeat-containing protein 10 OS=Homo sapiens OX=9606 GN=ARMC10 PE=1 SV=1 | | | | |
| C9JJ54 | WD repeat and FYVE domain-containing protein 1 (Fragment) OS=Homo sapiens OX=9606 GN=WDFY1 PE=1 SV=1 | 1.039 | 0.994361756 | 1.113 | 0.8 |
| Q96SK2 | Transmembrane protein 209 OS=Homo sapiens OX=9606 GN=TMEM209 PE=1 SV=2 | 1.411 | 0.835336742 | 0.89 | 0.9 |
| Q9GZZ9 | Ubiquitin-like modifier-activating enzyme 5 OS=Homo sapiens OX=9606 GN=UBA5 PE=1 SV=1 | 1.031 | 0.905470348 | 0.913 | 0.7 |
| Q7RTS9 | Dymeclin OS=Homo sapiens OX=9606 GN=DYM PE=1 SV=1 | 0.915 | 0.765235202 | 0.995 | 0.7 |
| I3L3H6 | Cell division cycle protein 27 homolog (Fragment) OS=Homo sapiens OX=9606 GN=CDC27 PE=1 SV=1 | 1.103 | 0.750023665 | 1.041 | 0.8 |
| Q9C0D5 | Protein TANC1 OS=Homo sapiens OX=9606 GN=TANC1 PE=1 SV=3 | 0.875 | 0.766667967 | 0.855 | 0.7 |
| Q13443 | Disintegrin and metalloproteinase domain-containing protein 9 OS=Homo sapiens OX=9606 GN=ADAM9 PE=1 SV=1 | 0.99 | 0.904500837 | 1.043 | 0.9 |
| Q13523 | Serine/threonine-protein kinase PRP4 homolog OS=Homo sapiens OX=9606 GN=PRPF4B PE=1 SV=3 | 1.063 | 0.827871327 | 0.915 | 0.7 |
| Q9H3K6 | BolA-like protein 2 OS=Homo sapiens OX=9606 GN=BOLA2 PE=1 SV=1 | 0.913 | 0.792509319 | 0.889 | 0.7 |
| Q96F86 | Enhancer of mRNA-decapping protein 3 OS=Homo sapiens OX=9606 GN=EDC3 PE=1 SV=1 | 1.01 | 0.918702104 | 1.011 | 0.8 |
| O15230 | Laminin subunit alpha-5 OS=Homo sapiens OX=9606 GN=LAMA5 PE=1 SV=8 | 1.013 | 0.910192239 | 0.895 | 0.7 |
| P45985 | Dual specificity mitogen-activated protein kinase kinase 4 OS=Homo sapiens OX=9606 GN=MAP2K4 PE=1 SV=1 | 0.854 | 0.763877559 | 0.874 | 0.7 |
| G8JLK1 | USE1-like protein OS=Homo sapiens OX=9606 GN=USE1 PE=1 SV=2 | 1.037 | 0.963059765 | 0.876 | 0.7 |
| Q969M3 | Protein YIPF5 OS=Homo sapiens OX=9606 GN=YIPF5 PE=1 SV=1 | 1.023 | 0.869995963 | 0.87 | 0.7 |
| Q96JC1 | Vam6/Vps39-like protein OS=Homo sapiens OX=9606 GN=VPS39 PE=1 SV=2 | 0.967 | 0.887475968 | 0.935 | 0.7 |
| O75582 | Ribosomal protein S6 kinase alpha-5 OS=Homo sapiens OX=9606 GN=RPS6KA5 PE=1 SV=1 | 0.954 | 0.680740328 | 0.854 | 0.7 |
| A0A087WVZ9 | DNA-directed RNA polymerase II subunit E OS=Homo sapiens OX=9606 GN=POLR2E PE=1 SV=1 | 0.867 | 0.750023665 | 1.032 | 0.8 |
| F8VVY5 | Death domain-containing protein CRADD OS=Homo sapiens OX=9606 GN=CRADD PE=1 SV=1 | 0.958 | 0.963059765 | 0.874 | 0.7 |
| H0Y9L5 | Exosome complex component RRP45 (Fragment) OS=Homo sapiens OX=9606 GN=EXOSC9 PE=1 SV=1 | 1.006 | 0.927148669 | 0.879 | 0.7 |
| P04183 | Thymidine kinase, cytosolic OS=Homo sapiens OX=9606 GN=TK1 PE=1 SV=2 | 0.918 | 0.954224541 | 1.363 | 0.7 |
| Q8WTS6 | Histone-lysine N-methyltransferase SETD7 OS=Homo sapiens OX=9606 GN=SETD7 PE=1 SV=1 | 0.972 | 0.894951666 | 0.823 | 0.7 |
| Q9H2V7 | Protein spinster homolog 1 OS=Homo sapiens OX=9606 GN=SPNS1 PE=1 SV=1 | 1.22 | 0.869995963 | 0.9 | 0.7 |
| A0A0U1RQT9 | Synaptophysin-like protein 1 (Fragment) OS=Homo sapiens OX=9606 GN=SYPL1 PE=1 SV=1 | 1.048 | 0.880371237 | 0.936 | 0.7 |
| Q8IZH2 | 5'-3' exoribonuclease 1 OS=Homo sapiens OX=9606 GN=XRN1 PE=1 SV=1 | 0.956 | 0.766667967 | 0.799 | 0.7 |
| O96005 | Cleft lip and palate transmembrane protein 1 OS=Homo sapiens OX=9606 GN=CLPTM1 PE=1 SV=1 | | | | |
| Q96CT7 | Coiled-coil domain-containing protein 124 OS=Homo sapiens OX=9606 GN=CCDC124 PE=1 SV=1 | 1.163 | 0.84941721 | 0.965 | 0.7 |
| Q68E01 | Integrator complex subunit 3 OS=Homo sapiens OX=9606 GN=INTS3 PE=1 SV=1 | 1.164 | 0.790808367 | 0.922 | 0.7 |
| Q9GZT8 | NIF3-like protein 1 OS=Homo sapiens OX=9606 GN=NIF3L1 PE=1 SV=2 | 1.045 | 0.780884711 | 0.972 | 0.8 |
| Q15814 | Tubulin-specific chaperone C OS=Homo sapiens OX=9606 GN=TBCC PE=1 SV=2 | 0.928 | 0.750023665 | 0.864 | 0.7 |
| H0Y5B0 | Band 4.1-like protein 2 (Fragment) OS=Homo sapiens OX=9606 GN=EPB41L2 PE=1 SV=1 | 1.186 | 0.765235202 | 1.036 | 0.9 |
| O43150 | Arf-GAP with SH3 domain, ANK repeat and PH domain-containing protein 2 OS=Homo sapiens OX=9606 GN=ASAP2 PE=1 SV=3 | 1.116 | 0.827451813 | 0.862 | 0.9 |
| H0YMG4 | E3 ubiquitin-protein ligase RNF31 (Fragment) OS=Homo sapiens OX=9606 GN=RNF31 PE=1 SV=2 | 0.928 | 0.924631778 | 1.032 | 0.9 |
| P61964 | WD repeat-containing protein 5 OS=Homo sapiens OX=9606 GN=WDR5 PE=1 SV=1 | 1.063 | 0.780884711 | 0.944 | 0.7 |
| Q9BXW7 | Haloacid dehalogenase-like hydrolase domain-containing 5 OS=Homo sapiens OX=9606 GN=HDHD5 PE=1 SV=1 | 1.067 | 0.79053595 | 0.941 | 0.7 |
| H9KV45 | Ubiquitin-conjugating enzyme E2 D3 OS=Homo sapiens OX=9606 GN=UBE2D3 PE=1 SV=1 | 0.965 | 0.887475968 | 1.089 | 0.7 |
| A0A2R8Y855 | SWI/SNF-related matrix-associated actin-dependent regulator of chromatin subfamily E member 1 OS=Homo sapiens OX=9606 GI | 0.95 | 0.906273459 | 0.996 | 0.7 |
| Q12974 | Protein tyrosine phosphatase type IVA 2 OS=Homo sapiens OX=9606 GN=PTP4A2 PE=1 SV=1 | 0.835 | 0.829390484 | 0.951 | 0.7 |
| Q9Y5K6 | CD2-associated protein OS=Homo sapiens OX=9606 GN=CD2AP PE=1 SV=1 | 1.003 | 0.908555656 | 0.921 | 0.7 |
| B5MCF9 | Pescadillo homolog OS=Homo sapiens OX=9606 GN=PES1 PE=1 SV=1 | 1.043 | 0.829390484 | 1.034 | 0.7 |
| Q96JQ0 | Protocadherin-16 OS=Homo sapiens OX=9606 GN=DCHS1 PE=1 SV=1 | 0.814 | 0.765235202 | 0.825 | 0.7 |
| O00483 | Cytochrome c oxidase subunit NDUFA4 OS=Homo sapiens OX=9606 GN=NDUFA4 PE=1 SV=1 | 0.971 | 0.902016292 | 0.995 | 0.9 |
| A0A0A0MQS3 | Leydig cell tumor 10 kDa protein homolog (Fragment) OS=Homo sapiens OX=9606 GN=C19orf53 PE=1 SV=1 | 1.089 | 0.765937763 | 1.071 | 0.7 |
| F5H2T0 | IkappaB kinase complex-associated protein OS=Homo sapiens OX=9606 GN=ELP1 PE=1 SV=1 | 1.138 | 0.878901226 | 0.798 | 0.7 |
| P15954 | Cytochrome c oxidase subunit 7C, mitochondrial OS=Homo sapiens OX=9606 GN=COX7C PE=1 SV=1 | 0.954 | 0.94475926 | 0.911 | 0.7 |
| Q8N3U4 | Cohesin subunit SA-2 OS=Homo sapiens OX=9606 GN=STAG2 PE=1 SV=3 | 1.022 | 0.989102644 | 0.892 | 0.7 |
| O15111 | Inhibitor of nuclear factor kappa-B kinase subunit alpha OS=Homo sapiens OX=9606 GN=CHUK PE=1 SV=2 | 1.092 | 0.887719165 | 0.891 | 0.7 |
| Q9H974 | Queuine tRNA-ribosyltransferase accessory subunit 2 OS=Homo sapiens OX=9606 GN=QTRT2 PE=1 SV=1 | 0.898 | 0.829390484 | 0.901 | 0.7 |
| Q9NXH9 | tRNA (guanine(26)-N(2))-dimethyltransferase OS=Homo sapiens OX=9606 GN=TRMT1 PE=1 SV=1 | 1.022 | 0.966586493 | 0.934 | 0.7 |
| S4R3Q6 | Vacuolar protein sorting-associated protein 26A OS=Homo sapiens OX=9606 GN=VPS26A PE=1 SV=1 | 0.825 | 0.765235202 | 0.879 | 0.7 |
| Q5VTR2 | E3 ubiquitin-protein ligase BRE1A OS=Homo sapiens OX=9606 GN=RNF20 PE=1 SV=2 | 1.148 | 0.758476925 | 0.85 | 0.7 |
| Q9UJX3 | Anaphase-promoting complex subunit 7 OS=Homo sapiens OX=9606 GN=ANAPC7 PE=1 SV=4 | 1.298 | 0.750023665 | 0.74 | 0.7 |
| Q96F24 | Nuclear receptor-binding factor 2 OS=Homo sapiens OX=9606 GN=NRBF2 PE=1 SV=1 | 1.244 | 0.750023665 | 0.934 | 0.7 |
| P14927 | Cytochrome b-c1 complex subunit 7 OS=Homo sapiens OX=9606 GN=UQCRB PE=1 SV=2 | 1.072 | 0.941166044 | 0.899 | 0.7 |
| H0YGT8 | Peroxisomal targeting signal 1 receptor (Fragment) OS=Homo sapiens OX=9606 GN=PEX5 PE=1 SV=1 | 1.46 | 0.750023665 | 0.875 | 0.7 |
| Q19T08 | Endothelial cell-specific chemotaxis regulator OS=Homo sapiens OX=9606 GN=ECSCR PE=1 SV=1 | 1.03 | 0.916363276 | 1.01 | 0.7 |
| Q9NYR9 | NF-kappa-B inhibitor-interacting Ras-like protein 2 OS=Homo sapiens OX=9606 GN=NKIRAS2 PE=1 SV=1 | | | | |
| A0A0A0MTB8 | WD repeat-containing protein 36 OS=Homo sapiens OX=9606 GN=WDR36 PE=1 SV=1 | 0.983 | 0.993362873 | 0.854 | 0.7 |
| Q10570 | Cleavage and polyadenylation specificity factor subunit 1 OS=Homo sapiens OX=9606 GN=CPSF1 PE=1 SV=2 | 1.036 | 0.87017509 | 0.934 | 0.7 |
| Q14137 | Ribosome biogenesis protein BOP1 OS=Homo sapiens OX=9606 GN=BOP1 PE=1 SV=2 | 100 | | 0.01 | |
| K7ER08 | RNA-binding protein 42 OS=Homo sapiens OX=9606 GN=RBM42 PE=1 SV=1 | 0.898 | 0.765235202 | 0.868 | 0.7 |

| | | | | | |
|---|---|---|---|---|---|
| H0Y9R3 | WD repeat-containing protein 26 (Fragment) OS=Homo sapiens OX=9606 GN=WDR26 PE=1 SV=1 | | | | |
| Q00013 | 55 kDa erythrocyte membrane protein OS=Homo sapiens OX=9606 GN=MPP1 PE=1 SV=2 | 0.891 | 0.766667967 | 0.888 | 0.7 |
| A0A087WYR0 | Signal recognition particle 19 kDa protein OS=Homo sapiens OX=9606 GN=SRP19 PE=1 SV=1 | 1.054 | 0.780884711 | 0.981 | 0.7 |
| A0A2R8Y7W5 | Cyclin-dependent kinase 13 (Fragment) OS=Homo sapiens OX=9606 GN=CDK13 PE=1 SV=1 | 1.398 | 0.765235202 | 1.072 | 0.9 |
| Q96ST3 | Paired amphipathic helix protein Sin3a OS=Homo sapiens OX=9606 GN=SIN3A PE=1 SV=2 | 1.243 | 0.750023665 | 1.062 | 0.8 |
| A0A590UJQ1 | E3 ubiquitin-protein ligase OS=Homo sapiens OX=9606 GN=ITCH PE=1 SV=1 | 0.99 | 0.886561233 | 0.971 | 0.7 |
| C9JST7 | Protein YIF1A (Fragment) OS=Homo sapiens OX=9606 GN=YIF1A PE=1 SV=2 | 1.064 | 0.750023665 | 0.945 | 0.7 |
| Q6UW63 | Protein O-glucosyltransferase 2 OS=Homo sapiens OX=9606 GN=POGLUT2 PE=1 SV=1 | 1.051 | 0.756287988 | 1.048 | 0.7 |
| Q07617 | Sperm-associated antigen 1 OS=Homo sapiens OX=9606 GN=SPAG1 PE=1 SV=3 | 1.188 | 0.750023665 | 0.993 | 0.9 |
| Q93096 | Protein tyrosine phosphatase type IVA 1 OS=Homo sapiens OX=9606 GN=PTP4A1 PE=1 SV=2 | 0.776 | 0.765235202 | 0.813 | 0.7 |
| W5RWE6 | PX domain-containing protein kinase-like protein OS=Homo sapiens OX=9606 GN=PXK PE=1 SV=1 | 0.941 | 0.794019775 | 0.924 | 0.7 |
| A0A494BZY7 | EEF1A lysine methyltransferase 2 OS=Homo sapiens OX=9606 GN=EEF1AKMT2 PE=1 SV=1 | 0.987 | 0.983953845 | 1.165 | 0.7 |
| Q7Z7C8 | Transcription initiation factor TFIID subunit 8 OS=Homo sapiens OX=9606 GN=TAF8 PE=1 SV=1 | 0.856 | 0.801385256 | 1.056 | 0.7 |
| Q9H8J5 | MANSC domain-containing protein 1 OS=Homo sapiens OX=9606 GN=MANSC1 PE=1 SV=1 | 0.955 | 0.969190229 | 1.04 | 0.[illegible] |
| Q96HD1 | Protein disulfide isomerase CRELD1 OS=Homo sapiens OX=9606 GN=CRELD1 PE=1 SV=3 | 1.018 | 0.984442117 | 0.902 | 0.7 |
| P05496 | ATP synthase F(0) complex subunit C1, mitochondrial OS=Homo sapiens OX=9606 GN=ATP5MC1 PE=1 SV=2 | 1.125 | 0.765235202 | 0.873 | 0.7 |
| A0A0C4DFP4 | N-glycanase 1 (Fragment) OS=Homo sapiens OX=9606 GN=NGLY1 PE=1 SV=1 | 1.008 | 0.9962822 | 0.878 | 0.7 |
| P19447 | General transcription and DNA repair factor IIH helicase subunit XPB OS=Homo sapiens OX=9606 GN=ERCC3 PE=1 SV=1 | 0.978 | 0.9962822 | 0.853 | 0.7 |
| Q9NPI6 | mRNA-decapping enzyme 1A OS=Homo sapiens OX=9606 GN=DCP1A PE=1 SV=3 | 0.909 | 0.978562349 | 0.935 | 0.8 |
| Q15751 | Probable E3 ubiquitin-protein ligase HERC1 OS=Homo sapiens OX=9606 GN=HERC1 PE=1 SV=2 | 0.907 | 0.750023665 | 0.757 | 0.7 |
| O15160 | DNA-directed RNA polymerases I and III subunit RPAC1 OS=Homo sapiens OX=9606 GN=POLR1C PE=1 SV=1 | 1.002 | 0.902016292 | 1.25 | 0.7 |
| Q6NUQ1 | RAD50-interacting protein 1 OS=Homo sapiens OX=9606 GN=RINT1 PE=1 SV=1 | 1.06 | 0.978693882 | 0.926 | 0.8 |
| H3BNT3 | Methyltransferase-like 26 (Fragment) OS=Homo sapiens OX=9606 GN=METTL26 PE=1 SV=1 | 0.99 | 0.882454148 | 0.895 | 0.7 |
| J3KNI1 | Component of oligomeric Golgi complex 4 OS=Homo sapiens OX=9606 GN=COG4 PE=1 SV=1 | 1.062 | 0.886561233 | 2.102 | 0.7 |
| Q13574 | Diacylglycerol kinase zeta OS=Homo sapiens OX=9606 GN=DGKZ PE=1 SV=4 | 0.984 | 0.901934028 | 1.055 | 0.7 |
| Q02978 | Mitochondrial 2-oxoglutarate/malate carrier protein OS=Homo sapiens OX=9606 GN=SLC25A11 PE=1 SV=3 | | | | |
| Q9NVG8 | TBC1 domain family member 13 OS=Homo sapiens OX=9606 GN=TBC1D13 PE=1 SV=3 | 1.098 | 0.765235202 | 0.961 | 0.9 |
| Q96EM0 | Trans-3-hydroxy-L-proline dehydratase OS=Homo sapiens OX=9606 GN=L3HYPDH PE=1 SV=2 | 0.912 | 0.939024443 | 0.978 | 0.7 |
| Q15269 | Periodic tryptophan protein 2 homolog OS=Homo sapiens OX=9606 GN=PWP2 PE=2 SV=2 | 1.103 | 0.840133488 | 0.94 | 0.7 |
| G3V2S6 | V-type proton ATPase subunit D OS=Homo sapiens OX=9606 GN=ATP6V1D PE=1 SV=1 | 1.143 | 0.765235202 | 0.979 | 0.8 |
| Q96FN4 | Copine-2 OS=Homo sapiens OX=9606 GN=CPNE2 PE=1 SV=3 | 1.007 | 0.987622091 | 0.92 | 0.7 |
| Q4R9M9 | Kinesin family member 1Bbeta isoform II OS=Homo sapiens OX=9606 GN=KIF1B PE=1 SV=1 | 1.003 | 0.984805107 | 1.01 | 0.8 |
| A0A494C0D4 | Protoporphyrinogen oxidase (Fragment) OS=Homo sapiens OX=9606 GN=PPOX PE=1 SV=1 | 1.057 | 0.827085425 | 0.926 | 0.8 |
| Q8ND04 | Protein SMG8 OS=Homo sapiens OX=9606 GN=SMG8 PE=1 SV=1 | 1.064 | 0.785314375 | 1.021 | 0.7 |
| O95059 | Ribonuclease P protein subunit p14 OS=Homo sapiens OX=9606 GN=RPP14 PE=1 SV=3 | 0.926 | 0.776314646 | 0.935 | 0.8 |
| F8WCN7 | Protein BUD31 homolog OS=Homo sapiens OX=9606 GN=BUD31 PE=1 SV=1 | 1.113 | 0.780884711 | 0.933 | 0.9 |
| Q15758 | Neutral amino acid transporter B(0) OS=Homo sapiens OX=9606 GN=SLC1A5 PE=1 SV=2 | 0.951 | 0.835808354 | 1.143 | 0.7 |
| J3KT75 | Mannose-P-dolichol utilization defect 1 protein (Fragment) OS=Homo sapiens OX=9606 GN=MPDU1 PE=1 SV=1 | 0.962 | 0.750023665 | 0.993 | 0.8 |
| A0A0J9YW33 | Syntaxin-3 OS=Homo sapiens OX=9606 GN=STX3 PE=1 SV=1 | 0.978 | 0.983953845 | 0.932 | 0.7 |
| Q17RB0 | Retrotransposon Gag-like protein 8B OS=Homo sapiens OX=9606 GN=RTL8B PE=1 SV=1 | 1.072 | 0.870355966 | 1.046 | 0.9 |
| Q6UXH1 | Protein disulfide isomerase CRELD2 OS=Homo sapiens OX=9606 GN=CRELD2 PE=1 SV=1 | 0.925 | 0.991211283 | 0.959 | 0.7 |
| P05771 | Protein kinase C beta type OS=Homo sapiens OX=9606 GN=PRKCB PE=1 SV=4 | 1.081 | 0.94475926 | 0.993 | 0.9 |
| O00501 | Claudin-5 OS=Homo sapiens OX=9606 GN=CLDN5 PE=1 SV=1 | 0.934 | 0.815614436 | 0.713 | 0.7 |
| P0CG30 | Glutathione S-transferase theta-2B OS=Homo sapiens OX=9606 GN=GSTT2B PE=1 SV=1 | 0.991 | 0.930855729 | 0.98 | 0.8 |
| Q8N3P4 | Vacuolar protein sorting-associated protein 8 homolog OS=Homo sapiens OX=9606 GN=VPS8 PE=1 SV=3 | 0.865 | 0.766667967 | 0.805 | 0.7 |
| F8W0M9 | Secernin-3 (Fragment) OS=Homo sapiens OX=9606 GN=SCRN3 PE=1 SV=1 | 0.997 | 0.885528205 | 0.904 | 0.7 |
| Q9NVH0 | Exonuclease 3'-5' domain-containing protein 2 OS=Homo sapiens OX=9606 GN=EXD2 PE=1 SV=2 | 1.013 | 0.908555656 | 0.893 | 0.7 |
| Q96TC7 | Regulator of microtubule dynamics protein 3 OS=Homo sapiens OX=9606 GN=RMDN3 PE=1 SV=2 | 1.081 | 0.860909277 | 0.978 | 0.8 |
| P0DN79 | Cystathionine beta-synthase-like protein OS=Homo sapiens OX=9606 GN=CBSL PE=1 SV=1 | 1.337 | 0.765235202 | 0.815 | 0.7 |
| P49902 | Cytosolic purine 5'-nucleotidase OS=Homo sapiens OX=9606 GN=NT5C2 PE=1 SV=1 | 0.808 | 0.750023665 | 0.852 | 0.7 |
| P18084 | Integrin beta-5 OS=Homo sapiens OX=9606 GN=ITGB5 PE=1 SV=1 | 1.146 | 0.899793601 | 0.955 | 0.8 |
| B8ZWD1 | Acyl-CoA-binding protein OS=Homo sapiens OX=9606 GN=DBI PE=1 SV=1 | 1.069 | 0.80706255 | 1.078 | 0.7 |
| Q02083 | N-acylethanolamine-hydrolyzing acid amidase OS=Homo sapiens OX=9606 GN=NAAA PE=1 SV=3 | 0.993 | 0.985330338 | 0.893 | 0.7 |
| A0A087X2B4 | Intraflagellar transport 20 homolog (Chlamydomonas), isoform CRA_b OS=Homo sapiens OX=9606 GN=IFT20 PE=1 SV=1 | 0.991 | 0.956978207 | 0.958 | 0.8 |
| P01023 | Alpha-2-macroglobulin OS=Homo sapiens OX=9606 GN=A2M PE=1 SV=3 | 1.459 | 0.902016292 | 0.958 | 0.[illegible] |
| A0A3B3ISV9 | ERI1 exoribonuclease 3 OS=Homo sapiens OX=9606 GN=ERI3 PE=1 SV=1 | 0.942 | 0.882454148 | 0.788 | 0.7 |
| E9PNF5 | Glutamine-dependent NAD(+) synthetase (Fragment) OS=Homo sapiens OX=9606 GN=NADSYN1 PE=1 SV=1 | 0.976 | 0.985324977 | 1.113 | 0.7 |
| Q13131 | 5'-AMP-activated protein kinase catalytic subunit alpha-1 OS=Homo sapiens OX=9606 GN=PRKAA1 PE=1 SV=4 | 1.022 | 0.987622091 | 0.806 | 0.7 |
| A0A1W2PNW1 | Signal transducer and activator of transcription (Fragment) OS=Homo sapiens OX=9606 GN=STAT6 PE=1 SV=2 | 1.038 | 0.750023665 | 1.06 | 0.7 |
| Q13438 | Protein OS-9 OS=Homo sapiens OX=9606 GN=OS9 PE=1 SV=1 | 1.007 | 0.968359197 | 0.976 | 0.7 |
| Q9P2W9 | Syntaxin-18 OS=Homo sapiens OX=9606 GN=STX18 PE=1 SV=1 | 0.985 | 0.906450037 | 0.927 | 0.7 |
| Q13496 | Myotubularin OS=Homo sapiens OX=9606 GN=MTM1 PE=1 SV=2 | 1.033 | 0.917008272 | 0.899 | 0.7 |
| Q9HAS0 | Protein Njmu-R1 OS=Homo sapiens OX=9606 GN=C17orf75 PE=1 SV=2 | | | | |
| A0A0C4DGQ8 | Dehydrogenase/reductase SDR family member 7B OS=Homo sapiens OX=9606 GN=DHRS7B PE=1 SV=1 | 0.911 | 0.840158726 | 0.81 | 0.7 |
| Q9Y3D9 | 28S ribosomal protein S23, mitochondrial OS=Homo sapiens OX=9606 GN=MRPS23 PE=1 SV=2 | 0.942 | 0.805488634 | 0.902 | 0.7 |
| Q9H8H0 | Nucleolar protein 11 OS=Homo sapiens OX=9606 GN=NOL11 PE=1 SV=1 | 0.994 | 0.896983349 | 0.942 | 0.7 |
| P55899 | IgG receptor FcRn large subunit p51 OS=Homo sapiens OX=9606 GN=FCGRT PE=1 SV=1 | 0.989 | 0.95071661 | 0.911 | 0.7 |
| A0A3B3ISV3 | Collagen alpha-1(IV) chain (Fragment) OS=Homo sapiens OX=9606 GN=COL4A1 PE=1 SV=1 | 0.917 | 0.839891753 | 0.927 | 0.7 |
| O75306 | NADH dehydrogenase [ubiquinone] iron-sulfur protein 2, mitochondrial OS=Homo sapiens OX=9606 GN=NDUFS2 PE=1 SV=2 | 1.096 | 0.904500837 | 0.692 | 0.7 |
| P82094 | TATA element modulatory factor OS=Homo sapiens OX=9606 GN=TMF1 PE=1 SV=2 | 0.853 | 0.765235202 | 0.869 | 0.7 |
| Q8TCJ2 | Dolichyl-diphosphooligosaccharide--protein glycosyltransferase subunit STT3B OS=Homo sapiens OX=9606 GN=STT3B PE=1 SV=1 | 0.965 | 0.906273459 | 1.003 | 0.8 |
| E7ESY4 | Metastasis-associated protein MTA1 OS=Homo sapiens OX=9606 GN=MTA1 PE=1 SV=1 | 1.095 | 0.765235202 | 0.876 | 0.7 |
| P24941 | Cyclin-dependent kinase 2 OS=Homo sapiens OX=9606 GN=CDK2 PE=1 SV=2 | | | | |
| F8VUB6 | Aladin OS=Homo sapiens OX=9606 GN=AAAS PE=1 SV=2 | 1.019 | 0.876510992 | 1.048 | 0.9 |
| Q92692 | Nectin-2 OS=Homo sapiens OX=9606 GN=NECTIN2 PE=1 SV=1 | 1.002 | 0.998388336 | 0.949 | 0.7 |
| Q9P2N2 | Rho GTPase-activating protein 28 OS=Homo sapiens OX=9606 GN=ARHGAP28 PE=1 SV=3 | 1.044 | 0.96647353 | 0.878 | 0.7 |
| Q6V0I7 | Protocadherin Fat 4 OS=Homo sapiens OX=9606 GN=FAT4 PE=1 SV=2 | 1.045 | 0.917008272 | 0.803 | 0.7 |
| Q13015 | Protein AF1q OS=Homo sapiens OX=9606 GN=MLLT11 PE=1 SV=1 | 1.062 | 0.934908644 | 0.998 | 0.8 |
| Q9HD26 | Golgi-associated PDZ and coiled-coil motif-containing protein OS=Homo sapiens OX=9606 GN=GOPC PE=1 SV=1 | 0.977 | 0.952913364 | 1.055 | 0.[illegible] |
| Q5TCZ7 | Copper homeostasis protein cutC homolog (Fragment) OS=Homo sapiens OX=9606 GN=CUTC PE=1 SV=1 | 0.964 | 0.766250742 | 0.867 | 0.7 |
| M0R2N5 | Very-long-chain enoyl-CoA reductase OS=Homo sapiens OX=9606 GN=TECR PE=1 SV=1 | 1.099 | 0.750023665 | 1 | 0.8 |
| O95071 | E3 ubiquitin-protein ligase UBR5 OS=Homo sapiens OX=9606 GN=UBR5 PE=1 SV=2 | | | | |
| Q8TAE8 | Growth arrest and DNA damage-inducible proteins-interacting protein 1 OS=Homo sapiens OX=9606 GN=GADD45GIP1 PE=1 SV=1 | 0.998 | 0.976601777 | 0.883 | 0.7 |
| P56381 | ATP synthase subunit epsilon, mitochondrial OS=Homo sapiens OX=9606 GN=ATP5F1E PE=1 SV=2 | 0.917 | 0.80706255 | 0.924 | 0.7 |
| Q9HBH5 | Retinol dehydrogenase 14 OS=Homo sapiens OX=9606 GN=RDH14 PE=1 SV=1 | 1.066 | 0.880371237 | 0.879 | 0.7 |
| Q96BY6 | Dedicator of cytokinesis protein 10 OS=Homo sapiens OX=9606 GN=DOCK10 PE=1 SV=3 | 1.06 | 0.80706255 | 0.93 | 0.7 |
| F8VRL4 | Diphosphoinositol polyphosphate phosphohydrolase 2 (Fragment) OS=Homo sapiens OX=9606 GN=NUDT4 PE=1 SV=1 | 1.003 | 0.956825234 | 1.07 | 0.8 |
| Q01433 | AMP deaminase 2 OS=Homo sapiens OX=9606 GN=AMPD2 PE=1 SV=2 | 1.034 | 0.904500837 | 1 | 0.9 |
| F5GZ28 | DNA ligase OS=Homo sapiens OX=9606 GN=LIG1 PE=1 SV=1 | | | | |
| A0A087WY58 | Serine/threonine-protein kinase Nek3 OS=Homo sapiens OX=9606 GN=NEK3 PE=1 SV=1 | 0.971 | 0.895022941 | 0.831 | 0.7 |
| Q32M88 | Protein-glucosylgalactosylhydroxylysine glucosidase OS=Homo sapiens OX=9606 GN=PGGHG PE=1 SV=2 | | | | |
| A0A024RCR6 | BCL2-associated athanogene 6 OS=Homo sapiens OX=9606 GN=BAG6 PE=1 SV=1 | 1.119 | 0.765235202 | 1.009 | 0.9 |
| Q6YHU6 | Thyroid adenoma-associated protein OS=Homo sapiens OX=9606 GN=THADA PE=1 SV=1 | 1.032 | 0.985324977 | 1.009 | 0.7 |
| Q14202 | Zinc finger MYM-type protein 3 OS=Homo sapiens OX=9606 GN=ZMYM3 PE=1 SV=2 | 1.162 | 0.805488634 | 0.886 | 0.7 |
| Q5SVL2 | Caspase-7 (Fragment) OS=Homo sapiens OX=9606 GN=CASP7 PE=1 SV=1 | 0.986 | 0.805488634 | 0.962 | 0.7 |
| Q13617 | Cullin-2 OS=Homo sapiens OX=9606 GN=CUL2 PE=1 SV=2 | 0.924 | 0.750023665 | 0.96 | 0.7 |

| | | | | | |
|---|---|---|---|---|---|
| K7EIJ4 | Ran-binding protein 3 (Fragment) OS=Homo sapiens OX=9606 GN=RANBP3 PE=1 SV=1 | 1.019 | 0.989102644 | 0.995 | 0.9 |
| Q13546 | Receptor-interacting serine/threonine-protein kinase 1 OS=Homo sapiens OX=9606 GN=RIPK1 PE=1 SV=3 | 0.983 | 0.966959217 | 0.78 | 0.7 |
| Q9NXD2 | Myotubularin-related protein 10 OS=Homo sapiens OX=9606 GN=MTMR10 PE=1 SV=3 | | | | |
| A0A5F9ZHU7 | DNA primase large subunit OS=Homo sapiens OX=9606 GN=PRIM2 PE=1 SV=1 | 0.778 | 0.827871327 | 0.815 | 0.7 |
| Q96I51 | RCC1-like G exchanging factor-like protein OS=Homo sapiens OX=9606 GN=RCC1L PE=1 SV=2 | 0.89 | 0.765937763 | 0.917 | 0.7 |
| B7ZAR9 | Pyruvate dehydrogenase phosphatase regulatory subunit, mitochondrial OS=Homo sapiens OX=9606 GN=PDPR PE=1 SV=1 | 1.062 | 0.986297807 | 0.89 | 0.7 |
| J3KP15 | Serine/arginine-rich splicing factor 2 (Fragment) OS=Homo sapiens OX=9606 GN=SRSF2 PE=1 SV=8 | 0.947 | 0.765235202 | 0.99 | 0.9 |
| K7ES11 | (E3-independent) E2 ubiquitin-conjugating enzyme (Fragment) OS=Homo sapiens OX=9606 GN=UBE2O PE=1 SV=1 | 1.081 | 0.765235202 | 1.067 | 0.7 |
| Q96MW1 | Coiled-coil domain-containing protein 43 OS=Homo sapiens OX=9606 GN=CCDC43 PE=1 SV=2 | 1.051 | 0.815069882 | 0.89 | 0.7 |
| I3L3I6 | NmrA-like family domain-containing protein 1 (Fragment) OS=Homo sapiens OX=9606 GN=NMRAL1 PE=1 SV=8 | 0.949 | 0.80706255 | 0.869 | 0.7 |
| A0A2R8Y658 | Ribonuclease (Fragment) OS=Homo sapiens OX=9606 GN=RNASEH2A PE=1 SV=1 | 0.674 | 0.750023665 | 0.834 | 0.7 |
| H0YDP9 | Receptor-type tyrosine-protein phosphatase kappa (Fragment) OS=Homo sapiens OX=9606 GN=PTPRK PE=1 SV=1 | 1.005 | 0.952913364 | 0.947 | 0.7 |
| A0A0A0MTR2 | Prolyl 3-hydroxylase OGFOD1 OS=Homo sapiens OX=9606 GN=OGFOD1 PE=1 SV=1 | 0.836 | 0.854902952 | 0.719 | 0.7 |
| C9JSR1 | Myotubularin-related protein 14 OS=Homo sapiens OX=9606 GN=MTMR14 PE=1 SV=1 | 0.955 | 0.790808367 | 0.941 | 0.7 |
| Q9H3H3 | UPF0696 protein C11orf68 OS=Homo sapiens OX=9606 GN=C11orf68 PE=1 SV=3 | 0.973 | 0.911709867 | 0.953 | 0.7 |
| Q8TDZ2 | [F-actin]-monooxygenase MICAL1 OS=Homo sapiens OX=9606 GN=MICAL1 PE=1 SV=2 | 1.118 | 0.827091161 | 1.046 | 0.9 |
| O14530 | Thioredoxin domain-containing protein 9 OS=Homo sapiens OX=9606 GN=TXNDC9 PE=1 SV=2 | 0.963 | 0.884192327 | 1.087 | 0.7 |
| O43181 | NADH dehydrogenase [ubiquinone] iron-sulfur protein 4, mitochondrial OS=Homo sapiens OX=9606 GN=NDUFS4 PE=1 SV=1 | 0.941 | 0.996578046 | 0.97 | 0.7 |
| Q5SZK5 | Wiskott-Aldrich syndrome protein family member 1 (Fragment) OS=Homo sapiens OX=9606 GN=WASF1 PE=1 SV=1 | 1.039 | 0.774484855 | 0.864 | 0.[illegible] |
| H0YE28 | Selenoprotein H (Fragment) OS=Homo sapiens OX=9606 GN=SELENOH PE=1 SV=1 | 1.106 | 0.839891753 | 0.889 | 0.7 |
| Q3KQU3 | MAP7 domain-containing protein 1 OS=Homo sapiens OX=9606 GN=MAP7D1 PE=1 SV=1 | 1.047 | 0.765235202 | 1 | 0.9 |
| P52701 | DNA mismatch repair protein Msh6 OS=Homo sapiens OX=9606 GN=MSH6 PE=1 SV=2 | 1.014 | 0.944284199 | 1.058 | 0.9 |
| Q8N357 | Solute carrier family 35 member F6 OS=Homo sapiens OX=9606 GN=SLC35F6 PE=1 SV=1 | 0.884 | 0.837569766 | 1.04 | 0.9 |
| O00217 | NADH dehydrogenase [ubiquinone] iron-sulfur protein 8, mitochondrial OS=Homo sapiens OX=9606 GN=NDUFS8 PE=1 SV=1 | 0.941 | 0.829989142 | 0.956 | 0.[illegible] |
| O75976 | Carboxypeptidase D OS=Homo sapiens OX=9606 GN=CPD PE=1 SV=2 | 1.086 | 0.974919191 | 0.969 | 0.7 |
| Q9NPJ6 | Mediator of RNA polymerase II transcription subunit 4 OS=Homo sapiens OX=9606 GN=MED4 PE=1 SV=1 | 0.989 | 0.955325044 | 0.823 | 0.7 |
| Q96S55 | ATPase WRNIP1 OS=Homo sapiens OX=9606 GN=WRNIP1 PE=1 SV=2 | 0.86 | 0.824784308 | 0.949 | 0.8 |
| H7C5M1 | RNA-binding protein 34 (Fragment) OS=Homo sapiens OX=9606 GN=RBM34 PE=1 SV=1 | 0.967 | 0.869564486 | 0.749 | 0.7 |
| Q7Z3C6 | Autophagy-related protein 9A OS=Homo sapiens OX=9606 GN=ATG9A PE=1 SV=3 | 0.958 | 0.864643132 | 0.799 | 0.8 |
| Q643R3 | Lysophospholipid acyltransferase LPCAT4 OS=Homo sapiens OX=9606 GN=LPCAT4 PE=1 SV=1 | 0.992 | 0.942843201 | 0.984 | 0.8 |
| P28676 | Grancalcin OS=Homo sapiens OX=9606 GN=GCA PE=1 SV=2 | 0.923 | 0.827451813 | 0.902 | 0.7 |
| Q9P2B2 | Prostaglandin F2 receptor negative regulator OS=Homo sapiens OX=9606 GN=PTGFRN PE=1 SV=2 | 0.981 | 0.854598401 | 0.951 | 0.7 |
| X6RK58 | Leucine-rich repeat-containing protein 20 (Fragment) OS=Homo sapiens OX=9606 GN=LRRC20 PE=1 SV=7 | 0.852 | 0.642276967 | 1.024 | 0.9 |
| Q96DV4 | 39S ribosomal protein L38, mitochondrial OS=Homo sapiens OX=9606 GN=MRPL38 PE=1 SV=2 | 1.04 | 0.994361756 | 0.888 | 0.7 |
| Q99747 | Gamma-soluble NSF attachment protein OS=Homo sapiens OX=9606 GN=NAPG PE=1 SV=1 | 0.943 | 0.797711953 | 1.01 | 0.7 |
| Q92783 | Signal transducing adapter molecule 1 OS=Homo sapiens OX=9606 GN=STAM PE=1 SV=3 | 1.041 | 0.80706255 | 0.933 | 0.7 |
| Q5JRA6 | Transport and Golgi organization protein 1 homolog OS=Homo sapiens OX=9606 GN=MIA3 PE=1 SV=1 | 1.014 | 0.942843201 | 0.935 | 0.7 |
| Q4G0N4 | NAD kinase 2, mitochondrial OS=Homo sapiens OX=9606 GN=NADK2 PE=1 SV=2 | 0.988 | 0.886561233 | 0.923 | 0.7 |
| Q9NZV5 | Selenoprotein N OS=Homo sapiens OX=9606 GN=SELENON PE=1 SV=5 | 1.48 | 0.801301898 | 0.636 | 0.7 |
| Q92609 | TBC1 domain family member 5 OS=Homo sapiens OX=9606 GN=TBC1D5 PE=1 SV=1 | 0.965 | 0.750023665 | 0.888 | 0.7 |
| Q9Y5X2 | Sorting nexin-8 OS=Homo sapiens OX=9606 GN=SNX8 PE=1 SV=1 | 0.996 | 0.905369216 | 0.814 | 0.7 |
| H7BZU5 | Neuroblastoma-amplified sequence (Fragment) OS=Homo sapiens OX=9606 GN=NBAS PE=1 SV=1 | | | | |
| P07305 | Histone H1.0 OS=Homo sapiens OX=9606 GN=H1-0 PE=1 SV=3 | 0.946 | 0.985324977 | 0.99 | 0.7 |
| H3BP35 | Diphosphomevalonate decarboxylase (Fragment) OS=Homo sapiens OX=9606 GN=MVD PE=1 SV=1 | 0.899 | 0.750023665 | 0.827 | 0.7 |
| Q2T9J0 | Peroxisomal leader peptide-processing protease OS=Homo sapiens OX=9606 GN=TYSND1 PE=1 SV=3 | 12.791 | 0.9962822 | 0.379 | |
| P46100 | Transcriptional regulator ATRX OS=Homo sapiens OX=9606 GN=ATRX PE=1 SV=5 | 1.007 | 0.876510992 | 0.852 | 0.7 |
| S4R3T2 | E3 ubiquitin-protein ligase listerin (Fragment) OS=Homo sapiens OX=9606 GN=LTN1 PE=1 SV=1 | | | | |
| E7EQ45 | AP-5 complex subunit mu-1 OS=Homo sapiens OX=9606 GN=AP5M1 PE=1 SV=1 | 1.094 | 0.996578046 | 0.831 | 0.7 |
| A0A3B3ISI3 | Phosphatidylinositide phosphatase SAC2 OS=Homo sapiens OX=9606 GN=INPP5F PE=1 SV=1 | 0.898 | 0.865970839 | 0.709 | 0.7 |
| A0A3B3ITR6 | Phenylalanyl-tRNA synthetase OS=Homo sapiens OX=9606 GN=FARS2 PE=1 SV=1 | 1.13 | 0.750023665 | 0.865 | 0.7 |
| Q7Z422 | SUZ domain-containing protein 1 OS=Homo sapiens OX=9606 GN=SZRD1 PE=1 SV=1 | 1.041 | 0.990981579 | 0.934 | 0.7 |
| Q9Y657 | Spindlin-1 OS=Homo sapiens OX=9606 GN=SPIN1 PE=1 SV=3 | 0.938 | 0.809976184 | 1.001 | 0.9 |
| Q5TBU2 | Adipogenesis regulatory factor (Fragment) OS=Homo sapiens OX=9606 GN=ADIRF PE=1 SV=1 | 1.03 | 0.806303314 | 1.028 | 0.8 |
| P15374 | Ubiquitin carboxyl-terminal hydrolase isozyme L3 OS=Homo sapiens OX=9606 GN=UCHL3 PE=1 SV=1 | 0.995 | 0.997738183 | 1.107 | 0.9 |
| P26006 | Integrin alpha-3 OS=Homo sapiens OX=9606 GN=ITGA3 PE=1 SV=5 | 1.027 | 0.827871327 | 0.848 | 0.7 |
| O14519 | Cyclin-dependent kinase 2-associated protein 1 OS=Homo sapiens OX=9606 GN=CDK2AP1 PE=1 SV=1 | 0.973 | 0.869305857 | 0.942 | 0.7 |
| A0A590UJW0 | DNA mismatch repair protein Msh3 OS=Homo sapiens OX=9606 GN=MSH3 PE=1 SV=1 | 0.985 | 0.984200071 | 0.785 | 0.7 |
| Q8NI22 | Multiple coagulation factor deficiency protein 2 OS=Homo sapiens OX=9606 GN=MCFD2 PE=1 SV=1 | 0.899 | 0.811788015 | 0.904 | 0.7 |
| Q96A35 | 39S ribosomal protein L24, mitochondrial OS=Homo sapiens OX=9606 GN=MRPL24 PE=1 SV=1 | 1.304 | 0.765235202 | 0.954 | 0.8 |
| Q96BP3 | Peptidylprolyl isomerase domain and WD repeat-containing protein 1 OS=Homo sapiens OX=9606 GN=PPWD1 PE=1 SV=1 | 0.964 | 0.929783322 | 0.817 | 0.7 |
| O94804 | Serine/threonine-protein kinase 10 OS=Homo sapiens OX=9606 GN=STK10 PE=1 SV=1 | | | | |
| O14656 | Torsin-1A OS=Homo sapiens OX=9606 GN=TOR1A PE=1 SV=1 | 1.15 | 0.703015003 | 0.975 | 0.8 |
| H7C096 | BCL2/adenovirus E1B 19 kDa protein-interacting protein 2 (Fragment) OS=Homo sapiens OX=9606 GN=BNIP2 PE=1 SV=1 | 0.881 | 0.985324977 | 0.776 | 0.7 |
| H7C5L6 | SRSF protein kinase 2 (Fragment) OS=Homo sapiens OX=9606 GN=SRPK2 PE=1 SV=1 | 1.023 | 0.829390484 | 0.975 | 0.8 |
| Q6ZT12 | E3 ubiquitin-protein ligase UBR3 OS=Homo sapiens OX=9606 GN=UBR3 PE=2 SV=2 | 0.901 | 0.790136102 | 1.03 | 0.9 |
| A0A6I8PIV6 | Adhesion G protein-coupled receptor L2 OS=Homo sapiens OX=9606 GN=ADGRL2 PE=4 SV=1 | 1.085 | 0.983953845 | 0.897 | 0.7 |
| Q96RT7 | Gamma-tubulin complex component 6 OS=Homo sapiens OX=9606 GN=TUBGCP6 PE=1 SV=3 | 1.673 | 0.750023665 | 1.02 | 0.7 |
| Q5T073 | Lysosomal acid lipase/cholesteryl ester hydrolase (Fragment) OS=Homo sapiens OX=9606 GN=LIPA PE=1 SV=1 | 0.929 | 0.809976184 | 0.925 | 0.7 |
| Q8IZQ1 | WD repeat and FYVE domain-containing protein 3 OS=Homo sapiens OX=9606 GN=WDFY3 PE=1 SV=2 | 0.987 | 0.917150381 | 0.902 | 0.7 |
| G3V158 | 2-deoxy-D-ribose 5-phosphate aldolase OS=Homo sapiens OX=9606 GN=DERA PE=1 SV=1 | 1.001 | 0.95980123 | 0.932 | 0.9 |
| Q96KP1 | Exocyst complex component 2 OS=Homo sapiens OX=9606 GN=EXOC2 PE=1 SV=1 | 0.944 | 0.866300162 | 1.005 | 0.9 |
| Q9UHY8 | Fasciculation and elongation protein zeta-2 OS=Homo sapiens OX=9606 GN=FEZ2 PE=1 SV=2 | 1.191 | 0.750023665 | 0.97 | 0.8 |
| C9JEL3 | Eukaryotic translation initiation factor 4E type 2 (Fragment) OS=Homo sapiens OX=9606 GN=EIF4E2 PE=1 SV=1 | 0.939 | 0.87038391 | 0.947 | 0.8 |
| P52735 | Guanine nucleotide exchange factor VAV2 OS=Homo sapiens OX=9606 GN=VAV2 PE=1 SV=2 | 1.049 | 0.829390484 | 0.782 | 0.7 |
| Q969U7 | Proteasome assembly chaperone 2 OS=Homo sapiens OX=9606 GN=PSMG2 PE=1 SV=1 | 1.085 | 0.776314646 | 1.116 | 0.7 |
| A0A0A0MTH9 | TATA-binding protein-associated factor 172 OS=Homo sapiens OX=9606 GN=BTAF1 PE=1 SV=1 | 0.703 | 0.765235202 | 8.581 | 0.7 |
| Q8WW12 | PEST proteolytic signal-containing nuclear protein OS=Homo sapiens OX=9606 GN=PCNP PE=1 SV=2 | 1.053 | 0.917008272 | 1.008 | 0.9 |
| Q96K76 | Ubiquitin carboxyl-terminal hydrolase 47 OS=Homo sapiens OX=9606 GN=USP47 PE=1 SV=3 | 0.907 | 0.827871327 | 0.989 | 0.[illegible] |
| O60504 | Vinexin OS=Homo sapiens OX=9606 GN=SORBS3 PE=1 SV=2 | 0.996 | 0.987622091 | 0.943 | 0.7 |
| Q15291 | Retinoblastoma-binding protein 5 OS=Homo sapiens OX=9606 GN=RBBP5 PE=1 SV=2 | 0.952 | 0.944284199 | 0.95 | 0.7 |
| Q92805 | Golgin subfamily A member 1 OS=Homo sapiens OX=9606 GN=GOLGA1 PE=1 SV=3 | 1.215 | 0.765235202 | 0.904 | 0.7 |
| A0A1W2PNR9 | Immediate early response 3-interacting protein 1 OS=Homo sapiens OX=9606 GN=IER3IP1 PE=1 SV=1 | 1.077 | 0.763877559 | 0.872 | 0.7 |
| Q8N3R9 | MAGUK p55 subfamily member 5 OS=Homo sapiens OX=9606 GN=MPP5 PE=1 SV=3 | 1.059 | 0.862833222 | 0.872 | 0.7 |
| A0A5F9ZHG6 | Ataxin-2 (Fragment) OS=Homo sapiens OX=9606 GN=ATXN2 PE=1 SV=1 | 1.034 | 0.77773062 | 0.87 | 0.7 |
| Q5T160 | Probable arginine--tRNA ligase, mitochondrial OS=Homo sapiens OX=9606 GN=RARS2 PE=1 SV=1 | 0.895 | 0.867483128 | 0.949 | 0.7 |
| O15037 | Protein KHNYN OS=Homo sapiens OX=9606 GN=KHNYN PE=1 SV=3 | 0.662 | 0.765235202 | 0.924 | 0.9 |
| Q9NWW6 | Nicotinamide riboside kinase 1 OS=Homo sapiens OX=9606 GN=NMRK1 PE=1 SV=1 | 1.205 | 0.903190191 | 0.955 | 0.7 |
| E5RG91 | Vacuolar protein sorting-associated protein 37A OS=Homo sapiens OX=9606 GN=VPS37A PE=1 SV=1 | 1.041 | 0.95980123 | 1.085 | 0.9 |
| Q9Y673 | Dolichyl-phosphate beta-glucosyltransferase OS=Homo sapiens OX=9606 GN=ALG5 PE=1 SV=1 | 1.066 | 0.858815413 | 0.904 | 0.8 |
| P27658 | Collagen alpha-1(VIII) chain OS=Homo sapiens OX=9606 GN=COL8A1 PE=1 SV=2 | 1.044 | 0.983953845 | 0.794 | 0.7 |
| Q99933 | BAG family molecular chaperone regulator 1 OS=Homo sapiens OX=9606 GN=BAG1 PE=1 SV=4 | 0.821 | 0.756287988 | 0.938 | 0.8 |
| Q9H553 | Alpha-1,3/1,6-mannosyltransferase ALG2 OS=Homo sapiens OX=9606 GN=ALG2 PE=1 SV=1 | 0.948 | 0.95980123 | 0.981 | 0.8 |
| O76071 | Probable cytosolic iron-sulfur protein assembly protein CIAO1 OS=Homo sapiens OX=9606 GN=CIAO1 PE=1 SV=1 | 0.974 | 0.754725464 | 0.922 | 0.7 |
| A0A2R8YE10 | Vacuolar protein sorting-associated protein 45 OS=Homo sapiens OX=9606 GN=VPS45 PE=1 SV=1 | 1.007 | 0.889450577 | 1.077 | 0.9 |
| Q9GZN8 | UPF0687 protein C20orf27 OS=Homo sapiens OX=9606 GN=C20orf27 PE=1 SV=3 | | | | |
| X6RKY7 | Cell division cycle protein 123 homolog (Fragment) OS=Homo sapiens OX=9606 GN=CDC123 PE=1 SV=1 | 0.937 | 0.763877559 | 0.999 | 0.[illegible] |

| | | | | | |
|---|---|---|---|---|---|
| Q8TED0 | U3 small nucleolar RNA-associated protein 15 homolog OS=Homo sapiens OX=9606 GN=UTP15 PE=1 SV=3 | 1.107 | 0.765235202 | 0.981 | 0.9 |
| A0A3B3ISP1 | Ankyrin repeat and SAM domain-containing protein 1A OS=Homo sapiens OX=9606 GN=ANKS1A PE=1 SV=1 | | | | |
| Q9NW13 | RNA-binding protein 28 OS=Homo sapiens OX=9606 GN=RBM28 PE=1 SV=3 | 1.032 | 0.984200071 | 1.03 | 0.8 |
| Q8IVF2 | Protein AHNAK2 OS=Homo sapiens OX=9606 GN=AHNAK2 PE=1 SV=2 | 1.131 | 0.780884711 | 1.024 | 0.8 |
| B4DJL6 | Integrator complex subunit 14 OS=Homo sapiens OX=9606 GN=INTS14 PE=1 SV=1 | 1.058 | 0.987438982 | 0.938 | 0.7 |
| K7ES46 | DNA excision repair protein ERCC-1 (Fragment) OS=Homo sapiens OX=9606 GN=ERCC1 PE=1 SV=1 | 0.986 | 0.904500837 | 0.921 | 0.9 |
| H7C0V0 | m-AAA protease-interacting protein 1, mitochondrial (Fragment) OS=Homo sapiens OX=9606 GN=MAIP1 PE=1 SV=1 | 0.871 | 0.780884711 | 0.971 | 0.7 |
| E9PRJ8 | Tetraspanin (Fragment) OS=Homo sapiens OX=9606 GN=CD81 PE=1 SV=1 | 1 | 0.908555656 | 0.876 | 0.7 |
| U3KQN5 | Ribosome production factor 2 homolog OS=Homo sapiens OX=9606 GN=RPF2 PE=1 SV=1 | 1.049 | 0.892311206 | 0.966 | 0.9 |
| Q9Y4C8 | Probable RNA-binding protein 19 OS=Homo sapiens OX=9606 GN=RBM19 PE=1 SV=3 | 0.99 | 0.963059765 | 1.013 | 0.[illegible] |
| Q6ZR19 | Receptor-type tyrosine-protein phosphatase beta (Fragment) OS=Homo sapiens OX=9606 GN=PTPRB PE=1 SV=1 | 0.996 | 0.913279108 | 1.048 | 0.9 |
| I3L3A0 | Cytosolic Fe-S cluster assembly factor NUBP1 (Fragment) OS=Homo sapiens OX=9606 GN=NUBP1 PE=1 SV=1 | 0.928 | 0.680740328 | 0.956 | 0.7 |
| P52298 | Nuclear cap-binding protein subunit 2 OS=Homo sapiens OX=9606 GN=NCBP2 PE=1 SV=1 | 0.967 | 0.827451813 | 0.838 | 0.7 |
| Q02763 | Angiopoietin-1 receptor OS=Homo sapiens OX=9606 GN=TEK PE=1 SV=2 | | | | |
| Q9H4X1 | Regulator of cell cycle RGCC OS=Homo sapiens OX=9606 GN=RGCC PE=1 SV=1 | 0.867 | 0.756287988 | 0.768 | 0.7 |
| A0A087X0S5 | Collagen alpha-1(VI) chain OS=Homo sapiens OX=9606 GN=COL6A1 PE=1 SV=1 | | | | |
| A0A6I8PIU5 | Heat shock 70 kDa protein 12A OS=Homo sapiens OX=9606 GN=HSPA12A PE=4 SV=1 | 1.028 | 0.965945681 | 0.838 | 0.7 |
| F8W9X7 | Coiled-coil domain-containing protein 93 OS=Homo sapiens OX=9606 GN=CCDC93 PE=1 SV=1 | 1.138 | 0.902016292 | 0.886 | 0.9 |
| Q9C0H6 | Kelch-like protein 4 OS=Homo sapiens OX=9606 GN=KLHL4 PE=1 SV=2 | 1.062 | 0.774484855 | 0.934 | 0.[illegible] |
| Q9H9Y6 | DNA-directed RNA polymerase I subunit RPA2 OS=Homo sapiens OX=9606 GN=POLR1B PE=1 SV=2 | 1.063 | 0.972073425 | 1.124 | 0.8 |
| Q05086 | Ubiquitin-protein ligase E3A OS=Homo sapiens OX=9606 GN=UBE3A PE=1 SV=4 | 1.1 | 0.837941467 | 0.949 | 0.7 |
| Q08378 | Golgin subfamily A member 3 OS=Homo sapiens OX=9606 GN=GOLGA3 PE=1 SV=2 | 0.963 | 0.902016292 | 0.756 | 0.7 |
| Q92503 | SEC14-like protein 1 OS=Homo sapiens OX=9606 GN=SEC14L1 PE=1 SV=2 | 1.293 | 0.750023665 | 1.329 | 0.9 |
| Q5QPM7 | Proteasome inhibitor PI31 subunit OS=Homo sapiens OX=9606 GN=PSMF1 PE=1 SV=2 | 0.889 | 0.765235202 | 1.055 | 0.9 |
| A6NEP9 | Tetraspanin (Fragment) OS=Homo sapiens OX=9606 GN=TSPAN14 PE=1 SV=2 | 0.942 | 0.765235202 | 0.979 | 0.7 |
| Q9NR19 | Acetyl-coenzyme A synthetase, cytoplasmic OS=Homo sapiens OX=9606 GN=ACSS2 PE=1 SV=1 | | | | |
| P19387 | DNA-directed RNA polymerase II subunit RPB3 OS=Homo sapiens OX=9606 GN=POLR2C PE=1 SV=2 | 0.992 | 0.913279108 | 0.946 | 0.9 |
| A0A5F9ZHM8 | Argininosuccinate lyase (Fragment) OS=Homo sapiens OX=9606 GN=ASL PE=1 SV=1 | 0.965 | 0.903226585 | 0.913 | 0.7 |
| Q5QP56 | Apoptosis regulator Bcl-X (Fragment) OS=Homo sapiens OX=9606 GN=BCL2L1 PE=1 SV=1 | 0.971 | 0.869995963 | 0.88 | 0.7 |
| Q9H173 | Nucleotide exchange factor SIL1 OS=Homo sapiens OX=9606 GN=SIL1 PE=1 SV=1 | 0.814 | 0.703015003 | 0.913 | 0.9 |
| P61009 | Signal peptidase complex subunit 3 OS=Homo sapiens OX=9606 GN=SPCS3 PE=1 SV=1 | 0.953 | 0.765235202 | 0.934 | 0.7 |
| O75818 | Ribonuclease P protein subunit p40 OS=Homo sapiens OX=9606 GN=RPP40 PE=1 SV=3 | 1.002 | 0.920144076 | 0.879 | 0.7 |
| Q86V48 | Leucine zipper protein 1 OS=Homo sapiens OX=9606 GN=LUZP1 PE=1 SV=2 | 0.987 | 0.944284199 | 0.937 | 0.7 |
| Q9H269 | Vacuolar protein sorting-associated protein 16 homolog OS=Homo sapiens OX=9606 GN=VPS16 PE=1 SV=2 | 1.099 | 0.765235202 | 0.985 | 0.[illegible] |
| E7EPT4 | NADH dehydrogenase [ubiquinone] flavoprotein 2, mitochondrial OS=Homo sapiens OX=9606 GN=NDUFV2 PE=1 SV=1 | 0.985 | 0.902016292 | 1.001 | 0.7 |
| Q9H3Z4 | DnaJ homolog subfamily C member 5 OS=Homo sapiens OX=9606 GN=DNAJC5 PE=1 SV=1 | 0.75 | 0.750023665 | 0.913 | 0.7 |
| H0YCV9 | CD44 antigen (Fragment) OS=Homo sapiens OX=9606 GN=CD44 PE=1 SV=2 | 0.946 | 0.750023665 | 1.008 | 0.7 |
| Q9P0S9 | Transmembrane protein 14C OS=Homo sapiens OX=9606 GN=TMEM14C PE=1 SV=1 | 0.983 | 0.953792095 | 0.673 | 0.7 |
| Q8N5N7 | 39S ribosomal protein L50, mitochondrial OS=Homo sapiens OX=9606 GN=MRPL50 PE=1 SV=2 | 1.051 | 0.892901616 | 0.831 | 0.7 |
| F5H4F1 | Cell cycle checkpoint control protein RAD9A (Fragment) OS=Homo sapiens OX=9606 GN=RAD9A PE=1 SV=1 | 0.941 | 0.750023665 | 0.864 | 0.7 |
| P60468 | Protein transport protein Sec61 subunit beta OS=Homo sapiens OX=9606 GN=SEC61B PE=1 SV=2 | 1.065 | 0.902016292 | 1.037 | 0.9 |
| K7EPR7 | Ubiquitin-conjugating enzyme E2 G1 (Fragment) OS=Homo sapiens OX=9606 GN=UBE2G1 PE=1 SV=1 | 0.996 | 0.956825234 | 0.839 | 0.7 |
| G3V325 | ATP5MF-PTCD1 readthrough OS=Homo sapiens OX=9606 GN=ATP5MF-PTCD1 PE=4 SV=1 | 0.969 | 0.905369216 | 0.866 | 0.7 |
| C9JJV6 | Myeloid-associated differentiation marker (Fragment) OS=Homo sapiens OX=9606 GN=MYADM PE=1 SV=8 | 1.008 | 0.868906519 | 0.957 | 0.7 |
| G3V1E1 | Sphingomyelin phosphodiesterase OS=Homo sapiens OX=9606 GN=SMPD1 PE=1 SV=1 | | | | |
| Q9H2J4 | Phosducin-like protein 3 OS=Homo sapiens OX=9606 GN=PDCL3 PE=1 SV=1 | 1.188 | 0.765235202 | 0.926 | 0.7 |
| Q9NU22 | Midasin OS=Homo sapiens OX=9606 GN=MDN1 PE=1 SV=2 | 0.983 | 0.900915273 | 0.963 | 0.[illegible] |
| P82673 | 28S ribosomal protein S35, mitochondrial OS=Homo sapiens OX=9606 GN=MRPS35 PE=1 SV=1 | 0.963 | 0.825684047 | 0.939 | 0.8 |
| U3KQB0 | DNA-directed RNA polymerase II subunit F OS=Homo sapiens OX=9606 GN=POLR2F PE=1 SV=1 | 1.001 | 0.978693882 | 0.899 | 0.7 |
| Q9NZB2 | Constitutive coactivator of PPAR-gamma-like protein 1 OS=Homo sapiens OX=9606 GN=FAM120A PE=1 SV=2 | 1.045 | 0.939024443 | 0.984 | 0.8 |
| H3BRS3 | Telomere length regulation protein TEL2 homolog OS=Homo sapiens OX=9606 GN=TELO2 PE=1 SV=1 | 0.726 | 0.750023665 | 1.072 | 0.7 |
| O75886 | Signal transducing adapter molecule 2 OS=Homo sapiens OX=9606 GN=STAM2 PE=1 SV=1 | 0.984 | 0.951324063 | 0.955 | 0.7 |
| V9GYH6 | Polyamine-modulated factor 1 (Fragment) OS=Homo sapiens OX=9606 GN=PMF1 PE=4 SV=1 | 1.043 | 0.996637279 | 0.924 | 0.7 |
| H0YJH7 | Lamina-associated polypeptide 2, isoforms beta/gamma (Fragment) OS=Homo sapiens OX=9606 GN=TMPO PE=1 SV=1 | 1.081 | 0.766667967 | 0.846 | 0.7 |
| G3XAJ6 | Raft-linking protein, isoform CRA_c OS=Homo sapiens OX=9606 GN=RFTN1 PE=1 SV=1 | 0.918 | 0.876510992 | 0.895 | 0.8 |
| Q8IUR0 | Trafficking protein particle complex subunit 5 OS=Homo sapiens OX=9606 GN=TRAPPC5 PE=1 SV=1 | 1.08 | 0.750023665 | 1.012 | 0.9 |
| Q96GQ7 | Probable ATP-dependent RNA helicase DDX27 OS=Homo sapiens OX=9606 GN=DDX27 PE=1 SV=2 | 1.069 | 0.750023665 | 1.075 | 0.7 |
| P56182 | Ribosomal RNA processing protein 1 homolog A OS=Homo sapiens OX=9606 GN=RRP1 PE=1 SV=1 | 1.022 | 0.994361756 | 0.931 | 0.7 |
| Q96MG7 | Non-structural maintenance of chromosomes element 3 homolog OS=Homo sapiens OX=9606 GN=NSMCE3 PE=1 SV=1 | 1.049 | 0.829390484 | 0.932 | 0.7 |
| A0A499FJY3 | RNA-binding protein 15 OS=Homo sapiens OX=9606 GN=RBM15 PE=1 SV=1 | 1.023 | 0.939024443 | 0.748 | 0.7 |
| H0Y9G5 | Wolframin OS=Homo sapiens OX=9606 GN=WFS1 PE=1 SV=2 | 1.296 | 0.765235202 | 0.8 | 0.7 |
| P32929 | Cystathionine gamma-lyase OS=Homo sapiens OX=9606 GN=CTH PE=1 SV=3 | 0.932 | 0.750023665 | 1.006 | 0.8 |
| F8WE91 | DNA (cytosine-5-)-methyltransferase OS=Homo sapiens OX=9606 GN=DNMT3A PE=1 SV=1 | 0.959 | 0.968693062 | 0.948 | 0.7 |
| Q08722 | Leukocyte surface antigen CD47 OS=Homo sapiens OX=9606 GN=CD47 PE=1 SV=1 | 0.947 | 0.790136102 | 0.973 | 0.7 |
| O00411 | DNA-directed RNA polymerase, mitochondrial OS=Homo sapiens OX=9606 GN=POLRMT PE=1 SV=2 | 1.078 | 0.857755811 | 1.009 | 0.9 |
| Q96N11 | Uncharacterized protein C7orf26 OS=Homo sapiens OX=9606 GN=C7orf26 PE=2 SV=1 | 1.059 | 0.849403601 | 0.91 | 0.7 |
| E9PG02 | Inactive C-alpha-formylglycine-generating enzyme 2 OS=Homo sapiens OX=9606 GN=SUMF2 PE=1 SV=1 | 0.924 | 0.809976184 | 0.929 | 0.7 |
| J3KQ41 | COP9 signalosome complex subunit 7b OS=Homo sapiens OX=9606 GN=COPS7B PE=1 SV=1 | 0.895 | 0.792509319 | 1.006 | 0.8 |
| B7Z5P7 | Leupaxin OS=Homo sapiens OX=9606 GN=LPXN PE=1 SV=1 | 1.154 | 0.762199181 | 0.885 | 0.7 |
| V9GZ56 | U6 snRNA-associated Sm-like protein LSm4 (Fragment) OS=Homo sapiens OX=9606 GN=LSM4 PE=1 SV=1 | 0.989 | 0.99634713 | 0.965 | 0.8 |
| F2Z2T0 | Rab-like protein 6 OS=Homo sapiens OX=9606 GN=RABL6 PE=1 SV=1 | 1.011 | 0.985324977 | 1.071 | 0.9 |
| Q6P158 | Putative ATP-dependent RNA helicase DHX57 OS=Homo sapiens OX=9606 GN=DHX57 PE=1 SV=2 | | | | |
| P46199 | Translation initiation factor IF-2, mitochondrial OS=Homo sapiens OX=9606 GN=MTIF2 PE=1 SV=2 | 0.968 | 0.827871327 | 0.922 | 0.7 |
| P00846 | ATP synthase subunit a OS=Homo sapiens OX=9606 GN=MT-ATP6 PE=1 SV=1 | 0.924 | 0.889450577 | 0.873 | 0.7 |
| Q9NP77 | RNA polymerase II subunit A C-terminal domain phosphatase SSU72 OS=Homo sapiens OX=9606 GN=SSU72 PE=1 SV=1 | 1.151 | 0.750023665 | 0.897 | 0.7 |
| Q14457 | Beclin-1 OS=Homo sapiens OX=9606 GN=BECN1 PE=1 SV=2 | 0.985 | 0.925929947 | 0.857 | 0.7 |
| Q9NX46 | ADP-ribose glycohydrolase ARH3 OS=Homo sapiens OX=9606 GN=ADPRS PE=1 SV=1 | 0.984 | 0.985324977 | 1.008 | 0.8 |
| Q96D46 | 60S ribosomal export protein NMD3 OS=Homo sapiens OX=9606 GN=NMD3 PE=1 SV=1 | 0.889 | 0.765235202 | 0.946 | 0.7 |
| Q8IYI6 | Exocyst complex component 8 OS=Homo sapiens OX=9606 GN=EXOC8 PE=1 SV=2 | 0.951 | 0.97688166 | 0.881 | 0.7 |
| Q96C01 | Protein FAM136A OS=Homo sapiens OX=9606 GN=FAM136A PE=1 SV=1 | 1.039 | 0.753211297 | 0.998 | 0.7 |
| Q96EY5 | Multivesicular body subunit 12A OS=Homo sapiens OX=9606 GN=MVB12A PE=1 SV=1 | 0.985 | 0.978562349 | 0.936 | 0.7 |
| O95218 | Zinc finger Ran-binding domain-containing protein 2 OS=Homo sapiens OX=9606 GN=ZRANB2 PE=1 SV=2 | 1.06 | 0.761475165 | 0.937 | 0.7 |
| A0A0D9SFH9 | Battenin OS=Homo sapiens OX=9606 GN=CLN3 PE=1 SV=1 | 1.154 | 0.968322408 | 0.922 | 0.8 |
| Q9NZV1 | Cysteine-rich motor neuron 1 protein OS=Homo sapiens OX=9606 GN=CRIM1 PE=1 SV=1 | 0.958 | 0.902016292 | 0.976 | 0.7 |
| A0A1B0GW95 | Protein IWS1 homolog (Fragment) OS=Homo sapiens OX=9606 GN=IWS1 PE=1 SV=1 | 1.137 | 0.785314375 | 1.016 | 0.9 |
| P82933 | 28S ribosomal protein S9, mitochondrial OS=Homo sapiens OX=9606 GN=MRPS9 PE=1 SV=2 | 1.034 | 0.920074608 | 0.882 | 0.7 |
| A0A087X2D5 | 39S ribosomal protein L45, mitochondrial OS=Homo sapiens OX=9606 GN=MRPL45 PE=1 SV=1 | 0.941 | 0.866054955 | 0.871 | 0.7 |
| Q8IVB5 | LIX1-like protein OS=Homo sapiens OX=9606 GN=LIX1L PE=2 SV=1 | 0.946 | 0.983953845 | 1.025 | 0.9 |
| F6VUX8 | Centromere protein J OS=Homo sapiens OX=9606 GN=CENPJ PE=1 SV=1 | 0.993 | 0.901934028 | 1.056 | 0.7 |
| A0A494C1M4 | 10-formyltetrahydrofolate dehydrogenase OS=Homo sapiens OX=9606 GN=ALDH1L2 PE=1 SV=1 | 0.815 | 0.750023665 | 0.813 | 0.[illegible] |
| Q8WUA4 | General transcription factor 3C polypeptide 2 OS=Homo sapiens OX=9606 GN=GTF3C2 PE=1 SV=2 | 1.012 | 0.902016292 | 0.873 | 0.[illegible] |
| Q14677 | Clathrin interactor 1 OS=Homo sapiens OX=9606 GN=CLINT1 PE=1 SV=1 | 1.039 | 0.80706255 | 1.025 | 0.8 |
| B1AKC9 | Receptor protein-tyrosine kinase OS=Homo sapiens OX=9606 GN=EPHB2 PE=1 SV=1 | 0.994 | 0.910192239 | 0.853 | 0.8 |
| Q92733 | Proline-rich protein PRCC OS=Homo sapiens OX=9606 GN=PRCC PE=1 SV=1 | 0.952 | 0.954224541 | 1.007 | 0.9 |
| O15294 | UDP-N-acetylglucosamine--peptide N-acetylglucosaminyltransferase 110 kDa subunit OS=Homo sapiens OX=9606 GN=OGT PE=1 [illegible] | 0.931 | 0.889482529 | 1 | 0.8 |
| Q6ZR64 | HBV PreS1-transactivated protein 1 OS=Homo sapiens OX=9606 GN=MXRA7 PE=1 SV=1 | 1.013 | 0.98478736 | 0.964 | 0.9 |

| | | | | | |
|---|---|---|---|---|---|
| P15291 | Beta-1,4-galactosyltransferase 1 OS=Homo sapiens OX=9606 GN=B4GALT1 PE=1 SV=5 | 1.045 | 0.907031173 | 0.898 | 0.7 |
| Q9UBV2 | Protein sel-1 homolog 1 OS=Homo sapiens OX=9606 GN=SEL1L PE=1 SV=3 | 0.735 | 0.912604818 | 0.768 | 0.7 |
| A0A0G2JQ76 | Granule-associated Rac and RHOG effector protein 1 (Fragment) OS=Homo sapiens OX=9606 GN=GARRE1 PE=1 SV=1 | 0.922 | 0.801799196 | 1.015 | 0.9 |
| Q13206 | Probable ATP-dependent RNA helicase DDX10 OS=Homo sapiens OX=9606 GN=DDX10 PE=1 SV=2 | 1.015 | 0.929783322 | 0.903 | 0.[illegible] |
| Q96HY7 | Probable 2-oxoglutarate dehydrogenase E1 component DHKTD1, mitochondrial OS=Homo sapiens OX=9606 GN=DHTKD1 PE=1 SV | 0.59 | 0.750023665 | 0.745 | 0.7 |
| P36269 | Glutathione hydrolase 5 proenzyme OS=Homo sapiens OX=9606 GN=GGT5 PE=1 SV=2 | 0.915 | 0.901934028 | 1.086 | 0.[illegible] |
| O75940 | Survival of motor neuron-related-splicing factor 30 OS=Homo sapiens OX=9606 GN=SMNDC1 PE=1 SV=1 | 0.981 | 0.985330338 | 0.941 | 0.7 |
| H7C009 | cGMP-dependent 3',5'-cyclic phosphodiesterase (Fragment) OS=Homo sapiens OX=9606 GN=PDE2A PE=1 SV=1 | | | | |
| C9JIK8 | Cysteine protease (Fragment) OS=Homo sapiens OX=9606 GN=ATG4B PE=1 SV=1 | 1.069 | 0.876510992 | 0.995 | 0.9 |
| Q9H5N1 | Rab GTPase-binding effector protein 2 OS=Homo sapiens OX=9606 GN=RABEP2 PE=1 SV=2 | 1.014 | 0.815069882 | 1.006 | 0.9 |
| M0QY97 | Zinc finger CCCH domain-containing protein 4 (Fragment) OS=Homo sapiens OX=9606 GN=ZC3H4 PE=1 SV=1 | 1.005 | 0.951324063 | 0.892 | 0.7 |
| Q9Y3R5 | Protein dopey-2 OS=Homo sapiens OX=9606 GN=DOP1B PE=1 SV=5 | 0.835 | 0.765235202 | 0.77 | 0.7 |
| P52435 | DNA-directed RNA polymerase II subunit RPB11-a OS=Homo sapiens OX=9606 GN=POLR2J PE=1 SV=1 | 0.856 | 0.917008272 | 0.945 | 0.7 |
| P82921 | 28S ribosomal protein S21, mitochondrial OS=Homo sapiens OX=9606 GN=MRPS21 PE=1 SV=3 | 1.04 | 0.892311206 | 0.939 | 0.7 |
| A0A087WXB0 | Secretory carrier-associated membrane protein OS=Homo sapiens OX=9606 GN=SCAMP1 PE=1 SV=1 | 1.005 | 0.917008272 | 0.888 | 0.[illegible] |
| Q86UU1 | Pleckstrin homology-like domain family B member 1 OS=Homo sapiens OX=9606 GN=PHLDB1 PE=1 SV=1 | 0.937 | 0.892311206 | 1.198 | 0.7 |
| Q6NY19 | KN motif and ankyrin repeat domain-containing protein 3 OS=Homo sapiens OX=9606 GN=KANK3 PE=1 SV=1 | 1.212 | 0.750023665 | 1.122 | 0.8 |
| I3L466 | Histone acetyltransferase (Fragment) OS=Homo sapiens OX=9606 GN=CREBBP PE=1 SV=2 | | | | |
| Q96D53 | Atypical kinase COQ8B, mitochondrial OS=Homo sapiens OX=9606 GN=COQ8B PE=1 SV=2 | 1.157 | 0.703015003 | 0.894 | 0.7 |
| O43505 | Beta-1,4-glucuronyltransferase 1 OS=Homo sapiens OX=9606 GN=B4GAT1 PE=1 SV=1 | 0.86 | 0.765235202 | 0.998 | 0.9 |
| A0A087WUZ6 | InaD-like protein OS=Homo sapiens OX=9606 GN=PATJ PE=1 SV=1 | 1.469 | 0.939929386 | 0.864 | 0.8 |
| A0A140TA84 | MICOS complex subunit MIC13 OS=Homo sapiens OX=9606 GN=MICOS13 PE=1 SV=1 | 1.076 | 0.765235202 | 0.813 | 0.7 |
| Q6ZRP7 | Sulfhydryl oxidase 2 OS=Homo sapiens OX=9606 GN=QSOX2 PE=1 SV=3 | 0.904 | 0.765235202 | 1.027 | 0.8 |
| Q9Y3C1 | Nucleolar protein 16 OS=Homo sapiens OX=9606 GN=NOP16 PE=1 SV=2 | 1.05 | 0.766667967 | 1.021 | 0.8 |
| Q9UPQ0 | LIM and calponin homology domains-containing protein 1 OS=Homo sapiens OX=9606 GN=LIMCH1 PE=1 SV=4 | 0.929 | 0.869995963 | 0.851 | 0.7 |
| Q9BZL1 | Ubiquitin-like protein 5 OS=Homo sapiens OX=9606 GN=UBL5 PE=1 SV=1 | 0.999 | 0.918702104 | 0.951 | 0.8 |
| G3V207 | Transmembrane and coiled-coil domain family 3, isoform CRA_c OS=Homo sapiens OX=9606 GN=TMCC3 PE=1 SV=1 | | | | |
| K7ELI9 | Stromal cell-derived factor 2 (Fragment) OS=Homo sapiens OX=9606 GN=SDF2 PE=1 SV=1 | 0.961 | 0.904500837 | 0.972 | 0.8 |
| Q00796 | Sorbitol dehydrogenase OS=Homo sapiens OX=9606 GN=SORD PE=1 SV=4 | 0.981 | 0.970974607 | 0.914 | 0.7 |
| C9JNU9 | Ubiquitin carboxyl-terminal hydrolase 4 OS=Homo sapiens OX=9606 GN=USP4 PE=1 SV=1 | 0.916 | 0.703015003 | 1.012 | 0.9 |
| Q9P265 | Disco-interacting protein 2 homolog B OS=Homo sapiens OX=9606 GN=DIP2B PE=1 SV=3 | 1.203 | 0.873607677 | 1.043 | 0.9 |
| P49356 | Protein farnesyltransferase subunit beta OS=Homo sapiens OX=9606 GN=FNTB PE=1 SV=1 | 0.965 | 0.953739985 | 0.929 | 0.7 |
| H7C543 | Activated leukocyte cell adhesion molecule (Fragment) OS=Homo sapiens OX=9606 GN=ALCAM PE=1 SV=1 | 0.995 | 0.995582922 | 1.065 | 0.8 |
| H3BM52 | ADP-ribosylation factor-like protein 2-binding protein (Fragment) OS=Homo sapiens OX=9606 GN=ARL2BP PE=1 SV=1 | 0.828 | 0.750023665 | 0.749 | 0.7 |
| Q15024 | Exosome complex component RRP42 OS=Homo sapiens OX=9606 GN=EXOSC7 PE=1 SV=3 | | | | |
| Q99757 | Thioredoxin, mitochondrial OS=Homo sapiens OX=9606 GN=TXN2 PE=1 SV=2 | 1.01 | 0.902885462 | 0.95 | 0.7 |
| P05204 | Non-histone chromosomal protein HMG-17 OS=Homo sapiens OX=9606 GN=HMGN2 PE=1 SV=3 | 0.982 | 0.957536424 | 0.972 | 0.9 |
| Q8IWX8 | Calcium homeostasis endoplasmic reticulum protein OS=Homo sapiens OX=9606 GN=CHERP PE=1 SV=3 | 0.903 | 0.9962822 | 0.765 | 0.7 |
| A2A2V1 | Major prion protein (Fragment) OS=Homo sapiens OX=9606 GN=PRNP PE=1 SV=1 | 0.918 | 0.765235202 | 0.799 | 0.7 |
| Q8NE01 | Metal transporter CNNM3 OS=Homo sapiens OX=9606 GN=CNNM3 PE=1 SV=1 | 1.129 | 0.829390484 | 0.609 | 0.7 |
| P62166 | Neuronal calcium sensor 1 OS=Homo sapiens OX=9606 GN=NCS1 PE=1 SV=2 | 1.153 | 0.815069882 | 0.91 | 0.7 |
| Q8IYU8 | Calcium uptake protein 2, mitochondrial OS=Homo sapiens OX=9606 GN=MICU2 PE=1 SV=2 | 0.944 | 0.766667967 | 0.916 | 0.7 |
| Q9NXV6 | CDKN2A-interacting protein OS=Homo sapiens OX=9606 GN=CDKN2AIP PE=1 SV=3 | 0.96 | 0.80706255 | 0.847 | 0.7 |
| Q6R327 | Rapamycin-insensitive companion of mTOR OS=Homo sapiens OX=9606 GN=RICTOR PE=1 SV=1 | 1.308 | 0.750023665 | 0.711 | 0.7 |
| V9GYM8 | Guanine nucleotide exchange factor H1 OS=Homo sapiens OX=9606 GN=ARHGEF2 PE=1 SV=1 | 1.008 | 0.926688896 | 0.958 | 0.7 |
| P15289 | Arylsulfatase A OS=Homo sapiens OX=9606 GN=ARSA PE=1 SV=3 | 0.991 | 0.985330338 | 0.876 | 0.7 |
| H0YCA5 | Spermatogenesis-associated protein 5-like protein 1 (Fragment) OS=Homo sapiens OX=9606 GN=SPATA5L1 PE=1 SV=1 | 1.075 | 0.895384791 | 0.981 | 0.7 |
| Q5VUD6 | Divergent protein kinase domain 1B OS=Homo sapiens OX=9606 GN=DIPK1B PE=1 SV=3 | 0.996 | 0.994361756 | 0.886 | 0.9 |
| A3KN83 | Protein strawberry notch homolog 1 OS=Homo sapiens OX=9606 GN=SBNO1 PE=1 SV=1 | 0.986 | 0.902016292 | 1.047 | 0.7 |
| E7ENY0 | Alpha-adducin OS=Homo sapiens OX=9606 GN=ADD1 PE=1 SV=1 | 1.122 | 0.750023665 | 1.097 | 0.7 |
| F8VWZ8 | Rho GTPase-activating protein 29 OS=Homo sapiens OX=9606 GN=ARHGAP29 PE=1 SV=1 | 0.93 | 0.904500837 | 0.906 | 0.7 |
| Q6P1N0 | Coiled-coil and C2 domain-containing protein 1A OS=Homo sapiens OX=9606 GN=CC2D1A PE=1 SV=1 | 1.118 | 0.889450577 | 0.956 | 0.7 |
| Q15102 | Platelet-activating factor acetylhydrolase IB subunit gamma OS=Homo sapiens OX=9606 GN=PAFAH1B3 PE=1 SV=1 | 0.993 | 0.972946678 | 0.956 | 0.7 |
| M0R3B2 | E3 ubiquitin-protein ligase NOSIP (Fragment) OS=Homo sapiens OX=9606 GN=NOSIP PE=1 SV=8 | 0.988 | 0.992258896 | 0.821 | 0.9 |
| Q96QG7 | Myotubularin-related protein 9 OS=Homo sapiens OX=9606 GN=MTMR9 PE=1 SV=1 | 0.972 | 0.949254122 | 1.078 | 0.8 |
| K7EPH3 | N-acetylglucosamine-1-phosphodiester alpha-N-acetylglucosaminidase (Fragment) OS=Homo sapiens OX=9606 GN=NAGPA PE=1 | 0.925 | 0.845656075 | 0.951 | 0.8 |
| Q9UEE9 | Craniofacial development protein 1 OS=Homo sapiens OX=9606 GN=CFDP1 PE=1 SV=1 | 1.04 | 0.765235202 | 0.945 | 0.7 |
| A0A3B3IS91 | DNA polymerase subunit gamma-1 OS=Homo sapiens OX=9606 GN=POLG PE=1 SV=1 | 1.034 | 0.99634713 | 1.059 | 0.9 |
| K7ER53 | Nuclear factor of-activated T-cells, cytoplasmic 1 OS=Homo sapiens OX=9606 GN=NFATC1 PE=1 SV=1 | 1.781 | 0.765235202 | 0.547 | 0.7 |
| Q9NUY8 | TBC1 domain family member 23 OS=Homo sapiens OX=9606 GN=TBC1D23 PE=1 SV=3 | 0.96 | 0.889865188 | 0.828 | 0.7 |
| A0A087WV30 | NF-kappa-B essential modulator (Fragment) OS=Homo sapiens OX=9606 GN=IKBKG PE=1 SV=1 | 0.987 | 0.924992773 | 0.863 | 0.7 |
| A0A3B3IU50 | Vacuolar protein sorting-associated protein 13C (Fragment) OS=Homo sapiens OX=9606 GN=VPS13C PE=1 SV=1 | 1.133 | 0.766667967 | 0.87 | 0.7 |
| P08236 | Beta-glucuronidase OS=Homo sapiens OX=9606 GN=GUSB PE=1 SV=2 | 0.977 | 0.921087329 | 0.841 | 0.7 |
| Q9NY12 | H/ACA ribonucleoprotein complex subunit 1 OS=Homo sapiens OX=9606 GN=GAR1 PE=1 SV=1 | 0.971 | 0.916035389 | 1.004 | 0.8 |
| A6NIR2 | Chromosome 1 open reading frame 41, isoform CRA_b OS=Homo sapiens OX=9606 GN=HSPB11 PE=1 SV=1 | 0.971 | 0.702711636 | 0.976 | 0.7 |
| Q9UNK0 | Syntaxin-8 OS=Homo sapiens OX=9606 GN=STX8 PE=1 SV=2 | 0.981 | 0.925659097 | 0.811 | 0.7 |
| Q13322 | Growth factor receptor-bound protein 10 OS=Homo sapiens OX=9606 GN=GRB10 PE=1 SV=2 | 1.05 | 0.902016292 | 1.102 | 0.9 |
| Q5JUP3 | Allograft inflammatory factor 1-like OS=Homo sapiens OX=9606 GN=AIF1L PE=1 SV=1 | 0.95 | 0.948916863 | 0.744 | 0.7 |
| H3BQ50 | Gamma-aminobutyric acid receptor-associated protein-like 2 OS=Homo sapiens OX=9606 GN=GABARAPL2 PE=1 SV=1 | 0.981 | 0.996637279 | 1.197 | 0.7 |
| P56378 | ATP synthase subunit ATP5MPL, mitochondrial OS=Homo sapiens OX=9606 GN=ATP5MPL PE=1 SV=1 | 0.943 | 0.829390484 | 0.956 | 0.8 |
| Q14012 | Calcium/calmodulin-dependent protein kinase type 1 OS=Homo sapiens OX=9606 GN=CAMK1 PE=1 SV=1 | 0.873 | 0.806305724 | 0.987 | 0.8 |
| Q2TAY7 | WD40 repeat-containing protein SMU1 OS=Homo sapiens OX=9606 GN=SMU1 PE=1 SV=2 | 1.064 | 0.780884711 | 0.964 | 0.7 |
| H0Y6Y8 | 39S ribosomal protein L43, mitochondrial (Fragment) OS=Homo sapiens OX=9606 GN=MRPL43 PE=1 SV=1 | 1.043 | 0.930855729 | 1.002 | 0.[illegible] |
| O95104 | SR-related and CTD-associated factor 4 OS=Homo sapiens OX=9606 GN=SCAF4 PE=1 SV=3 | 1.06 | 0.750023665 | 0.942 | 0.7 |
| P40306 | Proteasome subunit beta type-10 OS=Homo sapiens OX=9606 GN=PSMB10 PE=1 SV=1 | 0.982 | 0.820396985 | 0.995 | 0.8 |
| K7EP45 | ESCRT-II complex subunit VPS25 OS=Homo sapiens OX=9606 GN=VPS25 PE=1 SV=1 | 1.015 | 0.935312766 | 0.833 | 0.7 |
| Q9NR50 | Translation initiation factor eIF-2B subunit gamma OS=Homo sapiens OX=9606 GN=EIF2B3 PE=1 SV=1 | 0.924 | 0.921087329 | 0.635 | 0.7 |
| B5MD46 | TBC1 domain family member 10A OS=Homo sapiens OX=9606 GN=TBC1D10A PE=1 SV=1 | 0.979 | 0.961750547 | 0.907 | 0.7 |
| Q9Y316 | Protein MEMO1 OS=Homo sapiens OX=9606 GN=MEMO1 PE=1 SV=1 | 0.912 | 0.840158726 | 0.934 | 0.7 |
| A0A096LNZ9 | Ubiquitin-like protein ISG15 (Fragment) OS=Homo sapiens OX=9606 GN=ISG15 PE=1 SV=6 | 0.919 | 0.765235202 | 1.018 | 0.[illegible] |
| Q5VIR6 | Vacuolar protein sorting-associated protein 53 homolog OS=Homo sapiens OX=9606 GN=VPS53 PE=1 SV=2 | 1.032 | 0.827871327 | 1.121 | 0.8 |
| E9PNP3 | Mth938 domain-containing protein OS=Homo sapiens OX=9606 GN=AAMDC PE=1 SV=1 | 1.01 | 0.827871327 | 1.016 | 0.9 |
| Q96L35 | Receptor protein-tyrosine kinase OS=Homo sapiens OX=9606 GN=EPHB4 PE=1 SV=1 | 1.066 | 0.829390484 | 0.995 | 0.[illegible] |
| Q9P000 | COMM domain-containing protein 9 OS=Homo sapiens OX=9606 GN=COMMD9 PE=1 SV=2 | 0.953 | 0.750023665 | 1.013 | 0.9 |
| F8WCH9 | 2-hydroxyacyl-CoA lyase 1 OS=Homo sapiens OX=9606 GN=HACL1 PE=1 SV=1 | 1.571 | 0.889482529 | 0.709 | 0.7 |
| Q8TCT8 | Signal peptide peptidase-like 2A OS=Homo sapiens OX=9606 GN=SPPL2A PE=1 SV=2 | 0.953 | 0.910115867 | 0.743 | 0.7 |
| A0A0G2JRQ5 | Glutathione transferase OS=Homo sapiens OX=9606 GN=GSTT1 PE=1 SV=1 | 1.038 | 0.765235202 | 0.923 | 0.8 |
| Q14573 | Inositol 1,4,5-trisphosphate receptor type 3 OS=Homo sapiens OX=9606 GN=ITPR3 PE=1 SV=2 | 0.985 | 0.996578046 | 0.858 | 0.7 |
| Q14689 | Disco-interacting protein 2 homolog A OS=Homo sapiens OX=9606 GN=DIP2A PE=1 SV=2 | 1.022 | 0.908555656 | 0.91 | 0.7 |
| J3KP27 | Trafficking protein particle complex subunit OS=Homo sapiens OX=9606 GN=TRAPPC4 PE=1 SV=1 | 0.967 | 0.902016292 | 1.015 | 0.9 |
| F8WES2 | S-methyl-5'-thioadenosine phosphorylase OS=Homo sapiens OX=9606 GN=MTAP PE=4 SV=1 | 0.932 | 0.861400199 | 0.977 | 0.8 |
| Q9H9A6 | Leucine-rich repeat-containing protein 40 OS=Homo sapiens OX=9606 GN=LRRC40 PE=1 SV=1 | 1.023 | 0.80706255 | 1.024 | 0.9 |
| E9PHY0 | Lysosomal acid phosphatase OS=Homo sapiens OX=9606 GN=ACP2 PE=1 SV=1 | 0.986 | 0.756287988 | 0.883 | 0.7 |
| J3KMX3 | Alpha-fetoprotein OS=Homo sapiens OX=9606 GN=AFP PE=1 SV=1 | 1.063 | 0.996578046 | 0.875 | 0.7 |
| P41743 | Protein kinase C iota type OS=Homo sapiens OX=9606 GN=PRKCI PE=1 SV=2 | 0.807 | 0.765235202 | 1.039 | 0.7 |
| P98170 | E3 ubiquitin-protein ligase XIAP OS=Homo sapiens OX=9606 GN=XIAP PE=1 SV=2 | 1.237 | 0.765235202 | 0.936 | 0.8 |

| | | | | | |
|---|---|---|---|---|---|
| O15127 | Secretory carrier-associated membrane protein 2 OS=Homo sapiens OX=9606 GN=SCAMP2 PE=1 SV=2 | 0.899 | 0.785314375 | 0.929 | 0.7 |
| H7C0C7 | CCR4-NOT transcription complex subunit 11 (Fragment) OS=Homo sapiens OX=9606 GN=CNOT11 PE=1 SV=1 | 0.991 | 0.906273459 | 0.832 | 0.7 |
| P35251 | Replication factor C subunit 1 OS=Homo sapiens OX=9606 GN=RFC1 PE=1 SV=4 | 1.006 | 0.978003638 | 0.944 | 0.9 |
| H7BY55 | Complement decay-accelerating factor OS=Homo sapiens OX=9606 GN=CD55 PE=1 SV=2 | 0.961 | 0.902016292 | 0.96 | 0.9 |
| E5RHT9 | Transcription and mRNA export factor ENY2 OS=Homo sapiens OX=9606 GN=ENY2 PE=1 SV=1 | 0.961 | 0.906273459 | 1.061 | 0.7 |
| Q05932 | Folylpolyglutamate synthase, mitochondrial OS=Homo sapiens OX=9606 GN=FPGS PE=1 SV=3 | 0.769 | 0.917008272 | 0.565 | 0.7 |
| O75663 | TIP41-like protein OS=Homo sapiens OX=9606 GN=TIPRL PE=1 SV=2 | 0.963 | 0.765235202 | 0.963 | 0.9 |
| Q9H0R6 | Glutamyl-tRNA(Gln) amidotransferase subunit A, mitochondrial OS=Homo sapiens OX=9606 GN=QRSL1 PE=1 SV=2 | 0.848 | 0.80706255 | 0.98 | 0.9 |
| Q8WUF5 | RelA-associated inhibitor OS=Homo sapiens OX=9606 GN=PPP1R13L PE=1 SV=4 | 1.081 | 0.951537478 | 0.556 | |
| Q93050-1 | Isoform 2 of V-type proton ATPase 116 kDa subunit a1 OS=Homo sapiens OX=9606 GN=ATP6V0A1 | 0.988 | 0.856216225 | 0.837 | 0.7 |
| Q96CS3 | FAS-associated factor 2 OS=Homo sapiens OX=9606 GN=FAF2 PE=1 SV=2 | 0.913 | 0.895384791 | 0.796 | 0.8 |
| Q99836 | Myeloid differentiation primary response protein MyD88 OS=Homo sapiens OX=9606 GN=MYD88 PE=1 SV=1 | 1.105 | 0.642276967 | 1.174 | 0.7 |
| K7EJE1 | Complex I-B16.6 OS=Homo sapiens OX=9606 GN=NDUFA13 PE=3 SV=1 | 1.052 | 0.99319413 | 0.836 | 0.7 |
| A0A1B0GTG4 | Ras association domain-containing protein 5 OS=Homo sapiens OX=9606 GN=RASSF5 PE=1 SV=1 | | | | |
| O95759 | TBC1 domain family member 8 OS=Homo sapiens OX=9606 GN=TBC1D8 PE=1 SV=3 | 1.381 | 0.765235202 | 1.201 | 0.9 |
| Q6NUK1 | Calcium-binding mitochondrial carrier protein SCaMC-1 OS=Homo sapiens OX=9606 GN=SLC25A24 PE=1 SV=2 | 1.002 | 0.902016292 | 0.903 | 0.7 |
| H0Y5K5 | Endoplasmic reticulum-Golgi intermediate compartment protein 3 (Fragment) OS=Homo sapiens OX=9606 GN=ERGIC3 PE=1 SV=1 | 0.904 | 0.854598401 | 0.908 | 0.7 |
| Q7Z5G4 | Golgin subfamily A member 7 OS=Homo sapiens OX=9606 GN=GOLGA7 PE=1 SV=2 | 1.025 | 0.994361756 | 0.998 | 0.9 |
| M0QYV8 | Fukutin-related protein (Fragment) OS=Homo sapiens OX=9606 GN=FKRP PE=1 SV=1 | 0.098 | 0.898015977 | 0.454 | 0.7 |
| Q13423 | NAD(P) transhydrogenase, mitochondrial OS=Homo sapiens OX=9606 GN=NNT PE=1 SV=3 | 0.943 | 0.765235202 | 0.802 | 0.7 |
| Q86US8 | Telomerase-binding protein EST1A OS=Homo sapiens OX=9606 GN=SMG6 PE=1 SV=2 | 1.102 | 0.927756817 | 0.825 | 0.7 |
| A0A3B3ISV5 | Uncharacterized protein OS=Homo sapiens OX=9606 PE=4 SV=1 | | | | |
| P42684 | Tyrosine-protein kinase ABL2 OS=Homo sapiens OX=9606 GN=ABL2 PE=1 SV=1 | 1.036 | 0.951324063 | 0.852 | 0.9 |
| Q8NBQ5 | Estradiol 17-beta-dehydrogenase 11 OS=Homo sapiens OX=9606 GN=HSD17B11 PE=1 SV=3 | 0.993 | 0.994361756 | 0.834 | 0.7 |
| P03905 | NADH-ubiquinone oxidoreductase chain 4 OS=Homo sapiens OX=9606 GN=MT-ND4 PE=1 SV=1 | | | | |
| F8WCI5 | Protein C3orf33 OS=Homo sapiens OX=9606 GN=C3orf33 PE=1 SV=1 | 0.951 | 0.866300162 | 1.023 | 0.9 |
| O60331 | Phosphatidylinositol 4-phosphate 5-kinase type-1 gamma OS=Homo sapiens OX=9606 GN=PIP5K1C PE=1 SV=2 | | | | |
| B0QZ40 | COMM domain-containing protein 6 OS=Homo sapiens OX=9606 GN=COMMD6 PE=1 SV=1 | 0.886 | 0.765235202 | 0.956 | 0.8 |
| Q9UBD5 | Origin recognition complex subunit 3 OS=Homo sapiens OX=9606 GN=ORC3 PE=1 SV=1 | 1.149 | 0.765235202 | 1.087 | 0.9 |
| Q6P5S2 | Protein LEG1 homolog OS=Homo sapiens OX=9606 GN=LEG1 PE=1 SV=2 | 1.323 | 0.740168868 | 0.755 | 0.7 |
| Q9Y5Y7 | Lymphatic vessel endothelial hyaluronic acid receptor 1 OS=Homo sapiens OX=9606 GN=LYVE1 PE=1 SV=2 | 0.97 | 0.811611894 | 1.085 | 0.7 |
| E9PIE3 | Caveolae-associated protein 3 OS=Homo sapiens OX=9606 GN=CAVIN3 PE=1 SV=1 | 0.909 | 0.765235202 | 0.888 | 0.7 |
| Q9Y5S2 | Serine/threonine-protein kinase MRCK beta OS=Homo sapiens OX=9606 GN=CDC42BPB PE=1 SV=2 | 0.976 | 0.905396375 | 0.935 | 0.8 |
| O14874 | [3-methyl-2-oxobutanoate dehydrogenase [lipoamide]] kinase, mitochondrial OS=Homo sapiens OX=9606 GN=BCKDK PE=1 SV=2 | 0.839 | 0.943791335 | 0.787 | 0.9 |
| Q96IJ6 | Mannose-1-phosphate guanyltransferase alpha OS=Homo sapiens OX=9606 GN=GMPPA PE=1 SV=1 | 1.092 | 0.98478736 | 0.971 | 0.8 |
| Q9BRP8 | Partner of Y14 and mago OS=Homo sapiens OX=9606 GN=PYM1 PE=1 SV=1 | 1.013 | 0.998901726 | 0.79 | 0.7 |
| Q92504 | Zinc transporter SLC39A7 OS=Homo sapiens OX=9606 GN=SLC39A7 PE=1 SV=2 | | | | |
| Q9Y3T9 | Nucleolar complex protein 2 homolog OS=Homo sapiens OX=9606 GN=NOC2L PE=1 SV=4 | 1.01 | 0.95071661 | 0.959 | 0.7 |
| Q309B1 | Tripartite motif-containing protein 16-like protein OS=Homo sapiens OX=9606 GN=TRIM16L PE=2 SV=3 | 0.939 | 0.9962822 | 0.868 | 0.7 |
| Q70J99 | Protein unc-13 homolog D OS=Homo sapiens OX=9606 GN=UNC13D PE=1 SV=1 | 1.121 | 0.845356491 | 0.888 | 0.7 |
| A0A1W2PR36 | Guanidinoacetate N-methyltransferase OS=Homo sapiens OX=9606 GN=GAMT PE=1 SV=1 | 1.198 | 0.860909277 | 0.978 | 0.8 |
| H0YLT2 | Cyclin-dependent kinase 12 (Fragment) OS=Homo sapiens OX=9606 GN=CDK12 PE=1 SV=1 | 100 | | 0.01 | |
| Q8TEA8 | D-aminoacyl-tRNA deacylase 1 OS=Homo sapiens OX=9606 GN=DTD1 PE=1 SV=2 | 1.149 | 0.765235202 | 1.023 | 0.7 |
| F8VVT9 | Arf-GAP with GTPase, ANK repeat and PH domain-containing protein 2 OS=Homo sapiens OX=9606 GN=AGAP2 PE=1 SV=1 | 0.989 | 0.955325044 | 0.898 | 0.7 |
| E7ERP7 | Apolipoprotein E (Fragment) OS=Homo sapiens OX=9606 GN=APOE PE=1 SV=1 | 1.089 | 0.805488634 | 0.915 | 0.7 |
| A0A1B0GU74 | Voltage-dependent P/Q-type calcium channel subunit alpha OS=Homo sapiens OX=9606 GN=CACNA1A PE=1 SV=1 | 0.973 | 0.924631778 | 1.126 | 0.7 |
| O00115 | Deoxyribonuclease-2-alpha OS=Homo sapiens OX=9606 GN=DNASE2 PE=1 SV=2 | 0.989 | 0.985324977 | 0.976 | 0.8 |
| G8JLB3 | tRNA pseudouridine synthase A (Fragment) OS=Homo sapiens OX=9606 GN=PUS1 PE=1 SV=1 | 0.956 | 0.902016292 | 0.903 | 0.7 |
| P49406 | 39S ribosomal protein L19, mitochondrial OS=Homo sapiens OX=9606 GN=MRPL19 PE=1 SV=2 | 0.981 | 0.902016292 | 0.968 | 0.7 |
| Q9BTE3 | Mini-chromosome maintenance complex-binding protein OS=Homo sapiens OX=9606 GN=MCMBP PE=1 SV=2 | 0.954 | 0.80706255 | 1.234 | 0.7 |
| Q9NQS1 | Cell death regulator Aven OS=Homo sapiens OX=9606 GN=AVEN PE=1 SV=1 | 1.188 | 0.750023665 | 0.756 | 0.7 |
| Q9BT22 | Chitobiosyldiphosphodolichol beta-mannosyltransferase OS=Homo sapiens OX=9606 GN=ALG1 PE=1 SV=2 | 0.96 | 0.827871327 | 0.974 | 0.9 |
| A0A499FJI4 | RCR-type E3 ubiquitin transferase OS=Homo sapiens OX=9606 GN=MYCBP2 PE=1 SV=1 | | | | |
| Q8WWI5 | Choline transporter-like protein 1 OS=Homo sapiens OX=9606 GN=SLC44A1 PE=1 SV=1 | 0.939 | 0.805488634 | 0.946 | 0.7 |
| Q494V2 | Cilia- and flagella-associated protein 100 OS=Homo sapiens OX=9606 GN=CFAP100 PE=1 SV=1 | 1.411 | 0.949934729 | 0.882 | 0.9 |
| Q15654 | Thyroid receptor-interacting protein 6 OS=Homo sapiens OX=9606 GN=TRIP6 PE=1 SV=3 | | | | |
| Q8WW59 | SPRY domain-containing protein 4 OS=Homo sapiens OX=9606 GN=SPRYD4 PE=1 SV=2 | 0.879 | 0.925981427 | 0.859 | 0.7 |
| P00403 | Cytochrome c oxidase subunit 2 OS=Homo sapiens OX=9606 GN=MT-CO2 PE=1 SV=1 | 0.976 | 0.980556519 | 0.806 | 0.7 |
| Q8N5H7 | SH2 domain-containing protein 3C OS=Homo sapiens OX=9606 GN=SH2D3C PE=1 SV=1 | 1.025 | 0.954224541 | 1.412 | 0.7 |
| I3L4A1 | Charged multivesicular body protein 6 (Fragment) OS=Homo sapiens OX=9606 GN=CHMP6 PE=1 SV=1 | 1.059 | 0.985324977 | 0.925 | 0.8 |
| H3BTU7 | Katanin p80 WD40 repeat-containing subunit B1 (Fragment) OS=Homo sapiens OX=9606 GN=KATNB1 PE=1 SV=1 | 1.115 | 0.765235202 | 0.996 | 0.9 |
| E5RGY0 | Derlin OS=Homo sapiens OX=9606 GN=DERL1 PE=1 SV=1 | | | | |
| B7ZAX5 | N-acetylgalactosamine kinase OS=Homo sapiens OX=9606 GN=GALK2 PE=1 SV=1 | 0.957 | 0.845656075 | 0.936 | 0.7 |
| P62891 | 60S ribosomal protein L39 OS=Homo sapiens OX=9606 GN=RPL39 PE=1 SV=2 | 1.039 | 0.882454148 | 1.084 | 0.9 |
| Q8N0Z6 | Tetratricopeptide repeat protein 5 OS=Homo sapiens OX=9606 GN=TTC5 PE=1 SV=2 | 0.895 | 0.765235202 | 0.971 | 0.7 |
| P62312 | U6 snRNA-associated Sm-like protein LSm6 OS=Homo sapiens OX=9606 GN=LSM6 PE=1 SV=1 | 0.955 | 0.829390484 | 0.949 | 0.7 |
| A0A3B3IRR6 | UPF0488 protein C8orf33 OS=Homo sapiens OX=9606 GN=C8orf33 PE=1 SV=1 | 1.255 | 0.869995963 | 0.807 | 0.7 |
| Q9UJV9 | Probable ATP-dependent RNA helicase DDX41 OS=Homo sapiens OX=9606 GN=DDX41 PE=1 SV=2 | 0.967 | 0.87017509 | 1.042 | 0.7 |
| H0YKH9 | Alpha-mannosidase 2x (Fragment) OS=Homo sapiens OX=9606 GN=MAN2A2 PE=1 SV=1 | 1.017 | 0.904500837 | 0.974 | 0.9 |
| M0R2M4 | Zinc finger protein 551 (Fragment) OS=Homo sapiens OX=9606 GN=ZNF551 PE=1 SV=1 | 0.966 | 0.766250742 | 0.925 | 0.7 |
| Q969X5 | Endoplasmic reticulum-Golgi intermediate compartment protein 1 OS=Homo sapiens OX=9606 GN=ERGIC1 PE=1 SV=1 | | | | |
| Q29RF7 | Sister chromatid cohesion protein PDS5 homolog A OS=Homo sapiens OX=9606 GN=PDS5A PE=1 SV=1 | 1.296 | 0.765235202 | 0.999 | 0.7 |
| Q92600 | CCR4-NOT transcription complex subunit 9 OS=Homo sapiens OX=9606 GN=CNOT9 PE=1 SV=1 | 1.074 | 0.766667967 | 1.029 | 0.9 |
| Q9HCE0 | Ectopic P granules protein 5 homolog OS=Homo sapiens OX=9606 GN=EPG5 PE=1 SV=2 | 0.933 | 0.956825234 | 0.862 | 0.7 |
| Q8WVC6 | Dephospho-CoA kinase domain-containing protein OS=Homo sapiens OX=9606 GN=DCAKD PE=1 SV=1 | 0.878 | 0.870355966 | 0.903 | 0.7 |
| Q9P016 | Thymocyte nuclear protein 1 OS=Homo sapiens OX=9606 GN=THYN1 PE=1 SV=1 | 0.977 | 0.903226585 | 0.894 | 0.7 |
| O60637 | Tetraspanin-3 OS=Homo sapiens OX=9606 GN=TSPAN3 PE=2 SV=1 | 1.005 | 0.921087329 | 0.96 | 0.7 |
| J3QQN7 | Lysophospholipase D GDPD1 OS=Homo sapiens OX=9606 GN=GDPD1 PE=1 SV=1 | 1.06 | 0.904500837 | 1.065 | 0.7 |
| Q9HBW9 | Adhesion G protein-coupled receptor L4 OS=Homo sapiens OX=9606 GN=ADGRL4 PE=1 SV=3 | 1.424 | 0.750023665 | 1.216 | 0.7 |
| Q14571 | Inositol 1,4,5-trisphosphate receptor type 2 OS=Homo sapiens OX=9606 GN=ITPR2 PE=1 SV=2 | 0.995 | 0.887719165 | 1.088 | 0.7 |
| Q9HCS7 | Pre-mRNA-splicing factor SYF1 OS=Homo sapiens OX=9606 GN=XAB2 PE=1 SV=2 | 1.051 | 0.895384791 | 0.979 | 0.8 |
| Q15648 | Mediator of RNA polymerase II transcription subunit 1 OS=Homo sapiens OX=9606 GN=MED1 PE=1 SV=4 | 0.84 | 0.89681291 | 1.164 | 0.8 |
| Q14197 | Peptidyl-tRNA hydrolase ICT1, mitochondrial OS=Homo sapiens OX=9606 GN=MRPL58 PE=1 SV=1 | 1.11 | 0.838881659 | 0.886 | 0.7 |
| Q8WWQ2 | Inactive heparanase-2 OS=Homo sapiens OX=9606 GN=HPSE2 PE=1 SV=3 | 0.984 | 0.979006266 | 0.707 | 0.7 |
| H0Y6A0 | ADP-ribosylation factor GTPase-activating protein 3 (Fragment) OS=Homo sapiens OX=9606 GN=ARFGAP3 PE=1 SV=1 | 0.877 | 0.866300162 | 0.913 | 0.8 |
| G3V1B8 | Protein ZNRD2 OS=Homo sapiens OX=9606 GN=ZNRD2 PE=1 SV=1 | 1.001 | 0.95980123 | 1.065 | 0.7 |
| A0A1W2PS92 | 39S ribosomal protein L9, mitochondrial (Fragment) OS=Homo sapiens OX=9606 GN=MRPL9 PE=1 SV=1 | | | | |
| C9JXC1 | Ribonucleoside-diphosphate reductase (Fragment) OS=Homo sapiens OX=9606 GN=RRM2 PE=1 SV=1 | | | | |
| Q96Q11 | CCA tRNA nucleotidyltransferase 1, mitochondrial OS=Homo sapiens OX=9606 GN=TRNT1 PE=1 SV=2 | 1.013 | 0.921087329 | 0.977 | 0.7 |
| Q9H2C0 | Gigaxonin OS=Homo sapiens OX=9606 GN=GAN PE=1 SV=1 | 1.285 | 0.750023665 | 0.89 | 0.7 |
| Q01968 | Inositol polyphosphate 5-phosphatase OCRL OS=Homo sapiens OX=9606 GN=OCRL PE=1 SV=3 | | | | |
| P56962 | Syntaxin-17 OS=Homo sapiens OX=9606 GN=STX17 PE=1 SV=2 | 0.85 | 0.829390484 | 0.739 | 0.7 |
| Q9UHP3 | Ubiquitin carboxyl-terminal hydrolase 25 OS=Homo sapiens OX=9606 GN=USP25 PE=1 SV=4 | 0.937 | 0.889482529 | 0.727 | 0.7 |
| Q7L2J0 | 7SK snRNA methylphosphate capping enzyme OS=Homo sapiens OX=9606 GN=MEPCE PE=1 SV=1 | 100 | | | |
| A0A0A0MS30 | Protein PRRC2C OS=Homo sapiens OX=9606 GN=PRRC2C PE=1 SV=1 | 0.952 | 0.845656075 | 1.1 | 0.8 |
| Q9BX67 | Junctional adhesion molecule C OS=Homo sapiens OX=9606 GN=JAM3 PE=1 SV=1 | 0.925 | 0.765235202 | 0.825 | 0.7 |

| | | | | | |
|---|---|---|---|---|---|
| P42858 | Huntingtin OS=Homo sapiens OX=9606 GN=HTT PE=1 SV=2 | | | | |
| P27449 | V-type proton ATPase 16 kDa proteolipid subunit OS=Homo sapiens OX=9606 GN=ATP6V0C PE=1 SV=1 | 0.78 | 0.761475165 | 0.815 | 0.7 |
| Q5T6J7 | Probable gluconokinase OS=Homo sapiens OX=9606 GN=IDNK PE=1 SV=1 | 1.06 | 0.994361756 | 0.765 | 0.7 |
| Q12873 | Chromodomain-helicase-DNA-binding protein 3 OS=Homo sapiens OX=9606 GN=CHD3 PE=1 SV=3 | 1.206 | 0.785314375 | 0.95 | 0.9 |
| Q8N4N8 | Kinesin-like protein KIF2B OS=Homo sapiens OX=9606 GN=KIF2B PE=1 SV=3 | 1.084 | 0.901934028 | 0.803 | 0.7 |
| A0A140T947 | Pre-mRNA-splicing factor ATP-dependent RNA helicase DHX16 (Fragment) OS=Homo sapiens OX=9606 GN=DHX16 PE=1 SV=1 | 0.826 | 0.849526028 | 0.906 | 0.7 |
| E9PCW1 | Golgi SNAP receptor complex member 1 OS=Homo sapiens OX=9606 GN=GOSR1 PE=1 SV=1 | | | | |
| Q7L0Y3 | tRNA methyltransferase 10 homolog C OS=Homo sapiens OX=9606 GN=TRMT10C PE=1 SV=2 | 0.945 | 0.882454148 | 1.046 | 0.8 |
| A0A087WY31 | YTH domain-containing family protein 3 OS=Homo sapiens OX=9606 GN=YTHDF3 PE=1 SV=1 | 0.651 | 0.889482529 | 0.97 | 0.8 |
| Q8NBN3 | Transmembrane protein 87A OS=Homo sapiens OX=9606 GN=TMEM87A PE=1 SV=3 | 1.114 | 0.750023665 | 0.915 | 0.7 |
| Q8NEZ4 | Histone-lysine N-methyltransferase 2C OS=Homo sapiens OX=9606 GN=KMT2C PE=1 SV=3 | 0.998 | 0.987438982 | 0.989 | 0.9 |
| E1CEI4 | Gamma-ECS OS=Homo sapiens OX=9606 GN=GCLC PE=1 SV=1 | 1.187 | 0.815069882 | 0.725 | 0.7 |
| C9IYN9 | SET and MYND domain-containing protein 5 (Fragment) OS=Homo sapiens OX=9606 GN=SMYD5 PE=1 SV=1 | 0.876 | 0.905369216 | 1.018 | 0.8 |
| A0A087WV05 | Uncharacterized protein OS=Homo sapiens OX=9606 PE=3 SV=1 | 0.801 | 0.765235202 | 0.884 | 0.7 |
| A0A087WYV4 | Dehydrogenase/reductase SDR family member 11 OS=Homo sapiens OX=9606 GN=DHRS11 PE=1 SV=1 | | | | |
| D6REK3 | Spliceosome-associated protein CWC27 homolog OS=Homo sapiens OX=9606 GN=CWC27 PE=1 SV=1 | 0.985 | 0.938276447 | 0.885 | 0.7 |
| P50281 | Matrix metalloproteinase-14 OS=Homo sapiens OX=9606 GN=MMP14 PE=1 SV=3 | 0.995 | 0.902016292 | 0.987 | 0.[illegible] |
| Q9H501 | ESF1 homolog OS=Homo sapiens OX=9606 GN=ESF1 PE=1 SV=1 | | | | |
| H3BNI3 | RNA-binding protein with serine-rich domain 1 (Fragment) OS=Homo sapiens OX=9606 GN=RNPS1 PE=1 SV=1 | 0.997 | 0.917008272 | 0.924 | 0.7 |
| Q5TC63 | Growth hormone-regulated TBC protein 1 OS=Homo sapiens OX=9606 GN=GRTP1 PE=1 SV=4 | 0.975 | 0.766667967 | 0.965 | 0.8 |
| Q969V1 | Melanin-concentrating hormone receptor 2 OS=Homo sapiens OX=9606 GN=MCHR2 PE=1 SV=1 | 1.018 | 0.933097749 | 0.988 | 0.7 |
| Q8WWV3 | Reticulon-4-interacting protein 1, mitochondrial OS=Homo sapiens OX=9606 GN=RTN4IP1 PE=1 SV=2 | 1.032 | 0.854119562 | 0.957 | 0.8 |
| Q495W5 | Alpha-(1,3)-fucosyltransferase 11 OS=Homo sapiens OX=9606 GN=FUT11 PE=1 SV=1 | 1.038 | 0.792509319 | 0.954 | 0.7 |
| B8ZZV5 | 39S ribosomal protein L30, mitochondrial OS=Homo sapiens OX=9606 GN=MRPL30 PE=1 SV=1 | 0.892 | 0.754126086 | 0.89 | 0.7 |
| A0A087X1H5 | Arf-GAP with coiled-coil, ANK repeat and PH domain-containing protein 2 OS=Homo sapiens OX=9606 GN=ACAP2 PE=1 SV=1 | 1.145 | 0.750023665 | 1.101 | 0.7 |
| D6RAW2 | ADP-ribose pyrophosphatase, mitochondrial (Fragment) OS=Homo sapiens OX=9606 GN=NUDT9 PE=1 SV=1 | 0.971 | 0.847589821 | 0.966 | 0.7 |
| Q5H9U9 | Probable ATP-dependent RNA helicase DDX60-like OS=Homo sapiens OX=9606 GN=DDX60L PE=2 SV=2 | 1.124 | 0.782369209 | 0.651 | 0.7 |
| A0A0A0MRJ3 | Neuron navigator 1 OS=Homo sapiens OX=9606 GN=NAV1 PE=1 SV=1 | 1.213 | 0.765235202 | 1.075 | 0.7 |
| O15258 | Protein RER1 OS=Homo sapiens OX=9606 GN=RER1 PE=1 SV=1 | | | | |
| Q96HP4 | Oxidoreductase NAD-binding domain-containing protein 1 OS=Homo sapiens OX=9606 GN=OXNAD1 PE=1 SV=1 | 0.969 | 0.910115867 | 0.708 | 0.7 |
| A3KMH1 | von Willebrand factor A domain-containing protein 8 OS=Homo sapiens OX=9606 GN=VWA8 PE=1 SV=2 | 1.06 | 0.983953845 | 0.705 | 0.7 |
| Q14683 | Structural maintenance of chromosomes protein 1A OS=Homo sapiens OX=9606 GN=SMC1A PE=1 SV=2 | | | | |
| Q9UG63 | ATP-binding cassette sub-family F member 2 OS=Homo sapiens OX=9606 GN=ABCF2 PE=1 SV=2 | 1.038 | 0.849526028 | 0.952 | 0.7 |
| Q10471 | Polypeptide N-acetylgalactosaminyltransferase 2 OS=Homo sapiens OX=9606 GN=GALNT2 PE=1 SV=1 | | | | |
| Q9HAB8 | Phosphopantothenate--cysteine ligase OS=Homo sapiens OX=9606 GN=PPCS PE=1 SV=2 | 0.976 | 0.927756817 | 0.918 | 0.7 |
| A0A087WTA5 | Translation initiation factor eIF-2B subunit delta OS=Homo sapiens OX=9606 GN=EIF2B4 PE=1 SV=1 | | | | |
| A0A0C4DGB1 | rRNA adenine N(6)-methyltransferase OS=Homo sapiens OX=9606 GN=DIMT1 PE=1 SV=1 | 1.049 | 0.797038721 | 0.865 | 0.7 |
| P31152 | Mitogen-activated protein kinase 4 OS=Homo sapiens OX=9606 GN=MAPK4 PE=1 SV=2 | 0.661 | 0.750023665 | 0.887 | 0.7 |
| Q9Y6X9 | ATPase MORC2 OS=Homo sapiens OX=9606 GN=MORC2 PE=1 SV=2 | 1.132 | 0.774484855 | 0.865 | 0.7 |
| Q9UMS0 | NFU1 iron-sulfur cluster scaffold homolog, mitochondrial OS=Homo sapiens OX=9606 GN=NFU1 PE=1 SV=2 | | | | |
| H0YB34 | 2-iminobutanoate/2-iminopropanoate deaminase (Fragment) OS=Homo sapiens OX=9606 GN=RIDA PE=1 SV=1 | 1.066 | 0.883217803 | 0.88 | 0.7 |
| C9JXM3 | Rab-like protein 3 OS=Homo sapiens OX=9606 GN=RABL3 PE=1 SV=1 | 0.891 | 0.869995963 | 0.915 | 0.7 |
| P51798 | H(+)/Cl(-) exchange transporter 7 OS=Homo sapiens OX=9606 GN=CLCN7 PE=1 SV=2 | 1.046 | 0.87038391 | 0.936 | 0.7 |
| O00139 | Kinesin-like protein KIF2A OS=Homo sapiens OX=9606 GN=KIF2A PE=1 SV=3 | 0.935 | 0.750023665 | 0.991 | 0.8 |
| Q8N3V7 | Synaptopodin OS=Homo sapiens OX=9606 GN=SYNPO PE=1 SV=2 | 0.962 | 0.809976184 | 0.9 | 0.7 |
| A0A494C0A9 | Core-binding factor subunit beta OS=Homo sapiens OX=9606 GN=CBFB PE=1 SV=1 | 0.908 | 0.886561233 | 1.037 | 0.[illegible] |
| D6RIH3 | Transcription elongation factor A protein-like 4 (Fragment) OS=Homo sapiens OX=9606 GN=TCEAL4 PE=1 SV=1 | 1.085 | 0.765235202 | 0.922 | 0.7 |
| H7BZW6 | Histone deacetylase complex subunit SAP18 (Fragment) OS=Homo sapiens OX=9606 GN=SAP18 PE=1 SV=2 | 1.002 | 0.921897381 | 0.905 | 0.7 |
| P82650 | 28S ribosomal protein S22, mitochondrial OS=Homo sapiens OX=9606 GN=MRPS22 PE=1 SV=1 | 0.981 | 0.895384791 | 0.933 | 0.7 |
| P01042 | Kininogen-1 OS=Homo sapiens OX=9606 GN=KNG1 PE=1 SV=2 | 1.051 | 0.889482529 | 0.809 | 0.7 |
| X6R824 | Probable E3 ubiquitin-protein ligase HECTD2 OS=Homo sapiens OX=9606 GN=HECTD2 PE=1 SV=1 | | | | |
| O60673 | DNA polymerase zeta catalytic subunit OS=Homo sapiens OX=9606 GN=REV3L PE=1 SV=2 | 0.926 | 0.983953845 | 0.88 | 0.7 |
| Q9Y5V0 | Zinc finger protein 706 OS=Homo sapiens OX=9606 GN=ZNF706 PE=1 SV=1 | 1.025 | 0.829390484 | 1.105 | 0.7 |
| J3KMW0 | (Lyso)-N-acylphosphatidylethanolamine lipase OS=Homo sapiens OX=9606 GN=ABHD4 PE=1 SV=2 | | | | |
| O95429 | BAG family molecular chaperone regulator 4 OS=Homo sapiens OX=9606 GN=BAG4 PE=1 SV=1 | 1.009 | 0.980966594 | 0.948 | 0.7 |
| Q4J6C6 | Prolyl endopeptidase-like OS=Homo sapiens OX=9606 GN=PREPL PE=1 SV=1 | 0.971 | 0.953521381 | 0.949 | 0.7 |
| Q8N2C7 | Protein unc-80 homolog OS=Homo sapiens OX=9606 GN=UNC80 PE=1 SV=2 | 1.267 | 0.952913364 | 0.962 | 0.9 |
| Q7Z4F1 | Low-density lipoprotein receptor-related protein 10 OS=Homo sapiens OX=9606 GN=LRP10 PE=1 SV=2 | 1.126 | 0.829390484 | 0.877 | 0.8 |
| Q15050 | Ribosome biogenesis regulatory protein homolog OS=Homo sapiens OX=9606 GN=RRS1 PE=1 SV=2 | | | | |
| O95721 | Synaptosomal-associated protein 29 OS=Homo sapiens OX=9606 GN=SNAP29 PE=1 SV=1 | 1.01 | 0.992258896 | 1.005 | 0.8 |
| A0A075B6Q2 | Formin-binding protein 1-like (Fragment) OS=Homo sapiens OX=9606 GN=FNBP1L PE=1 SV=1 | 1.033 | 0.895384791 | 0.935 | 0.7 |
| P48651 | Phosphatidylserine synthase 1 OS=Homo sapiens OX=9606 GN=PTDSS1 PE=1 SV=1 | 1.024 | 0.978562349 | 1.012 | 0.9 |
| O43657 | Tetraspanin-6 OS=Homo sapiens OX=9606 GN=TSPAN6 PE=1 SV=1 | 0.858 | 0.765235202 | 0.886 | 0.7 |
| Q96PZ0 | Pseudouridylate synthase 7 homolog OS=Homo sapiens OX=9606 GN=PUS7 PE=1 SV=2 | | | | |
| O75037 | Kinesin-like protein KIF21B OS=Homo sapiens OX=9606 GN=KIF21B PE=1 SV=2 | 1.01 | 0.91273701 | 0.876 | 0.7 |
| P22004 | Bone morphogenetic protein 6 OS=Homo sapiens OX=9606 GN=BMP6 PE=1 SV=1 | | | | |
| H0YB53 | Protein MTSS 1 (Fragment) OS=Homo sapiens OX=9606 GN=MTSS1 PE=1 SV=1 | 1.079 | 0.983953845 | 0.986 | 0.9 |
| Q9Y535 | DNA-directed RNA polymerase III subunit RPC8 OS=Homo sapiens OX=9606 GN=POLR3H PE=1 SV=1 | 0.917 | 0.905369216 | 0.977 | 0.7 |
| B4DXR6 | Cyclin N-terminal domain containing, isoform CRA_b OS=Homo sapiens OX=9606 GN=CNTD1 PE=2 SV=1 | 0.975 | 0.859846322 | 0.979 | 0.9 |

ıbjected to the phosphoteomics analysis. GN: gene name. n = 4 cell preparations (2 female and 2 male cells).*P* -values < 0.05.

**Table S31A. Overrepresentated GO biological process in ITE-dysregulated phosphoproteins at 4hr.***

| Biological process | Reference gene # (20589) | Mapped Gene # | Expected Gene # | Fold Enrichment | Direction | Raw *P* value | FDR *P* Value |
|---|---|---|---|---|---|---|---|
| cytoskeleton organization (GO:0007010) | 1240 | 18 | 3.79 | 4.74 | + | 2.22E-08 | 3.48E-04 |
| regulation of cytoskeleton organization (GO:0051493) | 540 | 11 | 1.65 | 6.66 | + | 7.74E-07 | 2.43E-03 |
| regulation of cellular component organization (GO:0051128) | 2364 | 22 | 7.23 | 3.04 | + | 1.02E-06 | 2.65E-03 |
| negative regulation of cytoskeleton organization (GO:0051494) | 158 | 7 | 0.48 | 14.48 | + | 6.95E-07 | 2.72E-03 |
| negative regulation of biological process (GO:0048519) | 5314 | 35 | 16.26 | 2.15 | + | 6.84E-07 | 3.57E-03 |
| negative regulation of cellular process (GO:0048523) | 4732 | 33 | 14.48 | 2.28 | + | 4.72E-07 | 3.70E-03 |
| cellular localization (GO:0051641) | 2655 | 23 | 8.12 | 2.83 | + | 1.81E-06 | 4.05E-03 |
| regulation of organelle organization (GO:0033043) | 1189 | 15 | 3.64 | 4.12 | + | 2.42E-06 | 4.75E-03 |
| regulation of protein polymerization (GO:0032271) | 201 | 7 | 0.62 | 11.38 | + | 3.26E-06 | 5.68E-03 |
| cellular component organization (GO:0016043) | 5523 | 34 | 16.9 | 2.01 | + | 5.61E-06 | 7.99E-03 |
| cellular component organization or biogenesis (GO:0071840) | 5727 | 35 | 17.52 | 2 | + | 6.43E-06 | 8.41E-03 |
| regulation of cellular component biogenesis (GO:0044087) | 951 | 13 | 2.91 | 4.47 | + | 5.50E-06 | 8.62E-03 |
| negative regulation of supramolecular fiber organization (GO:1902904) | 162 | 6 | 0.5 | 12.1 | + | 1.23E-05 | 1.28E-02 |
| regulation of protein depolymerization (GO:1901879) | 92 | 5 | 0.28 | 17.76 | + | 1.17E-05 | 1.31E-02 |
| negative regulation of organelle organization (GO:0010639) | 345 | 8 | 1.06 | 7.58 | + | 1.15E-05 | 1.39E-02 |
| actin filament-based process (GO:0030029) | 611 | 10 | 1.87 | 5.35 | + | 1.71E-05 | 1.58E-02 |
| organelle organization (GO:0006996) | 3026 | 23 | 9.26 | 2.48 | + | 1.65E-05 | 1.62E-02 |
| negative regulation of programmed cell death (GO:0043069) | 918 | 12 | 2.81 | 4.27 | + | 2.07E-05 | 1.80E-02 |
| regulation of supramolecular fiber organization (GO:1902903) | 383 | 8 | 1.17 | 6.83 | + | 2.40E-05 | 1.98E-02 |
| cell division (GO:0051301) | 514 | 9 | 1.57 | 5.72 | + | 2.81E-05 | 2.10E-02 |
| cell death (GO:0008219) | 1108 | 13 | 3.39 | 3.83 | + | 2.74E-05 | 2.15E-02 |
| macromolecule localization (GO:0033036) | 2345 | 19 | 7.18 | 2.65 | + | 5.09E-05 | 2.66E-02 |
| regulation of programmed cell death (GO:0043067) | 1498 | 15 | 4.58 | 3.27 | + | 3.74E-05 | 2.67E-02 |
| establishment of localization in cell (GO:0051649) | 1752 | 16 | 5.36 | 2.98 | + | 5.88E-05 | 2.71E-02 |
| establishment or maintenance of cell polarity (GO:0007163) | 210 | 6 | 0.64 | 9.34 | + | 5.05E-05 | 2.73E-02 |
| negative regulation of cell death (GO:0060548) | 1022 | 12 | 3.13 | 3.84 | + | 5.83E-05 | 2.77E-02 |
| cellular macromolecule localization (GO:0070727) | 1925 | 17 | 5.89 | 2.89 | + | 4.95E-05 | 2.77E-02 |
| regulation of protein-containing complex disassembly (GO:0043244) | 130 | 5 | 0.4 | 12.57 | + | 5.75E-05 | 2.82E-02 |
| response to stress (GO:0006950) | 3466 | 24 | 10.61 | 2.26 | + | 5.58E-05 | 2.82E-02 |
| positive regulation of biological process (GO:0048518) | 6304 | 35 | 19.29 | 1.81 | + | 4.90E-05 | 2.84E-02 |
| protein localization (GO:0008104) | 1919 | 17 | 5.87 | 2.9 | + | 4.76E-05 | 2.87E-02 |
| negative regulation of protein kinase activity (GO:0006469) | 207 | 6 | 0.63 | 9.47 | + | 4.67E-05 | 2.93E-02 |
| actin cytoskeleton organization (GO:0030036) | 549 | 9 | 1.68 | 5.36 | + | 4.65E-05 | 3.04E-02 |
| negative regulation of macromolecule metabolic process (GO:0010605) | 2764 | 21 | 8.46 | 2.48 | + | 4.58E-05 | 3.13E-02 |

| | | | | | | | |
|---|---|---|---|---|---|---|---|
| supramolecular fiber organization (GO:0097435) | 580 | 9 | 1.77 | 5.07 | + | 7.04E-05 | 3.16E-02 |
| negative regulation of protein depolymerization (GO:1901880) | 70 | 4 | 0.21 | 18.67 | + | 7.71E-05 | 3.18E-02 |
| nitrogen compound transport (GO:0071705) | 1589 | 15 | 4.86 | 3.09 | + | 7.31E-05 | 3.18E-02 |
| establishment of localization (GO:0051234) | 3997 | 26 | 12.23 | 2.13 | + | 7.60E-05 | 3.22E-02 |
| negative regulation of apoptotic process (GO:0043066) | 898 | 11 | 2.75 | 4 | + | 8.54E-05 | 3.35E-02 |
| negative regulation of kinase activity (GO:0033673) | 231 | 6 | 0.71 | 8.49 | + | 8.43E-05 | 3.39E-02 |
| cell cycle (GO:0007049) | 1249 | 13 | 3.82 | 3.4 | + | 9.28E-05 | 3.55E-02 |
| localization (GO:0051179) | 4566 | 28 | 13.97 | 2 | + | 9.58E-05 | 3.58E-02 |
| negative regulation of protein polymerization (GO:0032272) | 76 | 4 | 0.23 | 17.2 | + | 1.05E-04 | 3.64E-02 |
| response to external stimulus (GO:0009605) | 2469 | 19 | 7.55 | 2.51 | + | 1.02E-04 | 3.64E-02 |
| positive regulation of cellular process (GO:0048522) | 5670 | 32 | 17.35 | 1.84 | + | 1.01E-04 | 3.68E-02 |
| regulation of cell death (GO:0010941) | 1654 | 15 | 5.06 | 2.96 | + | 1.14E-04 | 3.90E-02 |
| regulation of apoptotic process (GO:0042981) | 1468 | 14 | 4.49 | 3.12 | + | 1.20E-04 | 3.93E-02 |
| transport (GO:0006810) | 3840 | 25 | 11.75 | 2.13 | + | 1.18E-04 | 3.95E-02 |
| negative regulation of protein-containing complex disassembly (GO:0043[illegible] | 80 | 4 | 0.24 | 16.34 | + | 1.26E-04 | 4.04E-02 |
| regulation of cellular catabolic process (GO:0031329) | 786 | 10 | 2.41 | 4.16 | + | 1.37E-04 | 4.30E-02 |

*PANTHER was used to analyze overrepresented biological process for ITE-dysregulated phosphoproteins. Gene ontology (GO) analysis was performed using Statistical Overrepresentation test (Fisher's exact test, FDR $P < 0.05$). n = 4 cell preparations.

**Table S31B. Overrepresentated GO molecular function in ITE-dysregulated phosphoproteins at 4 hr.***

| Molecular function | Reference Gene # (20589) | Mapped Gene # | Expected Gene # | Fold Enrichment | Direction | Raw *P* value | FDR *P* Value |
|---|---|---|---|---|---|---|---|
| RNA binding (GO:0003723) | 1666 | 27 | 5.1 | 5.3 | + | 1.02E-13 | 5.09E-10 |
| cadherin binding (GO:0045296) | 325 | 13 | 0.99 | 13.07 | + | 2.41E-11 | 5.99E-08 |
| cell adhesion molecule binding (GO:0050839) | 552 | 13 | 1.69 | 7.7 | + | 1.25E-08 | 2.08E-05 |
| actin filament binding (GO:0051015) | 222 | 8 | 0.68 | 11.78 | + | 4.79E-07 | 5.96E-04 |
| enzyme binding (GO:0019899) | 2103 | 21 | 6.43 | 3.26 | + | 6.31E-07 | 6.27E-04 |
| protein-containing complex binding (GO:0044877) | 1312 | 16 | 4.01 | 3.99 | + | 1.60E-06 | 1.14E-03 |
| protein kinase binding (GO:0019901) | 708 | 12 | 2.17 | 5.54 | + | 1.54E-06 | 1.27E-03 |
| calmodulin binding (GO:0005516) | 201 | 7 | 0.62 | 11.38 | + | 3.26E-06 | 2.03E-03 |
| kinase binding (GO:0019900) | 792 | 12 | 2.42 | 4.95 | + | 4.79E-06 | 2.65E-03 |
| mRNA binding (GO:0003729) | 326 | 8 | 1 | 8.02 | + | 7.69E-06 | 3.48E-03 |
| nucleic acid binding (GO:0003676) | 4007 | 28 | 12.26 | 2.28 | + | 6.99E-06 | 3.48E-03 |
| protein binding (GO:0005515) | 14448 | 59 | 44.21 | 1.33 | + | 1.02E-05 | 3.92E-03 |
| actin binding (GO:0003779) | 451 | 9 | 1.38 | 6.52 | + | 1.02E-05 | 4.22E-03 |
| structural constituent of cytoskeleton (GO:0005200) | 109 | 5 | 0.33 | 14.99 | + | 2.56E-05 | 9.08E-03 |
| heterocyclic compound binding (GO:1901363) | 5983 | 34 | 18.31 | 1.86 | + | 3.87E-05 | 1.28E-02 |
| GTPase binding (GO:0051020) | 308 | 7 | 0.94 | 7.43 | + | 4.78E-05 | 1.49E-02 |
| organic cyclic compound binding (GO:0097159) | 6054 | 34 | 18.52 | 1.84 | + | 7.11E-05 | 2.08E-02 |
| kinase inhibitor activity (GO:0019210) | 75 | 4 | 0.23 | 17.43 | + | 9.95E-05 | 2.75E-02 |
| molecular adaptor activity (GO:0060090) | 488 | 8 | 1.49 | 5.36 | + | 1.27E-04 | 3.33E-02 |
| protein serine/threonine kinase inhibitor activity (GO:0030291) | 34 | 3 | 0.1 | 28.84 | + | 0.000195 | 0.0486 |

*PANTHER was used to analyze overrepresented molecular function for ITE-dysregulated phosphoproteins. Gene ontology (GO) analysis was performed using Statistical Overrepresentation test (Fisher's exact test, FDR $P < 0.05$). n = 4 cell preparations.

**Table S31C. Overrepresentated GO cell component in ITE-dysregulated phosphoproteins at 4 hr.***

| Cell component | Reference Gene # (20589) | Mapped Gene # | Expected Gene # | Fold Enrichment | Direction | Raw *P* value | FDR *P* Value |
|---|---|---|---|---|---|---|---|
| cytosol (GO:0005829) | 5458 | 44 | 16.7 | 2.63 | + | 9.72E-13 | 1.99E-09 |
| cell-substrate junction (GO:0030055) | 433 | 15 | 1.32 | 11.32 | + | 4.14E-12 | 2.82E-09 |
| focal adhesion (GO:0005925) | 423 | 15 | 1.29 | 11.59 | + | 3.00E-12 | 3.07E-09 |
| non-membrane-bounded organelle (GO:0043228) | 5226 | 41 | 15.99 | 2.56 | + | 4.27E-11 | 1.45E-08 |
| anchoring junction (GO:0070161) | 1336 | 22 | 4.09 | 5.38 | + | 3.24E-11 | 1.66E-08 |
| intracellular non-membrane-bounded organelle (GO:0043232) | 5225 | 41 | 15.99 | 2.56 | + | 4.24E-11 | 1.73E-08 |
| cytoskeleton (GO:0005856) | 2401 | 27 | 7.35 | 3.68 | + | 4.73E-10 | 1.38E-07 |
| cell junction (GO:0030054) | 2136 | 25 | 6.54 | 3.83 | + | 1.26E-09 | 3.21E-07 |
| supramolecular complex (GO:0099080) | 1410 | 20 | 4.31 | 4.64 | + | 4.27E-09 | 9.69E-07 |
| cell cortex (GO:0005938) | 317 | 10 | 0.97 | 10.31 | + | 5.25E-08 | 1.07E-05 |
| intracellular organelle (GO:0043229) | 13254 | 59 | 40.56 | 1.45 | + | 8.89E-08 | 1.65E-05 |
| cytoplasm (GO:0005737) | 12097 | 56 | 37.02 | 1.51 | + | 2.34E-07 | 3.99E-05 |
| microtubule cytoskeleton (GO:0015630) | 1374 | 25 | 4.2 | 4.04 | + | 5.70E-07 | 7.77E-05 |
| supramolecular fiber (GO:0099512) | 1047 | 15 | 3.2 | 4.68 | + | 5.03E-07 | 7.91E-05 |
| supramolecular polymer (GO:0099081) | 1056 | 15 | 3.23 | 4.64 | + | 5.60E-07 | 8.17E-05 |
| intracellular anatomical structure (GO:0005622) | 14945 | 61 | 45.73 | 1.33 | + | 1.15E-06 | 1.46E-04 |
| organelle (GO:0043226) | 14064 | 59 | 43.03 | 1.37 | + | 1.94E-06 | 2.33E-04 |
| actin cytoskeleton (GO:0015629) | 510 | 10 | 1.56 | 6.41 | + | 3.63E-06 | 4.12E-04 |
| cell-cell junction (GO:0005911) | 508 | 9 | 1.55 | 5.79 | + | 2.56E-05 | 2.76E-03 |
| cortical cytoskeleton (GO:0030863) | 111 | 5 | 0.34 | 14.72 | + | 2.78E-05 | 2.84E-03 |
| nucleus (GO:0005634) | 7682 | 40 | 23.51 | 1.7 | + | 3.54E-05 | 3.45E-03 |
| microtubule organizing center (GO:0005815) | 840 | 11 | 2.57 | 4.28 | + | 4.72E-05 | 4.38E-03 |
| polysome (GO:0005844) | 67 | 4 | 0.21 | 19.51 | + | 6.55E-05 | 5.82E-03 |
| vesicle (GO:0031982) | 3975 | 26 | 12.16 | 2.14 | + | 7.22E-05 | 5.90E-03 |
| intrinsic component of membrane (GO:0031224) | 5958 | 5 | 18.23 | 0.27 | - | 7.06E-05 | 6.01E-03 |
| intracellular membrane-bounded organelle (GO:0043231) | 12154 | 52 | 37.19 | 1.4 | + | 9.16E-05 | 7.20E-03 |
| integral component of membrane (GO:0016021) | 5799 | 5 | 17.74 | 0.28 | - | 1.15E-04 | 8.71E-03 |
| cortical actin cytoskeleton (GO:0030864) | 81 | 4 | 0.25 | 16.14 | + | 1.32E-04 | 9.65E-03 |
| extracellular membrane-bounded organelle (GO:0065010) | 2124 | 17 | 6.5 | 2.62 | + | 1.66E-04 | 9.99E-03 |
| polymeric cytoskeletal fiber (GO:0099513) | 808 | 10 | 2.47 | 4.04 | + | 1.71E-04 | 1.00E-02 |
| extracellular exosome (GO:0070062) | 2100 | 17 | 6.43 | 2.65 | + | 1.45E-04 | 1.02E-02 |
| extracellular organelle (GO:0043230) | 2124 | 17 | 6.5 | 2.62 | + | 1.66E-04 | 1.03E-02 |
| extracellular vesicle (GO:1903561) | 2123 | 17 | 6.5 | 2.62 | + | 1.65E-04 | 1.06E-02 |
| axon (GO:0030424) | 649 | 9 | 1.99 | 4.53 | + | 1.63E-04 | 1.07E-02 |

| | | | | | | | |
|---|---|---|---|---|---|---|---|
| cluster of actin-based cell projections (GO:0098862) | 163 | 5 | 0.5 | 10.02 | + | 1.61E-04 | 1.10E-02 |
| membrane-bounded organelle (GO:0043227) | 13230 | 54 | 40.48 | 1.33 | + | 1.93E-04 | 1.10E-02 |
| ribonucleoprotein complex (GO:1990904) | 714 | 9 | 2.18 | 4.12 | + | 3.28E-04 | 1.81E-02 |
| spindle (GO:0005819) | 428 | 7 | 1.31 | 5.35 | + | 3.51E-04 | 1.84E-02 |
| brush border (GO:0005903) | 105 | 4 | 0.32 | 12.45 | + | 3.44E-04 | 1.85E-02 |
| perinuclear region of cytoplasm (GO:0048471) | 742 | 9 | 2.27 | 3.96 | + | 4.32E-04 | 2.21E-02 |
| plasma membrane raft (GO:0044853) | 115 | 4 | 0.35 | 11.37 | + | 4.80E-04 | 2.39E-02 |
| nuclear lumen (GO:0031981) | 4494 | 26 | 13.75 | 1.89 | + | 5.97E-04 | 2.44E-02 |
| cAMP-dependent protein kinase complex (GO:0005952) | 10 | 2 | 0.03 | 65.36 | + | 5.93E-04 | 2.47E-02 |
| nucleotide-activated protein kinase complex (GO:0031588) | 10 | 2 | 0.03 | 65.36 | + | 5.93E-04 | 2.52E-02 |
| cell periphery (GO:0071944) | 6460 | 33 | 19.77 | 1.67 | + | 5.83E-04 | 2.53E-02 |
| membrane-enclosed lumen (GO:0031974) | 5614 | 30 | 17.18 | 1.75 | + | 5.81E-04 | 2.58E-02 |
| intracellular organelle lumen (GO:0070013) | 5614 | 30 | 17.18 | 1.75 | + | 5.81E-04 | 2.64E-02 |
| organelle lumen (GO:0043233) | 5614 | 30 | 17.18 | 1.75 | + | 5.81E-04 | 2.70E-02 |
| membrane microdomain (GO:0098857) | 332 | 6 | 1.02 | 5.91 | + | 5.70E-04 | 2.71E-02 |
| membrane raft (GO:0045121) | 331 | 6 | 1.01 | 5.92 | + | 5.61E-04 | 2.73E-02 |
| centrosome (GO:0005813) | 634 | 8 | 1.94 | 4.12 | + | 7.16E-04 | 2.87E-02 |
| myofibril (GO:0030016) | 235 | 5 | 0.72 | 6.95 | + | 8.26E-04 | 3.24E-02 |
| contractile fiber (GO:0043292) | 244 | 5 | 0.75 | 6.7 | + | 9.73E-04 | 3.75E-02 |
| cytoplasmic ribonucleoprotein granule (GO:0036464) | 250 | 5 | 0.76 | 6.54 | + | 1.08E-03 | 3.88E-02 |
| cell projection (GO:0042995) | 2391 | 17 | 7.32 | 2.32 | + | 1.07E-03 | 3.90E-02 |
| protein-containing complex (GO:0032991) | 5771 | 30 | 17.66 | 1.7 | + | 1.06E-03 | 3.93E-02 |
| plasma membrane bounded cell projection cytoplasm (GO:0032838) | 248 | 5 | 0.76 | 6.59 | + | 1.04E-03 | 3.95E-02 |
| intracellular vesicle (GO:0097708) | 2496 | 17 | 7.64 | 2.23 | + | 1.36E-03 | 4.65E-02 |
| costamere (GO:0043034) | 16 | 2 | 0.05 | 40.85 | + | 1.36E-03 | 4.70E-02 |
| plasma membrane region (GO:0098590) | 1257 | 11 | 3.85 | 2.86 | + | 1.44E-03 | 4.75E-02 |
| cytoplasmic vesicle (GO:0031410) | 2493 | 17 | 7.63 | 2.23 | + | 1.35E-03 | 4.76E-02 |
| ribonucleoprotein granule (GO:0035770) | 267 | 5 | 0.82 | 6.12 | + | 1.44E-03 | 4.82E-02 |

*PANTHER was used to analyze overrepresented cell component for ITE-dysregulated phosphoproteins. Gene ontology (GO) analysis was performed using Statistical Overrepresentation test (Fisher's exact test, FDR $P < 0.05$). n = 4 cell preparations.

**Table S31D. Overrepresentated GO protein class in ITE-dysregulated phosphoproteins at 4 hr.***

| Protein class | Reference Gene # (20589) | Mapped Gene # | Expected Gene # | Fold Enrichment | Direction | Raw *P* value | FDR *P* Value |
|---|---|---|---|---|---|---|---|
| cytoskeletal protein (PC00085) | 627 | 13 | 1.92 | 6.78 | + | 5.43E-08 | 1.06E-05 |
| actin or actin-binding cytoskeletal protein (PC00041) | 278 | 6 | 0.85 | 7.05 | + | 2.26E-04 | 2.21E-02 |

*PANTHER was used to analyze overrepresented protein class for ITE-dysregulated phosphoproteins. Gene ontology (GO) analysis was performed using Statistical Overrepresentation test (Fisher's exact test, FDR $P < 0.05$). n = 4 cell preparations.

**Table S31E. Overreentated GO Reactome pathway in ITE-dysregulated phosphoproteins at 4 hr.***

| Reactome pathway | Reference Gene # (20589) | Mapped Gene # | Expected Gene # | Fold Enrichment | Direction | Raw *P* value | FDR *P* Value |
|---|---|---|---|---|---|---|---|
| Signaling by Rho GTPases, Miro GTPases and RHOBTB3 (R-HSA-9716542) | 685 | 13 | 2.1 | 6.2 | + | 1.48E-07 | 1.85E-04 |
| Signaling by Rho GTPases (R-HSA-194315) | 670 | 13 | 2.05 | 6.34 | + | 1.15E-07 | 2.88E-04 |
| Infectious disease (R-HSA-5663205) | 895 | 14 | 2.74 | 5.11 | + | 4.63E-07 | 3.85E-04 |
| Aggrephagy (R-HSA-9646399) | 44 | 4 | 0.13 | 29.71 | + | 1.38E-05 | 8.62E-03 |
| Apoptotic execution phase (R-HSA-75153) | 51 | 4 | 0.16 | 25.63 | + | 2.39E-05 | 1.19E-02 |
| Mitotic Prometaphase (R-HSA-68877) | 199 | 6 | 0.61 | 9.85 | + | 3.77E-05 | 1.34E-02 |
| Disease (R-HSA-1643685) | 1679 | 16 | 5.14 | 3.11 | + | 3.53E-05 | 1.47E-02 |
| RHO GTPases activate PAKs (R-HSA-5627123) | 21 | 3 | 0.06 | 46.69 | + | 5.23E-05 | 1.63E-02 |
| Vesicle-mediated transport (R-HSA-5653656) | 725 | 10 | 2.22 | 4.51 | + | 7.11E-05 | 1.77E-02 |
| RHO GTPase cycle (R-HSA-9012999) | 448 | 8 | 1.37 | 5.84 | + | 7.10E-05 | 1.97E-02 |
| Selective autophagy (R-HSA-9663891) | 80 | 4 | 0.24 | 16.34 | + | 1.26E-04 | 2.42E-02 |
| Membrane Trafficking (R-HSA-199991) | 626 | 9 | 1.92 | 4.7 | + | 1.25E-04 | 2.59E-02 |
| M Phase (R-HSA-68886) | 375 | 7 | 1.15 | 6.1 | + | 1.59E-04 | 2.65E-02 |
| Signal Transduction (R-HSA-162582) | 2500 | 19 | 7.65 | 2.48 | + | 1.21E-04 | 2.74E-02 |
| Regulation of PLK1 Activity at G2/M Transition (R-HSA-2565942) | 85 | 4 | 0.26 | 15.38 | + | 1.58E-04 | 2.82E-02 |
| Cell Cycle, Mitotic (R-HSA-69278) | 517 | 8 | 1.58 | 5.06 | + | 1.88E-04 | 2.92E-02 |
| Apoptotic cleavage of cellular proteins (R-HSA-111465) | 37 | 3 | 0.11 | 26.5 | + | 2.47E-04 | 3.62E-02 |
| RHO GTPase Effectors (R-HSA-195258) | 288 | 6 | 0.88 | 6.81 | + | 2.72E-04 | 3.77E-02 |
| Diseases of signal transduction by growth factor receptors and second messengers (R-HSA-5663202) | 418 | 7 | 1.28 | 5.47 | + | 3.05E-04 | 3.80E-02 |
| HCMV Early Events (R-HSA-9609690) | 101 | 4 | 0.31 | 12.94 | + | 2.98E-04 | 3.92E-02 |

*PANTHER was used to analyze overrepresented reactome pathway for ITE-dysregulated phosphoproteins. Gene ontology (GO) analysis was performed using Statistical Overrepresentation test (Fisher's exact test, FDR $P < 0.05$). n = 4 cell preparations.

**Fig. 32A. Overrepresentated GO biological process in ITE-dysregulated phosphoproteins at 24hr.***

| Biological process | Reference gene # (20589) | Mapped Gene # | Expected Gene # | Fold Enrichment | Direction | Raw *P* value | FDR *P* Value |
|---|---|---|---|---|---|---|---|
| organelle organization (GO:0006996) | 3026 | 33 | 9.85 | 3.35 | + | 2.84E-11 | 2.22E-07 |
| cellular component organization or biogenesis (GO:0071840) | 5727 | 45 | 18.64 | 2.41 | + | 2.74E-11 | 4.29E-07 |
| cellular component organization (GO:0016043) | 5523 | 42 | 17.97 | 2.34 | + | 1.63E-09 | 8.52E-06 |
| regulation of organelle organization (GO:0033043) | 1189 | 19 | 3.87 | 4.91 | + | 5.34E-09 | 2.10E-05 |
| cellular component biogenesis (GO:0044085) | 2633 | 27 | 8.57 | 3.15 | + | 1.80E-08 | 5.64E-05 |
| chromosome organization (GO:0051276) | 444 | 12 | 1.44 | 8.31 | + | 2.32E-08 | 6.06E-05 |
| regulation of cellular component organization (GO:0051128) | 2364 | 25 | 7.69 | 3.25 | + | 4.29E-08 | 9.61E-05 |
| cytoskeleton organization (GO:0007010) | 1240 | 18 | 4.04 | 4.46 | + | 6.34E-08 | 1.24E-04 |
| chromatin remodeling (GO:0006338) | 323 | 10 | 1.05 | 9.51 | + | 1.14E-07 | 1.79E-04 |
| cell division (GO:0051301) | 514 | 12 | 1.67 | 7.17 | + | 1.10E-07 | 1.92E-04 |
| regulation of DNA metabolic process (GO:0051052) | 541 | 12 | 1.76 | 6.82 | + | 1.89E-07 | 2.69E-04 |
| negative regulation of organelle organization (GO:0010639) | 345 | 10 | 1.12 | 8.91 | + | 2.06E-07 | 2.69E-04 |
| cell cycle (GO:0007049) | 1249 | 17 | 4.06 | 4.18 | + | 3.95E-07 | 4.76E-04 |
| cell cycle process (GO:0022402) | 841 | 14 | 2.74 | 5.12 | + | 4.94E-07 | 4.85E-04 |
| cellular process (GO:0009987) | 15044 | 65 | 48.96 | 1.33 | + | 4.87E-07 | 5.09E-04 |
| chromosome condensation (GO:0030261) | 43 | 5 | 0.14 | 35.73 | + | 4.74E-07 | 5.31E-04 |
| negative regulation of DNA metabolic process (GO:0051053) | 148 | 7 | 0.48 | 14.53 | + | 6.97E-07 | 6.43E-04 |
| cellular component assembly (GO:0022607) | 2394 | 23 | 7.79 | 2.95 | + | 1.03E-06 | 8.95E-04 |
| negative regulation of cellular process (GO:0048523) | 4732 | 33 | 15.4 | 2.14 | + | 2.95E-06 | 2.44E-03 |
| chromatin organization (GO:0006325) | 585 | 11 | 1.9 | 5.78 | + | 3.12E-06 | 2.45E-03 |
| nucleosome organization (GO:0034728) | 125 | 6 | 0.41 | 14.75 | + | 4.22E-06 | 3.15E-03 |
| positive regulation of cellular process (GO:0048522) | 5670 | 36 | 18.45 | 1.95 | + | 6.86E-06 | 4.89E-03 |
| microtubule cytoskeleton organization (GO:0000226) | 547 | 10 | 1.78 | 5.62 | + | 1.17E-05 | 7.97E-03 |
| positive regulation of biological process (GO:0048518) | 6304 | 38 | 20.51 | 1.85 | + | 1.29E-05 | 8.43E-03 |
| nucleosome assembly (GO:0006334) | 93 | 5 | 0.3 | 16.52 | + | 1.66E-05 | 8.67E-03 |
| negative regulation of cellular component organization (GO:0051129) | 690 | 11 | 2.25 | 4.9 | + | 1.45E-05 | 8.73E-03 |
| regulation of protein metabolic process (GO:0051246) | 2587 | 22 | 8.42 | 2.61 | + | 1.41E-05 | 8.85E-03 |
| negative regulation of nitrogen compound metabolic process (GO:0051172) | 2409 | 21 | 7.84 | 2.68 | + | 1.64E-05 | 8.89E-03 |
| spindle organization (GO:0007051) | 160 | 6 | 0.52 | 11.52 | + | 1.64E-05 | 9.17E-03 |
| negative regulation of biological process (GO:0048519) | 5314 | 34 | 17.29 | 1.97 | + | 1.58E-05 | 9.18E-03 |
| cellular developmental process (GO:0048869) | 3542 | 26 | 11.53 | 2.26 | + | 2.66E-05 | 1.34E-02 |
| cell-cell junction organization (GO:0045216) | 176 | 6 | 0.57 | 10.48 | + | 2.75E-05 | 1.35E-02 |
| protein localization (GO:0008104) | 1919 | 18 | 6.24 | 2.88 | + | 3.08E-05 | 1.46E-02 |
| cellular macromolecule localization (GO:0070727) | 1925 | 18 | 6.26 | 2.87 | + | 3.21E-05 | 1.48E-02 |
| regulation of protein modification process (GO:0031399) | 1563 | 16 | 5.09 | 3.15 | + | 3.36E-05 | 1.51E-02 |
| regulation of cellular process (GO:0050794) | 11187 | 53 | 36.4 | 1.46 | + | 3.61E-05 | 1.57E-02 |
| negative regulation of macromolecule metabolic process (GO:0010605) | 2764 | 22 | 8.99 | 2.45 | + | 3.97E-05 | 1.68E-02 |
| DNA repair (GO:0006281) | 508 | 9 | 1.65 | 5.44 | + | 4.23E-05 | 1.75E-02 |
| cellular response to stress (GO:0033554) | 1599 | 16 | 5.2 | 3.07 | + | 4.42E-05 | 1.78E-02 |
| DNA metabolic process (GO:0006259) | 794 | 11 | 2.58 | 4.26 | + | 5.15E-05 | 2.02E-02 |

| | | | | | | | |
|---|---|---|---|---|---|---|---|
| cellular senescence (GO:0090398) | 60 | 4 | 0.2 | 20.49 | + | 5.55E-05 | 2.07E-02 |
| homotypic cell-cell adhesion (GO:0034109) | 60 | 4 | 0.2 | 20.49 | + | 5.55E-05 | 2.12E-02 |
| mitotic cell cycle process (GO:1903047) | 533 | 9 | 1.73 | 5.19 | + | 6.09E-05 | 2.17E-02 |
| microtubule-based process (GO:0007017) | 809 | 11 | 2.63 | 4.18 | + | 6.08E-05 | 2.22E-02 |
| cell-cell junction assembly (GO:0007043) | 126 | 5 | 0.41 | 12.19 | + | 6.70E-05 | 2.33E-02 |
| negative regulation of cellular metabolic process (GO:0031324) | 2264 | 19 | 7.37 | 2.58 | + | 7.99E-05 | 2.61E-02 |
| negative regulation of molecular function (GO:0044092) | 1150 | 13 | 3.74 | 3.47 | + | 7.89E-05 | 2.63E-02 |
| regulation of macromolecule metabolic process (GO:0060255) | 6214 | 36 | 20.22 | 1.78 | + | 7.85E-05 | 2.67E-02 |
| regulation of biological process (GO:0050789) | 11806 | 54 | 38.42 | 1.41 | + | 9.42E-05 | 3.01E-02 |
| cellular response to stimulus (GO:0051716) | 6569 | 37 | 21.38 | 1.73 | + | 1.04E-04 | 3.26E-02 |
| positive regulation of macromolecule metabolic process (GO:0010604) | 3542 | 25 | 11.53 | 2.17 | + | 1.17E-04 | 3.59E-02 |
| protein-DNA complex subunit organization (GO:0071824) | 233 | 6 | 0.76 | 7.91 | + | 1.25E-04 | 3.76E-02 |
| negative regulation of nucleobase-containing compound metabolic process (GO:0045934) | 1562 | 15 | 5.08 | 2.95 | + | 1.27E-04 | 3.77E-02 |
| positive regulation of cellular component organization (GO:0051130) | 1054 | 12 | 3.43 | 3.5 | + | 1.45E-04 | 4.21E-02 |
| negative regulation of metabolic process (GO:0009892) | 2988 | 22 | 9.72 | 2.26 | + | 1.55E-04 | 4.33E-02 |
| oncogene-induced cell senescence (GO:0090402) | 4 | 2 | 0.01 | > 100 | + | 1.54E-04 | 4.40E-02 |
| cellular response to DNA damage stimulus (GO:0006974) | 756 | 10 | 2.46 | 4.06 | + | 1.70E-04 | 4.67E-02 |
| negative regulation of chromosome organization (GO:2001251) | 82 | 4 | 0.27 | 14.99 | + | 1.76E-04 | 4.75E-02 |
| negative regulation of cytoskeleton organization (GO:0051494) | 158 | 5 | 0.51 | 9.72 | + | 1.87E-04 | 4.98E-02 |

*PANTHER was used to enrich biological process for ITE-dysregulated phosphoproteins.  Gene ontology (GO) analysis was performed using Statistical Overrepresentation test (Fisher's exact test, FDR $P < 0.05$).  n = 4 cell preparations.

**Table S32B. Overrepresentated GO molecular function in ITE-dysregulated phosphoproteins at 24 hr.***

| Molecular function | Reference Gene # (20589) | Mapped Gene # | Expected Gene # | Fold Enrichment | Direction | Raw *P* value | FDR *P* Value |
|---|---|---|---|---|---|---|---|
| RNA binding (GO:0003723) | 1666 | 28 | 5.42 | 5.16 | + | 7.83E-14 | 3.89E-10 |
| cadherin binding (GO:0045296) | 325 | 14 | 1.06 | 13.24 | + | 3.52E-12 | 8.74E-09 |
| enzyme binding (GO:0019899) | 2103 | 26 | 6.84 | 3.8 | + | 7.56E-10 | 9.40E-07 |
| organic cyclic compound binding (GO:0097159) | 6054 | 44 | 19.7 | 2.23 | + | 1.04E-09 | 1.03E-06 |
| heterocyclic compound binding (GO:1901363) | 5983 | 44 | 19.47 | 2.26 | + | 7.36E-10 | 1.22E-06 |
| cell adhesion molecule binding (GO:0050839) | 552 | 14 | 1.8 | 7.79 | + | 2.95E-09 | 2.45E-06 |
| protein domain specific binding (GO:0019904) | 689 | 15 | 2.24 | 6.69 | + | 5.59E-09 | 3.97E-06 |
| nucleic acid binding (GO:0003676) | 4007 | 33 | 13.04 | 2.53 | + | 4.43E-08 | 2.75E-05 |
| protein kinase binding (GO:0019901) | 708 | 14 | 2.3 | 6.08 | + | 6.30E-08 | 3.48E-05 |
| kinase binding (GO:0019900) | 792 | 14 | 2.58 | 5.43 | + | 2.43E-07 | 1.21E-04 |
| molecular adaptor activity (GO:0060090) | 488 | 11 | 1.59 | 6.93 | + | 5.55E-07 | 2.51E-04 |
| structural molecule activity (GO:0005198) | 796 | 13 | 2.59 | 5.02 | + | 1.66E-06 | 6.88E-04 |
| protein-containing complex binding (GO:0044877) | 1312 | 16 | 4.27 | 3.75 | + | 3.85E-06 | 1.47E-03 |
| protein binding (GO:0005515) | 14448 | 62 | 47.02 | 1.32 | + | 1.21E-05 | 4.02E-03 |
| nucleosomal DNA binding (GO:0031492) | 39 | 4 | 0.13 | 31.52 | + | 1.13E-05 | 4.03E-03 |
| transcription factor binding (GO:0008134) | 611 | 10 | 1.99 | 5.03 | + | 2.97E-05 | 9.24E-03 |
| chromatin DNA binding (GO:0031490) | 118 | 5 | 0.38 | 13.02 | + | 4.96E-05 | 1.45E-02 |
| structural constituent of chromatin (GO:0030527) | 66 | 4 | 0.21 | 18.62 | + | 7.89E-05 | 2.18E-02 |
| protein kinase inhibitor activity (GO:0004860) | 71 | 4 | 0.23 | 17.31 | + | 1.03E-04 | 2.34E-02 |
| ribonucleotide binding (GO:0032553) | 1926 | 17 | 6.27 | 2.71 | + | 1.15E-04 | 2.38E-02 |
| nucleosome binding (GO:0031491) | 69 | 4 | 0.22 | 17.81 | + | 9.30E-05 | 2.43E-02 |
| purine ribonucleotide binding (GO:0032555) | 1909 | 17 | 6.21 | 2.74 | + | 1.03E-04 | 2.44E-02 |
| purine nucleotide binding (GO:0017076) | 1923 | 17 | 6.26 | 2.72 | + | 1.13E-04 | 2.44E-02 |
| protein kinase regulator activity (GO:0019887) | 224 | 6 | 0.73 | 8.23 | + | 1.01E-04 | 2.51E-02 |
| identical protein binding (GO:0042802) | 2146 | 18 | 6.98 | 2.58 | + | 1.31E-04 | 2.51E-02 |
| kinase inhibitor activity (GO:0019210) | 75 | 4 | 0.24 | 16.39 | + | 1.26E-04 | 2.52E-02 |
| binding (GO:0005488) | 16594 | 65 | 54 | 1.2 | + | 1.39E-04 | 2.56E-02 |
| chromatin binding (GO:0003682) | 605 | 9 | 1.97 | 4.57 | + | 1.57E-04 | 2.79E-02 |
| kinase regulator activity (GO:0019207) | 252 | 6 | 0.82 | 7.32 | + | 1.89E-04 | 3.24E-02 |
| DNA-binding transcription factor binding (GO:0140297) | 488 | 8 | 1.59 | 5.04 | + | 1.97E-04 | 3.26E-02 |
| purine ribonucleoside triphosphate binding (GO:0035639) | 1839 | 16 | 5.98 | 2.67 | + | 2.26E-04 | 3.63E-02 |
| protein serine/threonine kinase inhibitor activity (GO:0030291) | 34 | 3 | 0.11 | 27.11 | + | 2.34E-04 | 3.64E-02 |

| | | | | | | | |
|---|---|---|---|---|---|---|---|
| microtubule binding (GO:0008017) | 274 | 6 | 0.89 | 6.73 | + | 2.93E-04 | 4.06E-02 |
| tubulin binding (GO:0015631) | 384 | 7 | 1.25 | 5.6 | + | 2.70E-04 | 4.07E-02 |
| carbohydrate derivative binding (GO:0097367) | 2287 | 18 | 7.44 | 2.42 | + | 2.91E-04 | 4.13E-02 |
| 5'-deoxyribose-5-phosphate lyase activity (GO:0051575) | 6 | 2 | 0.02 | > 100 | + | 2.87E-04 | 4.19E-02 |

*PANTHER was used to analyze overrepresented molecular function for ITE-dysregulated phosphoproteins. Gene ontology (GO) analysis was performed using Statistical Overrepresentation test (Fisher's exact test, FDR $P < 0.05$). n = 4 cell preparations.

**Table S32C. Overrepresentated GO cell component in ITE-dysregulated phosphoproteins at 24 hr.***

| Cell component | Reference Gene # (20589) | Mapped Gene # | Expected Gene # | Fold Enrichment | Direction | Raw *P* value | FDR *P* Value |
|---|---|---|---|---|---|---|---|
| non-membrane-bounded organelle (GO:0043228) | 5226 | 46 | 17.01 | 2.7 | + | 1.48E-13 | 1.51E-10 |
| intracellular non-membrane-bounded organelle (GO:0043232) | 5225 | 46 | 17 | 2.71 | + | 1.47E-13 | 3.00E-10 |
| anchoring junction (GO:0070161) | 1336 | 23 | 4.35 | 5.29 | + | 1.76E-11 | 1.20E-08 |
| cell junction (GO:0030054) | 2136 | 28 | 6.95 | 4.03 | + | 3.06E-11 | 1.56E-08 |
| focal adhesion (GO:0005925) | 423 | 14 | 1.38 | 10.17 | + | 1.03E-10 | 4.21E-08 |
| cell-substrate junction (GO:0030055) | 433 | 14 | 1.41 | 9.94 | + | 1.39E-10 | 4.73E-08 |
| cytoskeleton (GO:0005856) | 2401 | 28 | 7.81 | 3.58 | + | 4.62E-10 | 1.35E-07 |
| cytosol (GO:0005829) | 5458 | 41 | 17.76 | 2.31 | + | 3.77E-09 | 9.64E-07 |
| intracellular organelle (GO:0043229) | 13254 | 63 | 43.13 | 1.46 | + | 2.11E-08 | 4.31E-06 |
| intracellular anatomical structure (GO:0005622) | 14945 | 66 | 48.63 | 1.36 | + | 1.95E-08 | 4.43E-06 |
| microtubule cytoskeleton (GO:0015630) | 1374 | 19 | 4.47 | 4.25 | + | 5.31E-08 | 9.85E-06 |
| organelle (GO:0043226) | 14064 | 64 | 45.77 | 1.4 | + | 8.61E-08 | 1.47E-05 |
| heterochromatin (GO:0000792) | 80 | 6 | 0.26 | 23.05 | + | 3.55E-07 | 5.58E-05 |
| supramolecular complex (GO:0099080) | 1410 | 18 | 4.59 | 3.92 | + | 4.21E-07 | 6.14E-05 |
| nucleus (GO:0005634) | 7682 | 45 | 25 | 1.8 | + | 8.53E-07 | 1.09E-04 |
| cell-cell junction (GO:0005911) | 508 | 11 | 1.65 | 6.65 | + | 8.17E-07 | 1.11E-04 |
| adherens junction (GO:0005912) | 180 | 7 | 0.59 | 11.95 | + | 2.45E-06 | 2.94E-04 |
| perinuclear region of cytoplasm (GO:0048471) | 742 | 12 | 2.41 | 4.97 | + | 4.86E-06 | 5.23E-04 |
| intrinsic component of membrane (GO:0031224) | 5958 | 4 | 19.39 | 0.21 | - | 4.63E-06 | 5.26E-04 |
| integral component of membrane (GO:0016021) | 5799 | 4 | 18.87 | 0.21 | - | 7.95E-06 | 8.12E-04 |
| polymeric cytoskeletal fiber (GO:0099513) | 808 | 12 | 2.63 | 4.56 | + | 1.14E-05 | 1.05E-03 |
| spindle (GO:0005819) | 428 | 9 | 1.39 | 6.46 | + | 1.13E-05 | 1.10E-03 |
| membrane-bounded organelle (GO:0043227) | 13230 | 59 | 43.05 | 1.37 | + | 1.68E-05 | 1.49E-03 |
| supramolecular fiber (GO:0099512) | 1047 | 13 | 3.41 | 3.82 | + | 3.04E-05 | 2.59E-03 |
| supramolecular polymer (GO:0099081) | 1056 | 13 | 3.44 | 3.78 | + | 3.32E-05 | 2.71E-03 |
| protein-DNA complex (GO:0032993) | 192 | 6 | 0.62 | 9.6 | + | 4.41E-05 | 3.47E-03 |
| polysome (GO:0005844) | 67 | 4 | 0.22 | 18.35 | + | 8.34E-05 | 6.31E-03 |
| senescence-associated heterochromatin focus (GO:0035985) | 3 | 2 | 0.01 | > 100 | + | 1.03E-04 | 7.26E-03 |
| extracellular exosome (GO:0070062) | 2100 | 18 | 6.83 | 2.63 | + | 9.97E-05 | 7.27E-03 |
| extracellular membrane-bounded organelle (GO:0065010) | 2124 | 18 | 6.91 | 2.6 | + | 1.15E-04 | 7.36E-03 |
| extracellular organelle (GO:0043230) | 2124 | 18 | 6.91 | 2.6 | + | 1.15E-04 | 7.59E-03 |
| extracellular vesicle (GO:1903561) | 2123 | 18 | 6.91 | 2.61 | + | 1.15E-04 | 7.80E-03 |
| microtubule (GO:0005874) | 471 | 8 | 1.53 | 5.22 | + | 1.55E-04 | 9.32E-03 |
| pericentric heterochromatin (GO:0005721) | 29 | 3 | 0.09 | 31.79 | + | 1.51E-04 | 9.36E-03 |
| chromosome (GO:0005694) | 1852 | 16 | 6.03 | 2.65 | + | 2.45E-04 | 1.43E-02 |

| | | | | | | | |
|---|---|---|---|---|---|---|---|
| condensed chromosome (GO:0000793) | 273 | 6 | 0.89 | 6.75 | + | 2.88E-04 | 1.63E-02 |
| protein-containing complex (GO:0032991) | 5771 | 33 | 18.78 | 1.76 | + | 3.07E-04 | 1.70E-02 |
| microtubule organizing center (GO:0005815) | 840 | 10 | 2.73 | 3.66 | + | 3.90E-04 | 2.04E-02 |
| nucleolus (GO:0005730) | 1002 | 11 | 3.26 | 3.37 | + | 3.85E-04 | 2.07E-02 |
| ribonucleoprotein complex (GO:1990904) | 714 | 9 | 2.32 | 3.87 | + | 5.22E-04 | 2.67E-02 |
| zonula adherens (GO:0005915) | 9 | 2 | 0.03 | 68.29 | + | 5.59E-04 | 2.79E-02 |
| nuclear lumen (GO:0031981) | 4494 | 27 | 14.62 | 1.85 | + | 8.95E-04 | 4.35E-02 |

*PANTHER was used to analyze overrepresented cell component for ITE-dysregulated phosphoproteins. Gene ontology (GO) analysis was performed using Statistical Overrepresentation test (Fisher's exact test, FDR $P < 0.05$). n = 4 cell preparations.

**Table S32D. Overrepresentated GO protein class in ITE-dysregulated phosphoproteins at 24 hr.***

| Protein class | Reference Gene # (20589) | Mapped Gene # | Expected Gene # | Fold Enrichment | Direction | Raw *P* value | FDR *P* Value |
|---|---|---|---|---|---|---|---|
| non-motor microtubule binding protein (PC00166) | 86 | 5 | 0.28 | 17.87 | + | 1.16E-05 | 2.27E-03 |
| cytoskeletal protein (PC00085) | 627 | 9 | 2.04 | 4.41 | + | 2.04E-04 | 2.00E-02 |

*PANTHER was used to analyze overrepresented protein class for ITE-dysregulated phosphoproteins. Gene ontology (GO) analysis was performed using Statistical Overrepresentation test (Fisher's exact test, FDR $P < 0.05$). n = 4 cell preparations.

**Table S32E. Overrepresentated GO reactome pathway in ITE-dysregulated phosphoproteins at 24 hr.***

| Reactome pathway | Reference Gene # (20589) | Mapped Gene # | Expected Gene # | Fold Enrichment | Direction | Raw *P* value | FDR *P* Value |
|---|---|---|---|---|---|---|---|
| Apoptotic execution phase (R-HSA-75153) | 51 | 7 | 0.17 | 42.18 | + | 7.22E-10 | 1.80E-06 |
| Formation of Senescence-Associated Heterochromatin Foci (SAHF) (R-HSA-2559584) | 16 | 5 | 0.05 | 96.03 | + | 6.03E-09 | 7.51E-06 |
| DNA Damage/Telomere Stress Induced Senescence (R-HSA-2559586) | 61 | 6 | 0.2 | 30.23 | + | 7.92E-08 | 6.58E-05 |
| Apoptosis (R-HSA-109581) | 173 | 7 | 0.56 | 12.43 | + | 1.90E-06 | 1.18E-03 |
| Programmed Cell Death (R-HSA-5357801) | 202 | 7 | 0.66 | 10.65 | + | 5.11E-06 | 2.55E-03 |
| Cellular Senescence (R-HSA-2559583) | 164 | 6 | 0.53 | 11.24 | + | 1.87E-05 | 7.79E-03 |
| Cell Cycle (R-HSA-1640170) | 648 | 10 | 2.11 | 4.74 | + | 4.84E-05 | 1.73E-02 |
| Signaling by Rho GTPases (R-HSA-194315) | 670 | 10 | 2.18 | 4.59 | + | 6.38E-05 | 1.99E-02 |
| Signaling by Rho GTPases, Miro GTPases and RHOBTB3 (R-HSA-9716542) | 685 | 10 | 2.23 | 4.49 | + | 7.65E-05 | 2.12E-02 |
| Signal Transduction (R-HSA-162582) | 2500 | 20 | 8.14 | 2.46 | + | 9.50E-05 | 2.37E-02 |
| VEGFR2 mediated vascular permeability (R-HSA-5218920) | 27 | 3 | 0.09 | 34.14 | + | 1.24E-04 | 2.82E-02 |
| Apoptotic cleavage of cellular proteins (R-HSA-111465) | 37 | 3 | 0.12 | 24.92 | + | 2.96E-04 | 4.34E-02 |
| Gene and protein expression by JAK-STAT signaling after Interleukin-12 stimulation (R-HSA-8950505) | 37 | 3 | 0.12 | 24.92 | + | 2.96E-04 | 4.61E-02 |
| VEGFA-VEGFR2 Pathway (R-HSA-4420097) | 94 | 4 | 0.31 | 13.08 | + | 2.90E-04 | 4.83E-02 |
| M Phase (R-HSA-68886) | 375 | 7 | 1.22 | 5.74 | + | 2.35E-04 | 4.87E-02 |
| Cell Cycle, Mitotic (R-HSA-69278) | 517 | 8 | 1.68 | 4.76 | + | 2.89E-04 | 5.14E-02 |
| Disease (R-HSA-1643685) | 1679 | 15 | 5.46 | 2.75 | + | 2.80E-04 | 5.36E-02 |

*PANTHER was used to analyze overrepresented reactome pathway for ITE-dysregulated phosphoproteins. Gene ontology (GO) analysis was performed using Statistical Overrepresentation test (Fisher's exact test, FDR $P < 0.05$). n = 4 cell preparations.

Table S33. RT-qPCR primers for rat placentas.*

| Gene Name | Gene ID | Gene Description | Cat. no. |
|---|---|---|---|
| ACE2 | ENSRNOG00000031665 | Angiotensin I converting enzyme 2 | Rn01416293_m1 |
| ACTB | ENSRNOG00000034254 | Actin, beta | Rn01412977_g1 |
| AHR | ENSRNOG00000004342 | Aryl hydrocarbon receptor | Rn00565750_m1 |
| ANGPT1 | ENSRNOG00000005854 | Angiopoietin 1 | Rn01504818_m1 |
| CAV1 | ENSRNOG00000056836 | Caveolin 1 | Rn00755834_m1 |
| CYP1A1 | ENSRNOG00000019500 | Cytochrome P450, family 1, subfamily a, polypeptide 1 | Rn00487218_m1 |
| FGFR2 | ENSRNOG00000016374 | Fibroblast growth factor receptor 2 | Rn01269940_m1 |
| FLT1 | ENSRNOG00000000940 | FMS-related tyrosine kinase 1 | Rn01409533_m1 |
| GAPDH | ENSRNOG00000018630 | Glyceraldehyde-3-phosphate dehydrogenase | Rn01462662_g1 |
| HIF1A | ENSRNOG00000008292 | Hypoxia inducible factor 1 subunit alpha | Rn00577560_m1 |
| IL1B | ENSRNOG00000004649 | Interleukin 1 beta | Rn00580432_m1 |
| KDR | ENSRNOG00000046829 | Kinase insert domain receptor | Rn00564986_m1 |
| MYBPC3 | ENSRNOG00000012307 | Myosin binding protein C, cardiac | Rn01513426_m1 |
| RPLP0 | ENSRNOG00000001148 | Ribosomal protein lateral stalk subunit P0 | Rn03302271_gH |
| SRY | ENSRNOG00000062090 | Sex determining region Y | Rn04224592_u1 |
| TFRC | ENSRNOG00000001766 | Transferrin receptor | Rn01474701_m1 |
| TIE1 | ENSRNOG00000020173 | Tyrosine kinase with immunoglobulin-like and EGF-like domains 1 | Rn01417182_m1 |
| VCAM1 | ENSRNOG00000014333 | Vascular cell adhesion molecule 1 | Rn00563627_m1 |
| VEGFA | ENSRNOG00000019598 | Vascular endothelial growth factor A | Rn01511602_m1 |
| VWF | ENSRNOG00000019689 | Von Willebrand factor | Rn01492158_m1 |
| YWHAZ | ENSRNOG00000008195 | Tyrosine 3-monooxygenase/tryptophan 5-monooxygenase activation protein, zeta | Rn00755072_m1 |

*All Tagman primers were purchased from Applied Biosystems, Waltham, MA.

Table S34. RT-qPCR primers for HUVECs.*

| Gene Name | Gene ID | Gene Description | Cat. no. |
|---|---|---|---|
| ACTB | ENSG00000075624 | Actin beta | Hs01060665_g1 |
| AHR | ENSG00000106546 | Aryl hydrocarbon receptor | Hs00169233_m1 |
| AHRR | ENSG00000063438 | Aryl-hydrocarbon receptor repressor | Hs01005076_m1 |
| ALDH1A1 | ENSG00000165092 | Aldehyde dehydrogenase 1 family member A1 | Hs00946916_m1 |
| APOLD1 | ENSG00000178878 | Apolipoprotein L domain containing 1 | Hs00707371_s1 |
| AQP1 | ENSG00000240583 | Aquaporin 1 | Hs01028916_m1 |
| COL1A2 | ENSG00000164692 | Collagen type I alpha 2 chain | Hs01028956_m1 |
| CYP1A1 | ENSG00000140465 | Cytochrome P450 family 1 subfamily A member 1 | Hs01054796_g1 |
| GAPDH | ENSG00000111640 | glyceraldehyde-3-phosphate dehydrogenase | Hs04420632_g1 |
| IL6 | ENSG00000136244 | Interleukin 6 | Hs00174131_m1 |
| KDR | ENSG00000128052 | Kinase insert domain receptor | Hs00911700_m1 |
| LIPG | ENSG00000101670 | lipase G, endothelial type | Hs00195812_m1 |
| MMP1 | ENSG00000196611 | Matrix metallopeptidase 1 | Hs00899658_m1 |
| NOS3 | ENSG00000164867 | Nitric oxide synthase 3 | Hs01574659_m1 |
| POSTN | ENSG00000133110 | Periostin | Hs01566750_m1 |
| PTGS2 | ENSG00000073756 | Prostaglandin-endoperoxide synthase 2 | Hs00153133_m1 |
| VEGFA | ENSG00000112715 | Vascular endothelial growth factor A | Hs00900055_m1 |
| YWHAZ | ENSG00000164924 | Tyrosine 3-monooxygenase/tryptophan 5-monooxygenase activation protein zeta | Hs01122445_g1 |

*All Tagman primers were purchased from Applied Biosystems (Waltham, MA).
